# ISG15 Differentially Modulates Clade Ib and II MPXV Infection in MEF cells

**DOI:** 10.1101/2025.09.26.678780

**Authors:** Joseph Patrick McGrail, Irene Campaña Gómez, Antti Tuhkala, Salla Keskitalo, Markku Varjosalo, Adolfo García Sastre, Gustavo Palacios, Mari Paz Sanchez-Seco, Susana Guerra

## Abstract

The unprecedented human-to-human transmission of Clade IIb monkeypox virus (MPXV) during the 2022 outbreak has renewed focus on host determinants of viral fitness. Interferon-stimulated gene 15 (ISG15) encodes a ubiquitin-like protein with broad immunomodulatory functions, yet its role in MPXV infection remains unclear. Using representative strains from recent and historical outbreaks spanning Clades I and II, we show that ISG15 deficiency enhances viral replication and protein production in murine cells. Given that rodents are considered potential natural reservoirs of MPXV, these findings highlight the importance of studying murine models to understand virus–host interactions. Notably, the 2024 Democratic Republic of Congo strain displays reduced sensitivity to ISG15, suggesting clade-specific adaptation. ISG15 also influences viral immune evasion, as knockout cells infected with Clade II viruses expressed fewer immunomodulatory proteins and exhibited marked reductions in host protein phosphorylation. These results identify ISG15 as a determinant of MPXV infection and underscore evolutionary differences between clades.

## Introduction

The study of poxviruses has recently received an increase in attention due to the surge of the disease known as MPOX. MPOX is caused by the Monkeypox virus (MPXV), belonging to the *Poxviridae* family (1)(2)(3). Poxviruses possess an incredibly large genome, allowing them to have all of the necessary machinery to replicate in the cytoplasm (4)(5). They also have a complex morphogenetic cycle that results in the formation of two main infectious forms, Intracellular Mature Virus (MVs) and Extracellular Enveloped Viruses (EVs) (6)(7)(8). MVs are associated with host-to-host transmission and localized lesions, while EVs mediate the dissemination within the infected organism and cell-to-cell spread (9)(10)(11)(12). These characteristics manifest how the study of MPXV is rather complex.

Traditionally, MPXV has been classified in two distinct Clades. Clade I has mainly been linked to Central Africa and more severe symptoms, while Clade II is more prevalent in Western Africa and produces lesser symptoms (13)(14)(15). The strain responsible for the global 2022 outbreak that alarmed scientists worldwide belonged to Clade IIb and was characterized by an increased human-to-human transmissibility (16) (17). Last year, another important outbreak occurred in the Democratic Republic of Congo (DRC), produced by the newly identified Clade Ib, along with an uptick of a different variant of Clade Ia (18) (19). This is concerning since the more virulent Clade I strains are acquiring the increased HHT that was more prevalent in Clade IIb MPXV (19)(20). In addition, there is currently an oubtreak of Clade IIb in Sierra Leone that reinforces the dangers that MPXV can cause, especially since this is the clade responsible for the 2022 global MPOX outbreak (21).

Efforts to understand if potential virus-host interactions are what caused the increased HHT of these recent Clade Ib and IIb strains have led to the study of numerous host proteins (16)(22)(23). One such protein of interest is Interferon Stimulated Gene 15 (ISG15). ISG15 is a ubiquitin-like protein that conjugates reversibly to *de novo* proteins in a process known as ISGylation, with this process being linked to having a role in numerous important biological processes and antiviral activity (24)(25)(26)(27) It also can influence the cell response in its unconjugated form intracellularly, or possess immunomodulatory activity when secreted extracellularly (28)(29)(30)(31). Previous research has displayed its relevance in metabolism, cancer and exosome secretion (32)(33)(30)(34). In addition, ISG15 has proven to be a potentially useful vaccine adjuvant in MVA vaccine candidates (35)(36).

Our research has suggested a role for ISG15 in modulating numerous biological processes and replication of poxviruses during infection (33) (37) (38). Given the biomedical importance of MPXV, a member of the *Poxviridae* family, we now sought to determine whether ISG15 exerts similar effects during MPXV infection. Because rodents are considered potential reservoirs of MPXV, we employed murine embryonic fibroblasts (MEFs) as a relevant model system. To this end, we infected wild-type (WT) and ISG15-deficient MEFs with epidemiological isolates from the 2024 Democratic Republic of Congo outbreak (Clade Ib), the 2022 global outbreak (Clade IIb), and two earlier reference strains from Clade IIa (USA 2003 and WRAIR 7-61).This approach enabled us to assess the importance of ISG15 in MPXV evolution. A previous study from our laboratory had already examined these Clade II strains in WT MEFs, revealing slower viral growth, distinct plaque morphology and altered protein expression of the 2022 MPXV strain compared with older isolates, such as USA 2003. (39).

With respect to the role of ISG15, our results revealed increased accumulation of intracellular virus and viral proteins in ISG15-deficient cells infected with MPXV, as measured by viral titers, immunofluorescence, electron microscopy and western blotting. These effects were especially pronounced in the oldest reference strain, WRAIR. Interestingly, proteomic analysis showed that while most viral proteins were upregulated in ISG15 KO infections compared to WT, those involved in immune evasion were decreased in Clade II MPXV infections. This suggests that in the absence of ISG15, the virus may prioritize structural protein production over countering the already compromised immune response. Lastly, phosphoproteomic analysis of infected cell lysates revealed a general reduction in phosphorylation events and a drastic decrease in identified phosphosites in ISG15 KO cells. These findings highlight a critical role for ISG15 in regulating MPXV infection and suggest it may influence viral evolution and host interactions.

## Materials and Methods

### Cells and viruses

Immortalized WT MEFs and ISG 15-KO MEFs (kindly provided by K.P Knobeloch) were cultured in 1 g/L glucose Dulbecco Modified Eagle Medium (DMEM) supplemented with penicillin (100 U/ml; Sigma-Aldrich, St. Louis, MO, USA), streptomycin (100□μg/ml; Sigma-Aldrich), L-glutamine (2□mM; Sigma-Aldrich), non-essential amino acids (0.1□mM; Sigma-Aldrich), and 5% heat-inactivated fetal bovine serum (FBS) (Invitrogen Gibco). BSC40 (ATCC CRL-2761) cells were cultured in same conditions except they were supplemented with 10% FBS. Cell cultures were maintained at 37°C in a humidified incubator containing 5% CO_2_. 2024 MPXV was obtained from a clinical isolate of a Clade Ib infected patient. MpxV/PHAS-506/Passage-03/SWE/2024_09_11, Clade 1b has been provided by the Public Health Agency of Sweden. 2022 MPXV used for this work had been isolated from fluid (vesicle) content from a clinical skin (lesion) swab specimen (PV67610) and its titer previously obtained in BSC-40 cells (40). USA 2003 and WRAIR 7-61 were obtained through BEI Resources (NR-2500) (NR-27).

### Viral stock generation

All virus work was performed in a BSL3 facility by trained scientists wearing appropriate personnel protective equipment. 14 confluent BSC-40 F-175 were inoculated at MOI 0.01 with each MPXV strain. When cytopathic effect was evident, the infected cells were scraped in the flask medium with a cell scraper, and the resulting suspension transferred to sterile 20 ml centrifugation tubes. Infected cells were pelleted by centrifugation at 2000 xg and 4°C for 5 min. Supernatant was discarded, each pellet was resuspended in 1 ml of DMEM and added to the same centrifugation tube. After centrifugation, supernatant was discarded once again, and pellet was resuspended in 2-3 ml DMEM. Suspension was aliquoted, freeze-thaw cycle was conducted three times and titration was done by standard plaque assay.

### Virus titration

Intracellular and extracellular virus was quantified using standard plaque assay (41). Intracellular virus was titrated after cells were subjected to three freeze-thaw cycles.

### Protein analysis by Western blotting

Infected Immortalized MEFs lysates were acquired through addition of 100 microlitres of SDS-Laemmli sample buffer supplemented with 100mM dithiothreitol (DTT) to each M12 well plate. Cell lysates were boiled for 5 min, resolved by sodium dodecyl sulfate-polyacrylamide gel electrophoresis (SDS-PAGE) with Laemmli running buffer and transferred to polyvinylidene difluoride membranes (Merck-Millipore) in a Trans-Blot SD Semi-Dry Transfer Cell (Bio-Rad) according to the manufacturer’s recommendations. Membranes were blocked with 5% skim milk in PBS containing 0.1% Tween 20 (PBS-T) and incubated with the corresponding primary antibodies as indicated in the figure legends (Beta-actin was from Santa Cruz and F13 was kindly provided by Rafael Blasco) in 0.5% skim milk in PBS-T. Membranes were then washed with PBS-T and incubated with anti-rabbit, anti-mouse or anti-rat peroxidase-labeled antibodies (1:10,000; Sigma-Aldrich). After extensive washing with PBS-T, the immune complexes were detected by using Clarity Western ECL blot substrate (Bio-Rad) and a ChemiDoc XRS^+^ System (Bio-Rad), according to the manufacturer’s instructions.

### qPCR

MPXV Infected Immortalized WT and ISG15 KO MEFs lysates were collected at MOI 0.1 or MOI 1 and 21 hpi with the NucleoZOL reagent (MACHEREY-NAGEL, Düren, Germany) following the manufacturer’s instructions to obtain RNA. Subsequently, cDNA was obtained from 1 µg of RNA samples using High-Capacity cDNA Reverse Transcription Kit with protease inhibitor (Thermo Fisher Scientific, Waltham, MA, USA). The cDNA obtained was diluted 1:10 in endonuclease free water and the expression of F3 MPXV (E3 VACV Homologue) was analyzed with qPCRBIO SyGreen Mix Hi-ROX (PCR Biosystems, Oxford, UK). RT-qPCR was performed using StepOnePlus System™ (Thermo Fisher Scientific) following the manufacture protocol. Expression levels of *F3* were analyzed using specific oligonucleotides. Hypoxanthine-guanine phosphoribosyltransferase (*Hprt*) gene was used as a reference housekeeping gene. Six replicates were performed and quantified.

### Electron Microscopy

F-75 flask of MEF cells were seeded to obtain 90% confluence on day of infection. After standard infection procedure with each MPXV strain, medium was removed 21 hpi, cells were washed with PBS 1X and chemical fixation of cells with 5 ml of glutaraldehyde 2.5%-tannic acid 1% in 0.4M HEPES was performed. After 2 hours, cells were carefully scraped with cell scraper and passed to 20 ml centrifugation tube. Samples were centrifuged at 2000 rpm and 4 °C, and afterwards fixation buffer is removed leaving only a small amount and cell pellet. Pellet is resuspended in 1ml of in 0.4M HEPES and stored at 4 °C. Samples undergo postfixation with osmium tetroxide 1% in potassium ferricyanide, are washed, uranyl acetate in water 2% is used and washed once again. Samples are dehydrated with increasing concentrations of acetone and are infilatrated with increasing concetrations of resins (epon:acetone 50% and epon 100%). Lastly, samples are encapsulated and polymerized with 100% resin. Ultrathin sections (80 nm) of the resin embedded cells were obtained with an ultramicrotome (Leica EM UC7) and a 35° diamond knife (Diatome) and collected on 3 mm copper slot grids (Agar Scientific) coated with 1% formvar in chloroform. Sections on grids were post-stained in 1% Uranyl-acetate in 70% methanol for 5 min. followed by Reynolds Lead-citrate for 3 min. The sections were examined and imaged with a transmission electron microscope (TEM, JEM-JEOL2100+).

### Inmunofluorescence

Immortalized MEFs were grown on 12-mm-diameter glass coverslips in DMEM–5% FCS to a confluence of 40 to 50% and mock infected or infected at a multiplicity of infection (MOI) of 0.1 PFU/cell, 0.5 PFU/cell and 1 PFU/cell with the different Clade Ib/II MPXV strains. At 24hpi, cells were washed with PBS, fixed with 4% paraformaldehyde, permeabilized with 0.25% Triton X-100 in PBS for 30□min, and blocked in PBS with 10% FCS for 30□min at room temperature. Poli-VACV (Invitrogen, Cat. No.14-5758-82), a policlonal antibody was used as the primary antibody. Alexa Fluor 488-conjugated rabbit IgG antibodies (Invitrogen) were used as secondary antibodies. Cell nuclei were stained with 4′,6′-diamidino-2-phenylindole (DAPI; Sigma, 1:200). Fluorescentent microscopy was performed using an EVOS fluorescent microscope, images were collected and processed with LAS□X software (Leica, Wetzlar, Germany).

### Urea 8M Buffer Lysis of infected cells

Buffer was prepared by adding 12.02g of urea and 16 ml of 50 mM NH_4_HCO_3_. F-75 flask of MEF cells were seeded to obtain 90% confluence on day of infection. After standard infection procedure with each MPXV strain, medium was removed 21 hpi and 500 microliters of Urea 8M buffer was used to lyse cells with the help of a cell scraper. Lysate is transferred to Eppendorf’s and maintained at 4 °C.

### Tryptic digestion and desalting of samples

Total concentration of homogenates was measured with Bio-Rad Protein Assay Dye. 50 μg of total protein from each sample was carried forward for total proteomics). The urea concentration was diluted from 1M to 100 mM NH4HCO3. Then, samples were reduced with 5 mM Tris (2-carboxyethyl) phosphine hydrochloride, alkalinized with 10 mM idoacetamide) at room temperature, to digested with trypsin at 37 C for 16 hours using Modified Trypsin Grade Sequencing). After digestion, samples were acidified with 10% trifluoroacetic acid and desalted with BioPureSPN PROTO 300 C18 Mini columns according to the manufacturer’s instructions. After the samples were desalted, they were dried in a concentrating centrifuge. The dried peptides were reconstituted in 30 μl Buffer A (0.1% (vol/vol) TFA, 1% (vol/vol) acetonitriphyl in HPLC-grade water) and analyzed on a timsTOF Pro2 mass spectrometer.

### Mass spectrometry analysis

Desalted samples were analyzed using a hybrid trapped ion mobility quadrupole TOF mass spectrometer through a CaptiveSpray nano-electrospray ion source (Bruker Daltonics). MS analysis was performed in positive ion mode with the dia-PASEF method with samples with data optimized by data independent analysis (DIA) scan parameter. The DIA acquisition windows was assembled separately for phosphoenriched samples due to the known drift in mass and ion mobility of phospho-modified peptides. To analyze diaPASEF data, raw data (.d) will be processed with DIA-NN v18.0 (42) using spectral library generated by UniProt proteome.

### Statistical analysis of MS proteomics data

The input document to advance DIA data analysis was the DIA-NN Report.pg.matrix. For data pre-processing a proprietary R script was used. Raw intensity values were log2 transformed and normalized by median. Afterwards, missing values were inserted using QRILC imputation. For sample group comparisons, p-values were calculated using the student’s t-test using the Python package scipy (43) and adjusted using the Benjamini-Hockberg method through a Statsmodels package (44).

### Pathway analysis of proteomics data

ShinyGOv0.81 was used to measure biological process pathway enrichment after input of MPXV infected MEF cellular protein true values (45). Fold enrichment, FDR and n values were used to construct a plot with the use of GraphPad 9. Venn diagram analysis performed with InteractiVenn (46).

### Kinase enrichment analysis of proteomics data

Kinase enrichment analysis was conducted with Kinase Enrichment Analysis 3 (KEA3) web software on proteomic and phospho-proteomic data (47)

### Statistical analysis

One-way ANOVA tests were used to analyze the differences in mean values between groups with the use of GraphPad 9. All results are expressed as means ± the standard deviations (SD). Significant differences are defined as: *p ≤ 0.05; **p ≤ 0.005; ***p ≤ 0.001, ****p ≤ 0.0001.

## Results

### Viral progression is accelerated in Clade Ib/II MPXV infected ISG15 KO MEFs

To address the role of ISG15 in Clade Ib/II MPXV infection, we assessed viral mRNA expression by quantitative PCR using the F3 probe, a homolog of the VACV E3 gene, in WT and ISG15 deficient MEFs with the Clade Ib (2024 MPXV) and Clade II strains (2022 MPXV, USA 2003 and WRAIR). ISG15-deficient cells exhibited significantly higher viral mRNA levels compared to wild-type (WT) controls. Notably, this effect appeared less pronounced with the 2024 MPXV strain compared to earlier variants (Figure 1A).

**Figure 1.**
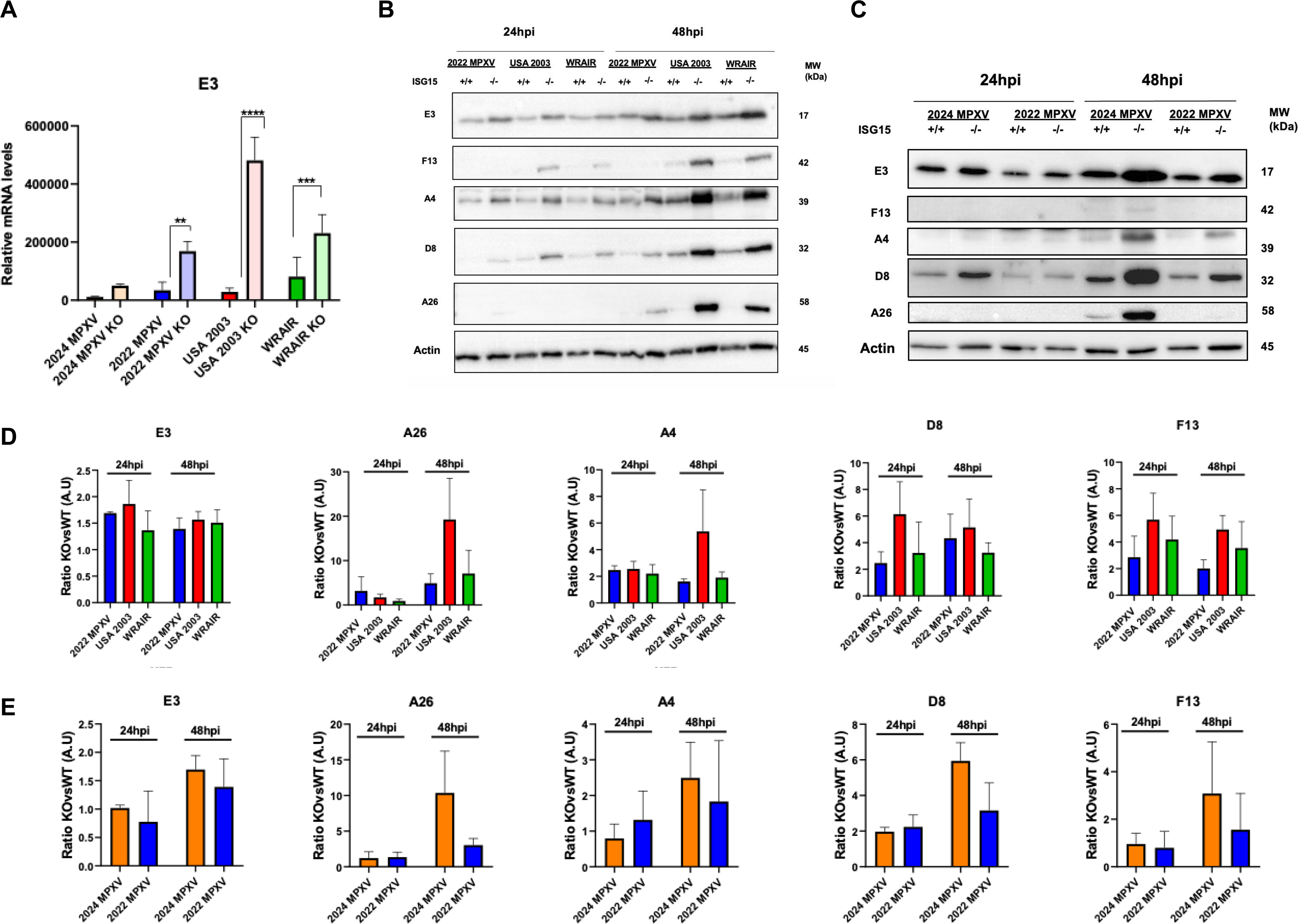
Loss of ISG15 enhances viral gene and protein production in Clade Ib and II MPXV infected MEFs. (A) Quantitative PCR analysis of F3 gene expression (VACV E3 homologue) in WT and ISG15-KO MEFs infected with four MPXV strains (MOI 0.1, 24 hpi). (B) WT and ISG15-KO MEFs infected with three Clade II MPXV strains (MOI 0.1); cell lysates collected at indicated times post-infection were analysed by immunoblotting using antibodies against VACV early protein E3, intermediate protein F13, and late proteins D8, A26 and A4. (C) As in (A), comparing the 2024 and 2022 MPXV strains. (D) Densitometric quantification of viral proteins from Clade II MPXV-infected WT and ISG15-KO MEFs, expressed as KO/WT ratios normalized to β-actin (n=3). (E) As in (D), focusing on the 2024 and 2022 strains. Molecular weights (kDa) are indicated based on protein standards.

In order to further investigate the impact of ISG15 on MPXV infection, we next analyzed viral protein production. From this point forward, MPXV viral proteins are referred to using their more commonly known VACV counterparts. Western blot analysis was performed on ISG15 knockout (KO) and wild-type (WT) MEFs infected with each Clade II MPXV strain at an MOI of 0.1, harvested at two timepoints (Figure 1B). A parallel comparison between the 2024 and 2022 MPXV strains was also conducted under identical conditions (Figure 1C). Blots employed antibodies against viral proteins expressed at different stages of the viral life cycle—early (E3), intermediate (F13), and late (A4, D8, A26)—confirmed consistently increased protein levels in ISG15-deficient cells for all MPXV strains and timepoints analyzed. Notably, the late protein A26 exhibited the most pronounced upregulation. Densitometric analysis of band intensities across three independent replicates supported these findings (Figures 1D) (Figure 1E). Together, these results suggest that ISG15 plays a critical role in restricting MPXV replication in MEFs, particularly through limiting viral mRNA levels and protein synthesis.

To further validate these findings and study its relevance in the viral cycle, we performed immunofluorescence on cells infected at low multiplicity of infection (MOI 0.1) and fixed at 24 hours post-infection (hpi). Staining with a polyclonal anti-VACV antibody revealed a greater number of infected cells in ISG15 knockout cultures, although 2024 MPXV benefitted the least from ISG15 absence (Figure 2A). In addition, we performed a low multiplicity infection (MOI 0.1) and a viral growth curve. We quantified intracellular titer to observe if differences in accumulation of MPXV virus titer could be observed. Results obtained indicate that absence of ISG15 in MEF cells leads to a more elevated amount of titer in all MPXV strains at 24 and 48hpi (Figure 2B). The strain that seems to benefit the most from the absence of ISG15 is WRAIR 7-61, with a statistically significant increase of intracellular titer in comparison to infection of WT cells, while 2024 MPXV is barely modified. However, when the experiments were conducted at a high multiplicity of infection (MOI 1), the differences in viral titers between WT and ISG15 KO cells were substantially reduced (Figure S1A), and viral protein levels appeared more similar across all MPXV strains, regardless of host genotype (Figure S1B). These findings suggest that a higher viral input can overcome the ISG15-mediated antiviral restriction observed in WT cells.

**Figure 2.**
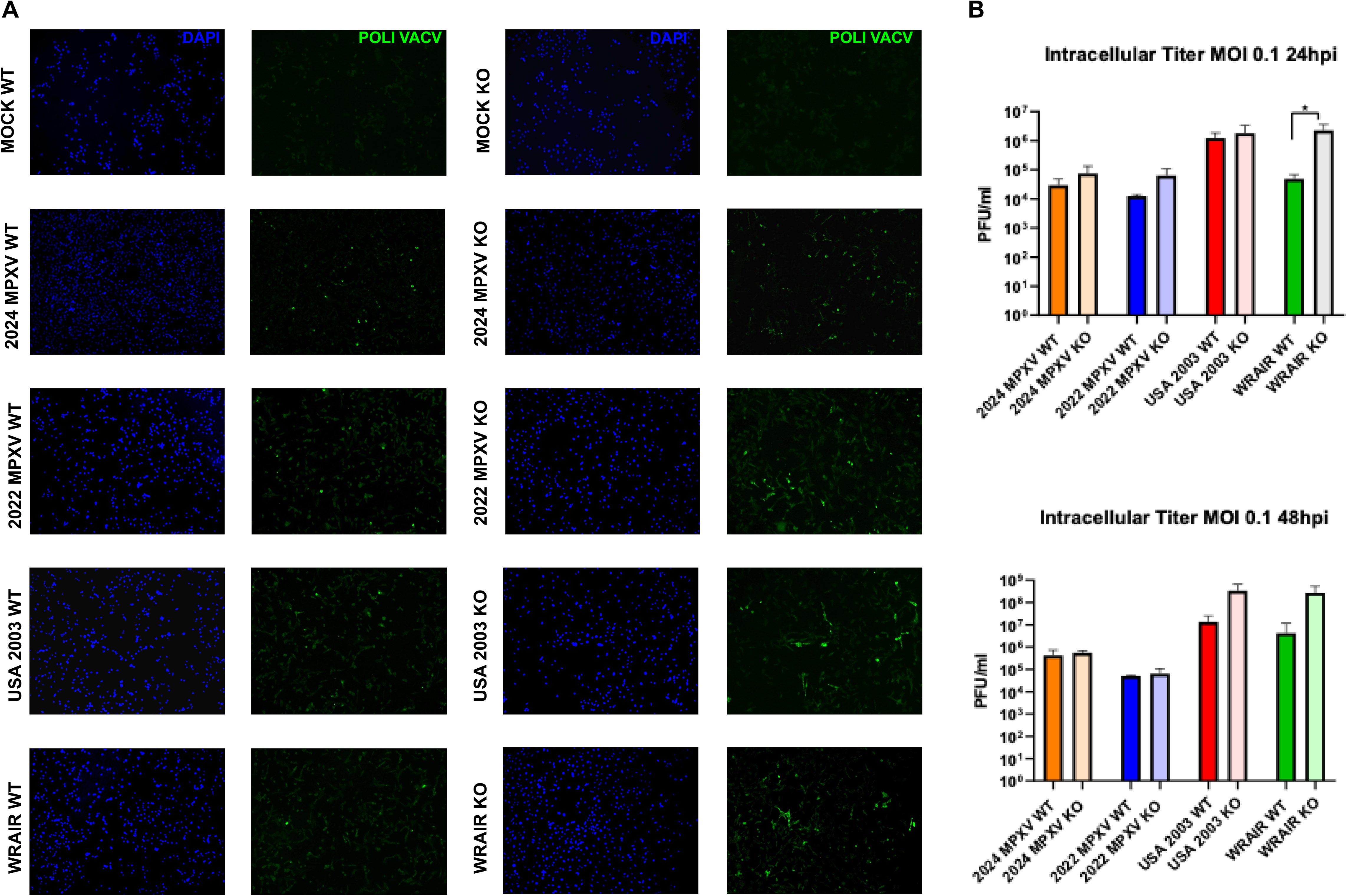
Clade Ib/II MPXV exhibits enhanced viral progression in ISG15 deficient MEFs. (A) Representative immunofluorescence images of Clade II MPXV-infected WT and ISG15 KO MEFs (MOI 0.1, 24 hpi). Infected cells were stained with polyclonal anti-VACV antibody (green), and nuclei with DAPI (blue). (B) WT and ISG15-KO MEFs were infected with four MPXV strains (MOI 0.1), and infectious particles in cell lysates (intracellular virus) were quantified by plaque assay at 24 and 48 hpi. Data represent mean ± SD from three independent experiments.

Collectively, these findings indicate that ISG15 acts as an important restriction factor against MPXV infection. However, this antiviral effect appears to be strain and multiplicity -dependent, with some MPXV variants, such as the 2024 strain, displaying reduced sensitivity to ISG15-mediated restriction.

### ISG15 modulates Clade Ib/II morphogenesis

To visualize differences in the production of MPXV viral forms, transmission electron microscopy (TEM) was performed. At 21 hours post-infection (hpi), following infection at a high multiplicity of infection (MOI 1), distinct poxvirus morphogenesis stages were identified across multiple cell profiles for each MPXV strain. ISG15 KO MEFs consistently displayed a higher abundance of viral forms across all strains, a finding further supported by quantification (Figures 3A). Interestingly, the effect of ISG15 deletion appeared to vary depending on the viral strain and the stage of the morphogenetic program affected. Early viral forms were particularly enriched in cells infected with the 2022 MPXV strain, whereas mature virions (MVs) and wrapped forms were more prominent in USA 2003 and WRAIR 7-61 (Figures 3B-C). These differences likely reflect strain-specific replication kinetics. In this regard, USA 2003 displayed the fastest progression through the viral cycle and 2022 MPXV the slowest.

**Figure 3.**
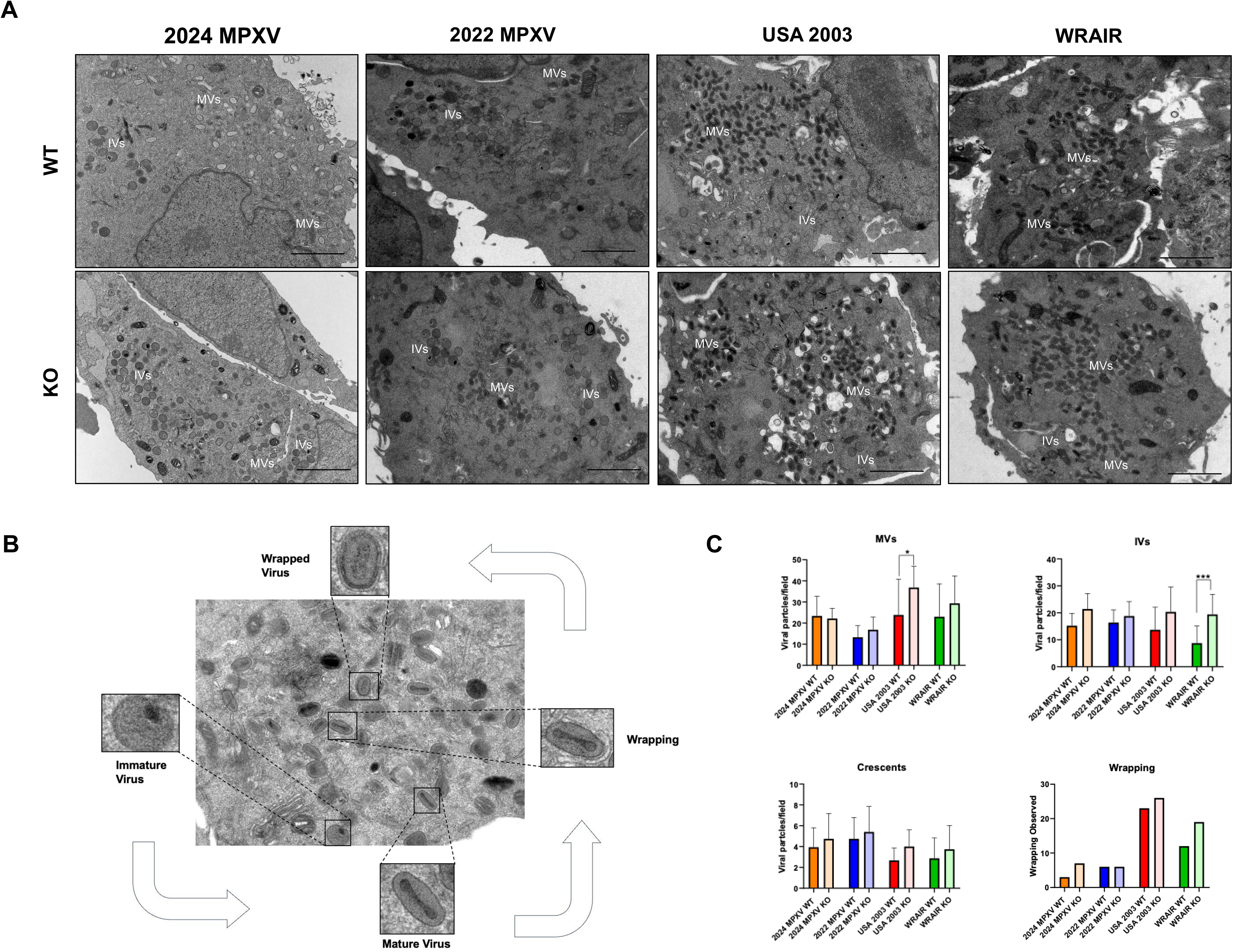
Electron microscopy reveal a role for ISG15 in MPXV morphogenesis. (A) Representative TEM images of Clade Ib/II MPXV-infected WT and ISG15 KO MEFs (MOI 1, 21 hpi). Scale bar: 2 μm. (B) TEM cell profile of a USA 2003 MPXV-infected ISG15 KO MEF showing individual viral forms: immature virus, mature virus, wrapping and wrapped virus. (C) Quantification of viral forms (mature virus [MV], immature virus [IV], crescents [C], Wrapping) in TEM cell profiles. Data from 15 cell profiles across three biological replicates.

Altogether, these observations suggest that ISG15 has an impact in the viral cycle, regulating MPXV morphogenesis in a strain-dependent manner, with its impact varying across distinct stages of the viral replication program.

### Proteomic Analysis Reveals ISG15-Dependent Regulation of Viral Proteins Across MPXV Strains

To investigate the impact of ISG15 on viral and host protein expression during MPXV infection, we performed quantitative proteomic analysis on WT and ISG15 KO MEFs infected with all MPXV strains used in this study, including both Clade Ib and Clade II viruses. Infections were performed under conditions identical to those of the previous ME experiment, using a high multiplicity of infection (MOI = 1), and samples were harvested at 21 hours post-infection (hpi) (Figure 4A). This approach allowed us to compare protein expression profiles across viral strains in the presence or absence of ISG15. Data-independent acquisition (DIA) identified a total of 5489, 5669, 5133 and 4774 proteins, of which 134, 1932, 2671 and 3031 were differentially expressed in 2024, 2022, USA 2003 and WRAIR ISG15 KO vs WT MEF infection (P ≤ 0.05, Log2 Fold Change ≥ 0.68, 4 biological replicates) (Figures 4 B–D). Strikingly, 2024 MPXV presented the least amount of significantly altered proteins when comparing ISG15 KO vs WT infection, reinforcing that this strain presents a reduced sensitivity to ISG15 absence. All MPXV proteins are reported according to their VACV Western Reserve homologues.

**Figure 4.**
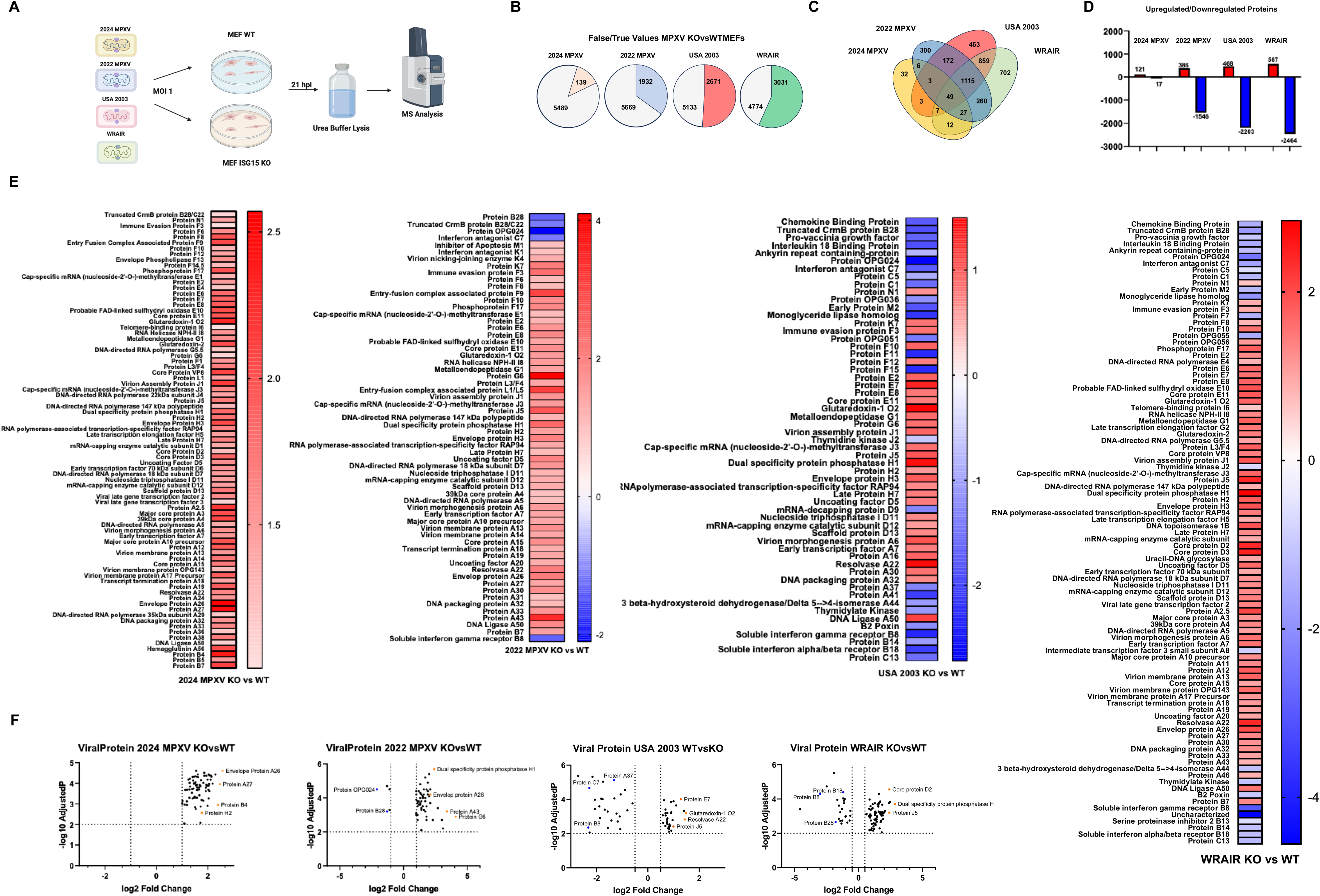
Proteomic analysis reveals differential viral protein expression in ISG15 KO MEFs infected with Clade Ib and II MPXV strains. (A) Experimental workflow for proteomic sample preparation and LC-MS/MS analysis. Four biological replicates were used. (B) Pie chart showing number of differentially expressed proteins in ISG15 KO vs WT MPXV infected MEFs compared to false values. (C) Venn diagram of differentially expressed proteins across Clade Ib and II infections. (D) Bar graph indicating the number of upregulated and downregulated proteins per strain. (E) Heat Map displaying Log2 Fold Change of differentially expressed viral proteins in ISG15 KO MEFs vs WT MEFs MPXV infection. (F) Volcano plot of differentially expressed viral proteins in ISG15 KOvsWT MPXV infections. Data shown meet both log_2_ fold change and –log_10_ adjusted p-value cutoffs.

We first analyzed differences in viral protein abundance between KO and WT conditions for each MPXV strain, presenting the data as heatmaps and volcano plots (Figures 4E-F)(Table S1). Among the four strains, WRAIR 7-61 showed the greatest increase in viral protein expression in ISG15 KO MEFs, consistent with previous growth kinetics and western blot results. Across all strains, a general increase in viral protein abundance was observed in the absence of ISG15. However, notable exceptions included several immune evasion proteins located at the genomic termini, which were reduced in ISG15 KO Clade II MPXV infections. This suggests that ISG15 may selectively regulate viral protein expression. This is feature that is not observed in ISG15 KO 2024 MPXV infection, where all proteins are increased regardless of being located at the genomic termini. This highlights potential differences between MPXV clades in response to ISG15 presence.

### ISG15 Deficiency Alters Significantly Host Cell Processes During Clade II MPXV Infection, but not Clade Ib

Beyond its effect on viral proteins, ISG15 deficiency led to substantial changes in host proteome composition during Clade II MPXV infection. Surprisingly, this was not at all the case in 2024 MPXV infected ISG15 KO MEFs. Only approximately 60 cellular proteins out of a total 139 proteins were differentially expressed during 2024 MPXV ISG15 KO infection. This contrasts with Clade II MPXV infection, where from 1850 to 2900 cellular proteins were significantly altered during ISG15 absence.

Biological process pathway analysis revealed that processes related to the mitotic cell cycle, cytoskeletal organization, and organelle dynamics were particularly affected in Clade II MPXV infection. Due to the low amount of differences in cellular protein amount in 2024 MPXV infection when ISG15 is absent, pathway analysis was not as robust but affected processes included cell motility, supramolecular fiber and actin cytoskeleton organization (Figures 5A-B).

**Figure 5.**
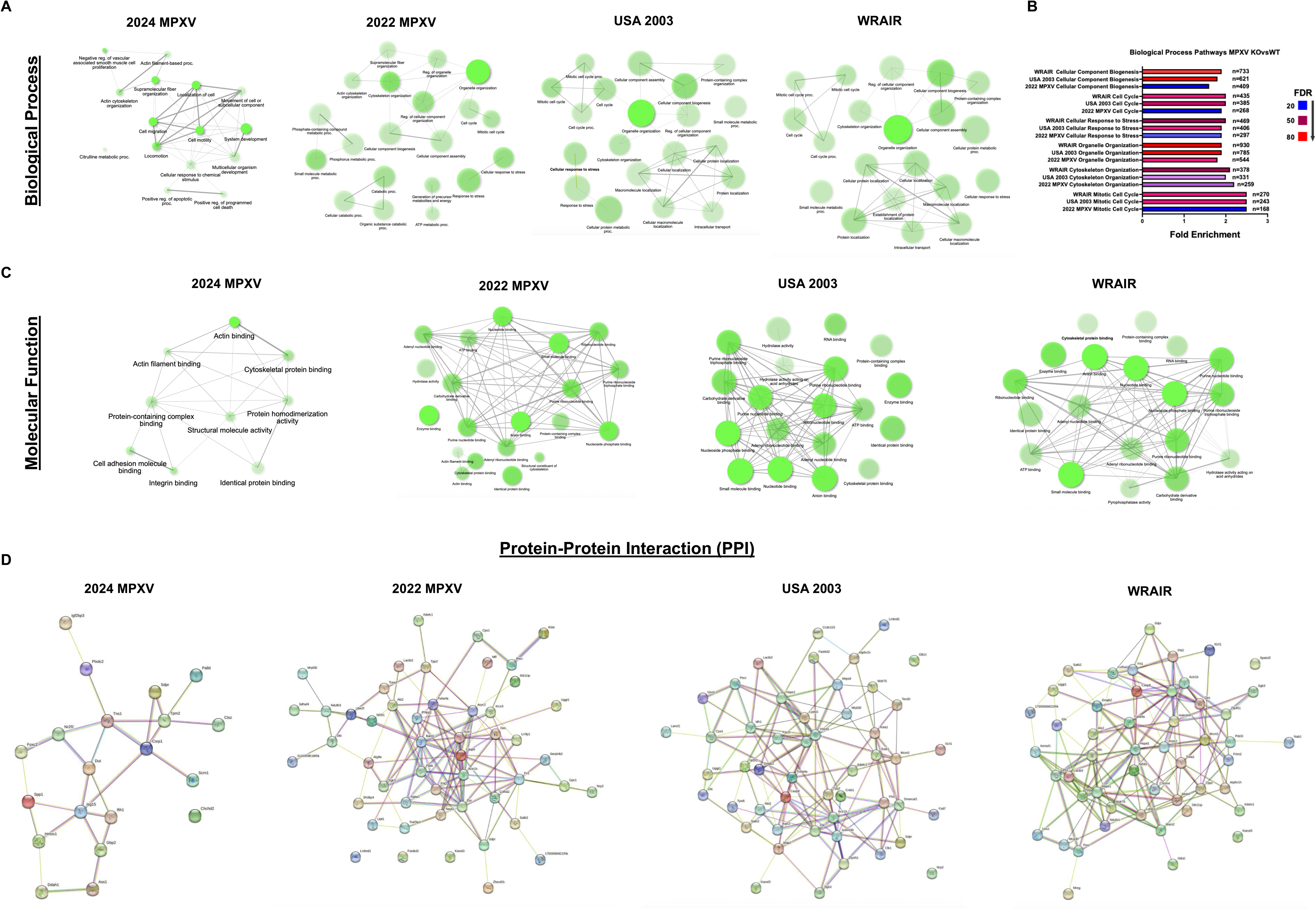
Significantly altered host protein expression in ISG15 KO MEFs infected with Clade II MPXV strains, but not Clade Ib. (A) Network graphs displaying the most enriched biological process pathways comparing ISG15 KO vs WT MPXV infected MEFs for each strain. n= 20 pathways. (B) Graph comparing the fold enrichment of common biological process pathways produced in each ISG15 KO vs WT MPXV strain infection. Fold enrichment indicates the number of proteins found to be associated with the indicated pathways. False discovery rate (FDR) has been colored blue to red, with blue indicating less trustworthy and red indicating more certainty in values calculated. N displays the number of proteins to be associated with the indicated pathway. (C) Network graphs displaying the most enriched molecular function pathways comparing ISG15 KO vs WT MPXV infected MEFs for each strain. n= 20 pathways. (D) Protein-protein interactions (PPI) of ISG15 KO vs WT MPXV infected MEFs for each strain. n = 50 proteins.

Molecular function pathway analysis displays that a common altered pathway between Clade II MPXV strains is nucleotide and nucleoside phosphate binding, indicating that ISG15 is involved in maintaining energy homeostasis during viral infection, as these processes are involved in ATP and NADPH generation (48). The only notable molecular functions of interest in Clade Ib MPXV infection are related to actin and cytoskeletal protein binding, correlating with biological process pathway results (Figure 5C). Studying the localization of these proteins informed us that the majority of the altered proteins are in the cytosol. Other sites of interest are non-membrane bounded organelles and mitochondrias where ISG15 has an important role during VACV infection (49) (33) (Figure S2). The cytoskeleton is also regulated by ISG15 after infection, as previously demonstrated (38).

Protein–protein interaction (PPI) analysis comparing ISG15 knockout (KO) and wild-type (WT) infections across multiple MPXV strains revealed key host factors potentially regulated by ISG15 during infection (Figure 5D)(Figure S3)(Table S2-S5). In the 2024 Clade Ib strain, ISG15 deficiency was associated with differential regulation of palladin, a protein involved in actin cytoskeleton organization, and forkhead box protein C2, a transcription factor implicated in cellular differentiation (50)(51). This highlights that although Clade Ib induces fewer global host protein changes compared to Clade II, it can still remodel host cellular processes in the absence of ISG15. In 2022 MPXV-infected ISG15 KO MEFs, alterations were observed in neuropilin-2, linked to viral entry, and Zfand2b, involved in protein homeostasis (52)(53). The USA 2003 strain induced a notable downregulation of interleukin-1 receptor-like 1 (IL1RL1), the receptor for IL-33, indicating that ISG15 may protect against MPXV-mediated suppression of innate immune responses (54). Additionally, this strain altered the expression of KAT8 regulatory NSL complex subunit 3 (Kansl3), a critical component of the NSL histone acetyltransferase complex involved in transcriptional regulation (55). Finally, WRAIR MPXV infection in ISG15 KO cells revealed changes in melanoregulin and NGFI-A-binding protein 1 (Nab1), proteins associated with intracellular trafficking and transcriptional repression, respectively (56)(57). Together, these findings underscore the broad regulatory role of ISG15 in limiting MPXV-induced manipulation of host cell machinery.

Together, these findings demonstrate that ISG15 plays a key regulatory role in host protein expression during Clade II MPXV infection, but not as significant in the case of Clade Ib infection, underscoring evolutionary differences between MPXV clades.

### ISG15 Absence Dampens Phosphorylation in Clade II MPXV Infection

Phosphorylation is a key post-translational modification involved in regulating numerous cellular processes (58)(59). To investigate how ISG15 deficiency influences phosphorylation during MPXV infection, we conducted a phospho-proteomic analysis comparing WT and ISG15 KO MEFs infected with Clade Ib and II MPXV strains. Previous research has also displayed that phosphorylation of specific viral proteins, such as SARS CoV-2 N protein, promoted its ISGylation, reinforcing the important relationship between ISG15 and phosphorylation events (60).

Total phosphorylated protein and phosphosites identification highlighted how 2024 MPXV differed from its Clade II brethren. In Clade II infected ISG15 KO cells, a reduction in phosphorylated proteins was visible, with this being more evident in USA 2003 infection. Notably, the number of phosphosites identified also decreased significantly across all Clade II MPXV strains in the absence of ISG15. This was not significant in 2024 MPXV, with even a slight increase in phosphorylation being observed in ISG15 KO cells (Figure 6A). Most of this is due to viral proteins being differentially phosphorylated in this strain. Volcano plots reinforced these results, with this effect being less pronounced in cells infected with 2022 MPXV and WRAIR 7-61, although a shift toward the left side of the plots was still observed, suggesting a trend toward reduced phosphorylation even if not statistically significant (Figure 6B).

**Figure 6.**
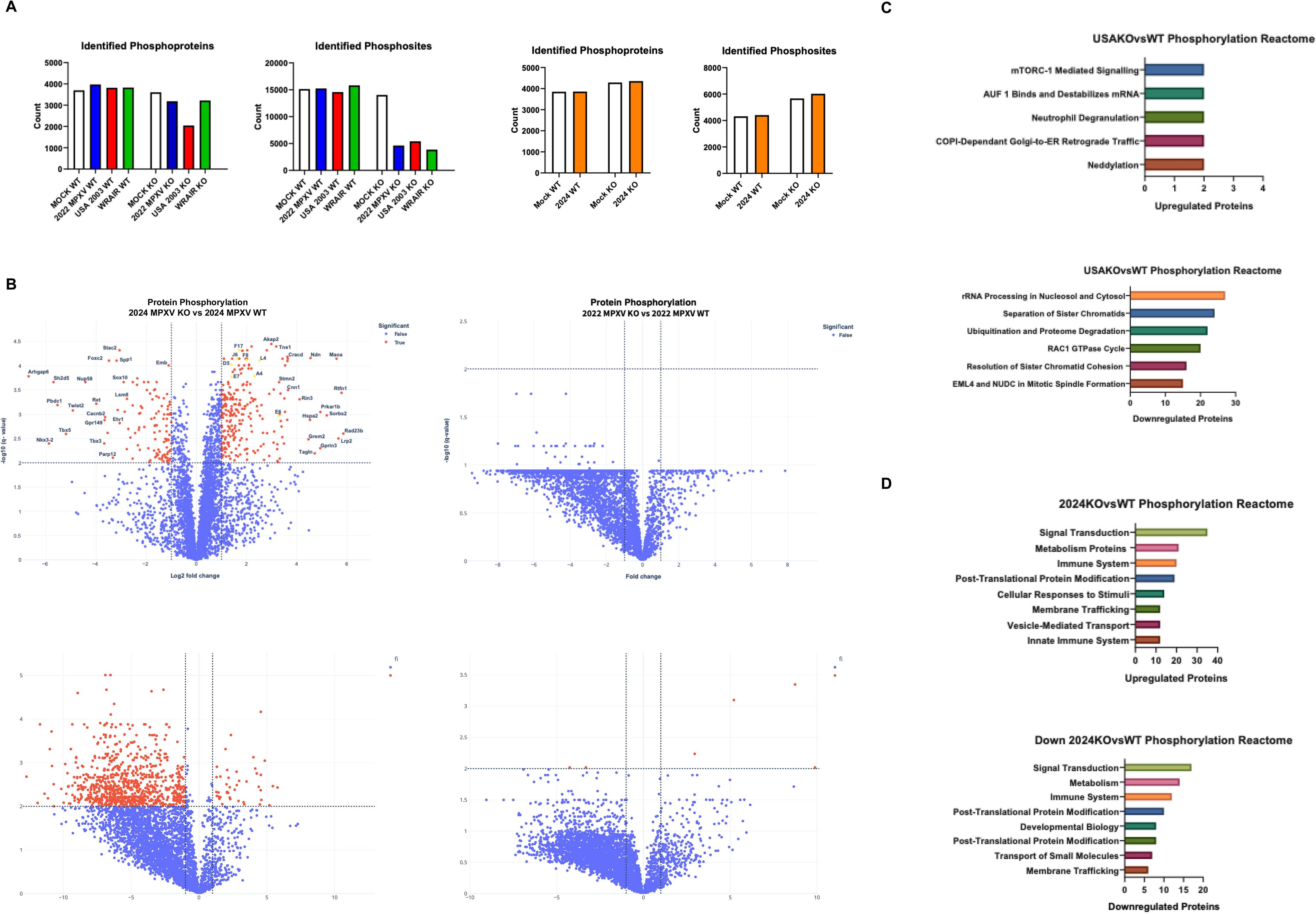
Phosphorylation is reduced during Clade II MPXV infection in ISG15 deficient MEFs. (A) Number of identified phosphoproteins and phosphosites in mock, WT and KO MPXV infected cells. (B) Volcano plots showing log_2_ fold change and - Log_10_ p-values of differentially phosphorylated proteins in ISG15 KO MEFs vs WT MPXV infected MEFs. (C) Pathway enrichment analysis of differentially phosphorylated proteins in USA 2003 and 2024 MPXV-infected KO vs WT MEFs.

Due to the higher number of differentially phosphorylated proteins in the 2024 MPXV and USA 2003 dataset, pathway analysis was focused on these strains. In USA 2003 infected ISG15 KO cells, increased phosphorylation was associated with reactome pathways such as Neddylation, COPI-mediated Golgi-to-ER retrograde transport, and neutrophil degranulation. In contrast, pathways with significantly reduced phosphorylation included mitotic spindle formation, sister chromatid cohesion and separation, ubiquitination, and proteasome-mediated degradation (Figure 6C). In 2024 MPXV regulated (up and down) phosphorylation pathways included signal transduction, metabolism, immune system and post translational modifications such as ubiquitin binding (Figure 6D). Biological process, molecular function and cellular component pathway analysis also displayed numerous differences between 2024 MPXV and USA 2003 phosphorylation in cells lacking ISG15 (Figure S6)

To validate our data we performed a Western blot analysis using anti-phospho-Tyrosine and anti-phospho-Serine antibodies corroborating the Clade II MPXV phospho-proteomic findings. A marked reduction in phosphorylation-associated bands was observed in ISG15 KO cells, particularly in USA 2003-infected samples (Figure S7A) confirming the phospho-proteomic data (Figures S7B–C) (Figure S8).

These findings suggest that ISG15 plays a central role in maintaining phosphorylation homeostasis during MPXV infection, with its absence leading to dysregulation of key cellular signaling pathways, specially in the context of USA 2003 MPXV.

### ISG15 regulates stress-activated and mitotic kinase cascades during MPXV infection

Further analysis was performed on our obtained proteomic and phospho-proteomic datasets through kinase enrichment analysis. Results obtained of proteome analysis indicated that relevant enriched kinases included MAP kinases (MAPK1, MAPK3, MAPK8, MAPK14), PLK1 and AURKA in MPXV infected ISG15 KO MEFs (Figure S9). This indicates an increased alteration in stress response and cell cycle control during infection when ISG15 is not present (61) (62) (63).

Phospho-proteomic analysis of the 2024 MPXV and 2003 USA strains further supported this hypothesis, highlighting kinases implicated in key cellular processes, including MAPK, BUB1B, CDK1, and EGFR (Fig. S10). Beyond its role in infection, ISG15 mediates the protective effects of interferon on DNA replication, promotes cell survival under stress, and contributes to therapy resistance, linking immune signaling to genome stability and disease outcomes (64). These findings suggest that ISG15 may regulate stress responses and cell cycle progression during pathogen infection, and that its absence could lead to errors in cell division.

## Discussion

Global increase in monkeypox (mpox) cases and human-to-human transmission (HHT) between 2022-2025 has raised major public health concerns (65)(66)(67)(68). While the Democratic Republic of Congo (DRC) continues to report high numbers of endemic cases, the molecular mechanisms driving the enhanced transmissibility of recent MPXV strains remain poorly understood (18)(69)(70). APOBEC3-mediated mutagenesis has been implicated in shaping the genome of circulating MPXV strains, potentially facilitating adaptation to the human host (71)(72). Another proposed mechanism involves a shift in the MV/EV ratio, whereby reduced production of EVs relative to MVs leads to more localized lesions and increased direct transmission, but reduced systemic virulence (73)(74).

Given its established antiviral role (75), we sought to investigate its contribution to MPXV infection dynamics using four strains representing distinct clades and temporal origins. We investigated the contribution of ISG15 to MPXV infection using four representative strains spanning distinct clades and time points: a 2024 Clade Ib human isolate linked to travel in the Democratic Republic of Congo, a 2022 Clade IIb isolate from the New York City outbreak, and two Clade IIa reference strains—the USA 2003 zoonotic isolate and the WRAIR strain from non-human primates not associated with human disease (76)(77)(78)(39). The absence of human-to-human transmissibility in the latter two strains provides a critical contrast for exploring evolutionary differences in viral adaptation (39)(79) .

Low MOI infections revealed that ISG15 knockout (KO) cells consistently supported higher viral gene and protein expression, in addition to higher intracellular viral titers. This feature was not as impactful in the recent 2024 MPXV infection. WRAIR, by contrast, exhibited the most pronounced increase, hinting that more recent MPXV strains may have evolved mechanisms to counteract ISG15’s antiviral effects. This characteristic is even more evident in the Clade Ib strain, as other experiments have displayed. High MOI infections minimized these differences, suggesting that viral load can overwhelm ISG15-mediated restriction.

Western blot analyses confirmed higher viral protein abundance in ISG15-deficient cells, independent of strain. Among these, A26 stood out with up to 20-fold higher expression in USA 2003 KO cells. Given that A26 impairs EV formation by retaining MVs intracellularly, its upregulation supports the notion that ISG15 absence promotes MV accumulation and hinders viral egress (80). Whether ISG15 directly regulates A26 via ISGylation or indirectly modulates its expression remains an open question.

Transmission electron microscopy (TEM) further supported these findings. A statistically significant increase in MVs was observed in ISG15 KO cells infected with USA 2003, with a similar trend in other strains. In contrast, the number of wrapped virions remained largely unchanged, implying that while early events in morphogenesis are accelerated in ISG15-deficient cells, progression to extracellular forms is impaired. Early structures like crescents and immature virions (IVs) were also more frequent in KO cells, suggesting faster early-stage replication but inefficient egress. Previous publications have used TEM as a tool to visualize how MPXV utilizes our cells to its advantage, with our intent to also study this characteristic in future experiments, for example alterations in organelles related to cell division (81)(40).

Proteomic analyses revealed a general increase in viral protein abundance in ISG15-deficient cells. However, a subset of immunomodulatory proteins encoded in the terminal genomic regions—including B28, B8, C7, and C13—were underrepresented in Clade II KO infections. One hypothesis is that, in the absence of ISG15, MPXV prioritizes structural protein synthesis over immune evasion due to a dampened host response. Alternatively, the rapid replication kinetics in KO cells may curtail early gene expression windows, explaining the reduced abundance of these early-expressed proteins. Absence of this key characteristic in 2024 MPXV highlights differences from Clade I strains, while overall viral protein levels remain elevated, showing that ISG15 still limits viral protein production in this less sensitive strain. Among the most abundantly produced viral proteins is A26, a pattern also observed in the 2022 MPXV and WRAIR strains, suggesting that ISG15 influences A26 abundance across all viral strains examined. In the case of the USA 2003 strain, this increase appears less pronounced, potentially reflecting differences in viral replication kinetics. However, when infections were conducted at low multiplicity of infection (MOI), the increase was consistently observed, supporting a role for ISG15 in regulating A26 levels. This finding further indicates that A26, a protein associated with mature virion presence, correlates with higher intracellular viral accumulation, in agreement with previous western blot analyses. Future studies are required to determine whether these differences arise from direct interactions with ISG15 or from indirect effects on viral transcriptional or translational programs.

Biological process enrichment analysis of host proteomes revealed major changes in cytoskeleton organization and cell cycle regulation in Clade II infected ISG15 KO cells. Disruption of centrosome biology, including centriole overamplification, has been reported during other viral infections and in VACV infection, which may indicate it also occurring in MPXV-infected cells (82)(83)(84). We identified several candidate proteins involved in centrosome dynamics among those most altered in ISG15 knockout compared with wild-type infections, highlighting them for further investigation. Proteomic profiling shows that 2024 infection induces negligible host protein changes in ISG15-deficient MEFs compared with wild-type cells. While 2024 MPXV preserves pathways common to other strains, including actin cytoskeleton organization, it involves fewer proteins. The reduced variability observed in Clade Ib in the absence of ISG15 further suggests that 2024 MPXV can evade ISG15-mediated restriction.

ISG15 has also been implicated in regulating protein phosphorylation, a critical process in both host signaling and viral replication (85)(86)(87). Phosphoprotein levels were reduced in Clade II–infected ISG15 KO MEFs relative to wild-type, with the effect most pronounced in the USA 2003 strain. Notably, the absence of ISG15 caused a striking reduction in phosphosites. The pathway analysis conducted on USA 2003 strain informed us that processes related to cell division had their phosphorylation reduced, correlating with the total protein pathway analysis. This highlights how ISG15 absence during MPXV infection causes an increase in mitotic cell cycle alterations. Other affected pathways included ubiquitination and proteosome mediated degradation, signaling that lack of ISG15 leads to dysfunctions in protein degradation which could be key in combating MPXV infection. Although 2024 MPXV presented less differences, ubiquitination was also an altered process, along with important cell division processes such as chromatin binding. While our current study did not directly assess phospho-proteomic changes, prior work suggests that ISG15 may influence kinase activity and substrate availability, potentially contributing to the altered phenotypes we observed (88)(89). Future work will address this using targeted phospho-specific assays.

Although murine systems do not fully recapitulate human MPXV infection, rodent models remain essential given their role as suspected viral reservoirs (90)(91). Importantly, murine and human ISG15 differ in function; human ISG15 can suppress continued ISG expression, whereas murine ISG15 does not (92)(93)(94). In humans, ISG15 deficiency often results in overactive immune responses that restrict viral replication—contrary to the enhanced infection seen in ISG15 KO MEFs (95). These species-specific effects underscore the importance of validating findings in human cells and tissues, particularly for viruses like poxviruses that encode potent immune evasion strategies (96)(97).

Recent studies suggest that standard laboratory mice are largely resistant to Clade IIb strains, possibly reflecting host adaptation toward humans (98)(99)(100). However, rats remain susceptible to infection, suggesting that murine resistance is strain-specific (101). Testing MPXV infection in ISG15 KO mouse strains (e.g., on C57BL/6 background) and comparing to permissive strains like CAST/EiJ could illuminate ISG15’s in vivo relevance. Recent studies have also widened our knowledge using other infection models, such as simian models simulating HIV and MPXV co-infection, neuronal organoids and human fibroblasts, allowing us to better comprehend differences in host infection dynamics (102) (103) (104).

Furthermore, ISGylomics approaches may help identify MPXV proteins targeted by ISG15, clarifying the mechanistic basis for its antiviral activity. ISG15 has already served as the basis for the development of broad mRNA based antivirals and been used as adjuvants in MVA based vaccines (105) (35) (36). This is of extreme necessity since recent evidence has displayed that existing antivirals are not very effective in countering MPXV infections (106). An example is Tecovirimitat, which displayed reduced effectivity in clinical trials and a low barrier to resistance (106)

(107). Recent progress in mpox therapeutics highlights monoclonal antibodies targeting MPXV proteins such as A35 and A16/G9, which provide enhanced protection during infection (108) (109). The use of approved drugs for other diseases has also been studied as an option, with research on plitidepsin displaying antiviral activity (110). Additional research to complement these finding would be essential to be prepared for future mpox outbreaks.

Together, our findings demonstrate that ISG15 modulates multiple aspects of MPXV infection, including intracellular replication, protein expression and host responses (Fig. 7). The enhanced replication and intracellular viral accumulation observed in ISG15-deficient cells establish ISG15 as a critical host factor restricting MPXV spread. We also reveal evolutionary differences between clades, with Clade Ib displaying reduced sensitivity to ISG15, potentially reflecting unique mechanisms that diminish its activity. Such features raise concern should Clade Ib strains continue to acquire enhanced human-to-human transmissibility. Defining the precise molecular targets of ISG15 may not only advance understanding of MPXV pathogenesis but also inform the development of ISG15-based antivirals or adjuvants.

**Figure 7.**
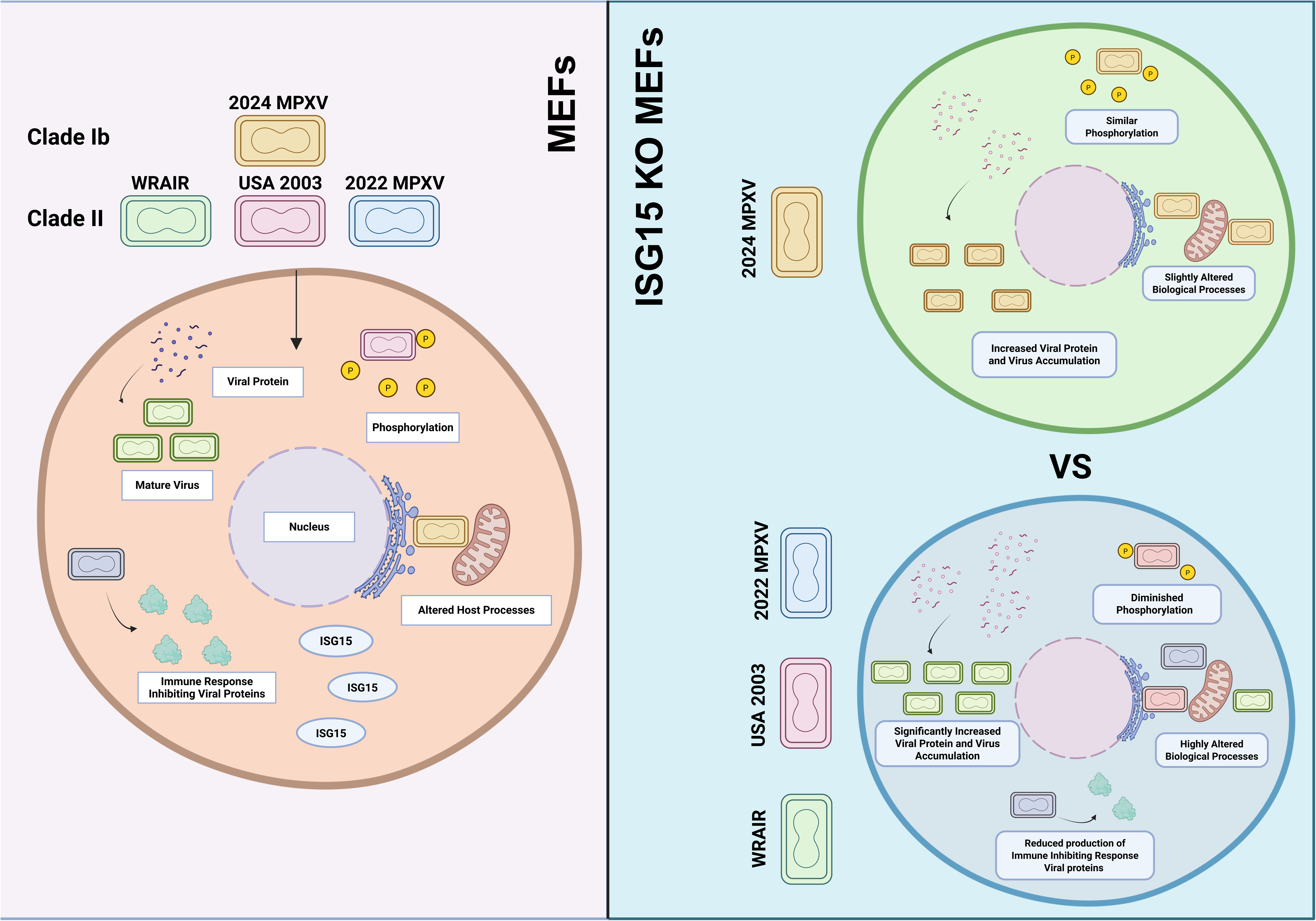
Schematic summary of phenotypic differences observed in Clade Ib and Clade II MPXV infected ISG15 KO MEFs. Created with BioRender (https://BioRender.com).

## ACKNOWLEDGEMENTS

We thank the ISIDORe consortium (Project *# ISID_55f9)*, Agencia Estatal Investigación and FEDER (PID2023-146351OB-I00, PID2020-117425RB-C21, PDC2021-121307-I00) for the funding of experiments. We thank Instruct-ERIC, Euro-BioImaging and Aryna Rybina for their role in management and support of unit services. We are also grateful for the help of the EMBL Electron Microscopy team (Viola Oorschot and Rachel Mellwig; Sample preparation and acquisition) and CNB Electron Microscopy team (Beatriz Martín Jouve and Juan Pablo; Sample preparation and acquisition). In addition, we would like to thank the Mount Sinai Pathogen Surveillance Program, Sara Sandoval and Celine Tárrega for their technical support during the development of these experiments, and Pilar Sánchez Gomez from the Instituto de Salud Carlos III for providing necessary infrastructure for the experiments.

## AUTHOR CONTRIBUTIONS

SG contributed to conception and design of the study. JPM and ILC performed the experiments. JPM, AT and SK performed the statistical analysis. APM, JDR, SG, MSS, AGS, SV and GP provided reagents. JPM and SG wrote the manuscript. All authors contributed to final manuscript revision, read and approved the submitted version.

## DATA AVAILABILITY STATEMENT

The data that support the findings of this study are available in the manuscript and in the supplementary material of this article. Proteomic and Phospho-proteomic data has been deposited in the PRIDE database (https://www.ebi.ac.uk/pride/) under the accession codes PXD068865. Source data is provided with this paper.

## REFERENCES

1. Martínez-Fernández DE, Fernández-Quezada D, Casillas-Muñoz FAG, Carrillo-Ballesteros FJ, Ortega-Prieto AM, Jimenez-Guardeño JM, Regla-Nava JA. 2023. Human Monkeypox: A Comprehensive Overview of Epidemiology, Pathogenesis, Diagnosis, Treatment, and Prevention Strategies. Pathogens 12:947.

2. Moss B. 2024. Understanding the biology of monkeypox virus to prevent future outbreaks. Nat Microbiol 9:1408–1416.

3. Bayer-Garner IB. 2005. Monkeypox virus: histologic, immunohistochemical and electron-microscopic findings. Journal of Cutaneous Pathology 32:28–34.

4. Schramm B, Locker JK. 2005. Cytoplasmic Organization of POXvirus DNA Replication. Traffic 6:839–846.

5. Yang Z, Gray M, Winter L. 2021. Why do poxviruses still matter? Cell & Bioscience 11:96.

6. Payne LG. 1980. Significance of extracellular enveloped virus in the in vitro and in vivo dissemination of vaccinia. J Gen Virol 50:89–100.

7. Law M, Smith GL. 2001. Antibody Neutralization of the Extracellular Enveloped Form of Vaccinia Virus. Virology 280:132–142.

8. Roberts KL, Smith GL. 2008. Vaccinia virus morphogenesis and dissemination. Trends in Microbiology 16:472–479.

9. Moss B. 2015. Poxvirus membrane biogenesis. Virology 479–480:619–626.

10. Roper RL, Wolffe EJ, Weisberg A, Moss B. 1998. The envelope protein encoded by the A33R gene is required for formation of actin-containing microvilli and efficient cell-to-cell spread of vaccinia virus. J Virol 72:4192–4204.

11. Schmidt FI, Bleck CKE, Helenius A, Mercer J. 2011. Vaccinia extracellular virions enter cells by macropinocytosis and acid-activated membrane rupture. EMBO J 30:3647–3661.

12. Bryk P, Brewer MG, Ward BM. 2018. Vaccinia Virus Phospholipase Protein F13 Promotes Rapid Entry of Extracellular Virions into Cells. J Virol 92:e02154–17.

13. Chen N, Li G, Liszewski MK, Atkinson JP, Jahrling PB, Feng Z, Schriewer J, Buck C, Wang C, Lefkowitz EJ, Esposito JJ, Harms T, Damon IK, Roper RL, Upton C, Buller RML. 2005. Virulence differences between monkeypox virus isolates from West Africa and the Congo basin. Virology 340:46–63.

14. Isidro J, Borges V, Pinto M, Sobral D, Santos JD, Nunes A, Mixão V, Ferreira R, Santos D, Duarte S, Vieira L, Borrego MJ, Núncio S, de Carvalho IL, Pelerito A, Cordeiro R, Gomes JP. 2022. Phylogenomic characterization and signs of microevolution in the 2022 multi-country outbreak of monkeypox virus. 8. Nat Med 28:1569–1572.

15. Patiño LH, Guerra S, Muñoz M, Luna N, Farrugia K, van de Guchte A, Khalil Z, Gonzalez-Reiche AS, Hernandez MM, Banu R, Shrestha P, Liggayu B, Firpo Betancourt A, Reich D, Cordon-Cardo C, Albrecht R, Pearl R, Simon V, Rooker A, Sordillo EM, van Bakel H, García-Sastre A, Bogunovic D, Palacios G, Paniz Mondolfi A, Ramírez JD. 2023. Phylogenetic landscape of Monkeypox Virus (MPV) during the early outbreak in New York City, 2022. Emerging Microbes & Infections 12:e2192830.

16. Gigante CM, Korber B, Seabolt MH, Wilkins K, Davidson W, Rao AK, Zhao H, Smith TG, Hughes CM, Minhaj F, Waltenburg MA, Theiler J, Smole S, Gallagher GR, Blythe D, Myers R, Schulte J, Stringer J, Lee P, Mendoza RM, Griffin-Thomas LA, Crain J, Murray J, Atkinson A, Gonzalez AH, Nash J, Batra D, Damon I, McQuiston J, Hutson CL, McCollum AM, Li Y. 2022. Multiple lineages of monkeypox virus detected in the United States, 2021–2022. Science 378:560–565.

17. Pekar JE, Wang Y, Wang JC, Shao Y, Taki F, Forgione LA, Amin H, Clabby T, Johnson K, Torian LV, Braunstein SL, Pathela P, Omoregie E, Hughes S, Suchard MA, Vasylyeva TI, Lemey P, Wertheim JO. 2025. Transmission dynamics of the 2022 mpox epidemic in New York City. Nat Med 10.1038/s41591-025-03526-9.

18. Vakaniaki EH, Kacita C, Kinganda-Lusamaki E, O’Toole Á, Wawina-Bokalanga T, Mukadi-Bamuleka D, Amuri-Aziza A, Malyamungu-Bubala N, Mweshi-Kumbana F, Mutimbwa-Mambo L, Belesi-Siangoli F, Mujula Y, Parker E, Muswamba-Kayembe P-C, Nundu SS, Lushima RS, Makangara-Cigolo J-C, Mulopo-Mukanya N, Pukuta-Simbu E, Akil-Bandali P, Kavunga H, Abdramane O, Brosius I, Bangwen E, Vercauteren K, Sam-Agudu NA, Mills EJ, Tshiani-Mbaya O, Hoff NA, Rimoin AW, Hensley LE, Kindrachuk J, Baxter C, de Oliveira T, Ayouba A, Peeters M, Delaporte E, Ahuka-Mundeke S, Mohr EL, Sullivan NJ, Muyembe-Tamfum J-J, Nachega JB, Rambaut A, Liesenborghs L, Mbala-Kingebeni P. 2024. Sustained human outbreak of a new MPXV clade I lineage in eastern Democratic Republic of the Congo. Nat Med 30:2791–2795.

19. Otieno JR, Ruis C, Onoja AB, Kuppalli K, Hoxha A, Nitsche A, Brinkmann A, Michel J, Mbala-Kingebeni P, Mukadi-Bamuleka D, Osman MM, Hussein H, Raja MA, Fotsing R, Herring BL, Keita M, Rico JM, Gresh L, Barakat A, Katawera V, Nahapetyan K, Naidoo D, Floto RA, Cunningham J, Van Kerkhove MD, Lewis RF, Subissi L. 2025. Global genomic surveillance of monkeypox virus. Nat Med 31:342–350.

20. Kibungu EM, Vakaniaki EH, Kinganda-Lusamaki E, Kalonji-Mukendi T, Pukuta E, Hoff NA, Bogoch II, Cevik M, Gonsalves GS, Hensley LE, Low N, Shaw SY, Schillberg E, Hunter M, Lunyanga L, Linsuke S, Madinga J, Peeters M, Cigolo J-CM, Ahuka-Mundeke S, Muyembe J-J, Rimoin AW, Kindrachuk J, Mbala-Kingebeni P, Lushima RS, International Mpox Research Consortium. 2024. Clade I-Associated Mpox Cases Associated with Sexual Contact, the Democratic Republic of the Congo. Emerg Infect Dis 30:172–176.

21. Mitjà O, Watson-Jones D, Choi EM, Jalloh MB, Sahr F. 2025. Clade IIb mpox outbreak in Sierra Leone. The Lancet 405:2274–2275.

22. Monzón S, Varona S, Negredo A, Vidal-Freire S, Patiño-Galindo JA, Ferressini-Gerpe N, Zaballos A, Orviz E, Ayerdi O, Muñoz-Gómez A, Delgado-Iribarren A, Estrada V, García C, Molero F, Sánchez-Mora P, Torres M, Vázquez A, Galán J-C, Torres I, Causse del Río M, Merino-Diaz L, López M, Galar A, Cardeñoso L, Gutiérrez A, Loras C, Escribano I, Alvarez-Argüelles ME, del Río L, Simón M, Meléndez MA, Camacho J, Herrero L, Jiménez P, Navarro-Rico ML, Jado I, Giannetti E, Kuhn JH, Sanchez-Lockhart M, Di Paola N, Kugelman JR, Guerra S, García-Sastre A, Cuesta I, Sánchez-Seco MP, Palacios G. 2024. Monkeypox virus genomic accordion strategies. Nat Commun 15:3059.

23. Wang L, Shang J, Weng S, Aliyari SR, Ji C, Cheng G, Wu A. 2023. Genomic annotation and molecular evolution of monkeypox virus outbreak in 2022. Journal of Medical Virology 95:e28036.

24. Zhang D, Zhang D-E. 2011. Interferon-Stimulated Gene 15 and the Protein ISGylation System. J Interferon Cytokine Res 31:119–130.

25. Perng Y-C, Lenschow DJ. 2018. ISG15 in antiviral immunity and beyond. Nat Rev Microbiol 16:423–439.

26. Dzimianski JV, Scholte FEM, Bergeron É, Pegan SD. 2019. ISG15: It’s Complicated. J Mol Biol 431:4203–4216.

27. Munnur D, Banducci-Karp A, Sanyal S. 2022. ISG15 driven cellular responses to virus infection. Biochem Soc Trans 50:1837–1846.

28. Álvarez E, Falqui M, Sin L, McGrail JP, Perdiguero B, Coloma R, Marcos-Villar L, Tárrega C, Esteban M, Gómez CE, Guerra S. 2024. Unveiling the Multifaceted Roles of ISG15: From Immunomodulation to Therapeutic Frontiers. Vaccines (Basel) 12:153.

29. Swaim CD, Scott AF, Canadeo LA, Huibregtse JM. 2017. Extracellular ISG15 Signals Cytokine Secretion through the LFA-1 Integrin Receptor. Mol Cell 68:581–590.e5.

30. Villarroya-Beltri C, Baixauli F, Mittelbrunn M, Fernández-Delgado I, Torralba D, Moreno-Gonzalo O, Baldanta S, Enrich C, Guerra S, Sánchez-Madrid F. 2016. ISGylation controls exosome secretion by promoting lysosomal degradation of MVB proteins. Nat Commun 7:13588.

31. Swaim CD, Canadeo LA, Monte KJ, Khanna S, Lenschow DJ, Huibregtse JM. 2020. Modulation of Extracellular ISG15 Signaling by Pathogens and Viral Effector Proteins. Cell Rep 31:107772.

32. Sainz B, Martín B, Tatari M, Heeschen C, Guerra S. 2014. ISG15 is a critical microenvironmental factor for pancreatic cancer stem cells. Cancer Res 74:7309–7320.

33. Albert M, Vázquez J, Falcón-Pérez JM, Balboa MA, Liesa M, Balsinde J, Guerra S. ISG15 Is a Novel Regulator of Lipid Metabolism during Vaccinia Virus Infection. Microbiol Spectr 10:e03893–22.

34. Nguyen H-M, Gaikwad S, Oladejo M, Agrawal MY, Srivastava SK, Wood LM. 2023. Interferon stimulated gene 15 (ISG15) in cancer: An update. Cancer Letters 556:216080.

35. Gómez CE, Perdiguero B, Falqui M, Marín MQ, Bécares M, Sorzano CÓS, García-Arriaza J, Esteban M, Guerra S. 2020. Enhancement of HIV-1 Env-Specific CD8 T Cell Responses Using Interferon-Stimulated Gene 15 as an Immune Adjuvant. J Virol 95:e01155–20.

36. Falqui M, Perdiguero B, Coloma R, Albert M, Marcos-Villar L, McGrail JP, Sorzano CÓS, Esteban M, Gómez CE, Guerra S. 2023. An MVA-based vector expressing cell-free ISG15 increases IFN-I production and improves HIV-1-specific CD8 T cell immune responses. Front Cell Infect Microbiol 13:1187193.

37. Guerra S, Cáceres A, Knobeloch K-P, Horak I, Esteban M. 2008. Vaccinia Virus E3 Protein Prevents the Antiviral Action of ISG15. PLOS Pathogens 4:e1000096.

38. Bécares M, Albert M, Tárrega C, Coloma R, Falqui M, Luhmann EK, Radoshevich L, Guerra S. ISG15 Is Required for the Dissemination of Vaccinia Virus Extracellular Virions. Microbiol Spectr 11:e04508–22.

39. McGrail JP, Mondolfi AP, Ramírez JD, Vidal S, García-Sastre A, Palacios G, Sanchez-Seco MP, Guerra S. 2024. Comparative Analysis of 2022 Outbreak MPXV and Previous Clade II MPXV. Journal of Medical Virology 96:e70023.

40. Paniz-Mondolfi A, Reidy J, Pagani N, Lednicky JA, McGrail JP, Kasminskaya Y, Patino LH, Garcia-Sastre A, Palacios G, Gonzalez-Reiche AS, van Bakel H, Firpo Betancourt A, Hernandez MM, Cordon-Cardo C, Simon V, Sordillo EM, Ramírez JD, Guerra S. 2023. Genomic and ultrastructural analysis of monkeypox virus in skin lesions and in human/animal infected cells reveals further morphofunctional insights into viral pathogenicity. J Med Virol 95:e28878.

41. Earl PL, Cooper N, Wyatt LS, Moss B, Carroll MW. 1998. Preparation of Cell Cultures and Vaccinia Virus Stocks. Current Protocols in Molecular Biology 43:16.16.1–16.16.13.

42. Demichev V, Messner CB, Vernardis SI, Lilley KS, Ralser M. 2020. DIA-NN: Neural networks and interference correction enable deep proteome coverage in high throughput. Nat Methods 17:41–44.

43. Virtanen P, Gommers R, Oliphant TE, Haberland M, Reddy T, Cournapeau D, Burovski E, Peterson P, Weckesser W, Bright J, van der Walt SJ, Brett M, Wilson J, Millman KJ, Mayorov N, Nelson ARJ, Jones E, Kern R, Larson E, Carey CJ, Polat İ, Feng Y, Moore EW, VanderPlas J, Laxalde D, Perktold J, Cimrman R, Henriksen I, Quintero EA, Harris CR, Archibald AM, Ribeiro AH, Pedregosa F, van Mulbregt P. 2020. SciPy 1.0: fundamental algorithms for scientific computing in Python. Nat Methods 17:261–272.

44. Seabold S, Perktold J. 2010. Statsmodels: Econometric and Statistical Modeling with Python, p. 92–96. *In* . Austin, Texas.

45. Ge SX, Jung D, Yao R. 2020. ShinyGO: a graphical gene-set enrichment tool for animals and plants. Bioinformatics 36:2628–2629.

46. Heberle H, Meirelles GV, da Silva FR, Telles GP, Minghim R. 2015. InteractiVenn: a web-based tool for the analysis of sets through Venn diagrams. BMC Bioinformatics 16:169.

47. Kuleshov MV, Xie Z, London ABK, Yang J, Evangelista JE, Lachmann A, Shu I, Torre D, Ma’ayan A. 2021. KEA3: improved kinase enrichment analysis via data integration. Nucleic Acids Res 49:W304–W316.

48. Pollak N, Dölle C, Ziegler M. 2007. The power to reduce: pyridine nucleotides – small molecules with a multitude of functions. Biochem J 402:205–218.

49. Baldanta S, Fernández-Escobar M, Acín-Perez R, Albert M, Camafeita E, Jorge I, Vázquez J, Enríquez JA, Guerra S. 2017. ISG15 governs mitochondrial function in macrophages following vaccinia virus infection. PLoS Pathog 13:e1006651.

50. Goicoechea SM, Arneman D, Otey CA. 2008. The role of palladin in actin organization and cell motility. Eur J Cell Biol 87:517–525.

51. Cannell IG, Sawicka K, Pearsall I, Wild SA, Deighton L, Pearsall SM, Lerda G, Joud F, Khan S, Bruna A, Simpson KL, Mulvey CM, Nugent F, Qosaj F, Bressan D, CRUK IMAXT Grand Challenge Team, Dive C, Caldas C, Hannon GJ. 2023. FOXC2 promotes vasculogenic mimicry and resistance to anti-angiogenic therapy. Cell Rep 42:112791.

52. Martinez-Martin N, Marcandalli J, Huang CS, Arthur CP, Perotti M, Foglierini M, Ho H, Dosey AM, Shriver S, Payandeh J, Leitner A, Lanzavecchia A, Perez L, Ciferri C. 2018. An Unbiased Screen for Human Cytomegalovirus Identifies Neuropilin-2 as a Central Viral Receptor. Cell 174:1158–1171.e19.

53. Lee DJ, Kim Y, Dinh PTN, Chung Y, Lee D, Kim Y, Lee SH, Choi I, Lee SH. 2023. Identification of Missense Variants Affecting Carcass Traits for Hanwoo Precision Breeding. Genes (Basel) 14:1839.

54. Cayrol C, Girard J-P. 2018. Interleukin-33 (IL-33): A nuclear cytokine from the IL-1 family. Immunological Reviews 281:154–168.

55. Chatterjee A, Seyfferth J, Lucci J, Gilsbach R, Preissl S, Böttinger L, Mårtensson CU, Panhale A, Stehle T, Kretz O, Sahyoun AH, Avilov S, Eimer S, Hein L, Pfanner N, Becker T, Akhtar A. 2016. MOF Acetyl Transferase Regulates Transcription and Respiration in Mitochondria. Cell 167:722–738.e23.

56. Ohbayashi N, Maruta Y, Ishida M, Fukuda M. 2012. Melanoregulin regulates retrograde melanosome transport through interaction with the RILP-p150Glued complex in melanocytes. J Cell Sci 125:1508–1518.

57. Russo MW, Sevetson BR, Milbrandt J. 1995. Identification of NAB1, a repressor of NGFI-A- and Krox20-mediated transcription. Proc Natl Acad Sci U S A 92:6873–6877.

58. Kyriakis JM. 2014. In the Beginning, There Was Protein Phosphorylation. Journal of Biological Chemistry 289:9460–9462.

59. Maller JL. 1993. On the importance of protein phosphorylation in cell cycle control. Mol Cell Biochem 127–128:267–281.

60. Zhu J, Liu G, Sayyad Z, Goins CM, Stauffer SR, Gack MU. ISGylation of the SARS-CoV-2 N protein by HERC5 impedes N oligomerization and thereby viral RNA synthesis. J Virol 98:e00869–24.

61. Zhang W, Liu HT. 2002. MAPK signal pathways in the regulation of cell proliferation in mammalian cells. Cell Res 12:9–18.

62. Chapagai D, Strebhardt K, Wyatt MD, McInnes C. 2025. Structural regulation of PLK1 activity: implications for cell cycle function and drug discovery. Cancer Gene Ther 32:608–621.

63. Du R, Huang C, Liu K, Li X, Dong Z. 2021. Targeting AURKA in Cancer: molecular mechanisms and opportunities for Cancer therapy. Mol Cancer 20:15.

64. Moro RN, Biswas U, Kharat SS, Duzanic FD, Das P, Stavrou M, Raso MC, Freire R, Chaudhuri AR, Sharan SK, Penengo L. 2023. Interferon restores replication fork stability and cell viability in BRCA-defective cells via ISG15. Nat Commun 14:6140.

65. Gnanaprakasam R, Keller M, Glassman R, El-Khoury MY, Chen DS, Feola N, Feldman J, Chaturvedi V. 2023. Mpox in the New York metropolitan area, Summer 2022. J Med Virol 95:e28699.

66. CDC. 2023. Mpox in the U.S. Centers for Disease Control and Prevention. https://www.cdc.gov/poxvirus/mpox/response/2022/world-map.html. Retrieved 21 March 2023.

67. Masirika LM, Udahemuka JC, Schuele L, Ndishimye P, Otani S, Mbiribindi JB, Marekani JM, Mambo LM, Bubala NM, Boter M, Nieuwenhuijse DF, Lang T, Kalalizi EB, Musabyimana JP, Aarestrup FM, Koopmans M, Oude Munnink BB, Siangoli FB. 2024. Ongoing mpox outbreak in Kamituga, South Kivu province, associated with monkeypox virus of a novel Clade I sub-lineage, Democratic Republic of the Congo, 2024. Euro Surveill 29:2400106.

68. Mathiang JR, Sahr F, Kamara IF, Ansumana R, Rogers MK, Lee SS, Bockarie MJ. 2025. Rapid Surge of Mpox in Sierra Leone: Public Health Challenges. International Journal of Infectious Diseases 0.

69. Alcamí A. Pathogenesis of the circulating mpox virus and its adaptation to humans. Proc Natl Acad Sci U S A 120:e2301662120.

70. Bunge EM, Hoet B, Chen L, Lienert F, Weidenthaler H, Baer LR, Steffen R. 2022. The changing epidemiology of human monkeypox—A potential threat? A systematic review. PLoS Negl Trop Dis 16:e0010141.

71. Ndodo N, Ashcroft J, Lewandowski K, Yinka-Ogunleye A, Chukwu C, Ahmad A, King D, Akinpelu A, Maluquer de Motes C, Ribeca P, Sumner RP, Rambaut A, Chester M, Maishman T, Bamidele O, Mba N, Babatunde O, Aruna O, Pullan ST, Gannon B, Brown CS, Ihekweazu C, Adetifa I, Ulaeto DO. 2023. Distinct monkeypox virus lineages co-circulating in humans before 2022. Nat Med 29:2317–2324.

72. O’Toole Á, Neher RA, Ndodo N, Borges V, Gannon B, Gomes JP, Groves N, King DJ, Maloney D, Lemey P, Lewandowski K, Loman N, Myers R, Omah IF, Suchard MA, Worobey M, Chand M, Ihekweazu C, Ulaeto D, Adetifa I, Rambaut A. 2023. APOBEC3 deaminase editing in mpox virus as evidence for sustained human transmission since at least 2016. Science 382:595–600.

73. Ulaeto DO, Dunning J, Carroll MW. 2022. Evolutionary implications of human transmission of monkeypox: the importance of sequencing multiple lesions. The Lancet Microbe 3:e639–e640.

74. Maluquer de Motes C, Ulaeto DO. 2025. Mpox poses an ever-increasing epidemic and pandemic risk. Nat Med 31:1743–1746.

75. Morales DJ, Lenschow DJ. 2013. The antiviral activities of ISG15. J Mol Biol 425:4995–5008.

76. Mcconnell SJ, Herman YF, Mattson DE, Erickson L. 1962. Monkey Pox Disease in Irradiated Cynomologous Monkeys. Nature 195:1128–1129.

77. Multistate Outbreak of Monkeypox --- Illinois, Indiana, and Wisconsin, 2003. https://www.cdc.gov/mmwr/preview/mmwrhtml/mm5223a1.htm. Retrieved 6 March 2024.

78. Reynolds MG, Yorita KL, Kuehnert MJ, Davidson WB, Huhn GD, Holman RC, Damon IK. 2006. Clinical Manifestations of Human Monkeypox Influenced by Route of Infection. The Journal of Infectious Diseases 194:773–780.

79. Di Giulio DB, Eckburg PB. 2004. Human monkeypox: an emerging zoonosis. Lancet Infect Dis 4:15–25.

80. Holley J, Sumner RP, Lant S, Ribeca P, Ulaeto D, Maluquer de Motes C. Engineered Promoter-Switched Viruses Reveal the Role of Poxvirus Maturation Protein A26 as a Negative Regulator of Viral Spread. J Virol 95:e01012–21.

81. Witt ASA, Trindade G de S, Souza FG de, Serafim MSM, da Costa AVB, Silva MVF, de Melo Iani FC, Rodrigues RAL, Kroon EG, Abrahão JS. 2023. Ultrastructural analysis of monkeypox virus replication in Vero cells. Journal of Medical Virology 95:e28536.

82. Peloponese J-M, Haller K, Miyazato A, Jeang K-T. 2005. Abnormal centrosome amplification in cells through the targeting of Ran-binding protein-1 by the human T cell leukemia virus type-1 Tax oncoprotein. Proceedings of the National Academy of Sciences 102:18974–18979.

83. Shumilov A, Tsai M-H, Schlosser YT, Kratz A-S, Bernhardt K, Fink S, Mizani T, Lin X, Jauch A, Mautner J, Kopp-Schneider A, Feederle R, Hoffmann I, Delecluse H-J. 2017. Epstein–Barr virus particles induce centrosome amplification and chromosomal instability. Nat Commun 8:14257.

84. Ploubidou A, Moreau V, Ashman K, Reckmann I, González C, Way M. 2000. Vaccinia virus infection disrupts microtubule organization and centrosome function. The EMBO Journal 19:3932–3944.

85. Okumura F, Okumura AJ, Uematsu K, Hatakeyama S, Zhang D-E, Kamura T. 2013. Activation of double-stranded RNA-activated protein kinase (PKR) by interferon-stimulated gene 15 (ISG15) modification down-regulates protein translation. J Biol Chem 288:2839–2847.

86. Park JH, Yang SW, Park JM, Ka SH, Kim J-H, Kong Y-Y, Jeon YJ, Seol JH, Chung CH. 2016. Positive feedback regulation of p53 transactivity by DNA damage-induced ISG15 modification. Nat Commun 7:12513.

87. Yángüez E, García-Culebras A, Frau A, Llompart C, Knobeloch K-P, Gutierrez-Erlandsson S, García-Sastre A, Esteban M, Nieto A, Guerra S. 2013. ISG15 Regulates Peritoneal Macrophages Functionality against Viral Infection. PLoS Pathog 9:e1003632.

88. Ganesan M, Poluektova LY, Tuma DJ, Kharbanda KK, Osna NA. 2016. Acetaldehyde Disrupts Interferon Alpha Signaling in Hepatitis C Virus Infected Liver Cells by Up-Regulating USP18. Alcohol Clin Exp Res 40:2329–2338.

89. Zhang M, Li J, Yan H, Huang J, Wang F, Liu T, Zeng L, Zhou F. 2021. ISGylation in Innate Antiviral Immunity and Pathogen Defense Responses: A Review. Front Cell Dev Biol 9.

90. Falendysz EA, Lopera JG, Rocke TE, Osorio JE. 2023. Monkeypox Virus in Animals: Current Knowledge of Viral Transmission and Pathogenesis in Wild Animal Reservoirs and Captive Animal Models. 4. Viruses 15:905.

91. Mbala-Kingebeni P, Rimoin AW, Kacita C, Liesenborghs L, Nachega JB, Kindrachuk J. 2024. The time is now (again) for mpox containment and elimination in Democratic Republic of the Congo. PLOS Glob Public Health 4:e0003171.

92. Bogunovic D, Byun M, Durfee LA, Abhyankar A, Sanal O, Mansouri D, Salem S, Radovanovic I, Grant AV, Adimi P, Mansouri N, Okada S, Bryant VL, Kong X-F, Kreins A, Velez MM, Boisson B, Khalilzadeh S, Ozcelik U, Darazam IA, Schoggins JW, Rice CM, Al-Muhsen S, Behr M, Vogt G, Puel A, Bustamante J, Gros P, Huibregtse JM, Abel L, Boisson-Dupuis S, Casanova J-L. 2012. Mycobacterial disease and impaired IFN-γ immunity in humans with inherited ISG15 deficiency. Science 337:1684–1688.

93. Hermann M, Bogunovic D. 2017. ISG15: In Sickness and in Health. Trends Immunol 38:79–93.

94. Martin-Fernandez M, Bravo García-Morato M, Gruber C, Murias Loza S, Malik MNH, Alsohime F, Alakeel A, Valdez R, Buta S, Buda G, Marti MA, Larralde M, Boisson B, Feito Rodriguez M, Qiu X, Chrabieh M, Al Ayed M, Al Muhsen S, Desai JV, Ferre EMN, Rosenzweig SD, Amador-Borrero B, Bravo-Gallego LY, Olmer R, Merkert S, Bret M, Sood AK, Al-rabiaah A, Temsah MH, Halwani R, Hernandez M, Pessler F, Casanova J-L, Bustamante J, Lionakis MS, Bogunovic D. 2020. Systemic Type I IFN Inflammation in Human ISG15 Deficiency Leads to Necrotizing Skin Lesions. Cell Reports 31:107633.

95. Speer SD, Li Z, Buta S, Payelle-Brogard B, Qian L, Vigant F, Rubino E, Gardner TJ, Wedeking T, Hermann M, Duehr J, Sanal O, Tezcan I, Mansouri N, Tabarsi P, Mansouri D, Francois-Newton V, Daussy CF, Rodriguez MR, Lenschow DJ, Freiberg AN, Tortorella D, Piehler J, Lee B, García-Sastre A, Pellegrini S, Bogunovic D. 2016. ISG15 deficiency and increased viral resistance in humans but not mice. Nat Commun 7:11496.

96. Hernaez B, Alcamí A. 2024. Poxvirus Immune Evasion. Annual Review of Immunology 42:551–584.

97. Meade N, Toreev HK, Chakrabarty RP, Hesser CR, Park C, Chandel NS, Walsh D. 2023. The poxvirus F17 protein counteracts mitochondrially orchestrated antiviral responses. Nat Commun 14:7889.

98. Earl PL, Americo JL, Moss B. 2012. Lethal Monkeypox Virus Infection of CAST/EiJ Mice Is Associated with a Deficient Gamma Interferon Response. J Virol 86:9105–9112.

99. Parker S, Buller RM. 2013. A review of experimental and natural infections of animals with monkeypox virus between 1958 and 2012. Future Virol 8:129–157.

100. Americo JL, Earl PL, Moss B. 2023. Virulence differences of mpox (monkeypox) virus clades I, IIa, and IIb.1 in a small animal model. Proceedings of the National Academy of Sciences 120:e2220415120.

101. Rockx B, Van Keulen L, Van Der Poel WHM, Ariën KK, Vercauteren K, Van De Water S, Harders-Westerveen J, Schreur PJW, Van Der Giessen J, Maas M, Stockhofe N, Liesenborghs L. 2024. Experimental monkeypox virus infection in rats. Cold Spring Harbor Laboratory 10.1101/2024.12.04.626788.

102. Zhang D, Liu J, Zhu L, Huang B, Cong Z, Li N, Zhang J, Chen T, Ma J, Lu J, Hou Y, Yang C, Peng W, Wei Q, Tan W, Yang J, Xue J. 2025. The pathogenicity and multi-organ proteomic profiles of Mpox virus infection in SIVmac239-infected rhesus macaques. Nat Commun 16:7653.

103. Huang Y, Bergant V, Grass V, Emslander Q, Hamad MS, Hubel P, Mergner J, Piras A, Krey K, Henrici A, Öllinger R, Tesfamariam YM, Dalla Rosa I, Bunse T, Sutter G, Ebert G, Schmidt FI, Way M, Rad R, Bowie AG, Protzer U, Pichlmair A. 2024. Multi-omics characterization of the monkeypox virus infection. Nat Commun 15:6778.

104. Schultz-Pernice I, Fahmi A, Brito F, Liniger M, Chiu Y-C, David T, Oliveira Esteves BI, Golomingi A, Zumkehr B, Gerber M, Jandrasits D, Züst R, Steiner S, Wotzkow C, Blank F, Engler OB, Summerfield A, Ruggli N, Baud D, Alves MP. 2025. Monkeypox virus spreads from cell-to-cell and leads to neuronal death in human neural organoids. Nat Commun 16:5376.

105. Akalu YT, Patel RS, Taft J, Canas-Arranz R, Geltman R, Richardson A, Buta S, Martin-Fernandez M, Sazeides C, Pearl RL, Mainkar G, Kurland AP, Rosberger H, Kang DD, Kurian AA, Kaur K, Altman J, Dong Y, Johnson JR, Zangi L, Lim JK, Albrecht RA, García-Sastre A, Rosenberg BR, Bogunovic D. 2025. An mRNA-based broad-spectrum antiviral inspired by ISG15 deficiency protects against viral infections in vitro and in vivo. Science Translational Medicine 17:eadx5758.

106. Lenharo M. 2024. Hopes dashed for drug aimed at monkeypox virus spreading in Africa. Nature 632:965.

107. Smith TG, Gigante CM, Wynn NT, Matheny A, Davidson W, Yang Y, Condori RE, O’Connell K, Kovar L, Williams TL, Yu YC, Petersen BW, Baird N, Lowe D, Li Y, Satheshkumar PS, Hutson CL. 2023. Tecovirimat Resistance in Mpox Patients, United States, 2022-2023. Emerg Infect Dis 29:2426–2432.

108. Fantin RF, Yuan M, Park S-C, Bozarth B, Cohn H, Ignacio M, Earl P, Civljak A, Laghlali G, Zhang D, Zhu X, Crandell J, Monteiro V, Clark JJ, Cotter C, Burkhardt M, Singh G, Warang P, Diego JG-B, Srivastava K, Lugo LA, Pischel L, Yildirim I, Omer SB, Silva D da, Krammer F, Bajic G, Simon V, Schotsaert M, Lucas C, Wilson IA, Moss B, Coelho CH. 2025. Human monoclonal antibodies targeting A35 protect from death caused by mpox. Cell 0.

109. Meola A, Vernuccio R, Battini L, Albericio G, Delgado P, Bamford R, Pokorny L, Broutin M, León AM, Gallien S, Gil M, Noriega MA, Guivel-Benhassine F, Porrot F, Postal J, Buchrieser J, Hubert M, Haouz A, Lafaye P, Esteban M, Hub JS, Mahévas M, Chappert P, Mercer J, Garcia-Arriaza J, Schwartz O, Guardado-Calvo P. 2025. Structural basis of poxvirus fusion regulation and anti-A16/G9 antibody-mediated neutralization and protection. Cell 0.

110. Albericio G, Rodríguez-Martín D, Avilés P, Cuevas C, Guillén-Navarro MJ, Noriega MA, Flores S, Sánchez-Cordón PJ, Astorgano D, Pérez P, Esteban M, García-Arriaza J. 2025. Functional characteristics of plitidepsin as an antiviral treatment against monkeypox virus infection. Antiviral Res 241:106238.

