## Supplementary figures and images for "ISG15 Differentially Modulates Clade Ib and II MPXV Infection in MEF cells"

### S1

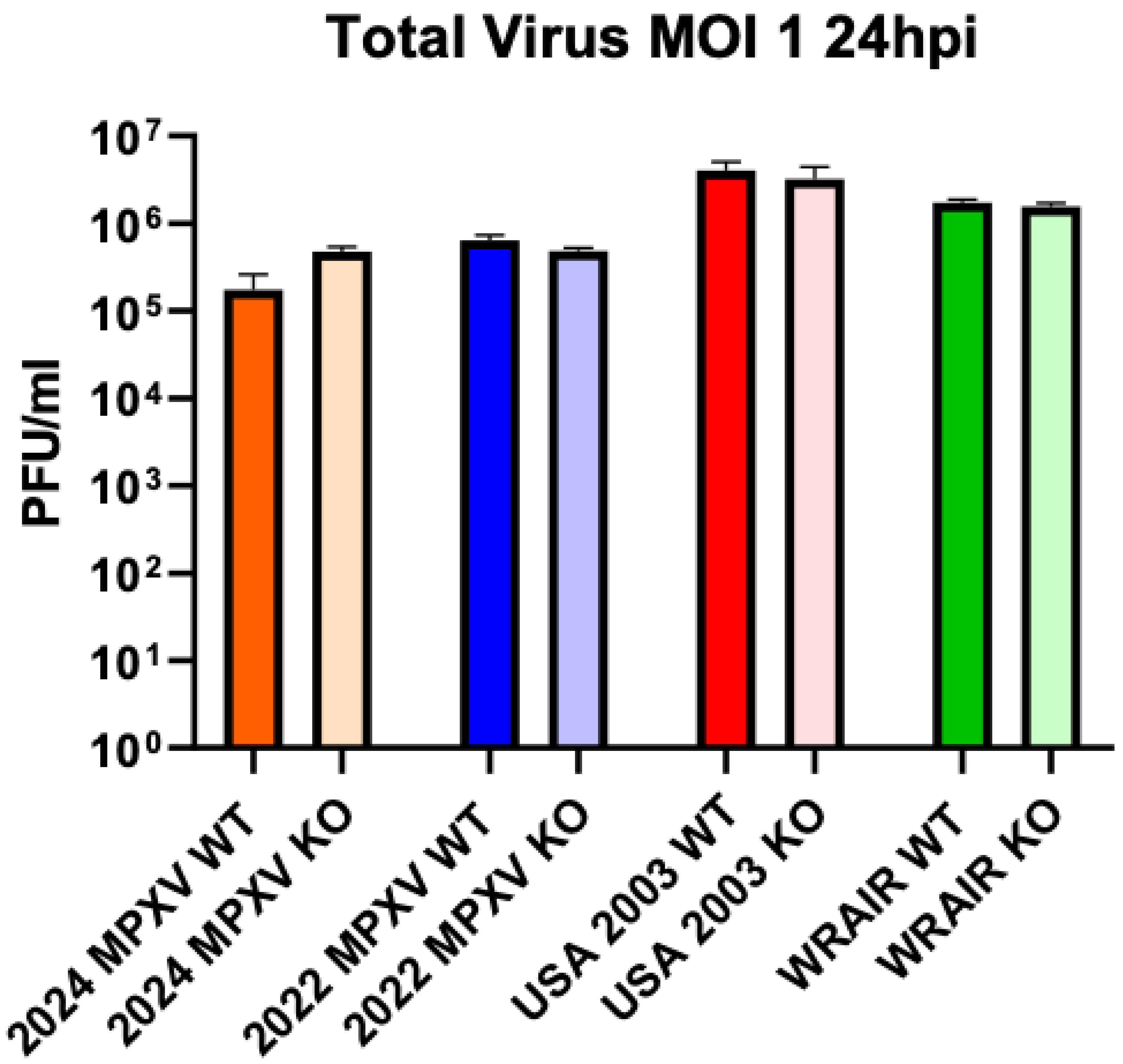

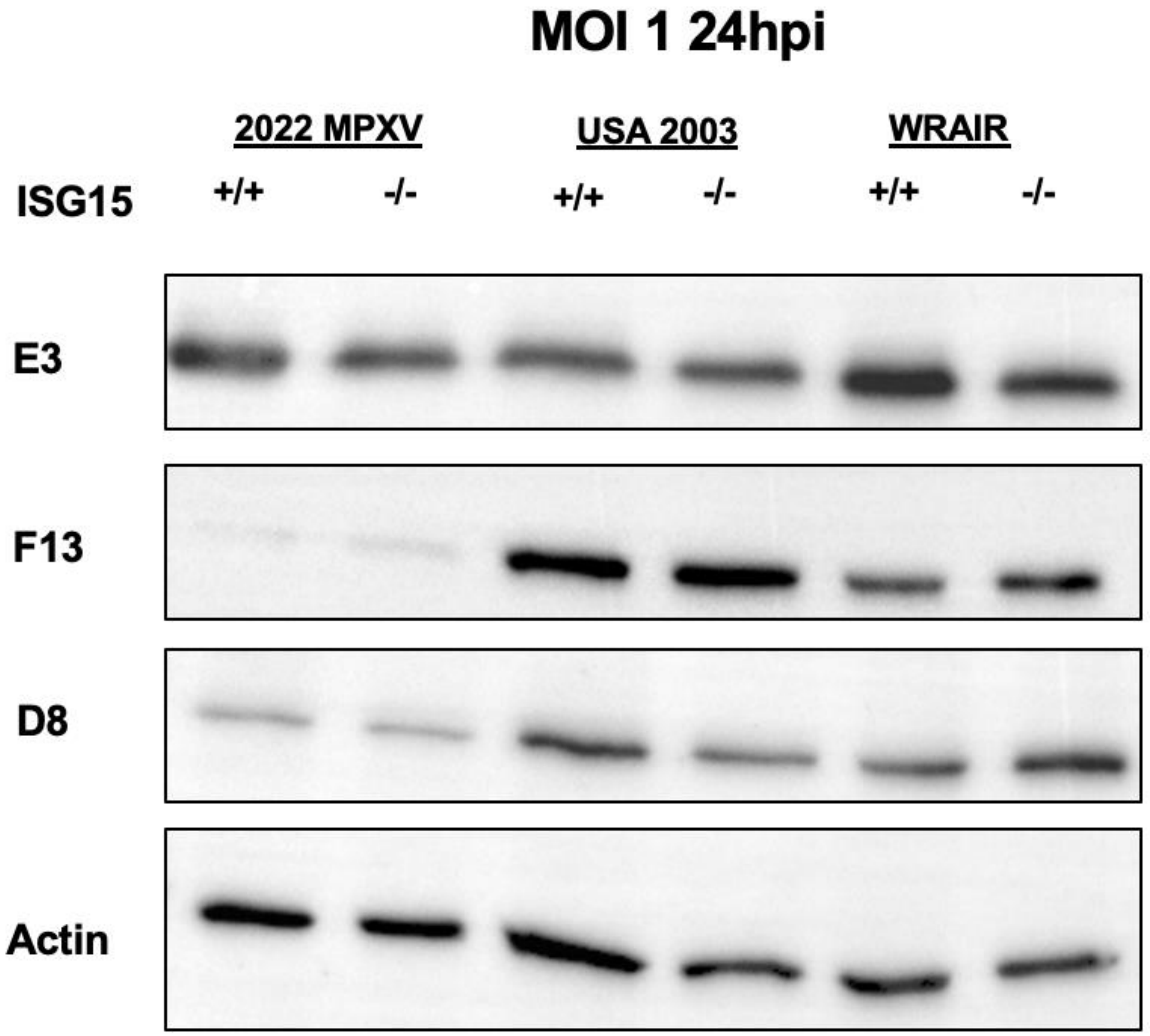

### S3

#### Cellular Component Network MPXV KOvsWT

**2024 MPXV**

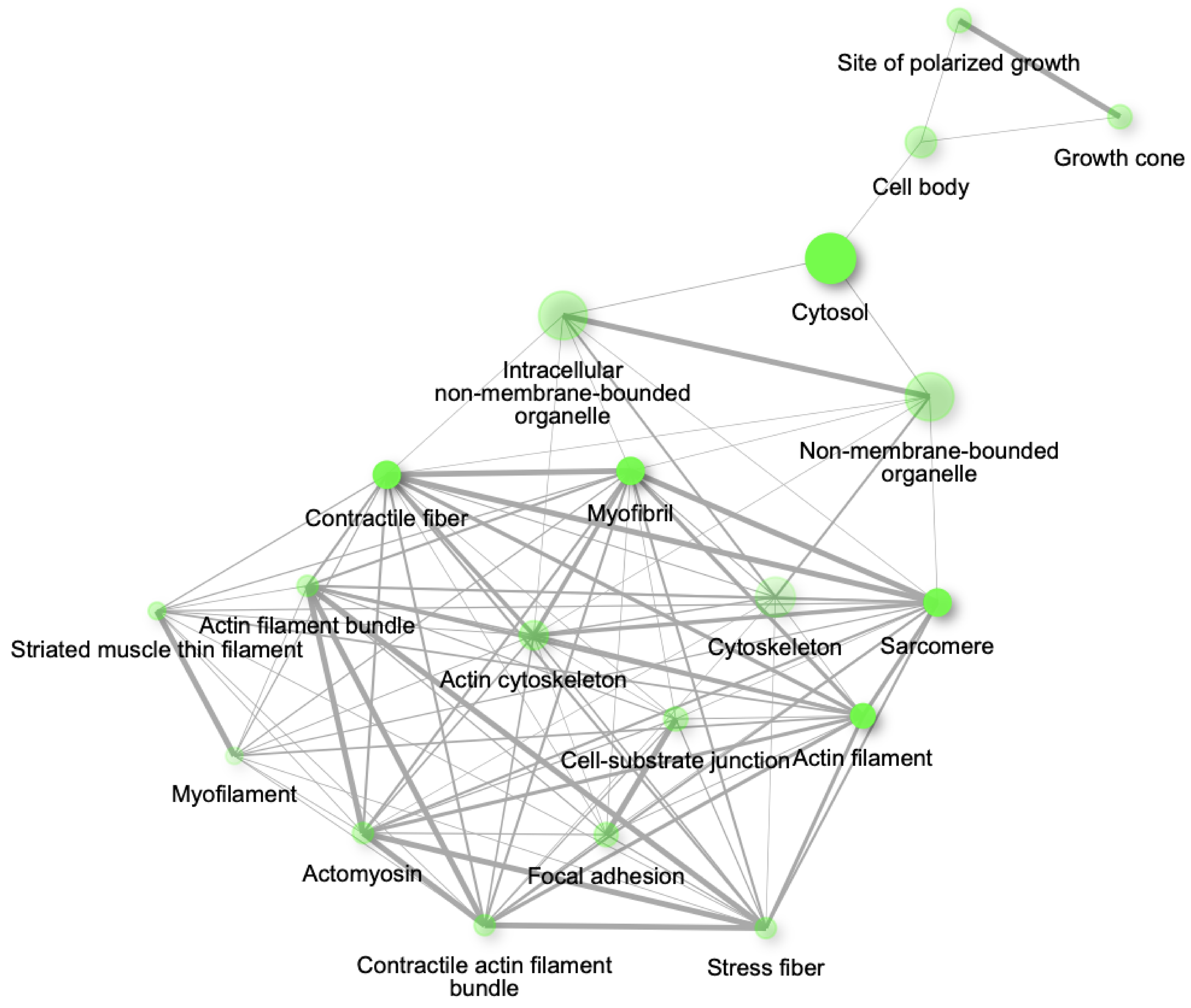


**USA 2003**

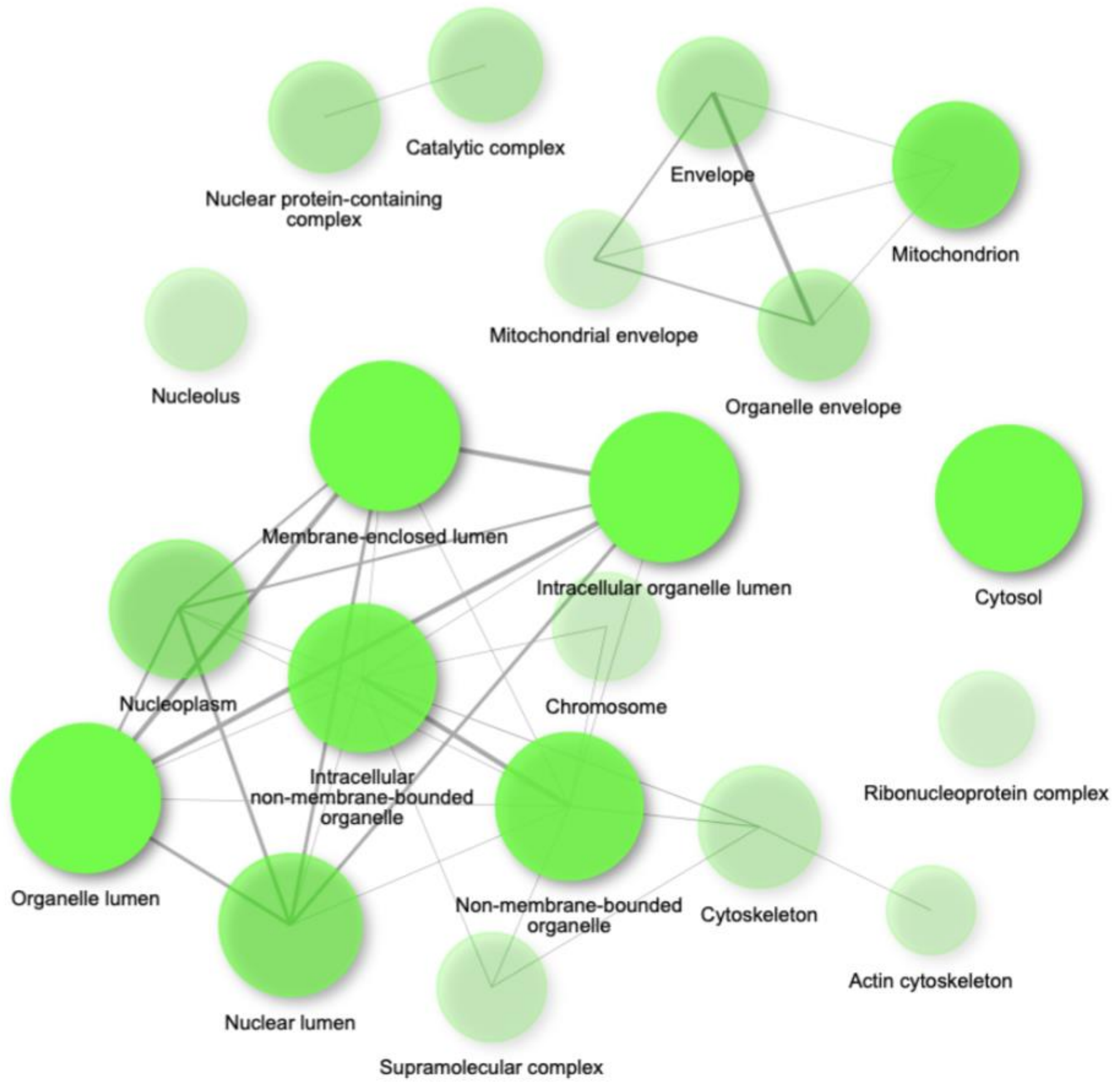


**2022 MPXV**

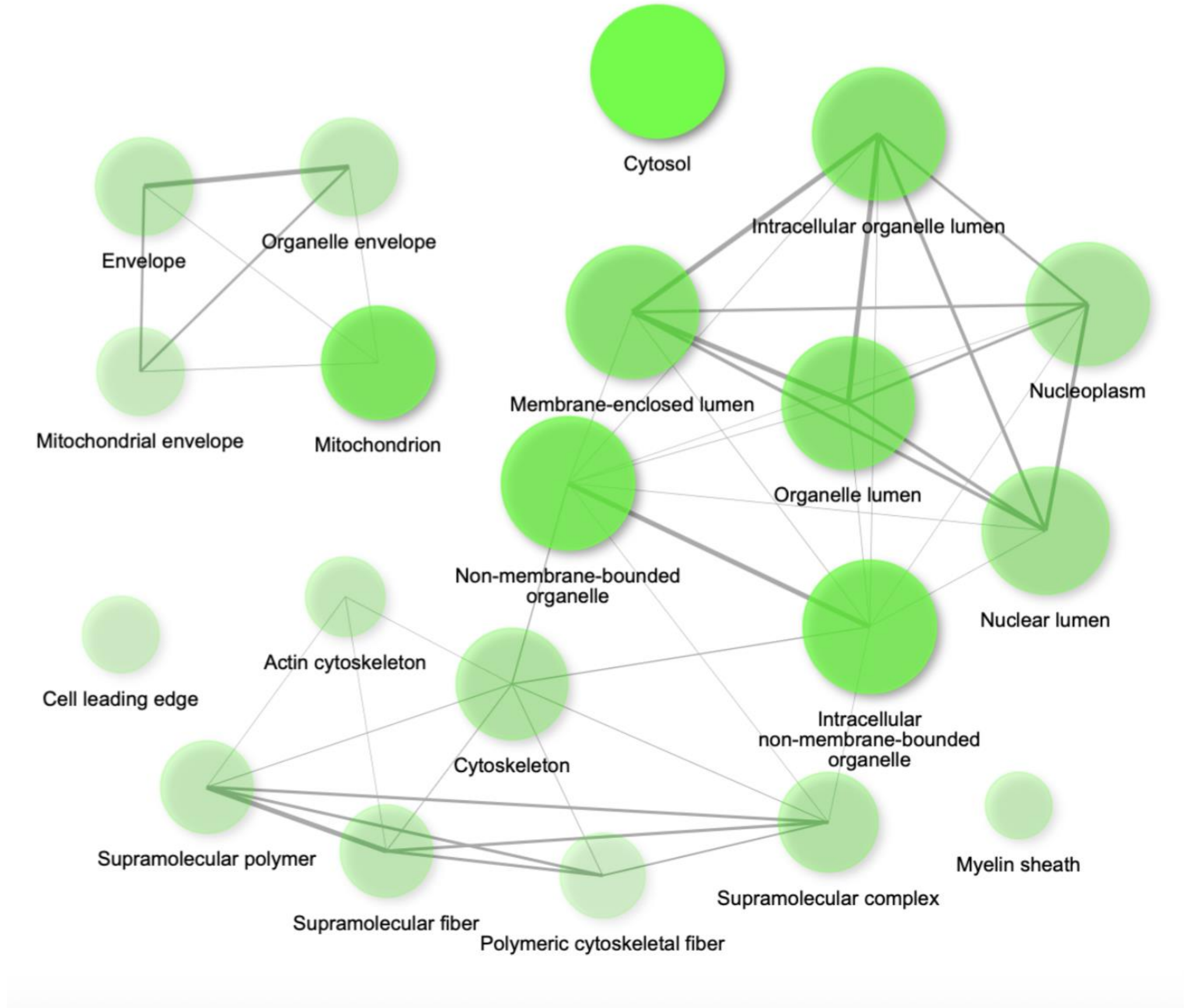


**WRAIR**

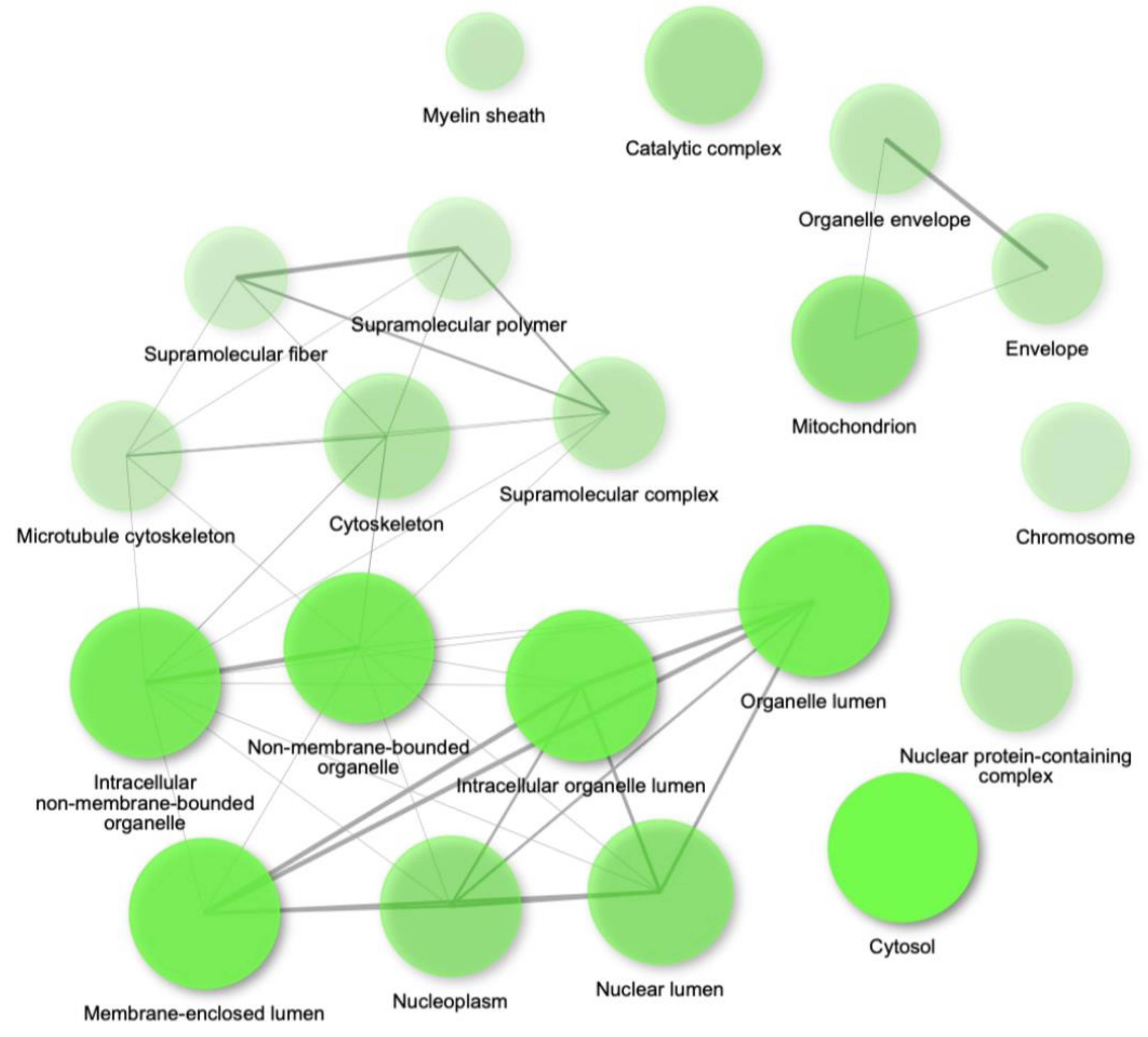

### S4

#### Protein-Protein Interactions MPXV KOvsWT (100 Proteins)

**2022 MPXV**

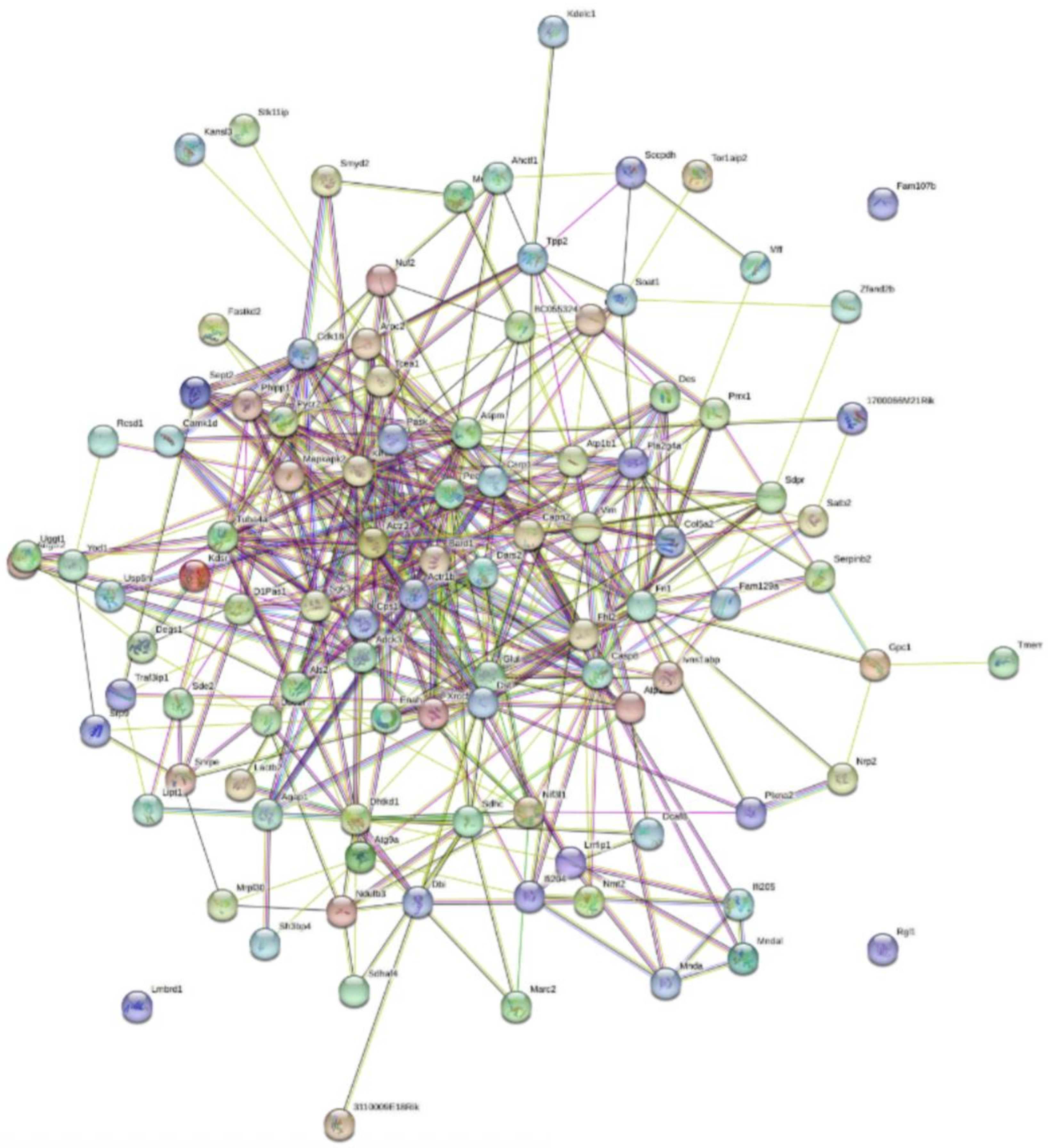

**USA 2003**

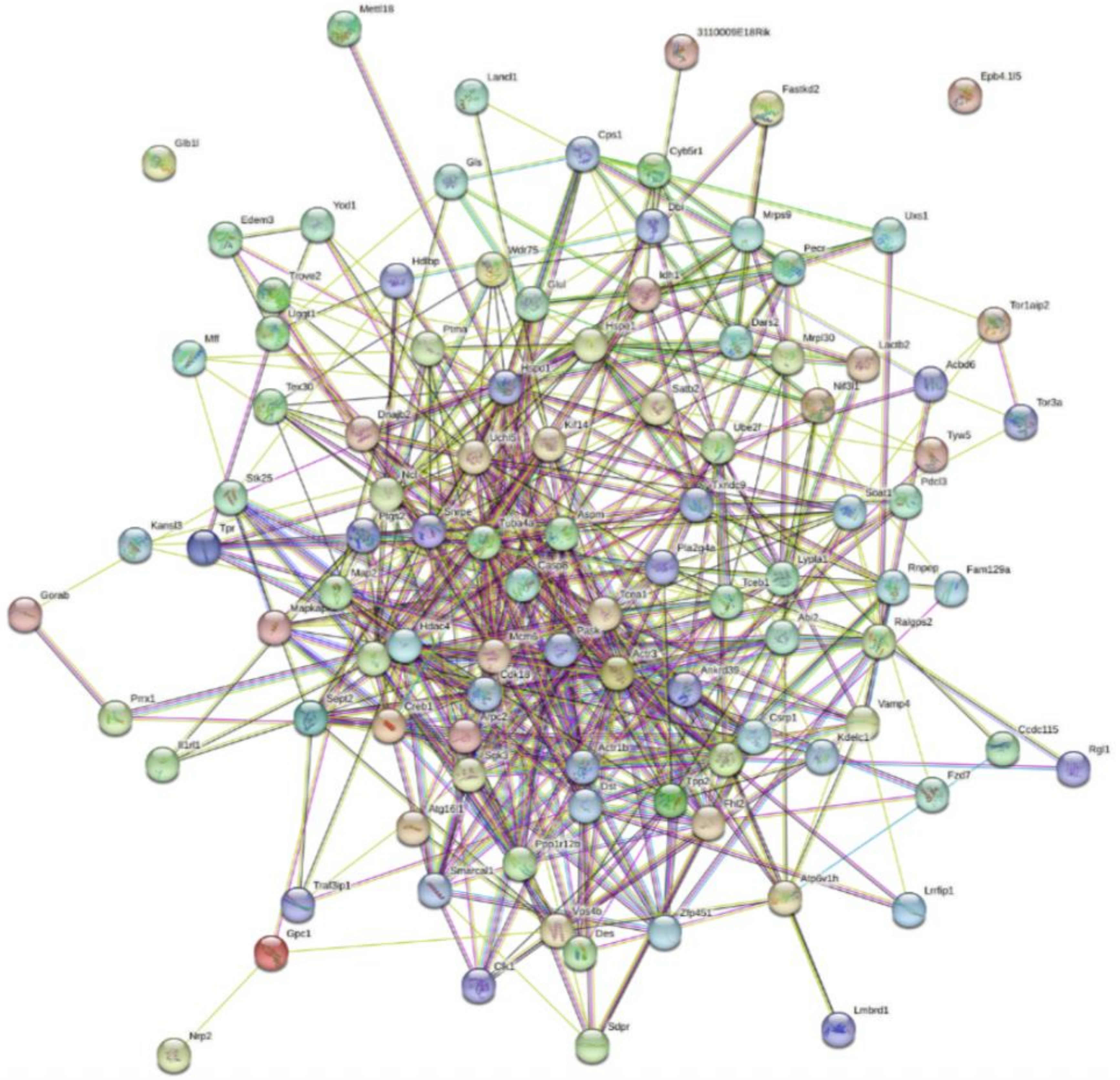

**WRAIR**

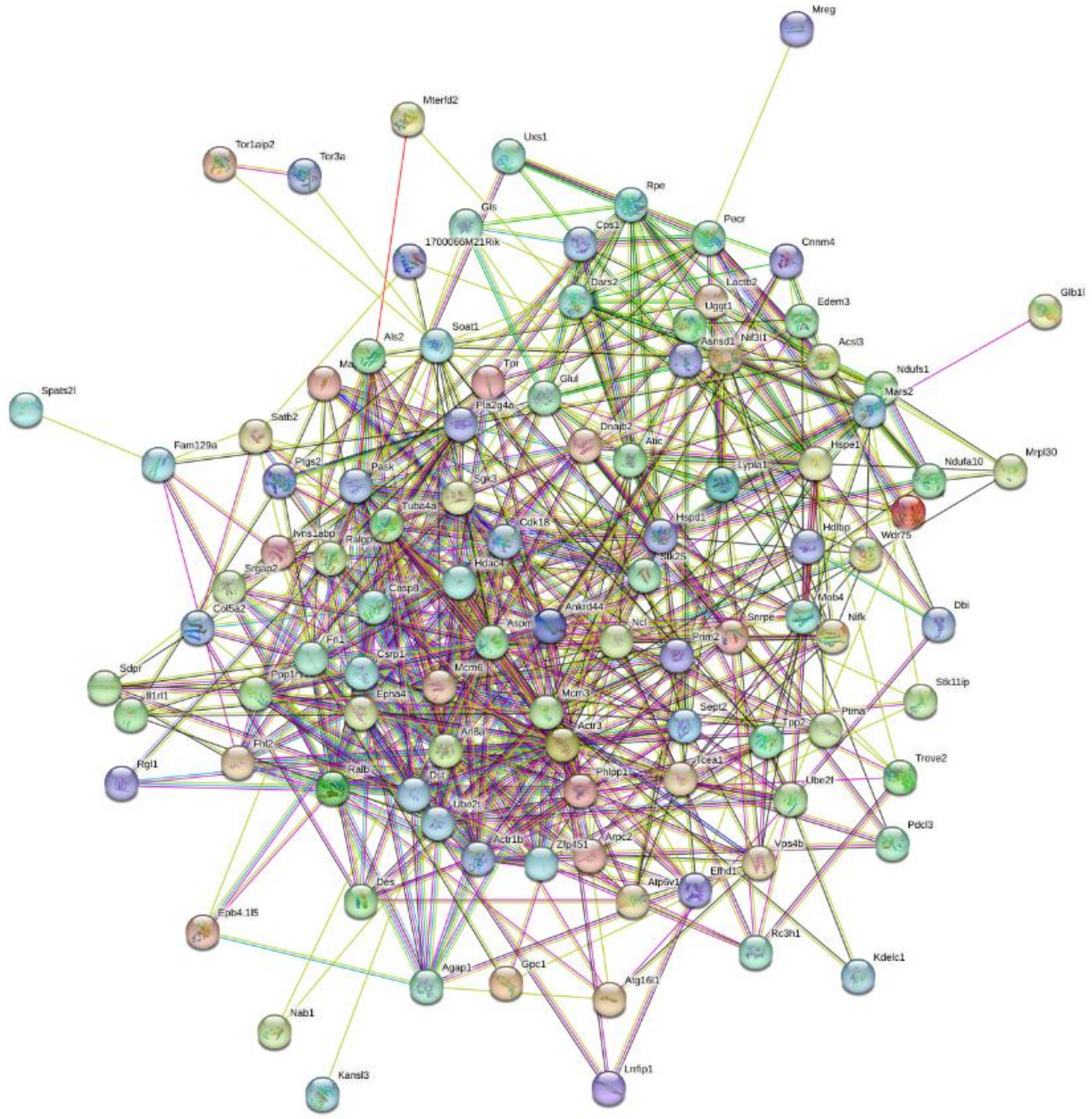

### S6

#### Phosphoproteomic Pathway Analysis KOvsWT

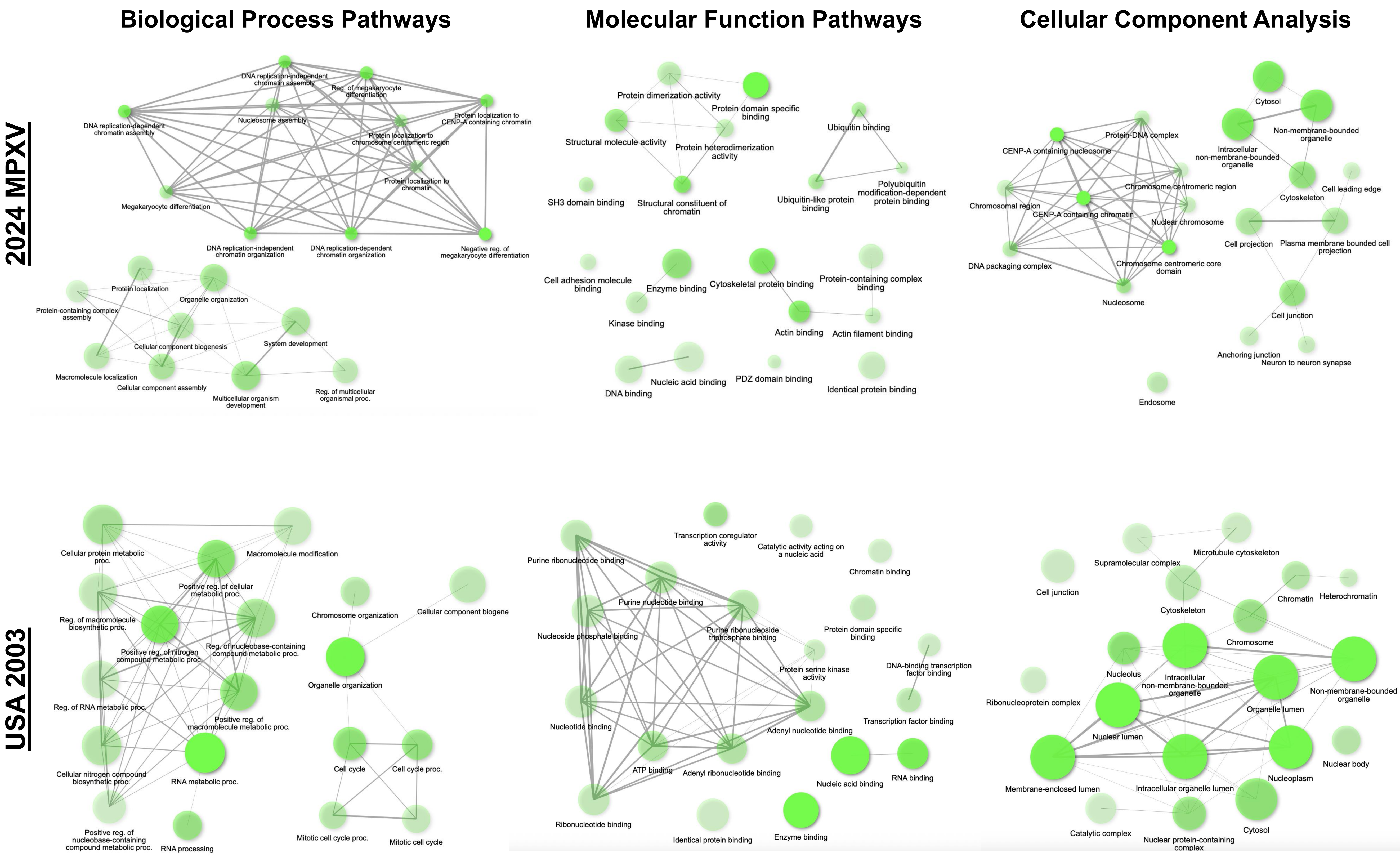

### S7

#### Clade II MPXV Phosphoproteomic QC Graphs

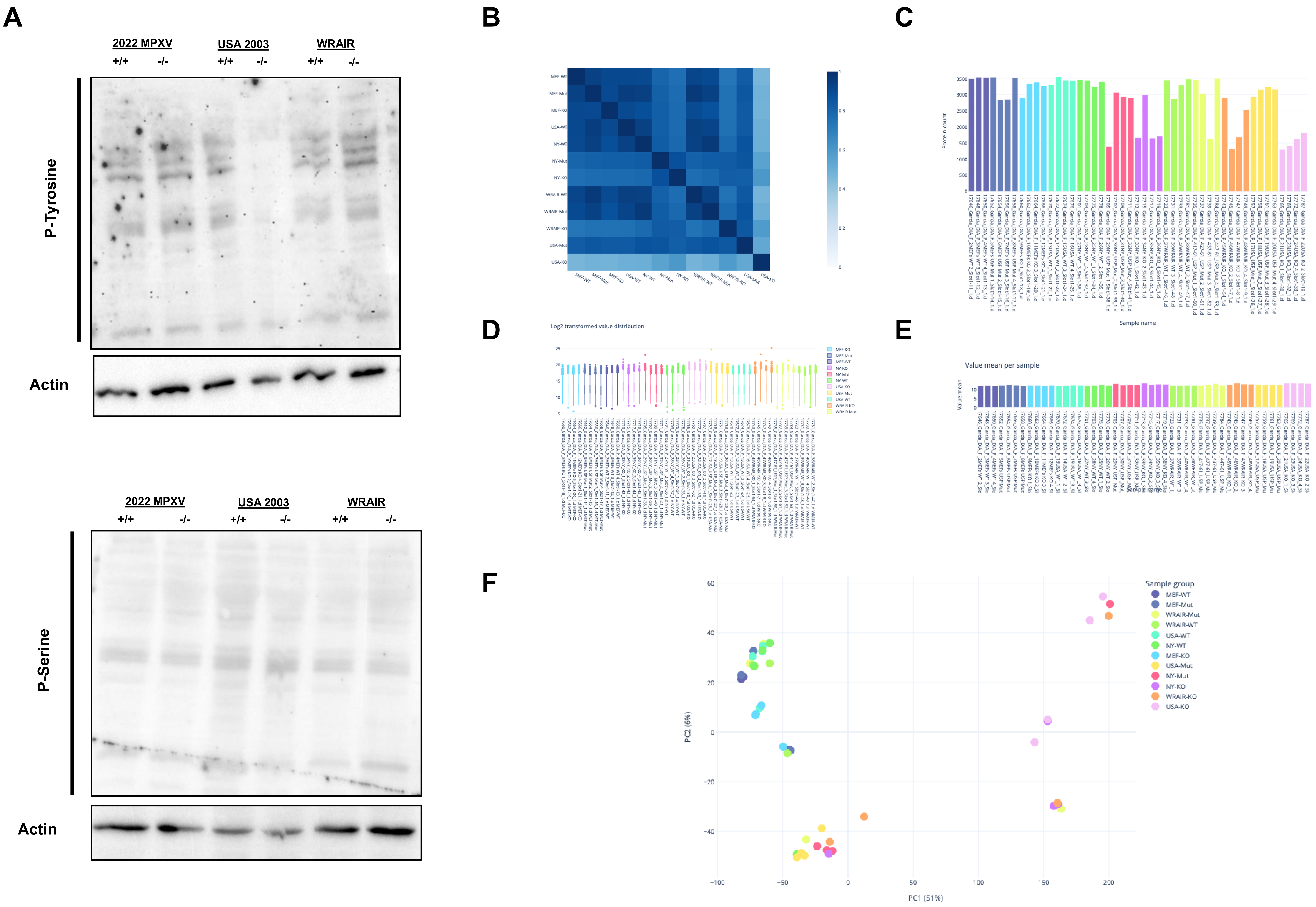

### S8

#### 2024 MPXV Phosphoproteomic QC Graphs

A

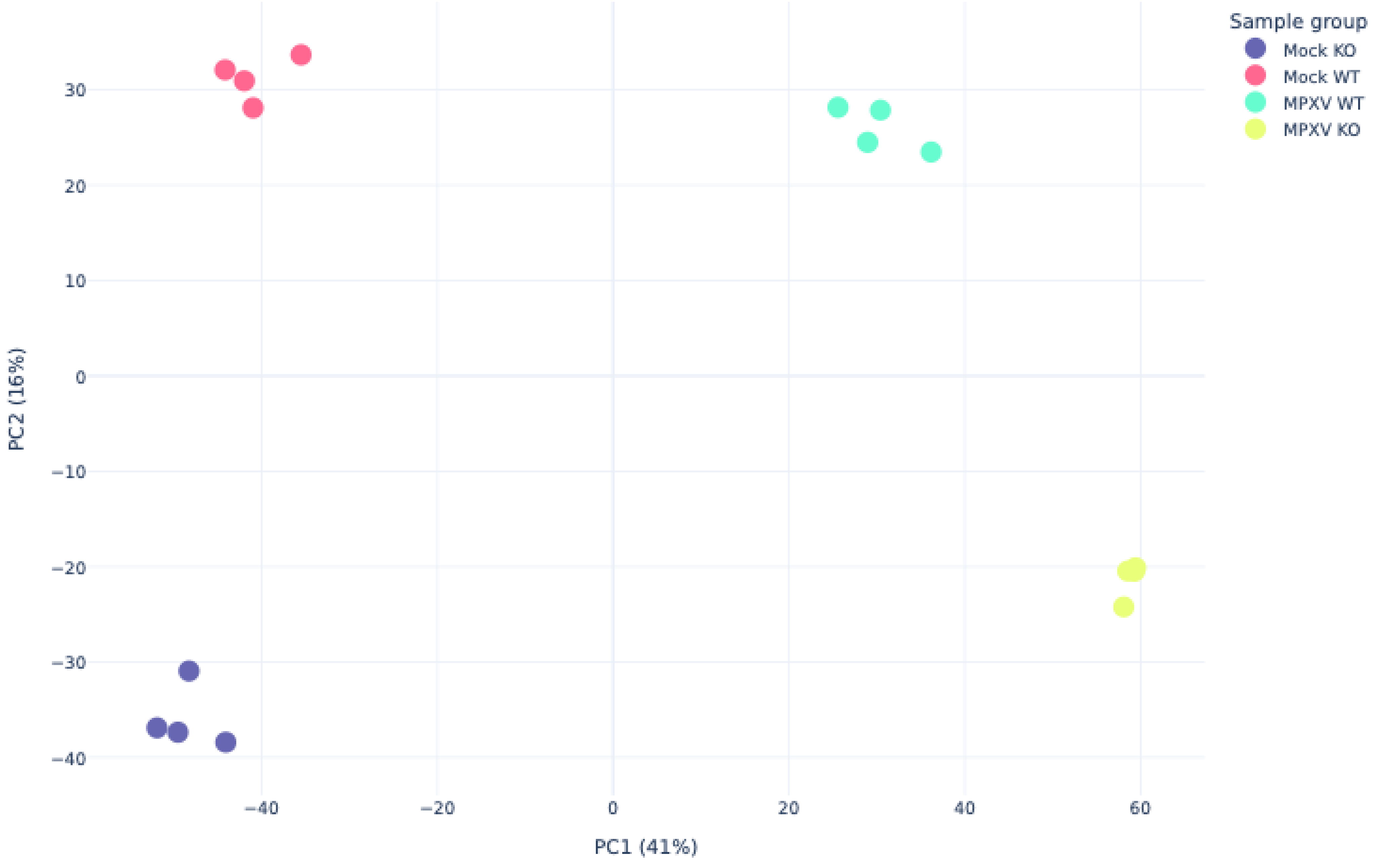


B

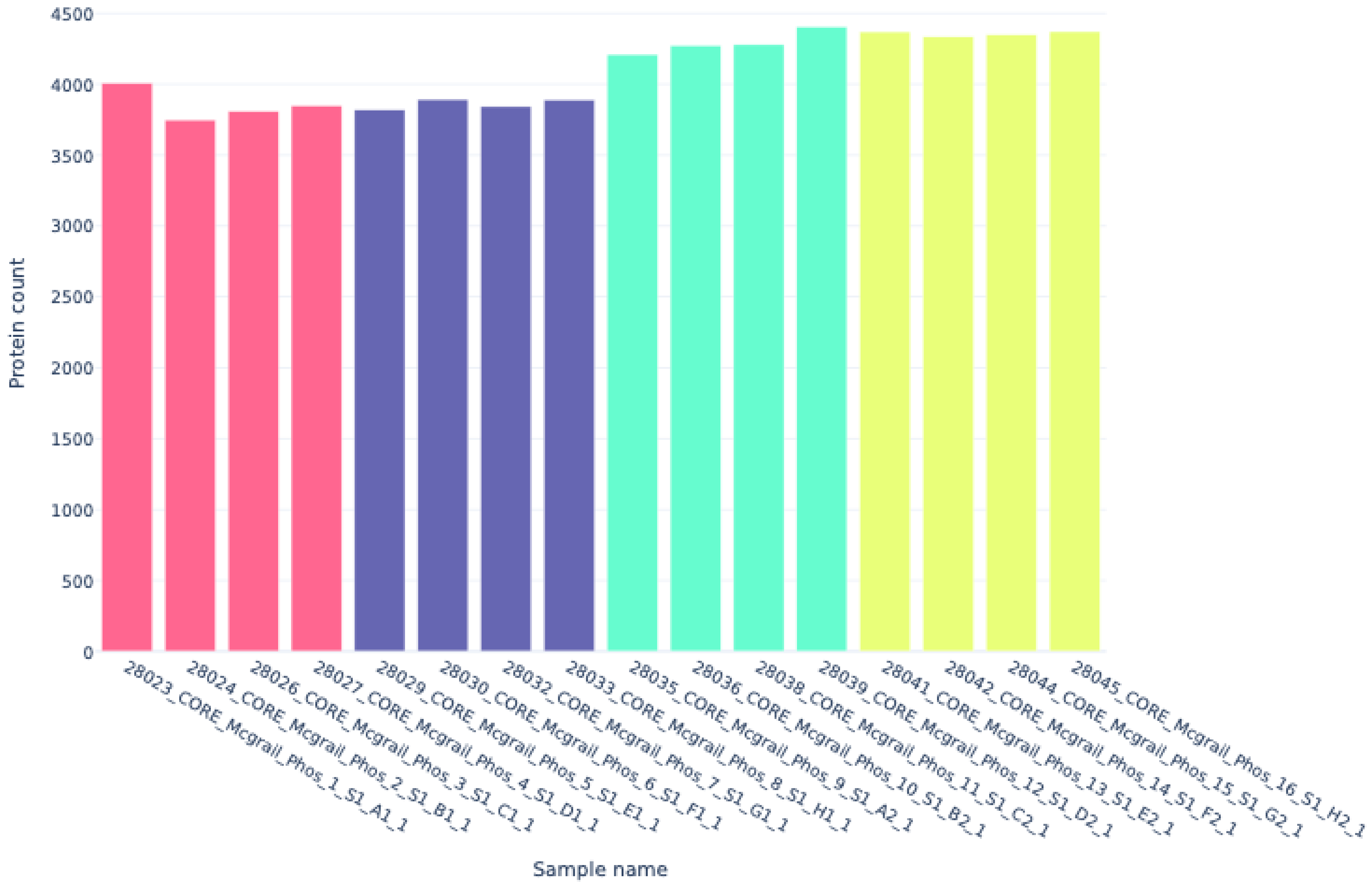


C

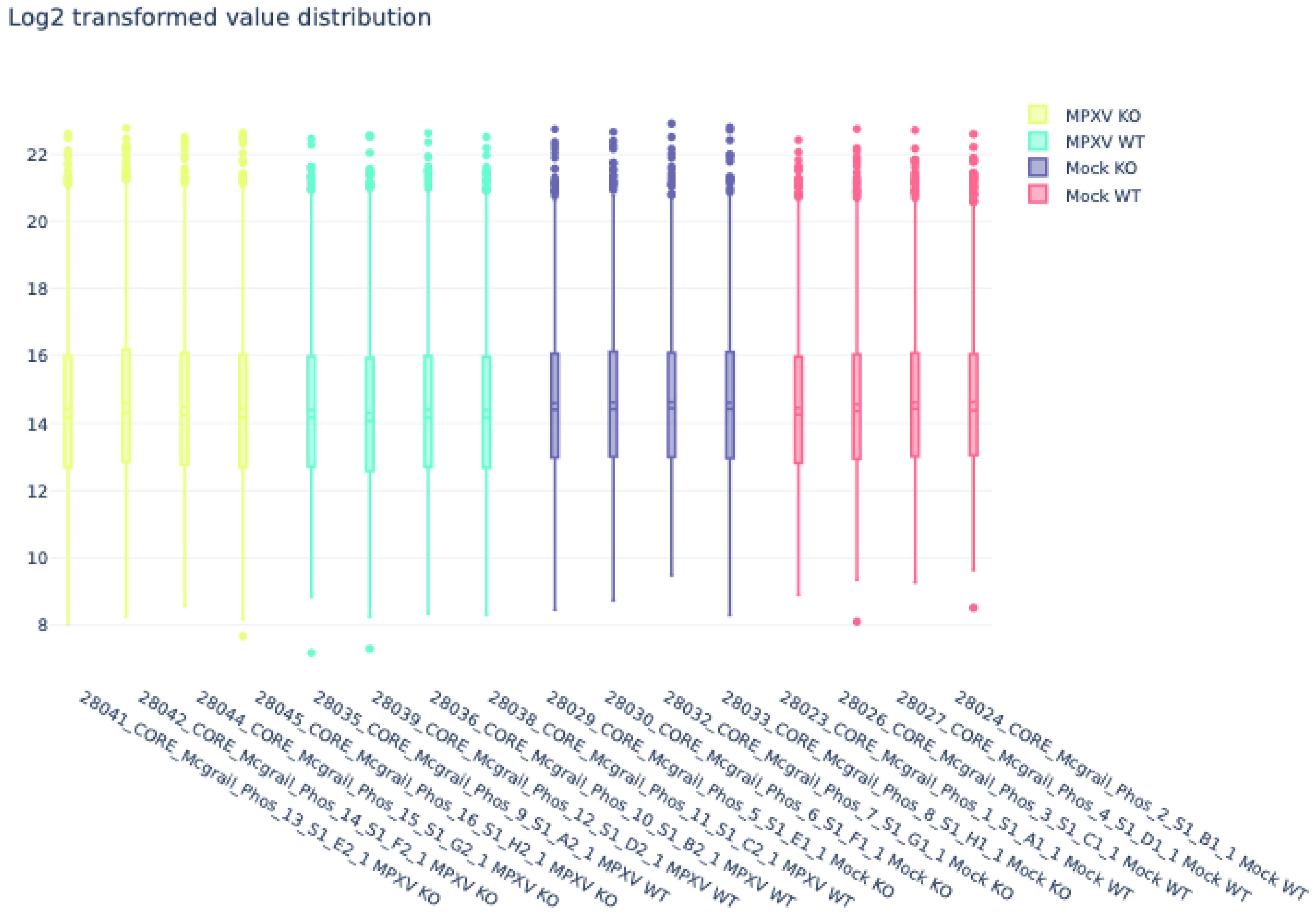


D

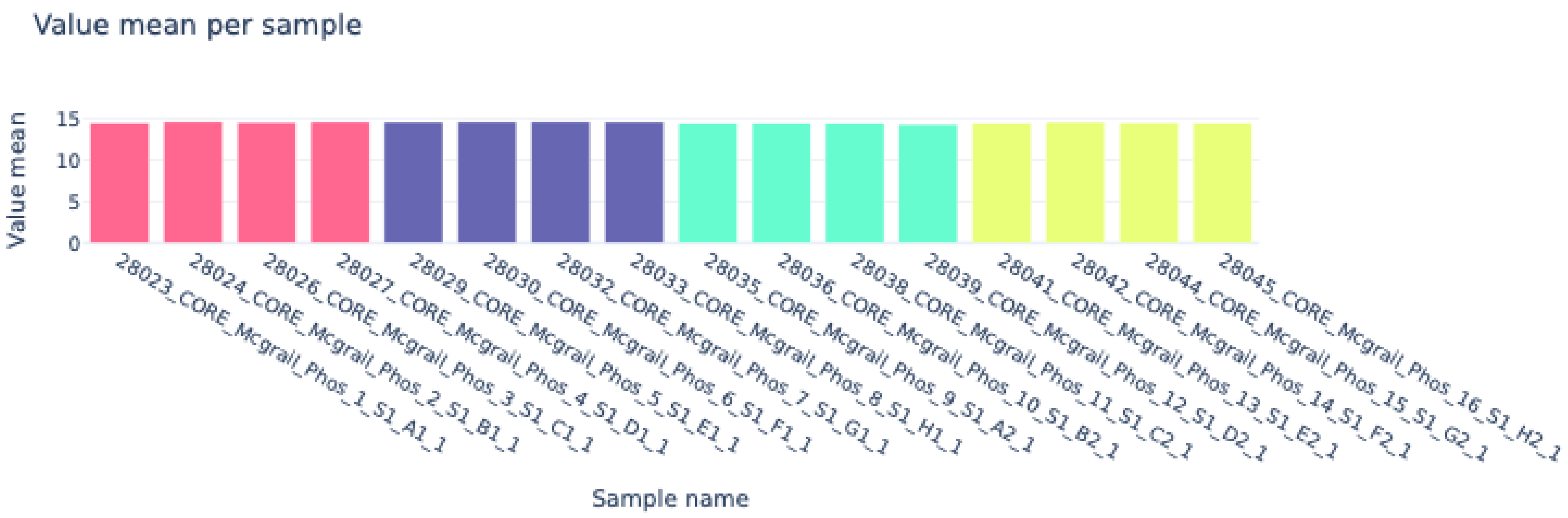

### S9

#### Kinase Enrichment Analysis KOvsWT

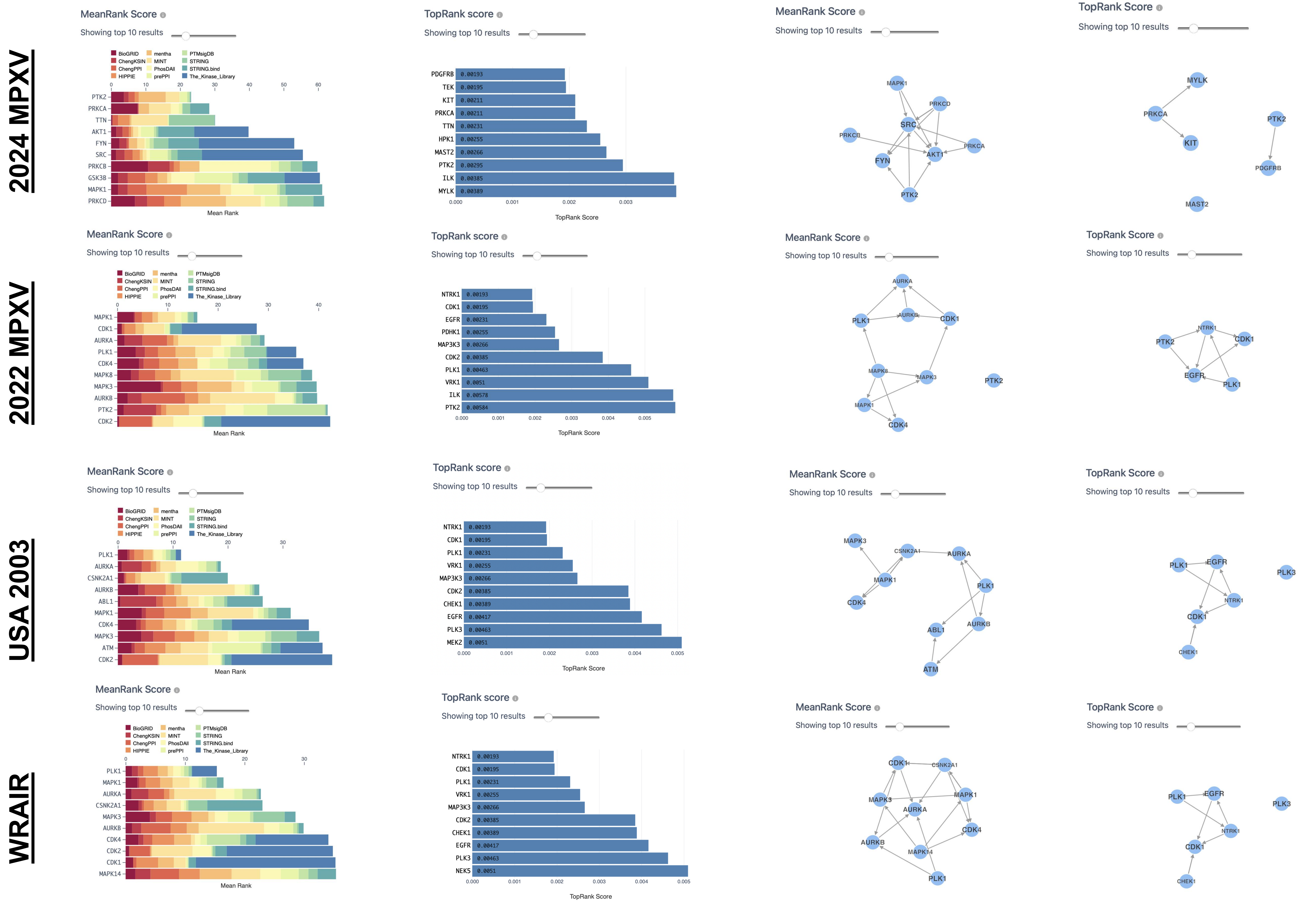

### S10

#### Phosphoproteomic Kinase Enrichment Analysis KOvsWT

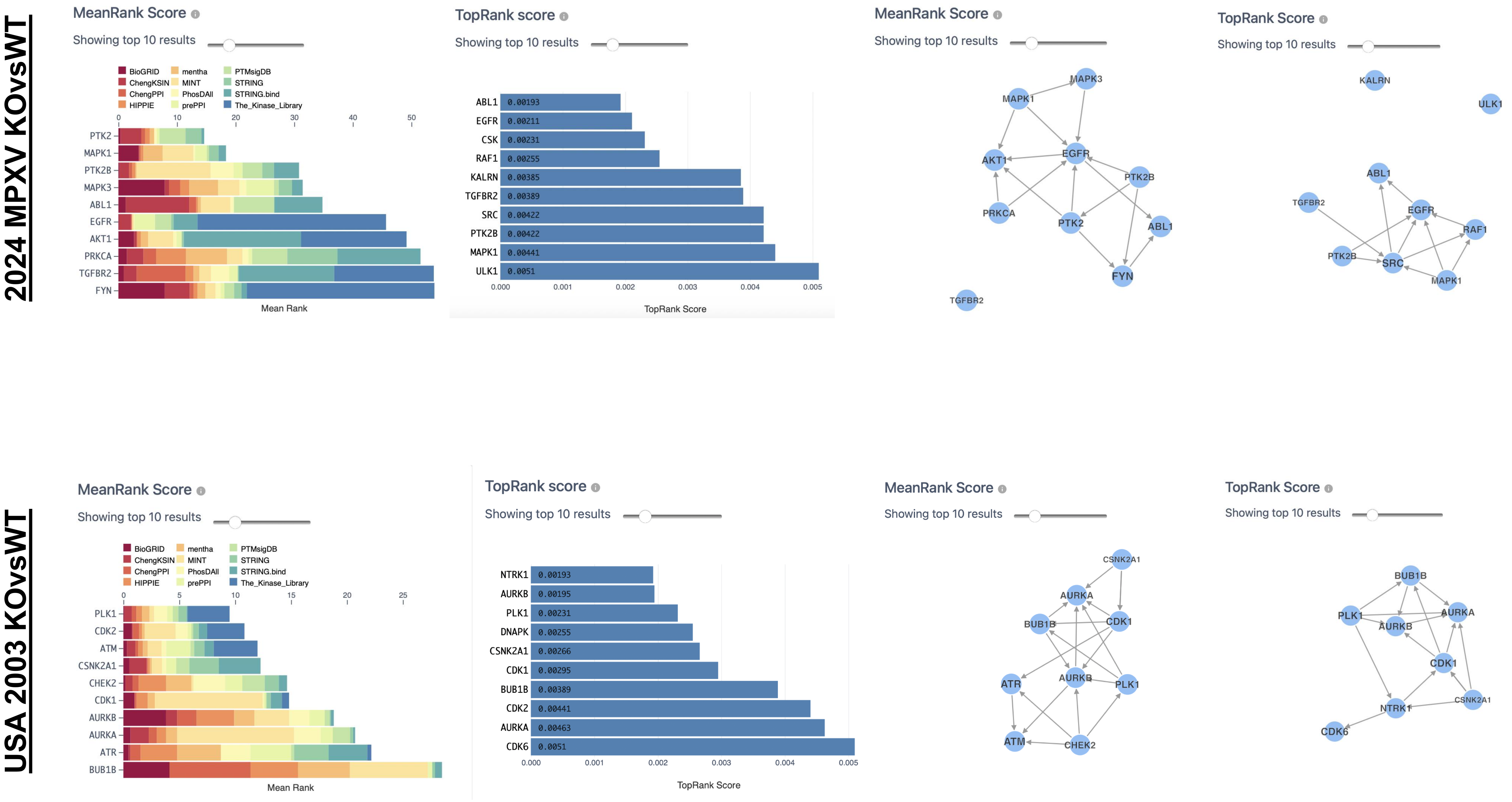
