## Supplementary material for "ISG15 Differentially Modulates Clade Ib and II MPXV Infection in MEF cells": S2

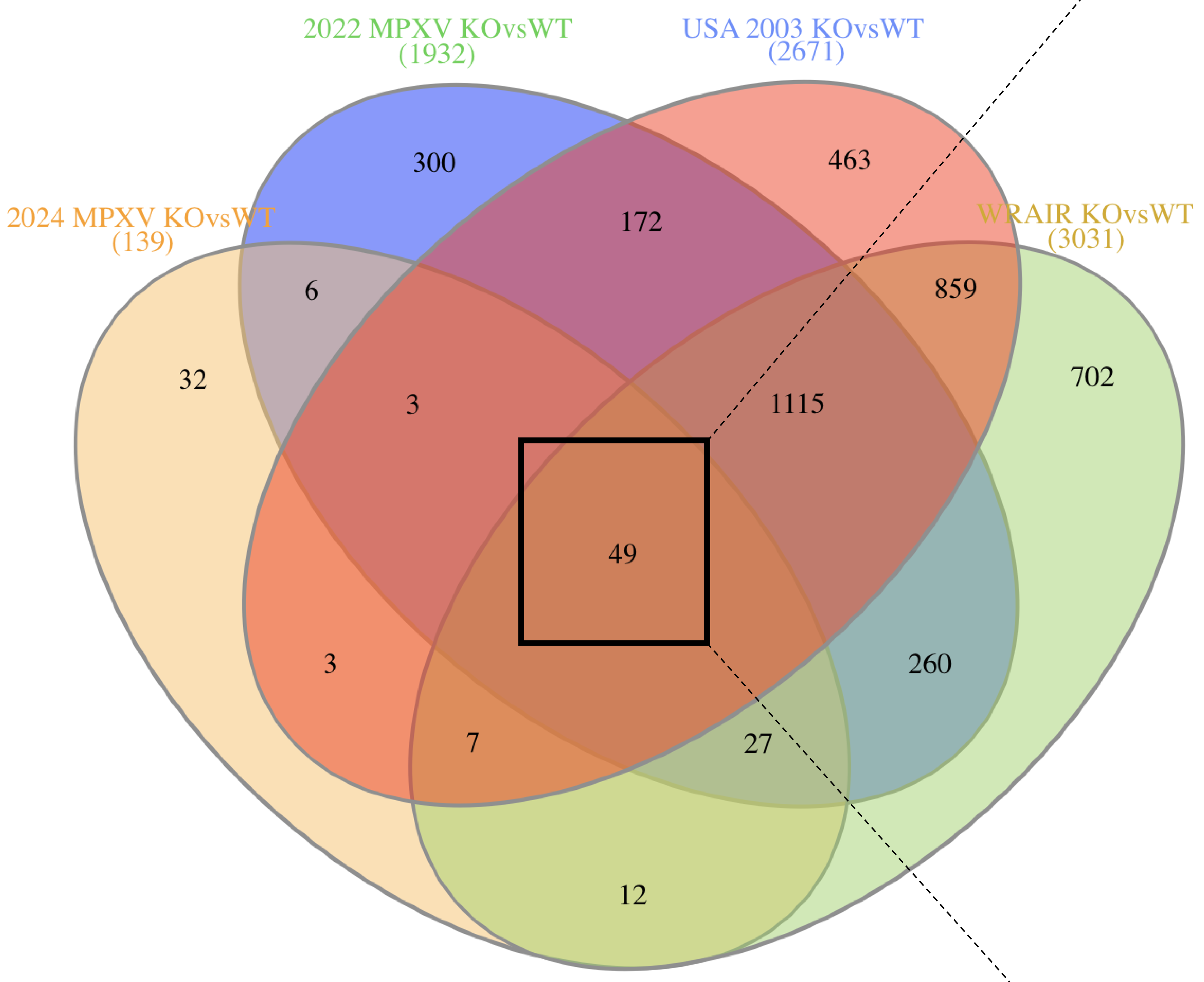


| Uniprot Code | Protein Name | 2024 KOvsWT Log2FC | 2022 KOvsWT Log2FC | USA KOvsWT Log2FC | WRAIR KOvsWT Log2FC |
|---|---|---|---|---|---|
| A0A7H0DN81 | Protein H4 | 1.7 | 1.4 | 0.7 | 2.1 |
| A0A7H0DMZ8 | Protein B28/C22 | 1.1 | -1.1 | -1.9 | -1.2 |
| A0A7H0DNA5 | Protein A7 | 1.5 | 1.4 | 0.8 | 1.7 |
| A0A7H0DNA4 | Protein A6 | 1.6 | 1.6 | 1.0 | 2.1 |
| A0A7H0DN84 | Protein H7 | 1.7 | 1.1 | 0.8 | 2.3 |
| A0A7H0DN97 | Protein D13 | 1.7 | 1.3 | 0.7 | 1.5 |
| A0A7H0DN95 | Protein D11 | 1.5 | 1.2 | 0.7 | 1.9 |
| A0A7H0DN89 | Protein D5 | 1.6 | 2.0 | 0.8 | 2.2 |
| A0A7H0DN96 | Protein D12 | 1.4 | 1.3 | 0.6 | 1.7 |
| A0A7H0DN80 | Protein H3 | 2.1 | 2.0 | 1.0 | 1.4 |
| A0A7H0DN78 | Protein H1 | 1.8 | 2.4 | 1.4 | 2.2 |
| A0A7H0DN42 | Protein E7 | 1.9 | 1.5 | 1.3 | 1.9 |
| A0A7H0DN43 | Protein E8 | 1.9 | 1.9 | 0.8 | 1.9 |
| P14901 | Hmox1 | 1.9 | -6.3 | -2.6 | -2.2 |
| A0A7H0DN46 | Protein E11 | 2.0 | 1.8 | 0.7 | 2.4 |
| A0A7H0DN48 | Protein O2 | 2.3 | 1.6 | 1.5 | 2.0 |
| A0A7H0DN79 | Protein H2 | 1.8 | 1.5 | 0.6 | 2.5 |
| A0A7H0DN57 | Protein G1 | 1.6 | 1.5 | 0.9 | 1.8 |
| A0A7H0DN37 | Protein E2 | 1.5 | 1.2 | 0.9 | 1.7 |
| A0A7H0DN72 | Protein J1 | 2.1 | 1.7 | 0.9 | 2.0 |
| A0A7H0DN74 | Protein J3 | 1.6 | 1.6 | 0.8 | 1.9 |
| A0A7H0DN76 | Protein J5 | 1.7 | 2.7 | 1.0 | 2.2 |
| A0A7H0DN20 | Protein F3 | 1.0 | 2.3 | 0.8 | 1.3 |
| A0A7H0DN27 | Protein F10 | 1.3 | 1.8 | 0.9 | 1.9 |
| Q7TPR4 | Actn1 | 1.0 | -1.9 | -1.7 | -1.74 |
| E9Q0S6 | Tns1 | 2.2 | -3.5 | -1.8 | -4.0 |
| Q64339 | ISG15 | -3.1 | -7.3 | -4.8 | -3.8 |
| Q91XV3 | Basp1 | 1.8 | -4.4 | -2.8 | -2.5 |
| Q9CZC8 | Scrn1 | 2.2 | 4.6 | 4.1 | 3.8 |
| Q9D1L0 | Chchd2 | -1.4 | -4.6 | -6.4 | -4.6 |
| Q9CPN8 | Igf2bp3 | 1.6 | 2.0 | 2.0 | 2.1 |
| M1LLA2 | Protein A1 | 1.6 | 1.5 | 0.6 | 1.8 |
| M1L502 | Protein G4 | 2.0 | 1.5 | 0.8 | 1.7 |
| M1KJ15 | Protein D2 | 1.2 | 2.6 | 1.5 | 2.1 |
| M1L543 | Protein A12 | 1.9 | 2.1 | 0.8 | 2.9 |
| Q63918 | Cavin2 | 1.3 | -3.5 | -5.5 | -4.5 |
| P58771 | Tpm1 | 1.0 | -2.4 | -2.4 | -1.9 |
| Q9QZL0 | Ripk3 | -1.3 | -7.6 | -7.0 | -8.1 |
| Q8R550 | Sh3kbp1 | -1.1 | -3.1 | -2.9 | -3.0 |
| M1LBQ5 | Protein D3 | 1.9 | 2.3 | 1.4 | 2.5 |
| P97447 | Fhl1 | 2.2 | 3.0 | 3.3 | 2.9 |
| A0A7H0DND9;Q8V4T7 | - | - | - | - | - |
| A0A7H0DND1 | Protein A32 | 1.5 | 1.1 | 0.7 | 2.2 |
| A0A7H0DNC1 | Protein A22 | 1.7 | 2.1 | 1.4 | 2.5 |
| A0A7H0DNE6 | Protein A50 | 1.1 | 1.9 | 1.1 | 1.1 |
| P47226 | Tes | -1.2 | -1.5 | -1.1 | -1.3 |
