## Supplementary material for "ISG15 Differentially Modulates Clade Ib and II MPXV Infection in MEF cells": S5

### Proteomic Analysis QC Graphs

**A**

**2022 MPXV**

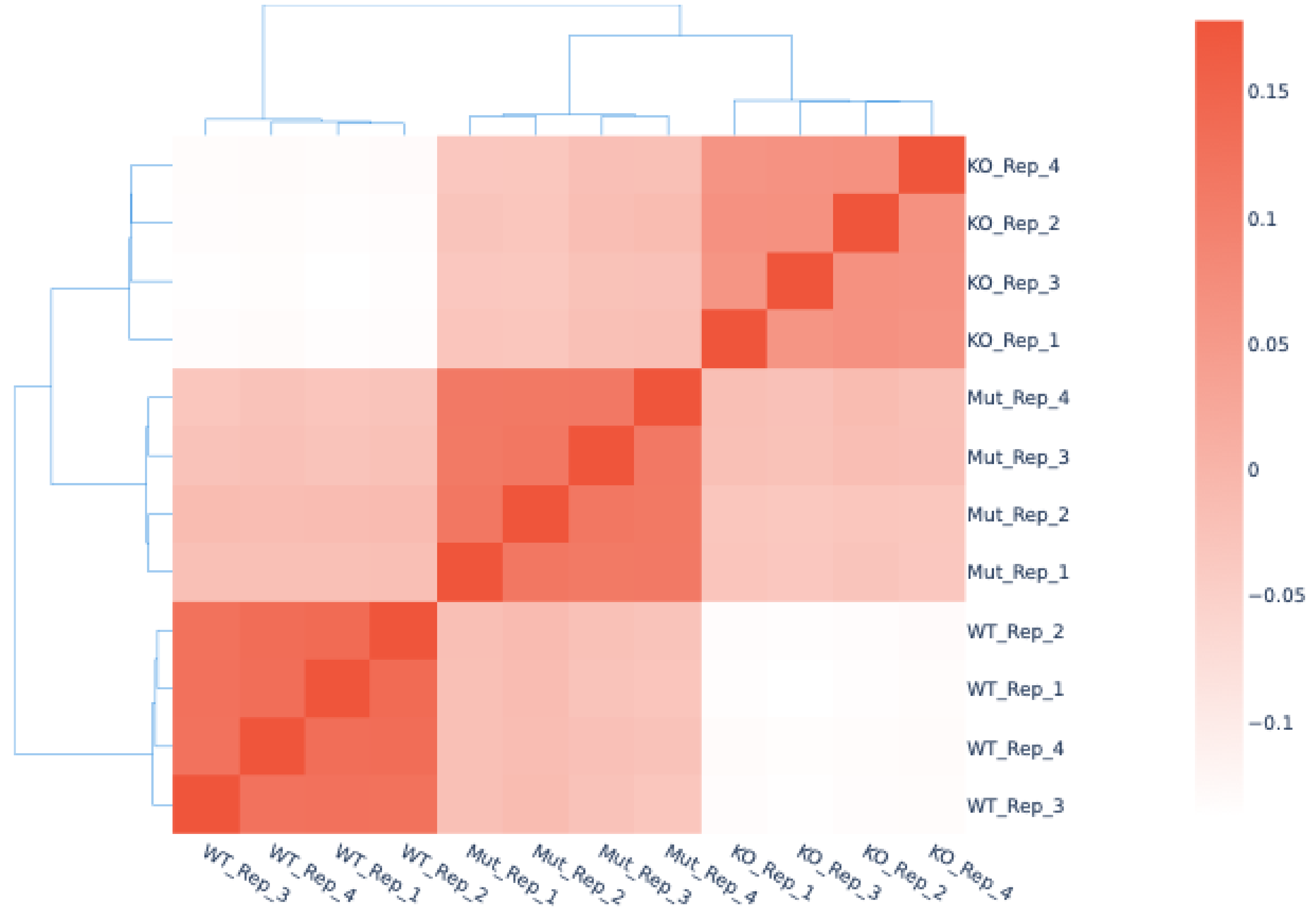


**B**

**USA 2003/ WRAIR**

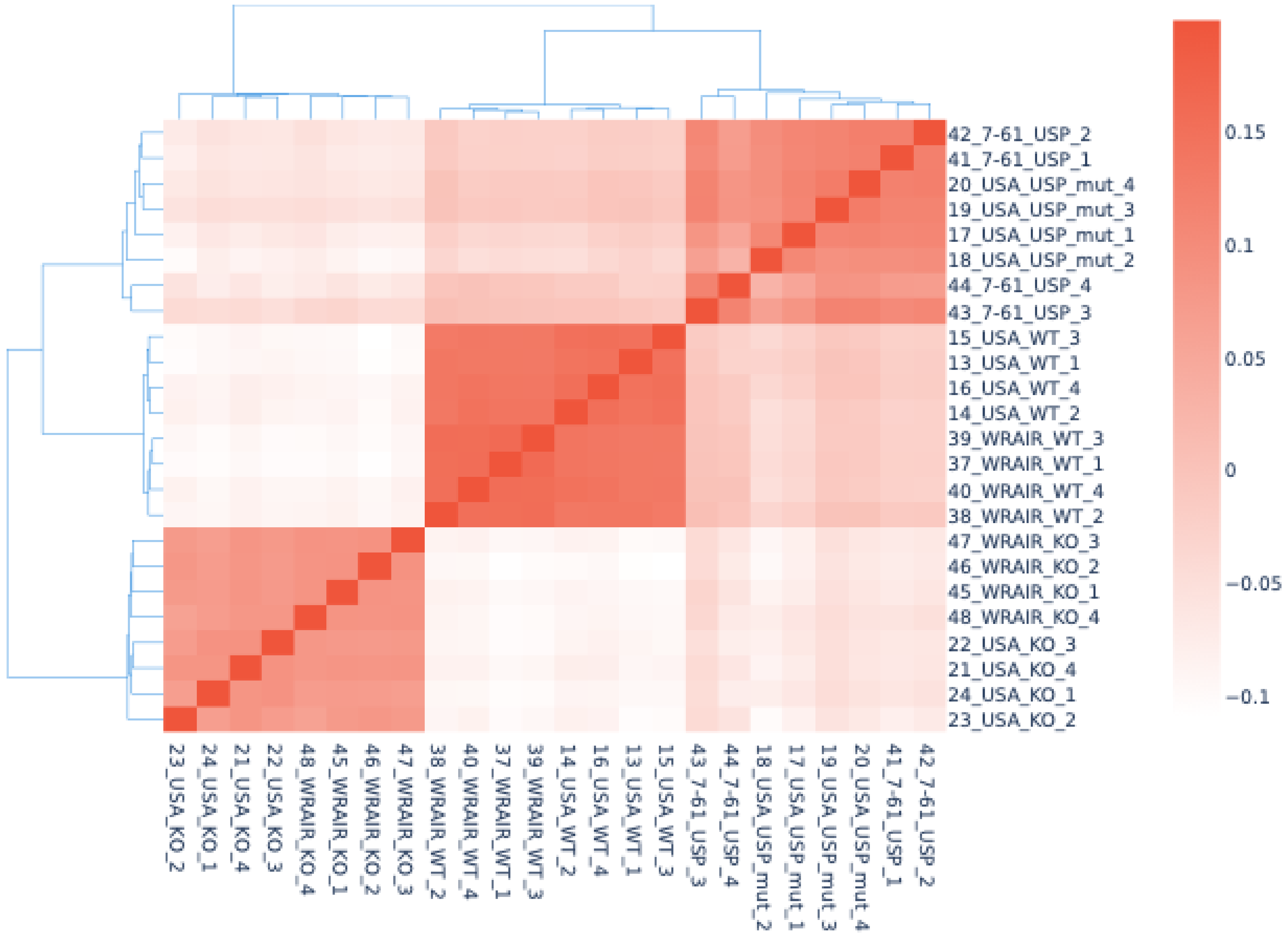


**C**

**2024 MPXV**

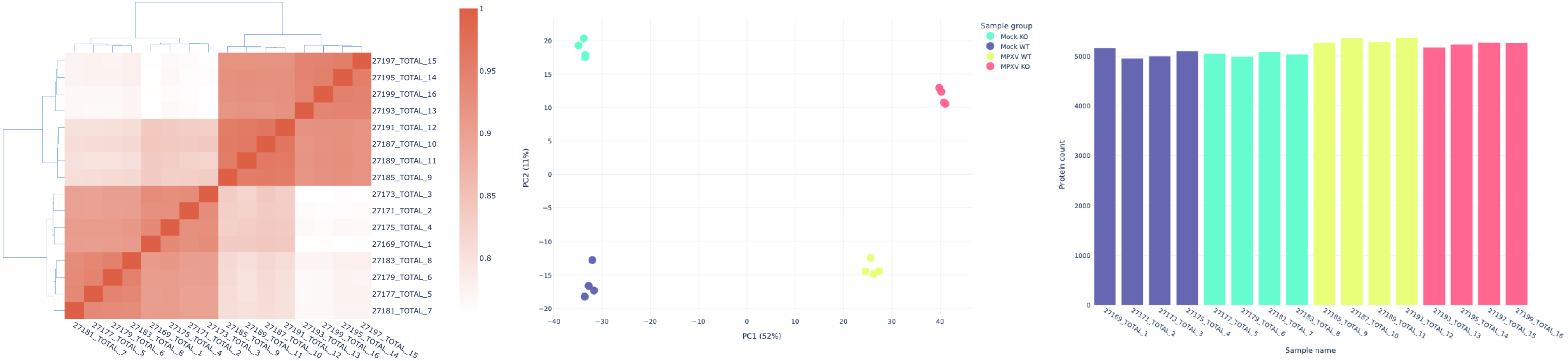


2024 MPXV Proteomics Total 1-4: Mock WT/ Total 5-8: Mock KO/ Total 9-12: 2024 WT/ Total 13-16: 2024 KO
