## Supplementary material for "ISG15 Differentially Modulates Clade Ib and II MPXV Infection in MEF cells": TableS1

| **Uniprot** | **Gene** | **Protein** | **2024**  **KOvsWT**  **FC** | **2024**  **KOvsWT**  **p Value**  **-Log10** | **2022**  **KOvsWT**  **FC** | **2022 KOvsWT**  **p Value**  **-Log10** | **USA**  **KOvsWT FC** | **USA KOvsWT p Value -Log10** | **761**  **KOvsWT FC** | **761**  **KOvsWT**  **p Value -Log10** |
| --- | --- | --- | --- | --- | --- | --- | --- | --- | --- | --- |
| P0DTM9 | OPG001 | Protein B29/C23 | 1.6 | 4.0 | N/S | N/S | -1.8 | 4.0 | -0.8 | 2.1 |
| P0DTN0 | OPG002 | Protein B28 | N/S | N/S | -1.3 | 3.2 | -1.9 | 2.9 | -1.4 | 2.8 |
| A0A7H0DNH0 | OPG003 MPXV | Protein OPG003 | N/S | N/S | N/S | N/S | N/S | N/S | 0.8 | 3.4 |
| A0A7H0DNG7 | OPG005 MPXV | Protein OPG005 | N/S | N/S | N/S | N/S | N/S | N/S | 0.9 | 4.1 |
| A0A7H0DMZ6 | OPG019 | Protein C11 | N/S | N/S | N/S | N/S | -1.7 | 3.4 | 0.8 | 2.2 |
| A0A7H0DMZ7 | OPG021 | Protein p28 | N/S | N/S | N/S | N/S | N/S | N/S | 1.5 | 3.7 |
| A0A7H0DMZ8 | OPG022 | Protein B28/C22 | 1.1 | 2.0 | -1.1 | 4.5 | -1.9 | 3.6 | -1.2 | 2.1 |
| A0A7H0DMZ9 | OPG023 | Protein MPXVgp011 | N/S | N/S | N/S | N/S | -0.8 | 3.4 | N/S | N/S |
| A0A7H0DN00 | OPG024 | Protein OPG024 | N/S | N/S | -2.1 | 4.5 | -2.7 | 5.4 | 1.4 | 3.4 |
| A0A7H0DN02 | OPG027 | Protein C7 | N/S | N/S | -1.1 | 3.3 | -1.7 | 5.0 | 1.0 | 2.4 |
| A0A7H0DN04 | OPG030 | Protein C5 | N/S | N/S | N/S | N/S | -0.9 | 4.6 | 0.7 | 3.9 |
| A0A7H0DN07 | OPG034 | Protein C1 | N/S | N/S | N/S | N/S | -1.6 | 3.3 | -1.0 | 2.6 |
| P0DTN4 | OPG035 | Protein N1 | 1.4 | 3.7 | N/S | N/S | 0.6 | 4.0 | 1.4 | 3.5 |
| A0A7H0DN09 | OPG036 MPXV | Protein OPG036 | N/S | N/S | N/S | N/S | -1.2 | 2.7 | N/S | N/S |
| A0A7H0DN10 | OPG037 | Protein M1 | N/S | N/S | 1.0 | 3.0 | N/S | N/S | N/S | N/S |
| A0A7H0DN11 | OPG038 | Protein M2 | N/S | N/S | N/S | N/S | -1.8 | 4.0 | -1.3 | 2.4 |
| A0A7H0DN12 | OPG039 | Protein K1 | N/S | N/S | 1.0 | 4.4 | N/S | N/S | N/S | N/S |
| A0A7H0DN15 | OPG042 | Protein K4 | N/S | N/S | 1.3 | 3.6 | N/S | N/S | N/S | N/S |
| A0A7H0DN16 | OPG043 MPXV | Protein OPG043 | N/S | N/S | N/S | N/S | -2.2 | 5.3 | N/S | N/S |
| A0A7H0DN17 | OPG044 | Protein K7 | N/S | N/S | N/S | N/S | 0.8 | 2.4 | 2.3 | 3.6 |
| A0A7H0DN20 | OPG047 | Protein F3 | 1.0 | 3.1 | 2.3 | 2.7 | 0.8 | 2.1 | 1.3 | 2.8 |
| A0A7H0DN23 | OPG050 | Protein F6 | 1.6 | 3.8 | 1.1 | 3.5 | N/S | N/S | 3.1 | 3.3 |
| A0A7H0DN24 | OPG051 MPXV | Protein OPG051 | N/S | N/S | N/S | N/S | -1.4 | 2.8 | N/S | N/S |
| A0A7H0DN25 | OPG052 | Protein F8 | 2.0 | 3.9 | 1.1 | 3.8 | N/S | N/S | 2.2 | 3.6 |
| A0A7H0DN26 | OPG053 | Protein F9 | 2.1 | 3.6 | 3.0 | 2.7 | N/S | N/S | 1.3 | 2.1 |
| A0A7H0DN27 | OPG054 | Protein F10 | 1.3 | 4.2 | 1.8 | 4.5 | 0.9 | 3.2 | 1.9 | 3.4 |
| A0A7H0DN28 | OPG055 | Protein F11 | N/S | N/S | N/S | N/S | -2.3 | 4.7 | -0.8 | 2.5 |
| A0A7H0DN29 | OPG056 | Protein F12 | 1.4 | 4.1 | N/S | N/S | 0.7 | 2.7 | 1.5 | 3.3 |
| A0A7H0DN30 | OPG057 | Protein F13 | 1.6 | 3.3 | N/S | N/S | N/S | N/S | 1.1 | 2.8 |
| A0A7H0DN32 | OPG059 | Protein F14 | 1.3 | 3.3 | N/S | N/S | N/S | N/S | N/S | N/S |
| A0A7H0DN33 | OPG060 | Protein F15 | N/S | N/S | N/S | N/S | -2.3 | 2.1 | N/S | N/S |
| A0A7H0DN35 | OPG062 | Protein F17 | 2.0 | 2.9 | 1.9 | 4.2 | N/S | N/S | 2.2 | 3.2 |
| A0A7H0DN36 | OPG063 | Protein E1 | 1.3 | 3.8 | 1.1 | 3.7 | N/S | N/S | 1.8 | 3.8 |
| A0A7H0DN37 | OPG064 | Protein E2 | 1.5 | 3.4 | 1.2 | 3.6 | 0.9 | 3.2 | 1.7 | 3.2 |
| A0A7H0DN39 | OPG066 | Protein E4 | 1.2 | 3.7 | N/S | N/S | N/S | N/S | 1.4 | 2.8 |
| A0A7H0DN41 | OPG068 | Protein E6 | 1.8 | 4.0 | 1.7 | 5.2 | N/S | N/S | 1.8 | 3.1 |
| A0A7H0DN42 | OPG069 | Protein E7 | 1.9 | 4.4 | 1.5 | 4.5 | 1.3 | 4.0 | 1.9 | 3.3 |
| A0A7H0DN43 | OPG070 | Protein E8 | 1.9 | 3.9 | 1.9 | 4.9 | 0.8 | 2.3 | 1.9 | 3.1 |
| A0A7H0DN44 | OPG071 | Protein E9 | N/S | N/S | N/S | N/S | N/S | N/S | 1.2 | 3.5 |
| A0A7H0DN45 | OPG072 | Protein E10 | 2.0 | 3.7 | 1.4 | 2.1 | N/S | N/S | N/S | N/S |
| A0A7H0DN46 | OPG073 | Protein E11 | 2.0 | 3.9 | 1.8 | 3.5 | 0.7 | 2.8 | 2.4 | 3.4 |
| A0A7H0DN48 | OPG075 | Protein O2 | 2.3 | 4.4 | 1.6 | 3.3 | 1.5 | 3.2 | 2.0 | 2.0 |
| M1L9M0 | OPG082 | Protein I6 | 1.0 | 3.0 | 1.0 | 3.5 | N/S | N/S | 1.4 | 4.0 |
| A0A7H0DN56 | OPG084 | Protein I8 | 1.5 | 3.9 | 1.6 | 3.3 | N/S | N/S | 2.2 | 3.2 |
| A0A7H0DN57 | OPG085 | Protein G1 | 1.6 | 3.7 | 1.5 | 4.0 | 0.9 | 2.5 | 1.8 | 2.3 |
| M1L9M3 | OPG087 | Protein G2 | N/S | N/S | 1.8 | 4.4 | N/S | N/S | 1.4 | 3.2 |
| M1L502 | OPG088 | Protein G4 | 2.0 | 4.4 | 1.5 | 3.5 | 0.8 | 3.4 | 1.7 | 2.5 |
| A0A7H0DN62 | OPG090 | Protein G5.5 | 1.2 | 3.1 | N/S | N/S | N/S | N/S | N/S | N/S |
| A0A7H0DN63 | OPG091 | Protein G6 | 1.1 | 2.0 | 4.1 | 2.9 | 0.8 | 2.5 | N/S | N/S |
| A0A7H0DN66 | OPG094 | Protein G9 | 1.5 | 3.3 | N/S | N/S | N/S | N/S | N/S | N/S |
| A0A7H0DN68 | OPG096 | Protein L2 | N/S | N/S | N/S | N/S | N/S | N/S | 0.6 | 2.8 |
| A0A7H0DN69 | OPG097 | Protein L3/F4 | 1.7 | 3.9 | 1.4 | 3.3 | N/S | N/S | 1.4 | 2.3 |
| M1L511 | OPG098 | Protein L4 | 2.1 | 4.2 | 1.6 | 4.6 | N/S | N/S | 1.9 | 3.1 |
| M1LBP0 | OPG099 | Protein L1 | 1.4 | 3.3 | 1.2 | 2.2 | N/S | N/S | N/S | N/S |
| A0A7H0DN71 | OPG099 | Protein L5 | N/S | N/S | 3.1 | 2.2 | N/S | N/S | N/S | N/S |
| A0A7H0DN72 | OPG100 | Protein J1 | 2.1 | 4.3 | 1.7 | 4.0 | 0.9 | 3.9 | 2.0 | 2.9 |
| A0A7H0DN73 | OPG101 | Protein J2 | N/S | N/S | N/S | N/S | -0.6 | 3.8 | N/S | N/S |
| A0A7H0DN74 | OPG102 | Protein J3 | 1.6 | 4.4 | 1.6 | 4.5 | 0.8 | 2.4 | 1.9 | 3.6 |
| M1L514 | OPG103 | Protein J4 | 1.4 | 4.3 |  |  | N/S | N/S | 1.0 | 2.4 |
| A0A7H0DN76 | OPG104 | Protein J5 | 1.7 | 3.9 | 2.7 | 3.4 | 1.0 | 2.4 | 2.2 | 2.3 |
| A0A7H0DN77 | OPG105 | Protein OPG105 | 1.3 | 4.0 | 1.2 | 3.8 | N/S | N/S | 2.4 | 3.8 |
| A0A7H0DN78 | OPG106 | Protein H1 | 1.8 | 3.2 | 2.4 | 5.7 | 1.4 | 3.3 | 2.2 | 2.9 |
| A0A7H0DN79 | OPG107 | Protein H2 | 1.8 | 2.6 | 1.5 | 4.1 | 0.6 | 2.6 | 2.5 | 2.3 |
| A0A7H0DN80 | OPG108 | Protein H3 | 2.1 | 4.0 | 2.0 | 4.1 | 1.0 | 2.8 | 1.4 | 2.2 |
| A0A7H0DN81 | OPG109 | Protein H4 | 1.7 | 3.8 | 1.4 | 3.8 | 0.7 | 3.2 | 2.1 | 3.1 |
| A0A7H0DN82 | OPG110 | Protein H5 | 1.5 | 4.0 | N/S | N/S | N/S | N/S | 1.0 | 2.4 |
| A0A7H0DN83 | OPG111 | Protein H6 | N/S | N/S | N/S | N/S | N/S | N/S | 1.9 | 3.0 |
| A0A7H0DN84 | OPG112 | Protein H7 | 1.7 | 3.4 | 1.1 | 2.7 | 0.8 | 3.2 | 2.3 | 3.5 |
| A0A7H0DN85 | OPG113 | Protein D1 | 1.4 | 3.7 | N/S | N/S | N/S | N/S | 1.3 | 3.4 |
| M1KJ15 | OPG114 | Protein D2 | 1.2 | 4.2 | 2.6 | 5.1 | 1.5 | 3.4 | 2.1 | 3.8 |
| M1LBQ5 | OPG115 | Protein D3 | 1.9 | 4.5 | 2.3 | 4.6 | 1.4 | 3.9 | 2.5 | 3.7 |
| M1LL92 | OPG116 | Protein D4 | 1.0 | 2.1 | N/S | N/S | N/S | N/S | 1.2 | 2.9 |
| A0A7H0DN89 | OPG117 | Protein D5 | 1.6 | 4.3 | 2.0 | 5.4 | 0.8 | 2.9 | 2.2 | 4.0 |
| A0A7H0DN90 | OPG118 | Protein D6 | 1.5 | 2.7 | N/S | N/S | N/S | N/S | 1.4 | 2.8 |
| A0A7H0DN91 | OPG119 | Protein D8 | 1.5 | 3.3 | 1.7 | 3.2 | N/S | N/S | N/S | N/S |
| A0A7H0DN93 | OPG121 | Protein D9 | N/S | N/S | N/S | N/S | -1.3 | 4.1 | N/S | N/S |
| A0A7H0DN95 | OPG123 | Protein D11 | 1.5 | 4.3 | 1.2 | 4.4 | 0.7 | 3.4 | 1.9 | 3.6 |
| A0A7H0DN96 | OPG124 | Protein D12 | 1.4 | 4.0 | 1.3 | 4.3 | 0.6 | 3.1 | 1.7 | 3.6 |
| A0A7H0DN97 | OPG125 | Protein D13 | 1.7 | 4.2 | 1.3 | 4.0 | 0.7 | 3.1 | 1.5 | 2.6 |
| M1LLA2 | OPG126 | Protein A1 | 1.6 | 3.7 | 1.5 | 4.7 | 0.6 | 3.5 | 1.8 | 2.9 |
| A0A7H0DN99 | OPG127 | Protein A2 | 1.1 | 3.0 | N/S | N/S | N/S | N/S | N/S | N/S |
| M1L535 | OPG128 | Protein A2.5 | 2.2 | 4.0 | N/S | N/S | N/S | N/S | N/S | N/S |
| M1KJ27 | OPG129 | Protein A3 | 2.0 | 3.9 | 1.3 | 3.8 | N/S | N/S | N/S | N/S |
| A0A7H0DNA2 | OPG130 | Protein A4 | 2.1 | 4.4 | 1.0 | 3.1 | N/S | N/S | 1.4 | 2.4 |
| A0A7H0DNA3 | OPG131 | Protein A5 | 1.4 | 3.5 | 1.0 | 3.9 | N/S | N/S | 1.3 | 3.5 |
| A0A7H0DNA4 | OPG132 | Protein A6 | 1.6 | 3.9 | 1.6 | 4.2 | 1.0 | 3.9 | 2.1 | 3.8 |
| A0A7H0DNA5 | OPG133 | Protein A7 | 1.5 | 4.3 | 1.4 | 4.2 | 0.8 | 2.6 | 1.7 | 3.9 |
| A0A7H0DNA8 | OPG136 | Protein A10 | 2.0 | 4.2 | 1.4 | 4.2 | N/S | N/S | N/S | N/S |
| M1L9Q3 | OPG137 | Protein A11 | 2.1 | 4.0 | 1.3 | 3.2 | N/S | N/S | 1.8 | 2.8 |
| M1L543 | OPG138 | Protein A12 | 1.9 | 4.4 | 2.1 | 4.2 | 0.9 | 2.1 | 2.9 | 3.2 |
| P0DTN1 | OPG139 | Protein A13 | 1.5 | 2.8 | 1.8 | 4.6 | N/S | N/S | 1.9 | 2.6 |
| A0A7H0DNB2 | OPG140 | Protein A14 | 1.8 | 3.2 | 1.4 | 2.1 | N/S | N/S | N/S | N/S |
| A0A7H0DNB4 | OPG142 | Protein A15 | 1.7 | 2.9 | 1.8 | 2.7 | N/S | N/S | N/S | N/S |
| A0A7H0DNB5 | OPG143 | Protein A16 | 1.3 | 3.7 | N/S | N/S | 0.9 | 2.2 | 1.7 | 2.2 |
| A0A7H0DNB6 | OPG144 | Protein A17 | 1.9 | 3.6 | N/S | N/S | N/S | N/S | N/S | N/S |
| A0A7H0DNB7 | OPG145 | Protein A18 | 1.4 | 4.3 | 1.3 | 4.8 | N/S | N/S | 1.7 | 3.6 |
| A0A7H0DNB8 | OPG146 | Protein A19 | 1.9 | 3.8 | 1.4 | 3.4 | N/S | N/S | 2.0 | 3.1 |
| A0A7H0DNC0 | OPG148 | Protein A20 | N/S | N/S | 1.3 | 3.7 | N/S | N/S | N/S | N/S |
| A0A7H0DNC1 | OPG149 | Protein A22 | 1.7 | 3.7 | 2.1 | 4.5 | 1.4 | 2.8 | 2.5 | 3.2 |
| A0A7H0DNC3 | OPG151 | Protein A24 | 1.5 | 4.1 | N/S | N/S | N/S | N/S | N/S | N/S |
| A0A7H0DNC4 | OPG153 | Protein A26 | 2.6 | 4.6 | 2.1 | 4.2 | N/S | N/S | 2.0 | 3.2 |
| P0DTN3 | OPG154 | Protein A27 | N/S | N/S | 1.4 | 5.2 | N/S | N/S | 1.6 | 2.9 |
| A0A7H0DNC7 | OPG156 | Protein A29 | 1.2 | 3.1 | N/S | N/S | N/S | N/S | 1.1 | 3.3 |
| A0A7H0DNC8 | OPG157 | Protein A30 | N/S | N/S | 1.8 | 2.9 | 0.8 | 2.3 | 2.2 | 3.4 |
| A0A7H0DND0 | OPG159 | Protein A31 | N/S | N/S | 1.0 | 4.2 | N/S | N/S | 1.2 | 4.2 |
| A0A7H0DND1 | OPG160 | Protein A32 | 1.5 | 4.6 | 1.1 | 3.0 | 0.7 | 2.4 | 2.2 | 3.7 |
| A0A7H0DND2 | OPG161 | Protein A33 | 1.6 | 4.4 | 1.7 | 3.8 | N/S | N/S | 1.7 | 3.1 |
| A0A7H0DND3 | OPG162 | Protein A34 | N/S | N/S | N/S | N/S | N/S | N/S | 1.5 | 2.7 |
| A0A7H0DND5 | OPG164 | Protein A36 | 1.2 | 3.0 | N/S | N/S | N/S | N/S | N/S | N/S |
| A0A7H0DND6 | OPG165 | Protein A37 | N/S | N/S | N/S | N/S | -1.3 | 5.1 | N/S | N/S |
| A0A7H0DND7 | OPG166 | Protein A38 | 1.7 | 2.1 | N/S | N/S | N/S | N/S | N/S | N/S |
| A0A7H0DND8 | OPG170 MPXV | Protein A41 | N/S | N/S | N/S | N/S | -2.1 | 3.4 | N/S | N/S |
| A0A7H0DNE0 | OPG172 | Protein A43 | N/S | N/S | 3.4 | 3.2 | N/S | N/S | N/S | N/S |
| A0A7H0DNE2 | OPG174 | Protein A44 | N/S | N/S | N/S | N/S | -1.1 | 2.3 | N/S | N/S |
| A0A7H0DNE4 | OPG176 | Protein A46 | N/S | N/S | N/S | N/S | N/S | N/S | 0.7 | 2.2 |
| A0A7H0DNE5 | OPG178 | Protein OPG178 | N/S | N/S | N/S | N/S | -1.1 | 3.0 | -0.7 | 3.0 |
| A0A7H0DNE6 | OPG180 | Protein A50 | 1.1 | 3.7 | 1.9 | 4.4 | 1.1 | 3.6 | 1.1 | 2.8 |
| Q8BEJ6 | OPG185 | Protein A56 | 2.0 | 3.3 | N/S | N/S | N/S | N/S | N/S | N/S |
| A0A7H0DNE9 | OPG187 | Protein B1 | N/S | N/S | N/S | N/S | N/S | N/S | 0.7 | 3.3 |
| A0A7H0DNF0 | OPG188 | Protein B2 | N/S | N/S | N/S | N/S | -1.2 | 3.2 | N/S | N/S |
| A0A7H0DNF1 | OPG189 | Protein B4 | 2.4 | 2.9 | N/S | N/S | N/S | N/S | N/S | N/S |
| P0DTN2 | OPG190 | Protein B5 | 1.5 | 4.0 | N/S | N/S | N/S | N/S | N/S | N/S |
| A0A7H0DNF4 | OPG192 | Protein B7 | 2.1 | 4.4 | 1.5 | 3.5 | N/S | N/S | N/S | N/S |
| A0A7H0DNF5 | OPG193 | Protein B8 | N/S | N/S | -1.3 | 4.7 | -2.3 | 2.4 | -2.7 | 3.6 |
| A0A7H0DNG0 | OPG200 | Protein B14 | N/S | N/S | N/S | N/S | -1.1 | 4.3 | N/S | N/S |
| A0A7H0DNG2 | OPG204 | Protein B18 | N/S | N/S | N/S | N/S | -2.1 | 4.0 | N/S | N/S |
| A0A7H0DNG4 | OPG205 | Protein C12 | N/S | N/S | N/S | N/S | N/S | N/S | 1.3 | 2.5 |
| A0A7H0DNG5 | OPG209 | Protein C13 | N/S | N/S | N/S | N/S | -1.6 | 2.3 | -1.1 | 2.3 |
