## Supplementary material for "ISG15 Differentially Modulates Clade Ib and II MPXV Infection in MEF cells": TableS2

| **Name** | **Gene** | **fold_change** | **p_value_adj_neg_log10** |
| --- | --- | --- | --- |
| Q64133 | Maoa | 5.41817468 | 2.64129563 |
| P37804 | Tagln | 4.20041172 | 5.636218694 |
| Q9WUU7 | Ctsz | 3.0348296 | 2.182667663 |
| Q08091 | Cnn1 | 2.98761663 | 2.418042874 |
| P39876 | Timp3 | 2.63817741 | 2.881673924 |
| A0A7H0DNC4 | OPG153 | 2.57040889 | 4.594333175 |
| A0A7H0DND9;Q8V4T7 | A0A7H0DND9;Q8V4T7 | 2.52912269 | 4.089291648 |
| P0DTN3 | OPG154 | 2.47203188 | 3.957396926 |
| P01831 | Thy1 | 2.44945154 | 2.868970775 |
| A0A7H0DNF1 | OPG189 | 2.39690046 | 2.948599146 |
| Q60760 | Grb10 | 2.37941809 | 4.594333175 |
| A0A7H0DN48 | OPG075 | 2.29442718 | 4.43256433 |
| Q9CZC8 | Scrn1 | 2.24456876 | 3.294985427 |
| M1L535 | OPG128 | 2.1989101 | 3.979132037 |
| E9Q0S6 | Tns1 | 2.18443868 | 3.308081882 |
| P97447 | Fhl1 | 2.15041046 | 5.614163567 |
| A0A7H0DN72 | OPG100 | 2.13642459 | 4.32448011 |
| A0A7H0DNF4 | OPG192 | 2.12997134 | 4.36155562 |
| M1L511 | OPG098 | 2.10899321 | 4.182783559 |
| A0A7H0DNA2 | OPG130 | 2.06723164 | 4.384817083 |
| M1L9Q3 | OPG137 | 2.06035379 | 3.979132037 |
| A0A7H0DN26 | OPG053 | 2.05847327 | 3.62341064 |
| A0A7H0DN80 | OPG108 | 2.05369344 | 3.966434162 |
| M1L502 | OPG088 | 2.03341237 | 4.431900757 |
| M1KJ27 | OPG129 | 2.03238607 | 3.882962794 |
| Q8BEJ6 | OPG185 | 2.03052923 | 3.256158713 |
| A0A7H0DN45 | OPG072 | 2.02126913 | 3.696541931 |
| A0A7H0DN35 | OPG062 | 1.99867607 | 2.904580117 |
| A0A7H0DNA8 | OPG136 | 1.99400235 | 4.182783559 |
| A0A7H0DN25 | OPG052 | 1.98033135 | 3.882962794 |
| A0A7H0DN46 | OPG073 | 1.96188867 | 3.882962794 |
| A0A7H0DN43 | OPG070 | 1.94368345 | 3.934998519 |
| P14901 | Hmox1 | 1.94019181 | 3.858583743 |
| M1L543 | OPG138 | 1.93952561 | 4.359070645 |
| A0A7H0DNB6 | OPG144 | 1.92336348 | 3.621114405 |
| A0A7H0DNB8 | OPG146 | 1.92316085 | 3.786915875 |
| Q9WTQ5 | Akap12 | 1.8893644 | 4.600895728 |
| A0A7H0DN42 | OPG069 | 1.8637745 | 4.359070645 |
| M1LBQ5 | OPG115 | 1.85741377 | 4.511037698 |
| Q91XV3 | Basp1 | 1.82524337 | 4.381576482 |
| A0A7H0DN41 | OPG068 | 1.8233813 | 4.022834163 |
| A0A7H0DN78 | OPG106 | 1.8169779 | 3.150459014 |
| A0A7H0DN79 | OPG107 | 1.7662099 | 2.564568817 |
| A0A7H0DNB2 | OPG140 | 1.76242 | 3.195276072 |
| P16460 | Ass1 | 1.74239159 | 4.182783559 |
| A0A7H0DN92;Q8V4Y0 | A0A7H0DN92;Q8V4Y0 | 1.73731391 | 3.369366232 |
| A0A7H0DN97 | OPG125 | 1.73243049 | 4.153315861 |
| A0A7H0DNB4 | OPG142 | 1.72048074 | 2.874897094 |
| A0A7H0DND7 | OPG166 | 1.71891742 | 2.080759569 |
| A0A7H0DN76 | OPG104 | 1.71506012 | 3.863768318 |
| A0A7H0DN69 | OPG097 | 1.69267338 | 3.916583132 |
| A0A7H0DN49;Q8V518 | A0A7H0DN49;Q8V518 | 1.67822105 | 4.356776313 |
| A0A7H0DN84 | OPG112 | 1.67389389 | 3.41760998 |
| A0A7H0DN81 | OPG109 | 1.65871518 | 3.806093394 |
| A0A7H0DNC1 | OPG149 | 1.65497269 | 3.682313261 |
| A0A7H0DND2 | OPG161 | 1.6262564 | 4.381576482 |
| A0A7H0DN74 | OPG102 | 1.60613721 | 4.431900757 |
| Q9CPN8 | Igf2bp3 | 1.60289814 | 3.457211918 |
| A0A7H0DN23 | OPG050 | 1.59122744 | 3.797397958 |
| M1LLA2 | OPG126 | 1.59061699 | 3.682313261 |
| A0A7H0DN57 | OPG085 | 1.57647219 | 3.682313261 |
| A0A7H0DN89 | OPG117 | 1.56963386 | 4.32448011 |
| A0A7H0DNA4 | OPG132 | 1.56809701 | 3.934998519 |
| A0A7H0DN30 | OPG057 | 1.5680919 | 3.308081882 |
| P0DTM9 | OPG001 | 1.55008347 | 3.988502573 |
| A0A7H0DND1 | OPG160 | 1.54145987 | 4.600895728 |
| A0A7H0DN90 | OPG118 | 1.52926933 | 2.749088544 |
| A0A7H0DN56 | OPG084 | 1.5282049 | 3.897068048 |
| A0A7H0DNC3 | OPG151 | 1.52022217 | 4.054346296 |
| A0A7H0DNA5 | OPG133 | 1.51486887 | 4.343573925 |
| P0DTN1 | OPG139 | 1.51302556 | 2.839094208 |
| P0DTN2 | OPG190 | 1.50444672 | 3.979132037 |
| A0A7H0DN66 | OPG094 | 1.50099169 | 3.311828359 |
| A0A7H0DN37 | OPG064 | 1.49298785 | 3.357756999 |
| A0A7H0DN91 | OPG119 | 1.46999038 | 3.319982798 |
| A0A7H0DN95 | OPG123 | 1.45654505 | 4.32448011 |
| A0A7H0DN82 | OPG110 | 1.45528722 | 3.988502573 |
| A0A7H0DN96 | OPG124 | 1.44767908 | 3.979132037 |
| P0DTN4 | OPG035 | 1.44362433 | 3.739027096 |
| A0A7H0DN29 | OPG056 | 1.4334019 | 4.136289444 |
| Q8K157 | Galm | 1.42667568 | 4.431900757 |
| A0A7H0DNB7 | OPG145 | 1.41465621 | 4.297472848 |
| M1L514 | OPG103 | 1.41355385 | 4.32448011 |
| M1LBP0 | OPG099 | 1.41296002 | 3.260980291 |
| A0A7H0DN85 | OPG113 | 1.38511162 | 3.724709305 |
| Q9DC11 | Plxdc2 | 1.38202981 | 3.535425666 |
| P13595 | Ncam1 | 1.38072123 | 4.359070645 |
| A0A7H0DNA3 | OPG131 | 1.37706483 | 3.543252026 |
| A0A7H0DN36 | OPG063 | 1.31189673 | 3.847693393 |
| A0A7H0DN27 | OPG054 | 1.30595553 | 4.226912075 |
| A0A7H0DNB5 | OPG143 | 1.28743211 | 3.70282362 |
| A0A7H0DN77 | OPG105 | 1.28672911 | 4.016925959 |
| P58774 | Tpm2 | 1.28089572 | 4.36155562 |
| P62746 | Rhob | 1.28020963 | 3.360865724 |
| A0A7H0DN32 | OPG059 | 1.27989594 | 3.308081882 |
| A0A7H0DNC6;Q8V4U9 | A0A7H0DNC6;Q8V4U9 | 1.27468613 | 2.805539107 |
| Q63918 | Cavin2 | 1.25846507 | 4.182783559 |
| Q9DCT8 | Crip2 | 1.24857879 | 3.139514843 |
| M1KJ15 | OPG114 | 1.24229405 | 4.182783559 |
| Q9CWS0 | Ddah1 | 1.22472719 | 3.882962794 |
| A0A7H0DNE3;Q8V4T3 | A0A7H0DNE3;Q8V4T3 | 1.2007954 | 3.815988236 |
| A0A7H0DN39 | OPG066 | 1.19912522 | 3.696541931 |
| A0A7H0DND5 | OPG164 | 1.19225118 | 2.958968907 |
| P97315 | Csrp1 | 1.18603303 | 3.260980291 |
| A0A7H0DN62 | OPG090 | 1.18545847 | 3.122078265 |
| A0A7H0DNC7 | OPG156 | 1.1811344 | 3.12479801 |
| **Q9ET54** | **Palld** | 1.16286757 | 5.350851487 |
| O88667 | Rrad | 1.12486797 | 3.399270145 |
| A0A7H0DNE6 | OPG180 | 1.12018306 | 3.682313261 |
| A0A7H0DN99 | OPG127 | 1.10645293 | 2.96662547 |
| P20352 | F3 | 1.10363855 | 3.269725094 |
| A0A7H0DN63 | OPG091 | 1.10354369 | 2.027439056 |
| Q8R5F7 | Ifih1 | 1.08457538 | 4.211765498 |
| A0A7H0DMZ8 | OPG022 | 1.07460985 | 2.0464144 |
| Q9CQQ4 | Gemin2 | 1.06272765 | 4.153315861 |
| P58771 | Tpm1 | 1.04562493 | 3.432532957 |
| P17879 | Hspa1b | 1.03152226 | 5.636218694 |
| M1LL92 | OPG116 | 1.02255826 | 2.12279394 |
| A0A7H0DN20 | OPG047 | 1.02034444 | 3.141507185 |
| M1L9M0 | OPG082 | 1.01377193 | 3.011317227 |
| Q7TPR4 | Actn1 | 1.00658347 | 4.8086869 |
| Q9QUJ7 | Acsl4 | -1.0016079 | 3.934998519 |
| O09131 | Gsto1 | -1.0417769 | 4.136289444 |
| Q9CQ43 | Dut | -1.0471969 | 4.359070645 |
| Q8R5M8 | Cadm1 | -1.1326186 | 3.038192297 |
| Q8R550 | Sh3kbp1 | -1.139656 | 3.682313261 |
| P47226 | Tes | -1.1649551 | 3.306811284 |
| Q9QZL0 | Ripk3 | -1.3185087 | 4.211765498 |
| Q9D1L0 | Chchd2 | -1.3614979 | 3.436866757 |
| P62274 | Rps29 | -1.4894242 | 2.863746779 |
| P10400 | Pol | -1.5922444 | 2.887849278 |
| Q61850 | **Foxc2** | -1.7157344 | 3.241440118 |
| Q9Z0E6 | Gbp2 | -1.80768 | 2.019131039 |
| P51912 | Slc1a5 | -1.8318717 | 2.342782585 |
| Q3V038 | Ttc9 | -1.8679184 | 3.308081882 |
| P11370 | Fv4 | -2.3543828 | 2.270380748 |
| P43135 | Nr2f2 | -2.5594778 | 2.774058564 |
| Q64339 | Isg15 | -3.0901943 | 2.757088574 |
| P10923 | Spp1 | -4.1122111 | 2.272471133 |

**Table S1.** *Protein signature of 2024 MPXV MOI 1 infected KO vs WT MEF lysates at 21 hours post-infection. Table lists the protein ID, the log2-fold change (FC) (MPXV infected WT MEFs vs infected ISG15 KO MEFs) of the levels of each protein, and the statistical significance (−log P value). Ordered from largest to smallest FC. In bold, mentioned proteins in manuscript.*
