## Supplementary material for "ISG15 Differentially Modulates Clade Ib and II MPXV Infection in MEF cells": TableS3

| **Name** | **Gene** | **fold_change** | **p_value_adj_neg_log10** |
| --- | --- | --- | --- |
| P31254 | Uba1y | 7.867019964 | 6.074962845 |
| P28651 | Ca8 | 7.690196954 | 2.811607306 |
| Q9Z0N2 | Eif2s3y | 7.346322445 | 5.456448271 |
| Q8VIN1 | Pbp2 | 6.95395233 | 5.956516689 |
| Q6WVG3 | Kctd12 | 6.375792765 | 5.304891004 |
| Q03526 | Itk | 6.217067032 | 2.987500292 |
| Q80TL0 | Ppm1e | 6.158605772 | 5.108152267 |
| P08553 | Nefm | 6.057049828 | 3.022102985 |
| P00920 | Ca2 | 5.982995899 | 2.482956727 |
| Q9DAJ5 | Dynlrb2 | 5.95198054 | 3.997648036 |
| P30275 | Ckmt1 | 5.845189936 | 4.790899605 |
| Q9EQF6 | Dpysl5 | 5.512879147 | 3.167655458 |
| Q91WT9 | Cbs | 5.452657761 | 3.942589051 |
| P16125 | Ldhb | 4.909164927 | 3.43038628 |
| Q9CZC8 | Scrn1 | 4.859598578 | 3.769758773 |
| P61458 | Pcbd1 | 4.674388522 | 2.238279886 |
| Q80ZN9 | Cox6b2 | 4.51367875 | 4.24444 |
| P19639 | Gstm3 | 4.497786918 | 5.439865673 |
| O55111 | Dsg2 | 4.377389441 | 5.054813455 |
| Q8JZV9 | Bdh2 | 4.33075469 | 3.379827241 |
| Q8R0P4 | Aamdc | 4.323382432 | 2.763653523 |
| Q9JHI5 | Ivd | 4.220503159 | 3.379827241 |
| P56371 | Rab4a | 4.210810191 | 3.876113878 |
| P58321 | Uchl4 | 4.185808654 | 3.028424232 |
| Q3U0J8 | Tbc1d2b | 4.164025855 | 2.804885859 |
| Q8K157 | Galm | 4.158929501 | 2.741562966 |
| Q62148 | Aldh1a2 | 4.14206587 | 4.061741488 |
| A0A7H0DN63 | OPG091 | 4.082876688 | 2.9206348 |
| Q91ZE0 | Tmlhe | 3.914602636 | 3.370554182 |
| Q61474 | Msi1 | 3.907879926 | 4.744180963 |
| Q9D5J6 | Shpk | 3.890427343 | 2.798671852 |
| Q9CZS1 | Aldh1b1 | 3.883204708 | 2.881584501 |
| P17183 | Eno2 | 3.84631606 | 3.674192251 |
| Q8C779 | Radx | 3.836203037 | 3.961950585 |
| B1AVY7 | Kif16b | 3.82627978 | 3.666380082 |
| Q45KJ6 | Lin28b | 3.80622306 | 2.421569949 |
| Q6PDS3 | Sarm1 | 3.800508948 | 3.624600882 |
| O35188 | Cx3cl1 | 3.790675336 | 3.495475425 |
| P14094 | Atp1b1 | 3.698560385 | 3.297777922 |
| Q8BTG3 | Tcp11l1 | 3.67866048 | 2.32596558 |
| Q9D711 | Pir | 3.65466387 | 4.445252632 |
| P16381 | D1Pas1 | 3.62486435 | 4.456645619 |
| P70207 | Plxna2 | 3.582123738 | 6.239839385 |
| Q8BGV0 | Nars2 | 3.536312359 | 4.12668646 |
| Q3UQ44 | Iqgap2 | 3.526616957 | 3.33703832 |
| Q9DD02 | Hikeshi | 3.469394009 | 4.535801556 |
| G5E8K5 | Ank3 | 3.46596296 | 2.138060078 |
| Q8VCK3 | Tubg2 | 3.445728065 | 2.614376375 |
| Q9DA03 | Lyrm7 | 3.405299333 | 4.293579113 |
| O70209 | Pdlim3 | 3.394727665 | 3.435186902 |
| A0A7H0DNE0 | OPG172 | 3.382731168 | 3.182811913 |
| Q9CQE5 | Rgs10 | 3.348372214 | 3.590737257 |
| P08551 | Nefl | 3.34463706 | 3.33931556 |
| Q9QYB2 | Dach1 | 3.338751547 | 2.216005514 |
| Q8K0E1 | Kctd15 | 3.322416169 | 3.048725776 |
| O55042 | Snca | 3.317649349 | 3.222844106 |
| Q60936 | Coq8a | 3.302028653 | 3.621765036 |
| Q9CPU0 | Glo1 | 3.29426484 | 5.685614681 |
| Q3U564 | Dcp1b | 3.197542257 | 3.889186008 |
| P55271 | Cdkn2b | 3.194060884 | 6.384665185 |
| Q9JME5 | Ap3b2 | 3.181859792 | 3.064046074 |
| A0A7H0DN71 | OPG099 | 3.146729064 | 2.191105515 |
| Q9QZW9 | Mnx1 | 3.132449901 | 3.763064603 |
| Q62441 | Tle4 | 3.118681453 | 3.706003395 |
| P97447 | Fhl1 | 3.038844728 | 6.006599674 |
| P97313 | Prkdc | 3.03158169 | 3.076882103 |
| Q9D114 | Hddc3 | 3.015332677 | 4.194377901 |
| P59024 | Fkbp14 | 3.007056122 | 2.304948085 |
| Q641K1 | Agtpbp1 | 3.004810301 | 3.257008203 |
| O55091 | Impact | 3.004605237 | 2.242685909 |
| Q9JI91 | Actn2 | 3.000473218 | 5.372594106 |
| A0A7H0DN26 | OPG053 | 2.992628144 | 2.71389782 |
| Q8CAV0 | Ybey | 2.987833823 | 2.578630348 |
| Q04447 | Ckb | 2.981148361 | 5.301412195 |
| Q60899 | Elavl2 | 2.980378806 | 4.315603446 |
| Q8R1G2 | Cmbl | 2.977596386 | 3.666683919 |
| Q5BKP2 | Usp13 | 2.973173562 | 2.646461564 |
| Q8CAK1 | Iba57 | 2.958423819 | 3.652026712 |
| Q8K4G5 | Ablim1 | 2.923115771 | 5.013779153 |
| Q3UYC0 | Ppm1h | 2.877643571 | 3.366594436 |
| Q3UIW5 | Rnf10 | 2.83702208 | 2.208909153 |
| Q8BRK8 | Prkaa2 | 2.836796341 | 2.574071954 |
| P11679 | Krt8 | 2.831578574 | 2.161530567 |
| Q9D1H6 | Ndufaf4 | 2.810738157 | 2.708533301 |
| Q9D920 | Borcs5 | 2.807919393 | 2.121702003 |
| O54788 | Dffb | 2.779296708 | 2.401073334 |
| Q8K4I3 | Arhgef6 | 2.741039042 | 2.309637363 |
| A3KMP2 | Ttc38 | 2.728831851 | 2.577322599 |
| Q149C2 | Traf3ip1 | 2.695864648 | 2.925820904 |
| A0A7H0DN76 | OPG104 | 2.672878864 | 3.389352956 |
| Q9D8X1 | Cutc | 2.671333737 | 2.406377776 |
| Q5RL51 | Gstcd | 2.665286462 | 3.033320458 |
| Q68FE2 | Atg9a | 2.653490941 | 4.813512893 |
| Q059Y8 | Dcst1 | 2.653360472 | 3.99418129 |
| Q9CQA9 | Ntpcr | 2.651753262 | 3.992350703 |
| Q9CQE0 | Rnf138 | 2.637705703 | 2.029423986 |
| Q7TPD6 | Raver2 | 2.629301388 | 3.127994206 |
| M1KJ15 | OPG114 | 2.592255955 | 5.146289744 |
| P24549 | Aldh1a1 | 2.590288097 | 3.48176744 |
| Q64442 | Sord | 2.582783252 | 2.232587355 |
| P97931 | Ung | 2.578367792 | 4.77291352 |
| Q91VN1 | Znf24 | 2.572121672 | 3.372488468 |
| Q7TPM6 | Fsd1 | 2.551940244 | 2.891798486 |
| Q91W43 | Gldc | 2.53224987 | 3.365638571 |
| P09066 | En2 | 2.52813304 | 2.925955849 |
| P49813 | Tmod1 | 2.524902903 | 2.56854113 |
| P15105 | Glul | 2.458055549 | 5.485443412 |
| Q9CYT6 | Cap2 | 2.457029103 | 3.223382626 |
| Q60611 | Satb1 | 2.452811792 | 2.137984003 |
| P41139 | Id4 | 2.448231413 | 4.345719625 |
| Q8C8N2 | Scai | 2.439962092 | 4.830246859 |
| A0A7H0DN78 | OPG106 | 2.438643957 | 5.658443155 |
| B1AXP6 | Tomm5 | 2.436527464 | 3.020074884 |
| P27790 | Cenpb | 2.394186926 | 3.182249906 |
| P48758 | Cbr1 | 2.374321268 | 3.751356221 |
| O88396 | Grpel2 | 2.373957109 | 2.599089035 |
| M1LBQ5 | OPG115 | 2.339914521 | 4.613213288 |
| P23475 | Xrcc6 | 2.328977544 | 4.534017238 |
| Q8VI93 | Oas3 | 2.305663321 | 4.399134168 |
| P0C605 | Prkg1 | 2.295907199 | 2.865400187 |
| Q9JKW0 | Arl6ip1 | 2.290700128 | 2.03499861 |
| A0A7H0DN20 | OPG047 | 2.289916375 | 2.724404259 |
| Q8C196 | Cps1 | 2.282816361 | 2.429831468 |
| Q8BGD6 | Slc38a9 | 2.271281955 | 2.437323488 |
| Q99MU3 | Adar | 2.246550607 | 4.491522958 |
| O89106 | Fhit | 2.224008006 | 3.271253463 |
| P10637 | Mapt | 2.218144682 | 2.506973682 |
| Q9R0Q6 | Arpc1a | 2.210400942 | 4.19568106 |
| Q61169 | Gata6 | 2.204350837 | 2.377316523 |
| P83510 | Tnik | 2.195845262 | 3.759733241 |
| E9PVD3 | Dchs1 | 2.188522758 | 2.036276324 |
| B1ARW8 |  | 2.185090709 | 2.046200026 |
| Q8C7Q4 | Rbm4 | 2.166282242 | 5.60091374 |
| Q9DB50 | Ap1s2 | 2.15901858 | 2.687923565 |
| Q32NZ6 | Tmc5 | 2.151000345 | 2.60833269 |
| Q9DBL2 | Gdap2 | 2.150872616 | 2.164723739 |
| Q8BYH3 | Trmt13 | 2.137005935 | 2.272301225 |
| Q9CQZ7 | Polr3k | 2.133988561 | 2.438245614 |
| O70583 | Mid1 | 2.129473537 | 3.240257247 |
| Q9ERE3 | Sgk3 | 2.109956929 | 2.058848697 |
| Q61599 | Arhgdib | 2.107361337 | 5.146289744 |
| A0A7H0DNC4 | OPG153 | 2.095311791 | 4.218376287 |
| A0A7H0DNC1 | OPG149 | 2.073695695 | 4.450296389 |
| M1L543 | OPG138 | 2.058593928 | 4.166111963 |
| A0A7H0DN17 | OPG044 | 2.053263147 | 3.667683569 |
| Q8CB27 | Yod1 | 2.0443341 | 4.586399308 |
| Q8BUL6 | Plekha1 | 2.044129917 | 2.341230564 |
| Q9CQ02 | Commd4 | 2.031688979 | 2.593165547 |
| Q9CPN8 | Igf2bp3 | 1.997294149 | 4.199705293 |
| P60521 | Gabarapl2 | 1.984717004 | 4.568429436 |
| Q64512 | Ptpn13 | 1.98211765 | 2.157283496 |
| Q9D7X8 | Ggct | 1.979432723 | 5.124297528 |
| A0A7H0DN80 | OPG108 | 1.962901825 | 4.129593044 |
| A0A7H0DN89 | OPG117 | 1.956833187 | 5.364449318 |
| Q9Z191 | Eya4 | 1.953605604 | 3.263888972 |
| Q9CWF2 | Tubb2b | 1.928832607 | 3.125568408 |
| A0A7H0DNE6 | OPG180 | 1.917932639 | 4.361587937 |
| A0A7H0DN43 | OPG070 | 1.914381831 | 4.90767091 |
| Q80W88 | Homez | 1.875923253 | 2.841458122 |
| Q8JZS0 | Lin7a | 1.867968354 | 2.742328255 |
| Q62420 | Sh3gl2 | 1.863355865 | 3.236515046 |
| A0A7H0DN35 | OPG062 | 1.856499745 | 4.236364123 |
| Q8BT51 | COA4 | 1.84983546 | 3.445111651 |
| P0DTN1 | OPG139 | 1.844975475 | 4.606924891 |
| Q9QYB8 | Add2 | 1.831311561 | 2.054483341 |
| Q8BYJ6 | Tbc1d4 | 1.828414815 | 2.564537551 |
| Q925B0 | Pawr | 1.827807465 | 4.617016377 |
| A0A7H0DN27 | OPG054 | 1.827618416 | 4.543557558 |
| P62855 | Rps26 | 1.818924702 | 2.413340227 |
| P11103 | Parp1 | 1.803232686 | 3.790609706 |
| Q8R001 | Mapre2 | 1.792584612 | 4.785385155 |
| A0A7H0DNB4 | OPG142 | 1.792437643 | 2.653446694 |
| M1L9M3 | OPG087 | 1.790086549 | 4.358854655 |
| P47964 | Rpl36 | 1.769205126 | 2.208909153 |
| A0A7H0DNC8 | OPG157 | 1.769045666 | 2.945541416 |
| A0A7H0DN46 | OPG073 | 1.756180482 | 3.538891335 |
| A0A7H0DN91 | OPG119 | 1.73716054 | 3.191053218 |
| A0A7H0DND2 | OPG161 | 1.727359801 | 3.837174782 |
| P60824 | Cirbp | 1.720495326 | 4.092282459 |
| Q9CR64 | Tmem167a | 1.71890778 | 2.71140787 |
| Q8CEZ0 | Kctd2 | 1.702711757 | 2.385758222 |
| A0A7H0DN41 | OPG068 | 1.702276826 | 5.155459614 |
| P62305 | Snrpe | 1.695599027 | 4.06026088 |
| A0A7H0DN72 | OPG100 | 1.67355944 | 4.022907829 |
| Q9D6T1 | Tube1 | 1.671466149 | 2.3649017 |
| O35984 | Pbx2 | 1.669849806 | 2.391972208 |
| Q9WV02 | Rbmx | 1.653709069 | 2.161530567 |
| Q9JKL4 | Ndufaf3 | 1.637648653 | 2.831905855 |
| Q03173 | Enah | 1.622429733 | 4.482379556 |
| Q9Z129 | Recql | 1.603974691 | 4.04388881 |
| M1L511 | OPG098 | 1.596626206 | 4.588259686 |
| Q9DB07 | Ift46 | 1.580972381 | 2.127534967 |
| Q3TAA7 | Stk11ip | 1.574428206 | 2.622324773 |
| Q9EQS9 | Igdcc4 | 1.57062703 | 3.780172873 |
| A0A7H0DN74 | OPG102 | 1.562372231 | 4.492631565 |
| A0A7H0DN56 | OPG084 | 1.561717671 | 3.346806955 |
| Q6ZWN5 | Rps9 | 1.560434084 | 2.829743475 |
| A0A7H0DN48 | OPG075 | 1.557839751 | 3.307307481 |
| Q78IS1 | Tmed3 | 1.55662152 | 2.119971815 |
| A0A7H0DNA4 | OPG132 | 1.552426671 | 4.194377901 |
| M1L502 | OPG088 | 1.544585266 | 3.508306196 |
| A0A7H0DNF4 | OPG192 | 1.541091942 | 3.53839266 |
| A0A7H0DN57 | OPG085 | 1.540605106 | 4.036374954 |
| A0A7H0DN79 | OPG107 | 1.521972408 | 4.086829476 |
| Q9D032 | Ssbp3 | 1.519680905 | 3.566779763 |
| Q9JJV2 | Pfn2 | 1.515629441 | 3.958727997 |
| A0A7H0DN42 | OPG069 | 1.511407874 | 4.479171456 |
| Q8BGS2 | Bola2 | 1.503543362 | 5.438059717 |
| Q8C0D0 | Trub1 | 1.499043956 | 2.745410003 |
| Q9CQF6 | Aasdhppt | 1.496309601 | 2.719893809 |
| Q8BGV8 | Mief1 | 1.482234769 | 2.725963974 |
| Q8BPK2 | Zcchc3 | 1.481189043 | 2.93283529 |
| Q8CCA0 | Dcun1d4 | 1.472219047 | 3.566779763 |
| Q05186 | Rcn1 | 1.470135134 | 4.082764858 |
| M1LLA2 | OPG126 | 1.465220664 | 4.689457015 |
| Q80V62 | Fancd2 | 1.449604246 | 3.77835938 |
| P0DTN3 | OPG154 | 1.449458862 | 5.165196611 |
| A0A7H0DN81 | OPG109 | 1.442529975 | 3.824685892 |
| Q9D0B0 | Srsf9 | 1.440624635 | 4.143111025 |
| Q8VIM9 | Irgq | 1.435041978 | 3.777104823 |
| Q8BJZ4 | Mrps35 | 1.433154134 | 3.191520923 |
| A0A7H0DNA8 | OPG136 | 1.430688231 | 4.20602303 |
| Q91VL8 | Terf2ip | 1.40503347 | 4.11567958 |
| Q924Z6 | Xpo6 | 1.399814353 | 3.545576685 |
| A0A7H0DNB2 | OPG140 | 1.39013363 | 2.056107392 |
| O88477 | Igf2bp1 | 1.388739398 | 5.699939096 |
| Q9Z130 | Hnrnpdl | 1.388115952 | 4.910967547 |
| A0A7H0DN69 | OPG097 | 1.384888005 | 3.260182598 |
| Q9CQ89 | Cuta | 1.381589938 | 3.773310299 |
| Q9Z2Z9 | Gfpt2 | 1.378466913 | 2.018444886 |
| A0A7H0DNB8 | OPG146 | 1.377849443 | 3.428979468 |
| A0A7H0DN45 | OPG072 | 1.377006891 | 2.127303156 |
| Q9D273 | Mmab | 1.366745549 | 3.79312351 |
| A0A7H0DNA5 | OPG133 | 1.357003926 | 4.197532132 |
| M1KJ27 | OPG129 | 1.344964812 | 3.761868005 |
| Q04841 | Mpg | 1.341524859 | 2.385879776 |
| Q5SU73 | Coil | 1.340486383 | 5.653371399 |
| M1L9Q3 | OPG175 | 1.327632235 | 3.182149927 |
| Q8R123 | Flad1 | 1.326136245 | 4.092505374 |
| Q99K51 | Pls3 | 1.31103241 | 5.693763805 |
| Q9D9H8 | Cb069 | 1.304651447 | 2.289898102 |
| Q08122 | Tle3 | 1.299743872 | 4.98391828 |
| Q8K0Z7 | Taco1 | 1.299404078 | 3.28854454 |
| A0A7H0DN97 | OPG125 | 1.295654494 | 4.041025008 |
| Q9D7S7 | Rpl22l1 | 1.295482924 | 3.926334063 |
| A0A7H0DNC0 | OPG148 | 1.286720103 | 3.664622717 |
| A0A7H0DN96 | OPG124 | 1.285783371 | 4.321174517 |
| Q922B1 | Macrod1 | 1.284698464 | 3.441552772 |
| Q8VI24 | Satb2 | 1.280253813 | 3.063354165 |
| Q61189 | Clns1a | 1.262214617 | 3.295499037 |
| P97364 | Sephs2 | 1.260207707 | 5.132566088 |
| A0A7H0DN15 | OPG042 | 1.256004658 | 3.635948014 |
| Q9CZT4 | Polr3e | 1.253952153 | 3.048308288 |
| Q8BHN1 | Txlng | 1.251661293 | 4.167868373 |
| A0A7H0DNB7 | OPG145 | 1.2515137 | 4.831192686 |
| M1LBP0 | OPG099 | 1.242071194 | 2.234076473 |
| Q9Z204 | Hnrnpc | 1.237493953 | 4.224054207 |
| A0A7H0DN37 | OPG064 | 1.230468604 | 3.572487984 |
| Q8BFY6 | Pef1 | 1.225353717 | 3.767177067 |
| Q9D1F3 | Eola1 | 1.221901611 | 2.071316885 |
| P56812 | Pdcd5 | 1.210925832 | 4.478713687 |
| A0A7H0DN95 | OPG123 | 1.209174578 | 4.410572856 |
| A0A7H0DN77 | OPG151 | 1.200479525 | 3.789675561 |
| P56671 | Maz | 1.195338903 | 2.738523635 |
| Q6PE54 | Dhx40 | 1.195307415 | 4.482379556 |
| Q9QY93 | Dctpp1 | 1.19380318 | 3.961950585 |
| Q3TC72 | Fahd2a | 1.193585317 | 2.799392317 |
| A2ATU0 | Dhtkd1 | 1.192637482 | 2.670258954 |
| Q91WD1 | Polr3d | 1.186497523 | 4.126845934 |
| Q05816 | Fabp5 | 1.185399918 | 3.302178621 |
| Q9Z1X4 | Ilf3 | 1.180738739 | 3.999130838 |
| Q9JL70 | Fanca | 1.178858662 | 2.576774461 |
| Q920Q6 | Msi2 | 1.16975357 | 5.564441488 |
| Q9ERA0 | Tfcp2 | 1.168857333 | 2.874392261 |
| Q9CYR0 | Ssbp1 | 1.166447987 | 2.128095175 |
| Q9D6M3 | Slc25a22 | 1.164089605 | 2.201156676 |
| Q8BH55 | Thnsl1 | 1.161398973 | 3.072581159 |
| A0A7H0DN84 | OPG112 | 1.139213723 | 2.658391358 |
| A0A7H0DND1 | OPG160 | 1.137472365 | 3.035139128 |
| Q8BMJ3 | Eif1ax | 1.13629262 | 2.787008916 |
| P11930 | Nudt19 | 1.126341492 | 2.416370706 |
| Q9EST4 | Psmg2 | 1.124489362 | 2.077691886 |
| Q9CXU9 | Eif1b | 1.10934232 | 2.622290554 |
| Q3V0K9 | Pls1 | 1.101994494 | 4.031828661 |
| Q6PCN7 | Hltf | 1.093235407 | 2.97182311 |
| Q9CXY6 | Ilf2 | 1.086761186 | 5.150538496 |
| Q2TPA8 | Hsdl2 | 1.085873107 | 4.117235238 |
| P59048 | Pdrg1 | 1.083823938 | 3.650530729 |
| Q45VK7 | Dync2h1 | 1.083010987 | 3.155213553 |
| Q9DD18 | Dtd1 | 1.081617519 | 2.016431981 |
| A0A7H0DN36 | OPG063 | 1.076285729 | 3.676867172 |
| A0A7H0DN25 | OPG052 | 1.074705978 | 3.752342019 |
| Q6DG52 | Churc1 | 1.069801961 | 3.304736627 |
| P62141 | Ppp1cb | 1.068163946 | 3.288391674 |
| A0A7H0DN23 | OPG050 | 1.065552354 | 3.477624463 |
| Q9ERI6 | Rdh14 | 1.045210644 | 2.183679058 |
| Q8K4F6 | Nsun5 | 1.041834721 | 2.30039889 |
| A0A7H0DN12 | OPG039 | 1.036056755 | 4.364040474 |
| A0A7H0DNA2 | OPG130 | 1.031404942 | 3.0651015 |
| A0A7H0DN10 | OPG037 | 1.030901805 | 2.963173982 |
| Q811J3 | Ireb2 | 1.030803212 | 4.073271516 |
| Q60668 | Hnrnpd | 1.028170673 | 4.229889932 |
| A0A7H0DND0 | OPG159 | 1.027061337 | 4.228189692 |
| Q8BIP0 | Dars2 | 1.026264015 | 3.333670833 |
| Q99J10 | Ctu1 | 1.020300986 | 2.049997399 |
| P56391 | Cox6b1 | 1.01957303 | 2.162871655 |
| A0A7H0DNA3 | OPG131 | 1.012193716 | 3.860283843 |
| Q8C167 | Prepl | 1.010130107 | 3.028865799 |
| Q8VDS4 | Rprd1a | 1.008248459 | 4.233409873 |
| M1L9M0 | OPG082 | 1.000333048 | 3.53839266 |
| E9Q634 | Myo1e | -1.00013495 | 3.001531555 |
| Q9JMA1 | Usp14 | -1.000561409 | 4.883446124 |
| Q8CJF7 | Ahctf1 | -1.000695402 | 3.457574441 |
| P17182 | Eno1 | -1.002145885 | 4.141007816 |
| Q3TC93 | Hs1bp3 | -1.002974765 | 3.186179563 |
| Q8BLN5 | Lss | -1.003810056 | 2.909903361 |
| O88878 | Zfand5 | -1.003950042 | 2.090299478 |
| Q9ERG2 | Strn3 | -1.004363534 | 5.305660207 |
| P47934 | Crat | -1.004878385 | 3.159490263 |
| F7BJB9 | Morc3 | -1.005009469 | 4.746578186 |
| P18653 | Rps6ka1 | -1.005158726 | 4.460119096 |
| Q6ZQF0 | Topbp1 | -1.006010482 | 3.379827241 |
| Q8K2H6 | Anapc10 | -1.00701682 | 2.075357669 |
| Q8BZR9 | Ncbp3 | -1.008225093 | 3.66754012 |
| P60670 | Nploc4 | -1.008310982 | 4.737436632 |
| Q8BMC4 | Nop9 | -1.008984561 | 3.737635644 |
| Q8CHI8 | Ep400 | -1.010012005 | 4.345533714 |
| Q8CCI5 | Rybp | -1.012262154 | 4.566740796 |
| Q922S8 | Kif2c | -1.012602618 | 3.432860982 |
| Q60710 | Samhd1 | -1.01342316 | 4.566740796 |
| Q61881 | Mcm7 | -1.013539217 | 5.32435078 |
| O55222 | Ilk | -1.015175259 | 4.847091878 |
| P16951 | Atf2 | -1.015377241 | 2.814807518 |
| Q9CQI3 | Gmfb | -1.016322789 | 2.525753525 |
| B2RQC6 | Cad | -1.016358191 | 4.602517587 |
| Q8BHL5 | Elmo2 | -1.016363503 | 4.846983301 |
| G5E8V9 | Arfip1 | -1.016424492 | 4.067545668 |
| P18760 | Cfl1 | -1.016731527 | 5.305288851 |
| Q91W36 | Usp3 | -1.018051955 | 2.812813424 |
| Q8BPY9 | Fignl1 | -1.018629509 | 2.493359043 |
| P40142 | Tkt | -1.019193841 | 3.440237675 |
| Q9JI10 | Stk3 | -1.019546769 | 4.651839245 |
| Q571I9 | Aldh16a1 | -1.020045623 | 4.429211747 |
| Q8CFI0 | Nedd4l | -1.020145855 | 3.457574441 |
| Q8C547 | Heatr5b | -1.02338893 | 4.439752883 |
| Q8K2C7 | Os9 | -1.025120413 | 2.614376375 |
| Q8R3L2 | Tcf25 | -1.028987687 | 2.727199454 |
| Q9D358 | Acp1 | -1.02929941 | 5.099297171 |
| Q5EG47 | Prkaa1 | -1.029495544 | 3.917593976 |
| Q80T85 | Dcaf5 | -1.031861672 | 2.50660225 |
| P08113 | Hsp90b1 | -1.032913378 | 4.263272019 |
| P56480 | Atp5f1b | -1.035177912 | 4.474124825 |
| P19182 | Ifrd1 | -1.037131181 | 2.824720432 |
| Q9CPY7 | Lap3 | -1.037146726 | 5.274807648 |
| P53996 | Cnbp | -1.039031205 | 5.134396157 |
| Q6ZWZ2 | Ube2r2 | -1.039031484 | 3.734639794 |
| Q9QYB1 | Clic4 | -1.040593941 | 4.259652185 |
| Q9CR00 | Psmd9 | -1.041072648 | 3.194460254 |
| P49718 | Mcm5 | -1.041214868 | 4.154766226 |
| Q62245 | Sos1 | -1.041229357 | 5.267753915 |
| Q9Z1E3 | Nfkbia | -1.041444549 | 3.077817815 |
| Q8C5W3 | Tbcel | -1.04253361 | 4.885828148 |
| P31938 | Map2k1 | -1.042685056 | 4.77579784 |
| Q9JHL1 | Nherf2 | -1.044789469 | 4.703028482 |
| Q8R0K9 | E2f4 | -1.046524641 | 3.247869799 |
| Q6PFE3 | Rad54b | -1.047475248 | 2.20974702 |
| Q60692 | Psmb6 | -1.047577788 | 2.341230564 |
| Q61584 | Fxr1 | -1.047931036 | 5.598055492 |
| G5E829 | Atp2b1 | -1.048045037 | 2.300322174 |
| Q6VN19 | Ranbp10 | -1.049046818 | 2.588760277 |
| Q9D832 | Dnajb4 | -1.049209975 | 3.479812827 |
| Q9R1X4 | Timeless | -1.052607274 | 3.802717195 |
| Q80TQ2 | Cyld | -1.052703004 | 3.34948799 |
| Q9Z248 | Aebp2 | -1.055536641 | 2.808099536 |
| Q8BQZ5 | Cpsf4 | -1.055750377 | 3.822801824 |
| Q9D6R2 | Idh3a | -1.055903526 | 4.645866109 |
| Q69ZA1 | Cdk13 | -1.057284982 | 4.092282459 |
| Q9CX86 | Hnrnpa0 | -1.057433423 | 4.218376287 |
| O55131 | Septin7 | -1.058012132 | 5.885144701 |
| A2AN08 | Ubr4 | -1.05881948 | 5.219426495 |
| Q80YV2 | Zc3hc1 | -1.058873042 | 3.548523338 |
| Q8BH48 | Ubap1 | -1.058915871 | 3.570100172 |
| Q80UK0 | Sestd1 | -1.059257648 | 2.064389141 |
| Q9QZ05 | Eif2ak4 | -1.059287502 | 3.826167348 |
| P59999 | Arpc4 | -1.061073172 | 4.021535489 |
| Q9Z2I9 | Sucla2 | -1.061995927 | 2.245556574 |
| Q8CIN4 | Pak2 | -1.06333354 | 4.132197229 |
| Q3UMT1 | Ppp1r12c | -1.063833073 | 2.291383317 |
| Q9ERD7 | Tubb3 | -1.066623255 | 3.468897239 |
| Q9CQJ1 | Higd2a | -1.066693285 | 2.074194113 |
| Q80UY2 | Kcmf1 | -1.06727617 | 3.610068128 |
| P53810 | Pitpna | -1.067363295 | 3.845464579 |
| Q80UV9 | Taf1 | -1.067762195 | 2.136757621 |
| A2AAY5 | Sh3pxd2b | -1.067786016 | 2.961486326 |
| Q9CS00 | Cactin | -1.067844283 | 3.8507144 |
| P0C0A3 | Chmp6 | -1.068394984 | 3.478932132 |
| Q8CI08 | Slain2 | -1.068794299 | 3.266238087 |
| Q8CHY6 | Gatad2a | -1.070510541 | 3.946362164 |
| Q924T7 | Rnf31 | -1.071159971 | 2.741841729 |
| Q6P5E4 | Uggt1 | -1.071493156 | 4.838625744 |
| Q9Z2M7 | Pmm2 | -1.072155172 | 4.260876391 |
| Q9QY76 | Vapb | -1.072477804 | 2.166244183 |
| Q08509 | Eps8 | -1.073342098 | 4.435219331 |
| A0A7H0DN02 | OPG027 | -1.074040653 | 3.261932566 |
| P97376 | Frg1 | -1.076402842 | 2.302722817 |
| P20029 | Hspa5 | -1.077219153 | 3.826167348 |
| Q5S006 | Lrrk2 | -1.077447941 | 2.059541817 |
| P54116 | Stom | -1.077707554 | 2.481058309 |
| Q9CXU1 | Med31 | -1.078082362 | 2.619877098 |
| Q8BK08 | Tmem11 | -1.078949 | 3.196657569 |
| Q8VCX5 | Micu1 | -1.082277801 | 4.008234337 |
| P13439 | Umps | -1.083239736 | 6.074962845 |
| Q5SSI6 | Utp18 | -1.083245067 | 2.19412277 |
| O54988 | Slk | -1.083306335 | 5.456448271 |
| Q8CBW3 | Abi1 | -1.083586489 | 4.57376385 |
| O54931 | Pakap | -1.083750114 | 3.971957233 |
| Q8BRN9 | Cc2d1b | -1.085384637 | 4.410517613 |
| Q61686 | Cbx5 | -1.085639635 | 4.937787386 |
| Q8C1A5 | Thop1 | -1.08563998 | 3.146700157 |
| Q9WTX8 | Mad1l1 | -1.087785887 | 4.620226859 |
| Q7TMB8 | Cyfip1 | -1.087812642 | 3.838549539 |
| Q8C4B4 | Unc119b | -1.088054133 | 2.472068902 |
| Q61083 | Map3k2 | -1.088326148 | 4.171488726 |
| P04184 | Tk1 | -1.088990214 | 3.541784102 |
| Q01320 | Top2a | -1.089619553 | 3.579283262 |
| P53564 | Cux1 | -1.089632403 | 3.049278708 |
| Q9D0M1 | Prpsap1 | -1.092807159 | 4.94750944 |
| Q9D0B6 | Pbdc1 | -1.092892245 | 4.054376181 |
| P61022 | Chp1 | -1.092999206 | 3.03431599 |
| O55023 | Impa1 | -1.094445885 | 2.919158465 |
| Q05BC3 | Eml1 | -1.094553507 | 2.883613623 |
| Q99PL5 | Rrbp1 | -1.095268644 | 3.773579754 |
| Q8BJU9 | Mtrf1l | -1.096399505 | 2.446462921 |
| Q61543 | Glg1 | -1.097058689 | 3.648164178 |
| P97452 | Bop1 | -1.097294759 | 3.774376841 |
| Q3TSG4 | Alkbh5 | -1.099222809 | 3.043655343 |
| Q9JHP7 | Poglut2 | -1.10082857 | 3.57318033 |
| Q80TZ9 | Rere | -1.102949197 | 2.945637321 |
| Q64727 | Vcl | -1.103109003 | 4.855585066 |
| Q8CJG0 | Ago2 | -1.104900383 | 5.124330148 |
| Q9D3E6 | Stag1 | -1.107169341 | 3.141566361 |
| Q8BPM2 | Map4k5 | -1.108864459 | 4.709437835 |
| Q8K2B3 | Sdha | -1.113971512 | 4.420495476 |
| Q922J3 | Clip1 | -1.116320474 | 4.231143282 |
| Q8K337 | Inpp5b | -1.11696239 | 2.428408294 |
| P08030 | Aprt | -1.117393915 | 3.174296158 |
| Q9D9K3 | Aven | -1.121644893 | 3.865078208 |
| P14873 | Map1b | -1.122113575 | 4.687420751 |
| P10711 | Tcea1 | -1.122192152 | 4.28081681 |
| P30415 | Nktr | -1.123315272 | 4.742705473 |
| Q9CVB6 | Arpc2 | -1.123474201 | 4.828924576 |
| Q99JY9 | Actr3 | -1.123558761 | 5.214663041 |
| A0A7H0DMZ8 | OPG022 | -1.125370204 | 4.491522958 |
| Q3UHF3 | Mier3 | -1.126646546 | 2.042628676 |
| P09041 | Pgk2 | -1.129600315 | 2.222502841 |
| Q8BZQ7 | Anapc2 | -1.130466976 | 2.996769446 |
| Q8K120 | Nfatc4 | -1.131022124 | 3.518332324 |
| Q91ZU6 | Dst | -1.131826504 | 5.062355206 |
| P47791 | Gsr | -1.132216296 | 4.580089942 |
| O88685 | Psmc3 | -1.132647492 | 5.229648104 |
| P27612 | Plaa | -1.134499708 | 5.712943736 |
| O88271 | Cfdp1 | -1.136093616 | 4.166065651 |
| P97363 | Sptlc2 | -1.138397446 | 3.576678788 |
| P56394 | Cox17 | -1.13842839 | 3.330107322 |
| Q8BKG3 | Ptk7 | -1.1385169 | 3.31089258 |
| Q8K2Q5 | Chchd7 | -1.138650665 | 3.505011352 |
| P26039 | Tln1 | -1.140222436 | 5.485443412 |
| Q9DCX2 | Atp5pd | -1.140355435 | 3.531278671 |
| Q5SV77 | Ggnbp2 | -1.14097087 | 3.33853278 |
| Q501J7 | Phactr4 | -1.142241759 | 3.161060623 |
| Q8VD62 | Bles03 | -1.142999419 | 3.802717195 |
| Q9D2H6 | Sp2 | -1.143215479 | 2.800437328 |
| Q91YR1 | Twf1 | -1.144292726 | 5.150538496 |
| Q9WU42 | Ncor2 | -1.145967096 | 3.468897239 |
| Q8BXK8 | Agap1 | -1.147069077 | 2.219549315 |
| Q64455 | Ptprj | -1.148513013 | 2.036276324 |
| Q3TB82 | Plekhf1 | -1.149650331 | 2.016431981 |
| P57776 | Eef1d | -1.154908852 | 5.32435078 |
| Q6PER3 | Mapre3 | -1.156214353 | 5.836217927 |
| Q9Z0N1 | Eif2s3x | -1.156335732 | 4.828924576 |
| Q8JZN5 | Acad9 | -1.156886171 | 2.72480391 |
| Q8C1S0 | Med19 | -1.156942527 | 3.297777922 |
| P61750 | Arf4 | -1.158652298 | 2.7100364 |
| Q60848 | Hells | -1.158674511 | 4.06026088 |
| Q9CW46 | Raver1 | -1.15881865 | 4.783501869 |
| Q68FE8 | Znf280d | -1.15983563 | 2.020965217 |
| Q3UE37 | Ube2z | -1.160176086 | 4.140710214 |
| Q9DAU1 | Cnpy3 | -1.160499474 | 4.321174517 |
| O55135 | Eif6 | -1.16138207 | 4.868473268 |
| Q6PCP5 | Mff | -1.161919074 | 2.396151081 |
| Q8K009 | Aldh1l2 | -1.162783327 | 4.020711843 |
| D3YXK2 | Safb | -1.163489304 | 3.418188002 |
| Q8BUY9 | Pggt1b | -1.163673149 | 2.213902803 |
| P61967 | Ap1s1 | -1.164414249 | 2.036196532 |
| Q9D787 | Ppil2 | -1.164775234 | 3.468897239 |
| Q61214 | Dyrk1a | -1.164810511 | 3.570100172 |
| P43346 | Dck | -1.166649694 | 3.588215931 |
| Q8K268 | Abcf3 | -1.167034013 | 4.94750944 |
| Q9EQ80 | Nif3l1 | -1.167921658 | 3.518332324 |
| P97477 | Aurka | -1.168198297 | 3.885865711 |
| E9Q5G3 | Kif23 | -1.169402624 | 3.746318656 |
| O88811 | Stam2 | -1.171693955 | 4.28081681 |
| P09411 | Pgk1 | -1.17494978 | 4.768744389 |
| Q8BHG1 | Nrdc | -1.175051481 | 5.229648104 |
| P35700 | Prdx1 | -1.178410498 | 2.915563167 |
| Q60876 | Eif4ebp1 | -1.181081982 | 2.507013207 |
| Q9JKR6 | Hyou1 | -1.181871472 | 3.73318606 |
| P21107 | Tpm3 | -1.183074142 | 3.18269284 |
| Q91ZR2 | Snx18 | -1.183123707 | 5.32435078 |
| Q80VJ2 | Sra1 | -1.18464766 | 4.603120306 |
| O70566 | Diaph2 | -1.185151408 | 4.237734352 |
| Q9D4D4 | Tktl2 | -1.185363893 | 2.273496896 |
| Q9Z0R6 | Itsn2 | -1.185871096 | 3.479812827 |
| Q9QZ23 | Nfu1 | -1.18687933 | 3.284501448 |
| P45878 | Fkbp2 | -1.186933718 | 4.258612036 |
| A2A9C3 | Szt2 | -1.187801716 | 3.401382477 |
| Q8BJM5 | Slc30a6 | -1.188750513 | 2.826825468 |
| Q91YW3 | Dnajc3 | -1.190539318 | 3.882209749 |
| Q9DBN5 | Lonp2 | -1.191406499 | 3.808665166 |
| Q810J8 | Zfyve1 | -1.19205717 | 2.971043268 |
| Q9D3D0 | Ttpal | -1.192122797 | 2.426046785 |
| P33215 | Nedd1 | -1.19234278 | 4.075092917 |
| Q9JL62 | Gltp | -1.19543037 | 3.403744172 |
| Q8K274 | Fn3krp | -1.1958886 | 2.67186203 |
| Q9CY18 | Snx7 | -1.196346875 | 5.573184379 |
| P62137 | Ppp1ca | -1.196376428 | 4.036374954 |
| Q8BG51 | Rhot1 | -1.196840583 | 3.909311167 |
| Q0KL02 | Trio | -1.197898631 | 4.910967547 |
| P35831 | Ptpn12 | -1.198733281 | 5.040454588 |
| Q9JLQ2 | Git2 | -1.200277732 | 3.767177067 |
| Q9ES46 | Parvb | -1.202240681 | 3.795884272 |
| Q0VEE6 | Znf800 | -1.202992135 | 2.191987132 |
| Q8C166 | Cpne1 | -1.204312046 | 3.418850572 |
| Q9R207 | Nbn | -1.205704149 | 3.987761945 |
| Q9DBJ1 | Pgam1 | -1.21327211 | 5.947495263 |
| O35887 | Calu | -1.213801683 | 4.063211844 |
| P62192 | Psmc1 | -1.213988367 | 5.404447568 |
| P56399 | Usp5 | -1.214815858 | 5.720581761 |
| Q3TNA1 | Xylb | -1.216338089 | 2.104540302 |
| Q3UFY7 | Nt5c3b | -1.217402559 | 2.574071954 |
| Q9JJ78 | Pbk | -1.217683892 | 3.127084139 |
| Q6PNC0 | Dmxl1 | -1.218212149 | 5.155459614 |
| Q9CY34 | Ube2f | -1.220986723 | 3.845464579 |
| Q9CZ44 | Nsfl1c | -1.222650771 | 4.366752465 |
| Q9QZM0 | Ubqln2 | -1.224559132 | 4.916992165 |
| Q9Z1S0 | Bub1b | -1.226578091 | 4.990428188 |
| Q99KH8 | Stk24 | -1.226590169 | 4.936841965 |
| Q9JJ28 | Flii | -1.228820969 | 4.548988157 |
| Q9QZ88 | Vps29 | -1.229050137 | 4.740853929 |
| Q922R5 | Ppp4r3b | -1.229054514 | 3.460326362 |
| Q6PHZ2 | Camk2d | -1.230999176 | 4.977733831 |
| P70441 | Nherf1 | -1.231654872 | 3.398939489 |
| Q9DCD0 | Pgd | -1.231756323 | 5.158952867 |
| Q8BWT5 | Dip2a | -1.232010982 | 3.213849536 |
| Q8BTY2 | Slc4a7 | -1.239276184 | 3.588606435 |
| P55194 | Sh3bp1 | -1.240068824 | 2.917089237 |
| Q9EPC1 | Parva | -1.240873296 | 4.114501893 |
| Q80XC6 | Nrde2 | -1.246984836 | 3.236920238 |
| Q6DVA0 | Lemd2 | -1.247992305 | 2.617979483 |
| Q91VX2 | Ubap2 | -1.248280922 | 5.947495263 |
| Q923T9 | Camk2g | -1.2499224 | 4.603958583 |
| Q8R149 | Bud13 | -1.252496337 | 3.027449461 |
| Q9WVS8 | Mapk7 | -1.254807052 | 2.865400187 |
| P70270 | Rad54l | -1.255656764 | 2.199314326 |
| Q91VH6 | Memo1 | -1.25584157 | 4.656736291 |
| Q5SF07 | Igf2bp2 | -1.255883769 | 4.118825136 |
| Q9CQ39 | Med21 | -1.256527439 | 3.883383866 |
| Q811D0 | Dlg1 | -1.256536843 | 4.580980642 |
| Q8CHU3 | Epn2 | -1.258125542 | 5.090854674 |
| Q9ER41 | Tor1b | -1.258268049 | 4.447949878 |
| Q9JII6 | Akr1a1 | -1.259264041 | 3.449303506 |
| Q8BJS4 | Sun2 | -1.260368018 | 2.551114425 |
| Q62188 | Dpysl3 | -1.261412622 | 4.695022215 |
| Q9DB77 | Uqcrc2 | -1.263464915 | 3.532585291 |
| Q8CFJ9 | Wdr24 | -1.265658115 | 3.341607078 |
| Q9CQ60 | Pgls | -1.267528831 | 4.011228075 |
| Q91W59 | Rbms1 | -1.267688469 | 4.544507578 |
| Q80WE4 | Kif20b | -1.269799257 | 3.800955401 |
| Q9ER73 | Elp4 | -1.270953786 | 3.035477383 |
| Q3UGR5 | Hdhd2 | -1.272892797 | 4.382488688 |
| Q8C0I4 | Epc2 | -1.274427446 | 2.280788353 |
| Q9CYN9 | Atp6ap2 | -1.276089262 | 4.073271516 |
| A0A7H0DNF5 | OPG193 | -1.280560076 | 4.6806093 |
| Q9QUR6 | Prep | -1.284679989 | 4.981870438 |
| P27641 | Xrcc5 | -1.284797847 | 2.445005617 |
| O88587 | Comt | -1.290661297 | 3.958627787 |
| Q8R146 | Apeh | -1.2909743 | 4.7892525 |
| P54071 | Idh2 | -1.292218942 | 5.274721676 |
| O35379 | Abcc1 | -1.294388891 | 2.936567484 |
| P09103 | P4hb | -1.294530484 | 4.828924576 |
| Q922E4 | Pcyt2 | -1.295236527 | 4.073271516 |
| P0DTN0 | OPG002 | -1.300836628 | 3.183339002 |
| Q922H2 | Pdk3 | -1.300937982 | 4.678988776 |
| Q8CIG3 | Kdm1b | -1.302546207 | 2.725963974 |
| Q91YS8 | Camk1 | -1.304218771 | 5.23619099 |
| Q9CZN7 | Shmt2 | -1.306624253 | 5.456448271 |
| P70349 | Hint1 | -1.306936062 | 5.069107293 |
| Q99ME2 | Wdr6 | -1.309437814 | 4.30079528 |
| Q3UE17 | Mex3d | -1.310414117 | 2.870388441 |
| Q61687 | Atrx | -1.312745712 | 4.763897782 |
| Q8BKX1 | Baiap2 | -1.314274153 | 5.217754918 |
| P14211 | Calr | -1.316912732 | 3.860640783 |
| Q9WUU8 | Tnip1 | -1.319418827 | 3.489270602 |
| O09172 | Gclm | -1.319614598 | 3.020074884 |
| Q9ERH4 | Nusap1 | -1.32058938 | 2.029109882 |
| Q8CHK4 | Kat5 | -1.322129276 | 2.925186801 |
| Q3TUA9 | Pomk | -1.324421549 | 2.217491687 |
| Q69Z38 | Peak1 | -1.328473602 | 2.998684483 |
| Q6P9L6 | Kif15 | -1.329097044 | 5.068810487 |
| P03958 | Ada | -1.333176681 | 3.963825463 |
| Q9CR86 | Carhsp1 | -1.335220053 | 4.768744389 |
| Q9JJN5 | Cpn1 | -1.335697915 | 2.225176318 |
| Q8CJ53 | Trip10 | -1.337867755 | 3.429140305 |
| P21981 | Tgm2 | -1.339781448 | 3.335413139 |
| Q9QWT9 | Kifc1 | -1.341919029 | 2.512243063 |
| Q8BP47 | Nars1 | -1.344471793 | 5.485443412 |
| P58871 | Tnks1bp1 | -1.345727305 | 4.745015107 |
| Q02819 | Nucb1 | -1.347826521 | 4.831304084 |
| P35293 | Rab18 | -1.348638639 | 4.060801174 |
| Q80Y50 | Camta2 | -1.350655357 | 2.105061757 |
| O70404 | Vamp8 | -1.352627953 | 3.4042273 |
| Q9DAR7 | Dcps | -1.353987202 | 5.086829807 |
| Q3UHJ0 | Aak1 | -1.35443884 | 5.130024203 |
| Q6RHR9 | Magi1 | -1.355413001 | 2.267469288 |
| Q8BVE8 | Nsd2 | -1.355736604 | 4.876898319 |
| P70404 | Idh3g | -1.356576734 | 4.665074767 |
| Q8BI72 | Cdkn2aip | -1.358236909 | 4.236364123 |
| Q6A0A9 | FAM120A | -1.359596508 | 6.53644885 |
| Q8K0W9 | Dph3 | -1.360188479 | 2.088979068 |
| Q9R1S8 | Capn7 | -1.360800898 | 2.305146348 |
| Q8BVI4 | Qdpr | -1.361201368 | 4.334919866 |
| Q920R0 | Als2 | -1.363840617 | 2.240000145 |
| O88456 | Capns1 | -1.363910053 | 4.065931001 |
| Q80UG5 | Septin9 | -1.364151961 | 6.457680354 |
| Q7TPV2 | Dzip3 | -1.364543824 | 3.61261773 |
| Q8R4U7 | Luzp1 | -1.365610297 | 2.65570546 |
| Q91W40 | Klc3 | -1.367003707 | 3.327504148 |
| Q9D1N9 | Mrpl21 | -1.372212265 | 2.379070025 |
| Q9Z2R6 | Unc119 | -1.374423295 | 3.42553264 |
| P47713 | Pla2g4a | -1.376774235 | 4.491522958 |
| Q9D662 | Sec23b | -1.37944295 | 5.159507659 |
| Q61029 | Tmpo | -1.381771345 | 2.263251505 |
| Q06180 | Ptpn2 | -1.3818358 | 3.128621352 |
| Q9DB00 | Gon4l | -1.382519603 | 3.00273901 |
| Q7TSG2 | Ctdp1 | -1.386428374 | 5.076280368 |
| Q9JK81 | Myg1 | -1.386917039 | 2.891798486 |
| Q8C3Y4 | Kntc1 | -1.391645612 | 3.672863989 |
| P46414 | Cdkn1b | -1.391827702 | 2.025525192 |
| Q64516 | Gk | -1.391971655 | 2.268922464 |
| Q920Q8 | Ivns1abp | -1.404447268 | 4.160464863 |
| O54984 | Get3 | -1.406201206 | 5.402247759 |
| P18242 | Ctsd | -1.408997969 | 5.081821909 |
| Q9Z2D1 | Mtmr2 | -1.416639566 | 3.418875336 |
| Q8VBT9 | Aspscr1 | -1.419405232 | 3.534652012 |
| Q9DBC7 | Prkar1a | -1.423709829 | 4.482379556 |
| Q01853 | Vcp | -1.424976569 | 5.364449318 |
| P52480 | Pkm | -1.427616643 | 5.720581761 |
| Q9DB20 | Atp5po | -1.428235004 | 3.030769383 |
| Q9D902 | Gtf2e2 | -1.429109387 | 4.154766226 |
| Q99MK9 | Rassf1 | -1.433107848 | 2.317649218 |
| Q8CB77 | Eloa | -1.436246994 | 3.594930165 |
| Q9JM14 | Nt5c | -1.438351077 | 4.154749573 |
| Q9EPV8 | Ubl5 | -1.441767439 | 3.801925542 |
| Q8CH72 | Trim32 | -1.444401483 | 3.54834489 |
| Q8CGK3 | Lonp1 | -1.446239675 | 4.382256 |
| Q9ES00 | Ube4b | -1.446386056 | 5.002220191 |
| P50544 | Acadvl | -1.446762333 | 3.195766442 |
| Q64674 | Srm | -1.451371837 | 5.9667263 |
| Q9CZX0 | Elp3 | -1.45259166 | 3.53839266 |
| O35350 | Capn1 | -1.456666529 | 4.166111963 |
| P70459 | Erf | -1.456816929 | 2.778109046 |
| O08664 | Bcl7c | -1.458003113 | 3.518702918 |
| Q8R059 | Gale | -1.458782955 | 3.653991622 |
| P98083 | Shc1 | -1.461978206 | 2.808099536 |
| P47199 | Cryz | -1.462622653 | 2.544854995 |
| B2RRE7 | Otud4 | -1.463880435 | 3.980153935 |
| Q91V12 | Acot7 | -1.464138152 | 5.60091374 |
| P47226 | Tes | -1.465202799 | 4.883446124 |
| Q8BTY1 | Kyat1 | -1.466202727 | 3.304919565 |
| O08638 | Myh11 | -1.482533854 | 3.695887977 |
| Q9D7I8 | Fam83d | -1.484150548 | 2.131229641 |
| Q8BW96 | Camk1d | -1.485992672 | 3.563353288 |
| O08784 | Tcof1 | -1.48872415 | 3.254486056 |
| Q9DBS9 | Osbpl3 | -1.488860529 | 4.091619504 |
| A6H8H2 | Dennd4c | -1.493992215 | 4.777690664 |
| Q80W47 | Wipi2 | -1.495320621 | 3.490564936 |
| Q9JMH9 | Myo18a | -1.497323326 | 3.477624463 |
| P27773 | Pdia3 | -1.498254412 | 4.566740796 |
| Q68FL4 | Ahcyl2 | -1.498967447 | 4.348653145 |
| P33174 | Kif4 | -1.501036915 | 3.86532435 |
| Q8BT07 | Cep55 | -1.502359686 | 3.752342019 |
| Q5FWK3 | Arhgap1 | -1.505082424 | 5.60091374 |
| Q3TJD7 | Pdlim7 | -1.508236606 | 5.253470976 |
| P51480 | Cdkn2a | -1.50844937 | 5.307006512 |
| A2RSY1 | Kansl3 | -1.509333654 | 2.332721499 |
| A2AMM0 | Cavin4 | -1.509448486 | 3.063022727 |
| O08532 | Cacna2d1 | -1.51203696 | 2.373077809 |
| P06801 | Me1 | -1.513357555 | 5.185028475 |
| Q9JHJ0 | Tmod3 | -1.514641893 | 4.601953238 |
| Q9ER81 | Tor1aip2 | -1.523985055 | 3.398144895 |
| P05480 | Src | -1.526767313 | 2.413104223 |
| P22315 | Fech | -1.527788079 | 2.166480359 |
| Q9DCN2 | Cyb5r3 | -1.529123973 | 2.471649265 |
| Q9Z0U1 | Tjp2 | -1.530217672 | 5.507289836 |
| P06151 | Ldha | -1.530303729 | 3.490956993 |
| Q8BUY5 | Timmdc1 | -1.530709058 | 2.860741448 |
| P60766 | Cdc42 | -1.533134578 | 4.166065651 |
| Q8CDM1 | Atad2 | -1.535504006 | 3.735632031 |
| P13864 | Dnmt1 | -1.537438447 | 4.401282148 |
| P37913 | Lig1 | -1.542297914 | 5.229648104 |
| Q68EF0 | Rab3ip | -1.547327782 | 2.732229178 |
| Q921F4 | Hnrnpll | -1.550198014 | 6.239839385 |
| Q06186 | Hbegf | -1.550755929 | 2.451282317 |
| P70445 | Eif4ebp2 | -1.552211234 | 2.816402739 |
| Q9R0P3 | Esd | -1.55286116 | 5.002220191 |
| Q99P72 | Rtn4 | -1.553473451 | 4.062858541 |
| Q91W89 | Man2c1 | -1.553820442 | 2.450808029 |
| Q8BM55 | Tmem214 | -1.554765659 | 2.323223396 |
| O08529 | Capn2 | -1.554977313 | 4.977733831 |
| Q8K1J6 | Trnt1 | -1.557575545 | 3.838072707 |
| Q8BH59 | Slc25a12 | -1.557809731 | 3.51019309 |
| Q99104 | Myo5a | -1.565685799 | 5.5821464 |
| Q9QZW0 | Atp11c | -1.570413153 | 3.532585291 |
| Q8K0B2 | Lmbrd1 | -1.573194637 | 2.143436206 |
| O70445 | Bard1 | -1.580104597 | 2.114041802 |
| Q9EPK5 | Wwtr1 | -1.581026398 | 2.625094 |
| Q8K1L5 | Ppp1r11 | -1.582694889 | 3.634513424 |
| Q99KS6 | Adprm | -1.585317225 | 5.255333746 |
| Q9D0M3 | Cyc1 | -1.588870336 | 3.030976578 |
| Q9D198 | Syf2 | -1.592790735 | 3.720801553 |
| Q8R3B1 | Plcd1 | -1.595875818 | 2.153101047 |
| Q07076 | Anxa7 | -1.596360729 | 2.368472841 |
| Q9WV30 | Nfat5 | -1.601456338 | 3.559197605 |
| Q8CBE3 | Wdr37 | -1.601517904 | 3.224410025 |
| Q920A5 | Scpep1 | -1.60226957 | 4.7892525 |
| Q8K3X4 | Irf2bpl | -1.60278085 | 3.597182542 |
| Q8CHG7 | Rapgef2 | -1.604932233 | 2.059296192 |
| Q8R4Y8 | Rttn | -1.606269924 | 3.546691444 |
| P59108 | Cpne2 | -1.608715324 | 3.297777922 |
| Q9Z1Q9 | Vars1 | -1.609296327 | 5.947495263 |
| Q99LD8 | Ddah2 | -1.609464026 | 5.364449318 |
| Q4VAA7 | Snx33 | -1.609498387 | 2.43168231 |
| Q9EP71 | Rai14 | -1.610423503 | 5.60091374 |
| Q9DBG9 | Tax1bp3 | -1.616539377 | 4.7363708 |
| Q9CX30 | Yif1b | -1.618673872 | 2.236022208 |
| Q8C8U0 | Ppfibp1 | -1.61934244 | 4.74201806 |
| Q8BGH7 | Cdc42se2 | -1.626761533 | 2.977204501 |
| Q8BL95 | Cfap298 | -1.627833181 | 3.840185978 |
| P08103 | Hck | -1.631756295 | 3.186179563 |
| Q62523 | Zyx | -1.631917309 | 3.297092711 |
| P52623 | Uck1 | -1.634175183 | 3.304141289 |
| Q9QYI4 | Dnajb12 | -1.639431124 | 2.252196179 |
| Q3V3Q7 | Pacs2 | -1.643501853 | 2.245955308 |
| Q8BU85 | Msrb3 | -1.645251364 | 3.624600882 |
| Q9WUM3 | Coro1b | -1.646919407 | 4.311198228 |
| Q9D1I5 | Mcee | -1.648901883 | 2.141728201 |
| Q8VHX2 | Edaradd | -1.649721941 | 4.067545668 |
| Q8VDZ4 | Zdhhc5 | -1.649943887 | 2.765338113 |
| P02469 | Lamb1 | -1.652055895 | 4.429211747 |
| Q8R3C6 | Rbm19 | -1.655288662 | 5.507289836 |
| Q8K3W3 | Casc3 | -1.655907616 | 2.310123767 |
| Q64514 | Tpp2 | -1.66280442 | 7.133103323 |
| L0N7N1 | Kif14 | -1.663623476 | 2.377316523 |
| P12388 | Serpinb2 | -1.665321281 | 3.975894136 |
| Q08775 | Runx2 | -1.667789924 | 2.020804017 |
| Q7TT50 | Cdc42bpb | -1.667864735 | 5.214864811 |
| O35385 | Ppef2 | -1.669146806 | 2.872894663 |
| Q9CQ92 | Fis1 | -1.672922992 | 4.94750944 |
| Q6PDH0 | Phldb1 | -1.673143282 | 2.763757235 |
| Q5PRF0 | Heatr5a | -1.681478064 | 3.570179599 |
| Q9DBS2 | Tprg1l | -1.683802786 | 2.038110416 |
| P31266 | Rbpj | -1.688954512 | 4.044351482 |
| O35874 | Slc1a4 | -1.694821427 | 2.451126966 |
| Q3TZZ7 | Esyt2 | -1.69573578 | 4.428869132 |
| Q8BSY0 | Asph | -1.707178302 | 2.938936844 |
| Q9CYL5 | Glipr2 | -1.708693916 | 2.865400187 |
| Q9WTK7 | Stk11 | -1.709581705 | 3.58454419 |
| P63028 | Tpt1 | -1.711750744 | 4.551003354 |
| Q924A2 | Cic | -1.716547226 | 2.348715313 |
| P50431 | Shmt1 | -1.720277093 | 4.535801556 |
| P19324 | Serpinh1 | -1.724778791 | 4.782877206 |
| P02798 | Mt2 | -1.725509265 | 4.783501869 |
| Q9R0A0 | Pex14 | -1.729144567 | 2.872895852 |
| Q3TAS6 | Emc10 | -1.731965024 | 2.141728201 |
| Q8BYB9 | Poglut1 | -1.735275641 | 4.031828661 |
| Q9D968 | Hcfc2 | -1.735618657 | 3.330107322 |
| P60710 | Actb | -1.735716886 | 4.536961494 |
| Q922J9 | Far1 | -1.744229108 | 2.984226327 |
| Q9Z1B3 | Plcb1 | -1.753599607 | 2.433034141 |
| Q6ZQ73 | Cand2 | -1.756100036 | 3.693186434 |
| Q9DBU0 | Tm9sf1 | -1.758099015 | 2.551523964 |
| Q9R0N0 | Galk1 | -1.758301114 | 4.10869914 |
| O08691 | Arg2 | -1.766742454 | 3.801002364 |
| Q8K0Q5 | Arhgap18 | -1.772889999 | 2.128561491 |
| P17809 | Slc2a1 | -1.773700008 | 2.865498045 |
| Q64430 | Atp7a | -1.775718987 | 2.948317873 |
| O54946 | Dnajb6 | -1.779442594 | 3.727470885 |
| P39428 | Traf1 | -1.782768061 | 2.095412107 |
| P61025 | Cks1b | -1.785620835 | 4.227615259 |
| P97384 | Anxa11 | -1.786009549 | 3.245348704 |
| Q03145 | Epha2 | -1.786461892 | 3.695887977 |
| Q61810 | Ltbp3 | -1.797850904 | 2.677621326 |
| O70433 | Fhl2 | -1.804670815 | 3.828987274 |
| Q99KI0 | Aco2 | -1.809246236 | 5.507289836 |
| Q8R127 | Sccpdh | -1.815812719 | 3.47470681 |
| P35601 | Rfc1 | -1.823278359 | 4.56537048 |
| Q91VW3 | Sh3bgrl3 | -1.826299585 | 6.070246952 |
| Q91W67 | Ubl7 | -1.826952769 | 6.081184158 |
| Q60716 | P4ha2 | -1.827253753 | 4.169506422 |
| Q9CWV6 | Prkrip1 | -1.827540635 | 4.943475205 |
| P16858 | Gapdh | -1.828789836 | 4.802990413 |
| Q91X58 | **Zfand2b** | -1.830685541 | 3.807027436 |
| Q6PB44 | Ptpn23 | -1.831632855 | 5.550356867 |
| Q69ZK6 | Jmjd1c | -1.837016076 | 2.697035914 |
| Q9WVL2 | Stat2 | -1.838069302 | 3.69198754 |
| P51432 | Plcb3 | -1.841287511 | 4.537539953 |
| Q62465 | Vat1 | -1.844045248 | 5.485443412 |
| Q60875 | Arhgef2 | -1.848528518 | 4.937787386 |
| Q00612 | G6pdx | -1.852557213 | 4.911396089 |
| Q9JK91 | Mlh1 | -1.860060632 | 2.431027761 |
| Q6ZWY3 | Rps27l | -1.862826356 | 4.619630995 |
| P70290 | Mpp1 | -1.863901114 | 4.620492661 |
| Q60715 | P4ha1 | -1.868329155 | 4.435219331 |
| Q61207 | Psap | -1.868679797 | 3.554111488 |
| Q8VHI3 | Pofut2 | -1.871782745 | 2.775681387 |
| O08601 | Mttp | -1.873686694 | 2.861883958 |
| Q7TPR4 | Actn1 | -1.875077224 | 6.34689266 |
| Q9QY66 | Znhit2 | -1.881171265 | 3.855861341 |
| Q8N7N5 | Dcaf8 | -1.884566022 | 4.64119677 |
| O70306 | Tbx15 | -1.887723462 | 2.101454453 |
| Q8R1F1 | Niban2 | -1.891007617 | 4.883446124 |
| Q0GNC1 | Inf2 | -1.894907428 | 4.250912104 |
| Q3TLI0 | Trappc10 | -1.895021168 | 2.883613623 |
| P23591 | Gfus | -1.899107067 | 5.146289744 |
| P08752 | Gnai2 | -1.901770094 | 3.124139346 |
| Q61739 | Itga6 | -1.902603572 | 3.699028907 |
| O08599 | Stxbp1 | -1.903112443 | 5.30196706 |
| Q91VH2 | Snx9 | -1.90386606 | 5.135241031 |
| Q80XC3 | Usp6nl | -1.906927085 | 2.948317873 |
| Q6ZQJ5 | Dna2 | -1.908148635 | 3.05552025 |
| O70475 | Ugdh | -1.912748664 | 5.553900853 |
| Q9Z247 | Fkbp9 | -1.916317676 | 3.593909852 |
| O35295 | Purb | -1.918950443 | 5.394683131 |
| P10922 | H1-0 | -1.922749146 | 2.918323723 |
| P43275 | H1-1 | -1.929004967 | 2.432802497 |
| Q3UZ18 | Ice2 | -1.932211204 | 2.68539894 |
| D3Z4S3 | Ptrhd1 | -1.93917003 | 3.220384332 |
| P56212 | Arpp19 | -1.943561297 | 2.997423249 |
| O09130 | Nfatc2ip | -1.944595019 | 2.52154807 |
| Q9D2R8 | Mrps33 | -1.945901745 | 2.630301856 |
| Q91ZV0 | Mia2 | -1.946438875 | 2.299286703 |
| P00375 | Dhfr | -1.9503603 | 4.747691102 |
| P56390 | Cks2 | -1.952066615 | 3.685324542 |
| Q3UMF0 | Cobll1 | -1.95359889 | 2.180465722 |
| Q9CQ65 | Mtap | -1.955755904 | 5.771121261 |
| Q60793 | Klf4 | -1.959912779 | 2.297037097 |
| Q921I6 | Sh3bp4 | -1.960982554 | 2.130988645 |
| Q91XE4 | Acy3 | -1.966447748 | 2.264563327 |
| Q9CQG3 | Zmynd19 | -1.969298041 | 2.044437593 |
| P97432 | Nbr1 | -1.97110537 | 2.474019304 |
| Q80TM9 | Nisch | -1.971778281 | 5.013779153 |
| Q8BNV1 | Trmt2a | -1.973860525 | 4.758922163 |
| Q9CWT3 | Snx10 | -1.977566984 | 3.994132267 |
| Q9D3D9 | Atp5f1d | -1.978668151 | 2.701107754 |
| Q8R4K2 | Irak4 | -1.986759309 | 2.455527183 |
| Q8K136 | Scnm1 | -1.997789895 | 2.562448668 |
| Q60597 | Ogdh | -2.006429249 | 5.638382919 |
| Q61490 | Alcam | -2.009034582 | 3.53839266 |
| Q91VW5 | Golga4 | -2.014444321 | 2.051201079 |
| P35123 | Usp4 | -2.01687413 | 5.456448271 |
| Q7SIG6 | Asap2 | -2.019080507 | 2.222502841 |
| Q99N85 | Mrps18a | -2.031726221 | 3.996800587 |
| Q9DB54 | Fam216a | -2.032057463 | 3.512154444 |
| Q9Z0R0 | Haspin | -2.034440598 | 2.16540525 |
| P81122 | Irs2 | -2.044381616 | 2.912748743 |
| Q69ZI1 | Sh3rf1 | -2.044626664 | 3.039410074 |
| P48036 | Anxa5 | -2.044645723 | 2.051201079 |
| Q3UHH1 | Zswim8 | -2.045476975 | 4.765542429 |
| Q3U0M1 | Trappc9 | -2.048960454 | 2.635777403 |
| P31324 | Prkar2b | -2.049262943 | 5.150538496 |
| Q3TMW1 | Ccdc102a | -2.049451667 | 4.235972649 |
| O88342 | Wdr1 | -2.059640009 | 5.615952409 |
| Q8R5A6 | Tbc1d22a | -2.060480918 | 2.913837835 |
| Q9QUR8 | Sema7a | -2.062483919 | 2.682880487 |
| Q9D1M7 | Fkbp11 | -2.072316921 | 2.331657096 |
| Q8VC03 | Eml3 | -2.081497117 | 2.06162451 |
| Q9CPQ5 | Cenpq | -2.083184776 | 2.037916081 |
| Q64092 | Tfe3 | -2.092128437 | 3.08515606 |
| P30681 | Hmgb2 | -2.094148973 | 5.940647608 |
| O08785 | Clock | -2.095963737 | 3.908210268 |
| P48760 | Fpgs | -2.107259117 | 2.197595772 |
| A0A7H0DN00 | OPG023 | -2.108177313 | 4.479171456 |
| Q8VDD5 | Myh9 | -2.109563021 | 6.239839385 |
| O35704 | Sptlc1 | -2.110011225 | 2.206046081 |
| Q64267 | Xpa | -2.110512767 | 2.396645904 |
| Q8BTM8 | Flna | -2.113943242 | 6.384665185 |
| P43277 | H1-3 | -2.118437125 | 2.296391706 |
| Q8C1Z7 | Bbs4 | -2.119797211 | 2.635038722 |
| Q91XB0 | Trex1 | -2.120199898 | 3.186391383 |
| Q5DTT3 | Tasor2 | -2.121273564 | 2.459657639 |
| Q8R3P2 | Dtx2 | -2.125524439 | 2.806357592 |
| Q61792 | Lasp1 | -2.133369475 | 5.846780448 |
| O70311 | Nmt2 | -2.135650092 | 3.860620749 |
| P61028 | Rab8b | -2.137564353 | 2.776534761 |
| Q5SXJ3 | Brip1 | -2.137603141 | 2.909934292 |
| Q9CQ20 | Mid1ip1 | -2.137813363 | 2.15373457 |
| P97329 | Kif20a | -2.137870732 | 5.057464996 |
| Q69ZF3 | Gba2 | -2.143034341 | 2.385143018 |
| Q9CQV4 | Retreg3 | -2.146448203 | 2.123680914 |
| Q91WK7 | Ankrd54 | -2.155109925 | 2.227418511 |
| P46718 | Pdcd2 | -2.156855481 | 2.611784357 |
| Q9CR14 | Fancl | -2.160666988 | 2.551214069 |
| Q9JIM1 | Slc29a1 | -2.161464556 | 2.881566838 |
| Q923Q2 | Stard13 | -2.16517311 | 2.106364922 |
| Q8BG73 | Sh3bgrl2 | -2.167063911 | 5.615952409 |
| P20152 | Vim | -2.167951016 | 5.551659592 |
| Q8K3Z9 | Pom121 | -2.186712354 | 2.118531383 |
| Q64311 | Ntan1 | -2.192245177 | 3.731438579 |
| O35621 | Pmm1 | -2.19501698 | 2.424931436 |
| Q8BJH1 | Zc2hc1a | -2.195716927 | 2.178600018 |
| Q810L3 | Chfr | -2.198584354 | 2.249071981 |
| Q76LS9 | Mindy1 | -2.199043868 | 3.860620749 |
| Q8VI36 | Pxn | -2.205326534 | 5.699939096 |
| Q9D084 | Cenps | -2.2063411 | 2.889329288 |
| F8VPU2 | Farp1 | -2.206573836 | 3.86038664 |
| Q8CAB8 | Castor2 | -2.20713297 | 2.263796242 |
| Q91WT8 | Rbm47 | -2.20790886 | 2.264080272 |
| Q8K212 | Pacs1 | -2.208927997 | 4.813039398 |
| Q9JJ11 | Tacc3 | -2.21059766 | 3.624245149 |
| Q9DB42 | Znf593 | -2.211359856 | 4.706971445 |
| Q52KR3 | Prune2 | -2.213336749 | 2.901017824 |
| Q80X90 | Flnb | -2.226472679 | 7.374803414 |
| Q9D771 | Pacc1 | -2.229544706 | 2.644501028 |
| Q9R059 | Fhl3 | -2.23129732 | 4.868941577 |
| Q8K2Y9 | Ccm2 | -2.232257014 | 3.341607078 |
| Q8BIG7 | Comtd1 | -2.248096907 | 3.941279595 |
| Q8BPZ8 | Abraxas1 | -2.248145853 | 2.165759082 |
| P12382 | Pfkl | -2.250041565 | 6.239839385 |
| Q9D8L5 | Ccdc91 | -2.250861572 | 2.70636947 |
| Q80W14 | Prpf40b | -2.255963422 | 2.912179584 |
| Q8CIL4 | Fsaf1 | -2.256949833 | 2.534608901 |
| Q3UMR5 | Mcu | -2.259834121 | 4.477032764 |
| Q9JLG8 | Capn15 | -2.265030625 | 2.545402136 |
| P21995 | Emb | -2.26609747 | 3.516720858 |
| Q9CXP8 | Gng10 | -2.267293763 | 2.701107754 |
| P38060 | Hmgcl | -2.267562181 | 3.871175789 |
| Q07113 | Igf2r | -2.277778593 | 3.130857045 |
| Q8CHE4 | Phlpp1 | -2.278943292 | 3.096180144 |
| Q8CDN6 | Txnl1 | -2.285215015 | 5.771121261 |
| Q9DBW3 | Natd1 | -2.291605638 | 2.002638517 |
| Q5SV80 | Myo19 | -2.307310682 | 2.607235217 |
| Q8C115 | Plekhh2 | -2.312975614 | 3.17698542 |
| Q5EE38 | Acd | -2.315137353 | 3.192666777 |
| Q9EQ15 | Gnb1l | -2.315853401 | 2.17720593 |
| P50637 | Tspo | -2.316429459 | 3.127354611 |
| Q8BHC4 | Dcakd | -2.323909292 | 2.287437595 |
| P27048 | Snrpb | -2.32447414 | 2.34360461 |
| Q7TPS0 | Rps6ka6 | -2.336722287 | 4.114501893 |
| O08553 | Dpysl2 | -2.337803369 | 5.947495263 |
| O09005 | Degs1 | -2.338015502 | 4.229889932 |
| Q9D0K2 | Oxct1 | -2.339758941 | 5.013779153 |
| Q9D8U7 | Dtwd1 | -2.341334537 | 3.376511645 |
| Q6P6I6 | Polr2m | -2.34325653 | 3.845464579 |
| Q6GV12 | Kdsr | -2.345663346 | 3.079858944 |
| Q6PE15 | Abhd10 | -2.347420341 | 3.732655182 |
| Q80U35 | Arhgef17 | -2.348026309 | 2.026507185 |
| Q922E6 | Fastkd2 | -2.349642307 | 2.438353755 |
| Q8BGR8 | Gskip | -2.353528538 | 3.29616882 |
| P11276 | Fn1 | -2.353826876 | 4.064178869 |
| Q3U4G3 | Xxylt1 | -2.359939401 | 2.581588194 |
| Q9JIA7 | Sphk2 | -2.360545024 | 2.292202421 |
| P54818 | Galc | -2.362830764 | 3.566779763 |
| P17095 | Hmga1 | -2.362946864 | 3.918871276 |
| P47754 | Capza2 | -2.373440446 | 6.133868173 |
| Q9ESW8 | Pgpep1 | -2.37544728 | 2.060577932 |
| Q9WTR5 | Cdh13 | -2.375588755 | 2.684026805 |
| Q0VGY8 | Tanc1 | -2.376227302 | 2.478109684 |
| Q9D8Z1 | Ascc1 | -2.378680332 | 2.049997399 |
| Q60949 | Tbc1d1 | -2.379994253 | 3.045940051 |
| P50543 | S100a11 | -2.380548164 | 4.847082718 |
| E9QAM5 | Helz2 | -2.383382752 | 3.468897239 |
| P52633 | Stat6 | -2.383488192 | 5.507289836 |
| Q07243 | Mtf1 | -2.384915965 | 2.104540302 |
| Q8K298 | Anln | -2.388691611 | 6.049262358 |
| Q9CR16 | Ppid | -2.391908246 | 6.116924515 |
| Q8C181 | Mbnl2 | -2.39404873 | 3.826167348 |
| P49138 | Mapkapk2 | -2.396989163 | 4.041025008 |
| Q8R205 | Zc3h10 | -2.401106901 | 2.996769446 |
| Q3TC33 | Ccdc127 | -2.405648607 | 2.272693692 |
| Q8R5C5 | Actr1b | -2.409297784 | 5.332618856 |
| A2ACJ2 | Faap100 | -2.411174007 | 3.675343822 |
| A2AI08 | Tprn | -2.411313996 | 2.742656824 |
| P97360 | Etv6 | -2.411925302 | 2.568857701 |
| Q8BH97 | Rcn3 | -2.412012833 | 3.603622505 |
| Q99K82 | Smox | -2.413136369 | 2.760578798 |
| Q8VC70 | Rbms2 | -2.413613156 | 3.865078208 |
| Q8K2J7 | Rell1 | -2.416297612 | 2.605047261 |
| P31750 | Akt1 | -2.430332002 | 4.746578186 |
| Q9CPY3 | Cdca5 | -2.431670431 | 2.129592365 |
| Q9CXH7 | Sgo1 | -2.4325677 | 2.508277903 |
| P17751 | Tpi1 | -2.433762873 | 5.598055492 |
| Q9D142 | Nudt14 | -2.435002423 | 2.607843713 |
| Q8VCS6 | Med9 | -2.436731494 | 4.832327748 |
| Q68FH4 | Galk2 | -2.437528417 | 2.007661813 |
| P58771 | Tpm1 | -2.43833775 | 5.046282868 |
| Q9CY50 | Ssr1 | -2.440849255 | 2.205897007 |
| Q9QUG2 | Polk | -2.441350182 | 3.002595911 |
| Q8K1C9 | Lrrc41 | -2.444590084 | 2.528293438 |
| P59222 | Scarf2 | -2.45238623 | 2.959935504 |
| Q8R0F3 | Sumf1 | -2.456321141 | 3.866001467 |
| Q8CCH7 | Zfpm2 | -2.458104968 | 3.386517214 |
| Q8BIA4 | Fbxw8 | -2.463582605 | 3.064046074 |
| Q99KW9 | Itfg1 | -2.466275664 | 3.259194958 |
| Q9CR75 | Tnfrsf12a | -2.466926358 | 2.186944492 |
| Q9WV32 | Arpc1b | -2.46767657 | 6.446737178 |
| P10518 | Alad | -2.468253514 | 5.162717733 |
| Q9CQA6 | Chchd1 | -2.469236363 | 2.939365786 |
| Q9WVJ9 | Efemp2 | -2.471003332 | 2.304948085 |
| E9Q784 | Zc3h13 | -2.476574756 | 2.511776441 |
| Q9CXC3 | Mgme1 | -2.485553225 | 2.578630348 |
| P70227 | Itpr3 | -2.486009263 | 2.25526171 |
| Q80W68 | Kirrel1 | -2.486865896 | 3.428442899 |
| O88986 | Gcat | -2.489385141 | 5.305288851 |
| Q922Q1 | Mtarc2 | -2.492870867 | 2.878955336 |
| P70255 | Nfic | -2.494586551 | 4.175600195 |
| Q61127 | Nab2 | -2.496213952 | 4.111456123 |
| P39053 | Dnm1 | -2.496593768 | 5.865710386 |
| Q922F4 | Tubb6 | -2.49701434 | 5.124297528 |
| Q9WUB0 | Rbck1 | -2.499336112 | 2.138011879 |
| Q64437 | Adh7 | -2.501528817 | 2.348572714 |
| A2ADA5 | Pusl1 | -2.503344859 | 2.097794545 |
| Q91ZS8 | Adarb1 | -2.505726171 | 3.53839266 |
| O35375 | **Nrp2** | -2.513769451 | 2.276972505 |
| Q9DD24 | Tceal9 | -2.51462436 | 3.499355994 |
| P11688 | Itga5 | -2.517988935 | 2.607843713 |
| Q01705 | Notch1 | -2.52286942 | 2.385758222 |
| Q9CZS3 | Cep20 | -2.527236195 | 2.304436226 |
| Q8C804 | Spice1 | -2.527914884 | 3.143039076 |
| Q64521 | Gpd2 | -2.529179918 | 5.258579507 |
| Q80WB5 | Ntaq1 | -2.531347698 | 2.441058526 |
| P07214 | Sparc | -2.531494709 | 5.298661973 |
| Q07417 | Acads | -2.535712848 | 5.699939096 |
| Q8BFW7 | Lpp | -2.53854845 | 5.653371399 |
| Q8R0S2 | Iqsec1 | -2.544394382 | 2.988935512 |
| Q3TGF2 | Fam107b | -2.549846723 | 5.595097028 |
| Q9Z1B7 | Mapk13 | -2.55355764 | 2.979655261 |
| Q9QY53 | Nphp1 | -2.556635393 | 3.141431077 |
| Q99LJ6 | Gpx7 | -2.558742952 | 2.088197728 |
| Q9WVL3 | Slc12a7 | -2.560184087 | 3.281969313 |
| Q99LB7 | Sardh | -2.561564365 | 2.431781762 |
| Q9CPY0 | Mrm2 | -2.562243264 | 2.104477093 |
| Q9CWY4 | Gemin7 | -2.563336405 | 2.184946133 |
| Q3UQ28 | Pxdn | -2.563477288 | 2.411307837 |
| Q61738 | Itga7 | -2.564092627 | 3.248555979 |
| Q9JL19 | Ncoa6 | -2.565712711 | 4.556503766 |
| Q8BZT9 | Lacc1 | -2.566720825 | 4.302626776 |
| Q9ERS5 | Plekha2 | -2.572744971 | 2.356897691 |
| P25322 | Ccnd1 | -2.578336377 | 2.551114425 |
| O54967 | Tnk2 | -2.587997204 | 2.436426571 |
| Q9WVK4 | Ehd1 | -2.589072147 | 6.074962845 |
| Q9CPV1 | Ska1 | -2.589205145 | 2.727626245 |
| P58059 | Mrps21 | -2.590417207 | 3.694254572 |
| Q9CY21 | Bud23 | -2.591996463 | 2.370280448 |
| O54791 | Maff | -2.592521404 | 2.79082311 |
| Q8BRM2 | Gorab | -2.593565751 | 2.578780342 |
| P09055 | Itgb1 | -2.593611983 | 4.102305266 |
| P53690 | Mmp14 | -2.595999267 | 2.341230564 |
| Q6IRU2 | Tpm4 | -2.597019118 | 4.75579062 |
| Q8C9X6 | Epc1 | -2.599568724 | 2.328159798 |
| Q8BMB0 | Emsy | -2.603887651 | 2.146422706 |
| P41731 | Cd63 | -2.604456401 | 2.231501868 |
| P26041 | Msn | -2.607365472 | 5.937472075 |
| Q640N1 | Aebp1 | -2.609049258 | 2.048798538 |
| Q9WTI7 | Myo1c | -2.61293621 | 4.939704172 |
| Q99K41 | Emilin1 | -2.62431809 | 2.178039988 |
| Q8K370 | Acad10 | -2.624512505 | 2.06837047 |
| Q9D0I6 | Wdsub1 | -2.627249684 | 2.074483872 |
| P22366 | Myd88 | -2.628286133 | 4.053629942 |
| Q7TPV4 | Mybbp1a | -2.635889141 | 4.746578186 |
| Q9CWT6 | Ddx28 | -2.639117654 | 3.072112706 |
| O89086 | Rbm3 | -2.650209598 | 5.058244064 |
| Q9D2R0 | Aacs | -2.659861289 | 5.229648104 |
| P27577 | Ets1 | -2.661247803 | 2.761951863 |
| Q8BYM8 | Cars2 | -2.676377471 | 2.976891719 |
| Q9DBR4 | Apbb2 | -2.677746183 | 4.09964401 |
| P62500 | Tsc22d1 | -2.67917774 | 2.259830195 |
| Q66GT5 | Ptpmt1 | -2.681089643 | 2.763653523 |
| Q9CQ91 | Ndufa3 | -2.685062908 | 2.378186823 |
| Q91YH5 | Atl3 | -2.685515688 | 2.20039387 |
| Q923K4 | Gtpbp3 | -2.687978926 | 2.389833093 |
| Q8CJ40 | Crocc | -2.694169787 | 2.614034464 |
| Q8K004 | Spata2 | -2.6962747 | 3.173963292 |
| Q3U1G5 | Isg20l2 | -2.697837643 | 2.259594926 |
| Q8K2B0 | P3h4 | -2.700589428 | 2.083043383 |
| P15806 | Tcf3 | -2.701024756 | 4.024185819 |
| P47738 | Aldh2 | -2.702506908 | 5.248871757 |
| Q8BP40 | Acp6 | -2.704253413 | 3.468897239 |
| Q6P5E8 | Dgkq | -2.704862519 | 2.18181174 |
| Q8K214 | Scmh1 | -2.709543395 | 3.152048374 |
| Q921E2 | Rab31 | -2.71144135 | 2.038400384 |
| O70325 | Gpx4 | -2.71307433 | 4.859020075 |
| Q8R5A3 | Apbb1ip | -2.713900985 | 2.217547278 |
| Q9CRW3 |  | -2.714335294 | 2.202293884 |
| P0C8B4 | Gon7 | -2.718232228 | 4.541171323 |
| Q99J08 | Sec14l2 | -2.719672452 | 3.07058594 |
| Q08093 | Cnn2 | -2.723414526 | 5.595097028 |
| O35126 | Atn1 | -2.735324959 | 2.780257452 |
| Q9CZB0 | Sdhc | -2.736019484 | 2.077691886 |
| Q3UIU2 | Ndufb6 | -2.736297668 | 2.382933932 |
| Q8VHT7 | Gtf3a | -2.737111153 | 2.334333379 |
| Q8R5A0 | Smyd2 | -2.744648799 | 3.240257247 |
| Q99MK8 | Grk2 | -2.745932986 | 3.786222622 |
| Q924Z4 | Cers2 | -2.747363364 | 2.944483168 |
| Q62266 | Sprr1a | -2.748292901 | 4.082764858 |
| P09450 | Junb | -2.750721843 | 2.090632987 |
| Q8JZS6 | N4bp2l2 | -2.752002421 | 2.329380965 |
| Q8BR90 | Rimoc1 | -2.753440109 | 3.596922766 |
| Q8BZB2 | Ppcdc | -2.758056149 | 2.461744439 |
| Q8C163 | Exog | -2.762944541 | 2.707735765 |
| Q9CQQ0 | Smim8 | -2.76744268 | 2.406729122 |
| Q9DBM1 | Gpatch1 | -2.768659992 | 2.836584457 |
| Q3U2K0 | Fam193b | -2.784999877 | 3.487278267 |
| P43135 | Nr2f2 | -2.789319857 | 2.982906239 |
| Q8BKR5 | Ppp1r37 | -2.792229196 | 2.26789247 |
| Q8R086 | Suox | -2.794944301 | 3.022102985 |
| P56375 | Acyp2 | -2.803768915 | 3.720647741 |
| Q9Z0P4 | Palm | -2.807304673 | 3.121230455 |
| P18052 | Ptpra | -2.80981543 | 2.878510747 |
| Q6NXW6 | Rad17 | -2.810623098 | 2.988371141 |
| P49817 | Cav1 | -2.811033441 | 2.592694811 |
| O08734 | Bak1 | -2.813738672 | 3.18346552 |
| Q7TSH3 | Znf516 | -2.814170236 | 2.026322963 |
| Q5SSK3 | Tefm | -2.815006183 | 3.591424562 |
| Q6F3F9 | Adgrg6 | -2.815340285 | 2.451854664 |
| P28033 | Cebpb | -2.815699798 | 2.892234104 |
| Q3U829 | Ap5z1 | -2.816095892 | 2.775681387 |
| Q571F8 | Gls2 | -2.817588805 | 2.888554909 |
| P50429 | Arsb | -2.817649516 | 2.727043859 |
| P35569 | Irs1 | -2.827119249 | 2.901017824 |
| Q9D009 | Lipt2 | -2.83455689 | 2.471597184 |
| Q8C3X8 | Lmf2 | -2.836129645 | 2.425297192 |
| O70252 | Hmox2 | -2.840863131 | 2.427725614 |
| Q8CGB6 | Tns2 | -2.844816878 | 2.176716027 |
| O88939 | Zbtb7a | -2.848579129 | 5.258686477 |
| Q924C6 | Loxl4 | -2.857924209 | 2.464078451 |
| Q99M15 | Pstpip2 | -2.858682861 | 3.40253602 |
| Q99JR5 | Tinagl1 | -2.860313689 | 3.192753146 |
| P97821 | Ctsc | -2.864792734 | 2.713154663 |
| O08539 | Bin1 | -2.865447379 | 6.770926025 |
| Q9QYF1 | Rdh11 | -2.867144091 | 2.946037477 |
| P70451 | Fer | -2.868366092 | 2.154248447 |
| Q80TA9 | Epg5 | -2.869808183 | 2.227418511 |
| Q99N20 | Brms1 | -2.870986636 | 2.061807783 |
| Q8CFI5 | Pars2 | -2.872401219 | 2.279961754 |
| Q8R080 | Gtse1 | -2.873589971 | 4.055275362 |
| Q5DTX6 | Jcad | -2.87878228 | 4.166065651 |
| Q75NR7 | Recql4 | -2.879155251 | 2.346810595 |
| P47740 | Aldh3a2 | -2.887137148 | 3.178972106 |
| Q8K371 | Amotl2 | -2.889523881 | 2.418932866 |
| Q8C0Z1 | Fam234a | -2.8946953 | 3.344965728 |
| P54761 | Ephb4 | -2.902794393 | 2.326059743 |
| Q91ZX7 | Lrp1 | -2.905013269 | 4.053024637 |
| Q9CZG3 | Commd8 | -2.911097754 | 4.11103279 |
| Q8BH04 | Pck2 | -2.913001206 | 4.099880595 |
| Q8R2Y0 | Abhd6 | -2.914183594 | 2.030804546 |
| Q8R0F5 | Rbmx2 | -2.920323039 | 3.872540851 |
| P09925 | Surf1 | -2.924234422 | 3.496475607 |
| Q9CRY7 | Gdpd1 | -2.925654461 | 2.13208685 |
| Q8BJM3 | R3hcc1l | -2.926897776 | 2.189351193 |
| Q7TNE3 | Spag7 | -2.927168411 | 2.894937113 |
| Q920A7 | Afg3l1 | -2.927273883 | 4.237734352 |
| Q9QWR8 | Naga | -2.92826223 | 3.979103932 |
| Q8JZU0 | Nudt13 | -2.928823708 | 2.066681278 |
| A2A7S8 | Nhsl3 | -2.929785812 | 3.349552547 |
| O88196 | Ttc3 | -2.930903547 | 3.020206524 |
| Q99JZ7 | Errfi1 | -2.936515283 | 3.333044656 |
| Q9WV76 | Ap4b1 | -2.9386149 | 2.255832439 |
| Q8K1K4 | Cenpi | -2.94321164 | 3.191884281 |
| A2A6T1 | Cdr2l | -2.943895479 | 3.015810841 |
| Q8BHE0 | Prr11 | -2.944729227 | 2.019100837 |
| Q8BQ47 | Cnpy4 | -2.950243678 | 4.086829476 |
| Q91VT4 | Cbr4 | -2.954548275 | 2.348942663 |
| Q8VC57 | Kctd5 | -2.959526407 | 3.188140581 |
| Q78IK4 | Apool | -2.959796127 | 3.305862891 |
| Q8VHY0 | Cspg4 | -2.980811264 | 2.652539418 |
| Q3UW53 | Niban1 | -2.983477395 | 2.82834688 |
| O35954 | Pitpnm1 | -2.984606333 | 2.4644355 |
| Q6PDX6 | Rnf220 | -2.988460808 | 4.031828661 |
| Q9JHJ3 | Glmp | -2.993283722 | 2.069517557 |
| Q3URQ7 | Mthfsd | -2.994359333 | 3.141566361 |
| Q7TSJ2 | Map6 | -2.998209873 | 2.334333379 |
| Q8K2J0 | Plcd3 | -2.998909207 | 2.099097227 |
| Q9CZT5 | Vasn | -3.001856645 | 2.542179822 |
| Q3UYI5 | Rgl3 | -3.011446959 | 3.43875609 |
| P58468 | Slx9 | -3.019442184 | 4.464720182 |
| Q9ET22 | Dpp7 | -3.024118712 | 4.910095805 |
| Q9QXS1 | Plec | -3.024369017 | 6.12004253 |
| Q499E4 | Dzip1l | -3.026745755 | 2.6248095 |
| Q99L00 | Haus8 | -3.027263398 | 3.003092714 |
| Q921Y4 | Mfsd5 | -3.02970975 | 4.98716896 |
| Q9D937 |  | -3.035394502 | 3.749840769 |
| Q91YI0 | Asl | -3.040267848 | 5.485443412 |
| Q9D9M5 | Phospho2 | -3.041329303 | 2.492795681 |
| P49935 | Ctsh | -3.043348881 | 2.707377792 |
| P48410 | Abcd1 | -3.051550704 | 4.459394778 |
| Q810S1 | Mcub | -3.057077233 | 2.463598998 |
| Q3TBT3 | Sting1 | -3.057736595 | 2.381722552 |
| A2AI05 | Ndor1 | -3.060903568 | 2.306649594 |
| Q8BMA5 | Npat | -3.061111529 | 2.560872422 |
| Q8K215 | Lyrm4 | -3.069530332 | 2.208178881 |
| P56542 | Dnase2 | -3.070188164 | 3.001862973 |
| Q8R550 | Sh3kbp1 | -3.073943936 | 4.738660704 |
| Q99MR1 | Gigyf1 | -3.076839421 | 2.918650745 |
| A0A7N9VSG0 | Afg2b | -3.076957659 | 2.466428773 |
| Q64337 | Sqstm1 | -3.076989132 | 6.12004253 |
| B7ZNG4 | Troap | -3.082430611 | 2.695327351 |
| Q9D8S3 | Arfgap3 | -3.084348057 | 2.176181428 |
| Q60695 | Rgl1 | -3.084655442 | 2.229317119 |
| P02340 | Tp53 | -3.085654279 | 3.597182542 |
| Q9CY62 | Rnf181 | -3.091227082 | 3.647032308 |
| P08074 | Cbr2 | -3.093011276 | 2.390446986 |
| Q9CR59 | Gadd45gip1 | -3.097828124 | 2.656274925 |
| P70671 | Irf3 | -3.103207131 | 3.365771643 |
| Q9JK92 | Hspb8 | -3.104394608 | 3.00273901 |
| Q8K1J5 | Sde2 | -3.104398322 | 3.299645564 |
| Q5NC05 | Ttf2 | -3.112371483 | 3.704371678 |
| Q8BZ20 | Parp12 | -3.114733844 | 2.101176546 |
| P22935 | Crabp2 | -3.117131265 | 5.214901068 |
| Q60I26 | Als2cl | -3.11937228 | 2.905001229 |
| Q8CHQ0 | Fbxo4 | -3.123528326 | 3.379827241 |
| Q61292 | Lamb2 | -3.125461745 | 3.821881907 |
| Q8R2Q8 | Bst2 | -3.128335212 | 2.545218711 |
| Q3V1H1 | Ckap2 | -3.129886532 | 3.521310689 |
| Q9D3P8 | Plgrkt | -3.131137838 | 4.566740796 |
| Q9CRC3 |  | -3.142732525 | 2.282632136 |
| Q91WC9 | Daglb | -3.149577007 | 2.136784799 |
| A2AUY4 | Baz2b | -3.150681438 | 3.8298213 |
| Q99J79 | Ddb2 | -3.159810111 | 3.191358675 |
| Q3TW96 | Uap1l1 | -3.164604103 | 4.687420751 |
| Q8R3K3 | Ptcd2 | -3.164827464 | 3.33931556 |
| Q8K039 |  | -3.164932601 | 3.294153079 |
| Q8BIW1 | Prune1 | -3.165492946 | 2.732922898 |
| Q80TS3 | Adgrl3 | -3.169979178 | 2.119784657 |
| Q91W61 | Fbxl15 | -3.170381567 | 3.438821128 |
| Q91WS0 | Cisd1 | -3.175541448 | 2.411359418 |
| Q9WUQ2 | Preb | -3.177084448 | 2.697035914 |
| Q6PFD6 | Kif18b | -3.177778601 | 2.436583847 |
| Q6A058 | Armcx2 | -3.181272849 | 2.514039263 |
| Q8C854 | Myef2 | -3.1815224 | 3.622128636 |
| Q8K387 | Usp45 | -3.191787401 | 2.482956727 |
| Q8VCM4 | Lipt1 | -3.196866465 | 2.761951863 |
| Q9JHE7 | Tssc4 | -3.205007237 | 3.271656488 |
| Q3UHQ6 | Dop1b | -3.207916772 | 2.201863603 |
| Q8R2Y2 | Mcam | -3.210160011 | 3.000126733 |
| Q8VEH5 | Epm2aip1 | -3.210843789 | 2.775681387 |
| Q9CX53 | Gemin6 | -3.211552952 | 2.343903604 |
| Q9D2G5 | Synj2 | -3.214088457 | 2.139607888 |
| O88207 | Col5a1 | -3.215471581 | 2.231501868 |
| Q9CR70 | Lage3 | -3.219748075 | 4.763897782 |
| Q8VCP8 | Ak6 | -3.223053844 | 2.570798114 |
| Q9CQY2 | Ramac | -3.229997136 | 3.958627787 |
| Q8C5P5 | Nt5dc1 | -3.23223021 | 5.551659592 |
| P40630 | Tfam | -3.23294563 | 2.503144935 |
| Q8R344 | Ccdc12 | -3.240958818 | 3.240257247 |
| Q8VI33 | Taf9 | -3.247027041 | 2.151951934 |
| E9Q9R9 | Dlg5 | -3.253599003 | 3.580777672 |
| Q9CQJ7 | Pttg1 | -3.258088153 | 3.030000236 |
| Q8BLY7 | Hps6 | -3.264856119 | 3.60108972 |
| Q8CBY0 | Gatc | -3.267161332 | 2.603110878 |
| Q9WVL0 | Gstz1 | -3.268365467 | 2.418932866 |
| Q9CQZ6 | Ndufb3 | -3.271415854 | 2.599089035 |
| Q8BTI9 | Pik3cb | -3.271927759 | 2.040237574 |
| Q9Z0Z3 | Skp2 | -3.277625704 | 2.361065623 |
| Q9D281 | Fam114a1 | -3.280254993 | 2.89842208 |
| Q9D8T2 | Gsdmd | -3.285398305 | 2.001641421 |
| P48678 | Lmna | -3.287397364 | 6.457680354 |
| P10605 | Ctsb | -3.288391538 | 3.961950585 |
| O88824 | Jtb | -3.290465407 | 2.609993787 |
| Q8R4E9 | Cdt1 | -3.290898668 | 3.248559989 |
| Q60872 | Eif1a | -3.296548795 | 3.250722062 |
| P59110 | Senp1 | -3.297161133 | 2.986085399 |
| Q9CQJ8 | Ndufb9 | -3.297783019 | 3.100342019 |
| P29533 | Vcam1 | -3.299090472 | 2.12248305 |
| P85094 | Isoc2a | -3.306208822 | 3.729899237 |
| D3YZG8 | Mthfd2l | -3.312672246 | 3.063354165 |
| Q8VCW8 | Acsf2 | -3.321872144 | 3.333044656 |
| Q62086 | Pon2 | -3.323515323 | 2.660965388 |
| O89023 | Tpp1 | -3.333123161 | 2.176081228 |
| Q9DC71 | Mrps15 | -3.336841607 | 4.182352601 |
| Q91XC0 | Ajuba | -3.337648421 | 3.033950056 |
| Q9CX48 | Zcchc10 | -3.338708052 | 2.142390788 |
| Q62172 | Ralbp1 | -3.341993596 | 2.329405984 |
| Q9CPT4 | Mydgf | -3.34332322 | 3.6695606 |
| Q8BHW2 | Oscp1 | -3.343816856 | 2.020179245 |
| Q3TVC7 | Ccndbp1 | -3.352566537 | 2.514277145 |
| Q9CX66 | Nopchap1 | -3.358036406 | 2.67459499 |
| P97864 | Casp7 | -3.371696603 | 2.511197774 |
| Q8BGD8 | Coa6 | -3.372626902 | 4.164251715 |
| Q9CWY3 | Setd6 | -3.373779614 | 2.144782717 |
| Q8BYH7 | Tbc1d17 | -3.373805057 | 2.549077182 |
| Q9CR25 | Dph2 | -3.377993511 | 4.64119677 |
| Q80YR6 | Rbbp8 | -3.382431979 | 2.007375937 |
| Q9WTK5 | Nfkb2 | -3.388153239 | 5.229648104 |
| P98063 | Bmp1 | -3.388180344 | 2.141733483 |
| Q8C0J6 | Sowahc | -3.394751681 | 2.67459499 |
| P97783 | Mllt11 | -3.402795817 | 2.437323488 |
| Q3U962 | Col5a2 | -3.404211652 | 2.465043306 |
| P46656 | Fdx1 | -3.419348234 | 2.996769446 |
| Q9DCS2 | Mettl26 | -3.42126131 | 4.552874983 |
| Q99PL6 | Ubxn6 | -3.425405059 | 3.230227534 |
| Q9DCA2 | Mrps11 | -3.434401546 | 2.888655138 |
| P28653 | Bgn | -3.435285106 | 3.772641943 |
| Q64701 | Rbl1 | -3.436183763 | 2.091314165 |
| Q9QYR9 | Acot2 | -3.437030944 | 3.973149129 |
| Q80Z25 | Ofd1 | -3.444393327 | 4.019367725 |
| Q61602 | Gli3 | -3.445152957 | 2.091219913 |
| Q9CQC7 | Ndufb4 | -3.445731774 | 2.652516772 |
| Q9QZM4 | Tnfrsf10b | -3.449398439 | 2.741694642 |
| Q9CRA4 | Msmo1 | -3.449910103 | 3.304141289 |
| O35405 | Pld3 | -3.451391946 | 3.674948154 |
| Q9CXX9 | Cuedc2 | -3.455688388 | 2.004831179 |
| Q8C3S2 | Tango6 | -3.466725511 | 2.081655548 |
| Q99LJ7 | Rcbtb2 | -3.467668557 | 2.029109882 |
| Q9Z2H5 | Epb41l1 | -3.468324192 | 2.966477288 |
| E9Q0S6 | Tns1 | -3.471886766 | 2.534705491 |
| Q9CWX2 | Ndufaf1 | -3.47268118 | 4.943475205 |
| P0DOV1 | Ifi211 | -3.473650554 | 2.60170874 |
| P13020 | Gsn | -3.475079473 | 5.68757106 |
| Q9D773 | Mrpl2 | -3.485509527 | 2.803566205 |
| Q80XL6 | Acad11 | -3.495358376 | 2.987993512 |
| P63013 | Prrx1 | -3.495739439 | 2.820026914 |
| Q63918 | Cavin2 | -3.509415812 | 5.767990337 |
| Q9D8C4 | Ifi35 | -3.509685302 | 2.519190393 |
| Q9EPR5 | Sorcs2 | -3.521067392 | 2.122843156 |
| P30412 | Ppic | -3.521538079 | 2.084550583 |
| Q5SUQ9 | Ctc1 | -3.523635038 | 4.695250907 |
| Q8R3P0 | Aspa | -3.529261232 | 2.069508952 |
| Q9WV91 | Ptgfrn | -3.542989107 | 2.82739836 |
| Q8CIK8 | Rfwd3 | -3.544559163 | 2.944296456 |
| Q3UHA3 | Spg11 | -3.54986627 | 2.195826816 |
| Q9D853 | Eef1akmt2 | -3.550071904 | 2.301081758 |
| Q6NXJ0 | Wwc2 | -3.551580464 | 3.107847103 |
| P35288 | Rab23 | -3.555135449 | 2.911283063 |
| Q8CBY1 | Samd4a | -3.55547716 | 3.376284446 |
| Q8BGV4 | Tti2 | -3.560995261 | 2.584271195 |
| Q8C6I2 | Sdhaf2 | -3.565071387 | 2.219549315 |
| Q14AI6 | Rpusd3 | -3.565580663 | 3.162682885 |
| P97302 | Bach1 | -3.565603911 | 2.03689361 |
| Q9D842 | Aplf | -3.570613044 | 3.69198754 |
| Q99PG2 | Ogfr | -3.583457278 | 4.91904186 |
| Q8C551 | Rad51ap1 | -3.584444369 | 2.366984678 |
| Q8CC35 | Synpo | -3.584987342 | 3.62298569 |
| Q80UU2 | Rpp38 | -3.588931354 | 3.468897239 |
| O35988 | Sdc4 | -3.601411937 | 2.497849079 |
| Q8C2B3 | Hdac7 | -3.612755386 | 2.755259233 |
| Q71FD7 | Fblim1 | -3.617746546 | 2.080199552 |
| Q9CWP6 | Mospd2 | -3.62192072 | 3.604572758 |
| Q78HU3 | Mvb12a | -3.622723192 | 3.572487984 |
| Q9DCZ1 | Gmpr | -3.623337291 | 2.758694485 |
| Q8VC34 | Rpap2 | -3.626542325 | 3.423507934 |
| Q7TT23 | Dnaaf9 | -3.627495924 | 3.443867263 |
| O55101 | Syngr2 | -3.63811344 | 3.499355994 |
| Q9CZ09 | Mettl18 | -3.642885471 | 3.133586075 |
| P70271 | Pdlim4 | -3.643888283 | 5.079061664 |
| Q8R4N0 | Clybl | -3.645080807 | 2.574994995 |
| Q9CR02 | Tma16 | -3.646595319 | 2.525909383 |
| Q8K4R9 | Dlgap5 | -3.652468144 | 3.332852224 |
| Q8CE46 | Pus7l | -3.655753717 | 2.379070025 |
| Q6PGG2 | Gmip | -3.657627564 | 2.571700694 |
| Q6NSR8 | Npepl1 | -3.658883435 | 2.128622911 |
| P52624 | Upp1 | -3.663598604 | 2.86266947 |
| Q8BXA1 | Golim4 | -3.668378393 | 2.505492799 |
| P28650 | Adss1 | -3.679901745 | 2.518268986 |
| P59017 | Bcl2l13 | -3.692630986 | 5.712943736 |
| Q8BL80 | Arhgap22 | -3.692770582 | 4.0731418 |
| Q8CIC2 | Nup42 | -3.695454073 | 2.356897691 |
| Q8C1M2 | Znf428 | -3.701615288 | 5.771121261 |
| Q9CZX7 | Pip4p2 | -3.70185814 | 2.299286703 |
| Q9D727 | Pex39 | -3.706697653 | 2.538667718 |
| Q7TQ95 | Lnpk | -3.710013142 | 3.667683569 |
| Q8K203 | Neil3 | -3.71075244 | 2.614376375 |
| Q9DCT8 | Crip2 | -3.714525002 | 3.499973753 |
| Q91UZ5 | Impa2 | -3.716579578 | 2.246056034 |
| Q8VE38 | Oxnad1 | -3.7235109 | 2.661560405 |
| P47955 | Rplp1 | -3.723543842 | 5.005523934 |
| Q9D6Y7 | Msra | -3.725563444 | 3.211195322 |
| P81269 | Atf1 | -3.731915313 | 3.674948154 |
| Q9ESY9 | Ifi30 | -3.73383097 | 3.291165914 |
| O70161 | Pip5k1c | -3.737400148 | 2.85956943 |
| Q9CQZ5 | Ndufa6 | -3.740969296 | 3.297777922 |
| Q9EPB5 | Serhl | -3.743073114 | 3.93563975 |
| Q8BHA0 | Ino80c | -3.743206487 | 4.029912263 |
| P62748 | Hpcal1 | -3.744475591 | 2.342593891 |
| O88512 | Ap1g2 | -3.746823783 | 2.259409243 |
| Q924T3 | Xrcc4 | -3.750545367 | 4.577390206 |
| Q9D845 | Tex9 | -3.754237851 | 2.576975813 |
| Q9D2L9 | Fam111a | -3.754466026 | 2.283605437 |
| Q921Q7 | Rin1 | -3.75564467 | 2.462693616 |
| Q7TMC8 | Fcsk | -3.765247347 | 2.090632987 |
| Q9Z0H8 | Clip2 | -3.766181488 | 4.228189692 |
| Q8BMI4 | Gen1 | -3.766375135 | 3.569576811 |
| Q6ZPG2 | Wdr90 | -3.766716026 | 2.146901718 |
| P47930 | Fosl2 | -3.768521515 | 2.807631898 |
| P56213 | Gfer | -3.77492282 | 3.559427113 |
| Q8CES0 | Naa30 | -3.776303567 | 2.131229641 |
| Q9JJ89 | Ccdc86 | -3.778226528 | 2.198635668 |
| P20060 | Hexb | -3.781867857 | 2.868871943 |
| Q61037 | Tsc2 | -3.783648242 | 4.073271516 |
| Q9CZX5 | Pinx1 | -3.785367564 | 3.063354165 |
| Q9DCI9 | Mrpl32 | -3.788341766 | 3.295071978 |
| P70677 | Casp3 | -3.791118536 | 6.770926025 |
| Q64669 | Nqo1 | -3.799632672 | 4.166065651 |
| Q8BW00 | Ptrh1 | -3.802065077 | 5.640144956 |
| P02802 | Mt1 | -3.806839884 | 3.560910326 |
| Q9Z223 | Mocs2 | -3.807962039 | 2.825486206 |
| P70213 | Fv1 | -3.808678576 | 2.187814133 |
| Q8K4E0 | Alms1 | -3.816072295 | 2.814098851 |
| Q3UX61 | Naa11 | -3.816878061 | 2.027505716 |
| Q8BK75 | Elp6 | -3.819829345 | 2.348715313 |
| Q9Z0M5 | Lipa | -3.82616305 | 3.167224891 |
| Q3TV70 | Nr2c2ap | -3.827912492 | 2.058700041 |
| Q8C796 | Rcor2 | -3.828267728 | 3.98644725 |
| Q9DBE8 | Alg2 | -3.82902444 | 2.942528866 |
| Q99JV5 | Stard4 | -3.830403408 | 2.770790178 |
| Q9Z2Q2 | Knop1 | -3.831580297 | 3.165716342 |
| Q8BZA9 | Tigar | -3.832264719 | 3.825030232 |
| Q6AW69 | Cgnl1 | -3.834794703 | 2.672209836 |
| Q9D6H2 | Ift25 | -3.847132997 | 2.312706093 |
| Q9CWH5 | Trmt11 | -3.848347127 | 3.753310744 |
| O08908 | Pik3r2 | -3.862262123 | 4.746578186 |
| Q6PG16 | Hjurp | -3.870780685 | 2.089665856 |
| Q9DBE0 | Csad | -3.877489622 | 3.494296573 |
| P70193 | Lrig1 | -3.877814905 | 2.011579108 |
| Q8R3F5 | Mcat | -3.87863737 | 2.368472841 |
| P12265 | Gusb | -3.887695927 | 2.439886776 |
| Q8CG19 | Ltbp1 | -3.901664411 | 3.295499037 |
| P68368 | Tuba4a | -3.903337293 | 5.507289836 |
| Q91ZF0 | Dnajc24 | -3.903535452 | 3.457574441 |
| Q3U6N9 |  | -3.912939686 | 2.033649346 |
| Q8K1I7 | Wipf1 | -3.918475829 | 2.883423676 |
| Q8CIV8 | Tbce | -3.919978428 | 4.746578186 |
| Q80V94 | Ap4e1 | -3.920387645 | 2.872894663 |
| Q9QX60 | Dguok | -3.921045111 | 3.422186973 |
| Q9CPW3 | Mrpl54 | -3.922329583 | 3.485936022 |
| Q3U7U3 | Fbxo7 | -3.928784154 | 3.027215755 |
| Q9CQP3 | Chchd5 | -3.929458472 | 3.045680823 |
| Q3UGP9 | Lrrc58 | -3.943218158 | 2.710080888 |
| P46935 | Nedd4 | -3.944649044 | 6.01085686 |
| Q6PAL7 | Ahdc1 | -3.949684498 | 2.898487227 |
| Q8VHK1 | Caskin2 | -3.949828138 | 2.025356619 |
| P48432 | Sox2 | -3.955236083 | 2.933424401 |
| Q61263 | Soat1 | -3.96335481 | 2.727626245 |
| Q00493 | Cpe | -3.964346407 | 5.364449318 |
| Q8BK30 | Ndufv3 | -3.966356664 | 2.766284881 |
| Q8VC19 | Alas1 | -3.971113663 | 2.399118646 |
| P29351 | Ptpn6 | -3.974841165 | 2.294534854 |
| Q9CX60 | Lbh | -3.975849026 | 2.337156543 |
| Q8R3F9 | Tut1 | -3.978695485 | 2.868132631 |
| A6PWY4 | Wdr76 | -3.984333327 | 2.946333073 |
| P0DJF2 | Pet117 | -3.986779559 | 2.881954638 |
| P58058 | Nadk | -3.987194946 | 3.926334063 |
| P97352 | S100a13 | -3.989304279 | 5.32435078 |
| Q8C263 | Ska3 | -3.993814969 | 2.92363923 |
| P35762 | Cd81 | -3.994010824 | 2.049997399 |
| Q9CQB2 | Mcrip2 | -3.996745693 | 2.625431125 |
| Q9CQU5 | Zwint | -4.005620689 | 3.946362164 |
| Q64364 | Cdkn2a | -4.010662164 | 3.398939489 |
| Q99M04 | Lias | -4.015517983 | 2.871967257 |
| P19783 | Cox4i1 | -4.019838464 | 4.428869132 |
| Q9CW79 | Golga1 | -4.022361354 | 2.036548192 |
| Q99MZ7 | Pecr | -4.02434107 | 4.746578186 |
| Q8C761 | Dync2i1 | -4.037868267 | 4.117235238 |
| Q921S7 | Mrpl37 | -4.043327758 | 3.265529133 |
| O88667 | Rrad | -4.047727914 | 2.081618343 |
| Q8BYY4 | Ttc39b | -4.047828112 | 3.254486056 |
| Q8VHX6 | Flnc | -4.053057168 | 6.770926025 |
| Q8C3R1 | Brat1 | -4.053818304 | 3.629634653 |
| Q99JP4 | Cdc26 | -4.055015952 | 2.802164633 |
| P48453 | Ppp3cb | -4.056735833 | 3.697293623 |
| Q8CJ26 | Nradd | -4.065438744 | 3.304496923 |
| Q922G2 | Fam76a | -4.066887599 | 2.292580383 |
| Q62187 | Ttf1 | -4.07199733 | 2.285943245 |
| Q61510 | Trim25 | -4.08139476 | 3.174574434 |
| A2AG58 | Bclaf3 | -4.081831477 | 2.19038063 |
| Q99PP9 | Trim16 | -4.083692143 | 2.456457146 |
| Q8R0X7 | Sgpl1 | -4.08517324 | 2.98010563 |
| Q60855 | Ripk1 | -4.089188278 | 4.620492661 |
| Q8BXN9 | Tmem87a | -4.090199338 | 2.093280602 |
| Q99P69 | Nuf2 | -4.091378511 | 2.689247298 |
| Q9EQN3 | Tsc22d4 | -4.091378799 | 4.855585066 |
| P21460 | Cst3 | -4.104060386 | 3.294153079 |
| Q6GQW0 | Abtb3 | -4.11317949 | 3.514055956 |
| P70444 | Bid | -4.134768523 | 3.592569889 |
| Q8BFS6 | Cpped1 | -4.141632132 | 3.464251363 |
| Q8CDJ8 | Ston1 | -4.149326766 | 2.39468315 |
| Q8BVA5 | Ldah | -4.149710396 | 3.058699242 |
| Q9DCH2 | Pop7 | -4.163543916 | 2.348457799 |
| P08228 | Sod1 | -4.172132635 | 2.709449244 |
| Q9R008 | Mvk | -4.172156367 | 2.271257664 |
| P97863 | Nfib | -4.179145355 | 2.523641025 |
| O35640 | Anxa8 | -4.186030998 | 2.403720605 |
| Q8BKT8 | Haus7 | -4.188386225 | 3.372791097 |
| Q8VCN5 | Cth | -4.192652979 | 3.559604896 |
| Q99JP6 | Homer3 | -4.195619207 | 3.879038068 |
| Q8R164 | Bphl | -4.200732776 | 5.126307477 |
| Q9WU79 | Prodh | -4.20697034 | 2.397252825 |
| P26645 | Marcks | -4.208995649 | 5.430170735 |
| Q99L04 | Dhrs1 | -4.215151997 | 4.603958583 |
| Q9JLR9 | Higd1a | -4.219019204 | 3.580777672 |
| Q8BYU6 | Tor1aip2 | -4.236816698 | 2.38123484 |
| P07356 | Anxa2 | -4.24192502 | 4.440791204 |
| Q6PAT0 | Adat3 | -4.242590297 | 2.853666849 |
| Q8VDP3 | Mical1 | -4.250375667 | 2.68164708 |
| O70348 | Dxo | -4.259704331 | 2.795515976 |
| Q9CQ54 | Ndufc2 | -4.267316099 | 2.318689914 |
| Q61036 | Pak3 | -4.268400421 | 3.763064603 |
| Q8BK58 | Hspbap1 | -4.274635001 | 2.339848664 |
| Q9D1C3 | Pyurf | -4.288427929 | 3.849709867 |
| Q9CQZ1 | Hsbp1 | -4.291390281 | 5.332618856 |
| Q60778 | Nfkbib | -4.306201844 | 4.206443196 |
| Q8BHA3 | Dtd2 | -4.307209217 | 4.858477354 |
| Q7TME2 | Spag5 | -4.308289465 | 3.611514886 |
| P97370 | Atp1b3 | -4.321661227 | 2.49816569 |
| Q8K1S3 | Unc5b | -4.328816011 | 2.747181476 |
| Q8VCH8 | Ubxn4 | -4.331564322 | 3.752166389 |
| Q8CD10 | Micu2 | -4.334704986 | 3.723605069 |
| Q9CQ06 | Mrpl24 | -4.336749991 | 3.609848513 |
| P56183 | Rrp1 | -4.338373059 | 3.229957208 |
| Q9CWV0 | Malsu1 | -4.340737316 | 3.344662619 |
| Q80ZQ9 | Abitram | -4.371954273 | 4.76408557 |
| Q8BMD8 | Slc25a24 | -4.372073292 | 2.613693916 |
| Q99MS8 | Tpgs1 | -4.373073253 | 2.736592084 |
| P25085 | Il1rn | -4.379199368 | 2.793617345 |
| Q6NXY1 | Tbc1d31 | -4.389444394 | 4.037387088 |
| Q9D2Q3 | Paat | -4.395805242 | 2.983036605 |
| Q60772 | Cdkn2c | -4.400460742 | 3.468893624 |
| Q8VC52 | Rbpms2 | -4.408530443 | 2.964667842 |
| Q99J47 | Dhrs7b | -4.415107267 | 2.811008552 |
| Q8BFQ9 | Klhl42 | -4.420112096 | 2.418932866 |
| P97465 | Dok1 | -4.425668443 | 2.660009604 |
| Q9ESP1 | Sdf2l1 | -4.426468802 | 3.885865711 |
| Q91XV3 | Basp1 | -4.428022218 | 2.725472326 |
| Q3TZX8 | Nol9 | -4.432998135 | 2.231501868 |
| P59759 | Mrtfb | -4.450708885 | 3.284501448 |
| Q80WT5 | Aftph | -4.461488594 | 2.329405984 |
| P12032 | Timp1 | -4.469475915 | 4.216846902 |
| P28798 | Grn | -4.470310352 | 2.614376375 |
| Q8CEE6 | Pask | -4.480650601 | 3.229985925 |
| Q8R5F7 | Ifih1 | -4.497896605 | 3.022102985 |
| Q9Z0F6 | Rad9a | -4.499798818 | 2.27593457 |
| P49962 | Srp9 | -4.503813395 | 4.082302706 |
| P57722 | Pcbp3 | -4.514101995 | 4.017740737 |
| A2ALU4 | Shroom2 | -4.515837174 | 3.363006212 |
| Q9CR89 | Ergic2 | -4.523800662 | 2.962938435 |
| Q64008 | Rab34 | -4.528992243 | 3.845464579 |
| Q3TUH1 | Tamm41 | -4.535008713 | 3.468897239 |
| Q3UA16 | Spc25 | -4.53757828 | 2.111297062 |
| Q61508 | Ecm1 | -4.55758673 | 2.514504122 |
| Q8R570 | Snap47 | -4.562956432 | 2.643962663 |
| Q811L6 | Mast4 | -4.568787347 | 2.814098851 |
| Q3ULW8 | Parp3 | -4.571864134 | 2.70678455 |
| Q9WV54 | Asah1 | -4.571971951 | 2.620533408 |
| P43406 | Itgav | -4.575820734 | 3.912550108 |
| Q8BX02 | Kank2 | -4.600010478 | 2.217460596 |
| Q9CQ62 | Decr1 | -4.609062866 | 2.154682353 |
| Q99JN2 | Klhl22 | -4.622385799 | 3.666888638 |
| Q9QZM2 | Polg2 | -4.631538881 | 3.787995056 |
| P20065 | Tmsb4x | -4.634051433 | 5.947495263 |
| Q9D1L0 | Chchd2 | -4.635390801 | 3.677793121 |
| Q60870 | Reep5 | -4.642918982 | 3.03291843 |
| Q8K4M5 | Commd1 | -4.643754639 | 2.594816609 |
| Q60953 | Pml | -4.653549214 | 3.859179907 |
| Q3V4B5 | Commd6 | -4.654017913 | 2.446184447 |
| Q80UW2 | Fbxo2 | -4.657828205 | 2.273496896 |
| Q5PSV9 | Mdc1 | -4.66160977 | 3.130566128 |
| Q9D7N6 | Mrpl30 | -4.668733283 | 2.926514861 |
| Q9DB29 | Iah1 | -4.680526079 | 2.64425344 |
| Q8C142 | Ldlrap1 | -4.686573004 | 2.974404108 |
| Q3UGC7 | Eif3j1 | -4.68748894 | 3.710340101 |
| P35951 | Ldlr | -4.688216022 | 2.574071954 |
| Q9CQ75 | Ndufa2 | -4.696708966 | 2.582720803 |
| P16675 | Ctsa | -4.701754408 | 4.428194814 |
| P48755 | Fosl1 | -4.702350789 | 4.550247216 |
| Q8N9S3 | Ahsa2 | -4.742118639 | 2.346014403 |
| Q9QWF0 | Chaf1a | -4.746832143 | 3.186080822 |
| Q78WZ7 | Polr1f | -4.748275664 | 4.037387088 |
| Q99LY9 | Ndufs5 | -4.749929584 | 2.56468471 |
| Q8C0T5 | Sipa1l1 | -4.761256189 | 2.352015065 |
| B1AR13 | Cisd3 | -4.765635462 | 2.551783992 |
| Q80WR5 |  | -4.773211991 | 2.868871943 |
| Q99N87 | Mrps5 | -4.773709297 | 3.372488468 |
| P14069 | S100a6 | -4.773913104 | 2.979488945 |
| Q61398 | Pcolce | -4.778157197 | 4.603934239 |
| Q9QZF2 | Gpc1 | -4.778221876 | 3.248555979 |
| Q9Z127 | Slc7a5 | -4.786019515 | 3.438821128 |
| Q8K3D3 | Swi5 | -4.803560033 | 3.813454386 |
| P50428 | Arsa | -4.809380808 | 2.806641507 |
| Q8R1F6 | Hid1 | -4.82154205 | 4.566740796 |
| Q9CPV5 | Pmf1 | -4.827701531 | 2.860741448 |
| Q8BP78 | Fra10ac1 | -4.839296154 | 4.345719625 |
| Q9DCS3 | Mecr | -4.840879024 | 5.598055492 |
| Q99KS2 | Ngrn | -4.842934086 | 2.89842208 |
| O54724 | Cavin1 | -4.853908838 | 2.844492573 |
| Q3UZA1 | Rcsd1 | -4.861278844 | 2.551596499 |
| Q9R112 | Sqor | -4.861343214 | 4.263848903 |
| Q8R1I1 | Uqcr10 | -4.861361792 | 2.507258679 |
| Q9D7E3 | Ovca2 | -4.881690933 | 2.779242118 |
| Q8C650 | Septin10 | -4.908853445 | 2.159257609 |
| Q76KJ5 | Polr1g | -4.919800401 | 2.10103513 |
| Q5BU09 | Eapp | -4.931919003 | 4.647099295 |
| Q8BTE0 | Sdhaf4 | -4.938110398 | 2.970660182 |
| Q3TQQ9 | Firrm | -4.946778956 | 2.187118986 |
| P40240 | Cd9 | -4.955171098 | 2.632979968 |
| Q9D8U0 | Ifrd2 | -4.959433947 | 2.088297311 |
| Q9D8S9 | Bola1 | -4.972472307 | 2.56854113 |
| Q8BQ30 | Ppp1r18 | -4.977610489 | 4.02514467 |
| Q6Q899 | Rigi | -4.977885398 | 2.872895852 |
| Q99LW0 | Ankrd10 | -4.980347016 | 3.144964035 |
| Q8BLH7 | Hirip3 | -4.986500766 | 2.167049269 |
| P43883 | Plin2 | -4.98861323 | 3.41292277 |
| Q60767 | Ly75 | -5.00518762 | 2.190804197 |
| Q8K1C0 | Angel2 | -5.017112246 | 2.964092436 |
| Q8BGX2 | Timm29 | -5.018899523 | 3.341607078 |
| Q61733 | Mrps31 | -5.019238628 | 2.617449636 |
| Q69ZN7 | Myof | -5.019621308 | 5.395105809 |
| Q8K2M0 | Mrpl38 | -5.033419958 | 2.817212683 |
| P54923 | Adprh | -5.034015251 | 3.821881907 |
| Q91YR9 | Ptgr1 | -5.035179334 | 2.838117359 |
| Q8BSL7 | Arf2 | -5.037099697 | 3.186179563 |
| Q6P3Y5 | Znf280c | -5.041973656 | 4.617484307 |
| Q9CWG8 | Ndufaf7 | -5.044381833 | 3.33964369 |
| Q499E6 | Airim | -5.049082523 | 2.221690808 |
| Q9DAZ9 | Zfyve19 | -5.057623251 | 2.682922066 |
| Q69ZZ9 | Macf1 | -5.077119715 | 4.298653757 |
| Q61468 | Msln | -5.082693259 | 2.290001437 |
| Q14AI0 | DSCC1 | -5.092407501 | 2.054055913 |
| Q9D945 | Llph | -5.092724895 | 2.755222408 |
| Q9R0X4 | Acot9 | -5.095742224 | 2.939581562 |
| Q791N7 | Polr1h | -5.103139294 | 2.083043383 |
| Q8R151 | Znfx1 | -5.107011486 | 2.084849485 |
| Q8K4F5 | Abhd11 | -5.113477826 | 2.912179584 |
| P62965 | Crabp1 | -5.168433105 | 4.119469655 |
| Q6SKR2 | Hemk2 | -5.180769325 | 3.557568164 |
| Q91W92 | Cdc42ep1 | -5.185859468 | 2.57824685 |
| Q9R118 | Htra1 | -5.186978697 | 3.066585472 |
| Q9D6U8 | Fam162a | -5.229101537 | 2.097029252 |
| Q99K30 | Eps8l2 | -5.23008767 | 3.146700157 |
| Q91V61 | Sfxn3 | -5.241576311 | 2.681258589 |
| Q8C3F2 | Fam120c | -5.251306498 | 3.579449078 |
| Q80TM6 | R3hdm2 | -5.287361107 | 2.584649103 |
| Q8BHL4 | Gprc5a | -5.294571878 | 3.466733568 |
| Q9D3W4 | Gpn3 | -5.308919275 | 3.240497318 |
| Q8K2T1 | Nmral1 | -5.318582707 | 2.653186369 |
| P99025 | Gchfr | -5.342149141 | 2.60511992 |
| Q9DCT1 | Akr1e2 | -5.362795148 | 2.416802306 |
| Q61033 | Tmpo | -5.373493626 | 2.289898102 |
| Q9D786 | Haus5 | -5.380956525 | 3.045940051 |
| Q9R1Z7 | Pts | -5.381731636 | 2.629137098 |
| Q9CY45 | Eef1akmt1 | -5.387026344 | 3.399419979 |
| P37889 | Fbln2 | -5.396023773 | 3.194086671 |
| O70503 | Hsd17b12 | -5.411079249 | 2.874485849 |
| P97825 | Jpt1 | -5.415894551 | 3.969204766 |
| P70699 | Gaa | -5.426328283 | 3.826167348 |
| Q9D2Y4 | Mlkl | -5.43824359 | 3.598367259 |
| Q8VE94 | Fam110c | -5.455345732 | 5.253470976 |
| Q91V76 |  | -5.456494274 | 4.011228075 |
| Q8C8T8 | Tsr2 | -5.470567053 | 2.087831549 |
| P48024 | Eif1 | -5.48057702 | 4.812323805 |
| Q8VCR7 | Abhd14b | -5.485717846 | 2.87661709 |
| P35456 | Plaur | -5.505365859 | 2.571749104 |
| Q62470 | Itga3 | -5.50546417 | 3.076065522 |
| P10404 |  | -5.521415937 | 2.602000835 |
| P06797 | Ctsl | -5.539424976 | 4.081175116 |
| Q8VDT9 | Mrpl50 | -5.544208748 | 3.694641614 |
| Q8BGB5 | Limd2 | -5.580550222 | 2.569398422 |
| Q99KR3 | Lactb2 | -5.606814819 | 3.693186434 |
| P24860 | Ccnb1 | -5.666620942 | 3.045940051 |
| Q91YU8 | Ppan | -5.689545679 | 3.288665371 |
| Q8CJ27 | Aspm | -5.718216976 | 2.520480655 |
| Q8BK72 | Mrps27 | -5.724289618 | 3.173384795 |
| P56389 | Cda | -5.7253845 | 2.255627584 |
| P97315 | Csrp1 | -5.734469451 | 4.846861682 |
| Q3U155 | Ccdc174 | -5.750046195 | 2.217547278 |
| Q9D1R2 | Kti12 | -5.760143907 | 3.614088333 |
| Q8BUI3 | LRWD1 | -5.788130715 | 4.767056497 |
| O54974 | Lgals7 | -5.793489151 | 3.688585043 |
| Q9R1Q7 | Plp2 | -5.795965865 | 4.222233379 |
| O09159 | Man2b1 | -5.802014698 | 3.053989081 |
| P09926 | Surf2 | -5.81404643 | 3.265529133 |
| P52760 | Rida | -5.820033605 | 3.704371678 |
| Q9DCM0 | Ethe1 | -5.872809812 | 3.474427899 |
| P45377 | Akr1b8 | -5.895189614 | 3.860028789 |
| P81069 | Gabpb2 | -5.912851419 | 3.206875116 |
| P10107 | Anxa1 | -5.926591547 | 5.217932867 |
| Q8BSI6 | R3hcc1 | -5.939554543 | 4.404322347 |
| Q9JL35 | Hmgn5 | -5.982206056 | 4.49650809 |
| O89110 | Casp8 | -6.005901695 | 2.451282317 |
| P10923 | Spp1 | -6.018044321 | 2.590912276 |
| Q3TLP5 | Echdc2 | -6.072010732 | 3.509511101 |
| Q9CQS5 | Riok2 | -6.074829589 | 2.198635668 |
| Q9CQ18 | Rnaseh2c | -6.138212006 | 2.386970619 |
| Q6PDY2 | Ado | -6.171360098 | 2.780906281 |
| O08997 | Atox1 | -6.184717038 | 5.771121261 |
| Q9D083 | Spc24 | -6.217893973 | 3.195766442 |
| Q80ZS3 | Mrps26 | -6.237953274 | 3.795884272 |
| B1ARD6 | Slfn9 | -6.293417542 | 2.630599244 |
| P14901 | Hmox1 | -6.308895705 | 4.106041097 |
| Q3UPF5 | Zc3hav1 | -6.314088232 | 3.935528444 |
| P70202 | Lxn | -6.332660764 | 4.199705293 |
| P97450 | Atp5pf | -6.340443355 | 2.730795759 |
| P18572 | Bsg | -6.359033894 | 3.864979693 |
| Q9CXL3 | Chlsn | -6.361301224 | 3.462406627 |
| P48725 | Pcnt | -6.371739574 | 3.009891812 |
| P52927 | Hmga2 | -6.403314303 | 2.48046476 |
| Q9QZN4 | Fbxo6 | -6.434757014 | 3.999455073 |
| Q3TYS2 | Cybc1 | -6.436649735 | 3.307733158 |
| Q9CQM5 | Txndc17 | -6.437768367 | 5.325519695 |
| Q62426 | Cstb | -6.458610329 | 3.398144895 |
| Q80U04 | Pja2 | -6.502596151 | 4.237734352 |
| P60330 | Espl1 | -6.506064117 | 3.678162621 |
| P16045 | Lgals1 | -6.537163089 | 5.551659592 |
| P00405 | Mtco2 | -6.562901584 | 2.160148676 |
| P51125 | Cast | -6.594453977 | 2.786803346 |
| P56135 | Atp5mf | -6.665467427 | 4.863228482 |
| Q64105 | Spr | -6.715985654 | 2.545350893 |
| Q9D855 | Uqcrb | -6.735207859 | 3.016432923 |
| Q8BVY0 | Rsl1d1 | -6.761749018 | 2.273496896 |
| Q8BJY1 | Psmd5 | -6.857395187 | 5.617406177 |
| P98078 | Dab2 | -6.918260461 | 3.178972106 |
| Q80VJ3 | Dnph1 | -6.946699914 | 3.765148542 |
| P15208 | Insr | -7.036310664 | 3.92586922 |
| Q9CRB2 | Nhp2 | -7.05319834 | 4.552874983 |
| P08207 | S100a10 | -7.180790023 | 2.953572465 |
| P19157 | Gstp1 | -7.239083498 | 3.030000236 |
| P99028 | Uqcrh | -7.274199078 | 3.552456506 |
| Q64339 | Isg15 | -7.298209333 | 3.108808549 |
| P24452 | Capg | -7.344719594 | 2.064389141 |
| Q60854 | Serpinb6 | -7.361194214 | 5.551659592 |
| P15379 | Cd44 | -7.485239314 | 3.398974144 |
| Q9CPQ1 | Cox6c | -7.511940903 | 3.191187017 |
| Q9QZL0 | Ripk3 | -7.556166645 | 4.831192686 |
| P07091 | S100a4 | -7.628910793 | 3.981151966 |
| O35639 | Anxa3 | -7.649283409 | 2.805866988 |
| Q9CPQ8 | Atp5mg | -7.687597222 | 2.514504122 |
| O35215 | Ddt | -7.770390285 | 4.734409437 |
| P31786 | Dbi | -8.955695635 | 4.777690664 |
| O09131 | Gsto1 | -8.970589077 | 4.49650809 |
| P16110 | Lgals3 | -9.005892128 | 3.941279595 |

**Table S2.** *Protein signature of 2022 MPXV MOI 1 infected KO vs WT MEF lysates at 21 hours post-infection. Table lists the protein ID, the log2-fold change (FC) (MPXV infected WT MEFs vs infected ISG15 KO MEFs) of the levels of each protein, and the statistical significance (−log P value). Ordered from largest to smallest FC.*
