## Supplementary material for "ISG15 Differentially Modulates Clade Ib and II MPXV Infection in MEF cells": TableS4

| **Name** | **Gene** | **fold_change** | **p_value_adj_neg_log10** |
| --- | --- | --- | --- |
| Q9Z0N2 | Eif2s3y | 7.524719717 | 3.265922864 |
| P30275;Q6P8J7 | Ckmt1, Ckmt2 | 7.066626151 | 3.900154741 |
| Q64522 | H2ac21 | 7.012390021 | 2.516548125 |
| Q8VIN1 | Pbp2 | 6.939496831 | 3.572541618 |
| Q80TL0 | Ppm1e | 6.803674792 | 3.113400012 |
| Q9DAJ5 | Dynlrb2 | 6.692128027 | 3.052209902 |
| P28651 | Ca8 | 6.671443339 | 4.023016918 |
| P31254 | Uba1y | 6.092465675 | 3.355111563 |
| Q8VHG2 | Amot | 5.735189688 | 3.490650649 |
| Q91WT9 | Cbs | 5.631088507 | 5.073302231 |
| Q80ZN9 | Cox6b2 | 5.555679485 | 2.631558157 |
| P00920 | Ca2 | 5.3722911 | 2.478357657 |
| Q8VEB1 | Grk5 | 5.358935568 | 4.883945468 |
| P30275 | Nefm | 5.313434624 | 2.474698964 |
| P08553 | Ldhb | 5.260312872 | 4.392132228 |
| P16125 | Gstm3 | 5.223081992 | 5.481832748 |
| P19639 | Itk | 5.123797972 | 4.375749023 |
| Q03526 | Kif16b | 5.058019353 | 2.429255075 |
| B1AVY7 | Galm | 5.038728666 | 3.602282682 |
| Q8K157 | Pcbd1 | 4.92967148 | 3.984452126 |
| P61458 | Cx3cl1 | 4.857988606 | 3.861784354 |
| O35188 | Plxna2 | 4.731496402 | 4.795605059 |
| P70207 | Ca2 | 4.704402478 | 3.676721531 |
| Q9ET01 | Pygl | 4.682772326 | 3.995737309 |
| Q60611 | Satb1 | 4.669135509 | 2.80518561 |
| O55091 | Impact | 4.618309919 | 3.208294899 |
| Q8BRK8 | Prkaa2 | 4.323692272 | 2.59059432 |
| P08551 | Nefl | 4.315877028 | 2.176762317 |
| P58321 | Uchl4 | 4.287364404 | 4.406025734 |
| Q61474 | Msi1 | 4.277083839 | 3.115819943 |
| O55111 | Dsg2 | 4.249138859 | 3.149979902 |
| Q8JZV9 | Bdh2 | 4.235707728 | 2.533694711 |
| Q60936 | Coq8a | 4.220914978 | 3.00594908 |
| Q9D711 | Pir | 4.18552455 | 3.528249765 |
| Q9Z1N7 | Arid3b | 4.180896279 | 2.385166613 |
| Q6PDS3 | Sarm1 | 4.147579507 | 2.215363955 |
| Q9JHI5 | Ivd | 4.145983113 | 4.327341891 |
| Q9CZC8 | Scrn1 | 4.138544838 | 3.242440779 |
| Q58A65;Q9ESN9 | Spag8; Mapk8ip3 | 4.083685862 | 2.851468308 |
| Q04447;P07310 | Ckb;Ckm | 4.051402622 | 5.060390014 |
| P97367 | Meis2 | 4.042891575 | 3.704403367 |
| Q9EQF6 | Dpysl5 | 4.017616432 | 2.631948076 |
| Q9D5J6 | Shpk | 4.014324415 | 2.861838547 |
| Q8VCK3 | Tubg2 | 4.004696447 | 2.514610742 |
| Q64524 | H2bc21 | 3.995425232 | 3.892115295 |
| O55042;Q91ZZ3 | Snca;Snacb | 3.960640069 | 3.113400012 |
| Q8R0P4 | Aamdc | 3.854369092 | 2.514952522 |
| Q8R1G2 | Cmbl | 3.788612655 | 2.585742998 |
| O55042 | Pdlim3 | 3.773047295 | 2.550034836 |
| O70209 | Hdac6 | 3.752391092 | 3.691613061 |
| Q9Z2V5 | Radx | 3.751875939 | 2.031803138 |
| Q8C779 | Tcp11l1 | 3.660645993 | 2.824218045 |
| Q8BTG3 | Ap3b2 | 3.660169615 | 2.074528631 |
| Q9JME5 | Acot5 | 3.653307237 | 3.226528825 |
| Q8VHG2 | D1Pas1 | 3.637479388 | 2.094159125 |
| Q6Q2Z6 | Cdkn2b | 3.618092246 | 2.656749287 |
| P16381 | Cps1 | 3.609758607 | 3.034735344 |
| P55271 | Aamdc | 3.548671432 | 4.797847856 |
| Q8C196 | Cmbl | 3.519705272 | 2.862815487 |
| Q62440;Q62441;Q9WVB2 | Tle1;Tle3;Tle2 | 3.516699495 | 2.121202048 |
| Q9Z2G9 | Htatip2 | 3.504293985 | 4.062999703 |
| P28738;P33175 | Kif5c;Kif5a | 3.407358644 | 2.917882694 |
| Q9D114 | Hddc3 | 3.389638019 | 3.043860565 |
| P49813 | Tmod1 | 3.372138199 | 2.480551402 |
| P97447 | Fhl1 | 3.295774694 | 5.220959003 |
| P97931 | Ung | 3.204096578 | 2.745869418 |
| P70265 | Pfkb2 | 3.142024841 | 2.38818275 |
| Q9CQE0 | Rnf138 | 3.093237514 | 2.052402574 |
| Q9CQE5 | Rgs10 | 3.093053011 | 2.34252566 |
| Q62441 | Ina | 3.084129677 | 2.786056352 |
| P46660 | Aldh1a2 | 3.05801355 | 4.090704502 |
| Q62148 | Glo1 | 3.048270235 | 4.786088482 |
| Q9CPU0 | Arhgdib | 3.028986173 | 5.296676108 |
| Q61599 | Pck1 | 3.018571802 | 6.136656712 |
| Q9Z2V4 | Serpinb1c | 3.018553734 | 3.698857208 |
| Q5SV42 | Rnf138 | 2.999341526 | 2.285378897 |
| P22227 | Zfp42 | 2.967941046 | 2.337580435 |
| Q9DB07 | Ift46 | 2.95823115 | 3.113400012 |
| P41139 | Id4 | 2.935221669 | 3.376988398 |
| A3KMP2 | Ttc38 | 2.924188201 | 3.811777131 |
| Q60899 | Elavl2 | 2.915795221 | 3.049389513 |
| Q5SVQ0 | Kat7 | 2.898406087 | 2.467157893 |
| B1ARW8 | Ca122 | 2.886063464 | 3.998412163 |
| Q91ZE0 | Tmlhe | 2.862034104 | 2.534503568 |
| Q60611 | Mnx1 | 2.86100963 | 2.059190471 |
| Q9QZW9 | Kctd15 | 2.853311904 | 3.396273085 |
| Q8K0E1 | Dhtkd1 | 2.849788739 | 3.98600012 |
| Q04447 | Ank3 | 2.841636708 | 5.296676108 |
| A2ATU0 | Glul | 2.832202014 | 2.175671397 |
| G5E8K5 | Cutc | 2.827118308 | 2.31489916 |
| P15105 | Tmlhe | 2.8148128 | 4.971761795 |
| Q9D8X1 | Mnx1 | 2.810628013 | 3.851287414 |
| Q6P3D0;Q8VHN8 | Nudt16 | 2.806871145 | 2.792820137 |
| Q9DA03 | Lyrm7 | 2.783577087 | 4.420212838 |
| P33173 | Kif1a | 2.766149657 | 2.042620074 |
| Q91X97 | Ncald | 2.75257278 | 2.626676414 |
| Q9WU63 | Hebp2 | 2.708681875 | 3.632049898 |
| P23475 | Xrcc6 | 2.701792056 | 5.079252495 |
| Q9D1H6 | Ndufaf4 | 2.651990719 | 3.169344225 |
| Q9JIG8 | Praf2 | 2.643029985 | 2.254500207 |
| O70583 | Mid1 | 2.63940355 | 3.4043404 |
| Q9CQA9 | Ntpcr | 2.58222432 | 3.522348899 |
| Q8R238 | Sdsl | 2.555383927 | 2.163673951 |
| Q5XG69 | Fam169a | 2.498210347 | 3.696494684 |
| Q5RL51 | Gstcd | 2.494027352 | 2.414700468 |
| P16627 | Hspa1l | 2.490543388 | 3.867651428 |
| Q9CZS1 | Aldh1b1 | 2.475089819 | 3.236128002 |
| Q91W43 | Gldc | 2.470708598 | 2.515800416 |
| P27790 | Cenpb | 2.450695076 | 2.021260858 |
| Q9QYB8 | Add2 | 2.406245925 | 4.057208694 |
| Q8VI93 | Oas3 | 2.401060612 | 3.274412271 |
| P48758 | Cbr1 | 2.380521709 | 3.244695414 |
| P05213 | Tuba1b | 2.353581254 | 3.954965996 |
| P60824 | Cirbp | 2.331325883 | 4.793088184 |
| Q9DB50 | Ap1s2 | 2.317645508 | 2.466614384 |
| P0C027 | Nudt10 | 2.317113099 | 2.047324049 |
| Q9QY15 | Ddx25 | 2.311644935 | 4.057208694 |
| Q9ERE3 | Sgk3 | 2.283029803 | 2.463220879 |
| Q99MU3 | Adar | 2.26219108 | 4.770682066 |
| Q5BKP2 | Usp13 | 2.262145611 | 2.253402406 |
| P15626 | Gstm2 | 2.246591205 | 2.521000194 |
| Q641K1 | Agtpbp1 | 2.241440343 | 4.127697567 |
| Q7TPM6 | Fsd1 | 2.23362821 | 4.044096318 |
| P16627 | Hps1ab | 2.228833162 | 3.41680066 |
| Q9R0Q6 | Arpc1a | 2.217252561 | 5.139550781 |
| P63101 | Ywhaz | 2.180013143 | 2.196559115 |
| Q62420 | Sh3gl2 | 2.148175118 | 2.114798687 |
| O35984 | Pbx2 | 2.125209933 | 4.640089824 |
| Q7TPD6 | Raver2 | 2.110454894 | 2.000628934 |
| Q149C2 | Traf3ip1 | 2.065249093 | 3.272024426 |
| Q8BI84 | Mia3 | 2.051031074 | 4.795605059 |
| Q9D023 | Mpc2 | 2.045653462 | 2.862912095 |
| P68372;P99024;Q7TMM9;Q9CWF2;Q9D6F9;Q9ERD7 | Tubb | 2.040596149 | 3.414664586 |
| Q9Z2Z9 | Gfpt2 | 2.03367097 | 2.015833317 |
| Q9D7X8 | Ggct | 2.019969721 | 3.975693654 |
| Q9CWF2 | Zcchc3 | 2.016862565 | 3.051701664 |
| Q8BPK2 | Epb41l5 | 2.010581386 | 3.861419064 |
| Q8BGS1;Q9JMC8 | Epb41l4b | 2.007630137 | 3.788792631 |
| Q9Z129 | Recql | 1.981471189 | 4.183809614 |
| Q9WVL1 | Ap4s1 | 1.965541047 | 4.131815145 |
| Q9QXE0 | Hacl1 | 1.956763713 | 2.371290746 |
| Q9CPN8 | Igf2bp3 | 1.955007599 | 4.207432459 |
| Q9ET01 | Kctd2 | 1.950829337 | 3.144857264 |
| Q8CEZ0 | Pawr | 1.935574254 | 3.104467292 |
| Q925B0 | Gfpt2 | 1.929619018 | 4.016822721 |
| P23198;P83917 | Cbx3;Cbx1 | 1.885988621 | 3.863193969 |
| Q8C7Q4 | Rbm4 | 1.866769209 | 2.988286639 |
| Q9DB76 | Emc9 | 1.848244964 | 2.588346309 |
| O88477 | Lgf2bp1 | 1.844856132 | 3.676721531 |
| Q3UUI3 | Them4 | 1.831752176 | 2.006168341 |
| Q61233;Q99K51 | Lcp1; Pls3 | 1.821986007 | 4.259173414 |
| Q03173 | Enah | 1.805931853 | 4.709620529 |
| Q8BWP8 | B4gat1 | 1.798975039 | 3.170236202 |
| B1AXP6 | Tomm5 | 1.798973917 | 2.988942342 |
| Q9ERD6 | Ralgps2 | 1.778892387 | 3.621386629 |
| Q5SSL4;Q6PAJ1 | Abr;Bcr | 1.777863872 | 3.051114047 |
| P11103 | Parp1 | 1.733839568 | 2.930534036 |
| P00342;P06151 | Ldhc;Ldha | 1.72938075 | 4.311461067 |
| Q9D032 | Ssbp3 | 1.718410877 | 3.957686009 |
| Q8CB27 | Yod1 | 1.69642395 | 4.186900043 |
| Q8JZS0 | Lin7a | 1.69595845 | 3.114596801 |
| P83510 | Tnik | 1.674361509 | 2.103394084 |
| Q5SU73 | Coil | 1.658097458 | 3.769260716 |
| Q8R001 | Mapre2 | 1.640653497 | 4.50536728 |
| Q8R123 | Flad1 | 1.636832005 | 3.327491975 |
| Q9CQZ7 | Polr3k | 1.594390492 | 3.331322344 |
| Q9JL70 | Fanca | 1.589988629 | 3.836472853 |
| Q8CHY3 | Dym | 1.582588126 | 2.164765396 |
| Q8BYJ6 | Tbc1d4 | 1.576673549 | 3.676659831 |
| Q8K0Z7 | Taco1 | 1.563916871 | 4.657840006 |
| P97313 | Prkdc | 1.553856692 | 3.463450845 |
| Q9D394 | Rufy3 | 1.54077052 | 2.453894304 |
| Q80V62 | Fancd2 | 1.521975895 | 2.766842597 |
| P56812 | Pdcd5 | 1.519568867 | 4.309548293 |
| Q61189 | Clns1a | 1.519058584 | 3.236128002 |
| O89053 | Coro1a | 1.510971137 | 2.18427048 |
| A0A7H0DN48 | OPG075 | 1.493262365 | 3.190379089 |
| P97364 | Sephs2 | 1.489514606 | 2.259502065 |
| Q9CQF6 | Aasdhppt | 1.488348358 | 4.145287568 |
| Q9CXU9 | Eif1b | 1.481042641 | 4.166146413 |
| O70456 | Sfn | 1.477952623 | 3.981051698 |
| Q9ESW4 | Agk | 1.468360166 | 2.510314281 |
| M1KJ15 | OPG114 | 1.46491767 | 3.350995472 |
| Q922B1 | Macrod1 | 1.456617706 | 3.209815688 |
| Q8CAK1 | Iba57 | 1.442936027 | 3.71025346 |
| Q45VK7 | Dync2h1 | 1.440673342 | 3.073805414 |
| A0A7H0DNC1 | OPG149 | 1.438875048 | 2.829841609 |
| Q9QXA7 | Trim44 | 1.437685312 | 2.110403521 |
| M1LBQ5 | OPG115 | 1.433142394 | 3.948873913 |
| Q8VD33 | Sgtb | 1.430730486 | 2.550003929 |
| O55060 | Tpmt | 1.42922412 | 4.1255601 |
| A6H630 | Armt1 | 1.423616711 | 4.970683561 |
| A0A7H0DN78 | OPG106 | 1.411998893 | 3.333820654 |
| Q8VE92 | Rbm4b | 1.403646378 | 4.676729037 |
| Q08122 | Tle3 | 1.397123561 | 4.289966041 |
| Q8BHN1 | Txlng | 1.396798607 | 3.49155129 |
| Q3TIR3;Q80XE1 | Ric8a;Ric8b | 1.390354067 | 2.99217068 |
| Q99J10 | Ctu1 | 1.377834967 | 2.926513144 |
| O08848 | RO60 | 1.370721143 | 4.375749023 |
| A2AQ07 | Tubb1 | 1.355814045 | 3.844929779 |
| P61079;P61080;P62838 | Ube2d | 1.35101218 | 3.371940058 |
| Q9CQE3 | Mrps17 | 1.347950683 | 2.616773913 |
| P16460 | Ass1 | 1.346332607 | 3.56383271 |
| Q6DG52 | Churc1 | 1.329271558 | 3.57573772 |
| Q3V0K9 | Pls1 | 1.326428347 | 2.523164883 |
| Q9QXK3;Q9QZE5 | Copg2;Copg1 | 1.324781225 | 4.188397876 |
| Q9Z130 | Hnrnpdl | 1.313391492 | 3.687706685 |
| P62855 | Rps26 | 1.313165933 | 2.421789684 |
| P48774 | Gstm5 | 1.308355495 | 3.445313976 |
| Q8VED9 | Lgalsl | 1.287425134 | 2.092301937 |
| A0A7H0DN42 | OPG069 | 1.284998962 | 4.006897472 |
| E9Q394 | Akap13 | 1.284993955 | 2.688460386 |
| Q9Z1X4 | Ilf3 | 1.281466598 | 3.612529958 |
| Q6DIC0 | Smarca2 | 1.279984067 | 3.113400012 |
| Q99K46 | Usp11 | 1.274048435 | 2.087347798 |
| Q8BGS2 | Bola2 | 1.264941217 | 2.480778618 |
| Q8BZ21 | Kat6a | 1.260882773 | 2.012768634 |
| Q8C167 | Prepl | 1.242823014 | 4.053475595 |
| Q9CQ02 | Commd4 | 1.240829617 | 2.611965946 |
| Q8R2U4 | Ntmt1 | 1.238364925 | 4.590301888 |
| Q61699 | Hpsh1 | 1.232856613 | 2.7467421 |
| Q9CZT4 | Polr3e | 1.231038447 | 3.781421365 |
| Q91VN1 | Znf24 | 1.227596723 | 3.041645729 |
| O35969 | Gamt | 1.224343495 | 3.7075662 |
| P60840 | Ensa | 1.213513219 | 2.941838767 |
| P60521 | Gabarapl2 | 1.210188555 | 3.423447522 |
| Q80UM3;Q9DBB4 | Naa15;Naa16 | 1.199920114 | 4.145136909 |
| Q8VDS4 | Rprd1a | 1.199517336 | 4.103666555 |
| Q9D273 | Mmab | 1.18130081 | 3.064222985 |
| P07742;A0A7H0DN52 | Rrm1;OPG80 | 1.172189618 | 2.532456069 |
| Q8BJZ4 | Mrps35 | 1.167361009 | 3.532188422 |
| Q9EQS9 | Igdcc4 | 1.166836892 | 3.191697306 |
| Q9D0T2 | Dusp12 | 1.15507975 | 2.619031226 |
| Q9R0P9 | Uchl1 | 1.154997331 | 3.220594607 |
| Q8CA72 | Gan | 1.154946133 | 2.572458873 |
| Q8BIP0 | Dars2 | 1.154385666 | 4.04704904 |
| Q9QY93 | Dctpp1 | 1.148895015 | 4.044096318 |
| Q9JJV2 | Pfn2 | 1.147554985 | 3.401526691 |
| Q8BG93 | Nudt15 | 1.140306773 | 2.44222007 |
| Q9EST4 | Psmg2 | 1.132979676 | 3.487766872 |
| Q9ERA0 | Tfcp2 | 1.131076584 | 2.474289796 |
| Q3V0K9 | Macroh2a2 | 1.125531364 | 3.429279945 |
| Q8CCK0 | Pgm3 | 1.121110712 | 2.63362868 |
| Q9CYR6 | Fanci | 1.115735308 | 4.784426578 |
| Q8K368 | Pagr1a | 1.110011706 | 2.743997233 |
| Q99L02 | Rmdn1 | 1.108757272 | 3.483097391 |
| Q9DCV4 | Psmd10 | 1.107970286 | 3.418183112 |
| Q9Z2X2 | Nudt15 | 1.106657838 | 3.597742334 |
| A0A7H0DNE6 | OPG180 | 1.1054361 | 3.64585918 |
| Q9ERV1 | Mkrn2 | 1.100409826 | 2.276827081 |
| Q9R0P4 | Smap | 1.099163462 | 4.184940346 |
| Q9DD18 | Dtd1 | 1.095358001 | 3.2639429 |
| Q3UDK1 | Trafd1 | 1.093702337 | 4.005229598 |
| A0A7H0DND9;Q8V4T7 | OPG171 | 1.092556936 | 3.538507794 |
| Q920Q6 |  | 1.077481226 | 3.569322397 |
| Q8BUH1 | Txnl4b | 1.071907101 | 2.008610176 |
| Q3TCJ1 | Abraxas2 | 1.059559769 | 2.988859299 |
| Q9D6M3 | Slc25a22 | 1.058924967 | 4.062999703 |
| Q8CI70 | Lrrc20 | 1.057192105 | 2.688460386 |
| Q6PE54 | Dhx40 | 1.05541617 | 2.258856267 |
| Q91V64 | Isoc1 | 1.046887699 | 3.658574246 |
| Q8C5N5 | Pdcd2l | 1.04333613 | 3.469268937 |
| Q99KR6 | Rnf34 | 1.041771942 | 3.007994335 |
| Q9R0D8 | Wdr54 | 1.040146073 | 2.538926105 |
| Q91VL8 | Terf2ip | 1.035892727 | 3.187546034 |
| Q8VEJ9 | Vps4a | 1.029429175 | 3.387312331 |
| Q9CQ89 | Cuta | 1.027981601 | 3.469498339 |
| Q9QZX7 | Srr | 1.027250999 | 2.029824362 |
| P22907 | Hmbs | 1.024920631 | 3.165103191 |
| Q9CR09 | Ufc1 | 1.023227999 | 4.45736006 |
| Q80ZI6 | Lrsam1 | 1.021545894 | 2.582695782 |
| Q66JV4;Q80YR9 | Rbm12b | 1.017186454 | 2.8492975 |
| Q9CXJ1 | Ears2 | 1.015808325 | 2.692990992 |
| Q2TPA8 | Hsdl2 | 1.015173799 | 3.340672865 |
| Q05186 | Rcn1 | 1.014495322 | 3.674422346 |
| Q3U0V1;Q91WJ8 | Fubp | 1.011923257 | 3.266881356 |
| A0A7H0DNA4 | OPG132 | 1.000619383 | 3.894426896 |
| A0A7H0DN76 | OPG104 | 0.999615627 | 2.416508747 |
| Q9ESZ8 | Gtf2i | 0.996534958 | 2.353121513 |
| Q91WM1 | Strbp | 0.99407885 | 3.984452126 |
| Q9CZ62 | Cep97 | 0.993093843 | 2.03802988 |
| Q6RI63 | Fam120b | 0.990448476 | 2.408602828 |
| Q9D0B0 | Srsf9 | 0.987882164 | 4.101631499 |
| P52482;P52483;Q91W82 | Ube2e | 0.979887394 | 2.432948325 |
| Q64261 | Cdk6 | 0.978543353 | 3.020139817 |
| Q3TC72 | Fahd2a | 0.973876719 | 2.086552024 |
| Q9D1R9 | Rpl34 | 0.968636609 | 3.0646993 |
| Q64331 | Myo6 | 0.964795252 | 3.80055133 |
| A0A7H0DN80 | OPG108 | 0.963681378 | 2.785320803 |
| Q8VDM6 | Hnrnpul1 | 0.960483524 | 4.271958081 |
| Q8K4F6 | Nsun5 | 0.959233377 | 2.6286283 |
| Q9CWQ0 | Dph5 | 0.954046906 | 2.646631443 |
| O08580;Q61539 | Essr | 0.949084618 | 2.487425976 |
| Q8R4X3 | Rbm12 | 0.948530081 | 4.36798556 |
| Q91WM1 |  | 0.94492964 | 3.709933871 |
| Q8BMJ3 | Eif1ax | 0.939250014 | 3.469200436 |
| Q9CWW6 | Pin4 | 0.937197234 | 3.768338095 |
| Q8R3D1 | Tbc1d13 | 0.933023141 | 3.622028593 |
| Q7TSH2 | Phkb | 0.930495266 | 2.129970321 |
| P62137;P63087 | Ppp1c | 0.923290349 | 2.868266047 |
| Q9D8T7 | Slirp | 0.92322054 | 2.651263928 |
| P97823 | Lypla1 | 0.920563695 | 3.638277906 |
| P00493 | Hprt1 | 0.917439633 | 4.155788101 |
| Q9ESE1 | Lrba | 0.916905238 | 4.098156107 |
| A0A7H0DNB5 | OPG143 | 0.914728336 | 2.174809252 |
| A0A7H0DN27 | OPG054 | 0.914481827 | 3.249993247 |
| Q9CXY6 | Ilf2 | 0.908958799 | 3.418453048 |
| Q3UMY5 | Eml4 | 0.90890516 | 2.804591016 |
| A0A7H0DN37 | OPG064 | 0.907052398 | 3.204566354 |
| P59048 | Pdrg1 | 0.906922462 | 3.354031106 |
| Q8BGC0 | Htatsf1 | 0.906604606 | 2.94594141 |
| Q3UPH1 | Prrc1 | 0.903562792 | 3.597547394 |
| A0A7H0DN72 | OPG100 | 0.901863213 | 3.851287414 |
| P81269;Q01147 | Atf1;Creb1 | 0.898286218 | 2.75608081 |
| Q91WJ8 |  | 0.886708713 | 4.335067817 |
| M1L543 | OPG138 | 0.885735358 | 2.127845888 |
| P97855;P97379 | G3bp | 0.884557201 | 3.032937407 |
| P62305 | Snrpe | 0.87854115 | 2.120974394 |
| Q9Z204 | Hnrnpc | 0.877859024 | 3.142356764 |
| A0A7H0DN57 | OPG085 | 0.872419405 | 2.467980131 |
| Q99LZ3 | Gins4 | 0.869780205 | 2.481159532 |
| Q9EPN1 | Nbea | 0.869231437 | 2.98480045 |
| P23492 | Pnp | 0.867145378 | 4.309548293 |
| Q99P31 | Hspbp1 | 0.865163598 | 4.259173414 |
| O54879 | Hmgb3 | 0.862674804 | 3.569896355 |
| Q9Z0H4 | Celf2 | 0.862596285 | 3.786435915 |
| P48193 | Epb41 | 0.861258105 | 3.504348547 |
| Q11011 | Npepps | 0.85940093 | 4.870110187 |
| Q9EPK7;Q99NF8 | Ranbp17 | 0.858375174 | 3.007047092 |
| Q9CV28 | Mindy3 | 0.849298773 | 2.961162955 |
| P50096 | Impdh1 | 0.845399781 | 3.095062935 |
| A0A7H0DN63 | OPG091 | 0.840363237 | 2.538865412 |
| Q8K284 | Gtf3c1 | 0.83981003 | 2.949906369 |
| M1L9M3 | OPG087 | 0.83364987 | 2.455396259 |
| A0A7H0DN74 | OPG102 | 0.830770969 | 2.371953445 |
| A0A7H0DNC8 | OPG157 | 0.829351322 | 2.282334194 |
| Q8CI43 | Myl6b | 0.828424805 | 3.621386629 |
| Q3SXD3 | Hddc2 | 0.827091214 | 2.441829717 |
| Q6P8I6 | Cox11 | 0.825992561 | 2.408149227 |
| Q9CR26 | Vta1 | 0.823582375 | 2.193020382 |
| A0A7H0DN20 | OPG047 | 0.821669764 | 2.118192647 |
| Q8R088 | Golph3l | 0.820514955 | 2.696681303 |
| A0A7H0DN17 | OPG044 | 0.817021979 | 2.416508747 |
| Q60649 | Clpb | 0.816502847 | 3.426161924 |
| Q9D2M8 | Ube2v2 | 0.816119383 | 2.334217837 |
| Q8K2F8 | Lsm14a | 0.814968108 | 3.249256393 |
| Q8BFY6 | Pef1 | 0.810881581 | 2.721710672 |
| Q9D483 | Polr3c | 0.810184841 | 3.481282279 |
| P61082 | Ube2m | 0.809665894 | 2.687128094 |
| Q8C569 | Fam118b | 0.80606918 | 3.649179786 |
| Q3U0B3 | Dhrs11 | 0.801462057 | 2.086194492 |
| P70697 | Urod | 0.799864958 | 3.308919843 |
| Q8R3H9 | Ttc4 | 0.799758384 | 3.2942389 |
| Q9CQ36 | Pole4 | 0.794759629 | 2.727803932 |
| A2AKG8 | Focad | 0.792863269 | 4.375749023 |
| A0A7H0DN43 | OPG070 | 0.792835592 | 2.308449 |
| O89112 | Lancl1 | 0.792063264 | 2.521020956 |
| Q71RI9 | Kyat3 | 0.789643923 | 3.271383934 |
| Q9JI19 | Fibp | 0.787401779 | 2.714697819 |
| O35613 | Daxx | 0.783180139 | 2.964860861 |
| Q9WUD8 | Faim | 0.778985774 | 2.560390229 |
| Q922W5 | Pycr1 | 0.778442512 | 2.804591016 |
| Q91Z53 | Grhpr | 0.767252948 | 2.783332927 |
| Q9D6F9 | OPG117 | 0.766473461 | 3.737586396 |
| A0A7H0DN89 | Mcmbp | 0.764902392 | 2.896216874 |
| Q8R3C0 | OPG112 | 0.764691362 | 4.355577622 |
| A0A7H0DN84 | Scaf4 | 0.762713124 | 3.202141513 |
| Q7TSH6 | Uba5 | 0.762338974 | 2.862815487 |
| Q8VE47 | OPG133 | 0.761269395 | 4.289966041 |
| A0A7H0DNA5 | Cdk5 | 0.755515226 | 2.578701638 |
| P49615 | OPG088 | 0.754401652 | 3.326609431 |
| M1L502 | Vti1a | 0.753255565 | 3.370961872 |
| O89116 | OPG073 | 0.751706535 | 2.782746742 |
| A0A7H0DN46 | Ocrl | 0.749936059 | 2.81401318 |
| Q6NVF0 | Rad1 | 0.748708062 | 2.788040772 |
| Q9QWZ1 | OPG056 | 0.748173565 | 2.458892903 |
| A0A7H0DN29 | OPG102 | 0.741080077 | 2.68840453 |
| O35381;Q9EST5;Q64G17 | Anp32 | 0.732247242 | 2.814903357 |
| P23506 | Pcmt1 | 0.731603441 | 3.282091937 |
| P07742 | Vps33b | 0.729578164 | 3.518999364 |
| P59016 | Cux1 | 0.722664178 | 2.433910954 |
| P70403 | Rcor3 | 0.720884907 | 2.168397241 |
| Q6PGA0 | Syap1 | 0.715872215 | 2.265427007 |
| Q9D5V6 | Gne | 0.714522789 | 3.248067218 |
| Q91WG8 | Rnpep | 0.71381627 | 3.389660107 |
| Q8VCT3 | Cbx8 | 0.710490238 | 3.724672543 |
| Q9QXV1 | Arih2 | 0.709650604 | 2.794992087 |
| Q9Z1K6 | Idh1 | 0.706257302 | 2.859419819 |
| O88844 | Hdgfl3 | 0.704754728 | 3.532493065 |
| Q9JMG7 | Cpne3 | 0.700714005 | 2.464754478 |
| Q8BT60 | OPG125 | 0.699158305 | 2.68782804 |
| A0A7H0DN97 | Aldh9a1 | 0.69534777 | 3.060071879 |
| Q9JLJ2 | OPG109 | 0.69059579 | 2.86922444 |
| A0A7H0DN81 | OPG123 | 0.689825146 | 3.199205131 |
| P52482 | Prdx6 | 0.689286259 | 2.941785995 |
| A0A7H0DN95 | Pithd1 | 0.687193497 | 3.409255231 |
| O08709 | Ap1g1 | 0.68579756 | 3.163411674 |
| Q8BWR2 | OPG160 | 0.685554515 | 2.816564262 |
| P22892 | Ipo8 | 0.685346304 | 3.518814068 |
| A0A7H0DND1 | Ppp5c | 0.684113411 | 2.449125162 |
| Q7TMY7 | Psme3 | 0.683842823 | 3.479132714 |
| Q60676 | Paics | 0.683162722 | 3.548281161 |
| P61290 | Trmt10c | 0.682006662 | 4.728136463 |
| Q9DCL9 | Ddx17 | 0.680406826 | 4.146679376 |
| Q3UFY8 | Tia1 | 0.678637008 | 2.99217068 |
| Q501J6 | Pold3 | 0.676286309 | 4.65757873 |
| P52912 | Glmn | 0.675025428 | 3.41069737 |
| Q9EQ28 | Pcmt1 | 0.670832902 | 2.541778891 |
| Q8BZM1 | Vps33b | 0.666149844 | 3.571105141 |
| P27659;E9PWZ3 | Rpl3 | 0.664779984 | 2.568007313 |
| Q571H0 | Urb1 | 0.664520526 | 2.025391364 |
| Q924Z6 | Xpo6 | 0.662902056 | 2.253402406 |
| Q78JW9 | Ubfd1 | 0.662836534 | 4.682064338 |
| Q80TV8;Q8BRT1 | Clasp | 0.652230002 | 2.282442769 |
| P50136 | Bckdha | 0.649329728 | 2.145509204 |
| Q3ULD5 | Mccc2 | 0.645723385 | 2.415339234 |
| Q99LC8 | Eif2b1 | 0.645632438 | 2.867136516 |
| Q9CXG3 | Ppil4 | 0.643745913 | 4.748067278 |
| O35638 | Stag2 | 0.640951246 | 3.093163664 |
| Q9CXS4 | Cenpv | 0.640523729 | 2.615605779 |
| M1LLA2 | OPG126 | 0.639814237 | 3.525645116 |
| Q9CWS0 | Ddah1 | 0.639197373 | 2.821995971 |
| P60710;P63260;P68033;P68134 | Act | 0.638100054 | 2.104511142 |
| Q91WQ3 | Yars1 | 0.636897758 | 3.797660463 |
| Q8R2R9;Q9JKC8 | Ap3m | 0.633277713 | 2.464754478 |
| Q9Z2I0 | Letm1 | 0.631287323 | 2.093413636 |
| P0DTN4 | OPG035 | 0.630497156 | 3.954965996 |
| Q8K327 | Champ1 | 0.626006393 | 2.57357525 |
| Q921X6 | Polr3f | 0.625448959 | 2.899299599 |
| O35730 | Ring1 | 0.623205208 | 2.806190992 |
| P53811 | Pitpnb | 0.62315327 | 3.610206607 |
| Q60668 | Hnrnpd | 0.618380338 | 2.964945088 |
| Q8VHL1 | Setd7 | 0.616244897 | 2.644845442 |
| Q3U0K8 | Ogfod1 | 0.614211684 | 2.285001572 |
| Q3UGS4 | Mcrip1 | 0.610358897 | 2.003039395 |
| Q8R418 | Dicer1 | 0.60868343 | 3.601456746 |
| Q9JI11 | Stk4 | 0.604824724 | 2.690518564 |
| Q9D6T0 | Nosip | 0.604532746 | 2.759692035 |
| Q8BFR5 | Tufm | 0.602968979 | 2.168567578 |
| A0A7H0DN79 | OPG107 | 0.599879024 | 2.63362868 |
| Q8BIJ7 | Rufy1 | 0.599063961 | 4.047895433 |
| Q9DBS5 | Klc4 | 0.598228643 | 2.607798939 |
| P60487 | Pdxp | 0.597433016 | 2.755924323 |
| Q9D1P4 | Chorc1 | 0.596720083 | 3.033582657 |
| Q9DBF1 | Aldh7a1 | 0.596090836 | 2.892161268 |
| A2AUM9;Q62036 | Cep152;Cep131 | 0.595560519 | 2.838631729 |
| Q99LX0 | Park7 | 0.59464231 | 3.444039343 |
| Q6ZPJ3 | Ube2o | 0.594357494 | 3.585351237 |
| O08663 | Metap2 | 0.593128442 | 3.095744381 |
| P42225 | Stat1 | 0.592784264 | 2.33847112 |
| O88513 | Gmnn | 0.589594433 | 3.007196706 |
| Q6PAR5 | Gapvd1 | 0.587876986 | 3.458938845 |
| Q78ZA7 | Nap1l4 | 0.587038292 | 2.859999848 |
| Q9Z0E0 | Ncdn | 0.586370948 | 3.405424105 |
| A0A7H0DN96 | OPG124 | 0.585281094 | 3.06876465 |
| Q8BUR4 | Dock1 | -0.585921331 | 2.569590768 |
| Q8BU88 | Mrpl22 | -0.586478714 | 2.442298596 |
| P62627 | Dynlrb1 | -0.587307107 | 2.203480818 |
| Q99JF8 | Psip1 | -0.587482549 | 2.397847369 |
| P84091 | Ap2m1 | -0.587916513 | 3.630710148 |
| Q91YR7 | Prpf6 | -0.588226615 | 3.531831991 |
| Q9JHU4 | Dync1h1 | -0.58873378 | 3.389345991 |
| Q9D0N7 | Chaf1b | -0.589152387 | 2.94292198 |
| Q9D8B3 | Chmp4b | -0.589359004 | 3.82700091 |
| Q61188 | Ezh2 | -0.590717993 | 2.94292198 |
| Q8BHL5 | Elmo2 | -0.590868599 | 3.075550835 |
| Q8BML9 | Qars1 | -0.591348382 | 2.506110379 |
| Q9CXW4 | Rpl11 | -0.592631743 | 2.889301775 |
| O08638;Q61879;Q6URW6;Q8VDD5 | Myh | -0.593182399 | 2.09507904 |
| Q6ZQK5 | Acap2 | -0.593305343 | 4.071150216 |
| Q6ZQ93 | Usp34 | -0.593839267 | 2.805698535 |
| Q3UZ01 | Rnpc3 | -0.594190344 | 2.365518988 |
| A0A7H0DN73 | OPG101 | -0.59420518 | 3.769260716 |
| P42669 | Pura | -0.594519389 | 2.022830781 |
| P69566 | Ranbp9 | -0.594980537 | 3.664720867 |
| Q9Z1M8 | Ik | -0.595046524 | 2.25950994 |
| Q99M87 | Dnaja3 | -0.595406638 | 3.126998588 |
| P63154 | Crnkl1 | -0.595705292 | 4.146679376 |
| Q9JKF7 | Mrpl39 | -0.59578216 | 2.132116177 |
| P60335 | Pcbp1 | -0.597400789 | 3.502597915 |
| Q99KH8;Q9Z2W1 | Stk2 | -0.597595503 | 2.116927462 |
| Q64737 | Gart | -0.598817094 | 2.975971524 |
| P39447 | Tjp1 | -0.60066551 | 3.192449852 |
| Q91YP0 | L2hgdh | -0.601237162 | 2.713200525 |
| Q923D5 | Wbp11 | -0.601675698 | 2.641849541 |
| Q8BL66 | Eea1 | -0.601997304 | 3.201646269 |
| Q9D0D3 | Mtpap | -0.602018503 | 2.329728407 |
| Q8BVF2 | Pdcl3 | -0.602932206 | 2.766993347 |
| Q922H9 | Znf330 | -0.603228188 | 2.484339782 |
| O88712 | Ctbp1 | -0.603354749 | 3.170821894 |
| P05132 | Prkaca | -0.603519983 | 2.613564548 |
| Q8BXC6 | Commd2 | -0.604228548 | 2.581737222 |
| P60867 | Rps20 | -0.60518832 | 2.532456069 |
| Q99JY0 | Hadhb | -0.605763391 | 2.310358194 |
| Q9Z2A0 | Pdpk1 | -0.606049114 | 2.789848482 |
| O88587 | Comt | -0.606702846 | 3.051114047 |
| Q80Y17 | Llgl1 | -0.606712405 | 5.029275318 |
| Q9D9Z1 | Knstrn | -0.606758929 | 3.173768604 |
| Q9CZW5 | Tomm70 | -0.607004835 | 3.093163664 |
| Q99JW4 | Lims1 | -0.607023488 | 3.073620454 |
| P67778 | Phb1 | -0.607257169 | 2.015376571 |
| P53995 | Anapc1 | -0.60772483 | 2.780326516 |
| Q922U1 | Prpf3 | -0.608280079 | 2.580618701 |
| Q9QZ73 | Dcun1d1 | -0.608448856 | 2.131153879 |
| Q9JJG9 | Noa1 | -0.609532802 | 2.300766078 |
| Q8BJW6 | Eif2a | -0.609928621 | 2.639245147 |
| Q8BY71 | Hat1 | -0.610069614 | 3.671001717 |
| Q8BG17 | Nol12 | -0.610186766 | 2.445670076 |
| P59328 | Wdhd1 | -0.612115007 | 2.528629925 |
| Q8BZR9 | Ncbp3 | -0.612482191 | 2.188684889 |
| O55128 | Sap18 | -0.612539816 | 3.409818611 |
| P57780 | Actn4 | -0.613235374 | 3.098190734 |
| P62746;Q62159;Q9QUI0 | Rho | -0.613570254 | 2.673506622 |
| O70310 | Nmt1 | -0.614221687 | 3.525645116 |
| Q8BU31 | Rap2c | -0.614943538 | 2.481159532 |
| Q6PGB8;Q91ZW3 | Smarca | -0.615371034 | 2.973616699 |
| Q6NS46 | Pdcd11 | -0.616047544 | 3.069439574 |
| Q80U93 | Nup214 | -0.616812147 | 2.370467396 |
| P40336 | Vps26a | -0.618106389 | 2.930539457 |
| Q9EP97 | Senp3 | -0.618303735 | 3.409818611 |
| O09012 | Pex5 | -0.618437412 | 3.113400012 |
| Q6WKZ8 | Ubr2 | -0.618759933 | 3.819494354 |
| Q80X82 | Sympk | -0.61901726 | 2.624820406 |
| P51410 | Rpl9 | -0.619531719 | 3.190864341 |
| Q8CFQ3 | Aqr | -0.619636538 | 2.862815487 |
| P26350 | Ptma | -0.619706284 | 3.366472488 |
| P56873 | Znrd2 | -0.620001117 | 2.568720434 |
| P70297 | Stam | -0.620286833 | 3.825137647 |
| Q9ERU9 | Ranbp2 | -0.62110185 | 2.739332546 |
| P61161 | Actr2 | -0.621627404 | 4.066986485 |
| Q62311 | Taf6 | -0.622818795 | 4.067384495 |
| Q8CFI0 | Nedd4l | -0.622945805 | 2.839599271 |
| Q8R3L2 | Tcf25 | -0.623228633 | 2.938185144 |
| Q69ZZ9;Q9QXZ0 | Macf1 | -0.623233764 | 2.779973797 |
| Q6PCL9 | Papolg | -0.623556484 | 2.266011947 |
| P62245 | Rps15a | -0.624418426 | 2.153806708 |
| Q3U487 | Hectd3 | -0.624642502 | 3.249115164 |
| Q9JLQ0 | Cd2ap | -0.626367472 | 2.375181399 |
| Q91WN1 | Dnajc9 | -0.62672979 | 2.105942402 |
| P50516 | Atp6v1a | -0.627126454 | 3.961863085 |
| P46061 | Rangap1 | -0.627511689 | 4.504982838 |
| Q5EG47 | Prkaa1 | -0.627646187 | 2.108109386 |
| P21279 | Gnaq | -0.627716342 | 2.16162239 |
| Q60838 | Dvl2 | -0.628035321 | 2.729387407 |
| P59997 | Kdm2a | -0.628793891 | 3.190864341 |
| Q9D6J6 | Ndufv2 | -0.628886195 | 2.695365468 |
| Q9Z2N8 | Actl6a | -0.628913227 | 4.370906514 |
| Q99MN1 | Kars1 | -0.629373863 | 3.45079281 |
| Q3URQ0 | Tex10 | -0.630195021 | 2.881258881 |
| Q6A028 | Swap70 | -0.633479775 | 2.538926105 |
| Q8BH24 | Tm9sf4 | -0.633521605 | 2.298108872 |
| P35979 | Rpl12 | -0.633902257 | 3.453027624 |
| Q9ES28 | Arhgef7 | -0.63474375 | 2.408196209 |
| Q8VDR9 | Dock6 | -0.635135285 | 2.956310666 |
| Q8R1A4 | Dock7 | -0.636327528 | 2.855853664 |
| Q9WUB4 | Dctn6 | -0.637567688 | 2.160812749 |
| Q8BG81 | Poldip3 | -0.637650553 | 3.312843693 |
| Q99LI8 | Hgs | -0.638628012 | 3.568013035 |
| Q4FK66 | Prpf38a | -0.639358901 | 3.117089736 |
| Q8R317;Q9QZM0 | Ubqln | -0.640854809 | 2.155259866 |
| P52825 | Cpt2 | -0.641546818 | 2.231476653 |
| P70182 | Pip5k1a | -0.642003831 | 2.715583862 |
| Q6NZL6 | Tonsl | -0.643040227 | 2.371826232 |
| Q8BHJ5;Q9QXE7 | Tbl1x | -0.64334624 | 2.357028205 |
| Q9JLT4;Q9JMH6 | Txnrd | -0.643938964 | 3.58429209 |
| Q8K2Q0 | Commd9 | -0.644159629 | 2.399764025 |
| Q71LX4 | Tln2 | -0.645208378 | 2.176762317 |
| P28658 | Atxn10 | -0.645350459 | 3.073620454 |
| Q9CQA5 | Med4 | -0.645857294 | 3.108466315 |
| P59326;Q91YT7 | Ythdf | -0.646666357 | 2.151497626 |
| Q3UMC0 | Afg2a | -0.64835649 | 4.375749023 |
| P70336 | Rock2 | -0.648415009 | 3.511262909 |
| Q80VQ1 | Lrrc1 | -0.648811345 | 2.058979311 |
| Q8CC88 | Vwa8 | -0.648994774 | 2.16162239 |
| Q9CQ48 | Nudcd2 | -0.649385653 | 2.861053963 |
| O35493;P22518 | Clk4;Clk1 | -0.649868639 | 2.589781589 |
| Q6P9P6 | Kif11 | -0.649965643 | 2.900650697 |
| Q9WUP7 | Uchl5 | -0.650372302 | 2.668965216 |
| P42208 | Septin2 | -0.650377383 | 3.791968151 |
| Q8VE65 | Taf12 | -0.650390989 | 4.038227053 |
| A2RSJ4 | Bltp3b | -0.651562585 | 2.669660409 |
| Q9CYA6 | Zcchc8 | -0.652005928 | 4.014518561 |
| Q8R0G9 | Nup133 | -0.652371281 | 2.809307074 |
| Q6P5C5 | Smug1 | -0.653009251 | 2.737802089 |
| P97473 | Tarbp2 | -0.654787529 | 2.560593219 |
| O88487 | Dync1i2 | -0.65544432 | 3.327654332 |
| P50518;Q9D593 | Atp6v1e | -0.656124395 | 2.548119973 |
| P62814;Q91YH6 | Atp61vb | -0.658688588 | 3.564244288 |
| Q9JIX0 | Eny2 | -0.659006722 | 2.977710599 |
| O54946;Q9QYI5 | Dnajb | -0.659498641 | 2.607798939 |
| P28271 | Aco1 | -0.65978267 | 3.720374973 |
| Q9CZM2 | Rpl15 | -0.660420429 | 3.302835901 |
| P62484;Q8CBW3 | Abi | -0.660473265 | 3.528876855 |
| O09106;P70288 | Hdac | -0.661338753 | 2.784403975 |
| Q7TQK1 | Ints7 | -0.661362146 | 2.806746896 |
| P27659 | Nup58 | -0.661622341 | 4.380486543 |
| Q8R332 | Dctn5 | -0.661813149 | 2.993675976 |
| Q9QZB9 | Fntb | -0.661852015 | 2.527613829 |
| Q8K2I1 | Cdk9 | -0.66189501 | 2.381370037 |
| Q99J95 | Snrnp40 | -0.662583849 | 2.893378644 |
| Q6PE01 | Polr1a | -0.662894035 | 3.851287414 |
| O35134 | Aars1 | -0.662979723 | 4.460956923 |
| Q8BGQ7 | Nt5c3b | -0.663316918 | 3.451032441 |
| Q3UFY7 | Med18 | -0.663920539 | 3.408852529 |
| Q9CZ82 | Grm1 | -0.664365412 | 2.096452441 |
| P97772 | Cad | -0.664713957 | 2.058068633 |
| B2RQC6 | Ddx5 | -0.664731595 | 3.282513529 |
| Q61656 | Rad18 | -0.665565464 | 2.766993347 |
| Q9QXK2 | Cops7a | -0.665916877 | 2.13485549 |
| Q9CZ04 | Tfip11 | -0.666050897 | 3.216924547 |
| Q9ERA6 | Ints7 | -0.669129358 | 2.979866261 |
| P62715;P63330 | Ppp2c | -0.669494081 | 3.010311365 |
| P83940 | Eloc | -0.671090178 | 4.883945468 |
| Q91Z96 | Bmp2k | -0.673036291 | 2.467396305 |
| Q8K4P0 | Wdr33 | -0.67324632 | 2.147841467 |
| Q9D358 | Acp1 | -0.676647462 | 2.939719841 |
| Q6ZQK0 | Ncapd3 | -0.677645808 | 2.066817633 |
| O55106 | Strn | -0.679712937 | 3.191697306 |
| Q8C0E2 | Vps26b | -0.681443054 | 4.797847856 |
| Q5SNZ0 | Ccdc88a | -0.681701772 | 3.448504382 |
| P70333 | Hnrnph2 | -0.682241165 | 2.936061684 |
| Q9DCD2 | Xab2 | -0.6832231 | 4.221908298 |
| Q9CRB9 | Chchd3 | -0.683794543 | 2.310358194 |
| O88630 | Gosr1 | -0.683978032 | 2.178504997 |
| P63085 | Mapk1 | -0.684507175 | 3.469268937 |
| P70429 | Evl | -0.684727965 | 2.053734392 |
| Q3UHX2 | Pdap1 | -0.684995706 | 2.937571117 |
| Q5KU39 | Vps41 | -0.688933763 | 2.515032542 |
| S4R2K0 | Pdf | -0.688969213 | 2.333279465 |
| P51807 | Dynlt1 | -0.689312079 | 2.430673567 |
| Q9JI13 | Utp3 | -0.689841713 | 4.28411153 |
| Q8R2U0 | Seh1l | -0.690056219 | 3.41680066 |
| Q9CQ60 | Pgls | -0.690274364 | 2.373209732 |
| Q3THG9 | Aarsd1 | -0.690332571 | 2.099864419 |
| Q8C079 | Strip1 | -0.690660888 | 2.631576003 |
| Q8BUK6 | Hook3 | -0.690681072 | 3.227484059 |
| Q60769 | Tnfaip3 | -0.692037543 | 2.376486587 |
| P62821 | Rab1a | -0.692520125 | 2.017490165 |
| P04627 | Araf | -0.692964106 | 2.502835994 |
| P26638 | Sars1 | -0.69363749 | 4.077057136 |
| Q8K301 | Ddx52 | -0.693644366 | 3.285476335 |
| Q91WC0 | Setd3 | -0.693945901 | 3.425998962 |
| Q9D0D5 | Gtf2e1 | -0.694003571 | 4.338387509 |
| E9Q7M2 | Tsc22d2 | -0.695000391 | 2.474471661 |
| Q6PGG6 | Gnl3l | -0.695730371 | 3.154400925 |
| Q9D1H7 | Get4 | -0.696162426 | 3.481282279 |
| Q6ZWX6 | Eif2s1 | -0.697593247 | 3.56383271 |
| Q3UL36 | Arglu1 | -0.698506493 | 3.548281161 |
| Q8CHT0 | Aldh4a1 | -0.698648225 | 2.198011327 |
| Q61686 | Cbx5 | -0.699248307 | 2.339810668 |
| Q63810 | Ppp3r1 | -0.69980008 | 2.322250803 |
| O35350 | Capn1 | -0.700783077 | 3.633415352 |
| Q9ESX5 | Dkc1 | -0.701138211 | 2.947172678 |
| Q8K4B0 | Mta1 | -0.701853823 | 3.159174208 |
| A0A7H0DNC2;Q8V4V4 | OPG150 | -0.702293327 | 3.942234794 |
| Q9Z110 | Aldh18a1 | -0.702559191 | 2.84602203 |
| Q9WUM4;Q920M5 | Coro | -0.702683853 | 2.377435311 |
| Q3URD3 | Slmap | -0.703246476 | 2.771116166 |
| Q8K400 | Stxbp5 | -0.704566656 | 2.071715284 |
| Q80U95 | Ube3c | -0.704755894 | 2.696483044 |
| Q9WV80 | Snx1 | -0.70479216 | 3.813954616 |
| Q8CI51 | Pdlim5 | -0.704973154 | 2.642345438 |
| P17918 | Pcna | -0.705675047 | 2.826070899 |
| P56382 | Atp5f1e | -0.705710214 | 2.061042081 |
| Q80V86 | Ints8 | -0.706171617 | 2.953791144 |
| Q99ME9 | Gtpbp4 | -0.706196899 | 2.69007632 |
| Q91XU3 | Pip4k2c | -0.707155484 | 2.947130775 |
| Q9CQD1 | Rab5a | -0.707338442 | 2.823223225 |
| Q61753 | Phgdh | -0.708346647 | 3.698857208 |
| Q9CWU9 | Nup37 | -0.708474306 | 3.059581514 |
| Q9JHS4 | Clpx | -0.708836101 | 3.363017573 |
| Q60605 | Myl6 | -0.709107854 | 2.465090625 |
| Q8K2H2 | Otud6b | -0.709946945 | 2.834799649 |
| Q80TZ9 | Rere | -0.710042023 | 2.305174284 |
| P97470 | Ppp4c | -0.710079433 | 2.839075922 |
| P63005 | Pafah1b1 | -0.71036635 | 4.709620529 |
| Q9JL62 | Gltp | -0.710502253 | 2.111199451 |
| Q99K23 | Ufsp2 | -0.710825392 | 2.554472534 |
| Q8CE33 | Klhl11 | -0.711970892 | 2.017055844 |
| P41209;Q9R1K9 | Cetn | -0.712018072 | 2.138732816 |
| Q3UHX9 | Spout1 | -0.712749173 | 2.245574758 |
| Q99K28 | Arfgap2 | -0.712768381 | 2.400131292 |
| Q9JHR7 | Ide | -0.714235109 | 4.520849775 |
| Q8BWZ3 | Naa25 | -0.714448849 | 3.584289567 |
| Q8C0J2 | Atg16l1 | -0.714514337 | 2.404174138 |
| Q3THK3 | Gtf2f1 | -0.715346199 | 4.899776614 |
| Q9D8Y0 | Efhd2 | -0.716705793 | 2.066322179 |
| Q9ESU6 | Brd4 | -0.716897717 | 3.170236202 |
| Q9D061 | Acbd6 | -0.717081615 | 2.526828034 |
| Q8R480 | Nup85 | -0.717528898 | 3.688554386 |
| Q922X9 | Prmt7 | -0.717638998 | 3.709651557 |
| Q9QZB7 | Actr10 | -0.717862776 | 3.830497484 |
| Q9CRD2 | Emc2 | -0.717913511 | 3.523217974 |
| Q80UK8 | Ints2 | -0.718005444 | 2.692975644 |
| Q9D2U5 | Naa38 | -0.719782662 | 2.431211565 |
| E9Q5C9 | Nolc1 | -0.720257099 | 2.515217771 |
| O35226 | Psmd4 | -0.720528247 | 3.495288432 |
| Q99LN9 | Dohh | -0.720861859 | 2.536023361 |
| Q6PFD9 | Nup98 | -0.721378115 | 4.086979652 |
| Q9DBR7;Q8BG95 | Ppp1r12 | -0.721628047 | 3.223193377 |
| Q78PG9 | Ccdc25 | -0.72166548 | 3.041249319 |
| Q9R190 | Mta2 | -0.721756996 | 3.418453048 |
| Q61035 | Hars1 | -0.724016435 | 3.393290214 |
| Q9Z0N1 | Eif2s3x | -0.725320723 | 3.436794662 |
| Q8BIJ6 | Iars2 | -0.726731168 | 2.183960178 |
| Q8BIZ6 | Snip1 | -0.727261291 | 2.460998441 |
| Q6R891 | Ppp1r9b | -0.727410239 | 3.154947284 |
| Q3TDD9 | Ppp1r21 | -0.727566417 | 2.907852665 |
| Q9EPC1 | Parva | -0.728079474 | 3.525645116 |
| Q9DCN2 | Cyb5r3 | -0.729671188 | 2.216015376 |
| Q8BXQ2 | Pigt | -0.729855173 | 2.02395183 |
| P34152 | Ptk2 | -0.730954269 | 2.233214925 |
| Q9DBS1 | Tmem43 | -0.731054512 | 2.109343874 |
| Q3UE37 | Ube2z | -0.731822057 | 4.638422705 |
| Q9EP72 | Emc7 | -0.73219683 | 2.787041423 |
| P22682 | Cbl | -0.732387717 | 3.159174208 |
| P28652;Q923T9 | Camk2 | -0.732615126 | 2.172329793 |
| O88990;P57780;Q7TPR4;Q9JI91 | Actn | -0.733202188 | 3.170236202 |
| P47802 | Mtx1 | -0.733259026 | 2.446090414 |
| P53996 | Cnbp | -0.733404923 | 3.638277906 |
| Q922S8;Q8C0N1 | Kif2 | -0.734241232 | 3.198309702 |
| Q64727 | Vcl | -0.734819033 | 4.500546036 |
| O70252 | Hmox2 | -0.736148143 | 2.502835994 |
| Q91VJ4 | Stk38 | -0.736675457 | 2.579569481 |
| Q9CYI4 | Luc7l | -0.736844761 | 2.27550844 |
| P30999 | Ctnnd1 | -0.73725117 | 3.294352734 |
| Q9WVE8 | Pacsin2 | -0.737573758 | 3.427994675 |
| Q8BP48 | Metap1 | -0.738336351 | 3.078930263 |
| P31938 | Map2k1 | -0.738361283 | 3.177646571 |
| Q8R059 | Gale | -0.740435076 | 3.190864341 |
| Q8CHT3 | Ints5 | -0.740995657 | 2.81342393 |
| Q9Z1J3 | Nfs1 | -0.741093932 | 3.793850156 |
| Q3TIR1 | Trappc13 | -0.742816443 | 2.671752268 |
| Q569Z6 | Thrap3 | -0.743066755 | 3.899435712 |
| P33215 | Nedd1 | -0.744506128 | 2.944951665 |
| Q9EPU0 | Upf1 | -0.744593818 | 3.483558807 |
| Q99JX7 | Nxf1 | -0.744730716 | 3.005790611 |
| A0A7H0DN73;P04363 | OPG101 | -0.744865207 | 3.732363984 |
| Q9CPR7 | Sike1 | -0.745013525 | 3.28976424 |
| P09411 | Pgk1 | -0.745701745 | 3.274412271 |
| Q60520 | Sin3a | -0.74583768 | 2.550617457 |
| Q9JLC8 | Sacs | -0.745890597 | 2.444099785 |
| Q91W96 | Anapc4 | -0.746107516 | 4.207432459 |
| P62334 | Psmc6 | -0.746373733 | 5.695812265 |
| Q99J62 | Rfc4 | -0.747429708 | 3.769260716 |
| Q8BM72 | Hspa13 | -0.747486528 | 2.595912531 |
| Q8BTZ4 | Anapc5 | -0.747816135 | 2.507162967 |
| Q80XI4 | Pip4k2b | -0.748581079 | 3.051701664 |
| Q68FE6 | Ripor1 | -0.748641153 | 2.192765311 |
| Q9Z2Y8 | Plpbp | -0.74865168 | 2.663769551 |
| P61750 | Arf4 | -0.749962737 | 2.329930184 |
| Q8K5B2 | Mcfd2 | -0.751108786 | 2.370955186 |
| Q922R8 | Pdia6 | -0.751172658 | 2.350442459 |
| Q8BVE3 | Atp6v1h | -0.751962773 | 3.308919843 |
| P28740 | Kif2a | -0.754139614 | 2.358736505 |
| P97434 | Mprip | -0.754747464 | 2.908403409 |
| Q9CR51 | Atp6v1g1 | -0.755801312 | 2.450732767 |
| Q6PGH1 | Bud31 | -0.756337972 | 3.265922864 |
| P20664 | Prim1 | -0.757173556 | 2.126288629 |
| P40142 | Tkt | -0.758313871 | 3.676721531 |
| Q9D051 | Pdhb | -0.758869313 | 3.518814068 |
| Q8BQM4 | Heatr3 | -0.759302004 | 3.641463432 |
| Q9WUA3 | Pfkp | -0.759498594 | 3.899435712 |
| P59325 | Eif5 | -0.75975587 | 3.189864445 |
| O35286 | Dhx15 | -0.760550949 | 3.460925702 |
| Q99JT9 | Adi1 | -0.761633706 | 2.464787551 |
| Q9CQK7 | Rwdd1 | -0.762952455 | 3.114057143 |
| B1AQJ2 | Usp36 | -0.763574508 | 2.604551492 |
| P31230 | Aimp1 | -0.764138102 | 3.500036686 |
| Q8C166 | Cpne1 | -0.764857568 | 2.58842094 |
| Q9WV84 | Nme4 | -0.765099655 | 3.190864341 |
| B2RY56 | Rbm25 | -0.766066332 | 4.066442441 |
| Q9CPT5 | Nop16 | -0.767280019 | 3.063424856 |
| Q6IRU5 | Cltb | -0.767473799 | 3.538507794 |
| Q8BFR4 | Gns | -0.767495761 | 2.579165401 |
| Q99K70 | Rragc | -0.767842266 | 4.971384312 |
| Q8BH74 | Nup107 | -0.768219036 | 3.883740041 |
| Q8VE10 | Naa40 | -0.768228117 | 2.861838547 |
| O09110 | Map2k3 | -0.76824493 | 3.02812628 |
| Q61553 | Fscn1 | -0.768443061 | 4.146679376 |
| O35326 | Srsf5 | -0.768829414 | 3.995737309 |
| Q9CQC8 | Spg21 | -0.769917566 | 3.741281563 |
| Q8C050 | Rps6ka5 | -0.770671991 | 2.171586363 |
| P58404;Q9ERG2 | Stm4 | -0.770803783 | 3.144857264 |
| Q61879 | Slu7 | -0.771212608 | 4.196928254 |
| Q8BHJ9 | Tpr | -0.771649806 | 3.091688467 |
| F6ZDS4 | Msn | -0.771815177 | 4.145136909 |
| P26043 | Rdx | -0.772940627 | 2.848713965 |
| Q99M28 | Rnps1 | -0.77316654 | 3.212102835 |
| O35239 | Ptpn9 | -0.773500489 | 2.912609422 |
| F7BJB9 | Morc3 | -0.77490574 | 3.340631639 |
| P60670 | Nploc4 | -0.775287944 | 3.980014441 |
| Q99L45 | Eif2s2 | -0.775396589 | 3.812738126 |
| Q61024 | Asns | -0.775540981 | 4.455989016 |
| Q3V1V3 | Esf1 | -0.776043968 | 3.729277997 |
| A2AGT5 | Ckap5 | -0.77606226 | 2.031070715 |
| Q921M4 | Golga2 | -0.777400335 | 3.709651557 |
| P35700 | Prdx1 | -0.777961383 | 2.229964157 |
| Q99KE1 | Me2 | -0.778723217 | 2.575510739 |
| Q8VEH3 | Arl8a | -0.779205695 | 2.54179338 |
| P58501 | Paxbp1 | -0.779745108 | 3.25179994 |
| Q5H8C4 | Vps13a | -0.780040002 | 2.400131292 |
| Q9CQL4 | Mrpl20 | -0.780835842 | 2.024915451 |
| P97770 | Thumpd3 | -0.781197726 | 2.765463767 |
| Q9JIY2 | Cbll1 | -0.782336468 | 3.340795523 |
| P35283 | Rab12 | -0.782442041 | 2.026908819 |
| Q9WVR4 | Fxr2 | -0.782775812 | 3.543500057 |
| P18760 | Cfl1 | -0.782785337 | 3.981570786 |
| Q91VY9 | Znf622 | -0.783164863 | 3.332644078 |
| Q924H7 | Wac | -0.783300241 | 2.80155168 |
| Q8BMF4 | Dlat | -0.783351685 | 3.423447522 |
| Q7TSZ8;Q9DCM7 | Nacc | -0.783370223 | 3.056109998 |
| P26231 | Ctnna1 | -0.784649186 | 2.616773913 |
| O70305 | Atxn2 | -0.784703977 | 3.013328665 |
| P42567 | Eps15 | -0.786214642 | 3.749446852 |
| Q9ER00 | Stx12 | -0.788270563 | 2.311480779 |
| A2A6A1 | Gpatch8 | -0.789637379 | 2.537049375 |
| Q9D0S9 | Hint2 | -0.790168438 | 2.177098013 |
| Q9DBJ1 | Pgam1 | -0.79144737 | 3.206648935 |
| Q9CZU6 | Cs | -0.791622005 | 2.543147792 |
| Q8C5N3 | Cwc22 | -0.792532254 | 4.420212838 |
| O35435 | Dhodh | -0.793081339 | 2.110686627 |
| Q9Z1R2 | Bag6 | -0.793592413 | 3.418453048 |
| Q9QZD8 | Slc25a10 | -0.793781315 | 2.390981276 |
| O88848 | Arl6 | -0.794321382 | 2.031803138 |
| Q6PA06 | Atl2 | -0.794540335 | 2.013740726 |
| O70279 | Ess2 | -0.796164406 | 2.508647235 |
| A0A7H0DMZ9 | OPG023 | -0.796664682 | 3.363323548 |
| O55131 | Septin7 | -0.797457135 | 2.349729012 |
| P55264 | Adk | -0.79785652 | 3.638277906 |
| Q52KI8 | Srrm1 | -0.798008492 | 3.828422659 |
| Q8BHN3 | Ganab | -0.798279155 | 2.861053963 |
| Q8BPG6 | Sumf2 | -0.79840479 | 3.070704593 |
| A2AF47 | Dock11 | -0.799014179 | 2.349769081 |
| P97371 | Psme1 | -0.799637839 | 4.341673123 |
| Q8BRN9 | Cc2d1b | -0.800011548 | 3.7075662 |
| Q80YV2 | Zc3hc1 | -0.800336531 | 2.913773963 |
| Q9WV98 | Timm9 | -0.800810705 | 3.847368222 |
| Q8VCD5 | Med17 | -0.800900045 | 2.862815487 |
| P25799 | Nfkb1 | -0.800944677 | 2.937571117 |
| Q99LM9 | Tada1 | -0.801314171 | 2.72476652 |
| Q9WUL7 | Arl3 | -0.801614491 | 2.64370489 |
| P24788 | Cdk11b | -0.801884424 | 4.186900043 |
| P59999 | Arpc4 | -0.802312259 | 3.994663053 |
| Q80YA3 | Ddhd1 | -0.802639909 | 3.917715693 |
| P48754 | Brca1 | -0.803363618 | 2.148271591 |
| Q501J7 | Phactr4 | -0.803550364 | 2.758323834 |
| A8Y5H7 | Sec14l1 | -0.804700559 | 3.199205131 |
| Q61140 | Bcar1 | -0.805407427 | 3.06268404 |
| P53994 | Rab2a | -0.805505879 | 2.213254578 |
| Q2HXL6 | Edem3 | -0.805605336 | 4.000365787 |
| Q7TPD0 | Ints3 | -0.805686589 | 2.720164429 |
| P62331 | Arf6 | -0.806061129 | 3.636575469 |
| Q3TKT4 | Smarca4 | -0.808212977 | 2.59236829 |
| Q6P4S8 | Ints1 | -0.809845117 | 3.635823344 |
| P24369 | Ppib | -0.810290593 | 3.190379089 |
| P68404 | Prkcb | -0.811495817 | 2.86203495 |
| Q9Z1E3 | Nfkbia | -0.811661437 | 2.237221256 |
| P53810 | Pitpna | -0.812012685 | 2.962134462 |
| Q9CPY7 | Lap3 | -0.81332716 | 3.753531572 |
| Q3U821 | Wdr75 | -0.81340206 | 3.199205131 |
| P27612 | Plaa | -0.813877297 | 3.423447522 |
| Q8BW72 | Kdm4a | -0.814366045 | 3.082698915 |
| Q9CXY9 | Pigk | -0.81438551 | 2.257530294 |
| Q9JIK5 | Ddx21 | -0.817283023 | 3.483558807 |
| Q9CS00 | Cactin | -0.817450431 | 2.071715284 |
| Q9ESL4 | Map3k20 | -0.817859054 | 3.344322415 |
| Q7TMI3;Q8VDF2 | Uhrf | -0.819044609 | 2.691560551 |
| P54103 | Dnajc2 | -0.819993627 | 2.785026917 |
| O88643;Q8CIN4 | Pak | -0.820601043 | 3.828422659 |
| P97760 | Polr2c | -0.821266365 | 3.892853995 |
| P26039 | Tln1 | -0.822530338 | 4.007518869 |
| Q8R3V5 | Sh3glb2 | -0.823045581 | 3.322573561 |
| Q8BK67 | Rcc2 | -0.823274473 | 4.000400057 |
| Q9D7N3 | Mrps9 | -0.825147041 | 2.699003068 |
| P61021 | Rab5b | -0.825291558 | 2.683528066 |
| P53702 | Hccs | -0.826602559 | 2.668965216 |
| Q8BIG4 | Fbxo28 | -0.827275906 | 3.392525804 |
| Q8CEC0 | Nup88 | -0.827442095 | 2.498156125 |
| P51175 | Ppox | -0.827513114 | 3.004003469 |
| Q8VHR5 | Gatad2b | -0.828162425 | 4.237499095 |
| Q925E7 | Ppp2r2d | -0.828352131 | 2.867560047 |
| O88895 | Hdac3 | -0.829274842 | 3.978925047 |
| A2AN08 | Ubr4 | -0.829675392 | 3.125748157 |
| Q9QZQ1 | Afdn | -0.831124365 | 3.41336664 |
| O35218 | Cpsf2 | -0.831436971 | 4.50536728 |
| Q8K114 | Ints9 | -0.831786166 | 4.5059213 |
| P40124 | Cap1 | -0.831893256 | 4.146679376 |
| Q0VEJ0 | Cep76 | -0.833563084 | 3.163533454 |
| Q8R2M2 | Dnttip2 | -0.833590145 | 3.946660784 |
| Q91VM3 | Wdr45 | -0.833814099 | 2.096671238 |
| Q8VD62 | Bles03 | -0.835828108 | 3.914955128 |
| P63147;Q9Z255 | Ube2 | -0.836026142 | 2.619336745 |
| P62983 | Rps27a | -0.837027711 | 4.50536728 |
| P97477 | Aurka | -0.837189924 | 2.595450236 |
| Q8CBY8 | Dctn4 | -0.837604564 | 2.560390229 |
| O88967 | Yme1l1 | -0.838205035 | 2.654136147 |
| Q8BUL5 | Klhl7 | -0.838646961 | 2.632429206 |
| Q8BZQ7 | Anapc2 | -0.838845851 | 2.403594757 |
| Q9JHW4 | Eefsec | -0.839084701 | 3.177646571 |
| Q9CYG7 | Tomm34 | -0.839138119 | 2.723118691 |
| Q9CU65 | Zmym2 | -0.839538261 | 3.236128002 |
| Q8VEK6 | Ing3 | -0.839802817 | 2.729387407 |
| Q9DCD0 | Pgd | -0.83985271 | 3.072863147 |
| Q9CYL5 | Glipr2 | -0.840166631 | 2.465965462 |
| P11440 | Cdk1 | -0.840321161 | 3.450039081 |
| P56399 | Usp5 | -0.842453946 | 4.373790724 |
| P62073 | Timm10 | -0.842616618 | 3.023656822 |
| Q8CG48 | Smc2 | -0.843530543 | 4.599565553 |
| Q8BY87 | Usp47 | -0.846568961 | 3.563777056 |
| O55222 | Ilk | -0.84697683 | 3.946660784 |
| Q8C1A5 | Thop1 | -0.847068161 | 3.984452126 |
| Q91XD2 | Lims2 | -0.847779097 | 2.737260358 |
| Q9D7M1 | Gid8 | -0.847807464 | 3.267845897 |
| Q05CL8 | Larp7 | -0.848411306 | 2.944476753 |
| Q9DB96 | Ngdn | -0.848687724 | 3.130165145 |
| Q9JMH6 | Rsrc1 | -0.849425906 | 3.502762463 |
| Q9DBU6 | Foxp1 | -0.849476052 | 2.275467981 |
| P58462 | Epn2 | -0.849980689 | 2.207267155 |
| P62814 | Isy1 | -0.852486289 | 4.057208694 |
| Q8CHU3 | Fmr1 | -0.852958215 | 2.558508014 |
| Q69ZQ2 | Wdr55 | -0.853651541 | 3.570770227 |
| P35922 | Rbm5 | -0.854789989 | 3.667783416 |
| Q9CX97 | Akap10 | -0.855164094 | 2.707609543 |
| Q91YE7 | Mnat1 | -0.855808717 | 3.393290214 |
| O88845 | Gnb1 | -0.856582775 | 2.973923598 |
| P51949 | Pmm2 | -0.856599914 | 4.114545872 |
| P62874 | Rrp7a | -0.856636973 | 2.960131368 |
| Q9Z2M7 | Hk1 | -0.856655174 | 3.564244288 |
| Q9D1C9 | Tbcel | -0.857518232 | 5.104472624 |
| P17710 | Lrrc59 | -0.857860467 | 4.420212838 |
| Q8C5W3 | Hnrnpf | -0.859415344 | 3.869108017 |
| Q922Q8 | Nherf2 | -0.859645929 | 2.692919997 |
| Q9Z2X1 | Crlf1 | -0.859653818 | 2.49480855 |
| Q9JHL1 | Ptges2 | -0.860788762 | 2.69129318 |
| Q9JM58 | Gid8 | -0.861010004 | 2.153094537 |
| Q8BWM0 | Larp7 | -0.86166931 | 2.103887614 |
| P70236 | Map2k6 | -0.861682273 | 3.612529958 |
| Q8VCX5 | Micu1 | -0.862627789 | 2.155259866 |
| Q91YN1 | Fam118a | -0.862834436 | 2.794681001 |
| Q8R2N2 | Utp4 | -0.862998139 | 2.948429486 |
| Q9WTL7 | Lypla2 | -0.86311619 | 3.867651428 |
| Q91YU6 | Lzts2 | -0.864970559 | 2.521000194 |
| Q8VCY6 | Utp6 | -0.865212354 | 3.599403769 |
| Q8CDG3 | Vcpip1 | -0.865908661 | 3.706017463 |
| O08749 | Dld | -0.866282769 | 3.546072554 |
| Q8C547 | Heatr5b | -0.868567222 | 3.227484059 |
| Q91WM3 | Rrp9 | -0.868934121 | 3.327491975 |
| Q64516 | Gk | -0.869744549 | 2.906564399 |
| Q8VI33;Q6NZA9 | Taf9 | -0.870847215 | 3.230954519 |
| Q3UM18 | Lsg1 | -0.871317473 | 4.240819544 |
| Q05306 | Col10a1 | -0.871443041 | 2.258543949 |
| Q8BHT6 | B3glct | -0.871857367 | 2.946133554 |
| Q9D0V8 | Cinp | -0.873170027 | 2.63136356 |
| Q3TCH7 | Cul4a | -0.875386712 | 4.169079325 |
| Q9EQH2 | Erap1 | -0.875432971 | 2.831810871 |
| P62869 | Elob | -0.875867608 | 3.984452126 |
| Q9D1C1 | Ube2c | -0.875893937 | 3.692526592 |
| P47757 | Capzb | -0.876518243 | 4.495282144 |
| Q9CQA1 | Trappc5 | -0.877159462 | 2.771116166 |
| Q5HZJ0 | Drosha | -0.877699266 | 2.696483044 |
| P63038 | Hspd1 | -0.87880333 | 3.224087602 |
| B1AVZ0 | Uprt | -0.879609814 | 2.218087786 |
| O88271 | Cfdp1 | -0.881517262 | 3.114596801 |
| G5E8V9 | Arfip1 | -0.881530476 | 3.308074587 |
| Q8BP47 | Nars1 | -0.882248954 | 4.495282144 |
| Q9QY76 | Vapb | -0.882967714 | 3.212680824 |
| O54941 | Smarce1 | -0.883200492 | 4.450920196 |
| Q99MR8 | Mccc1 | -0.883812617 | 3.326609431 |
| Q9EQC5 | Scyl1 | -0.884236928 | 2.926513144 |
| Q8R0J7 | Vps37b | -0.88458248 | 3.731113495 |
| A0A7H0DN04 | OPG030 | -0.884592588 | 4.552426484 |
| P30415 | Nktr | -0.884968227 | 2.938185144 |
| Q9D0F3 | Lman1 | -0.885272075 | 2.486072083 |
| Q9R0B9 | Plod2 | -0.885840324 | 2.942952096 |
| P10126 | Eef1a1 | -0.88622456 | 4.42846902 |
| O54988 | Slk | -0.88677598 | 3.223650072 |
| P70349 | Hint1 | -0.886786714 | 2.883195616 |
| Q60775 | Elf1 | -0.886892916 | 2.359655349 |
| P63017 | Hspa8 | -0.888320287 | 4.421479989 |
| Q60865 | Caprin1 | -0.888409599 | 3.706017463 |
| Q921N6 | Ddx27 | -0.889362154 | 2.72446142 |
| Q8R060 | Zwilch | -0.891608644 | 4.289311731 |
| Q9QZH3 | Ppie | -0.892026582 | 2.131153879 |
| Q6PHZ5 | Rbm15b | -0.89318424 | 2.322625547 |
| O88456 | Capns1 | -0.896402189 | 3.073025804 |
| P43346 | Dck | -0.897392398 | 2.029465712 |
| Q9JLV6 | Pnkp | -0.897551328 | 3.998412163 |
| Q9CSN1 | Snw1 | -0.898415527 | 3.978014205 |
| P47809 | Map2k4 | -0.898907522 | 3.768338095 |
| Q00PI9 | Hnrnpul2 | -0.89895105 | 3.073315138 |
| Q9Z0R4;Q9Z0R6 | Itsn | -0.900253599 | 2.044781506 |
| A2RSX7 | Tyw5 | -0.901875638 | 2.266882684 |
| Q6PNC0 | Dmxl1 | -0.903236058 | 2.500491225 |
| Q9ESV0 | Ddx24 | -0.903740968 | 3.087977907 |
| Q99JB8 | Pacsin3 | -0.904667984 | 3.039873912 |
| P46467 | Vps4b | -0.905318291 | 3.678336612 |
| Q64012 | Raly | -0.905624433 | 2.627639944 |
| Q9D4C5 | Eaf1 | -0.907289466 | 3.106116846 |
| Q921C5 | Bicd2 | -0.907323804 | 3.284104295 |
| O55023 | Impa1 | -0.907336139 | 2.081824106 |
| Q99K01 | Pdxdc1 | -0.907772946 | 4.023016918 |
| Q8BH48 | Ubap1 | -0.908728269 | 3.056109998 |
| Q5FWK3 | Arhgap1 | -0.908931635 | 3.867651428 |
| Q6ZQ03 | Fnbp4 | -0.909260164 | 3.883740041 |
| P46938 | Yap1 | -0.911789785 | 3.570664676 |
| Q9DBD5 | Pelp1 | -0.911953035 | 3.957686009 |
| P17156 | Hspa | -0.912254059 | 2.140484176 |
| P63094;Q6R0H7 | Gnas | -0.912500699 | 2.381370037 |
| P10711 | Tcea1 | -0.912885275 | 4.420212838 |
| Q9JMA1 | Usp14 | -0.913140789 | 4.183422758 |
| P30051;P48301;P70210;Q62296 | Tead | -0.913787557 | 3.346240978 |
| P47754;P47753 | Capza | -0.914813731 | 2.358736505 |
| Q9Z0H1 | Wdr46 | -0.915211964 | 2.872434173 |
| Q61543 | Glg1 | -0.915476942 | 3.476267802 |
| Q9CQA3 | Sdhb | -0.916645272 | 2.820452409 |
| Q9WTK2 | Cdyl | -0.916921452 | 2.962525012 |
| Q80T69 | Rsbn1 | -0.917275431 | 2.11010434 |
| Q9D2H6 | Sp2 | -0.917994863 | 2.018609579 |
| Q9Z321 | Top3b | -0.91802557 | 2.940120245 |
| Q9R0H0 | Acox1 | -0.918849713 | 2.58712405 |
| Q9D4D4 | Tktl2 | -0.920612157 | 2.290121265 |
| Q9ET26 | Rnf114 | -0.921041366 | 2.217827765 |
| P62077 | Timm8b | -0.921615252 | 3.405424105 |
| Q8CHH9 | Septin8 | -0.922228781 | 3.778608262 |
| Q9QX47 | Son | -0.922524849 | 3.837753622 |
| Q7TNP2 | Ppp2r1b | -0.922685032 | 5.073302231 |
| P18653;P18654;Q9WUT3 | Rps6ka | -0.923478753 | 2.297769828 |
| Q6NZM9 | Hdac4 | -0.923870023 | 2.053008858 |
| Q6PAV2 | Herc4 | -0.924612021 | 3.464843055 |
| Q62351 | Tfrc | -0.925580498 | 2.413195735 |
| Q9WVF7 | Pole | -0.925749346 | 3.583446204 |
| Q9JKX6 | Nudt5 | -0.926092421 | 2.631780739 |
| P49138;Q3UMW7 | Mapkapk | -0.926776169 | 2.535621364 |
| Q8K2A7 | Ints10 | -0.92936436 | 3.597547394 |
| Q6PAC3 | Dcaf13 | -0.929560535 | 2.656749287 |
| Q3V3R1 | Mthfd1l | -0.930476708 | 3.602282682 |
| Q9D824 | Fip1l1 | -0.931854557 | 4.398916716 |
| Q9D0I8 | Mrto4 | -0.932146941 | 2.003039395 |
| O54931 | Pakap | -0.93445396 | 2.917882694 |
| Q99KX1 | Mlf2 | -0.934567385 | 4.182525814 |
| Q05BC3 | Eml1 | -0.937118823 | 2.408629108 |
| Q9JIX8 | Acin1 | -0.937440424 | 3.919916677 |
| Q61425 | Hadh | -0.937621276 | 2.950403017 |
| Q8K2C7 | Os9 | -0.940253258 | 3.032916515 |
| O35344 | Kpna3 | -0.940606415 | 3.244695414 |
| Q91Y86 | Mapk8 | -0.940784839 | 2.909825426 |
| Q8C1B7 | Mlf2 | -0.941963604 | 4.345852956 |
| Q6P1F6;Q8BG02 | Ppp2r2 | -0.942574789 | 2.462958639 |
| Q69ZA1 | Cdk13 | -0.942660242 | 5.139550781 |
| Q80V53 | Chst14 | -0.943087564 | 2.01395517 |
| Q6A026 | Pds5a | -0.943687863 | 3.257235115 |
| Q9CQW2;Q8VEH3 | Arl8b | -0.943923082 | 2.883195616 |
| Q922P9 | Glyr1 | -0.94475722 | 4.262042568 |
| Q8BP92 | Rcn2 | -0.946527407 | 3.971565037 |
| Q9CZ44 | Nsfl1c | -0.946681231 | 4.192211703 |
| Q6PDG5 | Smarcc2 | -0.948905376 | 3.727430683 |
| Q9Z0R6 | Pcca | -0.949486531 | 3.22009771 |
| Q91ZA3 | Hdlbp | -0.949762502 | 2.795917996 |
| Q8VDJ3 | Scfd2 | -0.950050948 | 3.789307581 |
| Q8BTY8 | Rad51 | -0.950241309 | 3.236890527 |
| Q08297 | Clpp | -0.951399705 | 3.15212545 |
| O88696 | Nln | -0.951445352 | 2.926513144 |
| Q91YP2 | Smdt1 | -0.951462766 | 4.43303 |
| Q9DB10 | Inpp5b | -0.952987703 | 2.280469806 |
| Q8K337 | Wdr37 | -0.953095458 | 3.108466315 |
| Q8CBE3 | Dgcr8 | -0.954208414 | 2.279780718 |
| Q9EQM6 | Mrpl28 | -0.955672651 | 4.478568205 |
| Q9D1B9 | Cux1 | -0.957736014 | 3.608367302 |
| P53564 | Gnai3 | -0.957957282 | 3.469498339 |
| Q9DC51 | Cfap298 | -0.958013338 | 2.476526856 |
| Q8BL95 | Trrap | -0.959058298 | 3.619292811 |
| Q80YV3 | Raver1 | -0.959777007 | 3.243765217 |
| Q9CW46 | Ppt1 | -0.96019943 | 4.450920196 |
| Q6PER3 | Atrx | -0.960502265 | 4.044096318 |
| O88531 | Glyr1 | -0.961601947 | 3.128065679 |
| Q61687 | Rcn2 | -0.962283411 | 4.398916716 |
| Q8C1B7;Q9R1T4 | Septin6 | -0.962310757 | 4.424556899 |
| Q922J3 | Clip1 | -0.963117199 | 4.854481453 |
| Q3UZ39 | Lrrfip1 | -0.963220144 | 3.405424105 |
| Q6PDQ2 | Chd4 | -0.963320964 | 3.051589975 |
| Q8R1F9 | Rpp40 | -0.964167825 | 3.231574546 |
| Q9QYB1 | Clic4 | -0.964330692 | 4.684011748 |
| Q9DC23 | Dnajc10 | -0.964746565 | 3.1408269 |
| P58774 | Tpm2 | -0.96593885 | 3.45935844 |
| Q59J78 | Ndufaf2 | -0.966187985 | 3.56332944 |
| Q8CH25 | Sltm | -0.9667247 | 2.458651456 |
| Q8K1Z0 | Coq9 | -0.966874203 | 3.9127205 |
| P24668 | M6pr | -0.968482097 | 2.335364202 |
| Q9D7W5 | Med8 | -0.968682003 | 2.953143619 |
| Q9JM13 | Rabgef1 | -0.968791162 | 2.563255468 |
| Q8BH43 | Wasf2 | -0.968931607 | 2.585179995 |
| Q7TPM1 | Prrc2b | -0.969516686 | 2.687981612 |
| Q9CXU0 | Med10 | -0.970009672 | 3.531309036 |
| Q9EQP2;Q9QXY6 | Ehd | -0.970023237 | 2.018662255 |
| Q3UGR5 | Hdhd2 | -0.970596387 | 2.522985622 |
| Q99020 | Hnrnpab | -0.971863692 | 4.023016918 |
| P33611 | Pola2 | -0.972486077 | 2.801212548 |
| Q8C4B4 | Unc119b | -0.974200685 | 2.288073043 |
| Q3TNH5 | Arb2a | -0.974965461 | 3.514521755 |
| Q9QZ88 | Vps29 | -0.975594884 | 2.965926043 |
| P60764;P63001 | Rac | -0.976623153 | 2.140484176 |
| Q3THE2 | Myl12b | -0.978620705 | 2.718785915 |
| P61358 | Rpl27 | -0.979431877 | 3.569322397 |
| Q8BMJ2 | Lars1 | -0.979844032 | 3.094412447 |
| Q91ZU1 | Asb6 | -0.981050251 | 2.480867158 |
| Q9CR00 | Psmd9 | -0.981842503 | 3.630576021 |
| Q4FZF3 | Ddx49 | -0.985082673 | 3.019047596 |
| P24270 | Cat | -0.985238539 | 3.82123377 |
| P17182 | Eno1 | -0.985371698 | 3.899435712 |
| E9Q634 | Myo1e | -0.985647999 | 3.799512501 |
| Q60710 | Samhd1 | -0.986005775 | 3.965103492 |
| Q8R2N0 | Ccdc59 | -0.986192045 | 2.037501142 |
| Q6ZWS8 | Spop | -0.986616064 | 2.375181399 |
| Q7TMB8 | Cyfip1 | -0.986923026 | 3.847368222 |
| P97798 | Neo1 | -0.987658614 | 2.596294264 |
| P97287 | Mcl1 | -0.988840515 | 2.950403017 |
| Q9CZX0 | Elp3 | -0.989600386 | 2.714697819 |
| Q924H2 | Med15 | -0.990493762 | 3.455729418 |
| Q91ZR2 | Snx18 | -0.990839261 | 2.619171431 |
| Q8BG05 | Hnrnpa3 | -0.990868315 | 4.104706659 |
| Q8BHB4 | Wdr3 | -0.990933205 | 3.620476644 |
| P62737;P63268;P68033;P68134 | Act | -0.991222183 | 2.690518564 |
| Q9Z2X8 | Keap1 | -0.992222813 | 3.606005412 |
| Q921I2 | Klhdc4 | -0.993496053 | 2.090520141 |
| Q9DBG9 | Tax1bp3 | -0.996380355 | 2.892161268 |
| Q99ME2 | Wdr6 | -0.996406297 | 2.634997802 |
| Q8R317;Q99NB8;Q9QZM0 | Ubqln | -0.997400885 | 2.810436278 |
| Q62422 | Ostf1 | -0.999688245 | 3.474031737 |
| Q8CDM8 | Fhip2a | -1.001698244 | 2.485863023 |
| Q8C3X2 | Ccdc90b | -1.002251699 | 2.547903714 |
| Q9JM76 | Arpc3 | -1.002736078 | 4.193143718 |
| Q8CFI7 | Polr2b | -1.003140932 | 3.957686009 |
| O35598 | Adam10 | -1.004064199 | 2.911189271 |
| Q61083 | Map3k2 | -1.006344532 | 2.758323834 |
| O09167 | Rpl21 | -1.009106731 | 2.702512005 |
| Q80YQ2 | Med23 | -1.009213609 | 3.195758038 |
| Q9JIH2 | Nup50 | -1.010113072 | 2.722695991 |
| D3YXK2 | Safb | -1.0103133 | 2.668965216 |
| Q6NZF1 | Zc3h11a | -1.01073075 | 2.458781834 |
| Q9D902 | Gtf2e2 | -1.011001318 | 3.17514254 |
| P70340 | Smad1 | -1.01105258 | 3.626685116 |
| Q91WA1 | Tipin | -1.01139556 | 2.241167729 |
| Q61595 | Ktn1 | -1.012815051 | 2.705529557 |
| Q3TJD7 | Pdlim7 | -1.014099427 | 4.17994215 |
| Q8BXR9;Q9DBS9 | Osbpl | -1.015249131 | 2.39188864 |
| Q99LP6 | Grpel1 | -1.018126213 | 4.412462579 |
| Q8CIM8 | Ints4 | -1.01929258 | 3.684082502 |
| Q9Z1X9 | Cdc45 | -1.01933654 | 2.346948075 |
| Q8BGT5 | Gpt2 | -1.019487692 | 3.461753457 |
| Q9D0M1 | Prpsap1 | -1.020159494 | 3.866200923 |
| P97440 | Slbp | -1.020486321 | 2.680374262 |
| Q9D1K2 | Atp6v1f | -1.020614107 | 2.259840574 |
| Q921F2 | Tardbp | -1.0213965 | 3.786435915 |
| Q8BK08 | Tmem11 | -1.02201112 | 2.581181909 |
| Q61584 | Fxr1 | -1.023260584 | 5.104472624 |
| Q8VDW0 | Ddx39a | -1.024031594 | 4.054244187 |
| Q8VDM4 | Psmd2 | -1.024306615 | 4.784426578 |
| Q8VD75 | Hip1 | -1.025369337 | 2.848449313 |
| Q80XP8 | Fam76b | -1.02563915 | 2.247655763 |
| Q8R4R6 | Nup35 | -1.026703571 | 3.578407085 |
| E9Q5G3 | Kif23 | -1.026811747 | 3.617119599 |
| P62874;P62880 | Gnb | -1.027355324 | 2.607274644 |
| Q99K74 | Med24 | -1.02753655 | 4.142909775 |
| Q9WTV7 | Rlim | -1.027748332 | 2.978783325 |
| P33609 | Pola1 | -1.028392659 | 2.883195616 |
| Q9D1D4 | Tmed10 | -1.028504985 | 2.171569403 |
| Q5EG47 | Cpsf1 | -1.029082651 | 3.228435791 |
| Q9EPU4 | Utp20 | -1.029219678 | 4.327341891 |
| Q5XG71 | Eif6 | -1.029900317 | 3.283807658 |
| O55135 | Med24 | -1.030215294 | 3.981570786 |
| A2BH40;E9Q4N7 | Arid | -1.032244329 | 2.92801555 |
| Q9CZA6 | Nde1 | -1.032419077 | 3.052309328 |
| Q8CJG0 | Ago2 | -1.032782515 | 2.848176091 |
| Q91VM9 | Ppa2 | -1.034265359 | 2.185509706 |
| Q61164 | Ctcf | -1.035379978 | 2.692919997 |
| Q9CVB6 | Arpc2 | -1.035621884 | 4.854481453 |
| O08532 | Cacna2d1 | -1.035637433 | 2.559507335 |
| P60766 | Cdc42 | -1.037305836 | 3.469498339 |
| Q8BPM2 | Map4k5 | -1.039779097 | 3.788792631 |
| Q6P5D8 | Smchd1 | -1.04166048 | 3.044430754 |
| P97496 | Smarcc1 | -1.04376473 | 3.375319845 |
| Q9R0E2 | Plod1 | -1.045790062 | 2.416517884 |
| P46471 | Psmc2 | -1.045878037 | 4.937696224 |
| O08992 | Sdcbp | -1.046006178 | 3.409818611 |
| O88685 | Psmc3 | -1.047067623 | 4.717524465 |
| P09041;P09411 | Pgk | -1.047617288 | 4.458131581 |
| P62962 | Pfn1 | -1.047623547 | 2.752901764 |
| Q99MR0 | Actl6b | -1.048810944 | 2.514610742 |
| P26040;P26041;P26043 | Ezr | -1.049602681 | 3.431515452 |
| P47226 | Tes | -1.050058115 | 3.786435915 |
| Q91YR1 | Twf1 | -1.051734271 | 2.955076333 |
| Q99JY9 | Actr3 | -1.052031503 | 4.801067323 |
| Q8K0D5 | Gfm1 | -1.052607665 | 4.717524465 |
| Q8BI72 | Cdkn2aip | -1.053387849 | 2.851468308 |
| Q9CXZ1 | Ndufs4 | -1.053492664 | 2.888708295 |
| A0A7H0DNG0 | OPG200 | -1.053870084 | 4.291650172 |
| Q99LE6 | Abcf2 | -1.054119505 | 3.90166834 |
| Q9CX11 | Utp23 | -1.054898183 | 2.897910906 |
| O08600 | Endog | -1.056861722 | 2.34252566 |
| Q61214;Q9Z188 | Dyrk1 | -1.056877153 | 2.843802671 |
| Q8R502 | Lrrc8c | -1.059489373 | 2.818465152 |
| A2RSY1 | Kansl3 | -1.059679894 | 2.69129318 |
| Q8K268 | Abcf3 | -1.061024287 | 3.914955128 |
| O70378 | Emc8 | -1.061284494 | 2.796883871 |
| Q8BSQ9 | Pbrm1 | -1.06347555 | 3.105829734 |
| P08775 | Polr2a | -1.064822496 | 4.035823747 |
| Q8BKZ9 | Pdhx | -1.064957026 | 3.048607951 |
| P47739;P47740 | Aldh3a | -1.065225702 | 2.550062839 |
| P24288 | Bcat1 | -1.068913249 | 3.469534795 |
| A0A7H0DNE2 | OPG174 | -1.070026904 | 2.261523885 |
| Q3UQN2 | Fcho2 | -1.07096326 | 4.057208694 |
| Q8BJW5 | Nol11 | -1.071524313 | 2.232745641 |
| P97452 | Bop1 | -1.073018939 | 3.190379089 |
| Q9DC48 | Cdc40 | -1.073250293 | 4.070814827 |
| Q80UY2 | Kcmf1 | -1.073395733 | 2.853380929 |
| Q9CZP0 | Ufsp1 | -1.073788911 | 2.450732767 |
| Q9EQ80 | Nif3l1 | -1.075369513 | 2.84459214 |
| Q8BIW9 | Chtf18 | -1.07585187 | 4.710034016 |
| P56480 | Atp5f1b | -1.076557661 | 2.359363145 |
| Q8BMC4 | Nop9 | -1.077546069 | 3.759520826 |
| P51655 | Gpc4 | -1.079623619 | 2.979372903 |
| Q99JR8 | Smarcd2 | -1.08041322 | 3.345372784 |
| Q00547 | Hmmr | -1.080655503 | 2.16162239 |
| Q99KB8 | Hagh | -1.081040416 | 3.851287414 |
| Q9JLQ2 | Git2 | -1.081468088 | 3.346240978 |
| Q3UZ39;Q91WK0 | Lrrfip2 | -1.083304676 | 3.073620454 |
| Q8K2Q5 | Chchd7 | -1.084498612 | 3.098190734 |
| Q6TYB5 | Fez2 | -1.085555884 | 3.867651428 |
| Q64707;Q62377 | Zrsr2 | -1.086681827 | 2.346475041 |
| A2ABV5 | Med14 | -1.08680455 | 4.406025734 |
| P16951 | Atf2 | -1.089342127 | 3.166785549 |
| Q7TT50 | Cdc42bpb | -1.08998566 | 2.806190992 |
| P13439 | Umps | -1.090277915 | 5.374374854 |
| Q922K7 | Nop2 | -1.091546401 | 3.124100243 |
| Q9D8C6 | Med11 | -1.092921089 | 3.000998021 |
| Q99LD8 | Ddah2 | -1.093742943 | 3.87933269 |
| Q6ZQL4 | Wdr43 | -1.094084729 | 3.418453048 |
| Q8R1V4 | Tmed4 | -1.094369039 | 2.247885409 |
| A0A7H0DNE5 | OPG178 | -1.094414676 | 3.002323111 |
| Q91ZU6 | Dst | -1.094768904 | 3.881835894 |
| Q9QUR6 | Prep | -1.095024561 | 4.141563665 |
| E9Q634;P70248 | Myo1f | -1.095629387 | 3.851287414 |
| Q9QUJ7 | Acsl4 | -1.095930822 | 4.077057136 |
| Q6VN19 | Ranbp10 | -1.098502268 | 3.978925047 |
| Q9JKP5;Q8R003 | Mbnl | -1.098997421 | 3.995737309 |
| P70441 | Nherf1 | -1.099035286 | 3.432177068 |
| Q8R034 | Anapc13 | -1.099838755 | 2.624820406 |
| Q5SRX1 | Tom1l2 | -1.101001231 | 3.197552951 |
| Q6ZQ73 | Cand2 | -1.103246731 | 3.455729418 |
| Q5BLK4 | Tut7 | -1.105397088 | 2.305241321 |
| Q8BH59 | Slc25a12 | -1.107228028 | 2.741728091 |
| Q3TMV7 | Pyroxd1 | -1.107658693 | 2.320283664 |
| P21107 | Tpm3 | -1.108475639 | 3.116747084 |
| Q9ER81 | Tor1aip2 | -1.110055852 | 2.903138241 |
| O09172 | Gclm | -1.110391913 | 3.779011727 |
| O08529 | Capn2 | -1.111099312 | 3.311841059 |
| Q64674 | Srm | -1.112961874 | 3.546072554 |
| P46638 | Rab11b | -1.114244759 | 3.625278972 |
| Q811D0;Q91XM9 | Dlg | -1.115337734 | 2.538733416 |
| Q80VJ2 | Sra1 | -1.116582798 | 3.851287414 |
| P97310 | Mcm2 | -1.118871529 | 3.159174208 |
| P58742 | Aaas | -1.120062455 | 2.164018515 |
| Q8BGY7 | Fam210a | -1.121305602 | 3.359298731 |
| P48024;Q9CXU9 | Eif1 | -1.121524593 | 4.200670667 |
| Q60722;Q61286 | Tc4 | -1.121815057 | 3.344322415 |
| Q6ZQF0 | Topbp1 | -1.12391846 | 3.031905797 |
| Q91YH5 | Atl3 | -1.124621542 | 2.128717944 |
| Q3UMG5 | Lrch2 | -1.125144761 | 3.039873912 |
| Q6IE82 | Jade3 | -1.128238793 | 4.112667745 |
| Q8BGS0 | Mak16 | -1.128431466 | 4.145136909 |
| D3Z4I3;Q62176 | Rbm | -1.129818649 | 3.512639993 |
| P38647 | Hspa9 | -1.1300398 | 3.7075662 |
| Q91VX2 | Ubap2 | -1.131461702 | 2.812385134 |
| B1AVH7 | Tbc1d2 | -1.131499157 | 2.418399703 |
| Q8QZT1 | Acat1 | -1.131657965 | 2.80968254 |
| Q9DAR7 | Dcps | -1.134858615 | 3.220594607 |
| Q91W36 | Usp3 | -1.135286823 | 3.596367803 |
| P05480 | Src | -1.135541499 | 3.266881356 |
| Q8BG51 | Rhot1 | -1.136481127 | 2.978017029 |
| P12815 | Pdcd6 | -1.136906766 | 4.057208694 |
| Q8BHG1 | Nrdc | -1.137047322 | 4.47529076 |
| Q8VBT9 | Aspscr1 | -1.140021609 | 3.15212545 |
| Q3UHB1 | Nt5dc3 | -1.140281862 | 4.562735348 |
| Q91WG5 | Prkag2 | -1.143100352 | 2.825911095 |
| O08856 | Ell | -1.143103527 | 2.779055685 |
| Q925J9 | Med1 | -1.144248042 | 3.692526592 |
| Q9R0Q6;Q9WV32 | Arpc | -1.146390456 | 3.063473027 |
| P35831 | Ptpn12 | -1.14657528 | 2.469280776 |
| Q01405;Q9D662 | Sec23 | -1.146972403 | 2.666181318 |
| Q99JY9;Q641P0 | Actr3 | -1.147980692 | 3.083243027 |
| Q8BG15 | Ctdspl2 | -1.149311627 | 3.509403361 |
| P09405 | Ncl | -1.149416812 | 4.306368382 |
| Q99PL5 | Rrbp1 | -1.150491022 | 2.371484749 |
| O89050 | Mkln1 | -1.151402403 | 3.439486415 |
| P12787 | Cox5a | -1.152110151 | 3.793611432 |
| Q5SV77 | Ggnbp2 | -1.152301506 | 2.974227176 |
| Q9ERR7 | Selenof | -1.152707238 | 3.14582837 |
| Q80U58 | Pum2 | -1.154209178 | 3.227440977 |
| Q62388 | Atm | -1.154762267 | 2.055314098 |
| Q3TC33 | Ccdc127 | -1.157156957 | 2.558818026 |
| O55176 | Pja1 | -1.15793146 | 2.003744017 |
| Q8C7V3 | Utp15 | -1.158041388 | 3.130165145 |
| P48962;P51881 | Slc25a | -1.15932471 | 2.778042453 |
| Q8VEH8 | Erlec1 | -1.15939281 | 3.780123575 |
| Q9CR86 | Carhsp1 | -1.159731571 | 2.140871428 |
| Q3TC93 | Hs1bp3 | -1.161213079 | 3.421396536 |
| Q9Z0H3 | Smarcb1 | -1.163818417 | 4.42846902 |
| P35822 | Ptprk | -1.164726773 | 2.245765358 |
| P62192 | Psmc1 | -1.164883532 | 4.564076775 |
| Q64433 | Hspe1 | -1.165337225 | 2.085715016 |
| P52623 | Uck1 | -1.1662332 | 3.041156903 |
| Q571E4 | Galns | -1.167925402 | 4.309928136 |
| Q4VA53 | Pds5b | -1.168347168 | 4.005229598 |
| Q91YS8 | Camk1 | -1.169054289 | 4.104706659 |
| Q9WTX8 | Mad1l1 | -1.170144406 | 4.289311731 |
| Q62188 | Dpysl3 | -1.170541148 | 4.390003343 |
| P70445 | Eif4ebp2 | -1.170995826 | 3.363017573 |
| Q8BQZ5 | Cpsf4 | -1.171301694 | 5.060390014 |
| Q64516;Q9WU65 | Gk2 | -1.171625053 | 2.97471665 |
| A0A7H0DN09 | OPG036 | -1.171700734 | 2.706328935 |
| Q33DR3 | Pdss2 | -1.17350452 | 2.890174672 |
| Q920D3 | Med28 | -1.173613351 | 2.612917689 |
| Q8BXL7 | Arfrp1 | -1.176484478 | 3.241225708 |
| Q3UHJ0 | Aak1 | -1.180515254 | 3.260771276 |
| Q6PGF3 | Med16 | -1.182120317 | 3.190864341 |
| P51863 | Atp6v0d1 | -1.182588457 | 3.027136856 |
| P35123;Q8R5H1 | Usp4;Usp15 | -1.183204363 | 2.84089245 |
| Q7TMF3 | Ndufa12 | -1.18466276 | 3.908265185 |
| Q91VH6 | Memo1 | -1.184664529 | 4.155788101 |
| Q9CY34 | Ube2f | -1.187295852 | 3.404627076 |
| P21278;P21279 | Gna | -1.188819641 | 2.81927235 |
| Q6PHZ2 | Camk2d | -1.189498811 | 4.500546036 |
| P26443 | Glud1 | -1.191772208 | 3.082698915 |
| P12382;P47857 | Pfk | -1.192115582 | 3.525645116 |
| Q80UG5 | Septin9 | -1.194751631 | 4.122513054 |
| Q01320 | Top2a | -1.194819237 | 2.640588003 |
| B2RSH2;P08752 | Gnai | -1.195260794 | 3.017087539 |
| O88811 | Stam2 | -1.197212133 | 3.352632447 |
| Q8BVD5 | Mpp7 | -1.198854669 | 4.074884235 |
| P63028 | Tpt1 | -1.200856435 | 2.783538025 |
| Q9CZL5 | Pcbd2 | -1.201148198 | 2.976916077 |
| P62075 | Timm13 | -1.201336279 | 3.273064116 |
| Q8K411 | Pitrm1 | -1.203012672 | 3.954965996 |
| P47791 | Gsr | -1.203604683 | 4.128456686 |
| Q62108;Q811D0;Q91XM9 | Dlg4 | -1.204651527 | 2.175058901 |
| Q9Z2D1 | Mtmr2 | -1.205194594 | 2.280970113 |
| O88738 | Birc6 | -1.20684834 | 3.962846066 |
| Q8C147;Q8R1A4 | Dock8 | -1.20738917 | 2.87611009 |
| P59108 | Cpne2 | -1.207699806 | 4.355577622 |
| Q80YR7 | Clspn | -1.208298365 | 2.824218045 |
| Q9D1N9 | Mrpl21 | -1.208473375 | 2.529105879 |
| O35945;P24549;P47738;Q62148;Q9JHW9;Q9CZS1 | Aldh1 | -1.211473991 | 3.159174208 |
| Q62523 | Zyx | -1.211529729 | 3.302366022 |
| P63094;Q6R0H7;Q8CGK7 | Gnal | -1.211549034 | 3.636575469 |
| Q9CY18 | Snx7 | -1.21222877 | 3.867651428 |
| Q03265 | Atp5f1a | -1.214673933 | 2.850077071 |
| P55194 | Sh3bp1 | -1.216396912 | 3.650759083 |
| P12849;Q9DBC7 | Prkar1 | -1.218240996 | 2.900774461 |
| Q922J3;Q9Z0H8 | Clip2 | -1.219756516 | 3.95361594 |
| A0A7H0DNF0 | OPG188 | -1.219923554 | 3.174932088 |
| P11798;P28652;Q6PHZ2;Q923T9 | Camk2a | -1.220269144 | 2.472715855 |
| Q9Z277 | Baz1b | -1.221110818 | 3.861784354 |
| Q9QWT9 | Kifc1 | -1.221856372 | 2.848799441 |
| Q8K2B3 | Sdha | -1.222528378 | 3.910004264 |
| Q9CXK8 | Nip7 | -1.223724761 | 3.781559558 |
| Q925H1 | Trps1 | -1.225525699 | 2.350343013 |
| O08553;Q62188 | Dpysl | -1.22703689 | 3.095093951 |
| Q922Q4 | Pycr2 | -1.234077164 | 5.079252495 |
| Q3TYX3 | Smyd5 | -1.234690144 | 2.451325013 |
| Q9JLZ3 | Auh | -1.238075085 | 3.619573707 |
| Q61103 | Dpf2 | -1.238942392 | 4.339632504 |
| P58871 | Tnks1bp1 | -1.241833364 | 3.212102835 |
| Q921X9 | Pdia5 | -1.2445024 | 2.904477274 |
| P52480 | Pkm | -1.244784276 | 4.684011748 |
| Q9JHC9 | Elf2 | -1.246335235 | 3.313247495 |
| Q8BVI4 | Qdpr | -1.249942461 | 4.128456686 |
| O08674;P31266 | Rbpj | -1.250432514 | 2.257530294 |
| Q91W59 | Rbms1 | -1.251320355 | 4.260158219 |
| Q9QY30 | Abcb11 | -1.252467974 | 2.373294681 |
| Q8R2E9 | Ero1b | -1.254842677 | 2.42986597 |
| Q01853 | Vcp | -1.256992619 | 5.396543964 |
| P43024 | Cox6a1 | -1.258250527 | 3.073025804 |
| Q9DBZ9 | Syde1 | -1.263683661 | 2.237869018 |
| Q9DCE5 | Pak1ip1 | -1.264154708 | 2.893378644 |
| Q6P9L6 | Kif15 | -1.264662144 | 3.867651428 |
| Q8R5J9 | Arl6ip5 | -1.265132189 | 2.295081946 |
| P49717 | Mcm4 | -1.266716309 | 4.112667745 |
| Q9CX86 | Hnrnpa0 | -1.266853493 | 4.146679376 |
| P47934 | Crat | -1.272525584 | 2.56254237 |
| Q5SF07 | Igf2bp2 | -1.274670425 | 4.044096318 |
| Q91XU0 | Wrnip1 | -1.27493069 | 3.736292763 |
| Q61586 | Gpam | -1.27849699 | 3.602814454 |
| Q9R112 | Sqor | -1.278649781 | 2.040495536 |
| Q9CQ39 | Med21 | -1.280902535 | 4.1611869 |
| Q6A0A9 | FAM120A | -1.280960774 | 3.832088521 |
| P62137 | Dnajb11 | -1.281529967 | 4.927022073 |
| Q99KV1 | Atp2b1 | -1.282582544 | 4.391071762 |
| G5E829 | Spag7 | -1.282721955 | 3.396273085 |
| Q7TNE3 | Paip2b | -1.283787139 | 3.484552981 |
| Q91W45 | Paip2b | -1.284093252 | 2.135475719 |
| P08113 | Hsp90b1 | -1.284534747 | 3.710054148 |
| Q6QI06 | Rictor | -1.284740505 | 2.610793407 |
| P97311 | Mcm6 | -1.284906373 | 4.669873401 |
| Q924T7 | Rnf31 | -1.285983681 | 3.177646571 |
| Q8VDY9 | Caap1 | -1.286480761 | 2.792750104 |
| O35379 | Abcc1 | -1.290175472 | 2.930150574 |
| Q9JKX4 | Aatf | -1.290622323 | 4.338387509 |
| Q922M3 | Kctd10 | -1.292471349 | 3.601456746 |
| Q8K3X4 | Irf2bpl | -1.293836798 | 3.608392105 |
| P13864 | Dnmt1 | -1.293938242 | 2.882508743 |
| P06801 | Me1 | -1.294016908 | 4.908373764 |
| Q8BNY6 | Ncs1 | -1.29840349 | 2.561188498 |
| Q8CCH7 | Zfpm2 | -1.299426943 | 2.159184717 |
| A0A7H0DN93 | OPG121 | -1.300414267 | 4.057560896 |
| Q8V4S4;A0A7H0DNF0 | OPG188 | -1.301811698 | 5.637526423 |
| Q9JJ28 | Flii | -1.302160179 | 3.731113495 |
| O89032 | Sh3pxd2a | -1.304915843 | 2.079072417 |
| Q8BKG3 | Ptk7 | -1.311132636 | 2.613564548 |
| Q9D6K8 | Fundc2 | -1.316922132 | 3.041156903 |
| A0A7H0DND6 | OPG165 | -1.317170078 | 5.123103562 |
| Q91W90 | Txndc5 | -1.318865549 | 4.398916716 |
| Q9JHP7 | Poglut2 | -1.320958168 | 3.270529768 |
| Q9CQF4 | Mtres1 | -1.321025257 | 2.740495271 |
| E9PVB5 | Ttc17 | -1.324245807 | 4.145287568 |
| Q99N85 | Mrps18a | -1.32427928 | 2.387576484 |
| Q8BHY2 | Noc4l | -1.324435411 | 2.352247636 |
| Q08509 | Eps8 | -1.327035992 | 3.006376566 |
| Q8R3R8 | Gabarapl1 | -1.328263868 | 2.152855134 |
| Q8CJ53 | Trip10 | -1.330832157 | 4.341982222 |
| A2APV2 | Fmnl2 | -1.332204987 | 2.236303067 |
| Q9JJA7 | Ccnl2 | -1.332703084 | 4.347223105 |
| Q9D7P6 | Iscu | -1.335214585 | 3.222191862 |
| P02798 | Mt2 | -1.336651939 | 3.899435712 |
| Q8BU85 | Msrb3 | -1.337301241 | 2.040424779 |
| P51480 | Cdkn2a | -1.339071965 | 3.711274029 |
| Q8BKX1 | Baiap2 | -1.340928646 | 4.057208694 |
| P97493 | Txn2 | -1.341431161 | 4.869878922 |
| Q9D832 | Dnajb4 | -1.341442808 | 3.253266322 |
| Q9R207 | Nbn | -1.341642039 | 2.306476937 |
| Q8C7H1 | Mmaa | -1.342859807 | 2.904477274 |
| P35293 | Rab18 | -1.343811202 | 2.561188498 |
| Q07076 | Anxa7 | -1.344549865 | 3.170236202 |
| Q8C1S0 | Med19 | -1.345152379 | 3.378524604 |
| P11157 | Rrm2 | -1.347141864 | 3.516067913 |
| Q6VNB8 | Wdfy3 | -1.354505881 | 2.06808028 |
| Q9WVA2 | Timm8a1 | -1.355602227 | 3.492822794 |
| Q7TSG2 | Ctdp1 | -1.355674724 | 4.336926621 |
| Q3THE2;Q9CQ19 | Myl | -1.356221811 | 2.967737016 |
| Q9JK81 | Myg1 | -1.360786508 | 2.810436278 |
| Q9JJ78 | Pbk | -1.363795267 | 3.159174208 |
| P05202 | Got2 | -1.365381158 | 2.23250424 |
| P97432 | Nbr1 | -1.367303019 | 2.771648359 |
| Q8BWT5 | Dip2a | -1.367404996 | 3.019023631 |
| E9PYK3 | Parp4 | -1.371817619 | 3.266881356 |
| P70404 | Idh3g | -1.376250987 | 3.22009771 |
| Q9WV30 | Nfat5 | -1.379898282 | 4.293238757 |
| Q9ER39 | Tor1a | -1.381050437 | 4.622325871 |
| Q91ZV0 | Mia2 | -1.38396185 | 2.311810639 |
| Q64514 | Tpp2 | -1.384478243 | 4.748067278 |
| Q9D787 | Ppil2 | -1.385351928 | 3.638277906 |
| Q91V12 | Acot7 | -1.390574815 | 4.971384312 |
| Q9DCX2 | Atp5pd | -1.391805539 | 3.405722143 |
| Q9CZR8 | Tsfm | -1.391892897 | 3.538507794 |
| Q60848 | Hells | -1.392082445 | 3.168835891 |
| Q02819 | Nucb1 | -1.393534506 | 4.412462579 |
| Q922R5 | Ppp4r3b | -1.39480709 | 3.399028185 |
| Q9CXI5 | Manf | -1.397109017 | 3.49155129 |
| Q9CQ92 | Fis1 | -1.400029326 | 3.981051698 |
| Q9EP71 | Rai14 | -1.401943321 | 5.453998375 |
| Q9ES97 | Rtn3 | -1.403747017 | 3.576564595 |
| P25206 | Mcm3 | -1.405931647 | 5.396543964 |
| P61022 | Chp1 | -1.406104691 | 2.907852665 |
| P51569 | Gla | -1.406278657 | 2.491874269 |
| Q9CY50 | Ssr1 | -1.407298531 | 2.006841952 |
| P47713 | Pla2g4a | -1.411209569 | 4.336926621 |
| Q8R1F1 | Niban2 | -1.411372609 | 4.402250522 |
| Q8R180 | Ero1a | -1.413496337 | 4.420212838 |
| Q8VE99 | Vma22 | -1.41431731 | 3.786435915 |
| Q8R149 | Bud13 | -1.41565139 | 3.80055133 |
| Q6P9J9 | Ano6 | -1.417925556 | 3.393290214 |
| Q9JMH9 | Myo18a | -1.418583566 | 2.399990927 |
| A0A7H0DN24 | OPG051 | -1.418611179 | 2.767256915 |
| Q6NVE9 | Got2 | -1.419020423 | 3.113400012 |
| Q8BGC4 | Ptgr3 | -1.424010223 | 2.42193795 |
| Q9D6R2 | Idh3a | -1.424592792 | 4.420212838 |
| Q61792 | Lasp1 | -1.426558436 | 4.128456686 |
| Q9JJ66 | Cdc20 | -1.428897308 | 2.207916612 |
| Q9D0M3 | Cyc1 | -1.431580546 | 2.883195616 |
| Q3TZZ7 | Esyt2 | -1.439351299 | 3.146076189 |
| Q60875 | Arhgef2 | -1.440878795 | 3.563777056 |
| P01901 | H2-K1 | -1.442078561 | 3.451032441 |
| A6H8H2 | Dennd4c | -1.442116579 | 3.000998021 |
| P62500;E9Q7M2 | Tsc22d | -1.443715008 | 3.304278574 |
| O54940 | Bnip2 | -1.446094138 | 3.564860196 |
| Q68FL4 | Ahcyl2 | -1.446168267 | 2.977301264 |
| Q8K3A0 | Hscb | -1.447663115 | 3.111237931 |
| Q3UE17 | Mex3d | -1.450746882 | 2.064028719 |
| B2RUR8;Q8R554 | Otud7 | -1.452218008 | 4.543625599 |
| P70290 | Mpp1 | -1.462276685 | 4.066442441 |
| Q6ZWY3 | Rps27l | -1.464789301 | 3.077833984 |
| P35441 | Thbs1 | -1.466698229 | 3.070760511 |
| P30681 | Hmgb2 | -1.468926513 | 3.729277997 |
| Q9WUM3 | Coro1b | -1.469401353 | 4.937696224 |
| Q4VAA7 | Snx33 | -1.469693214 | 3.017906209 |
| Q6PB44 | Ptpn23 | -1.471491967 | 3.667783416 |
| Q8BVE8 | Nsd2 | -1.471517201 | 4.493629003 |
| Q9CRW3 |  | -1.471589955 | 2.144793083 |
| Q8CH72 | Trim32 | -1.471669936 | 3.8077472 |
| Q9D3E6 | Stag1 | -1.473087243 | 4.420212838 |
| Q9D818 | Sapcd2 | -1.476095339 | 3.724672543 |
| Q0KL02 | Trio | -1.480510857 | 3.958986167 |
| P56212 | Arpp19 | -1.484432056 | 2.77569627 |
| P12023 | App | -1.485813994 | 2.885149019 |
| Q61881 | Mcm7 | -1.488065017 | 4.289311731 |
| Q80U72;Q80VQ1 | Scrib | -1.488395579 | 2.927605028 |
| A0A7H0DN02;Q8V566 | OPG027 | -1.496437618 | 3.768338095 |
| Q9JI10 | Stk3 | -1.497509425 | 3.304278574 |
| Q9JHJ0 | Tmod3 | -1.498169921 | 3.545440266 |
| P49718 | Mcm5 | -1.501104415 | 3.7075662 |
| Q8BJL0 | Smarcal1 | -1.502487514 | 2.279780718 |
| Q922H2 | Pdk3 | -1.504190094 | 4.479925882 |
| Q9EPV8 | Ubl5 | -1.504863186 | 3.326609431 |
| Q8VE73 | Cul7 | -1.507660129 | 4.237369263 |
| O08664 | Bcl7c | -1.508284479 | 3.292557965 |
| Q5NCR9 | Nsrp1 | -1.508709292 | 2.015195732 |
| P54071 | Idh2 | -1.512552803 | 3.726682166 |
| Q6PCP5 | Mff | -1.520786778 | 2.756444127 |
| P23591 | Gfus | -1.523247308 | 3.984452126 |
| B2RXS4 | Plxnb2 | -1.525522639 | 3.621386629 |
| Q3UHX0 | Nol8 | -1.5256228 | 3.040016404 |
| D3Z7P3 | Gls | -1.52869062 | 4.030098513 |
| Q8BG73 | Sh3bgrl2 | -1.529906029 | 3.828422659 |
| Q9WV55 | Vapa | -1.530085696 | 2.165669584 |
| Q921H8 | Acaa1a | -1.530693227 | 2.521415831 |
| P37913 | Lig1 | -1.536774982 | 4.398916716 |
| Q921E2 | Rab31 | -1.536835306 | 3.614642117 |
| Q8K1J6 | Trnt1 | -1.53775701 | 4.207432459 |
| Q9JM14 | Nt5c | -1.538551814 | 4.375749023 |
| O54984 | Get3 | -1.542053002 | 4.784426578 |
| Q80X90;Q8BTM8;Q8VHX6 | Fln | -1.545570984 | 5.123103562 |
| Q8BSY0 | Asph | -1.54704683 | 2.937737245 |
| Q9CYN9 | Atp6ap2 | -1.547907776 | 3.42895811 |
| Q9CQF9 | Pcyox1 | -1.550313177 | 3.026633427 |
| P35601 | Rfc1 | -1.552344171 | 3.998412163 |
| P19324 | Serpinh1 | -1.555667973 | 4.168276181 |
| A0A7H0DNG5 | OPG209 | -1.556424111 | 2.346065993 |
| Q9Z1S0 | Bub1b | -1.55686033 | 4.891028448 |
| P58137 | Acot8 | -1.557740268 | 3.596305297 |
| Q9WUB0 | Rbck1 | -1.558504042 | 3.58429209 |
| Q8R2Y8 | Ptrh2 | -1.558811865 | 4.301759602 |
| P57776 | Eef1d | -1.559932337 | 4.375749023 |
| Q9R0A0 | Pex14 | -1.560721866 | 3.307197121 |
| Q922J9 | Far1 | -1.561237916 | 2.769654623 |
| A0A7H0DN07 | OPG034 | -1.562649197 | 3.293221175 |
| A2CG49;Q0KL02 | Kalm | -1.563434777 | 3.011588645 |
| P39428 | Traf1 | -1.566374746 | 3.116689103 |
| Q8BRM2 | Gorab | -1.568706868 | 2.56254237 |
| E9Q414 | Apob | -1.569721898 | 2.829147331 |
| Q9Z0U1 | Tjp2 | -1.569917307 | 4.501489155 |
| P37040 | Por | -1.57073416 | 2.872380237 |
| Q91VW3 | Sh3bgrl3 | -1.573482642 | 4.603939091 |
| O35887 | Calu | -1.573918233 | 4.306368382 |
| Q9R0P3 | Esd | -1.576047856 | 4.912516946 |
| Q5PRF0 | Heatr5a | -1.578402356 | 4.103666555 |
| O88878 | Zfand5 | -1.580988984 | 2.276345612 |
| Q8VE22 | Mrps23 | -1.58752419 | 2.785784034 |
| Q8CFE6 | Slc38a2 | -1.592140086 | 2.786515787 |
| Q99P72 | Rtn4 | -1.593755417 | 4.136514413 |
| P58802 | Tbc1d10a | -1.595504787 | 3.486617254 |
| P18242 | Ctsd | -1.597303061 | 4.622325871 |
| Q9CQ65 | Mtap | -1.603263677 | 5.139550781 |
| Q6P5E4 | Uggt1 | -1.604120865 | 3.463538703 |
| Q9Z1Q9 | Vars1 | -1.607091344 | 4.341982222 |
| O70475 | Ugdh | -1.608183185 | 5.139550781 |
| Q9QZ23 | Nfu1 | -1.608961103 | 3.348189879 |
| B2RSH2;P08752;P18872;Q9DC51 | Gnao | -1.609084855 | 3.173768604 |
| Q9ES00 | Ube4b | -1.614513892 | 3.504574648 |
| Q8BIG7 | Comtd1 | -1.615451314 | 3.44494674 |
| Q99M54 | Cdca3 | -1.615902479 | 3.096421615 |
| Q91W67 | Ubl7 | -1.62393095 | 3.172071882 |
| P21107;P58771;P58774 | Tpm1 | -1.624626531 | 3.745879176 |
| Q8C8U0 | Ppfibp1 | -1.625141852 | 3.984452126 |
| Q9DAU1 | Cnpy3 | -1.625597484 | 3.74158152 |
| Q8BJU9 | Mtrf1l | -1.62564864 | 2.935122989 |
| Q8K4K6 | Pank1 | -1.631625497 | 2.373209732 |
| P45878 | Fkbp2 | -1.634885866 | 4.937696224 |
| O70201 | Birc5 | -1.635477419 | 2.120562774 |
| Q8CGK3 | Lonp1 | -1.636198011 | 3.914955128 |
| P97384 | Anxa11 | -1.638081701 | 2.218917016 |
| Q8BT07 | Cep55 | -1.638133902 | 2.579172374 |
| Q61072 | Adam9 | -1.643579431 | 2.241615401 |
| O35295 | Purb | -1.647626386 | 4.398916716 |
| Q3UFB2 | Znhit6 | -1.648180385 | 2.718269877 |
| Q3UHH1 | Zswim8 | -1.649509665 | 3.909415931 |
| Q3U4G3 | Xxylt1 | -1.650340908 | 2.571799115 |
| Q6PDH0 | Phldb1 | -1.651221999 | 3.710054148 |
| Q8BTY2 | Slc4a7 | -1.6517585 | 3.41069737 |
| P48036 | Anxa5 | -1.654362193 | 3.984452126 |
| Q9WTK7 | Stk11 | -1.656930871 | 2.418399703 |
| O70311 | Nmt2 | -1.658558871 | 2.790105411 |
| P14211 | Calr | -1.660090428 | 3.821678735 |
| Q9CRA4 | Msmo1 | -1.663859524 | 2.396457932 |
| Q9CZT8 | Rab3b | -1.664061977 | 2.740583076 |
| Q99LI2 | Clcc1 | -1.666784217 | 3.004003469 |
| Q91VH2 | Snx9 | -1.667393095 | 3.731306767 |
| O70480 | Vamp4 | -1.671434623 | 3.081485254 |
| Q8R1J9;P0C7W3 | Tor2a | -1.674153086 | 3.215995546 |
| A0A7H0DMZ6 | OPG019 | -1.674291598 | 3.403930181 |
| Q8BP86 | Snapc4 | -1.676221169 | 2.006982608 |
| P54116 | Stom | -1.67634817 | 2.950403017 |
| P52633 | Stat6 | -1.690259078 | 2.400131292 |
| P31324 | Prkar2b | -1.694840716 | 4.084292067 |
| Q99104 | Myo5a | -1.697048415 | 4.746108676 |
| Q9R0E1 | Plod3 | -1.701713657 | 4.640089824 |
| P14733;P21619;P48678 | Lmnb | -1.704534439 | 4.398916716 |
| Q9D7B6 | Acad8 | -1.705728755 | 3.307197121 |
| Q9R0Q4 | Morf4l2 | -1.707289298 | 2.386409001 |
| O70433 | Fhl2 | -1.709355135 | 3.271332629 |
| Q9CZJ2 | Hspa12b | -1.710072509 | 2.368096158 |
| Q921F4 | Hnrnpll | -1.71114562 | 5.396543964 |
| O70404 | Vamp8 | -1.711322551 | 3.497547341 |
| Q3TBT3 | Sting1 | -1.712195535 | 2.961979029 |
| Q9ER41 | Tor1b | -1.712723377 | 4.223165845 |
| Q80TM9 | Nisch | -1.715144654 | 4.05095065 |
| P52332 | Jak1 | -1.716353999 | 2.903628428 |
| O08599 | Stxbp1 | -1.721680395 | 4.324695846 |
| Q80W47 | Wipi2 | -1.722543468 | 3.946660784 |
| P20152;P31001 | Vim;Des | -1.728543228 | 3.363017573 |
| P98083;Q8BMC3 | Shc | -1.730444798 | 3.178403054 |
| P17095 | Hmga1 | -1.734730319 | 4.917613712 |
| Q8BH64;Q9WVK4 | Ehd | -1.736482003 | 3.226406609 |
| Q91YW3 | Dnajc3 | -1.737723565 | 4.657002098 |
| Q9D2X0 | Ankrd39 | -1.73840066 | 3.108390384 |
| Q99KI0 | Aco2 | -1.739060988 | 3.838459094 |
| Q8C6B2 | Rtkn | -1.744184757 | 2.420600106 |
| A0A7H0DN02 | OPG027 | -1.746261481 | 5.038593208 |
| Q8K009 | Aldh1l2 | -1.756589171 | 3.512639993 |
| Q6P2L7 | Golm2 | -1.760691122 | 2.791662215 |
| P0DTM9 | OPG001 | -1.761384578 | 4.012821866 |
| Q3UW53 | Niban1 | -1.76573839 | 4.688478355 |
| Q91XL3 | Uxs1 | -1.770556945 | 2.860120163 |
| Q9DB77 | Uqcrc2 | -1.771368879 | 2.840578782 |
| P50431 | Shmt1 | -1.784223278 | 4.668093374 |
| Q9CZN7 | Shmt2 | -1.788518689 | 4.869878922 |
| Q8K215 | Lyrm4 | -1.789914399 | 2.785747576 |
| Q9Z2Q6 | Septin5 | -1.79233596 | 4.487641205 |
| Q00612 | G6pdx | -1.793707578 | 4.986912996 |
| Q06348 | Prrx2 | -1.794342409 | 2.308640179 |
| Q60949 | Tbc1d1 | -1.794549622 | 4.077057136 |
| Q8BZJ7 | Dcun1d2 | -1.7951413 | 2.126288629 |
| Q8VHI3 | Pofut2 | -1.796483903 | 3.891924175 |
| P10833 | Rras | -1.797245837 | 2.911605619 |
| Q8VI36 | Pxn | -1.801572579 | 5.10293975 |
| Q9D8Z1 | Ascc1 | -1.802145932 | 2.747575157 |
| Q80VI1 | Trim56 | -1.805446648 | 4.327341891 |
| Q61207 | Psap | -1.807276327 | 2.157964732 |
| Q32P12 | Hrob | -1.81440982 | 2.07061604 |
| A0A7H0DN11 | OPG038 | -1.814463081 | 4.044096318 |
| Q9JKR6 | Hyou1 | -1.821004825 | 4.176105219 |
| E9Q0S6 | Tns1 | -1.828392706 | 2.89459226 |
| Q99JI1 | Mustn1 | -1.829838536 | 2.346065993 |
| Q9R059 | Fhl3 | -1.831756926 | 3.831087744 |
| Q8K212 | Pacs1 | -1.835714639 | 3.627907501 |
| Q91WS0 | Cisd1 | -1.837142928 | 3.858131573 |
| Q8BMD8 | Slc25a24 | -1.83914536 | 4.235658482 |
| Q9D924 | Isca1 | -1.842633322 | 2.532285204 |
| P09103 | P4hb | -1.845308167 | 4.27815874 |
| Q5DTT3 | Tasor2 | -1.855169718 | 2.400735378 |
| Q8JZV7 | Amdhd2 | -1.856955837 | 2.903969406 |
| O88342 | Wdr1 | -1.85942388 | 5.177481901 |
| P0DTN0 | OPG002 | -1.86031062 | 2.913817753 |
| Q8JZS6 | N4bp2l2 | -1.860313415 | 2.427053783 |
| Q9D975 | Srxn1 | -1.871771951 | 2.242861538 |
| P17751 | Tpi1 | -1.872319134 | 5.396543964 |
| Q80XC3 | Usp6nl | -1.872675342 | 3.405424105 |
| Q9JL26 | Fmnl1 | -1.876648297 | 2.297920977 |
| Q9CQF0 | Mrpl11 | -1.8798536 | 3.30312095 |
| Q9CQF8 | Mrpl57 | -1.886328033 | 2.529489142 |
| A0A7H0DMZ8 | OPG022 | -1.886978261 | 3.592536103 |
| Q9EPE9 | Atp13a1 | -1.895925936 | 2.806190992 |
| Q8BYB9 | Poglut1 | -1.89969772 | 3.41266248 |
| Q3TNA1 | Xylb | -1.899861769 | 2.517478028 |
| Q5ND34 | Wdr81 | -1.907650075 | 2.737260358 |
| F8VPU2 | Farp1 | -1.910961197 | 2.806190992 |
| Q9CZS3 | Cep20 | -1.911189051 | 3.170821894 |
| P21271;Q99104 | Myo5b | -1.915535554 | 4.482405687 |
| Q9EPQ7 | Stard5 | -1.921760454 | 2.110033383 |
| Q9CYA0 | Creld2 | -1.922759702 | 2.903969406 |
| Q9JJ11 | Tacc3 | -1.926015405 | 2.835544336 |
| Q3UMR5 | Mcu | -1.929433481 | 4.012821866 |
| Q5SVD0 | Rflnb | -1.929987266 | 2.46244566 |
| P01901;P14426;P14429;P14430;P14431 | H2-Q | -1.93083434 | 2.187613117 |
| Q9D2R0 | Aacs | -1.934181171 | 3.957686009 |
| Q6P4T0 | Atg2a | -1.954748905 | 3.432450438 |
| Q920L1 | Fads1 | -1.967505775 | 2.188077938 |
| P10518 | Alad | -1.967692142 | 4.265015616 |
| Q8K1L5 | Ppp1r11 | -1.970927933 | 2.211534341 |
| Q8VEB4 | Pla2g15 | -1.9824969 | 3.424108929 |
| Q69ZF3 | Gba2 | -1.983013057 | 3.199205131 |
| Q07113 | Igf2r | -1.984759803 | 2.811415814 |
| Q9QUR8 | Sema7a | -1.984925467 | 3.163816379 |
| Q8K4L3 | Svil | -1.991051222 | 2.038927622 |
| Q60790 | Rasa3 | -1.995083068 | 4.552991571 |
| Q8R127 | Sccpdh | -2.002633588 | 2.013639508 |
| Q8R3C6 | Rbm19 | -2.008634913 | 3.538091003 |
| Q3U2U7 | Mettl17 | -2.008919893 | 2.934103671 |
| Q0VEE6 | Znf800 | -2.009314432 | 2.377472486 |
| Q9QWR8 | Naga | -2.010862869 | 3.064245135 |
| P0CG15 | Chtf8 | -2.01106326 | 2.407484819 |
| O09130 | Nfatc2ip | -2.013804515 | 3.159174208 |
| Q9CR56 | Nkiras2 | -2.015446108 | 2.285307083 |
| O54818 | Tpd52l1 | -2.018506376 | 2.464139312 |
| Q7TN02 | Med26 | -2.021865159 | 3.797352736 |
| Q8R4K2 | Irak4 | -2.022122299 | 2.077090045 |
| Q9Z247 | Fkbp9 | -2.022928156 | 4.920451005 |
| Q62465 | Vat1 | -2.023099371 | 4.054244187 |
| P16858 | Gapdh | -2.02572807 | 4.937696224 |
| Q91V61 | Sfxn3 | -2.036326359 | 3.205822435 |
| Q9CWV6 | Prkrip1 | -2.03906514 | 4.89725508 |
| Q8C006 | Trim35 | -2.039610658 | 3.213187808 |
| Q60597 | Ogdh | -2.045711929 | 3.954965996 |
| Q8BHF7 | Pgs1 | -2.04612369 | 3.13319636 |
| Q9D8Y1 | Tmem126a | -2.05833134 | 2.598388611 |
| A0A7H0DNG2 | OPG204 | -2.060558226 | 3.984452126 |
| A0A7H0DND8 | OPG170 | -2.06461735 | 3.439486415 |
| Q64521 | Gpd2 | -2.065360086 | 4.525141712 |
| Q3TCN2 | Plbd2 | -2.066413153 | 3.322891625 |
| Q8N7N5 | Dcaf8 | -2.077638277 | 3.284104295 |
| Q8CDN6 | Txnl1 | -2.079427706 | 4.638422705 |
| P27773 | Pdia3 | -2.082429595 | 4.289311731 |
| O35711;Q8C8U0 | Ppfibp2 | -2.083325483 | 4.319684118 |
| Q62356 | Ftsl1 | -2.090547948 | 2.433910954 |
| Q07417 | Acads | -2.095376716 | 4.245810515 |
| Q8BG67 | Efr3a | -2.097352742 | 2.478357657 |
| P21107;P58771;P58774;Q6IRU2 | Tpm4 | -2.103622641 | 4.801316868 |
| C0HKE1;C0HKE2;C0HKE3;C0HKE4;C0HKE5;C0HKE6;C0HKE7;C0HKE8;C0HKE9;Q8BFU2;Q8CGP5;Q8CGP6;Q8CGP7;Q8R1M2 | H2ac | -2.104699526 | 3.69136396 |
| Q8BFW7 | Lpp | -2.108087162 | 4.057208694 |
| P38060 | Hmgcl | -2.113442707 | 2.705274173 |
| Q9R008 | Mvk | -2.114543697 | 4.971761795 |
| Q8BMB0 | Emsy | -2.127686279 | 2.582207783 |
| P31750 | Akt1 | -2.127847803 | 3.490650649 |
| Q8C804 | Spice1 | -2.128867559 | 2.648271745 |
| P50544 | Acadvl | -2.131406601 | 3.641463432 |
| O08553;P97427;Q62188 | Crmp1 | -2.133965932 | 3.41069737 |
| A2AGH6 | Med12 | -2.138470773 | 2.628231856 |
| O54784 | Dapk3 | -2.138999928 | 2.044781506 |
| Q8R5C5 | Actr1b | -2.139877761 | 5.396543964 |
| Q0GNC1 | Inf2 | -2.143630308 | 3.984452126 |
| P47754 | Preb | -2.143939944 | 5.374374854 |
| Q9WUQ2 | Znf451 | -2.149157755 | 2.216846934 |
| Q8C0P7 | Med12 | -2.150012488 | 2.288396779 |
| A0A7H0DN16 | OPG043 | -2.154020741 | 5.319916405 |
| Q61739 | Itga6 | -2.161369182 | 3.265922864 |
| Q8CI03 | Flywch1 | -2.164973503 | 2.833312305 |
| Q9EPK5 | Wwtr1 | -2.17615479 | 2.140541111 |
| P02469 | Lamb1 | -2.176230473 | 4.104706659 |
| Q60695 | Rgl1 | -2.180740238 | 2.498156125 |
| Q08093 | Cnn2 | -2.181527535 | 5.221563914 |
| Q9JI08 | Bin3 | -2.18159083 | 2.570389087 |
| Q8BNV1 | Trmt2a | -2.182993398 | 3.71025346 |
| O70348 | Dxo | -2.185057275 | 2.860120163 |
| Q80U35 | Arhgef17 | -2.186621702 | 3.272591953 |
| Q80TS3 | Adgrl3 | -2.188646247 | 3.584440137 |
| Q8BZT9 | Lacc1 | -2.191784426 | 2.472161899 |
| Q9CY34;Q3UWQ3 | Ube2fb | -2.192336633 | 2.904477274 |
| Q9ER38 | Tor3a | -2.193160069 | 2.094905564 |
| Q6NXW6 | Rad17 | -2.204389321 | 3.508171925 |
| Q9D0Y8 | Mrpl52 | -2.20461107 | 2.923260866 |
| Q9CQA0 | Cenpm | -2.210379019 | 3.010073237 |
| Q8BUY5 | Timmdc1 | -2.212827642 | 2.51092124 |
| P00375 | Dhfr | -2.216323691 | 5.11684144 |
| Q9CPQ5 | Cenpq | -2.217697135 | 2.561019011 |
| Q60716 | P4ha2 | -2.220024483 | 4.176105219 |
| Q9WTI7 | Myo1c | -2.220832139 | 4.417019114 |
| Q9DBY0 | Foxp4 | -2.221833351 | 2.167669092 |
| O54724 | Cavin1 | -2.223040805 | 3.531831991 |
| P17809 | Slc2a1 | -2.22693907 | 2.5641281 |
| Q9DB42 | Znf593 | -2.229351872 | 4.845499921 |
| Q8BHA0 | Ino80c | -2.230962334 | 3.867651428 |
| D0QMC3;P0DOV1;P0DOV2;Q8CGE8 | Mndal | -2.236725607 | 2.207082521 |
| Q80WW9 | Ddrgk1 | -2.240362097 | 2.42165276 |
| Q8BUH8 | Senp7 | -2.252218448 | 2.121874241 |
| Q9EPK6 | Sil1 | -2.253895077 | 2.976040341 |
| Q6P4S6 | Sik3 | -2.261812499 | 4.504982838 |
| Q9CR16 | Ppid | -2.263880103 | 5.076691685 |
| A0A7H0DN33 | OPG060 | -2.267397109 | 2.060359839 |
| Q60715 | P4ha1 | -2.268916732 | 4.470482531 |
| P23242 | Gja1 | -2.273322296 | 2.915333325 |
| A0A7H0DN28 | OPG055 | -2.273365401 | 4.665047678 |
| P97329 | Kif20a | -2.279299141 | 4.770682066 |
| Q6P6I6 | Polr2m | -2.281413143 | 2.450296339 |
| Q9CZT6 | Cmss1 | -2.289442848 | 4.448560909 |
| Q3TGF2 | Fam107b | -2.290164977 | 4.89725508 |
| Q8C854 | Myef2 | -2.290976572 | 5.123103562 |
| Q91ZS8 | Adarb1 | -2.304822718 | 2.836734866 |
| Q7TSJ2 | Map6 | -2.310356018 | 3.132348999 |
| Q8CIL4 | Fsaf1 | -2.313324726 | 2.716874543 |
| Q8BZB2 | Ppcdc | -2.321499054 | 2.619031226 |
| A0A7H0DNF5 | OPG193 | -2.322302088 | 2.358680433 |
| Q3TH73 | Ttyh2 | -2.323011118 | 2.016767463 |
| Q61387 | Cox7a2l | -2.325725502 | 2.535621364 |
| Q91XB0 | Trex1 | -2.328386229 | 3.307197121 |
| Q9CZ09 | Mettl18 | -2.329748011 | 2.286730459 |
| Q9CQW9 | Ifitm3 | -2.338273129 | 2.634997802 |
| Q8K3Z9 | Pom121 | -2.339443486 | 2.232665505 |
| P14404 | Mecom | -2.353329647 | 2.032851887 |
| Q3TJ91 | Llgl2 | -2.353581998 | 3.233389828 |
| Q6ZPF4 | Fmnl3 | -2.35761563 | 2.180449607 |
| Q9D820 | Prorsd1 | -2.361202131 | 2.313039325 |
| Q8VE96 | Slc35f6 | -2.364913873 | 2.072528361 |
| P09055 | Itgb1 | -2.368758906 | 3.537763843 |
| P07356 | Anxa2 | -2.373411799 | 2.722459648 |
| Q6ZPT1 | Klhl9 | -2.380027835 | 2.027387533 |
| Q8BK75 | Elp6 | -2.385968327 | 2.209314884 |
| Q8R0F3 | Sumf1 | -2.387359146 | 3.442540867 |
| Q80TA9 | Epg5 | -2.38906871 | 3.02812628 |
| O88667 | Rrad | -2.392463838 | 2.562676697 |
| Q99JP6 | Homer3 | -2.396619 | 3.15675784 |
| Q9WTR5 | Cdh13 | -2.401500867 | 3.448504382 |
| Q80WE4 | Kif20b | -2.416207038 | 4.50536728 |
| Q9CZI9 | Aen | -2.417121512 | 3.115382909 |
| A3KGW5 | Cercam | -2.42109136 | 2.447789608 |
| Q9R0X4;Q32MW3 | Acot | -2.423655265 | 3.709703699 |
| Q61165 | Slc9a1 | -2.434289364 | 2.303458108 |
| Q8BYW9 | Eogt | -2.439295174 | 4.224903831 |
| Q9DBR0 | Akap8 | -2.450531878 | 3.767525233 |
| Q9CXX9 | Cuedc2 | -2.465524063 | 2.400131292 |
| Q6AW69 | Cgnl1 | -2.467209809 | 2.688460386 |
| Q99KW3 | Triobp | -2.46913934 | 4.448560909 |
| Q61127 | Nab2 | -2.472703588 | 5.396543964 |
| Q3URQ7 | Mthfsd | -2.47409345 | 3.20511094 |
| Q99JV5 | Stard4 | -2.474846712 | 3.095062935 |
| Q69ZN7 | Myof | -2.475808833 | 3.004783786 |
| Q9CPW9 | Metap1d | -2.476636291 | 3.041645729 |
| Q8BGV4 | Tti2 | -2.481143319 | 2.091056947 |
| Q61490 | Alcam | -2.481541234 | 4.614254512 |
| O54791 | Maff | -2.482896032 | 3.527688739 |
| Q9CU65;Q3U2E2 | Zmym5 | -2.487672929 | 3.271203322 |
| P17515 | Cxcl10 | -2.491093922 | 2.311716527 |
| Q9R210 | Tfeb | -2.494552859 | 2.947130775 |
| P43275 | H1-1 | -2.495819652 | 2.207517989 |
| Q80X85 | Mrps7 | -2.507582381 | 2.092769002 |
| Q03145 | Epha2 | -2.514748052 | 4.121524539 |
| Q920A5 | Scpep1 | -2.518445719 | 4.962269298 |
| Q9DBV3 | Dhx34 | -2.52103948 | 2.245765358 |
| Q8BR90 | Rimoc1 | -2.523200428 | 2.120647794 |
| Q9DD24 | Tceal9 | -2.523542408 | 2.014702039 |
| Q8R151 | Znfx1 | -2.526539431 | 2.060359839 |
| Q8BUE4 | Aifm2 | -2.527890725 | 2.126266342 |
| Q80TZ3;Q99KY4 | Dnajc6 | -2.536823826 | 3.537763843 |
| Q91YI0 | Asl | -2.541526612 | 6.015023095 |
| P56375 | Acyp2 | -2.541612897 | 3.098190734 |
| Q62136 | Ptpn21 | -2.541993667 | 2.303458108 |
| O08911 | S100a11 | -2.549039734 | 2.061042081 |
| Q8R003 | Mapk12 | -2.552135167 | 2.230947269 |
| Q6PDX6 | Rnf220 | -2.553173012 | 3.838459094 |
| Q8CGB6 | Tns2 | -2.559044452 | 2.350343013 |
| P01897;P01899 | H2 | -2.559218948 | 2.231380055 |
| Q571F8 | Gls2 | -2.562344645 | 2.400396094 |
| P23780;Q8VC60 | Glb1 | -2.562582811 | 3.070760511 |
| Q9CWY4 | Gemin7 | -2.566452002 | 2.186669322 |
| P59110 | Senp1 | -2.568283564 | 2.976916077 |
| Q8CIE4 | Parp10 | -2.568475225 | 2.962501984 |
| Q9CR70 | Lage3 | -2.568790105 | 2.210518095 |
| Q7TPV2 | Dzip3 | -2.574799308 | 2.114301441 |
| P14901 | Hmox1 | -2.577246947 | 2.640241439 |
| Q8K298 | Anln | -2.578150435 | 4.309619089 |
| Q7TPM9 | Mtmr10 | -2.578968395 | 3.576564595 |
| P39053 | Dnm1 | -2.585971081 | 3.673949422 |
| Q9JIA7 | Sphk2 | -2.592364293 | 2.567765788 |
| Q80UY1 | Carnmt1 | -2.593319543 | 2.821512195 |
| Q9D0K2 | Oxct1 | -2.595133395 | 4.770682066 |
| Q9QXS1 | Plec | -2.595807084 | 4.237499095 |
| Q8CDM1 | Atad2 | -2.596181034 | 4.831912034 |
| Q6RUT7 | Uqcc4 | -2.605622729 | 2.03392213 |
| Q3TLS3 | Gdpgp1 | -2.619901683 | 2.068711412 |
| O54926 | Siva1 | -2.625925639 | 2.332054175 |
| P28033 | Cebpb | -2.628128693 | 2.508494782 |
| Q9D771 | Pacc1 | -2.628410294 | 2.621180355 |
| Q8K1K4 | Cenpi | -2.631665323 | 2.437394605 |
| Q8CJ40 | Crocc | -2.635470774 | 2.269770297 |
| O08539 | Bin1 | -2.637589917 | 4.614254512 |
| P02340 | Tp53 | -2.63773164 | 2.732694338 |
| Q8BH04 | Pck2 | -2.641479833 | 3.387539163 |
| P22935 | Crabp2 | -2.649932777 | 4.603647461 |
| Q64112 | Ifit2 | -2.656346153 | 2.274539999 |
| Q8BGR8 | Gskip | -2.656885048 | 2.32049535 |
| Q920A7 | Afg3l1 | -2.656988503 | 3.586261451 |
| Q3UJP5 | Cfap418 | -2.658111713 | 2.124648807 |
| Q7TSJ6 | Lats2 | -2.673066153 | 2.050180309 |
| O70309 | Itgb5 | -2.676932336 | 2.538926105 |
| Q91YT8 | Tmem63a | -2.682095195 | 2.091219084 |
| P11679;P15331;P20152;P31001;Q9DCV7 | Krt8 | -2.683184556 | 3.676659831 |
| Q69ZI1 | Sh3rf1 | -2.683805787 | 2.303458108 |
| Q99N84 | Mrps18b | -2.690704531 | 3.170821894 |
| Q5EE38 | Acd | -2.696432783 | 2.388897278 |
| Q8VCF0 | Mavs | -2.699358749 | 3.304278574 |
| P61028 | Rab8b | -2.706041044 | 2.657493006 |
| Q80YR6 | Rbbp8 | -2.706481055 | 2.729387407 |
| Q9DCH2 | Pop7 | -2.708834814 | 3.638277906 |
| A0A7H0DN00 | OPG023 | -2.711509695 | 5.373699526 |
| P47930 | Fosl2 | -2.711977711 | 5.123103562 |
| Q9CZT5 | Vasn | -2.712366415 | 3.293188608 |
| P14719 | Il1rl1 | -2.716760817 | 3.699556757 |
| P11214 | Plat | -2.721781751 | 2.18427048 |
| P07214 | Sparc | -2.724090681 | 4.237369263 |
| P08207 | S100a10 | -2.724268142 | 3.335427287 |
| Q8VC70 | Rbms2 | -2.727240422 | 2.594063476 |
| Q8VCH8 | Ubxn4 | -2.729869186 | 2.805425296 |
| P61025 | Cks1b | -2.736972908 | 4.42846902 |
| Q6PDY2 | Ado | -2.737271356 | 4.495282144 |
| Q8BSL7 | Arf2 | -2.745200267 | 3.379367716 |
| E9Q286 | Ice1 | -2.749608946 | 2.533625915 |
| Q8CAB8 | Castor2 | -2.752915814 | 2.151797823 |
| P22366 | Myd88 | -2.753270495 | 3.348189879 |
| Q9ESY9 | Ifi30 | -2.766222409 | 3.981051698 |
| Q80V94 | Ap4e1 | -2.767133297 | 3.638277906 |
| A2A7S8 | Nhsl3 | -2.776664641 | 2.804365214 |
| P18406 | Ccn1 | -2.784710255 | 2.601246575 |
| Q9EQK5 | Mvp | -2.787380851 | 3.158413485 |
| O89086 | Rbm3 | -2.792145829 | 5.012861156 |
| Q8BGD8 | Coa6 | -2.792593093 | 3.293192357 |
| Q9DB73 | Cyb5r1 | -2.793428923 | 2.202341331 |
| E9PY46 | Ift140 | -2.794082234 | 2.16577023 |
| Q8BTI9 | Pik3cb | -2.79434711 | 2.575872946 |
| Q8C181 | Mbnl2 | -2.796333309 | 4.748067278 |
| Q9WTK5 | Nfkb2 | -2.799156785 | 4.401166509 |
| Q61391 | Mme | -2.800384498 | 2.607985672 |
| P21300;P45377 | Ark1b | -2.805094652 | 3.03849206 |
| O08605 | Mknk1 | -2.80518512 | 3.584289567 |
| Q9CQX4 | Pclaf | -2.811361773 | 2.795533743 |
| Q5PSV9 | Mdc1 | -2.812032688 | 3.987319542 |
| Q9DCX1 | Mad2l1bp | -2.812327063 | 2.865957562 |
| Q9DCT5 | Sdf2 | -2.816889416 | 2.201015351 |
| Q91XV3 | Basp1 | -2.817994763 | 2.756537818 |
| Q9CQJ1 | Higd2a | -2.820817673 | 2.885477621 |
| O35657 | Neu1 | -2.821146018 | 4.311461067 |
| P70279 | Surf6 | -2.840583194 | 2.979866261 |
| Q8VC34 | Rpap2 | -2.842443827 | 2.516991292 |
| Q99MS8 | Tpgs1 | -2.850105908 | 2.428334118 |
| Q62470 | Itga3 | -2.850335916 | 2.759756136 |
| Q9EP53 | Tsc1 | -2.853113861 | 3.135420387 |
| Q91ZX7 | Lrp1 | -2.855881567 | 3.220594607 |
| Q922E6 | Fastkd2 | -2.857522406 | 3.639886829 |
| E9PVX6 | Mki67 | -2.860104613 | 2.174821446 |
| P06869 | Plau | -2.861449352 | 2.362591623 |
| Q8K0B2 | Lmbrd1 | -2.86366637 | 2.037876594 |
| Q9D142 | Nudt14 | -2.865853995 | 2.11484364 |
| P20357 | Map2 | -2.867907689 | 2.668604774 |
| Q8VCH0;Q921H8 | Acaa1b | -2.869045068 | 2.124925168 |
| A2AI08 | Tprn | -2.870807032 | 2.146618738 |
| P70671 | Irf3 | -2.872023861 | 2.758006104 |
| A2ACJ2 | Faap100 | -2.872343767 | 2.144793083 |
| Q9CWT6 | Ddx28 | -2.87535282 | 3.282057064 |
| P30412 | Ppic | -2.887187891 | 2.397847369 |
| D3Z4S3 | Ptrhd1 | -2.888639788 | 2.210689202 |
| P22437 | Ptgs1 | -2.892981553 | 3.810489334 |
| Q9QXW0 | Fbxl6 | -2.896011721 | 2.263489589 |
| P70451 | Fer | -2.897649267 | 2.656222168 |
| Q99JR5 | Tinagl1 | -2.902243888 | 2.852332223 |
| Q99LJ7 | Rcbtb2 | -2.902503092 | 3.019023631 |
| Q99L00 | Haus8 | -2.907522857 | 2.282595242 |
| Q3UGP9 | Lrrc58 | -2.907584413 | 2.09605943 |
| Q61090 | Fzd7 | -2.915722582 | 2.455615275 |
| Q80U49 | Cep170b | -2.918024806 | 2.680374262 |
| Q8BM55 | Tmem214 | -2.919101804 | 2.34796011 |
| Q07243 | Mtf1 | -2.92655198 | 2.210607724 |
| Q8K2I2 | Cchcr1 | -2.92891783 | 2.137672693 |
| Q8BMI4 | Gen1 | -2.930141682 | 2.757380554 |
| Q8R550 | Sh3kbp1 | -2.937079645 | 5.076691685 |
| Q9QZI8 | Serinc1 | -2.939003361 | 2.42593175 |
| O88986 | Gcat | -2.951429599 | 5.221563914 |
| D3YXK1 | Samd1 | -2.953572916 | 3.329731918 |
| Q04899 | Cdk18 | -2.955410311 | 2.357869642 |
| Q6AXC6 | Ddx11 | -2.956466323 | 2.387616273 |
| Q9JJX7 | Tdp2 | -2.962630224 | 3.23542358 |
| Q8BXA1 | Golim4 | -2.965220466 | 2.475313853 |
| Q04519 | Smpd1 | -2.966608377 | 2.613564548 |
| Q8VEB2 | Sav1 | -2.980712661 | 2.322731947 |
| Q8BQ47 | Cnpy4 | -2.985630736 | 4.89725508 |
| Q9DBR4 | Apbb2 | -2.987834102 | 3.052209902 |
| Q9CPY3 | Cdca5 | -2.992376821 | 2.615280044 |
| Q9CQZ1 | Hsbp1 | -2.996936734 | 2.370467396 |
| Q5SUE8 | Ankrd40 | -2.999396122 | 2.051239096 |
| Q8C0Z1 | Fam234a | -3.002489049 | 3.409818611 |
| Q8VCP8 | Ak6 | -3.008817356 | 2.944476753 |
| Q3TW96 | Uap1l1 | -3.011056521 | 5.079252495 |
| Q8K1I7 | Wipf1 | -3.011740704 | 2.405762451 |
| Q8BGK6 | Slc7a6 | -3.012260811 | 2.081345769 |
| P48967 | Cdc25c | -3.012910873 | 3.390346495 |
| Q5XG73 | Acbd5 | -3.016270625 | 2.472730001 |
| O35988 | Sdc4 | -3.01718866 | 2.322096162 |
| Q8R080 | Gtse1 | -3.018798154 | 2.841565638 |
| E9QAM5 | Helz2 | -3.018805395 | 3.621386629 |
| Q8R216 | Sirt4 | -3.020186754 | 2.840578782 |
| Q9D8S9 | Bola1 | -3.027397468 | 3.746285395 |
| Q9Z223 | Mocs2 | -3.035454681 | 2.550006804 |
| P43883 | Plin2 | -3.039837832 | 2.197253812 |
| O88322 | Nid2 | -3.0450318 | 2.804591016 |
| Q3UQ28 | Pxdn | -3.046536063 | 2.070963561 |
| Q8CG19 | Ltbp1 | -3.047636379 | 3.861419064 |
| Q8CC35 | Synpo | -3.057286069 | 2.955247479 |
| Q6S5J6 | Krit1 | -3.058856967 | 2.758323834 |
| Q61510 | Trim25 | -3.060638391 | 2.249233923 |
| P0DPD9;P0DPE0 | Eef1akmt4 | -3.062481696 | 2.256344679 |
| P48432 | Sox2 | -3.067807528 | 2.369450875 |
| Q9JJ89 | Ccdc86 | -3.072187954 | 3.851287414 |
| L0N7N1 | Kif14 | -3.072286353 | 2.11838146 |
| Q640N1 | Aebp1 | -3.079529948 | 2.858562347 |
| Q64092 | Tfe3 | -3.082820687 | 2.672424221 |
| Q9JL15 | Lgals8 | -3.08681987 | 2.155259866 |
| Q9CPY0 | Mrm2 | -3.087564021 | 3.027136856 |
| Q8CHI8 | Ep400 | -3.088993404 | 2.407484819 |
| Q9CRA7 | Dmac2l | -3.094890307 | 2.207517989 |
| Q8BK58 | Hspbap1 | -3.096928405 | 4.79352438 |
| P29268 | Ccn2 | -3.098366669 | 4.524952501 |
| P97864 | Casp7 | -3.101555263 | 2.155259866 |
| P70677 | Casp3 | -3.102352875 | 4.375749023 |
| P70213 | Fv1 | -3.103464571 | 2.544719253 |
| Q8CE46 | Pus7l | -3.10627565 | 3.233389828 |
| Q9WTS2 | Fut8 | -3.111580349 | 3.230529342 |
| Q8VE38 | Oxnad1 | -3.112172541 | 3.191697306 |
| Q8R2X8 | Blzf1 | -3.114714745 | 2.340383875 |
| P81122 | Irs2 | -3.119340101 | 2.510314281 |
| P03930 | Mtatp8 | -3.121570162 | 3.301726151 |
| P31955 | Areg | -3.129766493 | 3.574330547 |
| Q3U155 | Ccdc174 | -3.132312866 | 2.214333282 |
| P58059 | Mrps21 | -3.134367067 | 3.55288282 |
| P56542 | Dnase2 | -3.139559428 | 3.851287414 |
| C0HKG5;C0HKG6 | Rnaset2 | -3.143060746 | 3.032740992 |
| Q99LC9 | Pex6 | -3.14384859 | 2.251434636 |
| Q921Q7 | Rin1 | -3.150777379 | 2.066770354 |
| P70193 | Lrig1 | -3.15388017 | 3.387073777 |
| Q9CXH7 | Sgo1 | -3.156437791 | 4.341982222 |
| Q8R0W0;Q9QXS1 | Eppk1 | -3.160318115 | 4.146679376 |
| P82343 | Renbp | -3.16464088 | 4.854481453 |
| Q99KW9 | Itfg1 | -3.164946008 | 2.765648107 |
| P49935 | Ctsh | -3.169342313 | 3.078839402 |
| P06795 | Abcb1b | -3.175117031 | 2.032851887 |
| P41731 | Cd63 | -3.19192901 | 2.108044661 |
| P13020 | Gsn | -3.194676006 | 4.463939346 |
| Q6PG16 | Hjurp | -3.198487331 | 4.784426578 |
| Q99MK8 | Grk2 | -3.209117207 | 2.949325137 |
| Q9CQY2 | Ramac | -3.211261853 | 3.906419724 |
| Q9CR75 | Tnfrsf12a | -3.214064067 | 3.170392126 |
| Q8CDJ8 | Ston1 | -3.217132214 | 2.771084148 |
| Q8CIG9 | Fbxl8 | -3.218455677 | 2.859419819 |
| Q9CYH2 | Prxl2a | -3.222047968 | 2.827908648 |
| Q8CBY1 | Samd4a | -3.222236384 | 3.70641885 |
| Q6A0A9;Q8C3F2 | Fam120c | -3.22275455 | 2.210518095 |
| Q8R1G6 | Pdlim2 | -3.232781091 | 2.238471782 |
| Q9WVJ9 | Efemp2 | -3.234913129 | 2.683299832 |
| Q9D9M5 | Phospho2 | -3.239896198 | 2.857635802 |
| P58058 | Nadk | -3.244906956 | 3.095744381 |
| Q3V4B5 | Commd6 | -3.249349959 | 3.273064116 |
| Q64449 | Mrc2 | -3.251947066 | 2.925356735 |
| Q64701 | Rbl1 | -3.252499506 | 2.324688417 |
| Q9R118 | Htra1 | -3.253132514 | 3.27178775 |
| Q80ST9 | Lca5 | -3.256511726 | 2.099068243 |
| Q64337 | Sqstm1 | -3.257662308 | 5.396543964 |
| Q66GT5 | Ptpmt1 | -3.265232938 | 2.589529301 |
| Q9Z0Z3 | Skp2 | -3.266582095 | 2.197253812 |
| P23927 | Cryab | -3.267379106 | 4.184940346 |
| Q9QWF0 | Chaf1a | -3.267729731 | 2.41253279 |
| Q3TL26 | Tfb2m | -3.274384502 | 2.757380554 |
| Q8CB62 | Cntrob | -3.275442628 | 3.107017811 |
| B2RXV4 | Flvcr1 | -3.282870971 | 2.358736505 |
| Q99J47 | Dhrs7b | -3.299240091 | 3.825137647 |
| Q62172 | Ralbp1 | -3.308835033 | 2.621005294 |
| Q7TME2 | Spag5 | -3.309479185 | 3.434148701 |
| Q9CWY9 | Rpain | -3.310128276 | 2.792192728 |
| Q61554 | Fbn1 | -3.31198507 | 2.705942438 |
| O89051 | Itm2b | -3.31271639 | 2.176762317 |
| O35375 | Nrp2 | -3.314076241 | 2.807447192 |
| P0C8B4 | Gon7 | -3.327998727 | 2.911801121 |
| P97863 | Nfib | -3.32951403 | 3.602814454 |
| Q9D6J5 | Ndufb8 | -3.331033271 | 2.178504997 |
| Q9D1M7 | Fkbp11 | -3.333543959 | 2.300326574 |
| Q91XC0 | Ajuba | -3.334186528 | 2.987934867 |
| Q8CJ26 | Nradd | -3.334510856 | 2.646631443 |
| O88207 | Col5a1 | -3.336996007 | 2.304819018 |
| Q62187 | Ttf1 | -3.340344039 | 4.063243692 |
| Q99KC8 | Vwa5a | -3.341466872 | 2.579569481 |
| P28650 | Adss1 | -3.345988005 | 3.220594607 |
| Q8C080 | Snx16 | -3.350098682 | 2.038077891 |
| Q66T02 | Plekhg5 | -3.354569015 | 2.533240806 |
| Q923K4 | Gtpbp3 | -3.360039445 | 2.834502286 |
| B1ARD6 | Slfn9 | -3.361641863 | 3.724672543 |
| P50429 | Arsb | -3.364867262 | 3.599184677 |
| P97465 | Dok1 | -3.365183628 | 2.031070715 |
| Q3UMW8 | Cln5 | -3.378614562 | 3.236128002 |
| Q6P3B9 | Rbfa | -3.381295262 | 3.527986203 |
| B1ARD8;B1ARD6 | Slnf8 | -3.387499612 | 3.183423764 |
| P11352 | Gpx1 | -3.390897846 | 2.082751335 |
| Q3UDP0 | Wdr41 | -3.402395351 | 2.433910954 |
| Q9D842 | Aplf | -3.405749335 | 2.624820406 |
| Q6NXJ0 | Wwc2 | -3.417374924 | 3.198962116 |
| Q9JL19 | Ncoa6 | -3.419459822 | 2.989044105 |
| Q8K2X3 | Stn1 | -3.420669337 | 4.552426484 |
| Q8BGA9 | Oxa1l | -3.424940083 | 2.967737016 |
| O08901 | Bub1 | -3.425837038 | 2.867327412 |
| P12265 | Gusb | -3.430497315 | 4.059076307 |
| Q9CY21 | Bud23 | -3.433829626 | 2.393983058 |
| Q9CXP8 | Gng10 | -3.444553285 | 3.301258443 |
| Q80WC7 | Agfg2 | -3.446364282 | 2.002667464 |
| Q9R045 | Angptl2 | -3.452165178 | 2.018609579 |
| Q3V1H1 | Ckap2 | -3.454776225 | 3.093163664 |
| P70255 | Nfic | -3.456196214 | 2.459671448 |
| P47955 | Rplp1 | -3.456279705 | 2.773645908 |
| P56390 | Cks2 | -3.456320435 | 2.285361188 |
| Q8BFQ9 | Klhl42 | -3.462223813 | 2.998780928 |
| Q99MB2 | Mtfr1 | -3.468939138 | 2.827908648 |
| Q62179 | Sema4b | -3.469008501 | 2.514927893 |
| Q9D3D9 | Atp5f1d | -3.474608663 | 2.187613117 |
| Q8VC19 | Alas1 | -3.475310143 | 2.86922444 |
| Q8BZ20 | Parp12 | -3.475918906 | 3.027287353 |
| Q8BMA5 | Npat | -3.478180457 | 2.950403017 |
| Q8K4E0 | Alms1 | -3.478874245 | 2.101798941 |
| O70161 | Pip5k1c | -3.486473526 | 2.045354757 |
| Q9CQL7 | Mrfap1 | -3.491734745 | 2.861838547 |
| Q9DCT1 | Akr1e2 | -3.500013589 | 3.482910049 |
| Q99P69 | Nuf2 | -3.501191835 | 3.343972894 |
| Q8VDG7 | Pafah2 | -3.505210079 | 3.153634325 |
| Q8K354 | Cbr3 | -3.505475226 | 2.504905142 |
| Q9DBM1 | Gpatch1 | -3.50711751 | 2.830050823 |
| O55229 | Chkb | -3.50932188 | 2.103035021 |
| O09044;P60879 | Snap | -3.513289412 | 3.398995898 |
| O55101 | Syngr2 | -3.513612924 | 3.325177398 |
| Q8BW00 | Ptrh1 | -3.513657909 | 2.222788319 |
| Q9D8U0 | Ifrd2 | -3.514247962 | 2.037501142 |
| P54761 | Ephb4 | -3.520380658 | 2.727708125 |
| P70271 | Pdlim4 | -3.532463466 | 4.044096318 |
| Q8VCR7 | Abhd14b | -3.535347275 | 3.519888891 |
| Q9QYM8 | Cenph | -3.551236157 | 3.225911988 |
| Q9CQK1 | Znhit3 | -3.551416514 | 3.808527211 |
| Q9WVL3 | Slc12a7 | -3.554753032 | 2.441829717 |
| Q8BZA9 | Tigar | -3.560209018 | 2.926513144 |
| Q6Y685 | Tacc1 | -3.56032316 | 3.128065679 |
| Q08775 | Runx2 | -3.565779887 | 2.207517989 |
| P10605 | Ctsb | -3.56909565 | 3.91312523 |
| P59759 | Mrtfb | -3.573111096 | 3.13008838 |
| Q9CQ79 | Txndc9 | -3.576951287 | 2.207267155 |
| P14069 | S100a6 | -3.578808464 | 3.797352736 |
| P26645 | Marcks | -3.5975768 | 3.861784354 |
| Q52KR3 | Prune2 | -3.597898049 | 2.229964157 |
| Q9CQ91 | Ndufa3 | -3.600036787 | 2.188684889 |
| Q8K1S3 | Unc5b | -3.602824961 | 2.647219907 |
| Q3TEL6;Q9D074 | Rnf157;Mgm | -3.620887055 | 2.02395183 |
| Q9CR02 | Tma16 | -3.624933487 | 2.829147331 |
| Q9CZB0 | Sdhc | -3.62497813 | 2.490057776 |
| Q9CPU4 | Mgst3 | -3.625905383 | 3.269778293 |
| P08074 | Cbr2 | -3.627636818 | 3.495062877 |
| Q9CPP0 | Npm3 | -3.629784812 | 3.524496387 |
| Q3TUA9 | Pomk | -3.630599827 | 3.14342936 |
| P01887 | B2m | -3.632096399 | 2.063860194 |
| Q3TVC7 | Ccndbp1 | -3.639104417 | 3.072948477 |
| Q9D6Y7 | Msra | -3.640673512 | 3.163816379 |
| P56183 | Rrp1 | -3.651021403 | 3.453027624 |
| Q8K305 | Nsl1 | -3.658828956 | 2.183068573 |
| Q9QX60 | Dguok | -3.671925196 | 4.271958081 |
| P85094 | Isoc2a | -3.683181479 | 4.624204349 |
| Q6F3F9 | Adgrg6 | -3.685348078 | 3.861784354 |
| Q64364 | Cdkn2a | -3.685672229 | 3.396273085 |
| Q9DCZ1 | Gmpr | -3.694223855 | 2.170137013 |
| Q8R5A3 | Apbb1ip | -3.698911217 | 2.322937754 |
| Q9DBE0 | Csad | -3.707465366 | 4.148480313 |
| P50428 | Arsa | -3.713007418 | 4.057208694 |
| Q9WU79 | Prodh | -3.716117835 | 2.491874269 |
| P09450 | Junb | -3.717196193 | 2.67066014 |
| Q9CWY3 | Setd6 | -3.721582417 | 2.792820137 |
| Q5SUQ9 | Ctc1 | -3.735024274 | 3.211286938 |
| Q9DB28 | Pop5 | -3.74232012 | 2.625514369 |
| Q9CR46 | Ska2 | -3.74824156 | 2.356363373 |
| P97821 | Ctsc | -3.758723998 | 3.80055133 |
| Q924K8;Q9R190 | Mta | -3.760042046 | 2.162646484 |
| Q80UU2 | Rpp38 | -3.760348088 | 3.190864341 |
| Q9CQ88 | Tspan31 | -3.76325571 | 2.771084148 |
| Q8R1F6 | Hid1 | -3.763983911 | 3.207433374 |
| P0DW87 | Zftraf1 | -3.769983614 | 3.736261838 |
| Q8C1M2 | Znf428 | -3.772662277 | 2.794297039 |
| Q8BK30 | Ndufv3 | -3.776711697 | 3.78156547 |
| O09111 | Ndufb11 | -3.778853018 | 2.258725633 |
| Q8R0X7 | Sgpl1 | -3.781043655 | 3.132647159 |
| P25085 | Il1rn | -3.784720781 | 2.213614591 |
| A2AG58 | Bclaf3 | -3.785541607 | 3.987319542 |
| O54974 | Lgals7 | -3.793659388 | 4.240819544 |
| Q9D2L9 | Fam111a | -3.802054597 | 2.587052318 |
| Q8C163 | Exog | -3.804817896 | 3.512639993 |
| Q8R395 | Commd5 | -3.808865669 | 2.500491225 |
| P24638 | Acp2 | -3.809821034 | 4.854481453 |
| P03899 | mt-Nd3 | -3.815933812 | 2.118633363 |
| Q8BHA3 | Dtd2 | -3.819802525 | 2.567879683 |
| Q9CPV1 | Ska1 | -3.823453973 | 2.911801121 |
| Q8CD10 | Micu2 | -3.823680489 | 2.582695782 |
| Q99M15 | Pstpip2 | -3.826235036 | 2.677268517 |
| Q8BL80 | Arhgap22 | -3.83203655 | 3.938168629 |
| Q8R5A0 | Smyd2 | -3.833844868 | 2.268010785 |
| Q9CX53 | Gemin6 | -3.837734405 | 3.619292811 |
| Q810Q5 | Nmes1 | -3.840621847 | 2.177380478 |
| Q9D0L7 | Armc10 | -3.841986123 | 2.055088451 |
| Q9DCS2 | Mettl26 | -3.843148154 | 2.524110723 |
| P57724 | Pcbp4 | -3.844663114 | 4.083457527 |
| Q9JIM1 | Slc29a1 | -3.846856667 | 2.825911095 |
| P56389 | Cda | -3.856845701 | 2.997878126 |
| Q9Z0M5 | Lipa | -3.86179948 | 2.914309998 |
| P54818 | Galc | -3.865208053 | 2.274539999 |
| P37889 | Fbln2 | -3.867103495 | 2.314908407 |
| Q62087 | Pon3 | -3.880045466 | 2.63362868 |
| Q06335 | Aplp2 | -3.893864937 | 4.355577622 |
| Q924Z4 | Cers2 | -3.894064459 | 2.023983752 |
| Q8R2Q8 | Bst2 | -3.896701665 | 3.142356764 |
| Q8BVA5 | Ldah | -3.904508557 | 3.234619869 |
| Q6ZPG2 | Wdr90 | -3.904974989 | 3.904392999 |
| P11688 | Itga5 | -3.909456963 | 2.556486684 |
| Q64299 | Ccn3 | -3.910977009 | 2.199525381 |
| Q7TMR0 | Prcp | -3.911648091 | 2.481159532 |
| Q60772 | Cdkn2c | -3.911954902 | 3.441144696 |
| Q8K2B0 | P3h4 | -3.918532693 | 3.051624817 |
| A2AJI0 | Map7d1 | -3.927206452 | 2.49107692 |
| P05480;P06240;P39688;Q04736 | Lck;Fyn;Yes1 | -3.931382288 | 2.187613117 |
| P58468 | Slx9 | -3.932108433 | 2.586690022 |
| Q9CX66 | Nopchap1 | -3.942130741 | 2.855853664 |
| D3YZG8 | Mthfd2l | -3.943714429 | 2.732925919 |
| Q9JM96 | Cdc42ep4 | -3.943915242 | 2.624820406 |
| O08734 | Bak1 | -3.949532832 | 3.95584471 |
| Q69ZJ7 | Ric1 | -3.949559431 | 3.427994675 |
| Q3U1G5 | Isg20l2 | -3.956746117 | 2.344195598 |
| Q5DTX6 | Jcad | -3.957484218 | 2.440842988 |
| Q9Z0R9 | Fads2 | -3.957513663 | 2.850077071 |
| Q9CZX5 | Pinx1 | -3.959290053 | 2.784017085 |
| Q8BKT8 | Haus7 | -3.960096877 | 2.758323834 |
| Q61738 | Itga7 | -3.963802352 | 3.579504343 |
| Q80Z25 | Ofd1 | -3.964723445 | 2.099864419 |
| P63013 | Prrx1 | -3.972660217 | 3.173768604 |
| P21460 | Cst3 | -3.988272467 | 4.026739105 |
| Q9JHJ3 | Glmp | -3.992602389 | 2.860062223 |
| Q91ZF0 | Dnajc24 | -3.993854078 | 4.307955709 |
| Q99PP9 | Trim16 | -3.998210888 | 2.847560602 |
| P01897;P01899;P01900;P01901;P03991;P04223;P06339;P14426;P14427;P14428;P14429;P14430;P14431 | H2 | -4.005414252 | 5.396543964 |
| Q924T3 | Xrcc4 | -4.007890764 | 3.004057188 |
| P54923 | Adprh | -4.008084356 | 4.196230789 |
| Q9D3P8 | Plgrkt | -4.019788954 | 2.918724641 |
| Q8CEE6 | Pask | -4.020417423 | 2.333279465 |
| Q7TMC8 | Fcsk | -4.024381225 | 2.538733416 |
| Q5I1X5 | Ppp1r13l | -4.034206874 | 2.786158612 |
| Q8R2L5 | Mrps18c | -4.034504535 | 3.088513283 |
| Q91WK7 | Ankrd54 | -4.037575663 | 2.013716365 |
| Q9D3W4 | Gpn3 | -4.038855907 | 3.082073598 |
| Q8BP27 | Sfr1 | -4.062650602 | 2.5258886 |
| Q8R3F5 | Mcat | -4.064038823 | 2.823558969 |
| Q8K0C8 | Cox19 | -4.070748178 | 2.435246009 |
| Q8C3R1 | Brat1 | -4.080056191 | 2.584188957 |
| Q14AI6 | Rpusd3 | -4.083666771 | 2.898558299 |
| Q3UMF0 | Cobll1 | -4.086165991 | 2.414445428 |
| Q8R3P0 | Aspa | -4.088494671 | 2.178317478 |
| Q9CZX7 | Pip4p2 | -4.088951676 | 2.862815487 |
| Q9CXL3 | Chlsn | -4.091719497 | 4.166146413 |
| Q61846 | Melk | -4.100245075 | 4.341982222 |
| P27046;Q8BRK9 | Man2a | -4.101164495 | 2.396609659 |
| Q6PIU9 |  | -4.106425285 | 3.374147521 |
| Q99PG2 | Ogfr | -4.109401563 | 2.174821446 |
| Q99JP4 | Cdc26 | -4.114191082 | 3.073025804 |
| Q9CWH5 | Trmt11 | -4.115213475 | 2.384254544 |
| Q8C263 | Ska3 | -4.116018709 | 3.032916515 |
| Q8BYY4 | Ttc39b | -4.116996408 | 2.830588454 |
| Q99JZ7 | Errfi1 | -4.119685253 | 3.079967569 |
| Q9Z0R0 | Haspin | -4.122572906 | 2.19827728 |
| Q8CBY0 | Gatc | -4.124135045 | 2.322937754 |
| Q5FWI3 | Cemip2 | -4.127123741 | 2.177380478 |
| Q8BHE0 | Prr11 | -4.128859265 | 2.405762451 |
| Q9CWX2 | Ndufaf1 | -4.131855871 | 2.756586777 |
| Q8CD19 | Lancl3 | -4.133378077 | 2.93083158 |
| Q76N33 | Stambpl1 | -4.136397279 | 2.707949902 |
| Q9D7N6 | Mrpl30 | -4.13715652 | 3.082698915 |
| O09005 | Degs1 | -4.137972431 | 2.860120163 |
| Q8K1J5 | Sde2 | -4.141110842 | 3.082073598 |
| Q3V038 | Ttc9 | -4.142858253 | 2.027156257 |
| Q8VCS6 | Med9 | -4.148097634 | 2.962458667 |
| Q8BIA4 | Fbxw8 | -4.152162212 | 2.385010999 |
| Q8BWQ4 | Cmtr2 | -4.158090393 | 2.543047254 |
| P10649 | Gstm1 | -4.158735287 | 2.638166226 |
| Q921S7 | Mrpl37 | -4.161807303 | 3.478484469 |
| Q9QY53 | Nphp1 | -4.16232304 | 2.242766829 |
| Q9D009 | Lipt2 | -4.168158061 | 2.827687546 |
| Q9DC28 | Csnk1d | -4.169803461 | 3.423447522 |
| Q9CYW4 | Hdhd3 | -4.171244835 | 2.846306533 |
| Q78IK4 | Apool | -4.172519759 | 3.617119599 |
| Q8BHX3 | Cdca8 | -4.179141542 | 2.53307961 |
| Q99LB7 | Sardh | -4.180949694 | 2.161783269 |
| Q6PAT0 | Adat3 | -4.182642 | 4.899776614 |
| Q3UGC7 | Eif3j1 | -4.186157898 | 2.765648107 |
| Q6Q899 | Rigi | -4.188155785 | 2.311507773 |
| O54864 | Suv39h1 | -4.190451737 | 2.704155829 |
| Q8K1N2 | Phldb2 | -4.193862597 | 2.027354008 |
| P70444 | Bid | -4.196029675 | 3.40462166 |
| Q9CUN6 | Smurf1 | -4.196280837 | 2.513449437 |
| O88824 | Jtb | -4.197418038 | 4.525141712 |
| P24860 | Ccnb1 | -4.203540705 | 3.463538703 |
| Q9JKV5 | Scamp4 | -4.214156097 | 2.635996096 |
| P29391;P49945 | Ftl1 | -4.224947517 | 3.056109998 |
| Q8VE94 | Fam110c | -4.237284556 | 5.079252495 |
| Q9DCT8 | Crip2 | -4.23756695 | 2.476903583 |
| Q00993 | Axl | -4.238688312 | 2.20941017 |
| P52760 | Rida | -4.239432209 | 3.191724095 |
| P28798 | Grn | -4.252622288 | 2.30470553 |
| Q61036 | Pak3 | -4.263478511 | 3.173768604 |
| Q80UW2 | Fbxo2 | -4.267560505 | 2.318710286 |
| Q00493 | Cpe | -4.271472962 | 3.867651428 |
| Q3UZA1 | Rcsd1 | -4.275797026 | 2.841565638 |
| P15864 | H1-2 | -4.281983379 | 2.498775343 |
| Q8CIF4 | Btd | -4.284355655 | 2.159482896 |
| P52624 | Upp1 | -4.28528962 | 3.26521754 |
| Q9CX60 | Lbh | -4.287416902 | 3.354970619 |
| Q924C6 | Loxl4 | -4.291000443 | 2.698972346 |
| Q8R344 | Ccdc12 | -4.292695211 | 2.671307763 |
| Q8BYU6 | Tor1aip2 | -4.294523333 | 2.994769192 |
| Q9CPV5 | Pmf1 | -4.299528907 | 2.015833317 |
| Q8CFI5 | Pars2 | -4.301172885 | 4.854481453 |
| Q9D937 |  | -4.312637835 | 2.984438046 |
| P97315 | Csrp1 | -4.313111739 | 3.85427115 |
| Q9D1C3 | Pyurf | -4.320004189 | 3.588060931 |
| Q9CY28 | Gtpbp8 | -4.328636361 | 2.216758931 |
| Q8R086 | Suox | -4.339193266 | 2.46244566 |
| Q8R0F5 | Rbmx2 | -4.341394361 | 2.812385134 |
| Q99MS7 | Ehbp1l1 | -4.346104255 | 2.926004105 |
| O89023 | Tpp1 | -4.348187632 | 4.460956923 |
| Q7TPV4 | Mybbp1a | -4.354413061 | 2.139336494 |
| Q9WVL0 | Gstz1 | -4.356006655 | 3.052898318 |
| Q78WZ7 | Polr1f | -4.358941634 | 2.341530109 |
| B1AY10 | Nfx1 | -4.362501248 | 5.079252495 |
| Q61733 | Mrps31 | -4.371500912 | 3.441144696 |
| Q8BK35 | Nop53 | -4.379602349 | 2.406008981 |
| Q99KR3 | Lactb2 | -4.38159526 | 4.023016918 |
| Q9CPW3 | Mrpl54 | -4.415431173 | 3.334790356 |
| P17047 | Lamp2 | -4.421367509 | 3.29000737 |
| P15806 | Tcf3 | -4.437139496 | 2.188684889 |
| Q3UKC1 | Tax1bp1 | -4.45187219 | 2.873052325 |
| O08908 | Pik3r2 | -4.45424969 | 2.043536641 |
| Q9Z0F6 | Rad9a | -4.456621402 | 2.737802089 |
| Q8R4N0 | Clybl | -4.459006342 | 3.518814068 |
| Q76KJ5 | Polr1g | -4.459365263 | 2.274539999 |
| Q8K3D3 | Swi5 | -4.4665174 | 4.420212838 |
| Q9ESP1 | Sdf2l1 | -4.472119556 | 3.426070453 |
| O09117 | Sypl1 | -4.472222096 | 3.272294687 |
| Q6PE15 | Abhd10 | -4.474514993 | 2.84089245 |
| Q5SV80 | Myo19 | -4.48215565 | 2.51092124 |
| P62965 | Crabp1 | -4.482298203 | 4.186900043 |
| Q9CQ18 | Rnaseh2c | -4.490492205 | 4.937696224 |
| Q8CES0 | Naa30 | -4.491497499 | 2.08138427 |
| O55137;Q8BWN8;Q9QYR9;Q32Q92 | Acot | -4.493983988 | 2.910796499 |
| Q9CXC3 | Mgme1 | -4.503068545 | 3.562771072 |
| Q80Y50 | Camta2 | -4.506896126 | 2.538733416 |
| Q8CIV8 | Tbce | -4.507175283 | 2.818465152 |
| A2ALU4 | Shroom2 | -4.508144931 | 3.113941295 |
| Q9CYC5 | Dsn1 | -4.514874381 | 3.123086821 |
| Q91W92 | Cdc42ep1 | -4.523030971 | 3.988497832 |
| Q61508 | Ecm1 | -4.525905691 | 3.851287414 |
| O09159 | Man2b1 | -4.52986181 | 3.202141513 |
| Q8K4F5 | Abhd11 | -4.53459971 | 2.288518052 |
| P70255;P70257 | Nfix | -4.546675231 | 4.404571635 |
| Q3TUU5 | Tex30 | -4.54859831 | 3.074252394 |
| P20065 | Tmsb4x | -4.556659853 | 2.716769035 |
| A2ADA5 | Pusl1 | -4.558836785 | 2.987156379 |
| Q9WUJ8 | Orc6 | -4.565738142 | 2.467474654 |
| O35405 | Pld3 | -4.573297846 | 2.484339782 |
| Q9CQA6 | Chchd1 | -4.581547709 | 2.3641628 |
| Q8K2M0 | Mrpl38 | -4.591538331 | 4.916674528 |
| Q8C650 | Septin10 | -4.605908415 | 2.651441305 |
| Q791N7 | Polr1h | -4.611807604 | 3.153485825 |
| Q80ZQ9 | Abitram | -4.616631568 | 2.765983087 |
| Q9D727 | Pex39 | -4.619905051 | 2.473217337 |
| Q9CQL6 | Mrpl35 | -4.629526491 | 3.096742541 |
| Q8VBT0 | Tmx1 | -4.630783889 | 2.319998327 |
| Q3UFK8 | Frmd8 | -4.639889058 | 2.105900219 |
| Q14B71 | Cdca2 | -4.640687476 | 2.529654868 |
| Q60778 | Nfkbib | -4.641798768 | 2.178504997 |
| Q3ULW8 | Parp3 | -4.644070446 | 3.786435915 |
| Q8CIC2 | Nup42 | -4.660315279 | 2.631576003 |
| Q99KS2 | Ngrn | -4.666270978 | 2.387576484 |
| Q9DCI9 | Mrpl32 | -4.670096267 | 4.240819544 |
| Q9JLR9 | Higd1a | -4.671772646 | 5.060390014 |
| P12032 | Timp1 | -4.671908098 | 2.862815487 |
| Q8CJ27 | Aspm | -4.676747022 | 3.658496941 |
| Q05915 | Gch1 | -4.68937357 | 3.9127205 |
| Q8R570 | Snap47 | -4.716482623 | 2.3475059 |
| Q14AI0 | DSCC1 | -4.718982187 | 2.597925029 |
| Q60I26 | Als2cl | -4.719602261 | 3.04540736 |
| Q9CQP3 | Chchd5 | -4.722820665 | 3.405424105 |
| Q60767 | Ly75 | -4.730663651 | 4.927022073 |
| Q9CYZ6 | Rex1bd | -4.731621864 | 2.161612467 |
| O35639 | Anxa3 | -4.731686708 | 4.801316868 |
| Q9D6U8 | Fam162a | -4.733081445 | 3.121138114 |
| Q8VDT9 | Mrpl50 | -4.734191511 | 2.204338917 |
| O35954 | Pitpnm1 | -4.744889779 | 2.38070201 |
| Q8R035 | Mrpl58 | -4.745723991 | 3.056109998 |
| Q9CPT4 | Mydgf | -4.753535476 | 3.094795869 |
| Q64339 | Isg15 | -4.769570424 | 2.435587213 |
| Q499E6 | Airim | -4.785430585 | 2.062844428 |
| Q6PB51 | Ccdc117 | -4.789905412 | 2.069708328 |
| Q9CWG8 | Ndufaf7 | -4.801210398 | 3.073620454 |
| Q60953 | Pml | -4.80796581 | 3.954965996 |
| Q60739 | Bag1 | -4.815692384 | 2.635691481 |
| P55302 | Lrpap1 | -4.827622456 | 3.093163664 |
| P40630 | Tfam | -4.828079004 | 2.770376587 |
| Q9D2R8 | Mrps33 | -4.838676642 | 2.991240844 |
| Q61263 | Soat1 | -4.838811173 | 2.197253812 |
| P10107 | Anxa1 | -4.850332585 | 4.373730627 |
| P35288 | Rab23 | -4.875413275 | 2.816241062 |
| P99025 | Gchfr | -4.876747688 | 3.346240978 |
| P43277 | H1-3 | -4.876752526 | 2.386857078 |
| Q99MZ7 | Pecr | -4.896710735 | 4.402250522 |
| Q9QZF2 | Gpc1 | -4.898046688 | 2.139557192 |
| Q3UIU2 | Ndufb6 | -4.910830682 | 2.152252642 |
| P35456 | Plaur | -4.911431321 | 2.738406244 |
| Q61398 | Pcolce | -4.916624237 | 4.375749023 |
| Q8VI94 | Oasl1 | -4.920800988 | 2.676764785 |
| Q8K2T1 | Nmral1 | -4.935392762 | 2.317680506 |
| Q3TYS2 | Cybc1 | -4.937879012 | 3.032916515 |
| O88668 | Creg1 | -4.940443131 | 2.384756347 |
| Q9CQZ5 | Ndufa6 | -4.944808172 | 3.383651636 |
| Q80WT5 | Aftph | -4.982711569 | 3.212680824 |
| Q9DB29 | Iah1 | -4.98771007 | 2.771084148 |
| Q8BFS6 | Cpped1 | -4.994173509 | 2.280740671 |
| Q9ET22 | Dpp7 | -4.995733681 | 4.124831985 |
| Q08775;Q64131 | Runx3 | -4.999652643 | 2.304676625 |
| Q9D8B4 | Ndufa11 | -5.002911263 | 2.955857843 |
| Q8R164 | Bphl | -5.006510287 | 2.913817753 |
| Q9CQ54 | Ndufc2 | -5.010818896 | 2.636996655 |
| Q8VEH5 | Epm2aip1 | -5.043816318 | 2.155839428 |
| Q60872 | Eif1a | -5.056102838 | 3.490600993 |
| Q9CR25 | Dph2 | -5.065537061 | 2.771116166 |
| Q3TUH1 | Tamm41 | -5.066079782 | 3.781559558 |
| O88939 | Zbtb7a | -5.070491898 | 3.608367302 |
| Q99LY9 | Ndufs5 | -5.09600022 | 3.313028026 |
| Q6P3Y5 | Znf280c | -5.109070494 | 3.852779563 |
| Q91VN4 | Chchd6 | -5.119804108 | 2.702870322 |
| Q91YR9 | Ptgr1 | -5.126163421 | 3.327753528 |
| Q8BIW1 | Prune1 | -5.147527042 | 3.732444265 |
| Q8VBT6 | Apobr | -5.152335218 | 2.30034663 |
| Q8BN58 | Arhgap28 | -5.15880753 | 2.858562347 |
| Q8BGX2 | Timm29 | -5.160882096 | 2.709388354 |
| O70503 | Hsd17b12 | -5.162131413 | 2.690518564 |
| Q9Z0L0 | Tpbg | -5.180330261 | 2.248160356 |
| P48725 | Pcnt | -5.180646148 | 3.418453048 |
| Q91YU8 | Ppan | -5.197976827 | 3.284104295 |
| P16045 | Lgals1 | -5.208209972 | 6.042708413 |
| Q8BGB5 | Limd2 | -5.209741101 | 2.906564399 |
| P0DJF2 | Pet117 | -5.238710215 | 2.448293297 |
| Q8K039 |  | -5.245079808 | 2.11838146 |
| Q9QZN4 | Fbxo6 | -5.251015737 | 2.975971524 |
| Q9CQ06 | Mrpl24 | -5.258842441 | 3.597547394 |
| P97352 | S100a13 | -5.2815032 | 3.409255231 |
| P53690 | Mmp14 | -5.304377558 | 2.718785915 |
| Q91V76 |  | -5.319879232 | 2.46244566 |
| Q8C2E4 | Ptcd1 | -5.32067519 | 4.713431547 |
| P10404 |  | -5.325997984 | 2.906999965 |
| P56135 | Atp5mf | -5.336626528 | 2.199116428 |
| Q9D2Y4 | Mlkl | -5.344361353 | 3.276959002 |
| Q9CR89 | Ergic2 | -5.366434024 | 2.186859784 |
| Q9JMB0 | Gkap1 | -5.386435888 | 2.088530858 |
| Q9WV54 | Asah1 | -5.443490543 | 2.279162457 |
| O35658 | C1qbp | -5.445521208 | 4.081971138 |
| Q63918 | Cavin2 | -5.455285882 | 3.408535097 |
| O89110 | Casp8 | -5.480569305 | 2.793751377 |
| Q9QZM2 | Polg2 | -5.494525929 | 4.309548293 |
| Q78HU3 | Mvb12a | -5.523531761 | 2.154581882 |
| Q64105 | Spr | -5.527456525 | 3.566475112 |
| Q3UY34 | Custos | -5.539264864 | 3.2555745 |
| O35640 | Anxa8 | -5.54321613 | 2.84089245 |
| P10923 | Spp1 | -5.546591872 | 2.932332797 |
| Q8VC52 | Rbpms2 | -5.559796898 | 2.288396779 |
| Q9D2Q3 | Paat | -5.565345854 | 3.031388376 |
| Q3TZX8 | Nol9 | -5.577768636 | 3.190864341 |
| Q9D3U0 | Pus10 | -5.57977563 | 2.74199825 |
| Q9QZM4 | Tnfrsf10b | -5.591588945 | 2.39282243 |
| Q61337 | Bad | -5.620367617 | 2.69129318 |
| Q9CWY8 | Rnaseh2a | -5.627739893 | 2.936730216 |
| Q9D945 | Llph | -5.64129939 | 4.368235472 |
| P00405 | Mtco2 | -5.644577966 | 4.024161182 |
| Q9JL35 | Hmgn5 | -5.64574531 | 2.970223976 |
| Q8BQ30 | Ppp1r18 | -5.653230372 | 4.412462579 |
| Q9CQJ8 | Ndufb9 | -5.661117761 | 2.166106964 |
| B1AR13 | Cisd3 | -5.665994768 | 3.258654337 |
| P07091 | S100a4 | -5.696793701 | 2.695218279 |
| P19157;P46425 | Gstp | -5.713855763 | 6.136656712 |
| Q8BP78 | Fra10ac1 | -5.731318096 | 3.404627076 |
| Q99J99 | Mpst | -5.781770614 | 3.350341861 |
| Q62426 | Cstb | -5.782147283 | 5.396543964 |
| P57722 | Pcbp3 | -5.782546836 | 3.081485254 |
| Q9DC71 | Mrps15 | -5.795321128 | 2.202707763 |
| Q9D1J1 | Necap2 | -5.816738844 | 3.446856584 |
| P43276 | H1-5 | -5.830876471 | 2.029113263 |
| O35215 | Ddt | -5.839937791 | 5.396543964 |
| Q8BVY0 | Rsl1d1 | -5.840649062 | 2.831778923 |
| P17182;P17183 | Eno | -5.872594434 | 4.207562647 |
| P35762 | Cd81 | -5.910951881 | 2.499561755 |
| Q9CRB2 | Nhp2 | -5.923203892 | 2.272134535 |
| Q9CQ75 | Ndufa2 | -5.936304874 | 3.258127611 |
| P16858;Q64467 | Gapdhs | -5.939201708 | 6.136656712 |
| Q8C407 | Yipf4 | -5.949188483 | 4.341673123 |
| Q9D1R2 | Kti12 | -5.988587448 | 2.695494803 |
| P97825 | Jpt1 | -5.999497462 | 4.676729037 |
| Q61033 | Tmpo | -6.005567956 | 4.044096318 |
| Q9CQC7 | Ndufb4 | -6.034349767 | 5.079252495 |
| P49962 | Srp9 | -6.035358363 | 3.308919843 |
| P19783 | Cox4i1 | -6.061444461 | 3.851287414 |
| P60330 | Espl1 | -6.098973106 | 2.86697281 |
| Q3TLP5 | Echdc2 | -6.10711809 | 3.22009771 |
| P06797 | Ctsl | -6.113456983 | 4.182525814 |
| P46656 | Fdx1 | -6.170453634 | 3.078839402 |
| Q9DBG5 | Plin3 | -6.201896787 | 4.194544378 |
| P28653 | Bgn | -6.327947001 | 2.50883411 |
| Q9CQM5 | Txndc17 | -6.363985127 | 4.054244187 |
| Q80ZS3 | Mrps26 | -6.364535061 | 4.237499095 |
| Q8VCW8 | Acsf2 | -6.388998511 | 2.433910954 |
| O08997 | Atox1 | -6.397687899 | 2.778551694 |
| Q9D1L0 | Chchd2 | -6.419328316 | 3.143195443 |
| P48771 | Cox7a2 | -6.427277805 | 4.005229598 |
| Q80VJ3 | Dnph1 | -6.447913403 | 3.522157992 |
| Q99K30 | Eps8l2 | -6.490031903 | 3.159174208 |
| P40240 | Cd9 | -6.632543891 | 3.954965996 |
| P16110 | Lgals3 | -6.654452836 | 5.362287042 |
| Q8BHL4 | Gprc5a | -6.658614484 | 4.099484955 |
| P16675 | Ctsa | -6.666831619 | 2.627640202 |
| Q9Z127 | Slc7a5 | -6.680858038 | 3.124487367 |
| P31786 | Dbi | -6.685944132 | 4.844819557 |
| P99028 | Uqcrh | -6.862311566 | 6.316420544 |
| Q9R1Q7 | Plp2 | -6.864647929 | 4.520849775 |
| P48755 | Fosl1 | -6.911887118 | 4.412462579 |
| Q80U04 | Pja2 | -6.927656428 | 4.748067278 |
| Q9QZL0 | Ripk3 | -6.973940645 | 2.525968266 |
| P70202 | Lxn | -7.006501045 | 3.142336775 |
| P97450 | Atp5pf | -7.043122753 | 4.417019114 |
| P51174 | Acadl | -7.052965948 | 5.052194719 |
| Q8BJY1 | Psmd5 | -7.180482664 | 4.103666555 |
| P51125 | Cast | -7.181454586 | 2.548751065 |
| P28474;Q64437 | Adh | -7.194601224 | 5.11684144 |
| P15379 | Cd44 | -7.258179699 | 3.753531572 |
| Q9DCS3 | Mecr | -7.275111754 | 2.926513144 |
| Q9CPQ8 | Atp5mg | -7.305826919 | 4.368140153 |
| Q8R1I1 | Uqcr10 | -7.722826803 | 3.769260716 |
| P18572 | Bsg | -7.764839231 | 3.706483995 |
| Q05769 | Ptgs2 | -7.943284006 | 2.763584951 |
| P24452 | Capg | -8.155672704 | 3.793850156 |
| P97370 | Atp1b3 | -8.180058403 | 2.84263146 |
| Q9CPQ1 | Cox6c | -10.51545614 | 3.166785549 |

**Table S4.** *Protein signature of USA 2003 MOI 1 infected KO vs WT MEF lysates at 21 hours post-infection. Table lists the protein ID, gene name,, the log2-fold change (FC) (MPXV infected WT MEFs vs infected ISG15 KO MEFs) of the levels of each protein, and the statistical significance (−log P value). Ordered from largest to smallest FC.*
