## Supplementary material for "ISG15 Differentially Modulates Clade Ib and II MPXV Infection in MEF cells": TableS5

| **Name** | **Gene** | **fold_change** | **p_value_adj_neg_log10** |
| --- | --- | --- | --- |
| Q8VIN1 | Pbp2 | 7.944863745 | 4.163018778 |
| Q9Z0N2 | Eif2s3y | 7.460030331 | 4.523737453 |
| Q03526 | Itk | 7.434598628 | 3.84216487 |
| P00920 | Ca2 | 7.11287456 | 4.970406784 |
| Q80TL0 | Ppm1e | 6.60906591 | 3.822496452 |
| P30275;Q6P8J7 | Ckmt | 6.569622554 | 3.616465406 |
| Q9DAJ5 | Dynlrb2 | 6.515414274 | 3.311602057 |
| P31254 | Uba1y | 6.05249476 | 5.902972035 |
| P28651 | Ca8 | 5.971864756 | 4.970406784 |
| P58321 | Uchl4 | 5.877588921 | 4.370300403 |
| Q91WT9 | Cbs | 5.828807547 | 4.953649444 |
| Q8VHG2;Q9D4H4 | Amot | 5.565602268 | 3.941049284 |
| Q9ET01;Q9WUB3 | Pyg | 5.484319366 | 3.237311168 |
| Q80ZN9 | Cox6b2 | 5.270359846 | 3.714132432 |
| P08553 | Nefm | 5.247954671 | 6.236924249 |
| B1AVY7 | Kif16b | 5.201782103 | 3.454451089 |
| Q8JZV9 | Bdh2 | 4.985109482 | 3.734916066 |
| P19639 | Gstm3 | 4.904672482 | 2.929979595 |
| Q61166;Q8R001 | Mapre | 4.767659832 | 3.429574276 |
| Q9EQF6 | Dpysl5 | 4.74139701 | 3.66847644 |
| P16125 | Ldhb | 4.733625891 | 4.433360725 |
| Q6PIC6;Q6PIE5 | Atp1a | 4.719501442 | 2.583008559 |
| Q60611;Q8VI24 | Satb | 4.663015927 | 3.819171414 |
| Q9D1H6 | Ndufaf4 | 4.559716836 | 4.175334966 |
| Q8K157 | Galm | 4.526497951 | 5.297187852 |
| Q60936 | Coq8a | 4.390218978 | 3.349429573 |
| Q8K0U4 | Hspa12a | 4.358033065 | 2.739175354 |
| O55111 | Dsg2 | 4.308754337 | 4.433360725 |
| O55091 | Impact | 4.287101815 | 4.554485663 |
| Q99JT2 | Stk26 | 4.247088542 | 3.651629014 |
| Q91ZE0 | Tmlhe | 4.225031514 | 3.199062862 |
| O55042;Q91ZZ3 | Snc | 4.209158776 | 4.546417755 |
| Q6PDS3 | Sarm1 | 4.207668161 | 4.23902178 |
| P08551 | Nefl | 4.19707777 | 4.48142587 |
| P84075;Q91X97 | Hpca;Ncald | 4.171061535 | 3.626170426 |
| Q45KJ6 | Lin28b | 4.141020562 | 4.682174872 |
| Q99K67 | Aass | 4.110506382 | 3.146070578 |
| Q8R1G2 | Cmbl | 4.044546182 | 4.060997583 |
| Q6WVG3 | Kctd12 | 4.017216491 | 4.701803578 |
| Q61474 | Msi1 | 4.01684546 | 4.489637732 |
| Q9QZW9 | Mnx1 | 4.010003794 | 2.577204759 |
| Q6Q2Z6 | Acot5 | 3.98835916 | 4.491544855 |
| Q58A65;Q9ESN9 | Spag9;Mapk8ip3 | 3.987435483 | 3.862771613 |
| P22227;Q00899;Q3TTC2 | Zfp42;Yy1;Yy2 | 3.984605199 | 3.255352526 |
| Q6NVG5 | **Mreg** | 3.971584759 | 3.107744365 |
| A2AUC9 | Klhl41 | 3.966913919 | 3.130627884 |
| Q63ZW7 | Patj | 3.866365607 | 2.939120519 |
| Q9WU63 | Hebp2 | 3.842252241 | 2.108608447 |
| Q9CZC8 | Scrn1 | 3.803833663 | 2.121971772 |
| P16381 | D1Pas1 | 3.802409923 | 3.659032049 |
| Q9CQE5 | Rgs10 | 3.754838163 | 4.267487385 |
| O70209 | Pdlim3 | 3.75161994 | 4.716211672 |
| Q8BRK8 | Prkaa2 | 3.735244171 | 3.516752857 |
| P62141;P63087 | Ppp1c | 3.727931355 | 4.043650602 |
| Q9DA03 | Lyrm7 | 3.71971141 | 4.291116846 |
| Q9WV02 | Rbmx | 3.713883647 | 2.477790167 |
| Q9Z2V4 | Pck1 | 3.691878794 | 3.170403812 |
| Q04447;P07310 | Ckb;Ckm | 3.672969272 | 4.756852261 |
| Q62148 | Aldh1a2 | 3.661465778 | 4.163018778 |
| P15864;P43274;P43275;P43277 | H1 | 3.625414023 | 3.83578365 |
| Q3UQ44 | Iqgap2 | 3.589000747 | 4.489637732 |
| Q8C779 | Radx | 3.539428536 | 2.933607775 |
| P05213;P05214;P68368;P68369;P68373;Q9JJZ2;Q3UX10 | Tuba | 3.535629385 | 3.59715736 |
| Q9JME5 | Ap3b2 | 3.524015812 | 3.561899877 |
| P70207 | Plxna2 | 3.501802459 | 3.804907615 |
| O55042 | Sh3gl2 | 3.486811467 | 3.657043317 |
| Q62420 | Hdac6 | 3.469274776 | 2.420917946 |
| Q9Z2V5 | Ap3b2 | 3.450573372 | 2.836770973 |
| P16627;P17879;Q61696 | Hspa | 3.443795506 | 4.364010479 |
| Q9D114 | Hddc3 | 3.411232574 | 2.225967087 |
| B1AXP6 | Tomm5 | 3.393723285 | 4.163018778 |
| Q641K1 | Agtpbp1 | 3.377050945 | 2.512393059 |
| Q8K4I3 | Arhgef6 | 3.363319814 | 2.798876143 |
| P55271 | Cdkn2b | 3.355806474 | 5.569291115 |
| Q9D5J6 | Shpk | 3.335489496 | 2.67283144 |
| Q9CPU0 | Glo1 | 3.313875815 | 5.176252433 |
| Q9Z1N7 | Arid3b | 3.313275886 | 4.546805601 |
| Q64524;Q8CGP0;Q9D2U9 | H2bc2 | 3.306914096 | 4.125187546 |
| O35188 | Cx3cl1 | 3.274074652 | 4.375414722 |
| Q62431;Q9Z1N7 | Arid3a | 3.268620812 | 3.413278872 |
| Q9JHI5 | Ivd | 3.223961727 | 2.274475429 |
| S4R1M9 | Osbpl10 | 3.219743832 | 2.102290164 |
| Q91WJ7 | Spats2l | 3.218014541 | 2.810302394 |
| Q8BIL5 | Hook1 | 3.181706081 | 3.116033233 |
| Q80UW0 | Hs6st2 | 3.163814255 | 3.500033842 |
| P09066 | En2 | 3.148181148 | 3.93928595 |
| O08638;Q61879;Q6URW6 | Myh | 3.144390605 | 2.095267826 |
| Q8R0P4 | Aamdc | 3.141790986 | 5.034065923 |
| Q5XG69 | Fam169a | 3.105927682 | 3.682308114 |
| Q5BKP2 | Usp13 | 3.081224769 | 3.077815263 |
| Q9Z2Z9 | Gfpt2 | 3.077213636 | 2.323942502 |
| P97931 | Ung | 3.075088388 | 4.032981772 |
| Q9CYT6 | Cap2 | 3.064245565 | 4.174892384 |
| P23198;P83917 | Cbx | 3.057883826 | 2.630153951 |
| Q8C196 | Cps1 | 3.049332376 | 2.732194975 |
| P46660 | Ina | 3.037545751 | 2.351676847 |
| O70318;Q9WV92;Q9Z2H5 | Epb41l | 2.978404485 | 2.076542446 |
| P23475 | Xrcc6 | 2.969700891 | 5.297187852 |
| P97447 | Fhl1 | 2.942515669 | 4.664616004 |
| Q9D711 | Pir | 2.930708811 | 2.110792021 |
| Q8VCK3 | Tubg2 | 2.928126181 | 5.569291115 |
| Q8CHY3 | Dym | 2.911126963 | 3.432078766 |
| G5E8K5 | Ank3 | 2.873865289 | 3.199798048 |
| Q9CQA9 | Ntpcr | 2.865787933 | 5.983044507 |
| P68372;P99024;Q7TMM9;Q9CWF2;Q9D6F9;Q9ERD7 | Tubb | 2.863989323 | 4.982715035 |
| A0A7H0DN78 | OPG106 | 2.845922022 | 3.718578439 |
| Q5SSL4;Q6PAJ1 | Abr;Bcr | 2.811116035 | 3.748695666 |
| P24549 | Aldh1a1 | 2.754261819 | 4.002145646 |
| Q9ERD6 | Ralgps2 | 2.751452457 | 2.680315737 |
| P0C027;P0C028 | Nudt1 | 2.750343233 | 2.29522114 |
| A3KMP2 | Ttc38 | 2.741349823 | 4.324404823 |
| Q9CZS1 | Aldh1b1 | 2.714275168 | 5.259521696 |
| Q60899 | Elavl2 | 2.700133066 | 4.125187546 |
| P15105 | Glul | 2.699672646 | 4.740254439 |
| Q8JZS0 | Lin7a | 2.657838037 | 3.742789998 |
| Q91W43 | Gldc | 2.647156636 | 3.849663192 |
| Q99MU3 | Adar | 2.64348719 | 6.245955496 |
| Q64442 | Sord | 2.639527592 | 3.103680465 |
| O08791;Q8K4J2 | Ebf | 2.63575435 | 2.548383399 |
| Q9D8X1 | Cutc | 2.625001352 | 4.747116276 |
| Q8K0E1 | Kctd15 | 2.607654463 | 3.458827404 |
| Q8VC88 | Gca | 2.599173398 | 2.485863816 |
| Q8VI93 | Oas3 | 2.599113259 | 3.483467608 |
| Q7TPD6 | Raver2 | 2.565157897 | 2.745999975 |
| Q8BTG3 | Tcp11l1 | 2.512899374 | 2.910066901 |
| Q02956 | Prkcz | 2.510031553 | 3.578138165 |
| Q6P3D0;Q8VHN8 | Nudt | 2.505134579 | 5.297187852 |
| P11103 | Parp1 | 2.498232114 | 4.443807798 |
| Q9WVL1 | Ap4s1 | 2.495120928 | 4.033884815 |
| Q62441 | Tle4 | 2.492243748 | 2.564507046 |
| A2ATU0 | Dhtkd1 | 2.491652266 | 4.059745738 |
| O35984 | Pbx2 | 2.475780446 | 2.460794555 |
| M1LBQ5 | OPG115 | 2.4736593 | 3.82489943 |
| Q5RL51 | Gstcd | 2.451079022 | 2.674292378 |
| A0A7H0DN76 | OPG104 | 2.40491287 | 3.202162361 |
| M1KJ15 | OPG114 | 2.404170847 | 4.56336512 |
| P83741;Q3UH66;Q80UE6 | Wnk | 2.382765639 | 3.822122385 |
| A0A7H0DNC1 | OPG149 | 2.381804764 | 3.764782768 |
| Q9R0Q6 | Arpc1a | 2.35303739 | 5.661601374 |
| Q3UUI3 | Them4 | 2.334025549 | 3.581609683 |
| A0A7H0DND9;Q8V4T7 | OPG171 | 2.321727275 | 3.110889723 |
| P49813 | Tmod1 | 2.314078477 | 3.503021201 |
| Q8BPK2 | Zcchc3 | 2.244682182 | 4.489637732 |
| P00342;P06151;P16125 | Ldh | 2.235165325 | 4.953649444 |
| Q9QY15;Q61655 | Ddx | 2.208029924 | 3.734328814 |
| Q9Z191 | Eya4 | 2.12486014 | 3.406179482 |
| O70583 | Mid1 | 2.10087024 | 2.790629326 |
| Q9CPN8 | Igf2bp3 | 2.085868838 | 4.491544855 |
| Q9CR64 | Tmem167a | 2.05946668 | 3.613871478 |
| M1L535 | OPG128 | 2.056843044 | 3.295317908 |
| Q9D7X8 | Ggct | 2.041719723 | 4.127449222 |
| B1ARW8 |  | 2.021869307 | 2.495871832 |
| Q9JL70 | Fanca | 2.020198738 | 3.874884085 |
| A0A7H0DNC4 | OPG153 | 2.017769172 | 3.217306994 |
| P97313 | Prkdc | 2.012860673 | 2.434244463 |
| P07742;A0A7H0DN52 | Rmn1;OPG080 | 1.994647352 | 3.133929925 |
| Q9Z129 | Recql | 1.988743148 | 5.280410943 |
| A2AUM9;Q62036 | Cep1 | 1.975192256 | 3.560731626 |
| Q7TPM6 | Fsd1 | 1.973895401 | 2.056545023 |
| P61079;P61080;P62838 | Ube2d | 1.970466262 | 5.342388324 |
| Q80UM3;Q9DBB4 | Naa15;Naa16 | 1.958532802 | 4.774598006 |
| P16460 | Ass1 | 1.947250943 | 2.507354832 |
| Q8C7Q4 | Rbm4 | 1.946555211 | 3.996449161 |
| Q3V0K9;Q61233;Q99K51 | Pls1 | 1.936368958 | 2.744669344 |
| O88477;Q9CPN8 | Igf2bp1 | 1.935249428 | 4.158338398 |
| A0A7H0DN45 | OPG072 | 1.923418606 | 2.821233618 |
| A0A7H0DN83 | OPG111 | 1.917453565 | 3.165908239 |
| A0A7H0DN80 | OPG108 | 1.909813442 | 3.043006879 |
| Q8BJZ4 | Mrps35 | 1.907328028 | 3.58503414 |
| P62137;P63087 | Ppp1ca | 1.90249124 | 4.098866019 |
| Q3TIR3;Q80XE1 | Ric8 | 1.895228015 | 3.705070314 |
| Q5SU73 | Coil | 1.892881525 | 4.142594819 |
| A0A7H0DN72 | OPG100 | 1.888682718 | 3.227637961 |
| P30658 | Cbx2 | 1.865261416 | 2.044824543 |
| Q8K0Z7 | Taco1 | 1.861758186 | 3.460817988 |
| Q8BYJ6 | Tbc1d4 | 1.859947248 | 3.697761762 |
| A0A7H0DN46 | OPG073 | 1.855945406 | 3.673337794 |
| Q80V62 | Fancd2 | 1.853976028 | 4.467485047 |
| Q9CQF6 | Aasdhppt | 1.846743192 | 4.400805756 |
| Q9QXK3;Q9QZE5 | Copg | 1.843403777 | 4.622443967 |
| Q9CWI3 | Bccip | 1.834121854 | 3.460817988 |
| Q9ERE3 | Sgk3 | 1.82857645 | 4.23149235 |
| P83510 | Tnik | 1.827519919 | 2.705550265 |
| A0A7H0DN79 | OPG107 | 1.814897803 | 2.05439843 |
| M1L543 | OPG138 | 1.805484252 | 3.031138203 |
| A0A7H0DN35 | OPG062 | 1.804262722 | 3.138921808 |
| Q8JZM0 | Tfb1m | 1.803675328 | 3.325110565 |
| Q9DB30 | Phkg2 | 1.799280784 | 3.107159946 |
| Q61699;P48722;Q61316 | Hsp | 1.79139664 | 4.104100375 |
| Q9WUR9 | Ak4 | 1.782783883 | 3.542778652 |
| A0A7H0DN69 | OPG097 | 1.78177522 | 2.741473552 |
| Q9JKN6 | Nova1 | 1.779382167 | 2.786073512 |
| P48758 | Cbr1 | 1.777247977 | 2.337142805 |
| Q9Z2H7 | Gipc2 | 1.775083028 | 2.966291815 |
| P60521 | Gabarapl2 | 1.772121786 | 4.56955956 |
| A0A7H0DN57 | OPG085 | 1.770789254 | 3.094208894 |
| P60824 | Cirbp | 1.764581757 | 4.779878283 |
| P62880;P29387 | Gnb | 1.760934906 | 2.796107751 |
| A0A7H0DN43 | OPG070 | 1.737340179 | 3.242251331 |
| P0DTN1 | OPG139 | 1.731238115 | 2.653733218 |
| A6H630 | Armt1 | 1.729859016 | 4.356639324 |
| A0A7H0DN48 | OPG075 | 1.725547051 | 3.150041842 |
| O70456;P61982;P62259;P63101;P68254;P68510;Q9CQV8 | Sfn;Ywh | 1.711314055 | 4.209368071 |
| Q99J10 | Ctu1 | 1.704280765 | 4.803217743 |
| A0A7H0DN74 | OPG102 | 1.696314558 | 3.856980877 |
| Q9D9H8 |  | 1.694434325 | 2.312491176 |
| O08575 | Eya2 | 1.688359488 | 2.478085382 |
| Q03173 | Enah | 1.687641067 | 4.848390091 |
| Q8BGS2 | Bola2 | 1.681898139 | 4.581748837 |
| A0A7H0DN41 | OPG068 | 1.676880903 | 3.156361021 |
| P48962;P51881;Q3V132 | Slc25a | 1.666637319 | 2.42647913 |
| A0A7H0DN89 | OPG117 | 1.664566966 | 3.823581864 |
| Q64512 | Ptpn13 | 1.657389292 | 3.551836988 |
| A0A7H0DNB5 | OPG143 | 1.650949503 | 2.287992226 |
| Q9D032 | Ssbp3 | 1.616859643 | 4.220191571 |
| Q8BXK9;Q9QYB1;Q8BHB9 | Clic | 1.612595484 | 2.008657815 |
| A0A7H0DN42 | OPG069 | 1.594182548 | 4.301764195 |
| A0A7H0DNA4 | OPG132 | 1.586542878 | 3.760306828 |
| Q9QY93 | Dctpp1 | 1.583629399 | 3.622078421 |
| A0A7H0DN27 | OPG054 | 1.580968114 | 3.268331914 |
| Q8BGS1;Q9JMC8 | Epb41l | 1.577392703 | 2.981551498 |
| P97855;P97379 | G3bp | 1.576841406 | 3.73630179 |
| Q61189 | Clns1a | 1.569061428 | 5.013335147 |
| A0A7H0DNE6 | OPG180 | 1.56230487 | 3.66649026 |
| P15626 | Gstm2 | 1.556048015 | 3.254485846 |
| Q3TIW9;Q8BTJ4 | Enpp | 1.548673535 | 3.567021592 |
| A0A7H0DNA5 | OPG133 | 1.548244803 | 3.560940714 |
| P70340;P97454;Q62432;Q8BUN5;Q9JIW5 | Smad | 1.539351905 | 3.684682755 |
| O89106 | Fhit | 1.534070227 | 2.305739066 |
| Q8VIM9 | Irgq | 1.532350953 | 3.012231411 |
| Q9R0P9 | Uchl1 | 1.524131088 | 3.577626323 |
| P52482;P52483;Q91W82 | Ube2e | 1.51097343 | 2.51947099 |
| P62855 | Rps26 | 1.510881703 | 3.27031048 |
| P61922 | Abat | 1.509661959 | 2.159951152 |
| Q925B0 | Pawr | 1.506835593 | 3.638918751 |
| Q8VD33 | Sgtb | 1.499057159 | 2.856855788 |
| M1L9M3 | OPG087 | 1.498592702 | 3.682104489 |
| Q9CYX7 | Rrp15 | 1.462289636 | 3.044006115 |
| Q8BHN1 | Txlng | 1.459473516 | 4.701474809 |
| A0A7H0DN81 | OPG109 | 1.451631212 | 2.991877156 |
| Q9Z130 | Hnrnpdl | 1.450677016 | 4.472014909 |
| Q6ZWN5 | Rps9 | 1.440256062 | 4.338535926 |
| Q8K4F6 | Nsun5 | 1.436423331 | 3.920654828 |
| M1L502 | OPG088 | 1.434425892 | 2.676869643 |
| Q9CZT4 | Polr3e | 1.43402458 | 4.03401086 |
| O55060 | Tpmt | 1.431952325 | 3.913634814 |
| M1L511 | OPG098 | 1.425312595 | 2.913034744 |
| Q9EPK7;Q99NF8 | Xpo7 | 1.424161573 | 3.926270217 |
| Q9CQZ7 | Polr3k | 1.420443384 | 3.293772833 |
| Q6PE54 | Dhx40 | 1.417259344 | 3.646931545 |
| A0A7H0DN97 | OPG125 | 1.414728739 | 2.801800901 |
| Q9D0T2 | Dusp12 | 1.401473045 | 3.108814582 |
| Q3TC72 | Fahd2a | 1.401161236 | 5.297187852 |
| P56371 | Rab4a | 1.400772425 | 3.415288168 |
| Q8R123 | Flad1 | 1.399789756 | 4.774610965 |
| P62274 | Rps29 | 1.393875663 | 4.953649444 |
| M1LLA2 | OPG126 | 1.391551626 | 3.209034268 |
| A0A7H0DNE3;Q8V4T3 | OPG175 | 1.388422594 | 2.638233604 |
| P47964 | Rpl36 | 1.37828718 | 3.149826054 |
| Q922B1 | Macrod1 | 1.378065246 | 2.914687631 |
| Q8BI84 | Mia3 | 1.373083305 | 3.295317908 |
| P59470 | Polr3b | 1.357168208 | 2.659379512 |
| Q99JR1 | Sfxn1 | 1.35709799 | 3.274848418 |
| Q80UU9 | Pgrmc2 | 1.354329693 | 3.523018628 |
| P0DTN3 | OPG154 | 1.349533153 | 2.84225662 |
| Q8C5N5 | Pdcd2l | 1.346036144 | 3.828873872 |
| Q9ESW4 | Agk | 1.342262268 | 2.990675681 |
| Q45VK7 | Dync2h1 | 1.341388725 | 4.142594819 |
| A0A7H0DNA2 | OPG130 | 1.339756809 | 2.487983348 |
| A0A7H0DN56 | OPG084 | 1.331320128 | 2.472262881 |
| Q08122 | Tle3 | 1.330955494 | 4.778515777 |
| Q8BMJ3 | Eif1ax | 1.329553529 | 4.174482185 |
| Q8BUH1 | Txnl4b | 1.32851275 | 2.981551498 |
| Q8CCA0 | Dcun1d4 | 1.324499615 | 2.485863816 |
| P62305 | Snrpe | 1.317693718 | 2.84225662 |
| M1L9Q3 | OPG137 | 1.316518241 | 2.752460768 |
| A0A7H0DNF4 | OPG192 | 1.314764712 | 2.567330249 |
| Q8V4Y0;A0A7H0DN92 | OPG105 | 1.313837745 | 2.494411789 |
| Q571H0 | Urb1 | 1.305993279 | 2.879481802 |
| A0A7H0DN96 | OPG124 | 1.304440575 | 3.268331914 |
| P27659;E9PWZ3 | Rpl3 | 1.30406417 | 3.884878028 |
| Q9JKL4 | Ndufaf3 | 1.302055645 | 2.658972887 |
| A0A7H0DN37 | OPG064 | 1.290637557 | 3.058720411 |
| P60840 | Ensa | 1.289963474 | 3.623441504 |
| A0A7H0DN95 | OPG123 | 1.288528275 | 3.178514837 |
| Q9JIK9 | Mrps34 | 1.287155147 | 2.047485393 |
| Q9D8T7 | Slirp | 1.28644887 | 3.695788405 |
| Q9CXU9 | Eif1b | 1.28310677 | 4.08439046 |
| A0A7H0DN84 | OPG112 | 1.28299498 | 3.754598494 |
| O08848 | RO60 | 1.279232845 | 4.519134463 |
| Q922Q9 | Chid1 | 1.275478554 | 3.305263183 |
| A0A7H0DNA8 | OPG136 | 1.259088925 | 2.53852377 |
| Q8BIP0 | Dars2 | 1.252792541 | 4.639353751 |
| Q3UFY8 | Trmt10c | 1.252616924 | 4.48998414 |
| P55200 | Kmt2a | 1.24799588 | 2.186384531 |
| A0A7H0DND1 | OPG160 | 1.247865539 | 2.804767485 |
| Q9Z2Y3 | Homer1 | 1.233956259 | 2.728700876 |
| A0A7H0DNC8 | OPG157 | 1.210457523 | 2.248027033 |
| Q9QZX7 | Srr | 1.207663969 | 3.658404874 |
| Q9JJ94 | Ssna1 | 1.207345843 | 3.456228446 |
| Q8VED9 | Lgalsl | 1.200655874 | 2.542559771 |
| O35381;Q9EST5;Q64G17 | Anp32 | 1.200205636 | 3.050896642 |
| Q8CAK1 | Iba57 | 1.197550108 | 3.244279079 |
| Q9Z1X4 | Ilf3 | 1.194728466 | 4.56391268 |
| Q8C7Q4;Q8VE92 | Rbm4b | 1.190675152 | 5.15144695 |
| P56812 | Pdcd5 | 1.1862058 | 3.849663192 |
| Q6RI63 | Fam120b | 1.182265643 | 2.03557672 |
| Q64261 | Cdk6 | 1.181945927 | 4.010371626 |
| M1KJ27 | OPG129 | 1.180106396 | 2.586534278 |
| Q91V64 | Isoc1 | 1.179117803 | 4.158338398 |
| Q8C167 | Prepl | 1.177093759 | 3.394062731 |
| Q11136 | Pepd | 1.175764428 | 3.252609025 |
| A0A7H0DN91 | OPG119 | 1.172265184 | 2.40129726 |
| Q9D4J1;Q9D8Y0 | Efhd | 1.1715142 | 2.421221963 |
| Q8K368 | Fanci | 1.168088018 | 2.857591551 |
| Q9CXJ1 | Ears2 | 1.156163548 | 2.876443604 |
| Q9D880 | Timm50 | 1.152765926 | 2.53600612 |
| Q9CWW6 | Pin4 | 1.150935517 | 2.982187347 |
| Q9ERV1 | Mkrn2 | 1.150231094 | 4.416614507 |
| Q9D273 | Mmab | 1.148897252 | 2.604863179 |
| A0A7H0DNB6 | OPG144 | 1.146009216 | 2.155221938 |
| Q9JJI8 | Rpl38 | 1.138239691 | 4.56955956 |
| Q9JJV2 | Pfn2 | 1.138185007 | 4.521413266 |
| Q8BP67 | Rpl24 | 1.135356744 | 3.926071903 |
| Q99K46 | Usp11 | 1.129504093 | 2.977447971 |
| Q08122;Q62440;Q62441;Q9WVB2 | Tle1;Tle2 | 1.128676283 | 4.668674569 |
| Q9QXG2;Q9QZD5 | Chm | 1.124047467 | 3.146070578 |
| Q8CCK0 | Macroh2a2 | 1.12139497 | 3.494523014 |
| Q8C2P3 | Dus1l | 1.120762988 | 2.714725184 |
| Q3U2A8 | Vars2 | 1.114082314 | 3.059461219 |
| A0A7H0DNB7 | OPG145 | 1.109912353 | 3.695788405 |
| Q9EST4 | Psmg2 | 1.108669528 | 3.656310572 |
| Q8VCL2 | Sco2 | 1.108361652 | 2.130473032 |
| Q9DD18 | Dtd1 | 1.106918084 | 4.52620226 |
| A0A7H0DN29 | OPG056 | 1.105033559 | 3.474029774 |
| Q9Z2X2 | Psmd10 | 1.103274011 | 4.09102197 |
| Q8BYN3 | Itpk1 | 1.097113041 | 4.482015345 |
| P30681;P63158 | Hmgb | 1.09372515 | 2.789736065 |
| Q9DCV4 | Rmdn1 | 1.088148246 | 3.023177152 |
| Q8R2U4 | Ntmt1 | 1.08273857 | 4.344325268 |
| Q8BWJ3 | Phka2 | 1.080396483 | 2.228108284 |
| Q5DU02 | Usp22 | 1.06387437 | 3.962329852 |
| Q9CWQ0 | Dph5 | 1.061530556 | 3.215189126 |
| A0A7H0DNC3;Q8V4V3 | OPG151 | 1.057048504 | 2.670490748 |
| Q5SF07;Q9CPN8 | Igf2bp2 | 1.056483832 | 2.428964036 |
| Q3U0V1;Q91WJ8 | Khsrp | 1.055978074 | 5.280410943 |
| Q9ERA0 | Tfcp2 | 1.050934978 | 3.473924072 |
| Q9QXV1 | Cbx8 | 1.039515298 | 2.836793386 |
| A0A7H0DNA3 | OPG131 | 1.038816423 | 3.429109377 |
| A0A7H0DNB8 | OPG146 | 1.038428762 | 2.548740493 |
| P51912 | Slc1a5 | 1.034404607 | 3.368137934 |
| Q7TMY7 | Ipo8 | 1.033824545 | 4.355860661 |
| Q3UFF7 | Lyplal1 | 1.033813086 | 3.825666737 |
| A0A7H0DND2 | OPG161 | 1.032404539 | 2.28845801 |
| A0A7H0DN62 | OPG090 | 1.028076631 | 3.435424985 |
| Q3V0K9 | Strbp | 1.025466397 | 2.991427678 |
| Q91WM1 | Impdh1 | 1.02245265 | 4.125187546 |
| P50096 | Srsf9 | 1.021727007 | 3.264210042 |
| Q9D0B0 | Hsf1 | 1.021524158 | 4.29367792 |
| P38532 | Ndufa7 | 1.020631061 | 2.31633828 |
| Q9Z1P6 | Zcchc7 | 1.015304879 | 2.227432039 |
| B1AX39 | Cpne3 | 1.012778651 | 2.060618744 |
| Q9D6F9 | Xpo6 | 1.012662433 | 2.856855788 |
| Q8BT60 | Tfcp2 | 0.999975723 | 3.057579546 |
| Q924Z6 | Cbx8 | 0.992390632 | 3.722240743 |
| Q61474;Q920Q6 | Msi2 | 0.991341976 | 3.352807586 |
| A0A7H0DNC0 | OPG148 | 0.988925957 | 3.035150716 |
| Q91VL8 | Terf2ip | 0.986592587 | 3.864244963 |
| O70172 | Pip4k2a | 0.986232329 | 3.38904092 |
| A0A7H0DNB4 | OPG142 | 0.980226179 | 2.182944553 |
| A0A7H0DN90 | OPG118 | 0.97958503 | 2.749599229 |
| Q3SXD3 | Hddc2 | 0.974505332 | 3.103505744 |
| Q64331 | Myo6 | 0.97160026 | 4.396566281 |
| Q9ESZ8 | Gtf2i | 0.96242799 | 3.327407796 |
| Q9D6M3 | Slc25a22 | 0.962409415 | 2.985937986 |
| Q9CYR6 | Pgm3 | 0.958905831 | 4.147136808 |
| Q8VEJ9 | Vps4a | 0.956527683 | 3.943655471 |
| Q8VDW0;Q9Z1N5 | Ddx39 | 0.956506977 | 3.711019085 |
| Q8R088 | Golph3l | 0.955917848 | 3.223656013 |
| Q8BFY6 | Pef1 | 0.953855508 | 2.585699992 |
| Q3U0J8 | Tbc1d2b | 0.9517012 | 2.325883987 |
| P97823 | Lypla1 | 0.951361151 | 2.137947008 |
| M1LL92 | OPG116 | 0.950493085 | 3.36030188 |
| Q8CJ19 | Mical3 | 0.946035654 | 2.721442541 |
| A0A7H0DN20 | OPG047 | 0.944464883 | 3.58034661 |
| Q62406 | Irak1 | 0.942451682 | 2.08244737 |
| Q9CXS4 | Cenpv | 0.93954538 | 2.377340794 |
| Q499X9 | Mars2 | 0.937839934 | 2.474146882 |
| Q8K021 | Scamp1 | 0.935518656 | 3.273057947 |
| Q9D2M8 | Ube2v2 | 0.933339083 | 2.518934047 |
| A0A7H0DN85 | OPG113 | 0.932611061 | 3.023596436 |
| Q8VDS4 | Rprd1a | 0.928765217 | 4.043020992 |
| Q05186 | Rcn1 | 0.916599636 | 3.67654749 |
| Q99LG4 | Ttc5 | 0.913816862 | 3.564158238 |
| A0A7H0DN17 | OPG044 | 0.911794413 | 2.916313283 |
| P11031 | Sub1 | 0.911129983 | 2.263386189 |
| A0A7H0DN82 | OPG110 | 0.910049033 | 3.103505744 |
| Q8K284 | Gtf3c1 | 0.909048485 | 3.194698912 |
| Q8CA72 | Gan | 0.906687379 | 2.406921239 |
| Q99L02 | Pagr1a | 0.905175643 | 3.120665462 |
| P61967;Q9DB50 | Ap1s | 0.904630764 | 2.26124497 |
| P42225 | Stat1 | 0.904493329 | 3.653154839 |
| Q91Z53 | Grhpr | 0.895212563 | 3.745377227 |
| Q9ERI6 | Rdh14 | 0.892380363 | 2.225242145 |
| Q9CR09 | Ufc1 | 0.889768554 | 3.066288146 |
| Q3UMY5 | Eml4 | 0.88919598 | 3.884845377 |
| Q3TAA7 | Stk11ip | 0.88610713 | 2.71423185 |
| Q9DCN1 | Nudt12 | 0.884949402 | 2.764848976 |
| Q9CXY6 | Ilf2 | 0.877326076 | 3.468448964 |
| A0A7H0DNE4 | OPG176 | 0.871511943 | 3.971348663 |
| Q9ESE1 | Lrba | 0.86906529 | 4.03401086 |
| Q61205 | Pafah1b3 | 0.867321775 | 2.335657737 |
| A0A7H0DN77 | OPG105 | 0.866195383 | 2.172860219 |
| Q60649 | Clpb | 0.862008582 | 3.114095961 |
| Q9R0P4 | Smap | 0.861899669 | 3.27483205 |
| P61290 | Psme3 | 0.861693431 | 3.863357419 |
| A0A7H0DN49;Q8V518 | OPG077 | 0.858292334 | 2.119950165 |
| Q3TCJ1 | Abraxas2 | 0.858025592 | 3.216991014 |
| Q8R2R3 | Aagab | 0.857921239 | 2.133763518 |
| Q9D032;Q9CYZ8 | Ssbp2 | 0.857842222 | 2.097732248 |
| Q9CQV6 | Map1lc3b | 0.856840298 | 2.632970266 |
| A0A7H0DN39 | OPG066 | 0.856667918 | 2.194184232 |
| Q9CZ62 | Cep97 | 0.855187722 | 2.033227749 |
| Q99P31 | Hspbp1 | 0.853801305 | 4.09150297 |
| Q9CQ37 | Ube2t | 0.851997607 | 2.798516854 |
| P60710;P63260;P68033;P68134 | Act | 0.84909582 | 3.712375218 |
| Q9WUV0 | Orc5 | 0.847877912 | 3.032666154 |
| Q78RX3 | Smim12 | 0.845736534 | 2.264271484 |
| A0A7H0DNE0 | OPG172 | 0.844681136 | 3.429109377 |
| Q9Z2I0 | Letm1 | 0.844302358 | 2.368502272 |
| Q80ZI6 | Lrsam1 | 0.843449512 | 2.710782255 |
| Q8R3C0 | Mcmbp | 0.839123786 | 4.25764623 |
| Q9D4V7 | Rabl3 | 0.830240718 | 3.056806361 |
| Q8BTU1 | Cfap20 | 0.828857493 | 2.03067538 |
| Q5U3K5 | Rabl6 | 0.82846652 | 2.537959561 |
| Q99PM9 | Uck2 | 0.82508476 | 3.351231497 |
| Q9D7X3 | Dusp3 | 0.821551455 | 3.339160631 |
| P52019 | Sqle | 0.819946881 | 2.678935588 |
| Q9JI11 | Stk4 | 0.816758652 | 3.209034268 |
| Q91WD1 | Polr3d | 0.815672232 | 3.354520254 |
| Q9Z204 | Hnrnpc | 0.814194974 | 3.06014031 |
| Q9QXE0 | Hacl1 | 0.81404629 | 2.242469601 |
| Q9Z0E0 | Ncdn | 0.81199628 | 3.288531949 |
| Q9QUI0 | Rhoa | 0.81096295 | 2.179903101 |
| A0A7H0DN25 | OPG052 | 0.809780847 | 2.991877156 |
| Q8VDU0 | Gpsm2 | 0.809688673 | 3.217381827 |
| O54940;Q8BHE3 | Bnip2;Atcay | 0.809239789 | 2.180335101 |
| P04627;P28028;Q99N57 | Araf;Braf;Raf1 | 0.807604613 | 3.418439225 |
| Q6NVF0 | Ocrl | 0.803537669 | 3.695788405 |
| Q71RI9 | Kyat3 | 0.803491409 | 2.677001559 |
| Q9CQ89 | Cuta | 0.797551435 | 3.516768834 |
| Q11011 | Npepps | 0.796923653 | 4.112320344 |
| Q66JV4;Q80YR9 | Rbm12b | 0.7948672 | 2.433292602 |
| Q91YE3 | Egln1 | 0.791088548 | 2.767560019 |
| Q3ULD5 | Mccc2 | 0.788195184 | 2.08398285 |
| Q8VE95 |  | 0.783754941 | 2.284671465 |
| P70403 | Cux1 | 0.779974568 | 2.941450307 |
| O70152 | Dpm1 | 0.779951651 | 2.485863251 |
| Q8BZM1 | Glmn | 0.779894872 | 3.760306828 |
| Q9DB27;Q9CQ21 | Mcts | 0.779515502 | 2.431281459 |
| Q8R3D1 | Tbc1d13 | 0.779429214 | 4.667501416 |
| P22892 | Ap1g1 | 0.77551594 | 3.997243608 |
| P59016 | Vps33b | 0.775469567 | 2.869362674 |
| Q80W88 | Homez | 0.773770644 | 2.720055381 |
| Q9Z0H4 | Celf2 | 0.772170131 | 2.419129046 |
| Q61102 | Abcb7 | 0.764464887 | 2.097419308 |
| P11930 | Nudt19 | 0.763939203 | 2.138416925 |
| Q8VDM6 | Hnrnpul1 | 0.76359565 | 4.065637563 |
| Q8R4X3 | Rbm12 | 0.762865153 | 5.569291115 |
| Q3TXT3 | Inip | 0.762483177 | 3.382902526 |
| Q8BH55 | Thnsl1 | 0.758434136 | 2.299928228 |
| Q9DB85 | Rrp8 | 0.755078554 | 2.179903101 |
| Q8VI84 | Noc3l | 0.75343196 | 3.701195153 |
| P10639 | Txn | 0.752234081 | 2.863916699 |
| Q5ND52 | Mrm3 | 0.746015097 | 2.765282585 |
| P00493 | Hprt1 | 0.743940996 | 3.705070314 |
| Q8C5Q4 | Grsf1 | 0.730476045 | 2.372830825 |
| Q8R418 | Dicer1 | 0.728476809 | 2.850296099 |
| Q6PER3;Q8R001 | Mapre3 | 0.723486635 | 2.283884975 |
| G3XA57;Q9D620 | Rab11fip | 0.722376224 | 3.652137499 |
| Q8VBY2 | Camkk1 | 0.711125833 | 2.176895991 |
| Q80UM7 | Mogs | 0.707758847 | 2.140345641 |
| Q5U5Q9 | Uimc1 | 0.70521536 | 3.288978629 |
| Q9D483 | Polr3c | 0.69518211 | 2.741264625 |
| Q6PGA0 | Rcor3 | 0.695017957 | 2.871255946 |
| P22907 | Hmbs | 0.69456366 | 2.903558156 |
| Q91WG8 | Gne | 0.693185222 | 2.798383494 |
| Q9QY36 | Naa10 | 0.690148994 | 3.151413568 |
| O08663 | Metap2 | 0.688750782 | 3.440172329 |
| P70697 | Urod | 0.687166607 | 3.199062862 |
| P23506 | Pcmt1 | 0.684186587 | 3.473425837 |
| Q9Z1K6 | Arih2 | 0.681317558 | 2.923021959 |
| Q9D2C6 | Polr3h | 0.679515064 | 2.503905676 |
| Q8VC16 | Lrrc14 | 0.678411327 | 2.163910981 |
| Q8R3H9 | Ttc4 | 0.67810327 | 3.465348986 |
| Q9CQT7 | Desi1 | 0.677775048 | 3.045327451 |
| Q9EPN1;Q9ESE1 | Nbea | 0.675946507 | 2.804767485 |
| Q9D7S7 | Rpl22l1 | 0.674727426 | 2.559914934 |
| P50136 | Bckdha | 0.668407393 | 3.426139391 |
| Q8BZ36 | Rint1 | 0.663773715 | 2.020150611 |
| Q8C569 | Fam118b | 0.660562041 | 3.941049284 |
| Q921X6 | Polr3f | 0.66033147 | 3.83578365 |
| Q6DID3;Q7TSH6 | Scaf | 0.660159815 | 3.45509715 |
| Q60676 | Ppp5c | 0.656250196 | 4.045001285 |
| Q9DAJ4 | Wdr83 | 0.653280783 | 2.259858852 |
| Q3UDK1 | Trafd1 | 0.652754369 | 2.078990923 |
| Q9DBF7 | Cwc25 | 0.652498562 | 2.176114716 |
| Q9CXG3 | Ppil4 | 0.648806254 | 3.760114808 |
| Q9QYJ3 | Dnajb1 | 0.645639625 | 3.655950225 |
| Q7TND5 | Rpf1 | 0.638374159 | 2.624002121 |
| Q9JJT0 | Rcl1 | 0.63824459 | 3.456228446 |
| P0DTN4 | OPG035 | 0.63681896 | 3.133929925 |
| Q9D903 | Ebna1bp2 | 0.635018831 | 2.686730664 |
| Q61036;Q8CIN4 | Pak | 0.632496378 | 2.496135099 |
| Q3TJZ6 | Fam98a | 0.631015376 | 2.487983348 |
| P97801 | Smn1 | 0.628429865 | 2.985642353 |
| Q922H1 | Prmt3 | 0.625778919 | 2.275349069 |
| Q8K4L0 | Ddx54 | 0.623592253 | 3.548344642 |
| Q9CQQ4 | Gemin2 | 0.6190873 | 2.753005731 |
| Q9WUA6 | Akt3 | 0.618758645 | 2.105404739 |
| Q8BTW9 | Pak4 | 0.617618728 | 3.400220165 |
| M1L9M0 | OPG082 | 0.610098662 | 3.107757776 |
| P70280 | Vamp7 | 0.607573094 | 2.799466988 |
| Q60668 | Hnrnpd | 0.600302644 | 2.629855653 |
| Q8BFR5 | Tufm | 0.598411756 | 2.623425186 |
| Q6NZN0 | Rbm26 | 0.598180066 | 2.587253607 |
| Q8BGJ9;Q9D883 | U2af1 | 0.595202577 | 2.583402325 |
| P25425 | Pou2f1 | 0.592656255 | 2.630897889 |
| Q9JME5;Q9Z1T1 | Ap3b1 | 0.592579109 | 3.076525357 |
| Q9Z1G4 | Atp6v0a1 | 0.590616928 | 2.245861523 |
| O54879 | Hmgb3 | 0.589808604 | 2.696870264 |
| Q8BPG6 | Sumf2 | -0.585756912 | 2.457811077 |
| D3Z4I3;Q62176 | Rbm | -0.586345057 | 2.806595233 |
| Q8CI32 | Bag5 | -0.586581444 | 3.310489487 |
| Q8BHS3 | Rbm22 | -0.586618251 | 3.095340262 |
| Q9Z2U0 | Psma7 | -0.586937024 | 2.972316611 |
| Q8C525 | Mb21d2 | -0.587591439 | 2.151323894 |
| Q68FE6 | Ripor1 | -0.588924356 | 2.214410022 |
| Q99PT1 | Arhgdia | -0.59193179 | 3.136022283 |
| Q8CI61 | Bag4 | -0.592580868 | 3.546431381 |
| Q99MR0;Q9Z2N8 | Actl6 | -0.593081013 | 2.785058269 |
| P63085 | Mapk1 | -0.594389597 | 2.810307057 |
| Q80YV2 | Zc3hc1 | -0.594803579 | 3.863357419 |
| Q8R2U0 | Seh1l | -0.594983523 | 2.579614853 |
| P50518 | Atp6v1e1 | -0.595502722 | 3.364953294 |
| A2A791 | Zmym4 | -0.595609587 | 2.115885684 |
| Q9JIY2 | Cbll1 | -0.596211663 | 2.183009846 |
| P68037 | Ube2l3 | -0.596468226 | 2.579078009 |
| Q8R349 | Cdc16 | -0.596664112 | 3.954073685 |
| Q8CJF8;Q8CJF9;Q8CJG0;Q8CJG1 | Ago | -0.596684378 | 2.698843863 |
| P42230;P42232 | Stat5 | -0.598295355 | 2.434031632 |
| P63037 | Dnaja1 | -0.59888532 | 2.836770973 |
| Q80X82 | Sympk | -0.599098294 | 3.001046108 |
| Q9Z1K5 | Arih1 | -0.59950368 | 3.04624483 |
| Q9Z1R2 | Bag6 | -0.600057852 | 3.234858831 |
| P51150 | Rab7a | -0.601053331 | 2.601229478 |
| Q99NB8 | Ubqln4 | -0.601427563 | 2.86750694 |
| Q91YE6 | Ipo9 | -0.601473657 | 3.122178545 |
| E9Q236 | Abcc4 | -0.601770041 | 2.503905676 |
| Q8K5B2 | Mcfd2 | -0.601906449 | 2.375488257 |
| Q99JF8 | Psip1 | -0.601966571 | 3.847788825 |
| P49443 | Ppm1a | -0.602043125 | 2.787965472 |
| O54692 | Zw10 | -0.602647999 | 2.685585739 |
| Q9Z277 | Baz1b | -0.60577043 | 2.350662838 |
| Q571I9 | Aldh16a1 | -0.605785987 | 2.608118221 |
| Q9Z2U1 | Psma5 | -0.60631304 | 3.751479382 |
| Q99M28 | Rnps1 | -0.606437553 | 2.490021668 |
| Q8BGT7 | Smndc1 | -0.606500377 | 2.5668799 |
| Q9CWS4 | Ints11 | -0.607455612 | 3.564255074 |
| P70182 | Pip5k1a | -0.608755583 | 2.591741228 |
| Q8R307 | Vps18 | -0.609008687 | 2.900219868 |
| P80314 | Cct2 | -0.609150231 | 3.870828082 |
| Q8CF66 | Lamtor4 | -0.609696198 | 2.625936337 |
| Q3TDN2 | Faf2 | -0.610216 | 2.155587707 |
| Q8CGU1 | Calcoco1 | -0.610873104 | 2.15986315 |
| Q91YL3 | Uckl1 | -0.61098856 | 2.117734187 |
| Q8R0A0 | Gtf2f2 | -0.611543471 | 3.822122385 |
| Q8K4P0 | Wdr33 | -0.611787122 | 2.674847528 |
| Q7TMY4 | Thoc7 | -0.612271002 | 3.637487765 |
| Q9JHS4 | Clpx | -0.612402919 | 2.953147998 |
| Q9EQP2 | Ehd4 | -0.612568134 | 3.473924072 |
| Q7TPD1 | Fbxo11 | -0.613398183 | 2.288941948 |
| P59328 | Wdhd1 | -0.61364373 | 3.492310325 |
| Q9JLV6 | Pnkp | -0.613698772 | 2.399232123 |
| P60843;P10630 | Eif4a | -0.614364202 | 2.765229578 |
| P49025 | Cit | -0.614820232 | 2.769557247 |
| Q8C0E2 | Vps26b | -0.614995984 | 3.438878992 |
| P23198;Q61686 | Cbx5 | -0.615044391 | 3.246461291 |
| Q3V300 | Kif22 | -0.615364623 | 2.077484119 |
| Q9R190 | Mta2 | -0.615366084 | 2.675356552 |
| Q8R1J3 | Zcchc9 | -0.61555582 | 2.209559205 |
| Q9D8W5 | Psmd12 | -0.615736702 | 4.241090091 |
| Q6P8I4 | Pcnp | -0.616182671 | 2.713213212 |
| Q99JH1 | Rpp25l | -0.616375523 | 2.329241353 |
| Q924H7 | Wac | -0.616755508 | 3.120665462 |
| Q9CQG2 | Mettl16 | -0.61755372 | 2.256359783 |
| Q9D051 | Pdhb | -0.617568144 | 2.017436339 |
| O88967 | Yme1l1 | -0.617753988 | 2.024718323 |
| Q9CZ04 | Cops7a | -0.618661072 | 3.846460924 |
| Q8VCG3 | Wdr74 | -0.618880841 | 2.568465291 |
| Q99LG2 | Tnpo2 | -0.619056536 | 2.285619359 |
| Q8C2E7 | Washc5 | -0.619278594 | 3.040058451 |
| Q71LX4 | Tln2 | -0.61972467 | 2.631142022 |
| Q922P9 | Glyr1 | -0.619803812 | 2.248564417 |
| Q9D0D5 | Gtf2e1 | -0.619964309 | 3.045579831 |
| Q9QY76 | Vapb | -0.620625929 | 2.550110916 |
| P60710;P62737;P63260;P63268;P68033;P68134 | Acta2;Actg2 | -0.621109198 | 3.284265101 |
| Q99MN9 | Pccb | -0.621670853 | 2.455364796 |
| Q9QWY8 | Asap1 | -0.622222241 | 2.048750999 |
| Q8K4Z5 | Sf3a1 | -0.622489494 | 4.519134463 |
| Q8VEE0 | Rpe | -0.623516437 | 3.13889578 |
| Q8R323 | Rfc3 | -0.625505569 | 2.798876143 |
| Q9EP72 | Emc7 | -0.625670586 | 2.426645104 |
| Q8BGT5 | Gpt2 | -0.625690371 | 2.024709728 |
| Q9CQ43 | Dut | -0.62577397 | 2.652123322 |
| Q91WG4 | Elp2 | -0.626624967 | 3.836815499 |
| Q8R1Q8 | Dync1li1 | -0.627011378 | 4.102143461 |
| Q9JHR7 | Ide | -0.628775894 | 3.647210969 |
| P47968 | Rpia | -0.628900793 | 2.494506048 |
| A0A7H0DN04 | OPG030 | -0.629950658 | 3.312153122 |
| Q9CRB9 | Chchd3 | -0.630047875 | 2.728612609 |
| P05132;P68181 | Prkac | -0.630165639 | 2.97865981 |
| P97434 | Mprip | -0.630257751 | 3.357585047 |
| Q8K057 | Ift80 | -0.630697452 | 2.950248419 |
| Q9Z2X8 | Keap1 | -0.631558167 | 3.506834339 |
| Q8BTZ4 | Anapc5 | -0.632001273 | 2.655774066 |
| Q9CX34 | Sugt1 | -0.63235694 | 2.99337286 |
| P46467;Q8BPY9;Q8VEJ9 | Vps4b;Flgnl1 | -0.63285283 | 2.754624667 |
| P53995 | Anapc1 | -0.634724761 | 3.714132432 |
| Q04207 | Rela | -0.634930915 | 2.901093035 |
| Q8BTI7 | Ankrd52 | -0.635177002 | 3.333867882 |
| P63154 | Crnkl1 | -0.635387673 | 3.31025339 |
| Q60996 | Ppp2r5c | -0.635427688 | 2.306305277 |
| A0A7H0DNF9 | OPG199 | -0.637156109 | 3.028129832 |
| Q9R0E2 | Plod1 | -0.639235849 | 2.759131709 |
| Q9WUM4 | Coro1c | -0.639812369 | 3.106574663 |
| P62700 | Ypel5 | -0.640129524 | 3.414332145 |
| O88895 | Hdac3 | -0.640271952 | 2.847412954 |
| P60898 | Polr2i | -0.640499331 | 2.934910735 |
| Q5SVQ0 | Kat7 | -0.640541229 | 2.956812909 |
| Q6ZQ93 | Usp34 | -0.640797596 | 3.010351125 |
| Q61753 | Phgdh | -0.641365206 | 3.570876313 |
| P08556;P32883;Q61411 | Nras;Kras;Hras | -0.641480692 | 2.427316311 |
| Q78PG9 | Ccdc25 | -0.641966573 | 2.372830825 |
| P34152 | Ptk2 | -0.642187106 | 2.519303924 |
| P46938 | Yap1 | -0.642377583 | 3.057579546 |
| Q9D4H2 | Gcc1 | -0.642543117 | 3.462215544 |
| A2AWA9 | Rabgap1 | -0.643261239 | 2.998966427 |
| P63001 | Rac1 | -0.64328963 | 2.430196074 |
| Q7TPM1 | Prrc2b | -0.647267098 | 2.470767518 |
| Q8K4B0 | Mta1 | -0.647590231 | 2.857256837 |
| P99027 | Rplp2 | -0.649198804 | 4.202329354 |
| Q8R1Q8;Q6PDL0 | Dync1li2 | -0.649657285 | 2.345947374 |
| Q0VGB7 | Ppp4r2 | -0.650071285 | 3.890260886 |
| A2A5R2 | Arfgef2 | -0.650074614 | 3.780615316 |
| P63147;Q9Z255 | Ube2 | -0.650479068 | 2.159709038 |
| C0HKD8;C0HKD9 | Mfap1 | -0.651390638 | 4.56391268 |
| Q8K400 | Stxbp5 | -0.651543894 | 2.439950537 |
| P16381;Q61496;Q62095;Q62167 | Ddx4;Ddx3y;Ddx3x | -0.652385545 | 2.083073453 |
| Q9Z110 | Aldh18a1 | -0.652907419 | 2.384002132 |
| Q9Z1X9 | Cdc45 | -0.653005401 | 3.376809059 |
| Q9DBG7 | Srpra | -0.6534418 | 2.004265547 |
| O88559 | Men1 | -0.654351981 | 2.711937036 |
| Q8R480 | Nup85 | -0.654386382 | 3.51982275 |
| Q8BWQ6 | Vps35l | -0.654666698 | 3.489216846 |
| Q61578 | Fdxr | -0.655056745 | 2.292740225 |
| Q8K2Q9 | Shtn1 | -0.655778241 | 3.330352982 |
| Q6ZWX6 | Eif2s1 | -0.655928578 | 4.133548754 |
| Q9ESX5 | Dkc1 | -0.656208081 | 2.89371796 |
| Q99LP6 | Grpel1 | -0.656674411 | 2.272834774 |
| Q9CQE6 | Asf1a | -0.656911446 | 5.375668682 |
| Q8C033 | Arhgef10 | -0.657097676 | 2.806595233 |
| P70399 | Tp53bp1 | -0.657665755 | 3.052719313 |
| P70175 | Dlg3 | -0.65771348 | 3.118279675 |
| Q8BIF2;Q8BP71;Q9JJ43 | Rbfox | -0.658139688 | 4.004647294 |
| P35279;P61294 | Rab6 | -0.658976794 | 2.106035552 |
| C0HKE1;C0HKE2;C0HKE3;C0HKE4;C0HKE5;C0HKE6;C0HKE7;C0HKE8;C0HKE9;Q64523;Q6GSS7;Q8BFU2;Q8CGP5;Q8CGP6;Q8CGP7;Q8R1M2 | H2ac;Hist1h2 | -0.659356235 | 2.846971599 |
| O09110 | Map2k3 | -0.660211246 | 2.76338963 |
| Q3TCH7 | Cul4a | -0.6612798 | 3.457450454 |
| Q8BR65 | Suds3 | -0.661963624 | 2.662274334 |
| P57780 | Actn4 | -0.662039853 | 3.3751259 |
| Q91Z67 | Srgap2 | -0.662734852 | 2.507036881 |
| Q8C156 | Ncaph | -0.664093252 | 2.689078847 |
| Q99L45 | Eif2s2 | -0.664299219 | 3.528855408 |
| Q9QXE7 | Tbl1x | -0.665679888 | 4.03401086 |
| O70400 | Pdlim1 | -0.666688345 | 4.138485766 |
| Q91VD9 | Ndufs1 | -0.666731688 | 2.494680987 |
| Q9R1K9 | Cetn2 | -0.667053758 | 2.26026944 |
| Q8K339 | Kin | -0.667246924 | 2.420589529 |
| Q91WN1 | Dnajc9 | -0.667746851 | 2.686238416 |
| P62715;P63330 | Ppp2c | -0.668418758 | 2.979066167 |
| Q8BJU0 | Ube2v1 | -0.670193155 | 2.321509786 |
| Q9JL62 | Gltp | -0.670310458 | 2.215759631 |
| P61600 | Naa20 | -0.670373679 | 3.565496518 |
| Q03147 | Cdk7 | -0.670732533 | 3.583601983 |
| Q8BP48 | Metap1 | -0.672945678 | 3.723247426 |
| Q9CWJ9 | Atic | -0.673173754 | 3.650678066 |
| Q99K28 | Arfgap2 | -0.673544944 | 4.833270278 |
| Q8K2D6 | Dctd | -0.673691102 | 2.487084617 |
| Q6PE01 | Snrnp40 | -0.674157334 | 3.828873872 |
| Q921C5 | Bicd2 | -0.674267124 | 3.611079173 |
| Q9WUU9 | Mcm3ap | -0.67498936 | 3.513415023 |
| Q6PDQ2 | Chd4 | -0.675304192 | 3.249148195 |
| Q8R2N2 | Utp4 | -0.675734672 | 3.270995882 |
| Q6A026 | Pds5a | -0.676410704 | 3.235785658 |
| O88522 | Ikbkg | -0.676617995 | 2.728612609 |
| Q8BY71 | Hat1 | -0.676835278 | 3.310489487 |
| Q3UGR5 | Hdhd2 | -0.676994114 | 2.440199156 |
| Q8K409 | Polb | -0.677116595 | 3.428809419 |
| Q61025 | Ift20 | -0.677198824 | 2.635276417 |
| Q3URQ0 | Tex10 | -0.677482342 | 2.059970881 |
| Q9ESV0 | Ddx24 | -0.679031041 | 2.702739421 |
| P28658 | Atxn10 | -0.679610818 | 2.572386455 |
| Q80U93 | Nup214 | -0.681075709 | 3.107744365 |
| Q9R099 | Tbl2 | -0.681549369 | 2.023422968 |
| P16056 | Met | -0.682927272 | 2.4781011 |
| P63038 | Hspd1 | -0.683373741 | 2.726048838 |
| Q9D0L8 | Rnmt | -0.685229304 | 2.480510811 |
| Q7TMI3 | Uhrf2 | -0.685632248 | 2.240724487 |
| Q8BHJ5;Q9QXE7 | Tbl1xr1 | -0.685658284 | 3.191252304 |
| Q8BWZ3 | Naa25 | -0.685844965 | 2.523756919 |
| Q3UMC0 | Afg2a | -0.686315436 | 3.660537061 |
| O70310 | Nmt1 | -0.687064751 | 3.466323412 |
| Q9CSN1 | Snw1 | -0.68712477 | 3.221188158 |
| Q9EQH3 | Vps35 | -0.687484602 | 3.474029774 |
| Q8BZX4 | Srek1 | -0.687705168 | 3.618149593 |
| Q9EQM6 | Dgcr8 | -0.687945921 | 3.750865948 |
| O09110;P70236 | Map2k6 | -0.688284463 | 3.36409455 |
| Q923D5 | Wbp11 | -0.689989464 | 3.943029687 |
| Q99KY4 | Gak | -0.690901682 | 3.71344449 |
| Q8K3X4;E9Q1P8 | Ir2bp | -0.690955968 | 2.017570472 |
| A0A7H0DN73 | OPG101 | -0.692237526 | 2.747639929 |
| P55258 | Rab8a | -0.692459073 | 2.494984685 |
| P31938;Q63932 | Map2k | -0.692646714 | 3.324918476 |
| Q99KH8;Q9Z2W1 | Stk2 | -0.692871575 | 3.418366443 |
| P47809 | Map2k4 | -0.692992003 | 3.004741019 |
| Q9ESK4 | Ing2 | -0.693283348 | 2.234777409 |
| P60335 | Pcbp1 | -0.694282319 | 4.933474852 |
| O70126 | Aurkb | -0.695248967 | 2.412584214 |
| Q8BVE3 | Atp6v1h | -0.695316716 | 3.482256243 |
| Q91XU3 | Pip4k2c | -0.69545919 | 2.174608653 |
| Q9DCB8 | Isca2 | -0.695924125 | 2.52110936 |
| P62334 | Psmc6 | -0.696264654 | 4.060997583 |
| P58929 | Gmeb2 | -0.696375422 | 2.417306531 |
| P42208 | Septin2 | -0.696528135 | 3.486483846 |
| Q8CH18 | Ccar1 | -0.696667431 | 3.581609683 |
| Q99JW4 | Lims1 | -0.697338848 | 2.862690165 |
| Q9CPR7 | Sike1 | -0.697586706 | 2.932248799 |
| Q8R332 | Nup58 | -0.698537167 | 2.869634113 |
| Q91Y44;Q9ESU6;Q8K2F0 | Brd | -0.698772882 | 2.443747976 |
| Q8BJW6 | Eif2a | -0.69939292 | 4.489390806 |
| Q8VEH6 | Zng1 | -0.699577375 | 2.606270351 |
| P39447 | Tjp1 | -0.700254354 | 3.488954172 |
| Q8VEJ1 | Gpn2 | -0.700508807 | 2.910902767 |
| P51807 | Dynlt1 | -0.700666804 | 3.166866133 |
| Q99KK7 | Dpp3 | -0.700798621 | 3.306812666 |
| Q99LM9 | Tada1 | -0.701008756 | 2.843619388 |
| P62869 | Elob | -0.7014496 | 3.997067313 |
| Q62203 | Sf3a2 | -0.701701786 | 3.413819275 |
| Q8BG81 | Poldip3 | -0.701717395 | 4.037684761 |
| P61021 | Rab5b | -0.702025347 | 3.462215544 |
| Q9R226 | Khdrbs3 | -0.702327636 | 2.229887069 |
| Q9EP89 | Lactb | -0.703251047 | 2.746011386 |
| Q9Z2G0 | Fem1b | -0.703421579 | 2.576610617 |
| Q60838 | Dvl2 | -0.70461417 | 3.477420694 |
| P62814 | Atp6v1b2 | -0.705169199 | 4.024317461 |
| Q9DCJ1 | Mlst8 | -0.705308063 | 4.430085309 |
| Q8BKI2 | Tnrc6b | -0.705659594 | 3.618382813 |
| Q9CQA5 | Med4 | -0.706117874 | 3.568753141 |
| Q8K0D5 | Gfm1 | -0.707171201 | 2.836793386 |
| Q0VEJ0 | Cep76 | -0.707515575 | 2.296323147 |
| Q9D898 | Arpc5l | -0.707531587 | 3.622104865 |
| Q61235 | Sntb2 | -0.708039521 | 2.564078804 |
| Q8CHT3 | Ints5 | -0.708332839 | 2.497201427 |
| Q91VJ4 | Stk38 | -0.70879763 | 2.246283683 |
| O55013 | Trappc3 | -0.708984853 | 2.531946768 |
| Q61048 | Wbp4 | -0.709188656 | 2.7782893 |
| Q8VCD5 | Med17 | -0.710252393 | 3.138962129 |
| B9EKI3 | Tmf1 | -0.710417236 | 3.351231497 |
| Q8BMC4 | Nop9 | -0.710589729 | 2.061097248 |
| Q91VT1 | Nsmce2 | -0.710928632 | 2.68445133 |
| Q99J77 | Nans | -0.710940367 | 2.410587945 |
| O35226 | Psmd4 | -0.711538779 | 3.361085658 |
| Q922S8;Q8C0N1 | Kif2 | -0.711651856 | 2.52310294 |
| Q7TMX5 | Shq1 | -0.711699409 | 2.64993696 |
| Q9D6J6 | Ndufv2 | -0.711717608 | 2.406360254 |
| P58404;Q9ERG2 | Stm | -0.711733538 | 2.933393341 |
| Q9Z0X1 | Aifm1 | -0.712725711 | 2.413417836 |
| Q9JLT4;Q9JMH6 | Tnxrd | -0.713162057 | 2.801800901 |
| Q61206 | Pafah1b2 | -0.713955282 | 3.199286214 |
| Q9CZ82 | Med18 | -0.714371454 | 2.968670587 |
| Q9CZU6 | Cs | -0.715227115 | 2.423011267 |
| P67984 | Rpl22 | -0.715241898 | 2.964812876 |
| Q61035 | Hars1 | -0.715404597 | 3.441677862 |
| P84091 | Ap2m1 | -0.716174553 | 3.536492351 |
| Q8R3G1 | Ppp1r8 | -0.716561967 | 4.285322638 |
| Q6PGG6 | Gnl3l | -0.716590865 | 3.087402779 |
| P25799 | Nfkb1 | -0.717289149 | 3.462472292 |
| Q9CWU9 | Nup37 | -0.717836148 | 3.602774865 |
| Q8K1Z0 | Coq9 | -0.718045316 | 2.576610617 |
| Q9R1R2 | Trim3 | -0.718114279 | 2.170049693 |
| Q9JHS9 | Cwc15 | -0.718293244 | 4.098866019 |
| Q9WV98 | Timm9 | -0.718570098 | 2.755565545 |
| Q80TA6 | Mtmr12 | -0.719423976 | 3.022694938 |
| Q9Z0N1;Q9Z0N2 | Eif2s3x | -0.719702429 | 4.581748837 |
| Q61553 | Fscn1 | -0.720159662 | 4.091657855 |
| Q63829 | Commd3 | -0.72036296 | 3.624988825 |
| O35685 | Nudc | -0.721034646 | 3.928041713 |
| Q99KE1 | Me2 | -0.721074675 | 2.918949621 |
| P46061 | Rangap1 | -0.721808632 | 3.718578439 |
| P40336 | Vps26a | -0.722784171 | 3.253695642 |
| Q9JJC6 | Rilpl1 | -0.7228157 | 3.083717891 |
| P46638;P62492 | Rab11 | -0.7228663 | 3.107744365 |
| P69566 | Ranbp9 | -0.72405222 | 3.823581864 |
| P70336 | Rock2 | -0.724087544 | 3.562120404 |
| Q923B1 | Dbr1 | -0.725121151 | 3.355984575 |
| Q6IRU5 | Cltb | -0.72523321 | 4.142594819 |
| Q6PHQ8 | Naa35 | -0.726029309 | 2.559456518 |
| Q9WUK4 | Rfc2 | -0.726701388 | 4.533252445 |
| Q6P1F6;Q925E7 | Ppp2r2 | -0.727054116 | 2.65080434 |
| Q8C092 | Taf5 | -0.727551591 | 3.561009575 |
| Q6ZWR4;Q6P1F6;Q925E7 | Ppp2rb | -0.727649261 | 2.05046344 |
| P62331 | Arf6 | -0.727717022 | 3.474029774 |
| Q8C1A5 | Thop1 | -0.727794759 | 3.878515178 |
| P42567 | Eps15 | -0.728556397 | 4.639172554 |
| Q8BTI8 | Srrm2 | -0.729390352 | 2.406643687 |
| Q9DBT5 | Ampd2 | -0.73030195 | 3.218001956 |
| Q6A028 | Swap70 | -0.730833596 | 2.981551498 |
| Q3TIR1 | Trappc13 | -0.73114024 | 2.563306367 |
| P97440 | Slbp | -0.731522157 | 2.379108596 |
| O88990;P57780;Q7TPR4;Q9JI91 | Actn | -0.731625238 | 3.532501992 |
| Q9CQY1 | Atg12 | -0.732412939 | 2.14799969 |
| P59999 | Arpc4 | -0.732947147 | 3.104653351 |
| Q9WVQ5 | Apip | -0.733246608 | 2.415297853 |
| Q9D1K7 | Adissp | -0.733934941 | 3.044006115 |
| P30999 | Ctnnd1 | -0.734237944 | 4.033884815 |
| P24288 | Bcat1 | -0.734912957 | 2.852571139 |
| Q9WV80 | Snx1 | -0.735612406 | 4.696281617 |
| O09106;P70288 | Hdac | -0.73585713 | 3.506026962 |
| P61161 | Actr2 | -0.735861974 | 4.433360725 |
| Q9DAW9 | Cnn3 | -0.736015952 | 3.349429656 |
| Q91WM3 | Rrp9 | -0.736891661 | 2.857863665 |
| Q61598 | Gdi2 | -0.737294001 | 3.73598328 |
| O55128 | Sap18 | -0.738032125 | 3.65255687 |
| Q8R2M2 | Dnttip2 | -0.73895254 | 3.91854463 |
| P26638 | Sars1 | -0.740897792 | 3.471245769 |
| Q80YA3 | Ddhd1 | -0.741320616 | 3.280284657 |
| P53810 | Pitpna | -0.741720968 | 2.925013998 |
| Q9JIY5 | Htra2 | -0.744559479 | 2.156579735 |
| P18654 | Rps6ka3 | -0.745147678 | 3.328649359 |
| P0C090;Q4VGL6 | Rc3h | -0.745971236 | 2.768457938 |
| Q8K296 | Mtmr3 | -0.746008801 | 3.012231411 |
| Q01768 | Nme2 | -0.746428515 | 2.836120087 |
| Q9CQA3 | Sdhb | -0.746659845 | 2.220564651 |
| Q3UPL0 | Sec31a | -0.747271132 | 4.046706882 |
| Q8VEH8 | Erlec1 | -0.748229909 | 2.143924652 |
| Q9DBU6 | Rsrc1 | -0.749094744 | 2.109390051 |
| P22315 | Fech | -0.749724693 | 2.542405026 |
| Q9JMA1 | Usp14 | -0.750911218 | 3.696569095 |
| Q501J7 | Phactr4 | -0.751450398 | 2.372830825 |
| Q9D892 | Itpa | -0.75160473 | 3.403725202 |
| Q9WV60 | Gsk3b | -0.751880507 | 2.761649154 |
| Q3UJD6 | Usp19 | -0.751909267 | 3.255090432 |
| Q9Z2L7 | Crlf3 | -0.752027394 | 2.984763843 |
| Q61081 | Cdc37 | -0.752268357 | 4.536852705 |
| P35922;Q9WVR4 | Fmr1;Fxr2 | -0.753047863 | 2.749827115 |
| Q8C166 | Cpne1 | -0.753361259 | 2.925517539 |
| Q3V1V3 | Esf1 | -0.753463878 | 3.810996172 |
| Q7TPD0 | Ints3 | -0.753793054 | 3.647652241 |
| Q99JX7 | Nxf1 | -0.755593579 | 2.956422031 |
| P62835;Q99JI6 | Rap1 | -0.756196526 | 2.687059677 |
| C0HKE1;C0HKE2;C0HKE3;C0HKE4;C0HKE5;C0HKE6;C0HKE7;C0HKE8;C0HKE9;P27661;Q64523;Q6GSS7;Q8BFU2;Q8CGP5;Q8CGP6;Q8CGP7;Q8R1M2 | H2 | -0.756643221 | 2.912074778 |
| P62627 | Dynlrb1 | -0.757140264 | 3.27483205 |
| Q9CQD1 | Rab5a | -0.757412525 | 3.713868353 |
| O35239 | Ptpn9 | -0.758332377 | 2.306305277 |
| P97494 | Gclc | -0.759405177 | 2.71423185 |
| Q91VM3 | Wdr45 | -0.759506907 | 2.437519149 |
| Q7TMF3 | Ndufa12 | -0.759679912 | 2.355339801 |
| P61358 | Rpl27 | -0.759836753 | 3.713288757 |
| Q920D3 | Med28 | -0.759956645 | 2.362632618 |
| O08749 | Dld | -0.761110235 | 3.009245126 |
| P24369 | Ppib | -0.761723728 | 2.676556926 |
| O55143;Q64518 | Atp2a | -0.762195634 | 2.755979885 |
| Q9QZQ1 | Afdn | -0.762233341 | 3.085655806 |
| Q9DAS9 | Gng12 | -0.764960344 | 2.460794555 |
| O55106 | Strn | -0.765775717 | 3.672012816 |
| Q99JR8 | Smarcd2 | -0.765966662 | 3.254728226 |
| Q9QXZ0 | Macf1 | -0.766190633 | 2.888468551 |
| Q7TSV4 | Pgm2 | -0.768314652 | 2.799606547 |
| Q9CYG7 | Tomm34 | -0.768363071 | 2.623895764 |
| Q9Z0R6 | Itsn2 | -0.768452925 | 2.949155314 |
| P47802 | Mtx1 | -0.768837153 | 2.097419308 |
| Q9JI10;Q9JI11 | Stk | -0.768957725 | 3.748695666 |
| Q01320 | Top2a | -0.769465271 | 2.567944834 |
| Q569Z6 | Thrap3 | -0.770104625 | 3.305263183 |
| P62073 | Timm10 | -0.770433278 | 2.567571829 |
| Q99KX1 | Mlf2 | -0.773285914 | 2.548383399 |
| Q61122 | **Nab1** | -0.773584313 | 3.279006613 |
| Q922K7 | Nop2 | -0.774046781 | 3.429109377 |
| Q9WU78 | Pdcd6ip | -0.774186201 | 3.921191306 |
| Q8R2N0 | Ccdc59 | -0.774738058 | 3.002500033 |
| Q9D2U5 | Naa38 | -0.775252374 | 2.630897889 |
| F7BJB9 | Morc3 | -0.776251451 | 4.024935547 |
| O35343;O35344 | Kpna | -0.777575795 | 2.435673222 |
| Q9JIX8 | Acin1 | -0.777699762 | 3.91854463 |
| Q64327 | Mea1 | -0.778003198 | 2.53076099 |
| Q52KI8 | Srrm1 | -0.778233076 | 3.761064182 |
| Q5H8C4 | Vps13a | -0.778365963 | 2.541814047 |
| Q9WVE8 | Pacsin2 | -0.77858398 | 3.713032048 |
| Q8BPM2 | Map4k5 | -0.778594235 | 3.260593185 |
| P59325 | Eif5 | -0.779473387 | 3.847433276 |
| Q9D7W5 | Med8 | -0.780650769 | 2.948519767 |
| Q8BMF4 | Dlat | -0.78071433 | 2.226077201 |
| Q8BGF7 | Pan2 | -0.78080265 | 3.33509565 |
| Q8QZZ8;Q91YQ1;Q9CZE3 | Rab | -0.781170953 | 2.504964929 |
| Q3UHX2 | Pdap1 | -0.781385988 | 3.849663192 |
| Q61466;Q6P9Z1;Q99JR8 | Smarcd | -0.782774985 | 4.266968692 |
| Q9ERU9 | Ranbp2 | -0.782997569 | 3.898463943 |
| O08528 | Hk2 | -0.783299164 | 3.261064329 |
| P51949 | Mnat1 | -0.783508863 | 3.255998047 |
| P62874 | Gnb1 | -0.784030769 | 2.787105891 |
| O88879 | Apaf1 | -0.784309152 | 2.333002204 |
| Q80YV3 | Trrap | -0.784396596 | 4.228429098 |
| Q9CY97 | Ssu72 | -0.786745297 | 2.595216882 |
| Q99LE6 | Abcf2 | -0.788732158 | 3.052719313 |
| P62915 | Gtf2b | -0.789434369 | 4.868785445 |
| Q4VC33 | Maea | -0.789533714 | 3.546298103 |
| P41209;Q9R1K9 | Cetn1 | -0.790573868 | 3.029090849 |
| E9Q816 | Cyp2w1 | -0.790724597 | 2.109774233 |
| Q5U430 | Ubr3 | -0.791260262 | 2.716313356 |
| Q6ZPE2 | Sbf1 | -0.791346538 | 2.447638823 |
| D3YYU8 | Obsl1 | -0.791356726 | 2.810307057 |
| Q8BRN9 | Cc2d1b | -0.791570155 | 2.531541849 |
| Q8C5N3 | Cwc22 | -0.791590213 | 4.759418207 |
| Q05816 | Fabp5 | -0.791593997 | 2.836793386 |
| Q6ZWZ2 | Ube2r2 | -0.792031655 | 3.791671347 |
| Q7TQK1 | Ints7 | -0.792319902 | 3.941049284 |
| Q9D1G2 | Pmvk | -0.792529753 | 2.196634021 |
| Q6PA06 | Atl2 | -0.793697545 | 2.110239521 |
| Q5HZJ0 | Drosha | -0.793708633 | 3.240849705 |
| Q99LI8 | Hgs | -0.794774664 | 3.340876839 |
| Q9CQW2;Q8VEH3 | Arl8 | -0.794913093 | 3.245626802 |
| P46471 | Psmc2 | -0.795201064 | 4.673220337 |
| P22682 | Cbl | -0.795444193 | 3.061743443 |
| Q80UK8 | Ints2 | -0.795745063 | 2.743914748 |
| Q9QZB9 | Dctn5 | -0.796978791 | 4.48142587 |
| Q8C0J2 | Atg16l1 | -0.7973497 | 3.488469275 |
| O35127 | Grcc10 | -0.798695832 | 2.829308012 |
| Q922V4 | Plrg1 | -0.798776292 | 2.972163495 |
| A0A7H0DNC2;Q8V4V4 | OPG150 | -0.799611284 | 4.033862974 |
| P97496 | Smarcc1 | -0.800126349 | 3.593040908 |
| Q9CQK7 | Rwdd1 | -0.800201491 | 2.662505611 |
| P62821 | Rab1a | -0.800950031 | 2.460794555 |
| Q8CC88 | Vwa8 | -0.801125436 | 3.483467608 |
| Q91Z67;Q91Z69 | Srgap1 | -0.801402679 | 2.481667214 |
| O55222 | Ilk | -0.801909156 | 3.884684868 |
| Q9JIH2 | Nup50 | -0.802269175 | 2.722489644 |
| Q6TYB5 | Fez2 | -0.802409096 | 2.720632501 |
| P53564 | Cux1 | -0.804197371 | 3.013180793 |
| Q4FK66 | Prpf38a | -0.805155921 | 3.717640664 |
| Q3UM18 | Lsg1 | -0.806270867 | 4.003533613 |
| Q922X9 | Prmt7 | -0.807039659 | 2.686185798 |
| Q8VHE0 | Sec63 | -0.807116267 | 2.229448164 |
| Q9CYA6 | Zcchc8 | -0.807396316 | 5.705887197 |
| Q8BUK6 | Hook3 | -0.807490184 | 4.32367048 |
| Q9WUA3 | Pfkp | -0.808094527 | 3.761064182 |
| Q9CT10 | Ranbp3 | -0.808333402 | 3.491492343 |
| P59708 | Sf3b6 | -0.808501577 | 3.474029774 |
| Q8BYH7 | Tbc1d17 | -0.808530767 | 2.884288074 |
| P48754 | Brca1 | -0.808541626 | 3.114676111 |
| Q63810 | Ppp3r1 | -0.808560623 | 3.391700465 |
| Q9DC16 | Ergic1 | -0.808630254 | 2.166538626 |
| Q80Y17 | Llgl1 | -0.809232788 | 4.94399702 |
| Q9CQC8 | Spg21 | -0.809835026 | 2.872212454 |
| Q99ME2 | Wdr6 | -0.810327973 | 3.166646893 |
| Q8BY87 | Usp47 | -0.810735397 | 2.677063302 |
| Q9EPU0 | Upf1 | -0.811071121 | 4.366797517 |
| Q60710 | Samhd1 | -0.811604727 | 3.753674363 |
| Q8BH74 | Nup107 | -0.811987077 | 3.465393733 |
| O54946;Q9QYI5 | Dnajb | -0.812111681 | 3.015158646 |
| O54931 | Pakap | -0.812658709 | 3.748695666 |
| Q6PFD9 | Nup98 | -0.812765733 | 3.564633027 |
| Q6PEB6 | Mob4 | -0.812813021 | 3.36409455 |
| Q8C050 | Rps6ka5 | -0.812969103 | 3.071025651 |
| Q8BL66 | Eea1 | -0.813180271 | 3.318032339 |
| P62843 | Rps15 | -0.814062562 | 2.888193088 |
| Q99J62 | Rfc4 | -0.81431631 | 4.993940729 |
| Q9D358 | Acp1 | -0.814434997 | 3.049164468 |
| P97376 | Frg1 | -0.814467725 | 3.800835871 |
| Q8C1B7;Q9R1T4 | Septin11;Septin6 | -0.814473491 | 4.047882757 |
| P0C0A3 | Chmp6 | -0.815786457 | 2.019910435 |
| Q8QZS3 | Flcn | -0.818749889 | 2.606575535 |
| P24788 | Cdk11b | -0.819587982 | 4.574594509 |
| Q91ZU1 | Asb6 | -0.819607985 | 4.002145646 |
| Q8BLR9 | Hif1an | -0.820919565 | 2.11628558 |
| Q8BGD9 | Eif4b | -0.822013745 | 3.159640133 |
| Q80T85 | Dcaf5 | -0.822036045 | 2.698843863 |
| Q8BGQ7 | Aars1 | -0.822756963 | 3.199311583 |
| Q8C079 | Strip1 | -0.823045137 | 2.693909499 |
| Q60865 | Caprin1 | -0.823067945 | 4.74230084 |
| P97287 | Mcl1 | -0.823961247 | 2.081812412 |
| Q99J95 | Cdk9 | -0.824022343 | 3.558554591 |
| Q8CG48 | Smc2 | -0.824946427 | 4.931096713 |
| Q64012 | Raly | -0.825102225 | 2.900219868 |
| Q8QZY9 | Sf3b4 | -0.825233755 | 2.548727809 |
| Q62422 | Ostf1 | -0.825423986 | 3.317161291 |
| Q8R3V5 | Sh3glb2 | -0.825667473 | 3.700075902 |
| P16381;Q501J6;Q61656;Q62095;Q62167 | Ddx17;Ddx5 | -0.825775237 | 2.602265728 |
| P49717 | Mcm4 | -0.826212254 | 4.443807798 |
| Q9CR60 | Golt1b | -0.826662383 | 3.320182088 |
| Q8CI51 | Pdlim5 | -0.826693534 | 4.112998478 |
| Q921G7 | Etfdh | -0.827103528 | 2.919643939 |
| Q922J3 | Clip1 | -0.827250981 | 4.148398213 |
| Q921N6 | Ddx27 | -0.827432384 | 2.942286148 |
| O88487 | Dync1i2 | -0.828479936 | 3.07202973 |
| P70297 | Stam | -0.829268492 | 3.492310325 |
| O55236 | Rngtt | -0.829476 | 2.749557574 |
| P52825 | Cpt2 | -0.829981298 | 2.415249225 |
| Q91WG2 | Rabep2 | -0.830178488 | 3.465704771 |
| Q7TMK6 | Hook2 | -0.830343161 | 3.076257705 |
| Q8R1A4 | Dock7 | -0.83035842 | 3.61447752 |
| Q8BYR2 | Lats1 | -0.831134999 | 3.146070578 |
| P17742 | Ppia | -0.831460866 | 3.975394642 |
| O88271 | Cfdp1 | -0.83323931 | 2.941602987 |
| P10853;P10854;P70696;Q64475;Q64478;Q64524;Q64525;Q6ZWY9;Q8CGP0;Q8CGP1;Q8CGP2;Q9D2U9 | H2bc | -0.833345018 | 2.25745568 |
| Q9JLZ3 | Auh | -0.834740232 | 2.210510866 |
| Q3UE37 | Ube2z | -0.835376191 | 4.177210817 |
| Q9EQC5 | Scyl1 | -0.836059262 | 4.759418207 |
| Q7M6Y3 | Picalm | -0.836860115 | 3.740372647 |
| Q61029 | Tmpo | -0.836900642 | 2.085005107 |
| P60764;P60766;P63001;P84096;Q05144;Q8R527;Q9ER71 | Rac3;Cdc42;Rhog;Rac2;Rhoj;Rhoq | -0.837706836 | 2.476407463 |
| Q3THK3 | Gtf2f1 | -0.83800799 | 4.055120904 |
| Q9CYN9 | Atp6ap2 | -0.838494653 | 2.140345641 |
| Q8VDM4 | Psmd2 | -0.838534658 | 4.739970804 |
| P27046 | Man2a1 | -0.839445536 | 2.394687499 |
| Q8VI33;Q6NZA9 | Taf9 | -0.839598846 | 2.419086564 |
| Q8BLN5 | Lss | -0.839846721 | 3.120665462 |
| P18654;Q7TPS0 | Rps6ka6 | -0.839877202 | 2.48796244 |
| Q8VIJ8 | Nprl3 | -0.840361937 | 2.07655851 |
| O88643;Q8CIN4 | Pak1 | -0.841110575 | 3.279559674 |
| P62075 | Timm13 | -0.843100948 | 2.54384277 |
| Q99MN1 | Kars1 | -0.844063655 | 3.491492343 |
| Q9WVF7 | Pole | -0.844403095 | 3.414093521 |
| Q8BML9 | Qars1 | -0.844749711 | 2.820044585 |
| Q80XP8 | Fam76b | -0.845339333 | 4.046822367 |
| Q8CIG3 | Kdm1b | -0.845642705 | 2.014685359 |
| Q3U821 | Wdr75 | -0.846093087 | 3.117119536 |
| Q61164 | Ctcf | -0.847437066 | 2.066582482 |
| P41230;Q62240 | Kdm5 | -0.851371122 | 2.998966427 |
| Q9EPU4 | Cpsf1 | -0.851446817 | 4.294483376 |
| Q922E4 | Pcyt2 | -0.85161171 | 2.503905676 |
| P20918 | Plg | -0.852351297 | 2.663279666 |
| Q9QZ05 | Eif2ak4 | -0.852527046 | 2.713213212 |
| Q9Z0R4;Q9Z0R6 | Itsn1 | -0.852561932 | 2.361934814 |
| Q9CZX0 | Elp3 | -0.852707189 | 2.52759988 |
| Q9D706 | Rpap3 | -0.853138719 | 4.004922876 |
| Q9WVM1 | Racgap1 | -0.854817733 | 3.120665462 |
| Q8R0J7 | Vps37b | -0.85498385 | 3.262187402 |
| E9Q634;P70248 | Myo1e | -0.855157087 | 3.310489487 |
| Q6P4S8 | Ints1 | -0.856150649 | 4.176298047 |
| Q9Z321 | Top3b | -0.856230921 | 2.764848976 |
| P97493 | Txn2 | -0.857466915 | 3.352807586 |
| O88587 | Comt | -0.857510948 | 2.638527816 |
| Q9D1C1 | Ube2c | -0.858391663 | 3.107744365 |
| Q61771 | Kif3b | -0.859685015 | 2.93620878 |
| Q99KP3 | Cryl1 | -0.860810566 | 2.208967499 |
| P14873;Q9QYR6 | Map1 | -0.861926831 | 2.316124792 |
| Q9DBH5 | Lman2 | -0.862657844 | 2.973811502 |
| Q60972 | Rbbp4 | -0.863789281 | 3.996432538 |
| Q3UHB1 | Nt5dc3 | -0.863929316 | 3.395742822 |
| P62983 | Rps27a | -0.865288921 | 3.906997904 |
| P08003 | Pdia4 | -0.865671454 | 2.667004994 |
| Q8K268 | Abcf3 | -0.866615807 | 3.298830889 |
| P20664 | Prim1 | -0.86730283 | 3.91854463 |
| P17182;P17183;P21550 | Eno | -0.867735202 | 2.56522747 |
| Q924H2 | Med15 | -0.86779497 | 3.491492343 |
| P62077 | Timm8b | -0.868258747 | 2.465611893 |
| B2RY56 | Rbm25 | -0.868488248 | 3.691110732 |
| Q62083 | Pick1 | -0.869456383 | 2.712832241 |
| P18653;P18654;Q9WUT3 | Rps6ka | -0.869989471 | 2.761835778 |
| Q9WUI1 | Mapk11 | -0.870282544 | 2.036596281 |
| Q61543 | Glg1 | -0.87187071 | 2.54005747 |
| Q8K2Q0 | Commd9 | -0.872138669 | 3.551836988 |
| Q9JHW4 | Eefsec | -0.872302642 | 4.034126349 |
| A0A7H0DNA6 | OPG134 | -0.872447118 | 2.627370457 |
| Q9CR20 | Ier3ip1 | -0.873952785 | 3.107744365 |
| Q8CFE3;Q6PGA0 | Rcor1 | -0.874608013 | 3.565850416 |
| Q8QZX2 | Haus3 | -0.874664702 | 2.471495032 |
| Q3URD3 | Slmap | -0.87484033 | 3.018235962 |
| Q3TNH5 | Arb2a | -0.875173331 | 2.225697931 |
| Q8K2A7 | Ints10 | -0.875332575 | 3.921222483 |
| Q8R317;Q99NB8;Q9QZM0 | Ubqln | -0.875782501 | 4.214115181 |
| Q70FJ1 | Akap9 | -0.875922633 | 2.167369187 |
| Q9DBD5 | Pelp1 | -0.876478458 | 3.474029774 |
| O35435 | Dhodh | -0.876811526 | 2.130016841 |
| Q8CFI7 | Polr2b | -0.8776163 | 3.465838366 |
| P12382;P47857 | Pfk | -0.880522423 | 3.750865948 |
| Q7TNP2 | Ppp2r1b | -0.88053064 | 4.661074824 |
| Q60605 | Myl6 | -0.880861717 | 2.976004592 |
| O35218 | Cpsf2 | -0.881013548 | 4.942701901 |
| Q8CBW3 | Abi1 | -0.881736167 | 3.776732633 |
| Q9DCF9 | Ssr3 | -0.88178491 | 2.587836068 |
| Q9CU65 | Zmym2 | -0.884058913 | 3.582925274 |
| O88447;O88448;Q9DBS5 | Klc | -0.884076991 | 3.782514136 |
| Q8CIM8 | Ints4 | -0.88418524 | 3.022890723 |
| P53994 | Rab2a | -0.884492122 | 5.259521696 |
| Q8VHR5 | Gatad2b | -0.885029408 | 3.740372647 |
| O54941 | Smarce1 | -0.885115126 | 4.620967825 |
| Q8BG51 | Rhot1 | -0.885853598 | 2.389256597 |
| Q8CFI0 | Nedd4l | -0.885984694 | 3.410674765 |
| Q91XD2;Q99JW4 | Lims2 | -0.888166169 | 3.051913576 |
| Q99K23 | Ufsp2 | -0.889226484 | 3.107159946 |
| P63328 | Ppp3ca | -0.890191853 | 3.462472292 |
| P18760 | Cfl1 | -0.892690318 | 5.16574648 |
| A2AN08 | Ubr4 | -0.893932519 | 4.040348819 |
| F6ZDS4 | Tpr | -0.894721722 | 3.795200494 |
| Q8BMJ2 | Lars1 | -0.897068424 | 2.942640609 |
| Q9CY27 | Tecr | -0.897307061 | 3.073341531 |
| Q8CI08 | Slain2 | -0.897364519 | 3.483467608 |
| Q80XI4 | Pip4k2b | -0.897748548 | 3.838138631 |
| Q69ZQ2 | Isy1 | -0.898101259 | 3.492921131 |
| Q9DBJ1 | Pgam1 | -0.898126619 | 3.732476255 |
| Q8VD62 | Bles03 | -0.898338204 | 4.396566281 |
| P26350 | Ptma | -0.900618904 | 3.583487661 |
| Q9Z1E3 | Nfkbia | -0.90123747 | 2.417596365 |
| Q3V3R1 | Mthfd1l | -0.9030192 | 2.998966427 |
| A8Y5H7 | Sec14l1 | -0.904874084 | 4.491544855 |
| Q9ER00 | Stx12 | -0.904978702 | 3.283276416 |
| Q7TMB8 | Cyfip1 | -0.906427869 | 3.860201804 |
| O54988 | Slk | -0.90732757 | 3.218848999 |
| Q8C5W3 | Tbcel | -0.9073456 | 2.951497041 |
| B2RSH2;P08752;Q9DC51 | Gnai | -0.908203571 | 2.686730664 |
| O70279 | Ess2 | -0.908720175 | 3.79698936 |
| Q9ESL4 | Map3k20 | -0.908872744 | 3.428161622 |
| Q9CXU0 | Med10 | -0.909797301 | 3.209034268 |
| Q9WTK2 | Cdyl | -0.911401483 | 4.138032906 |
| Q8K2H2 | Otud6b | -0.9116761 | 3.537822397 |
| P49718 | Mcm5 | -0.911776292 | 4.838901889 |
| Q6NZF1 | Zc3h11a | -0.913216417 | 2.066170347 |
| P60670 | Nploc4 | -0.913566952 | 4.751391646 |
| Q8R4Z4 | Etv3 | -0.913891138 | 2.156468178 |
| P35123;Q8R5H1 | Usp4;Usp15 | -0.914747986 | 3.166866133 |
| Q9D0M1 | Prpsap1 | -0.916944402 | 3.513971357 |
| Q9WTL7 | Lypla2 | -0.918321063 | 3.389169626 |
| Q8VD75 | Hip1 | -0.919393547 | 4.316975038 |
| O35075 | Vps26c | -0.919983687 | 3.208703866 |
| Q61214;Q9Z188 | Dyrk1 | -0.920098167 | 2.291801445 |
| Q8K411 | Pitrm1 | -0.920496181 | 3.023175123 |
| Q8R2E9 | Ero1b | -0.920744258 | 2.556660777 |
| Q6WKZ8 | Ubr2 | -0.922095651 | 3.507268386 |
| Q8BH24 | Tm9sf4 | -0.922579137 | 2.283239569 |
| Q8C547 | Heatr5b | -0.922869992 | 3.456520582 |
| Q61083 | Map3k2 | -0.922939693 | 2.744669344 |
| Q9D787 | Ppil2 | -0.923476384 | 3.685082013 |
| Q9CQ39 | Med21 | -0.923923756 | 4.209368071 |
| Q64727 | Vcl | -0.924436344 | 4.990275012 |
| P39053;P39054;Q8BZ98 | Dnm | -0.92495768 | 3.057579546 |
| Q9JM13 | Rabgef1 | -0.925341288 | 3.847788825 |
| P51863 | Atp6v0d1 | -0.925404377 | 2.280705928 |
| Q8C9S4 | Ccdc186 | -0.925876758 | 3.035555702 |
| Q5EG47 | Prkaa1 | -0.926241856 | 3.851773076 |
| P40124 | Cap1 | -0.92637017 | 4.467536582 |
| P31230 | Aimp1 | -0.926865228 | 3.536492351 |
| O88845 | Akap10 | -0.928618209 | 3.317161291 |
| Q8VDJ3 | Hdlbp | -0.928724976 | 3.825683356 |
| Q921M4 | Golga2 | -0.929597778 | 4.233861669 |
| Q8C7H1 | Mmaa | -0.930387352 | 2.151442724 |
| Q99020 | Hnrnpab | -0.931646321 | 4.317142144 |
| P97760 | Polr2c | -0.931980895 | 4.039848135 |
| Q8BTY8 | Scfd2 | -0.932009369 | 2.537111473 |
| B2RQC6 | Cad | -0.932516587 | 4.930033579 |
| Q6PAC3 | Dcaf13 | -0.932715835 | 2.912074778 |
| Q9CR86 | Carhsp1 | -0.932746904 | 2.297972291 |
| Q6PAV2 | Herc4 | -0.933262915 | 3.669598294 |
| Q8BHL5 | Elmo2 | -0.933383182 | 3.43363605 |
| Q9DBC3 | Cmtr1 | -0.933835597 | 2.468111942 |
| A2AR02 | Ppig | -0.934953023 | 2.832640344 |
| P62192 | Psmc1 | -0.935685327 | 3.476733355 |
| P58462 | Foxp1 | -0.937733709 | 4.054889529 |
| P97363 | Sptlc2 | -0.938499339 | 2.700595061 |
| Q8CDG3 | Vcpip1 | -0.938499382 | 4.149489159 |
| P17879;P63017;Q61696 | Hspa8 | -0.938507512 | 4.057291445 |
| O88685 | Psmc3 | -0.938924366 | 4.698569796 |
| Q99LB2 | Dhrs4 | -0.939438371 | 2.188064293 |
| Q6P9J9 | Ano6 | -0.939652874 | 2.147166548 |
| Q9WUL7 | Arl3 | -0.939825666 | 4.074900775 |
| Q9D1K2 | Atp6v1f | -0.940390345 | 2.763700414 |
| Q8BUY9 | Pggt1b | -0.941040011 | 3.028149287 |
| Q99LN9 | Dohh | -0.941148093 | 3.15226893 |
| Q9D0J4 | Arl2 | -0.942294805 | 3.61204835 |
| Q9ES28 | Arhgef7 | -0.943280215 | 4.065637563 |
| Q9CX11 | Utp23 | -0.943432525 | 3.22459565 |
| Q62245 | Sos1 | -0.943555061 | 2.956924558 |
| Q8R4R6 | Nup35 | -0.944738087 | 3.807294588 |
| Q6R891 | Ppp1r9b | -0.946099476 | 3.566802794 |
| Q5ND34 | Wdr81 | -0.946987701 | 2.198872899 |
| P28652;Q923T9 | Camk2 | -0.947423486 | 3.259143974 |
| P33609 | Pola1 | -0.94770952 | 3.646428308 |
| Q3UYH7;Q99MK8 | Grk | -0.947916043 | 3.875722057 |
| Q8CH02 | Sugp1 | -0.948028379 | 2.60045472 |
| Q9CXV1 | Sdhd | -0.948105541 | 2.71423185 |
| P63005 | Pafah1b1 | -0.949345381 | 4.747116276 |
| Q5BL07 | Pex1 | -0.951683421 | 2.936432032 |
| P27612 | Plaa | -0.953312092 | 4.210092636 |
| Q9Z2Y8 | Plpbp | -0.953705337 | 2.951371057 |
| Q921X9 | Pdia5 | -0.956736026 | 3.130158439 |
| Q99K70 | Rragc | -0.960104955 | 3.645437068 |
| G5E8V9 | Arfip1 | -0.960170862 | 3.167342424 |
| Q8BH43 | Wasf2 | -0.960201988 | 3.078870242 |
| P47791 | Gsr | -0.960802862 | 3.854068366 |
| Q9R1X4 | Timeless | -0.961949536 | 3.281615871 |
| Q8V4S4;A0A7H0DNF0 | OPG188 | -0.963524302 | 4.34012424 |
| Q8BZQ7 | Anapc2 | -0.965458861 | 3.059461219 |
| Q99JB8 | Pacsin3 | -0.965823347 | 3.82489943 |
| Q6ZQL4 | Wdr43 | -0.966327879 | 3.918011144 |
| P17710 | Hk1 | -0.967409914 | 3.69515796 |
| P0DW87 | Zftraf1 | -0.970155408 | 2.203140004 |
| Q5FWK3 | Arhgap1 | -0.970648201 | 3.953530856 |
| Q9EQP2;Q9QXY6 | Ehd3 | -0.970920622 | 3.208047213 |
| P51175 | Ppox | -0.973200055 | 3.414384507 |
| P09411 | Pgk1 | -0.974284584 | 4.525833123 |
| Q8BQR4 | Kansl2 | -0.974386944 | 2.179274564 |
| P55264 | Adk | -0.974529479 | 3.403725202 |
| Q9D6K8 | Fundc2 | -0.974887122 | 3.296058319 |
| P27661;Q64522 | H2ac21 | -0.975350417 | 2.764468928 |
| Q99K74 | Med24 | -0.97551995 | 4.405087119 |
| Q9R0N0 | Galk1 | -0.976210502 | 2.394408491 |
| Q91W36 | Usp3 | -0.978570011 | 2.665818316 |
| Q920Q8 | Ivns1abp | -0.98033965 | 3.18581557 |
| Q925J9 | Med1 | -0.980430672 | 4.931096713 |
| Q64707;Q62377 | Zrsr2 | -0.980675288 | 2.307484049 |
| Q6PDG5 | Smarcc2 | -0.980878718 | 4.287483935 |
| Q9JMH6 | Vamp8 | -0.984505093 | 4.586411796 |
| O70404 | Psme1 | -0.985536286 | 3.285859828 |
| P97371 | Prim2 | -0.98570815 | 3.563717775 |
| P33610 | Rbm5 | -0.985960617 | 2.91100363 |
| Q91YE7 | Pgd | -0.98597333 | 4.02463907 |
| Q9DCD0 | Srsf5 | -0.985985978 | 4.091657855 |
| O35326 | Dmxl1 | -0.9860328 | 5.00222714 |
| Q6PNC0 | Acap2 | -0.986377879 | 3.35690514 |
| Q6ZQK5 | Os9 | -0.986970431 | 3.881349607 |
| Q8K2C7 | Msantd2 | -0.987065219 | 2.360695759 |
| Q6NZR2 | Xab2 | -0.987391409 | 3.869017621 |
| Q9DCD2 | Slu7 | -0.988318509 | 4.282565377 |
| Q8BHJ9 | Necap1 | -0.98883771 | 3.998251092 |
| Q9CR95 | Pmm2 | -0.989519578 | 2.896896438 |
| Q9Z2M7 | Dek | -0.990559613 | 5.8532722 |
| Q7TNV0 | Trio | -0.99135329 | 3.838138631 |
| Q0KL02 | OPG019 | -0.992297148 | 3.88014652 |
| A0A7H0DMZ6 | Ganab | -0.992820368 | 2.990059465 |
| Q8BHN3 | Capzb | -0.992954505 | 3.820461367 |
| P47757 | Smarcc2 | -0.992990388 | 4.95256605 |
| Q8C147;Q8R1A4 | Dock8 | -0.99356947 | 2.291801445 |
| O55131 | Septin7 | -0.994290824 | 4.485296484 |
| P63094;Q6R0H7;Q8CGK7 | Gnas | -0.995230546 | 3.109972546 |
| Q7TMF2 | Eri1 | -0.995930132 | 3.540578473 |
| Q8VCX5 | Micu1 | -0.996558985 | 2.291921696 |
| Q8BVF2 | Pdcl3 | -0.997290426 | 3.710448679 |
| A2BH40;E9Q4N7 | Arid1 | -0.997928427 | 2.252783524 |
| Q80T69 | Rsbn1 | -0.998917646 | 3.296058319 |
| Q922R8 | Pdia6 | -0.998957825 | 2.8594699 |
| Q8BH64;Q9WVK4 | Ehd | -1.001757577 | 2.629592962 |
| Q8C1B7;Q8CHH9 | Septin8 | -1.001916977 | 3.558554591 |
| Q8CE33 | Klhl11 | -1.003320491 | 3.350378663 |
| O35385 | Ppef2 | -1.004142234 | 3.012231411 |
| Q920R0 | Als2 | -1.00429428 | 2.595398235 |
| Q6P3E7 | Hdac10 | -1.006460841 | 2.329961069 |
| Q8BFR4 | Gns | -1.008004138 | 2.927609989 |
| P26039 | Tln1 | -1.008093625 | 4.482015345 |
| Q64318 | Zeb1 | -1.008328592 | 2.102597499 |
| O88738 | Birc6 | -1.008522477 | 4.194733453 |
| Q9Z2D1 | Mtmr2 | -1.010610738 | 2.785058269 |
| Q9DC23 | Dnajc10 | -1.011256203 | 2.75744129 |
| Q64436;Q6PIC6;Q6PIE5;Q8VDN2;Q9WV27;Q9Z1W8 | Atp | -1.012171122 | 2.710782255 |
| P11157 | Rrm2 | -1.013113234 | 4.453233937 |
| Q07139 | Ect2 | -1.013411418 | 2.020505064 |
| A0A7H0DNF0 | OPG188 | -1.01389084 | 2.971127483 |
| E9Q7M2 | Tsc22d2 | -1.017028076 | 3.022694938 |
| Q9CS00 | Cactin | -1.017322249 | 2.386764957 |
| Q8R060 | Zwilch | -1.018155857 | 3.238150255 |
| O54836 | Zmat3 | -1.019031675 | 3.942094589 |
| P42669 | Pura | -1.019613445 | 4.491544855 |
| Q922Q4 | Pycr2 | -1.020011962 | 2.739175354 |
| O88696 | Clpp | -1.020205032 | 2.958300441 |
| Q9EQ80 | Nif3l1 | -1.020722791 | 2.998966427 |
| P48453;P63328 | Ppp3cb | -1.020822832 | 3.107159946 |
| Q9QXT0 | Cnpy2 | -1.021583205 | 3.061743443 |
| Q505B7 | Zbtb8os | -1.021841359 | 2.375134642 |
| O89050 | Mkln1 | -1.022301761 | 4.34012424 |
| Q61112 | Sdf4 | -1.023054708 | 3.506732362 |
| Q8BKG3 | Ptk7 | -1.023117195 | 2.631684292 |
| Q8VDR9 | Dock6 | -1.023810597 | 4.542362531 |
| Q6ZQ03 | Fnbp4 | -1.024392106 | 3.707708155 |
| Q8K114 | Ints9 | -1.024699566 | 4.48142587 |
| Q9D0F3 | Lman1 | -1.025284756 | 2.591741228 |
| Q91ZR2 | Snx18 | -1.025511902 | 2.337379753 |
| Q9ERR7 | Selenof | -1.025616473 | 3.793487755 |
| O70305 | Atxn2 | -1.02605918 | 4.020475075 |
| Q9WV68 | Decr2 | -1.02849841 | 4.010371626 |
| Q9R0H0 | Acox1 | -1.029826683 | 4.048271194 |
| P28741;P33174 | Kif | -1.029922567 | 3.054854173 |
| P17918 | Pcna | -1.031008183 | 3.118766539 |
| Q9QZ88 | Vps29 | -1.031069685 | 3.921222483 |
| Q3UE31 |  | -1.031170964 | 2.976004592 |
| Q8R361 | Rab11fip5 | -1.034290747 | 3.212402718 |
| P40142 | Tkt | -1.035568416 | 5.259521696 |
| Q05CL8 | Larp7 | -1.037853126 | 4.395080478 |
| Q8BFS9 | Asnsd1 | -1.038606874 | 2.198574489 |
| P13439 | Umps | -1.039435607 | 4.333580364 |
| Q9CVB6 | Arpc2 | -1.039672529 | 4.735413384 |
| O35598 | Adam10 | -1.040065551 | 2.486237814 |
| P09041;P09411 | Pgk2 | -1.04131213 | 4.163018778 |
| P58871 | Tnks1bp1 | -1.04149737 | 3.324398395 |
| P97310 | Mcm2 | -1.042645199 | 4.192134858 |
| Q922M3 | Kctd10 | -1.042980757 | 2.731634489 |
| Q9EPC1 | Parva | -1.043817913 | 4.698560058 |
| Q61595 | Ktn1 | -1.045047218 | 2.52310294 |
| Q99K01 | Pdxdc1 | -1.045161072 | 3.902032311 |
| Q8VDY9 | Caap1 | -1.045650104 | 2.527433329 |
| Q8BU31 | Rap2c | -1.046226523 | 2.572496784 |
| P56399 | Usp5 | -1.046232122 | 4.622358186 |
| Q8BVD5 | Mpp7 | -1.047293224 | 4.301764195 |
| P35601 | Rfc1 | -1.047480007 | 2.311609463 |
| Q925H1 | Trps1 | -1.047749343 | 3.908383398 |
| Q04735;Q8K0D0;Q04899 | Cdk | -1.048152175 | 2.878993608 |
| P70441 | Nherf1 | -1.04945224 | 3.668381916 |
| Q8BGQ1 | Vipas39 | -1.049459023 | 2.399433359 |
| Q8VEE4 | Rpa1 | -1.049601753 | 4.378689432 |
| Q2HXL6 | Edem3 | -1.050019017 | 3.626653919 |
| Q8BK67 | Rcc2 | -1.050488901 | 5.013335147 |
| Q9CZ44 | Nsfl1c | -1.050720103 | 4.144758147 |
| O70480 | Vamp4 | -1.051160958 | 2.589329059 |
| Q9CR00 | Psmd9 | -1.052181427 | 3.673337794 |
| Q924T7 | Rnf31 | -1.053382664 | 3.611079173 |
| Q9DC48 | Cdc40 | -1.053502276 | 4.82406096 |
| Q8BGS0 | Mak16 | -1.05355208 | 3.265290242 |
| Q9D7P6 | Iscu | -1.054158042 | 3.50834372 |
| P53996 | Cnbp | -1.054364704 | 3.73124246 |
| Q61584 | Fxr1 | -1.055329363 | 5.540673463 |
| Q91VH6 | Memo1 | -1.056947363 | 3.599509382 |
| Q08297 | Rad51 | -1.05868356 | 3.678652402 |
| Q9CY18 | Snx7 | -1.059775087 | 3.402353655 |
| Q91XL3 | Uxs1 | -1.060558149 | 3.546298103 |
| P13864 | Dnmt1 | -1.060863795 | 2.932248799 |
| Q9EQH2 | Erap1 | -1.061299846 | 2.778162488 |
| P11798;Q6PHZ2 | Camk2 | -1.061422767 | 2.979211095 |
| P97302 | Bach1 | -1.062421728 | 2.183527933 |
| P08775 | Polr2a | -1.064630425 | 4.235930649 |
| Q3UL36 | Arglu1 | -1.065465378 | 3.714630109 |
| Q8VCB2 | Med25 | -1.066160644 | 2.312321731 |
| Q3TDD9 | Ppp1r21 | -1.066812561 | 2.604983142 |
| Q8BG15 | Ctdspl2 | -1.066898693 | 3.159670248 |
| Q9JJ28 | Flii | -1.068171388 | 4.166303027 |
| Q05BC3 | Eml1 | -1.068603699 | 2.949155314 |
| Q9JJ78 | Pbk | -1.070616226 | 2.435377326 |
| Q3THG9 | Aarsd1 | -1.071254287 | 3.269862196 |
| Q9DBR7 | Ppp1r12a | -1.071747597 | 4.82406096 |
| Q6P5C5 | Smug1 | -1.072538727 | 2.787105891 |
| Q62189 | Snrpa | -1.072597253 | 2.085290323 |
| P25206 | Mcm3 | -1.073102847 | 5.255932293 |
| O55135 | Eif6 | -1.073467402 | 4.014143757 |
| Q8BWM0 | Ptges2 | -1.073820704 | 3.121987081 |
| Q69ZA1 | Cdk13 | -1.074211095 | 3.227923018 |
| Q6PGF3 | Med16 | -1.074785808 | 4.010371626 |
| P11440;P97377;Q80YP0 | Cdk1;Cdk2;Cdk3 | -1.075524131 | 3.097525833 |
| Q9R0L6 | Pcm1 | -1.075710093 | 3.149826054 |
| A0A7H0DNE5 | OPG178 | -1.078687768 | 4.004933353 |
| Q8BQZ5 | Cpsf4 | -1.080840574 | 3.723501819 |
| Q9CPY7 | Lap3 | -1.081270377 | 4.489390806 |
| Q8BVU0 | Lrch3 | -1.082756032 | 4.110268549 |
| Q3UZ39 | Lrrfip1 | -1.084423608 | 3.50834372 |
| Q9CXK8 | Nip7 | -1.085073484 | 3.717290297 |
| Q9D902 | Gtf2e2 | -1.085174788 | 3.150797652 |
| Q8R034 | Anapc13 | -1.085495841 | 3.654931923 |
| A2A6A1 | Gpatch8 | -1.086940188 | 3.310489487 |
| Q6P5D8 | Smchd1 | -1.088067069 | 2.279285107 |
| Q80YQ2 | Med23 | -1.088728592 | 4.036377352 |
| Q5SQX6 | Cyfip2 | -1.088847751 | 3.822122385 |
| Q8CCK0;Q9QZQ8 | Macroh2a1 | -1.089526675 | 2.789013575 |
| Q6P1F6;Q8BG02 | Ppp2r2c | -1.089901201 | 3.216991014 |
| A2ABV5 | Med14 | -1.090612394 | 4.811630857 |
| P70333 | Hnrnph2 | -1.0914216 | 3.270143596 |
| Q9DBR7;Q8BG95 | Ppp1r12b | -1.092520248 | 3.991501482 |
| Q9JHP7 | Poglut2 | -1.093436756 | 3.75194599 |
| Q9CQ28 | Dph6 | -1.09530822 | 2.07655851 |
| Q9WVA2 | Timm8a1 | -1.096147907 | 2.601229478 |
| Q9Z2X1 | Hnrnpf | -1.09810692 | 4.53718775 |
| Q9ER39 | Tor1a | -1.098820037 | 3.548344642 |
| P21126 | Ubl4a | -1.101031791 | 3.634704616 |
| P12815 | Pdcd6 | -1.101199743 | 4.735413384 |
| P26443 | Glud1 | -1.102398398 | 3.768184482 |
| Q6A0A9 | FAM120A | -1.102642224 | 3.931235291 |
| O09172 | Gclm | -1.102948804 | 2.611069571 |
| Q3UQN2 | Fcho2 | -1.104535769 | 3.492310325 |
| Q9CX97 | Wdr55 | -1.105890439 | 2.63168358 |
| Q9JM76 | Arpc3 | -1.106248906 | 3.676032902 |
| P31750;Q60823;Q9WUA6 | Akt | -1.107029218 | 2.335939917 |
| A0A7H0DNG0 | OPG200 | -1.108838172 | 3.76084779 |
| Q62388 | Atm | -1.111190461 | 2.649316474 |
| A1A5B6 | Tbc1d25 | -1.112040998 | 2.378662126 |
| Q9QUR6 | Prep | -1.113098823 | 4.645336194 |
| Q7TT50 | Cdc42bpb | -1.113711853 | 4.451296921 |
| P38647 | Hspa9 | -1.11581821 | 3.164373362 |
| Q99JY9 | Actr3 | -1.116483296 | 5.394910166 |
| Q91XY4 | Pcdhga4 | -1.118329553 | 3.022890723 |
| Q921F2 | Tardbp | -1.11840181 | 4.730785014 |
| O70378 | Emc8 | -1.11845918 | 4.360655937 |
| P58742 | Aaas | -1.121003836 | 3.712046777 |
| P47739;P47740 | Aldh3a | -1.123203028 | 3.590757142 |
| A2AGH6 | Med12 | -1.124193323 | 2.181404133 |
| Q9R0A0 | Pex14 | -1.125202633 | 4.699228201 |
| P21278;P21279 | Gna | -1.125922439 | 3.08449826 |
| P58774 | Tpm2 | -1.12637785 | 4.821066334 |
| Q9D3E6 | Stag1 | -1.126869444 | 3.576342833 |
| Q8BNY6 | Ncs1 | -1.127138062 | 4.163018778 |
| Q91YS8 | Camk1 | -1.128705108 | 3.445781674 |
| Q8R5K2 | Usp33 | -1.128990997 | 2.705998176 |
| Q7TT45;Q99K70 | Rragd | -1.13082912 | 2.442189343 |
| Q5SRX1 | Tom1l2 | -1.133428322 | 3.819388623 |
| Q3UHJ0 | Aak1 | -1.133961195 | 3.743428691 |
| Q9JII6 | Akr1a1 | -1.134182175 | 3.592576569 |
| Q64433 | Hspe1 | -1.134434328 | 2.433292602 |
| P97311 | Mcm6 | -1.134773682 | 4.903962268 |
| A0A7H0DNE2 | OPG174 | -1.136838748 | 3.908383398 |
| P10711 | Tcea1 | -1.137557286 | 4.838901889 |
| Q62188 | Dpysl3 | -1.138099648 | 4.6160186 |
| P08249 | Mdh2 | -1.138374027 | 2.744669344 |
| Q6IE82 | Jade3 | -1.138451853 | 4.043650602 |
| Q8CHU3 | Epn2 | -1.138656058 | 3.553812509 |
| Q61687 | Atrx | -1.139199663 | 4.415156422 |
| Q4FZF3 | Ddx49 | -1.140105595 | 3.870007399 |
| Q9JKX4 | Aatf | -1.140640287 | 3.870828082 |
| Q8BHG1 | Nrdc | -1.145254797 | 4.237911274 |
| Q9CPU4 | Mgst3 | -1.145542091 | 2.17895369 |
| Q80U58 | Pum2 | -1.145693532 | 4.022680847 |
| Q8R3B1 | Plcd1 | -1.147465295 | 2.787105891 |
| P47934 | Crat | -1.148511921 | 2.678065491 |
| Q91YR1 | Twf1 | -1.150918043 | 3.223247692 |
| P53702 | Hccs | -1.153492078 | 3.356771671 |
| Q60605;Q8CI43 | Myl6b | -1.15390216 | 4.614974883 |
| Q91WA1 | Tipin | -1.15519534 | 3.714132432 |
| Q9Z0H3 | Smarcb1 | -1.158218884 | 4.56955956 |
| Q8CEC0 | Nup88 | -1.15846855 | 3.718969764 |
| Q3U182 | Crtc2 | -1.158821138 | 2.128760068 |
| Q60848 | Hells | -1.159251487 | 4.194733453 |
| Q9D0M3 | Cyc1 | -1.159445372 | 2.953113879 |
| Q80U72;Q80VQ1 | Scrib;Lrrc1 | -1.160271853 | 2.956924558 |
| Q9QUJ7 | Acsl4 | -1.160448826 | 5.144782839 |
| E9Q5G3 | Kif23 | -1.162812643 | 4.338535926 |
| O35945;P24549;P47738;Q62148;Q9JHW9;Q9CZS1 | Aldh | -1.163154254 | 3.135956355 |
| P55258;P61027;P61028;P62821;Q9D1G1;Q9DD03;Q8K386 | Rab | -1.163403608 | 2.159951152 |
| Q8K2Q5 | Chchd7 | -1.164193595 | 3.285437583 |
| Q68FL4 | Ahcyl2 | -1.167180922 | 3.879153496 |
| Q8CHE4 | Phlpp1 | -1.167184543 | 2.100246727 |
| Q8K4L3 | Svil | -1.168711988 | 3.488954172 |
| Q8K120 | Nfatc4 | -1.169047001 | 2.980338351 |
| Q64674 | Srm | -1.169139627 | 3.795659315 |
| P21981 | Tgm2 | -1.171261502 | 3.176905021 |
| Q9QXK2 | Rad18 | -1.1717772 | 3.060725734 |
| P10126 | Eef1a1 | -1.172217143 | 4.183060809 |
| Q91ZA3 | Pcca | -1.173487949 | 3.682247118 |
| Q6NVE9 | Pptc7 | -1.174282338 | 2.610165011 |
| Q6ZQ73 | Cand2 | -1.178555684 | 3.187541123 |
| Q8R5J9 | Arl6ip5 | -1.17889191 | 3.110889723 |
| Q6A009 | Ltn1 | -1.180722851 | 3.265318659 |
| O88456 | Capns1 | -1.184749942 | 4.498261722 |
| Q9R0E1 | Plod3 | -1.185329519 | 3.454972469 |
| Q9CYL5 | Glipr2 | -1.188372687 | 3.474029774 |
| A0A7H0DNG2 | OPG204 | -1.188586968 | 4.396566281 |
| O35648 | Cetn3 | -1.188682747 | 3.310489487 |
| Q3U7R1 | Esyt1 | -1.189359386 | 3.474029774 |
| Q8BH95 | Echs1 | -1.190419015 | 2.802438178 |
| D3Z7P3 | Gls | -1.190711214 | 3.564255074 |
| Q9D8C6 | Med11 | -1.191751116 | 3.268331914 |
| O88783 | F5 | -1.191959845 | 2.226067807 |
| Q9ES46 | Parvb | -1.192403563 | 3.052272778 |
| Q9D1D4 | Tmed10 | -1.192551858 | 3.030340882 |
| A0A7H0DMZ9 | OPG023 | -1.194580481 | 4.036776454 |
| P97477 | Aurka | -1.196196281 | 3.420118455 |
| Q9QX47 | Son | -1.200987718 | 2.850099865 |
| Q8BG05 | Hnrnpa3 | -1.202388591 | 4.634910335 |
| A2RSY1 | Kansl3 | -1.202766864 | 2.431281459 |
| Q3THE2 | Myl12b | -1.203172976 | 4.740770147 |
| Q9D8S4 | Rexo2 | -1.20531 | 3.825301669 |
| Q8C4B4 | Unc119b | -1.207262082 | 4.033884815 |
| P04184 | Tk1 | -1.207770076 | 3.463663909 |
| Q6ZWY3 | Rps27l | -1.208287039 | 4.399169536 |
| Q6Y5D8 | Arhgap10 | -1.20951483 | 2.580218546 |
| P05480 | Src | -1.210707333 | 2.224849622 |
| Q922Q4;Q922W5 | Pycr1 | -1.211501427 | 3.372426719 |
| P98083 | Shc1 | -1.214072885 | 3.556306265 |
| Q61586 | Gpam | -1.214692762 | 3.239118909 |
| Q62108;Q811D0;Q91XM9 | Dlg | -1.217646605 | 3.449232155 |
| Q3UZ39;Q91WK0 | Lrrfip2 | -1.218198625 | 3.972076717 |
| P12023 | App | -1.220075165 | 3.260593185 |
| Q8BKX1 | Baiap2 | -1.222106584 | 5.016249569 |
| Q922R5 | Ppp4r3b | -1.222849605 | 3.441677862 |
| B2RUR8;Q8R554 | Otud7 | -1.224193204 | 2.787105891 |
| Q9Z2R6 | Unc119 | -1.22644865 | 3.396035889 |
| Q8BVE8 | Nsd2 | -1.2267593 | 3.429313103 |
| E9PYK3 | Parp4 | -1.228338008 | 2.252520944 |
| Q8BH59 | Slc25a12 | -1.230028675 | 2.309225271 |
| Q03265 | Atp5f1a | -1.231544148 | 3.867663368 |
| Q8K2B3 | Sdha | -1.233040248 | 3.118011836 |
| Q6NZM9 | Hdac4 | -1.233213702 | 3.145816414 |
| Q9WUZ9 | Entpd5 | -1.233759618 | 2.85637436 |
| Q61881 | Mcm7 | -1.234479563 | 5.394910166 |
| Q06180 | Ptpn2 | -1.235126513 | 4.170902486 |
| Q8R059 | Gale | -1.242589451 | 4.357250411 |
| Q8BP47 | Nars1 | -1.246734949 | 4.233861669 |
| Q8R0K9 | E2f4 | -1.246979947 | 2.016983972 |
| Q9D824 | Fip1l1 | -1.247254467 | 4.328589847 |
| Q99KV1 | Dnajb11 | -1.247725593 | 3.454451089 |
| Q9JK81 | Myg1 | -1.250536523 | 4.267487385 |
| Q91ZU6 | Dst | -1.252623434 | 4.158338398 |
| Q8VI56 | Lrp4 | -1.25412249 | 2.388120626 |
| Q9CQ60 | Pgls | -1.255257213 | 3.561808738 |
| P08113;P11499 | Hsp90 | -1.255683918 | 2.561173679 |
| Q9CR57 | Rpl14 | -1.256146929 | 2.685709591 |
| P35831 | Ptpn12 | -1.256481778 | 5.902972035 |
| P12787 | Cox5a | -1.258876964 | 2.943391789 |
| Q8R0G9 | Nup133 | -1.259333069 | 3.564158238 |
| Q06185 | Atp5me | -1.259394386 | 2.278628982 |
| Q922H2 | Pdk3 | -1.261127292 | 3.600182778 |
| Q8BK08 | Tmem11 | -1.261671382 | 3.43861713 |
| P56480 | Atp5f1b | -1.262294586 | 3.847788825 |
| Q6ZQF0 | Topbp1 | -1.263068637 | 2.851835136 |
| A0A7H0DNG5 | OPG209 | -1.263214331 | 2.740317771 |
| Q91XU0 | Wrnip1 | -1.263533298 | 4.682174872 |
| Q3TC93 | Hs1bp3 | -1.264940508 | 3.850285735 |
| P09405 | Ncl | -1.266465609 | 5.20140885 |
| Q3TJD7 | Pdlim7 | -1.267793674 | 4.622443967 |
| Q5PRF0 | Heatr5a | -1.268011589 | 2.563413437 |
| Q6VN19 | Ranbp10 | -1.273325253 | 3.714630109 |
| A0A7H0DN11 | OPG038 | -1.273519269 | 3.420298723 |
| Q922J3;Q9Z0H8 | Clip2 | -1.274440645 | 4.112320344 |
| Q9QY30 | Abcb11 | -1.275955046 | 2.69058362 |
| Q9ES00 | Ube4b | -1.276299534 | 3.858693928 |
| O35207;Q9CPY4 | Cdk2ap | -1.278488148 | 4.473898187 |
| Q9D662 | Sec23b | -1.278726628 | 4.491544855 |
| Q9WU42 | Ncor2 | -1.278983001 | 3.536492351 |
| Q9CW46 | Raver1 | -1.279736115 | 4.640508668 |
| P30415 | Nktr | -1.281641758 | 4.511962317 |
| Q9R0Q6;Q9WV32 | Arpc1 | -1.282118521 | 4.176774851 |
| Q9CQX8 | Kgd4 | -1.284567115 | 3.799207139 |
| Q8CFE6 | Slc38a2 | -1.285564244 | 2.669462907 |
| Q8R180 | Ero1a | -1.286015101 | 3.715037591 |
| Q91VW3 | Sh3bgrl3 | -1.286559871 | 4.137061257 |
| Q9D7J4 | Cox20 | -1.286878588 | 2.063096743 |
| P97452 | Bop1 | -1.289252341 | 2.505273218 |
| Q6EDY6 | Carmil1 | -1.289707158 | 2.367747037 |
| Q9Z0J0 | Npc2 | -1.291905951 | 3.393597188 |
| P62962 | Pfn1 | -1.292244088 | 4.720963244 |
| P56212 | Arpp19 | -1.294763233 | 2.033227749 |
| G5E829 | Atp2b1 | -1.294934346 | 2.737113842 |
| Q08509 | Eps8 | -1.295890251 | 3.08449826 |
| Q9JHL1 | Nherf2 | -1.296379938 | 3.996449161 |
| Q60775 | Elf1 | -1.297633509 | 2.829872463 |
| P59108 | Cpne2 | -1.298602029 | 3.2412017 |
| Q9CZR8 | Tsfm | -1.298756871 | 2.85112369 |
| Q9CZW4;Q9QUJ7 | Acsl3 | -1.300317088 | 2.927960093 |
| Q8C0D4 | Arhgap12 | -1.30054963 | 2.600090358 |
| Q8CJ53 | Trip10 | -1.300628918 | 3.763954438 |
| Q9JMH9 | Myo18a | -1.300768821 | 3.13889578 |
| Q8VC03 | Eml3 | -1.301105606 | 2.061155948 |
| Q9DBS9 | Osbpl3 | -1.304172639 | 3.799634409 |
| Q91W59 | Rbms1 | -1.306728832 | 4.033884815 |
| P70445 | Eif4ebp2 | -1.307619795 | 3.287162148 |
| Q69ZF7 | Cnnm4 | -1.307755347 | 2.63789268 |
| Q9QWT9 | Kifc1 | -1.311581894 | 2.583402325 |
| Q80UY2 | Kcmf1 | -1.312865129 | 3.105319513 |
| O88531 | Ppt1 | -1.31559748 | 4.366797517 |
| Q9ET26 | Rnf114 | -1.316368797 | 3.27483205 |
| Q8C3Y4 | Kntc1 | -1.316968229 | 3.414093521 |
| P26040;P26041;P26043 | Ezr;Msn;Rdx | -1.319114742 | 4.864588747 |
| Q6P9P6 | Kif11 | -1.321226632 | 4.993940729 |
| O35379 | Abcc1 | -1.322887032 | 3.459200339 |
| Q8R2Y0 | Abhd6 | -1.322907136 | 2.258942628 |
| F8VPZ9 | Bicra | -1.323505709 | 2.996586809 |
| Q6P9L6 | Kif15 | -1.324216643 | 4.82406096 |
| Q99MR8 | Mccc1 | -1.324628512 | 2.030750736 |
| P24270 | Cat | -1.324995395 | 3.460817988 |
| Q8CIE4 | Parp10 | -1.326746354 | 2.724453764 |
| Q3UMG5 | Lrch2 | -1.32785565 | 3.999644043 |
| P47713 | Pla2g4a | -1.328028404 | 4.747116276 |
| P12849;Q9DBC7 | Prkar1 | -1.328298985 | 3.105050321 |
| Q91ZV0 | Mia2 | -1.331796929 | 2.15328049 |
| Q80VJ2 | Sra1 | -1.33201311 | 4.119034618 |
| Q8BIW9 | Chtf18 | -1.332316469 | 5.540673463 |
| Q00547 | Hmmr | -1.332526366 | 4.210829287 |
| Q9JLQ2 | Git2 | -1.332656875 | 3.654931923 |
| A0A7H0DN07 | OPG034 | -1.333080351 | 2.567944834 |
| Q9CY50 | Ssr1 | -1.336682702 | 3.477107387 |
| Q9DAP7 | Asf1b | -1.336963738 | 3.795200494 |
| P47226 | Tes | -1.336965212 | 4.183060809 |
| Q91VX2 | Ubap2 | -1.33738685 | 4.401394469 |
| Q99JY9;Q641P0 | Actr3b | -1.338504303 | 2.929979595 |
| Q61103 | Dpf2 | -1.339469938 | 3.808540344 |
| Q8BVI4 | Qdpr | -1.342161118 | 4.151296843 |
| P06537 | Nr3c1 | -1.34345145 | 2.494680987 |
| Q8CIF4 | Btd | -1.34487511 | 2.297621212 |
| Q8VBT9 | Aspscr1 | -1.345147444 | 3.978733079 |
| O08756 | Hsd17b10 | -1.345244349 | 2.509637238 |
| Q01405;Q9D662 | Sec23a | -1.345969049 | 3.585967196 |
| Q9D6R2 | Idh3a | -1.346765985 | 3.846460924 |
| Q8R1F1 | Niban2 | -1.347823148 | 4.141212591 |
| Q00PI9 | Hnrnpul2 | -1.348264536 | 4.170902486 |
| P43024 | Cox6a1 | -1.349862429 | 2.716595031 |
| O08674;P31266 | Rbpj | -1.351994211 | 4.333580364 |
| O08529 | Capn2 | -1.356213656 | 4.167373342 |
| P70349 | Hint1 | -1.356727136 | 3.019322326 |
| Q8BI72 | Cdkn2aip | -1.357733566 | 2.130024851 |
| Q07231 | Zscan21 | -1.362729325 | 2.290631496 |
| Q9WUU8 | Tnip1 | -1.363494292 | 2.157969571 |
| Q8R1V4 | Tmed4 | -1.368272797 | 3.491492343 |
| P51569 | Gla | -1.374663605 | 3.536492351 |
| Q8R3C6 | Rbm19 | -1.378232978 | 4.430085309 |
| Q99N85 | Mrps18a | -1.378604676 | 3.276734323 |
| Q5FWH2 | Unkl | -1.378630875 | 2.655774066 |
| Q3UFY0 | Rrp36 | -1.38096669 | 2.063456046 |
| Q02819 | Nucb1 | -1.381662014 | 3.804907615 |
| Q8JZN5 | Acad9 | -1.383061935 | 3.187541123 |
| P16951 | Atf2 | -1.3836788 | 2.671731443 |
| Q8R4U7 | Luzp1 | -1.384939998 | 4.183060809 |
| Q8C1S0 | Med19 | -1.385992188 | 4.03401086 |
| O35350 | Capn1 | -1.386914609 | 2.674711265 |
| Q08879 | Fbln1 | -1.388148174 | 2.29522114 |
| Q8BXR9;Q9DBS9 | Osbpl6 | -1.389851706 | 3.839765569 |
| Q8K1J6 | Trnt1 | -1.390118353 | 3.737109404 |
| Q61879;Q8VDD5 | Myh9 | -1.392061167 | 4.716211672 |
| Q01853 | Vcp | -1.397641085 | 5.038716555 |
| Q8C8U0 | Ppfibp1 | -1.402884759 | 4.620967825 |
| Q80UG5 | Septin9 | -1.402959252 | 4.430085309 |
| P54071 | Idh2 | -1.403812537 | 3.537822397 |
| Q99KB8 | Hagh | -1.405005669 | 3.943655471 |
| Q9EPQ7 | Stard5 | -1.405145832 | 2.280444367 |
| P52480 | Pkm | -1.406793254 | 4.931096713 |
| Q9CY66 | Gar1 | -1.40733359 | 4.233211323 |
| A0A7H0DN24 | OPG051 | -1.409059567 | 2.716595031 |
| P19536 | Cox5b | -1.410147797 | 3.25453213 |
| Q9EP71 | Rai14 | -1.415373753 | 4.685587194 |
| Q3TQB2 | Foxred1 | -1.418758043 | 2.119950165 |
| Q61576 | Fkbp10 | -1.419228089 | 2.096917004 |
| Q7TSG2 | Ctdp1 | -1.423925227 | 4.386402606 |
| P0DTM9 | OPG001 | -1.424530231 | 3.227923018 |
| Q9ES97 | Rtn3 | -1.424916067 | 3.920788675 |
| Q8BJU9 | Mtrf1l | -1.428615274 | 4.198784682 |
| O08600 | Endog | -1.429307732 | 2.477558067 |
| Q91W90 | Txndc5 | -1.429399825 | 3.065379107 |
| Q9D6K7 | Ttc33 | -1.43029283 | 2.564078804 |
| Q9Z1S0 | Bub1b | -1.430848868 | 3.940061563 |
| Q9WVL2 | Stat2 | -1.433697917 | 2.697386722 |
| Q02384;Q62245 | Sos2 | -1.436082505 | 4.501393133 |
| Q80X90;Q8BTM8;Q8VHX6 | Fln | -1.436541638 | 4.233861669 |
| Q9DCK4 | Znf414 | -1.438337591 | 2.171275073 |
| Q8K371 | Amotl2 | -1.438644426 | 3.557118976 |
| Q9CQT5 | Pomp | -1.443365331 | 2.95817039 |
| Q9CZT6 | Cmss1 | -1.443451117 | 3.915553082 |
| P36536;Q9CQC9 | Sar1 | -1.446690693 | 2.998966427 |
| Q69Z38 | Peak1 | -1.448814699 | 3.474029774 |
| O54984 | Get3 | -1.449598386 | 5.260100747 |
| Q9CZJ2 | Hspa12b | -1.452382832 | 2.899968608 |
| P35700 | Prdx1 | -1.458000589 | 3.114095961 |
| Q9D1N9 | Mrpl21 | -1.458840357 | 2.952520495 |
| P21107 | Tpm3 | -1.458929044 | 4.796539026 |
| Q80TA9 | Epg5 | -1.459925157 | 2.28856965 |
| P63321;Q9JIW9 | Rala;Ralb | -1.464777634 | 2.433368881 |
| Q8BP92 | Rcn2 | -1.47308186 | 3.429313103 |
| Q9JHC9 | Elf2 | -1.473083234 | 2.490139335 |
| P05622 | Pdgfrb | -1.474394003 | 2.475396316 |
| P08103 | Hck | -1.47510735 | 2.580200308 |
| P61022 | Chp1 | -1.475256624 | 3.305263183 |
| Q9CZA6 | Nde1 | -1.476674855 | 3.332729152 |
| Q07076 | Anxa7 | -1.478445974 | 4.213349624 |
| Q9QZ23 | Nfu1 | -1.479634495 | 3.473758 |
| Q9WTX8 | Mad1l1 | -1.480286452 | 4.326842125 |
| Q9DCX2 | Atp5pd | -1.481257717 | 3.159670248 |
| P06801 | Me1 | -1.482449622 | 4.329005527 |
| O54967 | Tnk2 | -1.484091116 | 4.244015654 |
| P35285;Q921E2 | Rab22a;Rab31 | -1.48441316 | 3.514271611 |
| P23798 | Pcgf2 | -1.486883645 | 2.745999975 |
| Q9D0S9 | Hint2 | -1.487389776 | 3.260371005 |
| Q9ER81 | Tor1aip2 | -1.489428899 | 3.804907615 |
| Q9D7J9 | Echdc3 | -1.492082411 | 2.047217074 |
| P37913 | Lig1 | -1.492695595 | 4.850280042 |
| Q3UE17 | Mex3d | -1.493468971 | 3.528855408 |
| Q8CHY6 | Gatad2a | -1.494007693 | 3.751295849 |
| Q9JII7 | Zfand2a | -1.51000305 | 2.796107751 |
| P48024;Q9CXU9 | Eif1 | -1.510994812 | 4.740770147 |
| Q99JY0 | Hadhb | -1.5120702 | 3.272696139 |
| Q8VCF0 | Mavs | -1.517180603 | 2.561480169 |
| Q91X51 | Gorasp1 | -1.51777616 | 3.120665462 |
| Q9WVA2;Q4FZG7 | Timm8a2 | -1.520464986 | 2.180335101 |
| Q9CX86 | Hnrnpa0 | -1.520820241 | 3.701535962 |
| P18242 | Ctsd | -1.521454782 | 3.750750582 |
| A0A7H0DN28 | OPG055 | -1.521981112 | 3.615897537 |
| Q8VE22 | Mrps23 | -1.522277863 | 3.351231497 |
| Q9CRA4 | Msmo1 | -1.523464787 | 2.750239545 |
| A0A7H0DN02;Q8V566 | OPG027 | -1.526518414 | 3.710710676 |
| Q91V12 | Acot7 | -1.528061252 | 4.716211672 |
| P70404 | Idh3g | -1.528499566 | 3.828930153 |
| Q9DAU1 | Cnpy3 | -1.529248224 | 3.941049284 |
| A0A7H0DMZ8 | OPG022 | -1.529869531 | 2.85401555 |
| Q9D0J8 | Ptms | -1.530387378 | 2.899713863 |
| Q8CH72 | Trim32 | -1.531188753 | 3.730343253 |
| Q7TS68 | Nsun6 | -1.532466259 | 2.342642171 |
| Q9CXI5 | Manf | -1.533967837 | 3.43363605 |
| P55194 | Sh3bp1 | -1.538805322 | 4.215216437 |
| P47915 | Rpl29 | -1.545708821 | 4.982715035 |
| Q8BMD8 | Slc25a24 | -1.5471134 | 3.342725054 |
| Q8JZU0 | Nudt13 | -1.547403449 | 2.773140659 |
| Q9WVB0 | Rbpms | -1.550205337 | 3.919867659 |
| Q9JM14 | Nt5c | -1.550471732 | 3.931993449 |
| Q80VP0 | Tecpr1 | -1.551694807 | 2.442488669 |
| Q3UMR5 | Mcu | -1.552240029 | 2.713110507 |
| Q80TS3 | Adgrl3 | -1.557737625 | 2.995913541 |
| Q8VHI3 | Pofut2 | -1.558632637 | 3.159798797 |
| Q8K1L5 | Ppp1r11 | -1.558818433 | 3.483503256 |
| E9PVB5 | Ttc17 | -1.562412061 | 3.240849705 |
| P35293 | Rab18 | -1.564027941 | 4.003533613 |
| Q60722;Q61286 | Tcf4;Tcf12 | -1.565159268 | 2.847742274 |
| P21107;P58771 | Tpm1 | -1.566636149 | 2.857863665 |
| Q69ZF3 | Gba2 | -1.567082617 | 2.082131595 |
| Q8K009 | Aldh1l2 | -1.572311233 | 3.013648734 |
| Q9JM05 | Pias4 | -1.5736679 | 2.046789648 |
| Q3TLS3 | Gdpgp1 | -1.574748306 | 2.37899756 |
| Q6P5E4 | Uggt1 | -1.575900215 | 3.849663192 |
| Q8BHC9 | Fut11 | -1.576808402 | 3.296058319 |
| Q6ZPF4;A2APV2 | Fmnl3;Fmnl2 | -1.577406161 | 3.637487765 |
| Q80W47 | Wipi2 | -1.578698457 | 4.002145646 |
| Q64514 | Tpp2 | -1.581395155 | 4.82406096 |
| Q9R207 | Nbn | -1.581411327 | 2.066034165 |
| Q9CY34 | Ube2f | -1.582602112 | 4.344982086 |
| P45878 | Fkbp2 | -1.582978326 | 3.682247118 |
| Q99LD8 | Ddah2 | -1.583292242 | 3.897420779 |
| Q6PDH0 | Phldb1 | -1.583360159 | 2.742336452 |
| Q71FD7 | Fblim1 | -1.583367059 | 4.131715835 |
| P05202 | Got2 | -1.584783264 | 2.368675173 |
| Q9JI39 | Abcb10 | -1.586559659 | 2.740292714 |
| Q8R0A7 | Kiaa0513 | -1.587666764 | 2.372017595 |
| B2RXS4 | Plxnb2 | -1.588012381 | 3.401577307 |
| Q3V3Q7 | Pacs2 | -1.590571165 | 2.777946029 |
| Q8BGC4 | Ptgr3 | -1.590720064 | 3.913657467 |
| Q9CQ92 | Fis1 | -1.591554026 | 4.639353751 |
| Q91YW3 | Dnajc3 | -1.59207528 | 3.561899877 |
| Q9Z0U1 | Tjp2 | -1.593729535 | 4.933474852 |
| Q497V5 | Srbd1 | -1.598739453 | 2.119950165 |
| P58137 | Acot8 | -1.60633226 | 2.677112056 |
| Q3TCN2 | Plbd2 | -1.607645948 | 2.623895764 |
| Q99104 | Myo5a | -1.607743997 | 4.560638611 |
| Q9DAR7 | Dcps | -1.608169531 | 3.962567042 |
| P08113 | Cep55 | -1.609059553 | 3.761064182 |
| Q8BT07 | Mrtfa | -1.609945907 | 3.432727018 |
| Q9JI10 | Zswim8 | -1.612863639 | 4.538681767 |
| Q8K4J6 | Lonp1 | -1.615121183 | 2.305570849 |
| Q3UHH1 | Nisch | -1.624845171 | 4.040348819 |
| Q8CGK3 | Phldb1 | -1.635533001 | 3.660537061 |
| Q80TM9 | Fblim1 | -1.63675778 | 3.637718366 |
| P19324 | Serpinh1 | -1.637996932 | 4.388408184 |
| Q60949 | Tbc1d1 | -1.638513943 | 3.967728853 |
| O88673 | Dgka | -1.641145245 | 2.38888588 |
| Q9DBS2 | Tprg1l | -1.642835405 | 2.32171417 |
| P57776 | Eef1d | -1.643057627 | 5.014660892 |
| O88878 | Zfand5 | -1.644078571 | 2.90448372 |
| P63028 | Tpt1 | -1.646626454 | 3.687393625 |
| A0A7H0DN02 | OPG027 | -1.648789494 | 3.953530856 |
| P52623 | Uck1 | -1.651130401 | 4.34012424 |
| Q6P5E8 | Dgkq | -1.65168508 | 2.37635589 |
| Q8BTY2 | Slc4a7 | -1.651928328 | 3.022113643 |
| P11276 | Fn1 | -1.652411846 | 2.625735216 |
| P59438 | Hps5 | -1.662264375 | 2.12890443 |
| Q3THE2;Q9CQ19 | Myl9 | -1.662402913 | 4.436054321 |
| P23591 | Gfus | -1.664866669 | 3.773164927 |
| Q922J9 | Far1 | -1.666339592 | 4.214069853 |
| O08664 | Bcl7c | -1.670283977 | 3.540578473 |
| Q0HA38 | Ttc21b | -1.673730415 | 2.121110042 |
| Q6AXC6 | Ddx11 | -1.674010732 | 3.990139814 |
| Q9WV30 | Nfat5 | -1.67401423 | 3.860367231 |
| Q8BWQ4 | Cmtr2 | -1.675266077 | 3.103505744 |
| Q91VU0 | Fam3c | -1.681340449 | 2.121902189 |
| Q91W67 | Ubl7 | -1.682433198 | 3.91854463 |
| Q8BYB9 | Poglut1 | -1.683070808 | 3.847795768 |
| Q8VD12 | Znf385a | -1.685228917 | 2.24977912 |
| Q9CWV6 | Prkrip1 | -1.685328779 | 4.316975038 |
| Q9EPV8 | Ubl5 | -1.688831158 | 4.607199918 |
| P18406 | Ccn1 | -1.690818429 | 2.023774141 |
| O55023 | Impa1 | -1.693546281 | 2.388429345 |
| Q9WUM3 | Coro1b | -1.698149053 | 5.144782839 |
| Q99LS1 | Mmadhc | -1.699972854 | 2.319228589 |
| Q9JKR6 | Hyou1 | -1.706491903 | 3.64642223 |
| P08030 | Aprt | -1.706743845 | 2.35595023 |
| Q9JHJ0 | Tmod3 | -1.707879964 | 4.430085309 |
| P62500;E9Q7M2 | Tsc22d1 | -1.712575334 | 4.477974072 |
| P14733;P21619;P48678 | Lmn | -1.713555018 | 4.430085309 |
| Q8CI03 | Flywch1 | -1.713847822 | 2.037608817 |
| Q9D8W7 | Ociad2 | -1.718554676 | 2.942192398 |
| Q6NXH3 | Gpbp1 | -1.719762526 | 2.103919628 |
| O35887 | Calu | -1.722069736 | 4.430085309 |
| Q921F4 | Hnrnpll | -1.725315263 | 6.445062849 |
| P52633 | Stat6 | -1.727101359 | 2.791812936 |
| Q91VH2 | Snx9 | -1.727609654 | 4.620967825 |
| P54754;P54761 | Ephb | -1.729800325 | 2.433537738 |
| O08599 | Stxbp1 | -1.731538898 | 4.601001249 |
| P31324 | Prkar2b | -1.732622248 | 4.043650602 |
| Q80XC3 | Usp6nl | -1.737548011 | 2.539359339 |
| P19096 | Fasn | -1.740956916 | 2.230390972 |
| Q9ER41 | Tor1b | -1.742908528 | 3.673337794 |
| Q07243 | Mtf1 | -1.747678867 | 2.690540748 |
| Q9CZN7 | Shmt2 | -1.748884199 | 3.79698936 |
| Q9Z1B3 | Plcb1 | -1.751033325 | 2.258972241 |
| Q00612 | G6pdx | -1.751106112 | 4.940371175 |
| P35441 | Thbs1 | -1.751684539 | 3.010661047 |
| Q9R1Z8 | Sorbs3 | -1.757679839 | 2.056346244 |
| Q6DFW0 | C9orf72 | -1.758126381 | 3.714630109 |
| Q91WT8 | Rbm47 | -1.7582503 | 3.616701249 |
| P70290 | Mpp1 | -1.759134992 | 4.040348819 |
| Q3TTL0 |  | -1.760413071 | 2.187316456 |
| P48036 | Anxa5 | -1.760453752 | 4.804351636 |
| Q9CQ65 | Mtap | -1.76058835 | 4.66362192 |
| Q60875 | Arhgef2 | -1.764092595 | 4.6344451 |
| Q67FY2 | Bcl9l | -1.764524019 | 2.88076794 |
| Q9DBG9 | Tax1bp3 | -1.766017508 | 4.126660061 |
| Q6P2L7 | Golm2 | -1.766365165 | 2.835400987 |
| Q8C006 | Dpysl2 | -1.770577942 | 2.311422065 |
| Q9Z1Q9 | Trim35 | -1.773940103 | 4.433360725 |
| Q6PB44 | Vars1 | -1.780757682 | 4.662886389 |
| P30681 | Ptpn23 | -1.783061109 | 4.417793068 |
| Q9D7B6 | Acad8 | -1.783720632 | 3.864244963 |
| Q99KI0 | Aco2 | -1.784878387 | 3.931235291 |
| Q99N20 | Brms1 | -1.785677733 | 3.207601799 |
| P0DTN0 | OPG002 | -1.78718555 | 2.658945589 |
| Q9D4D4 | Tktl2 | -1.792629767 | 3.179676886 |
| Q7TSJ6 | Lats2 | -1.801763664 | 2.687520922 |
| Q3TMX7 | Qsox2 | -1.804278754 | 2.221087166 |
| Q9D168 | Ints12 | -1.805635906 | 2.197531921 |
| Q9DAK9 | Phpt1 | -1.805941694 | 2.348746971 |
| Q8K214 | Scmh1 | -1.809525829 | 2.107542168 |
| Q8BH97 | Rcn3 | -1.809984605 | 3.272901787 |
| Q9D968 | Hcfc2 | -1.811341983 | 3.420298723 |
| B9EJ80 | Pdzd8 | -1.811635502 | 2.052128931 |
| P99029 | Prdx5 | -1.812864379 | 2.83985443 |
| Q9WUQ2 | Preb | -1.816521091 | 2.737389095 |
| P51480 | Cdkn2a | -1.818571319 | 4.040348819 |
| Q8R2Y8 | Ptrh2 | -1.822568761 | 3.619124736 |
| P21271;Q99104 | Myo5b | -1.823236234 | 4.477974072 |
| O35680 | Mrps12 | -1.827165779 | 3.260762867 |
| O35295 | Purb | -1.827312519 | 5.144782839 |
| Q9Z2Q6 | Septin5 | -1.830066532 | 4.03522281 |
| Q3TJM4 | Cenpt | -1.841374935 | 2.316124792 |
| Q6P9R4 | Arhgef18 | -1.841834073 | 2.220336027 |
| Q60692 | Psmb6 | -1.843613432 | 2.589163827 |
| Q61792 | Lasp1 | -1.845513636 | 4.531460466 |
| Q91W45 | Paip2b | -1.848938095 | 2.146239518 |
| P70227 | Itpr3 | -1.851018696 | 2.621827645 |
| D3YXK2 | Safb | -1.852698433 | 3.964407007 |
| Q921G6 | Lrch4 | -1.858918113 | 2.332943176 |
| Q3U829 | Ap5z1 | -1.868749218 | 2.491956998 |
| Q04863 | Relb | -1.868800818 | 3.042434817 |
| Q9CRC3 |  | -1.873065585 | 2.445760501 |
| P52480;P53657 | Pklr | -1.873349469 | 2.308741664 |
| Q8BG73 | Sh3bgrl2 | -1.874312869 | 3.847788825 |
| Q8R1K4 | Phykpl | -1.886495085 | 3.199311583 |
| O70475 | Ugdh | -1.887224948 | 5.280410943 |
| Q9CXP8 | Gng10 | -1.889192911 | 3.086587137 |
| Q9Z247 | Fkbp9 | -1.892230218 | 4.034126349 |
| Q03145 | Epha2 | -1.892844494 | 4.56391268 |
| P09103 | P4hb | -1.893887599 | 4.982715035 |
| Q91Z22 | Tmem123 | -1.896875172 | 3.04550426 |
| O55137;Q6Q2Z6;Q8BWN8;Q9QYR7;Q9QYR9;Q32Q92 | Acot | -1.909767837 | 2.122652319 |
| Q80U62 | Rubcn | -1.912842004 | 2.305470698 |
| P02469 | Lamb1 | -1.91463851 | 4.138808266 |
| Q68EF0 | Rab3ip | -1.916054374 | 2.787020721 |
| Q9D8S3 | Arfgap3 | -1.92027504 | 2.113033469 |
| Q9CQB5 | Cisd2 | -1.923110895 | 4.003533613 |
| A0A7H0DN16 | OPG043 | -1.927053144 | 5.52172333 |
| Q3TLI0 | Trappc10 | -1.933985915 | 2.306784033 |
| Q60597 | Ogdh | -1.935652156 | 4.144709934 |
| Q9CZP0 | Ufsp1 | -1.936328929 | 2.25745568 |
| Q8BSY0 | Asph | -1.93980127 | 3.683439887 |
| Q5HZK1 | Cep44 | -1.942249286 | 2.465852631 |
| Q3UFB2 | Znhit6 | -1.944919403 | 4.520467189 |
| P14211 | Calr | -1.945680708 | 3.853392838 |
| P29533 | Vcam1 | -1.948733313 | 2.250472338 |
| Q8R5F7 | Ifih1 | -1.949570718 | 4.121971107 |
| Q61084 | Map3k3 | -1.952027478 | 2.019082336 |
| Q99P72 | Rtn4 | -1.953618725 | 4.735413384 |
| P10518 | Alad | -1.954762482 | 3.879153496 |
| Q0VGY8 | Tanc1 | -1.958385102 | 2.247938357 |
| Q07113 | Igf2r | -1.958583223 | 3.941049284 |
| Q9CXE7 | Tmed5 | -1.959250646 | 3.154710621 |
| P09925 | Surf1 | -1.960681655 | 2.592475755 |
| O09130 | Nfatc2ip | -1.96242942 | 2.166679645 |
| P42337 | Pik3ca | -1.969976267 | 3.213324564 |
| Q91YH5 | Atl3 | -1.97402695 | 4.519134463 |
| Q9D975 | Srxn1 | -1.974942071 | 2.206305787 |
| O54926 | Siva1 | -1.977503738 | 3.032666154 |
| Q8N7N5 | Dcaf8 | -1.989980321 | 3.681192526 |
| Q921U8 | Smtn | -1.991517819 | 2.82119914 |
| O35427 | Crcp | -1.991616023 | 3.751479382 |
| Q8CEG5 | Ccdc28b | -1.997374511 | 2.138946278 |
| Q9D8B7 | Jam3 | -1.99772678 | 2.33322968 |
| Q9D967 | Mdp1 | -2.000774464 | 2.955893465 |
| O88342 | Wdr1 | -2.003170824 | 4.868785445 |
| Q920A5 | Scpep1 | -2.007246867 | 3.594493902 |
| Q3U2U7 | Mettl17 | -2.008973711 | 2.798876143 |
| Q8BHW2 | Oscp1 | -2.01063535 | 2.434031632 |
| Q3TGF2 | Fam107b | -2.011810051 | 4.186350718 |
| Q3TZZ7 | Esyt2 | -2.016898474 | 3.264210042 |
| P61750;P84084 | Arf4;Arf5 | -2.016998427 | 3.16726783 |
| Q8BNV1 | Trmt2a | -2.019986191 | 4.189519893 |
| Q9D8L5 | Ccdc91 | -2.021129883 | 2.128874285 |
| Q4VAA7 | Snx33 | -2.024594418 | 2.039567746 |
| A6H8H2 | Dennd4c | -2.03451164 | 3.729335111 |
| P50431 | Shmt1 | -2.036860323 | 4.698560058 |
| Q9ESY9 | Ifi30 | -2.038355967 | 3.705168426 |
| Q8VC70 | Rbms2 | -2.048719557 | 4.017127025 |
| Q9R059 | Fhl3 | -2.052222284 | 5.335553116 |
| Q9CQF0 | Mrpl11 | -2.055657434 | 4.142594819 |
| Q923B0 | Ggact | -2.057036269 | 2.25448424 |
| Q9DB42 | Znf593 | -2.05972399 | 4.746441307 |
| Q8BJM3 | R3hcc1l | -2.062457949 | 2.69058362 |
| Q3U4G3 | Xxylt1 | -2.064360915 | 2.839000993 |
| Q8BW96;Q91YS8 | Camk1d | -2.064682864 | 4.205916826 |
| P54116 | Stom | -2.071158806 | 3.483299217 |
| Q8R5C5 | Actr1b | -2.071256551 | 5.902972035 |
| P49817;P51637 | Cav1;Cav3 | -2.073714167 | 3.676032902 |
| Q8BP22 | Cibar1 | -2.075252747 | 3.429109377 |
| Q8VI36 | Pxn | -2.077796472 | 5.569291115 |
| Q8R4E9 | Cdt1 | -2.080318853 | 2.096291035 |
| Q60716 | P4ha2 | -2.085035605 | 4.56391268 |
| Q99LB0 | Dnttip1 | -2.094159522 | 3.06003506 |
| Q8R151 | Znfx1 | -2.097217064 | 2.85112369 |
| Q8CDN6 | Txnl1 | -2.101925608 | 4.736528849 |
| P17095 | Hmga1 | -2.103090992 | 4.709390902 |
| O88196 | Ttc3 | -2.107551283 | 2.417850939 |
| Q9R1Z7 | Pts | -2.108033927 | 4.433360725 |
| Q8BU33 | Ilvbl | -2.109453909 | 2.127985254 |
| Q8VE73 | Cul7 | -2.110192481 | 2.422825572 |
| Q91WS0 | Cisd1 | -2.115290012 | 2.921757372 |
| Q9JJ00 | Plscr1 | -2.115542155 | 2.406398606 |
| Q9CYW4 | Hdhd3 | -2.12514661 | 2.003527723 |
| Q80V26 | Bpnt2 | -2.130305494 | 2.660195254 |
| Q80X85 | Mrps7 | -2.13715537 | 3.451750727 |
| Q60651 | Klra4 | -2.138135372 | 3.545543697 |
| Q9JJ11 | Tacc3 | -2.140714956 | 2.28497868 |
| Q08775 | Runx2 | -2.148091347 | 2.032318049 |
| Q5FWI3 | Cemip2 | -2.149084897 | 2.744669344 |
| Q8R0F3 | Sumf1 | -2.164740501 | 4.453233937 |
| Q6NXM2 | Rcbtb1 | -2.164921014 | 2.917523923 |
| Q9ER38 | Tor3a | -2.167048517 | 2.036596281 |
| Q8CDM1 | Atad2 | -2.170714681 | 4.443807798 |
| P14901 | Hmox1 | -2.173292682 | 3.853392838 |
| Q05915 | Gch1 | -2.174582501 | 2.019082336 |
| P17751 | Tpi1 | -2.17934083 | 5.394910166 |
| Q8C522 | Endod1 | -2.179591841 | 2.005006902 |
| A2AI08 | Tprn | -2.180817284 | 2.373115675 |
| Q80WE4 | Kif20b | -2.185577641 | 2.647584791 |
| Q9R0P3 | Esd | -2.197739391 | 4.885133704 |
| Q64521 | Gpd2 | -2.199625536 | 4.046706882 |
| Q62465 | Vat1 | -2.200780411 | 4.443807798 |
| Q6P7W0 | Senp6 | -2.201749074 | 2.117734187 |
| P17809 | Slc2a1 | -2.203915622 | 3.849663192 |
| Q91W61 | Fbxl15 | -2.204490173 | 2.360545213 |
| Q9D2G5 | Synj2 | -2.204893463 | 2.283308347 |
| Q8K212 | Pacs1 | -2.21296763 | 3.737223033 |
| Q60715 | P4ha1 | -2.219835065 | 4.838901889 |
| Q99PL6 | Ubxn6 | -2.219940802 | 3.492310325 |
| P20152 | Vim | -2.222836676 | 4.886177338 |
| P47754 | Capza2 | -2.226697279 | 5.56236107 |
| Q8BSI6 | R3hcc1 | -2.228236752 | 2.904237223 |
| Q3TEL6;Q9D074 | Rnf157;Mgm1 | -2.230500797 | 2.23485496 |
| O70433 | Fhl2 | -2.230677667 | 6.245955496 |
| Q8BXA1 | Golim4 | -2.23101946 | 2.233941912 |
| Q9D9K3 | Aven | -2.235169868 | 2.118699255 |
| P00375 | Dhfr | -2.235361406 | 4.931096713 |
| P14069 | S100a6 | -2.240304898 | 2.745512984 |
| Q64430 | Atp7a | -2.240554086 | 3.454972469 |
| Q6A058 | Armcx2 | -2.241838998 | 2.512356424 |
| Q99M54 | Cdca3 | -2.246749652 | 2.388783479 |
| Q3TUA9 | Pomk | -2.251315511 | 2.500658341 |
| Q0GNC1 | Inf2 | -2.251635909 | 3.725671756 |
| Q9EPK6 | Sil1 | -2.254662404 | 2.720055381 |
| E9Q0S6;Q5SSZ5;Q8CGB6 | Tns | -2.259269679 | 4.838901889 |
| Q8BRM2 | Gorab | -2.261513409 | 2.444469485 |
| Q8VC34 | Rpap2 | -2.262697907 | 2.51779497 |
| Q8C854 | Myef2 | -2.263089788 | 4.933474852 |
| Q91WC9 | Daglb | -2.265717023 | 2.455520959 |
| Q91ZS8 | Adarb1 | -2.285092264 | 4.333580364 |
| Q8K3P5 | Cnot6 | -2.286890707 | 2.028316903 |
| Q8BZS9 | Dhx32 | -2.29533189 | 2.019206414 |
| Q8CHK3 | Mboat7 | -2.295894825 | 2.118699255 |
| Q9Z2Q2 | Knop1 | -2.299600758 | 3.864244963 |
| Q9CZB0 | Sdhc | -2.305287541 | 3.045327451 |
| Q8K004 | Spata2 | -2.305375492 | 2.097732248 |
| Q9CR16 | Ppid | -2.308981049 | 4.982715035 |
| P10922 | H1-0 | -2.311600861 | 2.442708803 |
| P20152;P31001 | Des | -2.325056059 | 4.664616004 |
| Q5SUE8 | Ankrd40 | -2.325171511 | 2.066262684 |
| Q9D404 | Oxsm | -2.334032768 | 2.273165605 |
| Q9JK92 | Hspb8 | -2.335229673 | 3.992177544 |
| Q8VEB4 | Pla2g15 | -2.34138177 | 3.340876839 |
| Q9JIA7 | Sphk2 | -2.341535798 | 2.434031632 |
| P15331;P20152 | Prph | -2.341891805 | 3.513971357 |
| Q8BFW7 | Lpp | -2.342814915 | 4.453233937 |
| Q8JZX3 | Poc1a | -2.356370651 | 3.658727211 |
| Q80X90;Q8VHX6 | Cchcr1 | -2.35822102 | 5.56236107 |
| Q8K2I2 | Cchcr1 | -2.358361347 | 2.337453991 |
| P25322 | Ccnd1 | -2.358459632 | 2.482992247 |
| Q05895 | Thbs3 | -2.361554379 | 2.098774712 |
| Q9R112 | Sqor | -2.362077633 | 3.976172752 |
| Q8CIK8 | Rfwd3 | -2.362704563 | 2.207717731 |
| Q3V1H1 | Ckap2 | -2.364407556 | 3.889228954 |
| Q5NC05 | Ttf2 | -2.365216085 | 2.575135503 |
| Q9CQQ0 | Smim8 | -2.365609463 | 2.531369085 |
| Q8C3W1 |  | -2.377371444 | 2.992749711 |
| Q9WTK7 | Stk11 | -2.377846972 | 2.306445648 |
| Q08093 | Cnn2 | -2.378181748 | 4.586411796 |
| Q8C5L6 | Inpp5k | -2.381009643 | 2.083073453 |
| P97384 | Anxa11 | -2.386401315 | 3.172647072 |
| Q6ZPT1 | Klhl9 | -2.391410256 | 2.560511723 |
| Q810S1 | Mcub | -2.394249225 | 4.07646372 |
| P23780;Q8VC60 | Glb1 | -2.398834479 | 3.869017621 |
| Q499E4 | Dzip1l | -2.400830083 | 2.76338963 |
| Q9CQH3 | Ndufb5 | -2.405340014 | 2.331929692 |
| Q91Z49 | Fyttd1 | -2.414322042 | 3.423025771 |
| Q9D8S9 | Bola1 | -2.420523871 | 2.201876491 |
| Q8CAB8 | Castor2 | -2.421951632 | 2.089906628 |
| Q3UK10 | B9d2 | -2.435327362 | 3.399189558 |
| P27773 | Pdia3 | -2.440401797 | 3.820461367 |
| Q8BFR6 | Zfand1 | -2.444378707 | 2.840294868 |
| Q9JHH9 | Copz2 | -2.450613006 | 2.119950165 |
| Q9D0K2 | Oxct1 | -2.456174405 | 4.142594819 |
| Q99KW3 | Triobp | -2.459349136 | 5.705887197 |
| Q8CES0 | Naa30 | -2.460130387 | 2.134120772 |
| Q8R2X8 | Blzf1 | -2.460897217 | 2.23190828 |
| Q5SVD0 | Cdkn1a | -2.463427942 | 2.386764957 |
| Q9QYF1 | Rflnb | -2.468817649 | 2.2117295 |
| Q6P3B9 | Rdh11 | -2.470140136 | 2.885880465 |
| Q8BGV4 | Rbfa | -2.470900349 | 3.457450454 |
| O54724 | Tti2 | -2.477940695 | 3.943655471 |
| Q9CQ20 | Cavin1 | -2.480534677 | 2.227432039 |
| Q3UQ28 | Mid1ip1 | -2.481897079 | 5.00222714 |
| Q9D7I8 | Pxdn | -2.483487517 | 2.929542755 |
| Q9CPV1 | Fam83d | -2.485037591 | 3.613871478 |
| A0A7H0DN00 | Ska1 | -2.486375154 | 4.735413384 |
| Q61468 | OPG023 | -2.489145742 | 3.403725202 |
| Q6P4S6 | Msln | -2.494291023 | 2.284566702 |
| P97465 | Sik3 | -2.495013133 | 2.021248948 |
| Q80WW9 | Dok1 | -2.497464271 | 2.943391789 |
| Q9CWY4 | Ddrgk1 | -2.497767244 | 2.793211004 |
| Q9D2R8 | Gemin7 | -2.498419845 | 2.907374786 |
| O88512 | Mrps33 | -2.499935308 | 2.246559202 |
| Q9D2L9 | Ap1g2 | -2.500628847 | 3.125614093 |
| Q8CD10 | Fam111a | -2.508801942 | 3.138962129 |
| Q64267 | Micu2 | -2.51287154 | 2.248325858 |
| Q91XV3 | Xpa | -2.519535611 | 3.15226893 |
| Q8CGB6 | Basp1 | -2.5210902 | 2.464452299 |
| Q3UHC0 | Tnrc6c | -2.524550186 | 3.491838582 |
| Q6PFD6 | Kif18b | -2.524792058 | 3.077354391 |
| Q80YX1 | Tnc | -2.526332209 | 3.460817988 |
| Q8BGD8 | Coa6 | -2.533115699 | 3.696975432 |
| Q9CQZ1 | Hsbp1 | -2.533919427 | 3.941049284 |
| Q6NXY1 | Tbc1d31 | -2.538874057 | 2.254601834 |
| P16858 | Gapdh | -2.53921374 | 4.546805601 |
| P11688 | Itga5 | -2.543250212 | 4.430085309 |
| Q3USW5 | Foxred2 | -2.552107142 | 3.162674133 |
| Q8R0F5 | Rbmx2 | -2.556606508 | 3.020638008 |
| Q8R0K4 | Ccdc137 | -2.556805105 | 2.202920992 |
| Q9WTI7 | Myo1c | -2.557989348 | 4.622443967 |
| Q8BY89 | Slc44a2 | -2.562932636 | 2.646747397 |
| Q80UY1 | Carnmt1 | -2.568011911 | 2.804739128 |
| Q99LE1 | Rilpl2 | -2.570716935 | 2.533709888 |
| Q8VCP8 | Ak6 | -2.570901016 | 2.977447971 |
| Q9QXW0 | Fbxl6 | -2.573521817 | 2.149727283 |
| Q9DCH2 | Pop7 | -2.574310313 | 2.707470251 |
| Q61490 | Alcam | -2.577364856 | 4.488772042 |
| Q9R0X4;Q32MW3 | Acot9;Acot10 | -2.580264914 | 4.838901889 |
| Q91W18 | Tdrd3 | -2.580727649 | 3.182719283 |
| P09055 | Itgb1 | -2.5860077 | 4.586895624 |
| Q9CPP0 | Npm3 | -2.589795344 | 3.023473523 |
| Q9WU79 | Prodh | -2.59056863 | 3.81976436 |
| O08539 | Bin1 | -2.59179513 | 5.259521696 |
| A2ACJ2 | Faap100 | -2.592112199 | 2.337379753 |
| P50543 | S100a11 | -2.595594613 | 5.280410943 |
| Q8QZY6 | Tspan14 | -2.596109269 | 2.40859633 |
| Q8BIA4 | Fbxw8 | -2.610789798 | 3.795659315 |
| Q9D3W4 | Gpn3 | -2.612234216 | 2.942297297 |
| Q8BUE4 | Aifm2 | -2.612942939 | 4.046784378 |
| Q63870 | Col7a1 | -2.616054091 | 2.250970482 |
| P10852 | Slc3a2 | -2.617590636 | 3.116531034 |
| Q9D0C1 | Rnf115 | -2.620033138 | 3.911821591 |
| Q99KW9 | Itfg1 | -2.621591767 | 3.457450454 |
| Q9D8C3 | Alg13 | -2.628188266 | 2.296411533 |
| A2ADA5 | Pusl1 | -2.636045596 | 3.3506553 |
| Q8K305 | Nsl1 | -2.637093677 | 3.981409025 |
| Q8BMI4 | Gen1 | -2.637554233 | 4.144709934 |
| Q3TAS6 | Emc10 | -2.644015177 | 4.443807798 |
| Q8C6I2 | Sdhaf2 | -2.646595234 | 2.400471711 |
| Q9D142 | Nudt14 | -2.646881243 | 2.756943713 |
| Q9QY66 | Znhit2 | -2.64937213 | 2.283884975 |
| Q9WTR5 | Cdh13 | -2.653591653 | 2.082449995 |
| P10404;P11370 | Fv4 | -2.659396172 | 2.777946029 |
| Q91XE4 | Acy3 | -2.660199688 | 2.066217266 |
| Q9DB77 | Uqcrc2 | -2.661530156 | 3.705070314 |
| P97429 | Anxa4 | -2.661692629 | 4.177909425 |
| Q8R395 | Commd5 | -2.661736617 | 3.558554591 |
| Q61207 | Psap | -2.661996918 | 2.888193088 |
| Q8C0P7 | Znf451 | -2.662589329 | 3.83324051 |
| Q99L85 | Elp5 | -2.665455373 | 2.708645358 |
| Q9DBW3 | Natd1 | -2.670111963 | 2.117734187 |
| A2AM29 | Mllt3 | -2.670344354 | 2.980688013 |
| P07214 | Sparc | -2.673096478 | 4.931096713 |
| Q80Z37 | Topors | -2.673173015 | 2.247938357 |
| P18760;P45591 | Cfl | -2.674860226 | 3.264177327 |
| Q9DA19 | Cir1 | -2.677910174 | 2.641817994 |
| O09111 | Ndufb11 | -2.681840005 | 2.335968435 |
| Q8CGF1 | Arhgap29 | -2.686376382 | 2.12860797 |
| Q9JI08 | Bin3 | -2.688494607 | 2.119950165 |
| Q91WT4 | Dnajc17 | -2.68865949 | 3.491492343 |
| Q8K248 | Hpdl | -2.6909499 | 2.548448832 |
| Q99J47 | Dhrs7b | -2.697030305 | 3.474029774 |
| Q9Z223 | Mocs2 | -2.697291959 | 2.52310294 |
| Q3TBT3 | Sting1 | -2.700392923 | 3.62611643 |
| O88986 | Gcat | -2.702347042 | 4.443807798 |
| Q8BLH7 | Hirip3 | -2.715305275 | 2.979356407 |
| P40338 | Vhl | -2.722172535 | 2.034735386 |
| Q6GV12 | Kdsr | -2.72266886 | 2.061052835 |
| P54818 | Galc | -2.722851691 | 2.705698586 |
| P09528 | Fth1 | -2.723330236 | 2.093643614 |
| Q8JZV7 | Amdhd2 | -2.723880802 | 2.172710533 |
| Q7TPW1 | Nexn | -2.724678825 | 4.546805601 |
| Q924Z4 | Cers2 | -2.727092993 | 2.455348403 |
| Q9Z0R0 | Haspin | -2.734293204 | 2.096497827 |
| P47930 | Fosl2 | -2.739387769 | 4.701803578 |
| Q5HZH2 | Tsr3 | -2.740861702 | 3.823581864 |
| Q9DAX9 | Appbp2 | -2.74300451 | 2.516966102 |
| Q9DCT5 | Sdf2 | -2.74468369 | 2.063720716 |
| Q3U155 | Ccdc174 | -2.746908198 | 2.918107542 |
| Q9CR02 | Tma16 | -2.754388037 | 2.939326175 |
| P22935 | Crabp2 | -2.756172242 | 4.716211672 |
| Q9CZ09 | Mettl18 | -2.76239221 | 2.710782255 |
| Q02013 | Aqp1 | -2.765617189 | 2.30814192 |
| Q9D8Z1 | Ascc1 | -2.766125714 | 3.477130914 |
| P10833 | Rras | -2.774322403 | 2.850099865 |
| Q8BK58 | Hspbap1 | -2.776447347 | 4.034126349 |
| P03899 | mt-Nd3 | -2.776906993 | 2.039947742 |
| Q7TN02 | Med26 | -2.790490268 | 2.104857259 |
| Q9CPY0 | Mrm2 | -2.792602597 | 2.270263314 |
| Q9DAZ9 | Zfyve19 | -2.793315763 | 2.457811077 |
| Q9CPQ5 | Cenpq | -2.793920145 | 2.728700876 |
| Q9Z0L8 | Ggh | -2.798437362 | 3.930398719 |
| Q9CQ13 | Coprs | -2.798811802 | 3.417682121 |
| Q9CZI9 | Aen | -2.799233925 | 2.279416165 |
| Q8R2Y2 | Mcam | -2.800239257 | 2.389552795 |
| P97329 | Kif20a | -2.802849368 | 3.599567534 |
| Q62425 | Ndufa4 | -2.806137661 | 3.492310325 |
| P56390 | Cks2 | -2.81371003 | 3.333741648 |
| Q8K298 | Anln | -2.81614469 | 4.406131863 |
| Q4U4S6;Q9ERG0 | Xirp2 | -2.818321551 | 4.283768543 |
| Q924C6 | Loxl4 | -2.82130833 | 3.908383398 |
| Q8C551 | Rad51ap1 | -2.821912569 | 2.601787636 |
| Q9WUJ8 | Orc6 | -2.821948858 | 4.267781879 |
| Q9R1K5 | Fzr1 | -2.822653546 | 3.717352883 |
| Q9CWY3 | Setd6 | -2.824858792 | 2.70558575 |
| Q8BH04 | Pck2 | -2.825972313 | 4.586411796 |
| Q8R5A3 | Apbb1ip | -2.834806883 | 2.507659262 |
| Q9JJG0 | Tacc2 | -2.835101691 | 2.78149508 |
| Q64311 | Ntan1 | -2.845651627 | 2.22639028 |
| Q52KR3 | Prune2 | -2.849610492 | 2.076200959 |
| P09450 | Junb | -2.850186506 | 2.159709038 |
| Q9D0L7 | Armc10 | -2.852288535 | 3.414332145 |
| Q9QXS1 | Plec | -2.85470788 | 5.297187852 |
| Q61165 | Slc9a1 | -2.859297816 | 2.225967087 |
| Q8CIB9 | Esco2 | -2.860148306 | 2.183729035 |
| Q8BHA3 | Dtd2 | -2.863983653 | 3.53581653 |
| P06795 | Abcb1b | -2.864312496 | 4.050428806 |
| Q91VW5 | Golga4 | -2.871090467 | 2.051606281 |
| Q920E5 | Fdps | -2.873557277 | 4.451296921 |
| Q80W68 | Kirrel1 | -2.877050589 | 2.435505642 |
| Q8K136 | Scnm1 | -2.882516907 | 2.507354832 |
| O54818 | Tpd52l1 | -2.883052406 | 2.130024851 |
| Q8BHA0 | Ino80c | -2.889167396 | 2.49147034 |
| Q920A7 | Afg3l1 | -2.893490799 | 2.061561021 |
| P37040 | Por | -2.897236656 | 2.520133137 |
| Q80YR6 | Rbbp8 | -2.898897581 | 2.569681605 |
| Q91XB0 | Trex1 | -2.907671412 | 3.148537551 |
| Q8K273 | Mmgt1 | -2.909169926 | 2.020150611 |
| Q8C0M9 | Asrgl1 | -2.913977231 | 2.23485496 |
| P38060 | Hmgcl | -2.919807452 | 3.61425479 |
| Q8VE38 | Oxnad1 | -2.924507711 | 2.252707609 |
| O35492 | Clk3 | -2.926625317 | 2.428964036 |
| Q61263 | Soat1 | -2.930651228 | 3.107744365 |
| Q07417 | Acads | -2.931886877 | 4.443807798 |
| Q64437 | Adh7 | -2.933664054 | 2.213954046 |
| P70193 | Lrig1 | -2.935730663 | 2.462451693 |
| Q91YI0 | Asl | -2.939058118 | 4.673220337 |
| E9QAM5 | Helz2 | -2.940190478 | 2.294246642 |
| Q9CW79 | Golga1 | -2.941447045 | 3.920654828 |
| O08795 | Prkcsh | -2.945754785 | 2.086573012 |
| P58468 | Slx9 | -2.949336365 | 2.764848976 |
| Q8C0L6 | Paox | -2.952753614 | 2.372472076 |
| Q3UDP0 | Wdr41 | -2.958605153 | 2.35595023 |
| Q9JJX7 | Tdp2 | -2.9593685 | 2.047291019 |
| Q8BZT9 | Lacc1 | -2.963477647 | 3.647652241 |
| O54791 | Maff | -2.966375961 | 3.638918751 |
| Q9D0Y8 | Mrpl52 | -2.969775286 | 2.334918361 |
| Q8CC35 | Synpo | -2.971644312 | 2.141881152 |
| Q3TW96 | Uap1l1 | -2.974318972 | 4.402676241 |
| Q91ZX7 | Lrp1 | -2.975454574 | 3.999644043 |
| P03930 | Mtatp8 | -2.977992734 | 3.616701249 |
| P48410 | Abcd1 | -2.990015591 | 2.153550713 |
| Q9EQK5 | Mvp | -2.991193948 | 2.89371796 |
| Q8C181 | Mbnl2 | -2.992841187 | 3.146070578 |
| Q8CD19 | Lancl3 | -2.993279049 | 3.507268386 |
| Q61127 | Nab2 | -2.995996649 | 4.443807798 |
| Q8R550 | Sh3kbp1 | -2.99736931 | 3.339161001 |
| A0A7H0DNF5 | OPG193 | -2.998485332 | 4.300020713 |
| Q9D8U0 | Ifrd2 | -2.999657304 | 2.730331965 |
| Q8BHX3 | Cdca8 | -3.010732099 | 3.048093182 |
| Q8K3A0 | Hscb | -3.013598406 | 2.847494558 |
| Q99LR1 | Abhd12 | -3.022386529 | 2.433882306 |
| B1ARD6 | Slfn9 | -3.02349482 | 3.057579546 |
| P55302 | Lrpap1 | -3.024110682 | 4.6160186 |
| Q9D7K2 | Ten1 | -3.025748888 | 4.202329354 |
| Q9CYZ6 | Rex1bd | -3.025982406 | 3.398972681 |
| Q8BGR8 | Gskip | -3.026737247 | 2.660951667 |
| Q9D727 | Pex39 | -3.037310644 | 3.776839772 |
| Q8CC12 | Cdan1 | -3.038555636 | 2.956678904 |
| Q8K330 | Ssh3 | -3.044265016 | 2.181896398 |
| Q91UZ5 | Impa2 | -3.045398296 | 2.370810158 |
| P0DOV1 | Ifi211 | -3.051311745 | 3.900558998 |
| Q3UX61 | Naa11 | -3.052027014 | 3.701195153 |
| P40630 | Tfam | -3.054888113 | 2.592806947 |
| Q8VCE4 |  | -3.057586748 | 2.001176464 |
| Q8BXN9 | Tmem87a | -3.058322704 | 3.066288146 |
| Q9CZS3 | Cep20 | -3.058416623 | 2.911206772 |
| Q6P0X2 | Znf511 | -3.062675741 | 2.106035552 |
| Q5U5V2 | Hykk | -3.066229246 | 3.010661047 |
| Q9Z0R9 | Fads2 | -3.069328662 | 2.391133095 |
| Q00993 | Axl | -3.069482871 | 3.616460155 |
| Q62523 | Zyx | -3.07806485 | 4.935839333 |
| P70255 | Nfic | -3.079208641 | 2.8629281 |
| P07356 | Anxa2 | -3.082417865 | 4.903962268 |
| Q9EP53 | Tsc1 | -3.082776212 | 2.861477286 |
| Q9JL15 | Lgals8 | -3.085379071 | 2.479826172 |
| P56375 | Acyp2 | -3.090488365 | 2.117734187 |
| Q9EPK5 | Wwtr1 | -3.096068736 | 2.804767485 |
| Q9ET22 | Dpp7 | -3.111263681 | 3.460817988 |
| Q6PG16 | Hjurp | -3.125855725 | 2.257710116 |
| Q99P69 | Nuf2 | -3.12903391 | 2.017570472 |
| P46935;Q8CFI0 | Nedd4 | -3.133057904 | 4.48142587 |
| Q8R216 | Sirt4 | -3.141492674 | 2.40859633 |
| Q8BZB2 | Ppcdc | -3.143584649 | 2.279573711 |
| Q8CHQ0 | Fbxo4 | -3.152594125 | 3.302918845 |
| P24638 | Acp2 | -3.152664995 | 2.786103692 |
| P06869 | Plau | -3.153944936 | 4.020320572 |
| Q9D2R0 | Aacs | -3.156490915 | 4.463349979 |
| P41731 | Cd63 | -3.164203302 | 2.655774066 |
| Q99MS8 | Tpgs1 | -3.166351577 | 2.176114716 |
| P70699 | Gaa | -3.16741421 | 2.731634489 |
| Q9WTK5 | Nfkb2 | -3.16850449 | 4.769346497 |
| Q923Q2 | Stard13 | -3.172460364 | 2.126468004 |
| P01897;P01899;P01900;P01901;P03991;P04223;P06339;P14426;P14427;P14428;P14429;P14430;P14431 | H2 | -3.17455227 | 2.547392204 |
| Q9JKV5 | Scamp4 | -3.17561806 | 2.954232629 |
| Q61510 | Trim25 | -3.175809766 | 2.839000993 |
| O70348 | Dxo | -3.176097832 | 2.433537738 |
| A2A7S8 | Nhsl3 | -3.177229852 | 2.159951152 |
| Q3UFK8 | Frmd8 | -3.180630796 | 2.537959561 |
| Q91WK7 | Ankrd54 | -3.180873485 | 2.408252159 |
| O09000 | Ncoa3 | -3.185402411 | 2.226240267 |
| Q9CQL7 | Mrfap1 | -3.191226282 | 3.449232155 |
| O09174 | Amacr | -3.193370301 | 2.524343194 |
| P12850 | Cxcl1 | -3.19352776 | 3.146340191 |
| P61025 | Cks1b | -3.197652669 | 5.394910166 |
| Q91XC0 | Ajuba | -3.200848071 | 2.597420476 |
| Q9JJE7 | Fads3 | -3.201712846 | 2.439688546 |
| Q8BLY7 | Hps6 | -3.207055923 | 2.685553141 |
| Q9DCA2 | Mrps11 | -3.209427344 | 3.118279675 |
| Q9CZX7 | Pip4p2 | -3.213807331 | 4.186350718 |
| P46062 | Sipa1 | -3.215495862 | 2.422908127 |
| Q8VDP3 | Mical1 | -3.219033156 | 2.740317771 |
| O35671 | Itgb1bp1 | -3.219898994 | 2.595398235 |
| Q5DTX6 | Jcad | -3.228016324 | 2.206636945 |
| Q9D6H2 | Ift25 | -3.23118841 | 2.467463676 |
| P26645 | Marcks | -3.232098875 | 4.601001249 |
| O89086 | Rbm3 | -3.233641233 | 3.729385258 |
| Q8R3F5 | Mcat | -3.238064931 | 2.877797319 |
| P70255;P70257 | Nfix | -3.242071774 | 3.865642614 |
| Q60695 | Rgl1 | -3.243545319 | 2.136324802 |
| Q9R045 | Angptl2 | -3.249187194 | 2.449621786 |
| P58058 | Nadk | -3.250942267 | 3.460817988 |
| P97863 | Nfib | -3.25743849 | 2.58711636 |
| P97864 | Casp7 | -3.260465126 | 2.296411533 |
| P17047 | Lamp2 | -3.260656824 | 4.386402606 |
| Q8CBY1 | Samd4a | -3.266525244 | 3.407447327 |
| P70296 | Pebp1 | -3.273022681 | 2.577797759 |
| Q99JP6 | Homer3 | -3.273363907 | 2.620342399 |
| Q8BSP2 | Ncaph2 | -3.283956062 | 3.026827443 |
| Q8BUH8 | Senp7 | -3.286793029 | 2.023682351 |
| Q61503 | Nt5e | -3.295377008 | 3.255178832 |
| Q61391 | Mme | -3.298765928 | 2.465852631 |
| Q8BGH7 | Cdc42se2 | -3.301383379 | 2.030354266 |
| Q9JHJ3 | Glmp | -3.303669681 | 2.847494558 |
| Q9QZF2 | Gpc1 | -3.310380735 | 3.491492343 |
| P59941 | Sirt6 | -3.310444586 | 2.17038972 |
| Q8QZT1 | Acat1 | -3.313048188 | 4.682174872 |
| Q8K1C0 | Angel2 | -3.317040652 | 2.88268994 |
| Q9QWR8 | Naga | -3.320513108 | 2.882642228 |
| Q9Z0M5 | Lipa | -3.323356691 | 2.331929692 |
| Q62470 | Itga3 | -3.325299002 | 3.467028165 |
| Q8C142 | Ldlrap1 | -3.326550671 | 3.689452901 |
| Q64299 | Ccn3 | -3.327567974 | 3.464786034 |
| Q99MB2 | Mtfr1 | -3.32968526 | 2.723764442 |
| Q921K9 | Bcl7b | -3.330270018 | 2.651310071 |
| Q69ZN7;Q9ESD7 | Myof;Dysf | -3.330894662 | 2.344497964 |
| Q9CQF9 | Pcyox1 | -3.331553825 | 2.998966427 |
| Q8BW00 | Ptrh1 | -3.336930468 | 2.036226071 |
| Q75NR7 | Recql4 | -3.336957006 | 3.161148941 |
| Q99LC9 | Pex6 | -3.344945283 | 3.061953612 |
| Q61508 | Ecm1 | -3.347965527 | 3.06324362 |
| Q8R5A0 | Smyd2 | -3.351751335 | 2.392215652 |
| Q9CR59 | Gadd45gip1 | -3.360476804 | 2.379895868 |
| Q91W92 | Cdc42ep1 | -3.361676577 | 3.825318033 |
| O08967;P63034;Q9QX11 | Cyth | -3.364260647 | 2.441526503 |
| Q6NXW6 | Rad17 | -3.36464627 | 2.451365874 |
| Q9CZT8 | Rab3b | -3.364906672 | 2.335673222 |
| Q9D3D9 | Atp5f1d | -3.366638074 | 4.398373616 |
| Q76N33 | Stambpl1 | -3.368694521 | 3.001595404 |
| Q99L00 | Haus8 | -3.369672561 | 2.145692367 |
| P97784;Q9R194 | Cry1;Cry2 | -3.370457819 | 3.161148941 |
| Q9JHI2 | Adat1 | -3.371992428 | 4.102143461 |
| Q99JZ7 | Errfi1 | -3.374843603 | 2.465852631 |
| Q9CWY9 | Rpain | -3.376783426 | 2.5727015 |
| Q8VBT0 | Tmx1 | -3.379370755 | 2.940271383 |
| Q5EE38 | Acd | -3.381332111 | 3.1078441 |
| Q99PP9 | Trim16 | -3.383817374 | 3.20611795 |
| Q9R008 | Mvk | -3.39068374 | 2.444335896 |
| Q9D771 | Pacc1 | -3.400384801 | 3.021275001 |
| P22437;Q05769 | Ptgs1;Ptgs2 | -3.406283858 | 2.437910055 |
| Q6PDY2 | Ado | -3.406944331 | 2.64993696 |
| O35954 | Pitpnm1 | -3.411021208 | 2.324982144 |
| Q9CQY2 | Ramac | -3.413441845 | 3.202162361 |
| P49935 | Ctsh | -3.414034245 | 2.407330183 |
| P14719 | Il1rl1 | -3.415644761 | 2.558666806 |
| Q9JHN8 | Whr1 | -3.423400715 | 2.553549332 |
| Q60I26 | Als2cl | -3.425798461 | 2.674292378 |
| Q66GT5 | Ptpmt1 | -3.428091308 | 2.383287995 |
| Q8VDL4 | Adpgk | -3.428997258 | 2.927609989 |
| P50544 | Acadvl | -3.431517661 | 3.288531949 |
| Q99LB7 | Sardh | -3.440918079 | 2.68445133 |
| Q99JR5 | Tinagl1 | -3.447191991 | 2.423200171 |
| Q9DCZ1 | Gmpr | -3.454768666 | 3.180729511 |
| Q5U5M8 | Bloc1s3 | -3.455816223 | 2.046789648 |
| Q9CQX4 | Pclaf | -3.45679364 | 3.714630109 |
| Q78HU3 | Mvb12a | -3.464748044 | 3.403725202 |
| P70271 | Pdlim4 | -3.465243858 | 4.03401086 |
| Q80UU2 | Rpp38 | -3.465610317 | 2.862690165 |
| Q8CIG9 | Fbxl8 | -3.466334939 | 3.174917194 |
| Q68FH4 | Galk2 | -3.47067977 | 3.403725202 |
| Q3UA16 | Spc25 | -3.472608864 | 2.5829143 |
| P70202 | Lxn | -3.476759188 | 5.163589011 |
| Q99LC3 | Ndufa10 | -3.481684655 | 4.200200317 |
| Q80U49 | Cep170b | -3.482792733 | 2.419988616 |
| P02340 | Tp53 | -3.484774853 | 3.510054428 |
| Q8CB62 | Cntrob | -3.499987439 | 2.730150114 |
| Q3UHQ6 | Dop1b | -3.500216095 | 2.521994398 |
| Q5SSK3 | Tefm | -3.502862413 | 2.449167448 |
| Q80V94 | Ap4e1 | -3.504332879 | 5.691428711 |
| Q8VCH0;Q921H8 | Acaa1 | -3.509937243 | 2.825614264 |
| Q8VHK1 | Caskin2 | -3.512594862 | 2.537482152 |
| Q8BIG7 | Comtd1 | -3.521526801 | 2.072336366 |
| Q6NXJ0 | Wwc2 | -3.522165799 | 3.1798948 |
| Q8BQ47 | Cnpy4 | -3.531710414 | 5.16574648 |
| Q9D845 | Tex9 | -3.533565744 | 2.378218816 |
| Q60855 | Ripk1 | -3.55597094 | 2.012232019 |
| Q9CQR4 | Acot13 | -3.56505284 | 2.653637294 |
| Q3U962 | Col5a2 | -3.565983174 | 2.478868603 |
| B1ARD8;B1ARD6 | Slnf8 | -3.568329682 | 2.324060895 |
| P52760 | Rida | -3.570282775 | 3.658848198 |
| Q61554;Q61555 | Fbn1;Fbn2 | -3.574990889 | 3.696569095 |
| Q9D6Z0 | Alkbh7 | -3.577045133 | 3.491492343 |
| O88939 | Zbtb7a | -3.582569934 | 3.760306828 |
| P0C8B4 | Gon7 | -3.58486219 | 3.468448964 |
| Q60739 | Bag1 | -3.592616731 | 3.537822397 |
| Q8K3G5 | Vrk3 | -3.602904582 | 2.258972241 |
| P08207 | S100a10 | -3.603490302 | 4.741208574 |
| Q791N7 | Polr1h | -3.603866451 | 3.171904652 |
| Q6ZPG2 | Wdr90 | -3.606887364 | 2.479390447 |
| Q14AI6 | Rpusd3 | -3.619090782 | 2.706123832 |
| Q8BZA9 | Tigar | -3.62736404 | 2.709626057 |
| Q9D7N6 | Mrpl30 | -3.632745511 | 2.85615276 |
| Q64105 | Spr | -3.633378038 | 2.22891495 |
| Q9CYC5 | Dsn1 | -3.633734802 | 2.116966444 |
| Q91VE6 | Nifk | -3.636023097 | 2.182726659 |
| Q9D8B4 | Ndufa11 | -3.64182657 | 4.033884815 |
| O09044;P60879 | Snap23;Snap25 | -3.647890842 | 2.636898833 |
| Q8C8T8 | Tsr2 | -3.648322941 | 2.860483459 |
| Q9D1M7 | Fkbp11 | -3.652845189 | 2.821233618 |
| Q8R1G6 | Pdlim2 | -3.656090857 | 2.434031632 |
| Q64008 | Rab34 | -3.659441332 | 2.003992307 |
| O70161 | Pip5k1c | -3.665732317 | 2.617586593 |
| Q91WE1 | Snx15 | -3.666046133 | 2.44369799 |
| A2AG58 | Bclaf3 | -3.672843981 | 2.052128931 |
| Q9D853 | Eef1akmt2 | -3.673666907 | 2.64022117 |
| E9PY46 | Ift140 | -3.673880124 | 3.027560742 |
| Q61337 | Bad | -3.674755547 | 3.916107223 |
| Q3TVC7 | Tsr2 | -3.674869687 | 3.180729511 |
| Q8BYM8 | Fkbp11 | -3.682343615 | 2.701888523 |
| Q64337 | Pdlim2 | -3.683822983 | 4.821066334 |
| Q499E6 | Airim | -3.689098907 | 2.67228523 |
| P13020 | Gsn | -3.691692756 | 5.432912705 |
| Q9DB28 | Pop5 | -3.692348642 | 3.512397028 |
| P82343 | Renbp | -3.695210633 | 3.566802794 |
| Q6SKR2 | Hemk2 | -3.6996178 | 3.820461367 |
| Q8CBY0 | Gatc | -3.699708928 | 2.48796244 |
| P25085 | Il1rn | -3.704097952 | 2.745226175 |
| Q8R3F9 | Tut1 | -3.709667433 | 2.036770217 |
| P29351 | Ptpn6 | -3.71579581 | 3.163015411 |
| Q9D198 | Syf2 | -3.719058742 | 3.217547766 |
| Q8N9S3 | Ahsa2 | -3.727919778 | 3.057121343 |
| Q9Z0P4 | Palm | -3.729956888 | 3.369051266 |
| P54761 | Pars2 | -3.738091549 | 2.35595023 |
| Q8CFI5 | Rcbtb2 | -3.738094776 | 2.712746161 |
| Q99LJ7 | Tradd | -3.73883538 | 3.260593185 |
| Q3U0V2 | Cdkn2c | -3.751496796 | 2.632497164 |
| Q60772 | Ift122 | -3.758445185 | 4.142594819 |
| Q6NWV3 | Bid | -3.759677294 | 2.227432039 |
| Q05769 | Ndor1 | -3.767050855 | 4.317828985 |
| P70444 | Il1rn | -3.768580973 | 4.000178532 |
| A2AI05 | Tut1 | -3.770579679 | 3.268331914 |
| Q640M1;Q6EJB6 | Utp14 | -3.771783023 | 3.233471219 |
| P56542 | Dnase2 | -3.774283936 | 3.695788405 |
| Q8BK35 | Nop53 | -3.775392748 | 3.302313576 |
| Q7TNE3 | Spag7 | -3.78094589 | 3.008268744 |
| Q9CWT6 | Ddx28 | -3.784790678 | 2.044607208 |
| P10649 | Gstm1 | -3.795994613 | 4.180771116 |
| P98063 | Bmp1 | -3.799162724 | 3.81005291 |
| P49138 | Mapkapk2 | -3.7997655 | 3.856352223 |
| Q9CQL6 | Mrpl35 | -3.800602433 | 3.916107223 |
| Q91ZF0 | Dnajc24 | -3.801644887 | 3.240849705 |
| P0DOV1;P0DOV2 | Ifi204 | -3.805804375 | 3.371943773 |
| Q5PSV9 | Mdc1 | -3.806631516 | 3.454451089 |
| Q64339 | Isg15 | -3.810775423 | 5.297187852 |
| P22366 | Myd88 | -3.811357683 | 3.712661454 |
| Q9CYA0 | Creld2 | -3.811627613 | 4.756852261 |
| P35288 | Rab23 | -3.814228073 | 2.630569915 |
| Q9JL26 | Fmnl1 | -3.818190668 | 2.685585739 |
| P11679;P15331;P20152;P31001;Q9DCV7 | Krt8;Krt9 | -3.820469279 | 4.301764195 |
| Q9D009 | Lipt2 | -3.824247964 | 2.038878375 |
| Q8C407 | Yipf4 | -3.826790687 | 3.82215772 |
| Q9Z0L0 | Tpbg | -3.830942139 | 2.102290164 |
| Q922Q1 | Mtarc2 | -3.836896196 | 2.797355395 |
| Q80VI1 | Trim56 | -3.848089055 | 4.287039153 |
| Q8BXK8 | Agap1 | -3.853728664 | 2.771609785 |
| Q8K1I7 | Wipf1 | -3.860578994 | 2.102290164 |
| Q8BR63 | Fam177a1 | -3.868841687 | 2.239321663 |
| Q9JIM1 | Slc29a1 | -3.869281994 | 3.514510419 |
| Q8R3P0 | Aspa | -3.870955303 | 4.344325268 |
| Q7TME2 | Spag5 | -3.874006275 | 2.389978697 |
| P58059 | Mrps21 | -3.875049731 | 3.300717051 |
| Q6ZQJ5 | Dna2 | -3.881229222 | 2.464184129 |
| P28650 | Adss1 | -3.884521367 | 2.471495032 |
| Q9DC71 | Mrps15 | -3.886992192 | 2.64993696 |
| P18828 | Sdc1 | -3.887558906 | 2.368540357 |
| Q9Z0F6 | Rad9a | -3.889795441 | 2.971092202 |
| Q8BVA5 | Ldah | -3.891793392 | 3.816431211 |
| Q9JM96 | Cdc42ep4 | -3.894787773 | 2.298478587 |
| Q8BK30 | Ndufv3 | -3.900690042 | 3.531635505 |
| Q8R1F6 | Hid1 | -3.901974576 | 3.457450454 |
| Q8BYY4 | Ttc39b | -3.90441606 | 2.287729896 |
| Q8R2Q8 | Bst2 | -3.904717013 | 2.458870221 |
| Q9CRA7 | Dmac2l | -3.909076039 | 3.709482554 |
| Q7TMR0 | Prcp | -3.922416752 | 3.361470749 |
| Q91YU8 | Ppan | -3.926156857 | 3.193556332 |
| Q9CWH5 | Trmt11 | -3.92628435 | 4.30521988 |
| P63013 | Prrx1 | -3.929253557 | 3.36409455 |
| Q9EPR5 | Sorcs2 | -3.929865161 | 2.224816401 |
| Q64092 | Tfe3 | -3.930680298 | 2.601229478 |
| Q71FD5 | Znrf2 | -3.932295985 | 2.283884975 |
| Q8C5P5 | Nt5dc1 | -3.932534508 | 3.701535962 |
| Q9CXX9 | Cuedc2 | -3.933306849 | 2.141979038 |
| Q3V4B5 | Commd6 | -3.93667856 | 4.868785445 |
| Q6PGG2 | Gmip | -3.943827918 | 2.467417086 |
| O88668 | Creg1 | -3.947920802 | 3.429313103 |
| Q61387 | Cox7a2l | -3.950856446 | 2.416207182 |
| Q3URQ7 | Mthfsd | -3.954070234 | 3.594782061 |
| P97315 | Csrp1 | -3.955673904 | 4.622443967 |
| Q8K4M5 | Commd1 | -3.96043277 | 2.368286895 |
| Q9ESP1 | Sdf2l1 | -3.967341093 | 3.255090432 |
| Q7TPV2 | Dzip3 | -3.970018886 | 2.806774018 |
| Q8BP40 | Acp6 | -3.973614483 | 2.619246657 |
| P43883 | Plin2 | -3.974378376 | 4.12874643 |
| Q9CQ91 | Ndufa3 | -3.974907443 | 3.240750912 |
| Q76KJ5 | Polr1g | -3.997363824 | 2.221584164 |
| Q80Y50 | Camta2 | -4.004772853 | 3.474029774 |
| Q8R080 | Gtse1 | -4.005890905 | 2.251799219 |
| O88413 | Tulp3 | -4.008952506 | 3.234670093 |
| Q9CR75 | Tnfrsf12a | -4.016523447 | 2.671731443 |
| O09159 | Man2b1 | -4.031272494 | 3.491492343 |
| Q8C3S2 | Tango6 | -4.031601777 | 3.415288168 |
| Q5I1X5 | Ppp1r13l | -4.037763861 | 3.332067245 |
| Q3UJP5 | Cfap418 | -4.03988638 | 4.340704687 |
| O08734 | Bak1 | -4.040186844 | 2.576400606 |
| Q9CRY7 | Gdpd1 | -4.044966367 | 3.19910958 |
| Q8VDT9 | Mrpl50 | -4.046010315 | 3.620965596 |
| Q99M15 | Pstpip2 | -4.046467701 | 2.207717731 |
| P97352 | S100a13 | -4.04900721 | 3.828873872 |
| Q99K85 | Psat1 | -4.056392018 | 5.059329797 |
| Q8C2E4 | Ptcd1 | -4.0566188 | 2.141697567 |
| Q91YR9 | Ptgr1 | -4.058660636 | 4.060997583 |
| Q9CZG3 | Commd8 | -4.063496716 | 2.413118057 |
| Q78IK4 | Apool | -4.066578086 | 3.715037591 |
| P30412 | Ppic | -4.068019192 | 2.281500876 |
| P52624 | Upp1 | -4.068830878 | 4.759418207 |
| Q99KS2 | Ngrn | -4.071007616 | 4.215418811 |
| Q8CJ26 | Nradd | -4.072502303 | 3.149826054 |
| O35988 | Sdc4 | -4.072662093 | 2.853394911 |
| P06151 | Csad | -4.073404755 | 3.399360927 |
| Q9DBE0 | Zcchc10 | -4.075666129 | 3.782846274 |
| Q9CX48 | Nopchap1 | -4.076479651 | 2.413417836 |
| Q9CX66 | Kifc3 | -4.08517995 | 3.086587137 |
| O35231 | Hsd17b12 | -4.094925348 | 3.637487765 |
| O70503 | Cst3 | -4.099573157 | 2.577900209 |
| P21460 | Mrtfb | -4.099745679 | 2.604983142 |
| P59759 | H1-5 | -4.100586319 | 2.321509786 |
| P43276 | Mast4 | -4.100847436 | 3.647210969 |
| Q811L6 | Decr1 | -4.10210514 | 2.353536567 |
| Q9CQ62 | Malsu1 | -4.104369866 | 3.732564997 |
| Q9CWV0 | Serpinb6 | -4.105366724 | 2.510137343 |
| Q60854 | Tceal9 | -4.115031963 | 2.009936553 |
| Q9DD24 | Fam110c | -4.11521347 | 2.519303924 |
| Q8VE94 | Med9 | -4.118946267 | 3.761064182 |
| Q8VCS6 | Psat1 | -4.128101452 | 2.468928662 |
| P29391;P49945 | Ftl1 | -4.129091331 | 2.559830579 |
| Q91V76 |  | -4.129447022 | 3.11279876 |
| P10107 | Anxa1 | -4.133264688 | 4.622358186 |
| Q80Z25 | Ofd1 | -4.136836988 | 2.449621786 |
| O89023 | Tpp1 | -4.138700128 | 3.216991014 |
| O08601 | Mttp | -4.138953681 | 2.113020597 |
| B2RXR6 | Ankrd44 | -4.159533075 | 4.005379984 |
| O54784 | Dapk3 | -4.164267418 | 2.954900957 |
| P12032 | Timp1 | -4.16578167 | 3.424295286 |
| O89110 | Casp8 | -4.167397265 | 2.562657435 |
| Q8K039 |  | -4.177812875 | 2.745512984 |
| Q91W40 | Klc3 | -4.184409465 | 2.005476554 |
| Q9QZM4 | Tnfrsf10b | -4.187238152 | 4.406131863 |
| P24452 | Capg | -4.200623388 | 5.070163205 |
| Q8R5C8 | Zmynd11 | -4.203462844 | 3.324876083 |
| Q9DBV4 | Mxra8 | -4.211254573 | 2.170049693 |
| Q69ZZ9 | Macf1 | -4.214291799 | 3.627441842 |
| O35405 | Pld3 | -4.225168879 | 3.530709373 |
| Q9CR89 | Ergic2 | -4.233570104 | 2.739175354 |
| Q8K2M0 | Mrpl38 | -4.241512286 | 4.53993979 |
| Q9JMB0 | Gkap1 | -4.242344592 | 2.170872001 |
| Q3UW53 | Niban1 | -4.24877869 | 2.100946362 |
| P10605 | Ctsb | -4.251901613 | 4.982715035 |
| P37889 | Fbln2 | -4.253287828 | 2.673651415 |
| Q62087 | Pon3 | -4.257038473 | 2.881799895 |
| Q9D842 | Aplf | -4.258276311 | 2.155587707 |
| Q8R2L5 | Mrps18c | -4.269784512 | 3.645500577 |
| Q9CPW3 | Mrpl54 | -4.273487926 | 5.255932293 |
| Q62187 | Ttf1 | -4.277334474 | 3.050158706 |
| Q8R4N0 | Clybl | -4.278587101 | 4.222797296 |
| P54923 | Adprh | -4.278705673 | 3.823581864 |
| P09926 | Surf2 | -4.279404103 | 3.713926904 |
| Q99JP4 | Cdc26 | -4.283352724 | 2.071429066 |
| P28033 | Cebpb | -4.284779749 | 2.652123322 |
| P07091 | S100a4 | -4.292550571 | 3.915553082 |
| Q9JLR9 | Higd1a | -4.310867851 | 2.830545107 |
| P20065 | Tmsb4x | -4.312554047 | 4.144758147 |
| Q6F3F9 | Adgrg6 | -4.314029042 | 3.689452901 |
| Q8R164 | Bphl | -4.317696884 | 4.149489159 |
| Q9CPV5 | Pmf1 | -4.318408764 | 3.616465406 |
| Q9CY28 | Gtpbp8 | -4.329840405 | 2.417626105 |
| Q8K2J0 | Plcd3 | -4.331135648 | 2.623895764 |
| Q9DCS2 | Mettl26 | -4.333665466 | 2.836793386 |
| Q14AI0 | DSCC1 | -4.335166843 | 2.793102901 |
| Q8BFS6 | Cpped1 | -4.343205265 | 3.386841655 |
| B1AR13 | Cisd3 | -4.349992274 | 2.872212454 |
| Q3TUH1 | Tamm41 | -4.350652869 | 3.027253272 |
| Q8BHY2 | Noc4l | -4.364659092 | 3.362617685 |
| O08997 | Atox1 | -4.369633229 | 3.49553074 |
| Q9CR80 | Fam32a | -4.371560835 | 2.530037485 |
| Q8BHE0 | Prr11 | -4.379007853 | 2.539359339 |
| Q8CDJ8 | Ston1 | -4.383565635 | 2.374305668 |
| P35456 | Plaur | -4.386877967 | 3.67110646 |
| Q9EPL9 | Acox3 | -4.400690501 | 3.843530949 |
| Q80WC7 | Agfg2 | -4.403329909 | 2.149303108 |
| Q9D6Y7 | Msra | -4.408661757 | 3.209034268 |
| Q3UIU2 | Ndufb6 | -4.410349573 | 2.967792554 |
| Q9QWF0 | Chaf1a | -4.412808517 | 3.206572311 |
| Q9CQ54 | Ndufc2 | -4.425564093 | 4.111574101 |
| Q9CXL3 | Chlsn | -4.43462232 | 2.252520944 |
| Q3UMW8 | Cln5 | -4.437210761 | 2.922773442 |
| Q3UKC1 | Tax1bp1 | -4.438444544 | 2.901093035 |
| Q6Q899 | Rigi | -4.44502502 | 2.368286895 |
| Q99PG2 | Ogfr | -4.446833009 | 4.202329354 |
| Q6PE15 | Abhd10 | -4.451508759 | 2.117734187 |
| Q63918 | Cavin2 | -4.462821931 | 3.268774511 |
| Q9CQ06 | Mrpl24 | -4.463108536 | 2.294331735 |
| P02802 | Mt1 | -4.464444037 | 3.074356549 |
| Q7TMV3 | Fastkd5 | -4.465902585 | 3.032666154 |
| Q924T3 | Xrcc4 | -4.47063737 | 3.474029774 |
| Q8BHX1 | Haus1 | -4.481025985 | 2.797355395 |
| Q9CR25 | Dph2 | -4.492180061 | 4.215415614 |
| Q8C263 | Ska3 | -4.494004572 | 2.869070839 |
| Q5SUQ9 | Ctc1 | -4.495735193 | 3.965961608 |
| Q99MZ7 | Pecr | -4.495859907 | 3.647780156 |
| P54754;P54761;P54763;Q03137;Q03145;Q61772;Q62413;Q8CBF3 | Ephb;Epha | -4.497872211 | 2.51947099 |
| Q6S5J6 | Krit1 | -4.498650516 | 2.337177106 |
| P81069 | Gabpb2 | -4.501873566 | 2.751967148 |
| P48725 | Pcnt | -4.502541414 | 4.034578328 |
| Q64701 | Rbl1 | -4.511437343 | 2.134219874 |
| O88824 | Jtb | -4.511676596 | 2.801800901 |
| Q91VM5 | Rbmxl1 | -4.521395303 | 2.112712528 |
| Q9D6U8 | Fam162a | -4.525691359 | 3.862103011 |
| Q9D2Y4 | Mlkl | -4.527947757 | 3.440172329 |
| Q8BSL7 | Arf2 | -4.528093454 | 2.872906427 |
| Q9DBG5 | Plin3 | -4.533447698 | 3.073582798 |
| Q99KR3 | Lactb2 | -4.541667204 | 6.236924249 |
| P0DJF2 | Pet117 | -4.542187526 | 2.306305277 |
| Q7TPV4 | Mybbp1a | -4.545087057 | 3.772582368 |
| Q9CZ42 | Naxd | -4.550009634 | 3.264949565 |
| A0A7H0DNF6 | OPG195 | -4.553653044 | 3.406018147 |
| Q80WR5 |  | -4.573353932 | 2.698215858 |
| Q9DCX1 | Mad2l1bp | -4.586303549 | 2.484546304 |
| Q62426 | Cstb | -4.59086502 | 4.934479354 |
| P0DN34 | Ndufb1 | -4.593218346 | 3.027253272 |
| Q3TV70 | Nr2c2ap | -4.594659969 | 2.487983348 |
| Q9D2Q3 | Paat | -4.59827539 | 2.151979058 |
| P16045 | Lgals1 | -4.607339209 | 5.661601374 |
| Q3TL26 | Tfb2m | -4.607450615 | 2.688747951 |
| Q8CIC2 | Nup42 | -4.610643137 | 2.686730664 |
| Q9WVL0 | Gstz1 | -4.61273239 | 3.373359825 |
| Q8BYU6 | Tor1aip2 | -4.621036473 | 4.057184233 |
| Q6P3Y5 | Znf280c | -4.623941919 | 2.697386722 |
| Q8C1M2 | Znf428 | -4.628591944 | 2.864424984 |
| P51174 | Acadl | -4.637929812 | 4.433007491 |
| Q9D1L0 | Chchd2 | -4.648973547 | 3.141875904 |
| Q8C3R1 | Brat1 | -4.65246434 | 2.714030097 |
| Q9CZX5 | Pinx1 | -4.653107312 | 2.332382659 |
| Q99MS7 | Ehbp1l1 | -4.653432 | 2.347731792 |
| Q8CEE6 | Pask | -4.66161091 | 2.827398513 |
| Q99LW0 | Ankrd10 | -4.675020384 | 2.216814285 |
| O35639 | Anxa3 | -4.675758035 | 5.564310284 |
| Q9CXC3 | Mgme1 | -4.675794509 | 2.491956998 |
| P50428 | Arsa | -4.677354406 | 2.542580232 |
| Q99L04 | Dhrs1 | -4.682023838 | 5.123375005 |
| P50429 | Arsb | -4.686007854 | 2.316747046 |
| P45377 | Akr1b8 | -4.690547938 | 2.017436339 |
| Q3ULW8 | Parp3 | -4.691125937 | 4.488724814 |
| P10923 | Spp1 | -4.696546729 | 2.510137343 |
| Q78WZ7 | Polr1f | -4.715898044 | 3.537699421 |
| O55101 | Syngr2 | -4.717757533 | 2.971127483 |
| Q8BIW1 | Prune1 | -4.720055837 | 3.466526223 |
| Q9R060 | Nubp1 | -4.721983525 | 2.143924652 |
| Q61846 | Melk | -4.722981136 | 3.325380691 |
| Q9CQP3 | Chchd5 | -4.738000651 | 2.512988546 |
| P48432 | Sox2 | -4.745068146 | 2.926053741 |
| Q80UW2 | Fbxo2 | -4.748268362 | 2.301395469 |
| Q8BGB5 | Limd2 | -4.753371882 | 3.06014031 |
| Q9CQU5 | Zwint | -4.777707059 | 2.622056965 |
| Q8R570 | Snap47 | -4.781146142 | 2.292405579 |
| Q8BL80 | Arhgap22 | -4.787533399 | 2.514821415 |
| Q8VCR7 | Abhd14b | -4.799174825 | 3.106574663 |
| Q3UZA1 | Rcsd1 | -4.800611272 | 3.027751947 |
| Q8BVN4 | Mterf4 | -4.80222567 | 2.450998639 |
| Q8BVY0 | Rsl1d1 | -4.831976594 | 3.884878028 |
| Q91VI7 | Rnh1 | -4.869106307 | 4.759418207 |
| Q9CQC7 | Ndufb4 | -4.873757846 | 3.353565241 |
| P97821 | Ctsc | -4.874706274 | 3.168113927 |
| P15535 | B4galt1 | -4.879864599 | 2.051975847 |
| Q8CFV9 | Rfk | -4.886989129 | 3.589295737 |
| Q9CPT4 | Mydgf | -4.889530594 | 2.400456418 |
| Q9CQ18 | Rnaseh2c | -4.889659033 | 4.546417755 |
| Q9D937 |  | -4.91367512 | 2.64993696 |
| Q8BHL4 | Gprc5a | -4.918207385 | 5.569291115 |
| Q99N87 | Mrps5 | -4.924973356 | 4.970406784 |
| Q60778 | Nfkbib | -4.925443841 | 2.549783631 |
| Q9CWG8 | Ndufaf7 | -4.96406976 | 3.073438687 |
| Q6ZWR6 | Syne1 | -4.964553285 | 2.140211232 |
| Q9D1R2 | Kti12 | -4.969170379 | 4.56336512 |
| Q3TLP5 | Echdc2 | -4.973173368 | 3.246461291 |
| P49962 | Srp9 | -4.986836038 | 2.162329795 |
| Q8K124 | Plekho2 | -4.987169728 | 4.125187546 |
| Q8K1J5 | Sde2 | -4.99585894 | 4.12618681 |
| Q60953 | Pml | -5.017469823 | 3.175541775 |
| Q8K2T1 | Nmral1 | -5.027893674 | 3.285203531 |
| Q9Z2C5 | Mtm1 | -5.032352202 | 2.192351077 |
| Q62266 | Sprr1a | -5.049872818 | 2.416919694 |
| P19157;P46425 | Gstp | -5.056568182 | 5.902972035 |
| P48755 | Fosl1 | -5.073864525 | 3.924647691 |
| Q8C052 | Map1s | -5.076328649 | 2.063742219 |
| Q60767 | Ly75 | -5.087498822 | 3.36409455 |
| Q9EPB5 | Serhl | -5.137125325 | 2.64305926 |
| Q8R0X7 | Sgpl1 | -5.165118878 | 2.781239713 |
| P48771 | Cox7a2 | -5.167366952 | 3.227469905 |
| P70671 | Irf3 | -5.175051596 | 3.561899877 |
| Q61738 | Itga7 | -5.176701576 | 2.477091785 |
| Q9CWX2 | Ndufaf1 | -5.191843339 | 2.491956998 |
| P42125 | Eci1 | -5.200665614 | 4.607199918 |
| Q9DBE8 | Alg2 | -5.206599951 | 2.716595031 |
| O35658 | C1qbp | -5.219318617 | 5.386150482 |
| P00405 | Mtco2 | -5.244018493 | 4.56955956 |
| P40240 | Cd9 | -5.253395176 | 3.972076717 |
| P35762 | Cd81 | -5.263017232 | 2.994654892 |
| Q9D786 | Haus5 | -5.266331071 | 2.314664481 |
| Q8VI94 | Oasl1 | -5.278412648 | 3.822122385 |
| Q9DBR0 | Akap8 | -5.285938091 | 2.2042313 |
| Q8BJY1 | Psmd5 | -5.314213652 | 3.645432586 |
| P28653 | Bgn | -5.323602721 | 3.438612496 |
| P56135 | Atp5mf | -5.337100323 | 4.634425399 |
| Q9CX53 | Gemin6 | -5.341609139 | 2.04481945 |
| Q64364 | Cdkn2a | -5.359979361 | 3.017859178 |
| Q3UGC7 | Eif3j1 | -5.385450129 | 3.561899877 |
| Q99K30 | Eps8l2 | -5.385588659 | 3.503412423 |
| P12265 | Gusb | -5.387348869 | 2.2412059 |
| P52624;Q8CGR7 | Upp2 | -5.388439215 | 3.302313576 |
| Q8K3D3 | Swi5 | -5.405828435 | 3.292104905 |
| Q99J99 | Mpst | -5.410826697 | 3.571577841 |
| P35951 | Ldlr | -5.414049666 | 3.753090624 |
| P99028 | Uqcrh | -5.422269973 | 4.953073521 |
| Q9D1C3 | Pyurf | -5.426204694 | 3.137411878 |
| Q921S7 | Mrpl37 | -5.438267941 | 2.179903101 |
| Q91VN4 | Chchd6 | -5.439432042 | 3.250444941 |
| P60330 | Espl1 | -5.446860539 | 2.822631661 |
| Q9D9M5 | Phospho2 | -5.483125384 | 2.916475167 |
| Q9CRB2 | Nhp2 | -5.489758164 | 4.920086943 |
| Q9QZN4 | Fbxo6 | -5.499393197 | 3.806485266 |
| Q99LY9 | Ndufs5 | -5.502432921 | 3.673105973 |
| Q924T2 | Mrps2 | -5.50362277 | 3.637487765 |
| Q8BQ30 | Ppp1r18 | -5.506295211 | 4.970406784 |
| Q9DCS9 | Ndufb10 | -5.530497338 | 4.673220337 |
| Q8VBT6 | Apobr | -5.543994549 | 2.785537672 |
| P99025 | Gchfr | -5.546625932 | 3.103505744 |
| Q80ZQ9 | Abitram | -5.555577592 | 2.597608149 |
| O09117 | Sypl1 | -5.562824038 | 3.193556332 |
| Q6PIU9 |  | -5.572060201 | 2.576520234 |
| Q8C650 | Septin10 | -5.592092243 | 2.309225271 |
| Q9DB29 | Iah1 | -5.601633423 | 3.125560883 |
| Q8R035 | Mrpl58 | -5.620301949 | 2.870478821 |
| P97450 | Atp5pf | -5.623703714 | 3.344575093 |
| Q9Z127 | Slc7a5 | -5.62703809 | 3.442935524 |
| Q8BKT8 | Haus7 | -5.633567862 | 2.747639929 |
| Q3TZX8 | Nol9 | -5.633667942 | 3.167791656 |
| Q6PFX9 | Tnks | -5.636652336 | 2.844082067 |
| O35640 | Anxa8 | -5.65010599 | 4.453233937 |
| P46656 | Fdx1 | -5.667111514 | 2.453076358 |
| Q61398 | Pcolce | -5.670769339 | 2.832640344 |
| O54974 | Lgals7 | -5.680307736 | 2.250472338 |
| Q8R344 | Ccdc12 | -5.698488398 | 2.048333816 |
| Q8VC52 | Rbpms2 | -5.70093031 | 3.310489487 |
| P56389 | Cda | -5.703635758 | 2.474117409 |
| Q9DCT1 | Akr1e2 | -5.706897507 | 3.847788825 |
| P62965 | Crabp1 | -5.712076411 | 3.853392838 |
| Q0P678 | Zc3h18 | -5.718202022 | 4.931096713 |
| Q9DCS3 | Mecr | -5.733662311 | 2.768457938 |
| Q9JL35 | Hmgn5 | -5.740131326 | 2.916313283 |
| P24860 | Ccnb1 | -5.759809157 | 2.491956998 |
| Q8R1I1 | Uqcr10 | -5.803638028 | 3.038273344 |
| Q3UPF5 | Zc3hav1 | -5.81068374 | 2.49428361 |
| Q6P5H2 | Nes | -5.823694099 | 4.004922876 |
| Q8R422 | Cd109 | -5.823695538 | 3.460817988 |
| Q9QYM8 | Cenph | -5.829262552 | 3.864244963 |
| Q9CPQ1 | Cox6c | -5.840285962 | 3.64462598 |
| P08228 | Sod1 | -5.891048852 | 3.159073857 |
| P52927 | Hmga2 | -5.899612624 | 2.446380845 |
| P70677 | Casp3 | -5.915763677 | 2.25745568 |
| Q8BGX2 | Timm29 | -5.935805874 | 2.492968002 |
| Q9CQZ5 | Ndufa6 | -5.952982755 | 3.857827799 |
| Q9CQM5 | Txndc17 | -5.963669793 | 4.026340821 |
| Q9CQJ8 | Ndufb9 | -5.980613377 | 2.358309035 |
| P21956 | Mfge8 | -6.029316501 | 2.063229881 |
| P57722 | Pcbp3 | -6.058095418 | 2.829010464 |
| P06797 | Ctsl | -6.082752409 | 3.689452901 |
| P28798 | Grn | -6.082759483 | 2.724403382 |
| Q8CJ27 | Aspm | -6.104353595 | 3.020029169 |
| P56183 | Rrp1 | -6.119676815 | 3.118279675 |
| Q9R1Q7 | Plp2 | -6.12759506 | 4.324572957 |
| Q9D945 | Llph | -6.152202613 | 3.110383486 |
| Q9QZM2 | Polg2 | -6.167241628 | 2.764468928 |
| P31786 | Dbi | -6.193645433 | 5.297187852 |
| Q9WV54 | Asah1 | -6.210930506 | 2.12702023 |
| Q9CX60 | Lbh | -6.222921434 | 2.690436146 |
| Q9D083 | Spc24 | -6.294112063 | 3.507268386 |
| Q60872 | Eif1a | -6.300477685 | 2.442189343 |
| P97825 | Jpt1 | -6.310318576 | 3.773615569 |
| Q80VJ3 | Dnph1 | -6.373366651 | 4.333580364 |
| P97370 | Atp1b3 | -6.475076741 | 2.359225332 |
| Q9CPQ8 | Atp5mg | -6.499822633 | 4.213349624 |
| Q9CQ75 | Ndufa2 | -6.527983521 | 2.739640072 |
| P56213 | Gfer | -6.528587398 | 2.09285816 |
| P15379 | Cd44 | -6.562051152 | 3.383117176 |
| Q00493 | Cpe | -6.572398347 | 2.102538323 |
| P18572 | Bsg | -6.638117046 | 5.015593452 |
| Q80ZS3 | Mrps26 | -6.762360953 | 3.825318033 |
| P51125 | Cast | -6.774875496 | 4.803217743 |
| P16110 | Lgals3 | -6.802493497 | 5.569291115 |
| Q9CWY8 | Rnaseh2a | -6.905245908 | 2.801800901 |
| Q80U04 | Pja2 | -6.932286713 | 2.405020944 |
| O09131 | Gsto1 | -7.841946876 | 2.216805338 |
| Q8VCN5 | Cth | -7.913077042 | 3.020638008 |
| Q9QZL0 | Ripk3 | -8.078605617 | 3.259143974 |

**Table S5.** *Protein signature of WRAIR MOI 1 infected KO vs WT MEF lysates at 21 hours post-infection. Table lists the protein ID, gene name, the log2-fold change (FC) (MPXV infected WT MEFs vs infected ISG15 KO MEFs) of the levels of each protein, and the statistical significance (−log P value). Ordered from largest to smallest FC. Genes mentioned in manuscript highlighted in bold.*
