## Supplementary material for "ISG15 Differentially Modulates Clade Ib and II MPXV Infection in MEF cells": TableS6

| **Name** | **Gene** | **fold_change** | **p_value_adj_neg_log10** |
| --- | --- | --- | --- |
| Q61474 | Msi1 | 5.792516628 | 2.432072408 |
| Q6PHU5 | Sort1 | 5.441583339 | 2.463270389 |
| Q8BG16 | Slc6a15 | 5.191954035 | 2.025608437 |
| Q9QYB2 | Dach1 | 4.846550931 | 3.046837233 |
| P41139 | Id4 | 4.597443809 | 2.146825289 |
| P60469 | Ppfia3 | 4.560795151 | 2.719371343 |
| Q3UPL5 | Ag2 | 4.55890702 | 4.166200583 |
| Q8K0S2 | Erich5 | 4.548628443 | 2.370555783 |
| P83510 | Tnik | 4.488818999 | 2.044694 |
| Q925T6 | Grip1 | 4.398661396 | 2.449281532 |
| Q8BH53 | Cfap69 | 4.260222584 | 2.754255314 |
| Q8BX09 | Rbbp5 | 4.148832443 | 2.866897192 |
| Q69Z36 | Mex3b | 4.012105419 | 3.125645256 |
| Q8R0P4 | Aamdc | 3.50299079 | 2.569662402 |
| P11531 | Dmd | 3.340462764 | 2.047639381 |
| P33173 | Kif1a | 3.289080952 | 2.094402133 |
| Q8VHG2 | Amot | 2.956299627 | 2.584431334 |
| Q64512 | Ptpn13 | 2.913940724 | 2.123683944 |
| Q64524;Q8CGP0;Q9D2U9 | Q64524;Q8CGP0;Q9D2U9 | 2.784246105 | 2.487712064 |
| Q8BSK8 | Rps6kb1 | 2.76802983 | 2.794650002 |
| A0A7H0DN62 | OPG090 | 2.635808915 | 2.382084818 |
| Q3UIA2 | Arhgap17 | 2.35459134 | 3.634553343 |
| Q6PIC6;Q6PIE5 | Q6PIC6;Q6PIE5 | 2.30809825 | 2.166173317 |
| P63158 | Hmgb1 | 2.267498915 | 2.202224068 |
| Q9WVA4 | Tagln2 | 2.267385735 | 2.437611739 |
| Q02614 | Sap30bp | 2.179981657 | 2.655468012 |
| Q8BYJ6 | Tbc1d4 | 2.151775838 | 2.3115829 |
| Q9D920 | Borcs5 | 2.080360052 | 2.834316503 |
| Q60611 | Satb1 | 2.069458392 | 2.331138856 |
| Q80VP1 | Epn1 | 2.063512901 | 2.310430558 |
| Q69Z98 | Brsk2 | 2.031829728 | 2.504688943 |
| Q99L47 | St13 | 1.956024983 | 3.312162942 |
| Q9Z1E4 | Gys1 | 1.716655101 | 2.049445416 |
| Q9DBT3 | Ccdc97 | 1.577997756 | 2.209293514 |
| Q9QYY0 | Gab1 | 1.574890098 | 2.683201492 |
| Q8K1R7 | Nek9 | 1.502870392 | 2.171402685 |
| Q8K4Q0 | Rptor | 1.469667752 | 2.220388116 |
| Q8JZK9 | Hmgcs1 | 1.438298665 | 2.31784007 |
| Q60668 | Hnrnpd | 1.319304958 | 2.239462544 |
| P70265 | Pfkfb2 | 1.318494628 | 2.906411417 |
| Q3TXS7 | Psmd1 | 1.290974868 | 2.66975613 |
| Q5XJY5 | Arcn1 | 1.28234829 | 2.329539656 |
| P19426 | Nelfe | -1.01530559 | 2.421757818 |
| Q6NZR5 | Skic2 | -1.062592568 | 2.094402133 |
| Q5F2E8 | Taok1 | -1.067714793 | 2.123683944 |
| Q8R1A4 | Dock7 | -1.083078547 | 2.228103628 |
| P54227 | Stmn1 | -1.097803485 | 3.125645256 |
| G5E870 | Trip12 | -1.107282051 | 2.003124043 |
| Q5FWH2 | Unkl | -1.108960764 | 2.216285575 |
| Q99P69 | Nuf2 | -1.115569038 | 2.382084818 |
| Q9WU00 | Nrf1 | -1.125555106 | 2.45878218 |
| Q6PDQ2 | Chd4 | -1.133585788 | 2.129112415 |
| P63101 | Ywhaz | -1.142574831 | 2.341483885 |
| Q9EP97 | Senp3 | -1.154673098 | 2.101634545 |
| P24788 | Cdk11b | -1.166316732 | 2.633105775 |
| Q4JIM5 | Abl2 | -1.182114259 | 2.120831581 |
| Q9DAZ9 | Zfyve19 | -1.187141254 | 2.908658267 |
| A2AN08 | Ubr4 | -1.191717318 | 2.013323529 |
| Q80U72 | Scrib | -1.224721244 | 2.908658267 |
| Q8CFC7 | Clasrp | -1.229696448 | 2.31784007 |
| A2A5R2 | Arfgef2 | -1.240738219 | 2.215671286 |
| B1AZI6 | Thoc2 | -1.264926121 | 2.423419562 |
| Q08024 | Cbfb | -1.266896041 | 2.836680621 |
| Q61206 | Pafah1b2 | -1.270977758 | 2.204022132 |
| P53564 | Cux1 | -1.277552878 | 2.66975613 |
| P49117 | Nr2c2 | -1.281739143 | 2.66975613 |
| Q9WTK7 | Stk11 | -1.304523946 | 2.449281532 |
| Q99L90 | Mcrs1 | -1.311086114 | 2.422063203 |
| Q9ESU6 | Brd4 | -1.311764697 | 2.419004193 |
| Q8CDG3 | Vcpip1 | -1.321318483 | 2.422063203 |
| P08775 | Polr2a | -1.323722379 | 2.209293514 |
| P53996 | Cnbp | -1.32936962 | 2.402014711 |
| Q61937 | Npm1 | -1.349717994 | 2.159119331 |
| Q8BXK8 | Agap1 | -1.355511427 | 2.169692729 |
| Q9EPQ8 | Tcf20 | -1.356297009 | 2.245840037 |
| Q62188 | Dpysl3 | -1.362243949 | 2.134653847 |
| Q923B1 | Dbr1 | -1.365280025 | 2.880638009 |
| E9Q5C9 | Nolc1 | -1.370535235 | 2.401362163 |
| Q60575 | Kif1b | -1.373405111 | 2.385280897 |
| Q569Z6;Q8K019 | Q569Z6;Q8K019 | -1.407314896 | 2.077988818 |
| Q64092 | Tfe3 | -1.412251951 | 2.196917128 |
| Q61235 | Sntb2 | -1.423245587 | 2.719371343 |
| Q9DB00 | Gon4l | -1.423460085 | 2.450457162 |
| Q3THK7 | Gmps | -1.428789727 | 2.229003843 |
| A2AR02 | Ppig | -1.435542696 | 2.222510102 |
| Q6A065 | Cep170 | -1.437692584 | 2.288290441 |
| P18654 | Rps6ka3 | -1.439666731 | 2.66009481 |
| Q9D8Y0 | Efhd2 | -1.451182002 | 2.100070371 |
| P97305 | Nfatc3 | -1.452869701 | 2.570809194 |
| Q02053 | Uba1 | -1.456593445 | 2.589137733 |
| P23949;P23950 | P23949;P23950 | -1.468215008 | 2.503507207 |
| P09405 | Ncl | -1.473934412 | 2.754255314 |
| Q62418 | Dbnl | -1.474892292 | 2.107435535 |
| Q6NXH3 | Gpbp1 | -1.476374787 | 2.256746404 |
| B1AZP2 | Dlgap4 | -1.482260471 | 2.234439577 |
| Q9JIX8 | Acin1 | -1.490874934 | 2.793958437 |
| Q68FH0 | Pkp4 | -1.492677335 | 2.458655902 |
| Q8K1Y2 | Prkd3 | -1.493286128 | 2.138639356 |
| Q99JR8 | Smarcd2 | -1.51108735 | 2.564653124 |
| Q7TPM1 | Prrc2b | -1.556085627 | 2.66975613 |
| Q9CZS3 | Cep20 | -1.556458977 | 2.925004879 |
| Q80V24 | Vgll4 | -1.577580739 | 2.449281532 |
| Q8K4B0 | Mta1 | -1.579919249 | 2.127534484 |
| Q9JJ66 | Cdc20 | -1.580744549 | 2.400261522 |
| Q52KI8 | Srrm1 | -1.582559171 | 2.860685748 |
| Q80U58 | Pum2 | -1.588672648 | 2.421757818 |
| Q61214;Q9Z188 | Q61214;Q9Z188 | -1.589897005 | 3.478421489 |
| A2ABV5 | Med14 | -1.593920779 | 2.221098096 |
| Q6PER3 | Mapre3 | -1.596436763 | 2.220342972 |
| Q6PGH2 | Jpt2 | -1.597913196 | 2.141043037 |
| Q9CQY2 | Ramac | -1.601800702 | 2.171730395 |
| B2RY56 | Rbm25 | -1.60938798 | 2.109724375 |
| P09411 | Pgk1 | -1.612105508 | 2.471222381 |
| Q3U2S4 | Otud5 | -1.616373003 | 2.274399182 |
| Q8BZQ7 | Anapc2 | -1.644342899 | 2.753040552 |
| Q8BTI8 | Srrm2 | -1.649298291 | 2.300495474 |
| Q91XU0 | Wrnip1 | -1.650056263 | 2.202224068 |
| Q9DBR4 | Apbb2 | -1.666782151 | 2.468979685 |
| A2A432 | Cul4b | -1.668888469 | 2.050868963 |
| Q80TM6 | R3hdm2 | -1.694545026 | 2.487712064 |
| Q8BVE8 | Nsd2 | -1.719659568 | 2.3358728 |
| Q6KCD5 | Nipbl | -1.723509003 | 2.479019853 |
| Q8VDD5 | Myh9 | -1.763347522 | 2.208478541 |
| P62264 | Rps14 | -1.778790285 | 2.131163697 |
| P17751 | Tpi1 | -1.779468779 | 2.357319084 |
| Q61699 | Hsph1 | -1.788562917 | 2.138639356 |
| Q6NZF1 | Zc3h11a | -1.803474671 | 2.156835118 |
| Q58FA4 | E2f8 | -1.813704475 | 2.296983096 |
| O54967 | Tnk2 | -1.834698734 | 2.331138856 |
| A2A7S8 | Kiaa1522 | -1.857519264 | 2.232314605 |
| Q569Z6 | Thrap3 | -1.870386927 | 2.137771886 |
| P35601 | Rfc1 | -1.875594717 | 2.175065389 |
| Q91W92 | Cdc42ep1 | -1.877869816 | 2.044694 |
| Q8BJQ2 | Usp1 | -1.890254889 | 2.427523518 |
| Q61316 | Hspa4 | -1.933434772 | 2.589137733 |
| Q99LJ0 | Cttnbp2nl | -1.941456232 | 2.713336585 |
| Q60790 | Rasa3 | -1.964828338 | 2.299018706 |
| Q9JM52 | Mink1 | -2.028680869 | 2.66975613 |
| Q8BYR2 | Lats1 | -2.032941032 | 2.03708339 |
| P07607 | Tyms | -2.049231971 | 2.02077341 |
| Q6PFD9 | Nup98 | -2.089999312 | 2.442351494 |
| Q68EF0 | Rab3ip | -2.111274666 | 2.02077341 |
| Q8C8U0 | Ppfibp1 | -2.120251956 | 2.695773975 |
| P97434 | Mprip | -2.156128038 | 2.047639381 |
| Q8CIN4 | Pak2 | -2.188258699 | 3.81323893 |
| Q9WU42 | Ncor2 | -2.202130081 | 2.385280897 |
| O35492 | Clk3 | -2.247024174 | 2.405517051 |
| Q9JK23 | Psmg1 | -2.264696737 | 2.020085399 |
| P16858 | Gapdh | -2.279083548 | 2.218200604 |
| E9Q309 | Cep350 | -2.299398821 | 2.196917128 |
| P58058 | Nadk | -2.326731421 | 3.125645256 |
| Q9R1Z8 | Sorbs3 | -2.330821452 | 3.056534508 |
| Q8K2H3 | Fam13b | -2.341975793 | 2.379629323 |
| Q8C6G8 | Wdr26 | -2.359651531 | 2.146825289 |
| Q80UG5 | Septin9 | -2.374779425 | 3.879020042 |
| O55201 | Supt5h | -2.397725791 | 3.117083711 |
| Q80Z37 | Topors | -2.412892137 | 3.879020042 |
| Q04899 | Cdk18 | -2.415041501 | 2.422063203 |
| P35979 | Rpl12 | -2.415160279 | 2.382084818 |
| Q7TNE3 | Spag7 | -2.428735831 | 3.631044627 |
| O88667;P55041 | O88667;P55041 | -2.431673208 | 2.422063203 |
| P62137 | Ppp1ca | -2.442267761 | 3.202628219 |
| Q61329 | Zfhx3 | -2.456001553 | 2.66975613 |
| A2AAE1 | Bltp1 | -2.460868739 | 2.047982213 |
| Q8K3X4 | Irf2bpl | -2.462940373 | 2.596787818 |
| Q8BWW4 | Larp4 | -2.52059712 | 3.533059016 |
| Q9Z1D1 | Eif3g | -2.528535661 | 3.202628219 |
| P97310 | Mcm2 | -2.528613408 | 3.115075241 |
| Q9D8E6 | Rpl4 | -2.610857015 | 2.819485753 |
| Q80TG1 | Kansl1 | -2.612906632 | 2.171730395 |
| Q99MK9 | Rassf1 | -2.61730091 | 4.673185206 |
| O88573 | Aff1 | -2.618523336 | 2.032549429 |
| Q9EQM6 | Dgcr8 | -2.638126214 | 2.047289226 |
| Q8K327 | Champ1 | -2.64329088 | 2.570809194 |
| P17095 | Hmga1 | -2.660553176 | 2.07816416 |
| Q8R3P2 | Dtx2 | -2.685501827 | 2.007425467 |
| Q8BW94 | Dnah3 | -2.700950279 | 2.219485838 |
| Q6P9R4 | Arhgef18 | -2.713344159 | 2.080700432 |
| Q9JHU9 | Isyna1 | -2.737097972 | 2.2816518 |
| Q9CVD2 | Atxn3 | -2.742781236 | 2.793958437 |
| P39428 | Traf1 | -2.761159291 | 2.66975613 |
| Q9DBR0 | Akap8 | -2.769013637 | 2.466857064 |
| Q3TMW1 | Ccdc102a | -2.80720742 | 3.045560895 |
| Q9JHS4 | Clpx | -2.838292988 | 2.058605851 |
| F8VPU2 | Farp1 | -2.847034093 | 3.285947727 |
| Q80X90 | Flnb | -2.853140611 | 2.958806266 |
| P58871 | Tnks1bp1 | -2.859152173 | 2.24588311 |
| Q99KX1 | Mlf2 | -2.871749791 | 3.174581166 |
| P13808 | Slc4a2 | -2.878457226 | 2.354207866 |
| Q9DBR7 | Ppp1r12a | -2.886944797 | 2.131163697 |
| P20152 | Vim | -2.890570209 | 2.120831581 |
| Q69Z38 | Peak1 | -2.910673908 | 3.103165063 |
| P59222 | Scarf2 | -2.932387481 | 2.11624034 |
| Q4VBD9 | Gzf1 | -2.959112247 | 2.66009481 |
| Q6PIX5 | Rhbdf1 | -2.960489832 | 2.225383415 |
| Q8CEE0 | Cep57 | -2.974024193 | 3.202628219 |
| Q9D1C2 | Cby1 | -2.988160033 | 2.146825289 |
| Q9DBC7 | Prkar1a | -2.995193356 | 2.220388116 |
| Q6P1D7 | Slx4 | -2.998915663 | 2.215671286 |
| Q5ND34 | Wdr81 | -3.024150047 | 2.74785658 |
| O35381 | Anp32a | -3.033067081 | 2.185326073 |
| A2AGH6 | Med12 | -3.04710831 | 2.66975613 |
| Q9D030 | Twist2 | -3.053378542 | 2.31784007 |
| Q3UHX0 | Nol8 | -3.06068993 | 2.163421195 |
| Q9QVP9 | Ptk2b | -3.068624386 | 2.471222381 |
| Q8K4E0 | Alms1 | -3.073782585 | 2.715625435 |
| Q8BTM8 | Flna | -3.077373324 | 3.111043383 |
| Q8CCJ9 | Phf20l1 | -3.105207541 | 2.107008507 |
| O55023 | Impa1 | -3.12156428 | 2.66009481 |
| P97315 | Csrp1 | -3.123897288 | 3.618353804 |
| P58802 | Tbc1d10a | -3.135380575 | 3.419565228 |
| Q3TV65 | Mpnd | -3.141310176 | 2.068951052 |
| Q6PDH0 | Phldb1 | -3.159710598 | 2.713336585 |
| Q8C0T5 | Sipa1l1 | -3.180482121 | 2.932132558 |
| G5E829;Q9R0K7 | G5E829;Q9R0K7 | -3.181645043 | 2.333301996 |
| P41183 | Bcl6 | -3.182692923 | 2.118509547 |
| Q8C142 | Ldlrap1 | -3.196087642 | 3.534643934 |
| Q02819 | Nucb1 | -3.199232638 | 2.138639356 |
| Q9R0P5 | Dstn | -3.217177738 | 2.569662402 |
| P46062 | Sipa1 | -3.21935605 | 2.503507207 |
| Q3U2K0 | Fam193b | -3.22103274 | 2.395340014 |
| B2RQC6 | Cad | -3.229156335 | 2.92388003 |
| Q61127 | Nab2 | -3.234727408 | 3.261802713 |
| Q7TSC1 | Prrc2a | -3.265382497 | 2.077988818 |
| Q6PAL7 | Ahdc1 | -3.285354661 | 2.436983361 |
| Q3UKU1 | Ell2 | -3.285843719 | 2.220342972 |
| D3YXK2;Q80YR5 | D3YXK2;Q80YR5 | -3.301640886 | 3.634553343 |
| Q9JIH2 | Nup50 | -3.329956512 | 2.071286394 |
| Q6ZQB6 | Ppip5k2 | -3.334845352 | 2.043650727 |
| Q9QZN4 | Fbxo6 | -3.371882778 | 2.335313637 |
| Q8K0H5 | Taf10 | -3.375237973 | 2.927568233 |
| Q9CY64 | Blvra | -3.380032904 | 2.422063203 |
| Q9JII1 | Inpp5e | -3.380141587 | 2.355358443 |
| Q8CFE4 | Scyl2 | -3.380380564 | 2.43636786 |
| Q8CJ27 | Aspm | -3.388483785 | 2.66975613 |
| Q7TSZ8 | Nacc1 | -3.407758282 | 2.449281532 |
| Q9D7I8 | Fam83d | -3.411338217 | 2.438961756 |
| Q8BJ37 | Tdp1 | -3.425976927 | 2.142083822 |
| Q6P9Q6 | Fkbp15 | -3.430852163 | 2.455410988 |
| Q3UM83 | Klrg2 | -3.441760192 | 2.767022219 |
| O54864 | Suv39h1 | -3.446209345 | 3.631044627 |
| Q8R2M2 | Dnttip2 | -3.447375275 | 2.719834101 |
| Q9QXS1 | Plec | -3.498896361 | 4.636544233 |
| Q7TN31 | Aggf1 | -3.499229328 | 2.906411417 |
| Q9Z1S8 | Gab2 | -3.505776067 | 2.480199952 |
| Q8BGT0 | Ostm1 | -3.508149489 | 2.128566481 |
| Q5EBG6 | Hspb6 | -3.514383875 | 2.161742832 |
| P59178 | L3mbtl2 | -3.517773203 | 3.192702694 |
| Q61321 | Six4 | -3.519892811 | 2.31784007 |
| Q0VEE6 | Znf800 | -3.530286921 | 2.66975613 |
| Q80X82 | Sympk | -3.53330717 | 2.020579272 |
| Q91VR2 | Atp5f1c | -3.573200748 | 2.006560264 |
| B1AXD8 | Akirin2 | -3.57915912 | 2.970039085 |
| Q6RT24 | Cenpe | -3.579355532 | 2.011307451 |
| P97494 | Gclc | -3.588040687 | 2.45878218 |
| Q9D6Z1 | Nop56 | -3.59145611 | 2.819052836 |
| Q8BYH8 | Chd9 | -3.593968314 | 2.275090062 |
| O09039 | Sh2b3 | -3.617713448 | 2.564653124 |
| Q6EDY6 | Carmil1 | -3.61790633 | 3.81323893 |
| Q148V8 | Fam83h | -3.631616474 | 3.634553343 |
| Q62077 | Plcg1 | -3.63196874 | 2.290707121 |
| Q8C0E3 | Trim47 | -3.652586935 | 2.120831581 |
| Q9CS00 | Cactin | -3.659341816 | 2.66975613 |
| Q8C1S0 | Med19 | -3.66880177 | 2.719371343 |
| O70472 | Tmem131 | -3.680762577 | 2.331138856 |
| Q8C6B2 | Rtkn | -3.686354302 | 2.3358728 |
| B9EJ80 | Pdzd8 | -3.704120255 | 2.381110338 |
| Q8BIG4 | Fbxo28 | -3.741942662 | 3.117083711 |
| Q9ERV1 | Mkrn2 | -3.768509167 | 2.31784007 |
| Q62388 | Atm | -3.769789505 | 2.194359831 |
| Q3UFY0 | Rrp36 | -3.770877937 | 2.401362163 |
| P97329 | Kif20a | -3.779550671 | 2.463270389 |
| Q9D8T7 | Slirp | -3.781093518 | 2.220342972 |
| Q9D8S3 | Arfgap3 | -3.803315787 | 2.047639381 |
| Q6NZN1 | Pprc1 | -3.81051476 | 2.500143671 |
| P70297 | Stam | -3.816720381 | 3.125645256 |
| P48678 | Lmna | -3.825091512 | 2.908658267 |
| Q02085 | Snai1 | -3.832641812 | 2.927568233 |
| A0A338P6K9 | Qser1 | -3.872657348 | 2.719834101 |
| P59110 | Senp1 | -3.880351233 | 2.060190984 |
| Q8VC56 | Rnf8 | -3.892768295 | 2.66975613 |
| Q64012 | Raly | -3.909303399 | 3.064371483 |
| Q8C5H8 | Nadk2 | -3.930006996 | 2.340877628 |
| Q91Z38 | Ttc1 | -3.934584986 | 2.234439577 |
| Q9CZ09 | Mettl18 | -3.936340033 | 2.422063203 |
| O70475 | Ugdh | -3.940338113 | 3.157539749 |
| P97929 | Brca2 | -3.95422958 | 2.142083822 |
| Q9Z0H8 | Clip2 | -3.954754818 | 3.879020042 |
| P14733 | Lmnb1 | -3.955362648 | 3.348164707 |
| Q9Z206 | Net1 | -3.978986895 | 2.157608546 |
| Q8R1M0 | Hmces | -3.979907958 | 2.225383415 |
| P30285 | Cdk4 | -3.997562892 | 2.66975613 |
| Q99K43 | Prc1 | -4.018671145 | 2.149927762 |
| Q9CT10 | Ranbp3 | -4.034279972 | 2.283036797 |
| F7BJB9 | Morc3 | -4.037913623 | 3.879020042 |
| Q60749 | Khdrbs1 | -4.04529571 | 2.080249241 |
| Q9Z1Y4 | Trip6 | -4.06303423 | 2.225383415 |
| Q9JKS5 | Habp4 | -4.063415289 | 2.736708897 |
| Q8K3G5 | Vrk3 | -4.079263377 | 3.103165063 |
| Q5RJH6 | Smg7 | -4.101089001 | 2.902115475 |
| Q9CYH6 | Rrs1 | -4.102031889 | 2.119464109 |
| Q6NZP2 | Gpbp1l1 | -4.121729654 | 2.357319084 |
| Q8BG81 | Poldip3 | -4.13309093 | 2.149298796 |
| Q6NZL6 | Tonsl | -4.133266605 | 2.62502513 |
| Q91X58 | Zfand2b | -4.141782149 | 2.01799435 |
| Q9DBN4 | P33monox | -4.142813652 | 2.211855119 |
| Q6PGG6 | Gnl3l | -4.142824547 | 2.118509547 |
| Q9D1M0 | Sec13 | -4.153610879 | 2.190703399 |
| Q8BKX6 | Smg1 | -4.182784287 | 2.719834101 |
| Q8BUH8 | Senp7 | -4.183724816 | 2.065151367 |
| Q6PIU9 | YJ005_MOUSE | -4.193298473 | 2.318889651 |
| Q8C9S4 | Ccdc186 | -4.196733041 | 2.13150652 |
| Q4LDD4 | Arap1 | -4.217136136 | 2.121893903 |
| Q923J1 | Trpm7 | -4.23320792 | 2.169692729 |
| O08609 | Mlx | -4.236588313 | 2.274399182 |
| Q9ES34 | Ube3b | -4.250007936 | 3.133566514 |
| P55937 | Golga3 | -4.253040304 | 2.068261005 |
| Q8K363 | Ddx18 | -4.266819553 | 2.649845701 |
| B1AY10 | Nfx1 | -4.268569431 | 2.181555304 |
| P70195 | Psmb7 | -4.284260727 | 2.540662548 |
| Q8K2J7 | Rell1 | -4.285308731 | 2.213297566 |
| Q8R010 | Aimp2 | -4.291103712 | 2.75727589 |
| Q8BGC0 | Htatsf1 | -4.293437243 | 2.925004879 |
| Q9CZJ2 | Hspa12b | -4.311854476 | 2.223259033 |
| Q8C1D8 | Iws1 | -4.32327537 | 2.866897192 |
| Q9DCI3 | Stard3nl | -4.327496976 | 2.393907308 |
| P26687 | Twist1 | -4.330661202 | 2.123683944 |
| Q9D9Z5 | Dda1 | -4.336049093 | 2.196917128 |
| O70551 | Srpk1 | -4.341667055 | 3.302865581 |
| Q8K202 | Polr1e | -4.361151157 | 2.396496929 |
| Q8VHX6 | Flnc | -4.376227097 | 2.15743443 |
| Q99K85 | Psat1 | -4.378597738 | 2.225383415 |
| Q8CIB6 | Tmem230 | -4.392173151 | 2.109724375 |
| Q9Z2S7 | Tsc22d3 | -4.419772809 | 2.042236602 |
| Q61037 | Tsc2 | -4.423821389 | 2.355917173 |
| Q6PDZ2 | Taf1c | -4.424082928 | 2.478803522 |
| Q61083 | Map3k2 | -4.439152084 | 2.091610845 |
| Q9JLY0 | Socs6 | -4.457766982 | 2.113840169 |
| P63154 | Crnkl1 | -4.472256186 | 2.449281532 |
| Q924H2 | Med15 | -4.473271312 | 3.370025064 |
| O08796 | Eef2k | -4.475730657 | 2.325168992 |
| Q69ZR9 | Tasor | -4.477501944 | 2.480199952 |
| Q64124 | Scx | -4.485291965 | 2.514195222 |
| Q6ZQF0 | Topbp1 | -4.493637966 | 2.115416933 |
| Q8K039 | K1143_MOUSE | -4.498821399 | 2.204022132 |
| Q8C052 | Map1s | -4.510394919 | 2.720057627 |
| P35980 | Rpl18 | -4.517100498 | 2.514195222 |
| Q8C4M7 | Cenpu | -4.528431729 | 2.202224068 |
| Q921F4 | Hnrnpll | -4.550907916 | 2.821938098 |
| Q9DBY8 | Nvl | -4.556916882 | 3.206969243 |
| Q80U22 | Rusc2 | -4.56062026 | 2.573993279 |
| Q9WTI7 | Myo1c | -4.562916287 | 2.483584572 |
| Q68ED7 | Crtc1 | -4.564868505 | 2.359452799 |
| E9Q5G3 | Kif23 | -4.567847855 | 3.198552061 |
| Q9CZX7 | Pip4p2 | -4.576367066 | 2.131163697 |
| Q6P1G0 | Heatr6 | -4.583674368 | 3.117083711 |
| Q8K268 | Abcf3 | -4.584284146 | 2.421757818 |
| Q2TBA3 | Malt1 | -4.603057103 | 2.866897192 |
| Q8CIL4 | CA131_MOUSE | -4.609238278 | 3.631044627 |
| Q62419 | Sh3gl1 | -4.610065909 | 2.223411881 |
| Q9R207 | Nbn | -4.623048227 | 2.540662548 |
| O35954 | Pitpnm1 | -4.624963348 | 2.047289226 |
| Q8BHL8 | Psmf1 | -4.627639694 | 2.131163697 |
| Q571H0 | Urb1 | -4.62818431 | 2.138639356 |
| Q9DBN9 | Ddx59 | -4.64624286 | 2.02632166 |
| E9Q4N7 | Arid1b | -4.652055394 | 2.045923218 |
| Q8K3K8 | Optn | -4.652476771 | 2.231292805 |
| Q80TQ5 | Plekhm2 | -4.655249102 | 2.765981717 |
| Q9D5R2 | Wdr20 | -4.655607295 | 2.60283957 |
| Q9JL61 | Rfx5 | -4.660339299 | 2.449281532 |
| Q80TL7 | Mon2 | -4.660899193 | 2.906028522 |
| Q9ESX5 | Dkc1 | -4.669641311 | 2.540662548 |
| Q9CRA8 | Exosc5 | -4.676037851 | 2.146825289 |
| Q80WJ7 | Mtdh | -4.678585572 | 2.220388116 |
| P30658 | Cbx2 | -4.67953635 | 2.220342972 |
| Q91V83 | Tti1 | -4.683576585 | 2.574109442 |
| Q91VW5 | Golga4 | -4.703671637 | 2.570435245 |
| Q70FJ1 | Akap9 | -4.712172695 | 2.540662548 |
| Q91X51 | Gorasp1 | -4.718458882 | 2.299713585 |
| Q9WTV7 | Rlim | -4.724331154 | 3.115075241 |
| Q9Z108 | Stau1 | -4.729955256 | 2.138639356 |
| Q5SSH7 | Zzef1 | -4.732867371 | 2.109724375 |
| P62075 | Timm13 | -4.739668845 | 2.09140918 |
| Q8VED8 | Mtfr2 | -4.751270976 | 2.091610845 |
| Q8BQ30 | Ppp1r18 | -4.760941353 | 2.171402685 |
| P48725 | Pcnt | -4.771188555 | 3.481155404 |
| B1AVZ0 | Uprt | -4.779283928 | 2.518516279 |
| Q8BFY7 | Pimreg | -4.78161697 | 2.204022132 |
| Q6P9Q4 | Fhod1 | -4.785642282 | 2.471222381 |
| Q9Z1Q5 | Clic1 | -4.788202115 | 2.125238164 |
| Q61382 | Traf4 | -4.79080792 | 2.113840169 |
| Q9DCL8 | Ppp1r2 | -4.794870652 | 2.232726645 |
| Q9CYC5 | Dsn1 | -4.803043774 | 2.458655902 |
| Q9D0L7 | Armc10 | -4.815818115 | 3.060637864 |
| Q9CQI9 | Med30 | -4.822670039 | 2.003069305 |
| Q91VS8 | Farp2 | -4.828733766 | 2.708257645 |
| P27546 | Map4 | -4.838618025 | 2.812313755 |
| Q8VD65 | Pik3r4 | -4.84482372 | 2.117572798 |
| Q8R502 | Lrrc8c | -4.855204813 | 2.114375881 |
| Q8K411 | Pitrm1 | -4.857863132 | 2.209293514 |
| Q68ED3 | Tent4b | -4.875053314 | 2.579135505 |
| Q5SSZ5 | Tns3 | -4.87699944 | 3.061289501 |
| P10605 | Ctsb | -4.887643341 | 2.131163697 |
| Q60953 | Pml | -4.891929041 | 3.461126922 |
| Q6PGL7 | Washc2 | -4.893432363 | 3.856285928 |
| P54279 | Pms2 | -4.900712661 | 2.051523539 |
| Q6ZPG2 | Wdr90 | -4.906719892 | 2.278164902 |
| Q8R3F9 | Tut1 | -4.915342276 | 2.385280897 |
| Q9JLG8 | Capn15 | -4.916405076 | 2.142083822 |
| Q8BT14 | Cnot4 | -4.917082809 | 2.179660064 |
| Q8C115 | Plekhh2 | -4.917156787 | 2.762292398 |
| Q8K368 | Fanci | -4.930902055 | 2.171052822 |
| Q8BMK1 | Mettl2 | -4.981001675 | 2.589137733 |
| Q8BLK9 | Rps6kc1 | -4.987920134 | 2.542455775 |
| Q9D2L9 | Fam111a | -4.998158946 | 2.428963469 |
| Q5DU05 | Cep164 | -5.01171543 | 2.03565185 |
| Q9JI57 | Gtf2ird1 | -5.011969416 | 3.46209774 |
| Q59J78 | Ndufaf2 | -5.022116875 | 3.202628219 |
| A2A699 | Fam171a2 | -5.023310536 | 2.220342972 |
| Q62406 | Irak1 | -5.030571594 | 2.077988818 |
| Q3UQU0 | Brd9 | -5.032581991 | 2.045923218 |
| Q3UMQ8 | Naf1 | -5.038107518 | 2.382084818 |
| O35130 | Emg1 | -5.041118227 | 2.044694 |
| Q99JN2 | Klhl22 | -5.04116273 | 2.025608437 |
| P16254 | Srp14 | -5.046647946 | 2.091610845 |
| Q5XF89 | Atp13a3 | -5.04883213 | 2.044694 |
| Q8CES0 | Naa30 | -5.049771613 | 2.097427715 |
| Q9CR25 | Dph2 | -5.073781622 | 2.922992182 |
| Q3UD01 | Atxn7l3b | -5.087489018 | 2.086516969 |
| P39053 | Dnm1 | -5.103607876 | 2.13150652 |
| Q62433 | Ndrg1 | -5.107661664 | 3.775283316 |
| Q99P31 | Hspbp1 | -5.120236365 | 2.142083822 |
| Q8CB62 | Cntrob | -5.137032962 | 2.114912392 |
| Q8K2Z4 | Ncapd2 | -5.137651853 | 2.419524971 |
| P28658 | Atxn10 | -5.145102185 | 2.220342972 |
| Q8BJU0 | Sgta | -5.17448084 | 2.19957522 |
| Q8K389 | Cdk5rap2 | -5.184185643 | 2.589137733 |
| Q9QWF0 | Chaf1a | -5.18843927 | 2.514195222 |
| Q91WE1 | Snx15 | -5.18873849 | 2.908658267 |
| Q60738 | Slc30a1 | -5.205425245 | 2.17647695 |
| Q3UIR3 | Dtx3l | -5.206157809 | 2.560810135 |
| Q91ZW3 | Smarca5 | -5.207730595 | 2.463270389 |
| O08715 | Akap1 | -5.215721959 | 2.847235318 |
| Q9JJZ4 | Ube2j1 | -5.217706772 | 3.420600908 |
| P70210 | Tead3 | -5.218224247 | 3.157539749 |
| Q8VI75 | Ipo4 | -5.232146033 | 2.438961756 |
| Q68FE2 | Atg9a | -5.232288497 | 2.437505107 |
| Q5FWK3 | Arhgap1 | -5.233922998 | 3.879020042 |
| Q8VIG1 | Rest | -5.241262875 | 2.419524971 |
| E9Q555 | Rnf213 | -5.241811828 | 2.209293514 |
| Q91WM3 | Rrp9 | -5.243062188 | 2.556141213 |
| Q9JK91 | Mlh1 | -5.250229944 | 2.131097669 |
| Q91Z96 | Bmp2k | -5.251515015 | 2.471222381 |
| B1AX39 | Zcchc7 | -5.264387009 | 2.753040552 |
| Q9Z2Q2 | Knop1 | -5.265830204 | 2.540662548 |
| Q9CSP9 | Ttc14 | -5.272964246 | 2.099154021 |
| Q60596 | Xrcc1 | -5.279289113 | 2.001742648 |
| Q3UMT1 | Ppp1r12c | -5.28435313 | 2.202224068 |
| Q8BTT6 | Utp25 | -5.285420252 | 2.471222381 |
| Q921C3 | Brwd1 | -5.312334061 | 2.461437852 |
| Q64337 | Sqstm1 | -5.316466867 | 3.370025064 |
| P70372 | Elavl1 | -5.331088713 | 2.063752082 |
| P35831 | Ptpn12 | -5.341530049 | 2.698786852 |
| Q99LT0 | Dpy30 | -5.350910953 | 2.713336585 |
| Q8JZP9 | Gas2l1 | -5.357237431 | 2.218200604 |
| Q8C1Z8 | Trmt10a | -5.375058061 | 2.077988818 |
| Q9DBU3 | Riok3 | -5.38133904 | 2.622008844 |
| Q91XC8 | Dap | -5.381398187 | 2.308715717 |
| Q6NXK2 | Znf532 | -5.382129337 | 2.564653124 |
| Q91YE3 | Egln1 | -5.384528474 | 2.185326073 |
| Q8CI11 | Gnl3 | -5.418354921 | 2.754255314 |
| D2EAC2 | Zbed6 | -5.438102392 | 2.041797925 |
| O35639 | Anxa3 | -5.441205544 | 2.348073429 |
| Q9D8S9 | Bola1 | -5.445888816 | 2.307576297 |
| P70671 | Irf3 | -5.45181922 | 2.753040552 |
| Q3TZX8 | Nol9 | -5.461228846 | 2.818566102 |
| Q8VBT9 | Aspscr1 | -5.467999372 | 2.583347163 |
| P19246 | Nefh | -5.468587084 | 2.174225408 |
| Q80YR6 | Rbbp8 | -5.47120512 | 2.819485753 |
| Q76KJ5 | Polr1g | -5.478169162 | 2.630790556 |
| Q9QZM0 | Ubqln2 | -5.479015494 | 2.113840169 |
| Q60695 | Rgl1 | -5.48159037 | 2.31784007 |
| P62631 | Eef1a2 | -5.484016041 | 2.225383415 |
| Q9D4V4 | Taf1d | -5.487103334 | 2.630790556 |
| Q6NZC7 | Sec23ip | -5.490005775 | 3.6458806 |
| Q8C5Q4 | Grsf1 | -5.504206042 | 2.463270389 |
| Q8VDF2 | Uhrf1 | -5.53043282 | 2.630790556 |
| Q91WG4 | Elp2 | -5.531207864 | 2.026215266 |
| P63013 | Prrx1 | -5.53275751 | 2.503507207 |
| Q9WV89 | Stxbp4 | -5.547068554 | 2.438961756 |
| A2CG63 | Arid4b | -5.561114336 | 2.535375088 |
| Q9Z2H5 | Epb41l1 | -5.575256464 | 2.202224068 |
| Q2TBE6 | Pi4k2a | -5.614223095 | 2.565641282 |
| Q921T2 | Tor1aip1 | -5.618883657 | 3.008200367 |
| Q9D620 | Rab11fip1 | -5.621175686 | 2.131964832 |
| Q64267 | Xpa | -5.624882123 | 2.093188935 |
| O89084 | Pde4a | -5.626189315 | 2.185326073 |
| Q8BLB7 | L3mbtl3 | -5.628772914 | 2.754255314 |
| O35969 | Gamt | -5.637297187 | 2.063752082 |
| Q6P2L6 | Nsd3 | -5.640061229 | 2.74785658 |
| Q9ER72 | Cars1 | -5.664865114 | 2.047289226 |
| Q9R1T2 | Sae1 | -5.66725964 | 2.706714619 |
| Q80Y44 | Ddx10 | -5.668421468 | 2.209293514 |
| Q6P5D4 | Cep135 | -5.67081665 | 2.344534756 |
| Q5SWW4 | Med13 | -5.68380709 | 2.138639356 |
| Q9JIS8 | Slc12a4 | -5.685694173 | 2.748463369 |
| Q64364 | Cdkn2a | -5.688718488 | 2.267419587 |
| O55098 | Stk10 | -5.699760913 | 2.220388116 |
| Q8VCI5 | Pex19 | -5.705749048 | 2.288290441 |
| Q5I1X5 | Ppp1r13l | -5.706903946 | 2.331333146 |
| P59017 | Bcl2l13 | -5.724917018 | 2.66975613 |
| Q60848 | Hells | -5.734653953 | 2.380039316 |
| Q8R184 | Fam110a | -5.741123757 | 2.958806266 |
| Q80V94 | Ap4e1 | -5.744190181 | 2.209293514 |
| Q8K1E6 | Alkbh3 | -5.749275367 | 2.634526369 |
| Q8C551 | Rad51ap1 | -5.751713131 | 2.049445416 |
| Q61464 | Znf638 | -5.763013675 | 3.879020042 |
| Q91VH2 | Snx9 | -5.765775911 | 3.302865581 |
| Q9JJ11 | Tacc3 | -5.766047064 | 2.310430558 |
| Q9CQJ6 | Denr | -5.768146538 | 3.261802713 |
| Q6ZPR6 | Ibtk | -5.774670369 | 2.220388116 |
| Q80T85 | Dcaf5 | -5.779633235 | 2.196917128 |
| Q8CH72 | Trim32 | -5.785250374 | 2.097005166 |
| Q9CQJ7 | Pttg1 | -5.785327714 | 2.422063203 |
| Q9QXE2 | Poll | -5.792097669 | 3.879020042 |
| Q8CBY1 | Samd4a | -5.792490687 | 2.211855119 |
| O35134 | Polr1a | -5.809283139 | 2.310130185 |
| Q91W96 | Anapc4 | -5.811967657 | 2.131163697 |
| Q3UMU9 | Hdgfl2 | -5.832554843 | 2.299713585 |
| O08856 | Ell | -5.838180747 | 3.478421489 |
| Q80Y17 | Llgl1 | -5.843025001 | 2.121893903 |
| Q8VDP4 | Ccar2 | -5.854445924 | 2.897995547 |
| Q3U7R1 | Esyt1 | -5.876088948 | 2.574109442 |
| P70255 | Nfic | -5.880757945 | 2.835532102 |
| P97450 | Atp5pf | -5.8816774 | 2.131163697 |
| P48967 | Cdc25c | -5.889194035 | 2.003058237 |
| Q6NZM9;Q8BIY3;Q99N13;Q9Z2V6 | Q6NZM9;Q8BIY3;Q99N13;Q9Z2V6 | -5.907026651 | 2.211682414 |
| O88622 | Parg | -5.91537361 | 2.185326073 |
| O88852 | Tspyl1 | -5.922955963 | 2.382084818 |
| P35546 | Ret | -5.934859581 | 2.471222381 |
| Q62210 | Birc2 | -5.936422078 | 2.396496929 |
| Q921C5 | Bicd2 | -5.942027409 | 2.296424628 |
| Q9ESV0 | Ddx24 | -5.946972846 | 2.878688127 |
| Q69ZH9 | Arhgap23 | -5.950000862 | 2.208805225 |
| O70326 | Grem1 | -5.954200023 | 2.719834101 |
| Q08943 | Ssrp1 | -5.959422138 | 2.867671206 |
| P13864 | Dnmt1 | -5.967405135 | 2.31784007 |
| Q9D0E3 | Lysmd1 | -5.97157515 | 2.197601341 |
| Q62523 | Zyx | -5.97179646 | 3.192702694 |
| Q9JK92 | Hspb8 | -5.973389956 | 2.109724375 |
| Q6KAU8 | Atg16l2 | -5.980204149 | 2.866897192 |
| Q62136 | Ptpn21 | -5.983878595 | 2.211855119 |
| Q9Z204 | Hnrnpc | -5.988060455 | 2.142083822 |
| Q3ULW8 | Parp3 | -5.988401817 | 2.92388003 |
| O35613 | Daxx | -6.021592702 | 2.366889277 |
| P28798 | Grn | -6.039435935 | 2.925004879 |
| Q9DC28 | Csnk1d | -6.048620283 | 2.449281532 |
| Q7TME2 | Spag5 | -6.06116993 | 3.431956117 |
| P15920 | Atp6v0a2 | -6.070238907 | 2.091610845 |
| Q8BX17 | Gemin5 | -6.074387511 | 3.117083711 |
| Q3TWF6 | Wdr70 | -6.075858232 | 2.754255314 |
| Q80U93 | Nup214 | -6.084346687 | 2.119464109 |
| A2AHC3 | Camsap1 | -6.102031509 | 2.715625435 |
| Q9D7S7 | Rpl22l1 | -6.105241568 | 2.113840169 |
| P62983 | Rps27a | -6.112557429 | 3.192702694 |
| O08784 | Tcof1 | -6.127355885 | 3.418197535 |
| P16045 | Lgals1 | -6.130684061 | 2.044694 |
| O88587 | Comt | -6.131885108 | 3.813654957 |
| O70405 | Ulk1 | -6.132357549 | 2.380039316 |
| P30415 | Nktr | -6.134684649 | 2.290707121 |
| Q9D5D8 | Cdyl2 | -6.152226944 | 2.172608458 |
| Q9WUH1 | Tmem115 | -6.155479461 | 2.471222381 |
| O88685 | Psmc3 | -6.162961455 | 3.418197535 |
| Q8BGF7 | Pan2 | -6.163919399 | 2.161742832 |
| Q8BH93 | Mapk1ip1l | -6.175438091 | 2.210730568 |
| Q9Z1S0 | Bub1b | -6.179344659 | 2.906411417 |
| Q6PCQ0 | Iqce | -6.191495996 | 2.074043187 |
| Q60710 | Samhd1 | -6.193327313 | 3.610088144 |
| B2RY04 | Dock5 | -6.193415971 | 2.355358443 |
| Q9Z1Z2 | Strap | -6.195091275 | 2.256975068 |
| P49025 | Cit | -6.198334101 | 3.101715613 |
| P51125 | Cast | -6.202193186 | 2.719834101 |
| Q9CR16 | Ppid | -6.217543597 | 2.514195222 |
| Q61602 | Gli3 | -6.220553206 | 2.119464109 |
| Q0VGB7 | Ppp4r2 | -6.245857015 | 2.468979685 |
| Q6P5E8 | Dgkq | -6.248234617 | 2.263924306 |
| Q99P72 | Rtn4 | -6.254699449 | 4.347089841 |
| B7ZNG4 | Troap | -6.267888644 | 2.229230713 |
| Q8BHG1 | Nrdc | -6.270997715 | 2.179660064 |
| Q80U49 | Cep170b | -6.27320236 | 3.075617369 |
| P21619 | Lmnb2 | -6.278238201 | 2.463270389 |
| Q8BH48 | Ubap1 | -6.278609493 | 3.060637864 |
| Q5SW19 | Cluh | -6.279473804 | 2.31784007 |
| P55200 | Kmt2a | -6.290946861 | 2.128566481 |
| P18653 | Rps6ka1 | -6.3021998 | 2.87023758 |
| P97473 | Tarbp2 | -6.302934307 | 2.432072408 |
| P17183 | Eno2 | -6.303053741 | 2.565641833 |
| Q8CGB6 | Tns2 | -6.303144083 | 3.255122224 |
| Q811L6 | Mast4 | -6.317075598 | 2.118878566 |
| Q3TN34 | Micall2 | -6.323364536 | 2.395340014 |
| E9Q7D5 | Arhgef5 | -6.324671857 | 2.665999609 |
| Q61103 | Dpf2 | -6.356108632 | 2.275090062 |
| Q9CPU0 | Glo1 | -6.370194249 | 2.73514213 |
| Q61749 | Eif2b4 | -6.374812107 | 2.835532102 |
| Q8BVY0 | Rsl1d1 | -6.391423152 | 2.363578906 |
| Q8R015 | Bloc1s5 | -6.392438723 | 2.573993279 |
| Q56A08 | Gpkow | -6.400824511 | 3.879020042 |
| Q5U5Q9 | Uimc1 | -6.40259858 | 3.155465047 |
| P47941 | Crkl | -6.410953913 | 2.146825289 |
| Q8CDJ8 | Ston1 | -6.412142219 | 2.099154021 |
| F8VPZ9 | Bicra | -6.424382347 | 3.046837233 |
| Q8N9S3 | Ahsa2 | -6.433767706 | 2.203647259 |
| Q05CL8 | Larp7 | -6.435094891 | 2.416033462 |
| P20065 | Tmsb4x | -6.440165337 | 2.044459414 |
| Q64727 | Vcl | -6.441199109 | 2.043650727 |
| Q3UDW8 | Hgsnat | -6.451659792 | 2.209293514 |
| Q80XP8 | Fam76b | -6.453798982 | 2.107435535 |
| P97825 | Jpt1 | -6.458754394 | 3.022087211 |
| P39689 | Cdkn1a | -6.462357254 | 2.17647695 |
| Q6P1H6 | Ankle2 | -6.463921991 | 3.81323893 |
| Q8R317 | Ubqln1 | -6.468677952 | 2.66009481 |
| Q99MY8 | Ash1l | -6.479009409 | 2.66975613 |
| Q8C4J7 | Tbl3 | -6.487256816 | 2.719834101 |
| E1U8D0 | Soga1 | -6.487356436 | 2.471222381 |
| A2AG58 | Bclaf3 | -6.487899605 | 2.395340014 |
| Q6PAM1 | Txlna | -6.500692812 | 4.10463829 |
| Q8VC70 | Rbms2 | -6.504687453 | 2.484502856 |
| Q9CWY3 | Setd6 | -6.507528773 | 2.060190984 |
| Q99KK9 | Hars2 | -6.514253043 | 2.045923218 |
| Q9JLB0 | Pals2 | -6.516298629 | 2.31784007 |
| Q3UPF5 | Zc3hav1 | -6.523389744 | 5.008539953 |
| Q8BMQ3 | Bnc2 | -6.53542439 | 2.400261522 |
| Q9ESL4 | Map3k20 | -6.536203689 | 3.618353804 |
| Q3UMF0 | Cobll1 | -6.549909935 | 2.449281532 |
| Q91VU7 | Pus7 | -6.550318325 | 2.231520711 |
| Q8BMA5 | Npat | -6.560904317 | 2.044459414 |
| P70193 | Lrig1 | -6.563727167 | 3.044449054 |
| Q8CHP6 | Phc3 | -6.563977821 | 3.590913466 |
| Q76N33 | Stambpl1 | -6.565825849 | 3.111043383 |
| Q8BRM2 | Gorab | -6.565832739 | 2.432072408 |
| Q8CCH7 | Zfpm2 | -6.568381942 | 2.3115829 |
| Q8C0P0 | Mastl | -6.572473867 | 2.275090062 |
| P0C7T6 | Atxn1l | -6.575563006 | 2.142083822 |
| Q9WUF3 | Casp8ap2 | -6.58627671 | 2.202053533 |
| Q8BU85 | Msrb3 | -6.587779852 | 2.381110338 |
| E9PV82 | Fam53a | -6.601255392 | 2.13150652 |
| P10107 | Anxa1 | -6.606611138 | 2.060958987 |
| Q9D1J1 | Necap2 | -6.616127011 | 2.401855283 |
| P12367 | Prkar2a | -6.617118057 | 2.958806266 |
| Q9WUK4 | Rfc2 | -6.617649103 | 2.546288776 |
| Q8BLH7 | Hirip3 | -6.636760717 | 2.131180476 |
| Q3U3T8 | Wdr62 | -6.673730107 | 2.564653124 |
| Q8C6B9 | Rps19bp1 | -6.679993436 | 2.834316503 |
| Q9CYG7 | Tomm34 | -6.711114715 | 2.034640334 |
| Q99L48 | Nmd3 | -6.71319531 | 3.202628219 |
| Q9CPV5 | Pmf1 | -6.727851035 | 2.685864616 |
| Q3TYA6 | Mphosph8 | -6.728038932 | 2.642366431 |
| Q99LL5 | Pwp1 | -6.728718305 | 3.356622611 |
| Q3THG9 | Aarsd1 | -6.732374189 | 2.463270389 |
| Q9EQN3 | Tsc22d4 | -6.735911483 | 2.835532102 |
| Q9JIK5 | Ddx21 | -6.750979822 | 2.118878566 |
| Q14B71 | Cdca2 | -6.755996337 | 2.404009495 |
| Q3TFK5 | Gpatch4 | -6.760436982 | 2.290707121 |
| Q9CPY3 | Cdca5 | -6.763420626 | 3.157539749 |
| Q64707 | Zrsr1 | -6.768697065 | 2.159119331 |
| P46935 | Nedd4 | -6.783466917 | 2.209293514 |
| P37913 | Lig1 | -6.788881337 | 3.907318488 |
| Q921I2 | Klhdc4 | -6.793656154 | 2.123683944 |
| Q7TPV4 | Mybbp1a | -6.820209476 | 4.673185206 |
| P48754 | Brca1 | -6.82125093 | 2.41223108 |
| Q62417 | Sorbs1 | -6.824310544 | 2.220342972 |
| Q6NXN1 | Szrd1 | -6.840936796 | 2.020085399 |
| Q9CXL3 | CG050_MOUSE | -6.842990859 | 2.333301996 |
| Q3UC65 | Rsrp1 | -6.854857409 | 2.329539656 |
| Q9CPY7 | Lap3 | -6.862361008 | 2.07816416 |
| Q8BG48 | Stk17b | -6.863939904 | 2.514195222 |
| Q8C0I1 | Agps | -6.869093264 | 2.385280897 |
| A2AI08 | Tprn | -6.87181453 | 3.202628219 |
| Q8JZQ9 | Eif3b | -6.873374492 | 3.634553343 |
| Q9CZ42 | Naxd | -6.881678669 | 3.584127609 |
| Q61216 | Mre11 | -6.883699788 | 2.33466931 |
| P63073 | Eif4e | -6.885262499 | 2.633105775 |
| Q4V9W2 | Srek1ip1 | -6.891961621 | 2.047289226 |
| F8VPQ2 | Arid4a | -6.892833796 | 2.455410988 |
| Q8BU04 | Ubr7 | -6.896594155 | 2.051523539 |
| Q80U04 | Pja2 | -6.898117676 | 2.42866516 |
| P58468 | Slx9 | -6.899907163 | 3.879020042 |
| P46664 | Adss2 | -6.901338921 | 5.008539953 |
| Q3UJU9 | Rmdn3 | -6.904692137 | 2.202224068 |
| Q3UGS4 | Mcrip1 | -6.922223468 | 2.553737 |
| Q8R080 | Gtse1 | -6.926701771 | 3.115075241 |
| Q9Z0H1 | Wdr46 | -6.944657373 | 2.040744047 |
| Q9CWX9 | Ddx47 | -6.948182627 | 2.405517051 |
| Q8BX02 | Kank2 | -6.964510935 | 2.714373302 |
| Q8C080 | Snx16 | -6.964886102 | 2.835532102 |
| P28474 | Adh5 | -6.968976492 | 2.860685748 |
| Q8BP27 | Sfr1 | -6.975094875 | 3.856285928 |
| Q9JLV6 | Pnkp | -6.979144082 | 3.469514158 |
| Q9Z266 | Snapin | -6.989839175 | 2.220388116 |
| Q9D8U7 | Dtwd1 | -6.99743102 | 2.266905577 |
| Q61033 | Tmpo | -6.997851246 | 2.821938098 |
| Q01320 | Top2a | -7.003299217 | 3.643885847 |
| Q9EP82 | Wdr4 | -7.027434931 | 3.103165063 |
| Q8R1J3 | Zcchc9 | -7.049765478 | 2.642366431 |
| Q6PDL0 | Dync1li2 | -7.070810385 | 2.66975613 |
| Q9QXX8 | Nufip1 | -7.086838621 | 2.66975613 |
| Q69ZK6 | Jmjd1c | -7.093409569 | 2.589137733 |
| Q8R4U7 | Luzp1 | -7.093925255 | 2.429245678 |
| Q8K371 | Amotl2 | -7.095279676 | 3.111043383 |
| Q5ND52 | Mrm3 | -7.133440322 | 2.142989949 |
| Q61189 | Clns1a | -7.136349565 | 2.026423158 |
| Q99LQ1 | Mbip | -7.137999857 | 2.275090062 |
| O88792 | F11r | -7.162294496 | 2.077988818 |
| Q9QXD8 | Limd1 | -7.165214217 | 2.190703399 |
| Q8VE88 | Fam114a2 | -7.17233822 | 2.225383415 |
| Q8CIB9 | Esco2 | -7.17920503 | 2.044694 |
| Q5DU00 | Dcdc2 | -7.184714607 | 2.472019672 |
| Q9WTN3 | Srebf1 | -7.19925955 | 2.069250001 |
| Q68FL6 | Mars1 | -7.218832734 | 2.55581188 |
| Q9Z2E2 | Mbd1 | -7.231947616 | 2.175065389 |
| Q6PG16 | Hjurp | -7.238164184 | 2.091610845 |
| Q8BHL4 | Gprc5a | -7.246195384 | 2.573218386 |
| Q9JJ89 | Ccdc86 | -7.247774891 | 2.66975613 |
| P02340 | Tp53 | -7.249448038 | 3.558813616 |
| Q91W40 | Klc3 | -7.270722835 | 2.564653124 |
| P58686 | Bbln | -7.277372673 | 3.907318488 |
| Q66T02 | Plekhg5 | -7.281104209 | 2.171652661 |
| Q8K296 | Mtmr3 | -7.281713587 | 2.471222381 |
| Q9CX66 | Nopchap1 | -7.30954881 | 2.596787818 |
| Q9WU62 | Incenp | -7.310957504 | 2.119464109 |
| Q8VEB3 | MACIR | -7.337107612 | 2.381110338 |
| Q5PSV9 | Mdc1 | -7.34850033 | 3.117083711 |
| Q9ER69 | Wtap | -7.349779798 | 2.172319439 |
| Q8CJF7 | Ahctf1 | -7.352702725 | 2.120831581 |
| Q9CZW5 | Tomm70 | -7.354423455 | 2.343975823 |
| Q923U0 | Tom1l1 | -7.372869713 | 2.419524971 |
| P47963 | Rpl13 | -7.377008605 | 2.017737277 |
| Q03347 | Runx1 | -7.377881252 | 2.608179809 |
| Q922Y1 | Ubxn1 | -7.381612636 | 2.66975613 |
| O55028 | Bckdk | -7.388918692 | 2.080910816 |
| Q9QZL0 | Ripk3 | -7.434340384 | 3.461126922 |
| Q9JM93 | Arl6ip4 | -7.461702119 | 2.503507207 |
| Q8K298 | Anln | -7.47996166 | 3.431956117 |
| Q52KR3 | Prune2 | -7.481581075 | 2.66975613 |
| Q9Z2U4 | Elf4 | -7.481984213 | 2.17647695 |
| Q80U56 | Avl9 | -7.495964569 | 2.237080566 |
| Q91YK2 | Rrp1b | -7.501161731 | 2.874379683 |
| Q8K212 | Pacs1 | -7.508699433 | 2.156810072 |
| Q8BGB5 | Limd2 | -7.516788365 | 2.331333146 |
| Q7TT18 | Atf7ip | -7.530361943 | 2.589137733 |
| Q9JKP8 | Chrac1 | -7.535546971 | 2.179660064 |
| Q8CEC0 | Nup88 | -7.54453412 | 2.514195222 |
| Q6IE82 | Jade3 | -7.559213356 | 2.202224068 |
| O70133 | Dhx9 | -7.564896556 | 2.438961756 |
| E9Q9R9 | Dlg5 | -7.569769905 | 2.113840169 |
| Q61850 | Foxc2 | -7.589056473 | 2.302141674 |
| P30306 | Cdc25b | -7.596230746 | 2.185326073 |
| Q9JJY4 | Ddx20 | -7.602252928 | 2.471222381 |
| Q8C0R0 | Usp37 | -7.60330649 | 2.156990505 |
| Q3U2P1 | Sec24a | -7.606842923 | 2.202224068 |
| Q61160 | Fadd | -7.617524285 | 2.159486746 |
| Q9ERA6 | Tfip11 | -7.657938982 | 2.142989949 |
| Q80U35 | Arhgef17 | -7.667352551 | 3.115075241 |
| Q6AW69 | Cgnl1 | -7.673387221 | 2.719371343 |
| Q91XC0 | Ajuba | -7.680176954 | 3.152428716 |
| O35295 | Purb | -7.689260232 | 2.113840169 |
| Q3UJV1 | Ccdc61 | -7.691854986 | 2.131163697 |
| P35550 | Fbl | -7.711392577 | 2.380039316 |
| O88286 | Wiz | -7.714536926 | 2.41223108 |
| Q922D8 | Mthfd1 | -7.722501678 | 2.31479657 |
| P33174 | Kif4 | -7.737213764 | 3.590913466 |
| Q9QXG4 | Acss2 | -7.741916255 | 2.449281532 |
| Q8BIJ7 | Rufy1 | -7.77059292 | 2.487712064 |
| P59438 | Hps5 | -7.780905819 | 2.66009481 |
| Q9EPK5 | Wwtr1 | -7.806061966 | 2.07687452 |
| Q8BHX3 | Cdca8 | -7.8083982 | 2.652935656 |
| P70441 | Nherf1 | -7.822619775 | 2.171402685 |
| Q9CU65;Q3U2E2 | Q9CU65;Q3U2E2 | -7.825440064 | 3.046837233 |
| Q8CGF1 | Arhgap29 | -7.88145857 | 2.497997585 |
| Q6PB75 | Tent4a | -7.88605344 | 2.02632166 |
| Q80VI1 | Trim56 | -7.904350702 | 2.906411417 |
| Q9CWY8 | Rnaseh2a | -7.937918864 | 2.451306968 |
| P23804 | Mdm2 | -7.940148879 | 2.218200604 |
| Q9D2H6 | Sp2 | -7.943712036 | 2.380039316 |
| Q9Z2C4 | Mtmr1 | -7.949104623 | 2.397648279 |
| Q6P9P6 | Kif11 | -7.958666819 | 2.193796708 |
| Q8BH74 | Nup107 | -7.96021108 | 2.416850135 |
| Q6ZPF3 | Tiam2 | -7.964091512 | 2.274399182 |
| Q9JHW4 | Eefsec | -7.969970563 | 2.422063203 |
| Q9CR29 | Ccdc43 | -8.002892072 | 2.66009481 |
| Q61823 | Pdcd4 | -8.008503074 | 2.319866609 |
| Q9D2E2 | Toe1 | -8.036819829 | 2.120831581 |
| Q3TJD7 | Pdlim7 | -8.039108085 | 2.185747866 |
| Q7TN29 | Smap2 | -8.039155168 | 2.099280737 |
| Q8R3N1 | Nop14 | -8.072110541 | 2.198321186 |
| Q69ZN7 | Myof | -8.134379322 | 2.220342972 |
| Q80XI3 | Eif4g3 | -8.143879205 | 2.66975613 |
| Q99JP4 | Cdc26 | -8.143885692 | 2.177921613 |
| Q8C5N3 | Cwc22 | -8.149470637 | 2.256746404 |
| P55194 | Sh3bp1 | -8.185150103 | 2.376504033 |
| P83093 | Stim2 | -8.186691018 | 2.713336585 |
| Q62377 | Zrsr2 | -8.234191146 | 3.806743398 |
| Q3TX08 | Trmt1 | -8.235204649 | 2.153536377 |
| O08795 | Prkcsh | -8.257346912 | 2.179660064 |
| Q6DID3 | Scaf8 | -8.269942707 | 2.601100938 |
| Q8BL80 | Arhgap22 | -8.29839356 | 2.736708897 |
| Q9JJA4 | Wdr12 | -8.329720515 | 2.157282761 |
| Q5HZJ0 | Drosha | -8.352784543 | 2.148838242 |
| Q01341 | Adcy6 | -8.372981792 | 2.437505107 |
| Q9CZB3 | Thumpd2 | -8.377952323 | 2.630790556 |
| Q8BHE1 | Gemin8 | -8.451883908 | 2.427523518 |
| Q99PG2 | Ogfr | -8.457973788 | 3.879020042 |
| Q8BWY9 | Cip2a | -8.47307075 | 2.553737 |
| A2A6A1 | Gpatch8 | -8.510587636 | 2.719834101 |
| Q9D8C8 | Ppp1r35 | -8.522020276 | 2.438961756 |
| Q8BIW9 | Chtf18 | -8.543680643 | 2.908856606 |
| Q99K01 | Pdxdc1 | -8.550207432 | 2.381110338 |
| Q61846 | Melk | -8.614634425 | 2.927568233 |
| Q99MS7 | Ehbp1l1 | -8.64879523 | 2.397648279 |
| Q8K1J5 | Sde2 | -8.653798056 | 2.227126091 |
| Q80WE4 | Kif20b | -8.658974883 | 2.202224068 |
| Q9JJA7 | Ccnl2 | -8.675301441 | 2.449281532 |
| P33611 | Pola2 | -8.699393966 | 2.480199952 |
| Q91VM5 | Rbmxl1 | -8.6994308 | 2.174901515 |
| Q61188 | Ezh2 | -8.707981588 | 2.274399182 |
| P97360 | Etv6 | -8.752648245 | 2.66975613 |
| Q99KR3 | Lactb2 | -8.817469259 | 2.138639356 |
| Q8C0J6 | Sowahc | -8.851783145 | 2.400261522 |
| Q00PI9 | Hnrnpul2 | -8.874474113 | 2.091610845 |
| Q80WT5 | Aftph | -8.884057109 | 2.471222381 |
| O35551 | Rabep1 | -8.918270211 | 2.471222381 |
| Q99PL6 | Ubxn6 | -8.922442783 | 2.382084818 |
| Q9D0L8 | Rnmt | -8.933511772 | 2.16554504 |
| O70400 | Pdlim1 | -8.937492134 | 2.88604899 |
| Q61029;Q61033 | Q61029;Q61033 | -8.937553981 | 4.596940043 |
| Q61140 | Bcar1 | -8.974163716 | 2.66975613 |
| Q80TM9 | Nisch | -8.978180313 | 2.6494188 |
| P70296 | Pebp1 | -9.040100896 | 3.127920927 |
| Q8CHH9 | Septin8 | -9.060508662 | 3.110799061 |
| P04184 | Tk1 | -9.080132587 | 3.007246761 |
| P52927 | Hmga2 | -9.141078156 | 2.540662548 |
| Q9DBD5 | Pelp1 | -9.146688898 | 2.202224068 |
| O09044 | Snap23 | -9.164083137 | 2.130549409 |
| Q8C3R1 | Brat1 | -9.165387409 | 2.220388116 |
| Q9CQQ8 | Lsm7 | -9.174722551 | 2.194359831 |
| Q63918 | Cavin2 | -9.205132463 | 2.66975613 |
| Q501J7 | Phactr4 | -9.323310464 | 2.476452255 |
| Q9DB42 | Znf593 | -9.326370423 | 2.261550667 |
| Q8VDJ3 | Hdlbp | -9.336565893 | 3.879020042 |
| Q80UM3 | Naa15 | -9.447069263 | 3.312162942 |
| A6H619 | Phrf1 | -9.484672335 | 2.063803518 |
| Q99K30 | Eps8l2 | -9.570362531 | 2.064312807 |
| Q3UMC0 | Afg2a | -9.611147427 | 2.463270389 |
| Q8VBT0 | Tmx1 | -9.645607241 | 2.449281532 |
| Q60855 | Ripk1 | -9.692591573 | 2.759761158 |
| Q9D1R2 | Kti12 | -9.778353757 | 2.288996011 |
| P58404 | Strn4 | -9.860235703 | 2.970039085 |
| Q99MR1 | Gigyf1 | -9.972725678 | 2.08457692 |
| Q9Z2D6 | Mecp2 | -10.14993219 | 2.400261522 |
| P59759 | Mrtfb | -10.68597681 | 2.001742648 |
| Q9D168 | Ints12 | -10.70854756 | 2.506655111 |
| Q6P3Y5 | Znf280c | -10.8388928 | 2.925004879 |
| Q9D706 | Rpap3 | -10.86618311 | 3.715457053 |
| O35609 | Scamp3 | -10.99068892 | 3.312162942 |
| Q60876 | Eif4ebp1 | -11.13154535 | 2.185326073 |
| P33609 | Pola1 | -11.3871366 | 2.289916509 |
| P43346 | Dck | -11.71794221 | 3.879020042 |
| Q99J09 | Wdr77 | -11.89029538 | 2.077493006 |
| Q63932 | Map2k2 | -12.71347418 | 2.68085938 |
