## Supplementary material for "ISG15 Differentially Modulates Clade Ib and II MPXV Infection in MEF cells": TableS7

| **Name** | **Gene** | **fold_change** | **p_value_adj_neg_log10** |
| --- | --- | --- | --- |
| P54728 | Rad23b | 5.850802136 | 2.602069481 |
| Q6A0D4 | Rftn1 | 5.781040074 | 3.441387955 |
| A2ARV4 | Lrp2 | 5.658732072 | 2.500415242 |
| Q64133 | Maoa | 5.582343992 | 4.144785421 |
| Q3UTJ2 | Sorbs2 | 5.195531412 | 2.97651221 |
| P12849 | Prkar1b | 4.93917913 | 3.046039612 |
| Q8BWS5 | Gprin3 | 4.932283755 | 2.300478163 |
| P37804 | Tagln | 4.714888173 | 2.194600316 |
| P25233 | Ndn | 4.544161776 | 4.15819844 |
| P17156 | Hspa2 | 4.52422712 | 2.885013369 |
| O88273 | Grem2 | 4.465031738 | 2.482372558 |
| P59729 | Rin3 | 4.109056855 | 3.309211961 |
| Q08091 | Cnn1 | 3.649879181 | 3.49774668 |
| Q9CYH6 | Rrs1 | 3.640106352 | 4.144785421 |
| Q9QWV4 | Mlf1 | 3.626142749 | 4.095498649 |
| Q5PR69 | Cracd | 3.625708826 | 4.18443978 |
| Q64314 | Cd34 | 3.617117736 | 2.885013369 |
| Q6PH08 | Erc2 | 3.533522429 | 4.006318172 |
| Q99MQ1 | Bicc1 | 3.528043177 | 3.049816408 |
| Q7TN22 | Txndc16 | 3.504213488 | 3.325176965 |
| P14901 | Hmox1 | 3.486303392 | 2.396309227 |
| Q3UMB9 | Washc4 | 3.427548235 | 4.14256398 |
| Q562D6 | Trmo | 3.420582572 | 2.528680734 |
| Q69Z89 | Radil | 3.389811428 | 2.747658317 |
| Q6NZA9 | Taf9b | 3.358371699 | 2.860303098 |
| Q8R5A3 | Apbb1ip | 3.350027322 | 3.425888711 |
| P70166 | Cpeb1 | 3.322047851 | 2.087231702 |
| A0A7H0DN41 | OPG068 | 3.308221146 | 2.997704742 |
| P55821 | Stmn2 | 3.293040898 | 3.660337349 |
| Q9D6X5 | Slc52a3 | 3.275760322 | 2.971077461 |
| Q9R123 | Naa80 | 3.247056766 | 2.01533078 |
| Q0VGU4 | Vgf | 3.230317676 | 2.037648187 |
| Q9QUN9 | Dkk3 | 3.222225317 | 2.382819451 |
| Q7TMF2 | Eri1 | 3.218209813 | 2.281916398 |
| E9Q0S6 | Tns1 | 3.180600338 | 4.396948082 |
| Q3USB7 | Plcl1 | 3.096624397 | 2.644733917 |
| Q60760 | Grb10 | 3.081503131 | 3.509627917 |
| Q9CYT6 | Cap2 | 3.073536928 | 2.416237705 |
| Q9WTQ5 | Akap12 | 2.986093414 | 4.446806959 |
| O55091 | Impact | 2.902212973 | 3.419723155 |
| G3X912 | Sprtn | 2.898776096 | 2.141038613 |
| Q91X88 | Pomgnt1 | 2.817527404 | 2.610076138 |
| Q9CPN8 | Igf2bp3 | 2.804634753 | 4.317927216 |
| P97445 | Cacna1a | 2.694301036 | 2.058880328 |
| Q8R3B1 | Plcd1 | 2.673302683 | 2.383216067 |
| Q02819 | Nucb1 | 2.65081273 | 2.616508103 |
| O08999 | Ltbp2 | 2.633527397 | 2.904706672 |
| Q9Z2Z9 | Gfpt2 | 2.59087076 | 3.188539872 |
| P37889 | Fbln2 | 2.56899134 | 2.340373718 |
| Q8BGW5 | Nabp1 | 2.529759709 | 2.971077461 |
| Q5SWP3 | Nacad | 2.516151253 | 4.103940296 |
| Q99N80 | Sytl1 | 2.446705476 | 2.255791719 |
| A3KGF9 | Ccdc9b | 2.425692866 | 3.068648592 |
| Q9Z1P7 | Kank3 | 2.410926869 | 3.164005544 |
| P53351 | Plk2 | 2.345297927 | 3.30694251 |
| P29533 | Vcam1 | 2.334608697 | 2.205416203 |
| Q05860 | Fmn1 | 2.313562296 | 3.782573839 |
| A0A7H0DN95 | OPG123 | 2.280228111 | 3.30694251 |
| P97333 | Nrp1 | 2.271636567 | 2.242414532 |
| A0A7H0DNC8 | OPG157 | 2.257076978 | 3.27197127 |
| P28474 | Adh5 | 2.206194663 | 2.642723699 |
| Q8BN59 | Larp6 | 2.197210756 | 3.332835054 |
| Q3UW53 | Niban1 | 2.196437348 | 4.396948082 |
| Q62219 | Tgfb1i1 | 2.187307823 | 3.941582527 |
| A0A7H0DN76 | OPG104 | 2.174755761 | 2.762714377 |
| P22893 | Zfp36 | 2.162192367 | 2.905731638 |
| Q80ZQ5 | Jazf1 | 2.150389831 | 3.017958809 |
| P13595 | Ncam1 | 2.149358512 | 3.242860957 |
| Q8R105 | Vps37c | 2.144729944 | 3.618684833 |
| A0A7H0DN37 | OPG064 | 2.12753312 | 3.441387955 |
| Q3UPL5 | Ag2 | 2.096357471 | 4.023433175 |
| P0DTN1 | OPG139 | 2.095123312 | 4.006318172 |
| M1L535 | OPG128 | 2.090014143 | 2.950250716 |
| A0A7H0DNA2 | OPG130 | 2.072871835 | 3.941582527 |
| Q91XV3 | Basp1 | 2.03718041 | 3.202058311 |
| Q9CZC8 | Scrn1 | 2.027264724 | 2.288762677 |
| Q8K4G5 | Ablim1 | 2.024016375 | 4.317927216 |
| A0A7H0DNC1 | OPG149 | 2.020827807 | 3.661943716 |
| Q80UN9 | Trit1 | 2.016822836 | 2.187619015 |
| P14115 | Rpl27a | 2.015876587 | 2.064095642 |
| Q64096 | Mcf2l | 2.013867123 | 4.14256398 |
| A0A7H0DN25 | OPG052 | 2.008789039 | 4.144785421 |
| M1L9Q3 | OPG137 | 2.00176021 | 3.6038722 |
| P28571 | Slc6a9 | 2.00166907 | 2.482372558 |
| M1L511 | OPG098 | 1.995707605 | 4.118675525 |
| A0A7H0DN27 | OPG054 | 1.980923441 | 2.281916398 |
| A0A7H0DN84 | OPG112 | 1.959499018 | 4.144785421 |
| A0A7H0DN80 | OPG108 | 1.953778092 | 3.166479573 |
| A0A7H0DNB9 | OPG147 | 1.93003289 | 2.269761496 |
| P58771 | Tpm1 | 1.927407548 | 3.164005544 |
| Q9ERS5 | Plekha2 | 1.916338215 | 3.110891075 |
| Q9Z191 | Eya4 | 1.904232484 | 4.006318172 |
| Q80U16 | Ripor2 | 1.886067781 | 3.498905257 |
| Q5DTU0 | Afap1l2 | 1.875438954 | 3.618684833 |
| M1L543 | OPG138 | 1.863258612 | 2.997704742 |
| P0DTM9 | OPG001 | 1.839009989 | 2.876556074 |
| Q99PP9 | Trim16 | 1.83783977 | 3.598869839 |
| A0A7H0DN35 | OPG062 | 1.833949803 | 4.308848214 |
| Q8BU85 | Msrb3 | 1.829614116 | 4.308848214 |
| Q9Z2V5 | Hdac6 | 1.825047361 | 3.941582527 |
| P17742 | Ppia | 1.824201423 | 3.113641481 |
| A0A7H0DN78 | OPG106 | 1.819975109 | 2.753941155 |
| A0A7H0DND2 | OPG161 | 1.813639803 | 2.996842248 |
| Q9D903 | Ebna1bp2 | 1.801496057 | 2.506446366 |
| A0A7H0DN42 | OPG069 | 1.779161577 | 3.840831393 |
| Q8BJS4 | Sun2 | 1.759635023 | 2.755217038 |
| Q60767 | Ly75 | 1.745787315 | 2.171836462 |
| A0A7H0DNB8 | OPG146 | 1.742170921 | 2.463515442 |
| A0A7H0DN49;Q8V518 | A0A7H0DN49;Q8V518 | 1.741975432 | 3.92091106 |
| A0A7H0DNA8 | OPG136 | 1.738136322 | 3.189175533 |
| M1KJ27 | OPG129 | 1.736730356 | 3.293287777 |
| P57787 | Slc16a3 | 1.720830614 | 2.873219838 |
| Q8C033 | Arhgef10 | 1.714225545 | 3.222920681 |
| P81122 | Irs2 | 1.713492806 | 4.317553551 |
| A0A7H0DN77 | OPG105 | 1.707651896 | 4.144785421 |
| P06803 | Pim1 | 1.700147907 | 2.361678087 |
| Q91X58 | Zfand2b | 1.696357126 | 2.670338735 |
| Q9D281 | Fam114a1 | 1.689633992 | 3.195085725 |
| Q61553 | Fscn1 | 1.688721063 | 4.006318172 |
| A0A7H0DN43 | OPG070 | 1.677973452 | 3.04090982 |
| A0A7H0DN90 | OPG118 | 1.656525897 | 2.131411024 |
| A0A7H0DNA3 | OPG131 | 1.643067129 | 3.498905257 |
| M1L9M0 | OPG082 | 1.632169215 | 3.087285308 |
| Q91YD6 | Vill | 1.621963596 | 4.144785421 |
| Q01063 | Pde4d | 1.593877716 | 3.087285308 |
| A0A7H0DN74 | OPG102 | 1.587972305 | 3.30694251 |
| A0A7H0DN57 | OPG085 | 1.586059025 | 3.499695209 |
| P54726 | Rad23a | 1.57883856 | 2.288762677 |
| P62806 | H4c1; H4c2; H4c3; H4c4; H4c6; H4c8; H4c9; H4c11; H4c12; Hist1h4m; H4c14; H4c16 | 1.576341038 | 2.965070464 |
| Q8BI21 | Rnf38 | 1.575831112 | 2.152448721 |
| Q8VDV3 | Rab3il1 | 1.574798793 | 3.100204631 |
| Q9JKF6 | Nectin1 | 1.572276551 | 2.651535612 |
| Q62394 | Znf185 | 1.563162559 | 3.205853679 |
| E9Q137 | Tex264 | 1.545940108 | 2.762841129 |
| Q61391 | Mme | 1.541876988 | 2.52569591 |
| A0A7H0DN19 | OPG046 | 1.526886522 | 2.711956482 |
| Q8K3K8 | Optn | 1.520957228 | 3.92091106 |
| Q8CE50 | Snx30 | 1.518372372 | 2.051421266 |
| Q6P9P8 | Dennd2c | 1.512275407 | 3.015735396 |
| O88746 | Tom1 | 1.51153245 | 3.205180239 |
| Q921I6 | Sh3bp4 | 1.50580126 | 3.425888711 |
| Q9R0A5 | Nek3 | 1.503111478 | 2.500415242 |
| A0A7H0DNB2 | OPG140 | 1.489870115 | 2.469045944 |
| D3YXJ0 | Dgkh | 1.48746031 | 2.594444994 |
| Q7TPW1 | Nexn | 1.486234421 | 2.61225799 |
| M1L9M3 | OPG087 | 1.48141049 | 2.992158601 |
| A0A7H0DN30 | OPG057 | 1.472111829 | 3.307968799 |
| P55041 | Gem | 1.457961319 | 2.562887456 |
| Q91W69 | Epn3 | 1.449603878 | 3.873650475 |
| Q60973 | Rbbp7 | 1.447955068 | 2.97651221 |
| Q9ET54 | Palld | 1.447257628 | 3.354367453 |
| Q9QZS8 | Sh2d3c | 1.440600993 | 2.677691126 |
| Q8R5F7 | Ifih1 | 1.438403101 | 3.30694251 |
| P97315 | Csrp1 | 1.422536156 | 4.144785421 |
| Q8BRV5 | Kiaa1671 | 1.413673706 | 3.070421594 |
| Q91VH2 | Snx9 | 1.412745442 | 3.768567729 |
| Q80TF3 | Pcdh19 | 1.408875624 | 2.764752338 |
| Q3ZK22 | Vezt | 1.40479161 | 2.649006669 |
| A0A7H0DNB6 | OPG144 | 1.401870156 | 3.164005544 |
| A0A7H0DN89 | OPG117 | 1.396972639 | 4.021059643 |
| Q80UE6 | Wnk4 | 1.395176552 | 2.98844401 |
| Q6PB44 | Ptpn23 | 1.394988114 | 2.212100705 |
| A0A7H0DN72 | OPG100 | 1.389165894 | 3.111815085 |
| A0A7H0DNC3 | OPG151 | 1.368782988 | 3.403097154 |
| Q8BMK4 | Ckap4 | 1.367011876 | 2.876556074 |
| A0A7H0DN15 | OPG042 | 1.366212537 | 3.332835054 |
| B1AVH7 | Tbc1d2 | 1.365121401 | 3.020820623 |
| P97303 | Bach2 | 1.358694543 | 2.971077461 |
| O35618 | Mdm4 | 1.353175881 | 2.544293098 |
| P98199 | Atp8b2 | 1.352087151 | 2.082888455 |
| P70444 | Bid | 1.348117553 | 3.04090982 |
| A0A7H0DN97 | OPG125 | 1.345746453 | 3.30694251 |
| Q8CFD4 | Snx8 | 1.337955072 | 2.971077461 |
| Q9D2V7 | Coro7 | 1.335440418 | 3.555567533 |
| Q8VED9 | Lgalsl | 1.330439171 | 2.698482717 |
| O88643 | Pak1 | 1.32905901 | 2.843571424 |
| Q8BGT7 | Smndc1 | 1.311874089 | 3.425888711 |
| Q7TN75 | Peg10 | 1.311592761 | 2.734785034 |
| A0A7H0DN92;Q8V4Y0 | A0A7H0DN92;Q8V4Y0 | 1.301553251 | 2.340294 |
| P26041 | Msn | 1.298224742 | 2.138875715 |
| P98203 | Arvcf | 1.289685079 | 3.443050861 |
| Q06186 | Hbegf | 1.278212741 | 2.416237705 |
| Q60988 | Stil | 1.276329484 | 3.624120707 |
| Q7TMK6 | Hook2 | 1.272767215 | 2.386754641 |
| Q9WVS7 | Map2k5 | 1.269963683 | 2.579894829 |
| Q9R1E0 | Foxo1 | 1.269006102 | 3.068648592 |
| Q5PR68 | Cep112 | 1.267641157 | 2.376967247 |
| Q63918 | Cavin2 | 1.264466416 | 2.753941155 |
| P59242 | Cgn | 1.262904144 | 3.661943716 |
| A0A7H0DN62 | OPG090 | 1.255475568 | 2.650494932 |
| Q69Z66 | Mysm1 | 1.238069898 | 2.416237705 |
| O88573 | Aff1 | 1.225464061 | 3.309211961 |
| Q9JLC8 | Sacs | 1.223784627 | 3.332835054 |
| P57080 | Usp25 | 1.213826621 | 2.863892912 |
| O54918 | Bcl2l11 | 1.193741995 | 2.284345197 |
| Q3U1F9 | Pag1 | 1.193346252 | 3.344960725 |
| Q9D3S3 | Snx29 | 1.18896618 | 3.189175533 |
| Q9EPM5 | Sync | 1.187973431 | 2.986859729 |
| G3XA57 | Rab11fip2 | 1.179382514 | 2.129915196 |
| A0A7H0DN82 | OPG110 | 1.173349131 | 3.087285308 |
| Q3U0J8 | Tbc1d2b | 1.169410306 | 3.164005544 |
| Q9QZS3 | Numb | 1.168580804 | 2.530498008 |
| Q8C739 | Fam110b | 1.159916867 | 3.216208563 |
| Q61234 | Snta1 | 1.151800253 | 3.205180239 |
| P33215 | Nedd1 | 1.147065811 | 2.506112875 |
| Q8BIG4 | Fbxo28 | 1.146782935 | 2.218611353 |
| Q99MI1 | Erc1 | 1.145483326 | 2.482372558 |
| O70479 | Tnfaip1 | 1.144593048 | 3.384399705 |
| Q8C080 | Snx16 | 1.143211888 | 2.312448857 |
| Q64727 | Vcl | 1.140992316 | 3.349327713 |
| Q62082 | Myl10 | 1.126563706 | 2.661800962 |
| O08584 | Klf6 | 1.124954425 | 3.327511175 |
| Q3ULZ2 | Fhdc1 | 1.124287511 | 3.293287777 |
| A0A7H0DNG4 | OPG205 | 1.112542703 | 2.282880362 |
| Q8BMI0 | Fbxo38 | 1.109408327 | 2.401086919 |
| P43274 | H1-4 | 1.104807709 | 3.406379031 |
| P70295 | Aup1 | 1.098227989 | 3.183135385 |
| A0A7H0DND5 | OPG164 | 1.096422363 | 3.164005544 |
| Q8K1N2 | Phldb2 | 1.094876992 | 4.144785421 |
| Q9QYY0 | Gab1 | 1.094275061 | 3.013887794 |
| Q62433 | Ndrg1 | 1.075132003 | 2.644733917 |
| Q9JIP4 | Panx1 | 1.07444968 | 2.418111852 |
| P20357 | Map2 | 1.070890645 | 3.164005544 |
| O35682 | Myadm | 1.066760095 | 2.156117889 |
| Q9R1X5 | Abcc5 | 1.065299184 | 3.601946827 |
| P61967 | Ap1s1 | 1.06218475 | 3.154392116 |
| P15037 | Ets2 | 1.061354143 | 2.929637984 |
| Q7TSY8 | Sgo2 | 1.059946497 | 2.542721171 |
| P61079 | Ube2d3 | 1.055970473 | 3.303903991 |
| Q60769 | Tnfaip3 | 1.050514413 | 3.49774668 |
| Q9ESK9 | Rb1cc1 | 1.042919956 | 3.15677242 |
| P97433 | Arhgef28 | 1.040953743 | 3.40056193 |
| P62700 | Ypel5 | 1.03195242 | 2.255624337 |
| Q9D032 | Ssbp3 | 1.031715453 | 2.110489265 |
| Q62441 | Tle4 | 1.023592593 | 2.3968504 |
| P26039 | Tln1 | 1.023440971 | 3.411987449 |
| Q8VC56 | Rnf8 | 1.023403794 | 2.971085586 |
| Q3UMF0 | Cobll1 | 1.023124918 | 3.131712819 |
| Q9CQ80 | Vps25 | 1.021875563 | 3.029596997 |
| Q0VGY8 | Tanc1 | 1.019149076 | 2.997704742 |
| Q6P549 | Inppl1 | 1.016696558 | 2.270738411 |
| Q8BH07 | Arl6ip6 | 1.015133696 | 2.310058933 |
| P28658 | Atxn10 | 1.014169954 | 2.317514 |
| Q6XUX1 | Dstyk | 1.013165368 | 2.548747915 |
| Q9Z0E8 | Slc22a5 | 1.011330882 | 2.737204442 |
| P82343 | Renbp | 1.008434308 | 2.76762081 |
| P06537 | Nr3c1 | 1.003909881 | 3.415772866 |
| Q8CHP0 | Zc3h3 | 1.002242168 | 2.967739929 |
| Q9D5D8 | Cdyl2 | -1.00298522 | 2.838265752 |
| Q5XG73 | Acbd5 | -1.003503322 | 2.401086919 |
| Q6NZK5 | Kiaa1328 | -1.019498214 | 2.997704742 |
| Q91YD9 | Wasl | -1.026126024 | 3.020820623 |
| Q9Z1B7 | Mapk13 | -1.034384384 | 2.263883093 |
| P17095 | Hmga1 | -1.0448906 | 2.085860358 |
| Q3TTA7 | Cblb | -1.064350317 | 3.257426242 |
| B1AWL2 | Znf462 | -1.064482567 | 3.202058311 |
| Q8BPY9 | Fignl1 | -1.076901393 | 2.575740861 |
| Q80YD1 | Supv3l1 | -1.079401158 | 2.047715697 |
| Q99KX1 | Mlf2 | -1.083779991 | 2.997704742 |
| O35973 | Per1 | -1.085210241 | 2.235868472 |
| P70677 | Casp3 | -1.104134908 | 2.469045944 |
| P21995 | Emb | -1.105248497 | 4.006318172 |
| Q9EP97 | Senp3 | -1.134524171 | 3.164005544 |
| Q9R0H0 | Acox1 | -1.135463117 | 2.217505567 |
| Q5U465 | Ccdc125 | -1.138173249 | 2.482372558 |
| O35231 | Kifc3 | -1.143546354 | 2.009605634 |
| Q9QZL0 | Ripk3 | -1.143592057 | 2.091900279 |
| Q91YT7 | Ythdf2 | -1.143923943 | 3.101915904 |
| Q8JZZ7 | Adgrl2 | -1.149405386 | 2.997704742 |
| Q99JY8 | Plpp3 | -1.160864463 | 2.178415519 |
| Q922D8 | Mthfd1 | -1.169650913 | 3.415772866 |
| Q9DB25 | Alg5 | -1.170396406 | 2.042082106 |
| P48453 | Ppp3cb | -1.177083182 | 2.905731638 |
| Q811I0 | Atpaf1 | -1.179689846 | 2.31693854 |
| Q03173 | Enah | -1.19113568 | 2.645938512 |
| Q99M07 | Coa5 | -1.193642005 | 2.762841129 |
| A0A7H0DN47 | OPG074 | -1.199385555 | 2.453968142 |
| O55003 | Bnip3 | -1.211784835 | 2.486017612 |
| O35149 | Slc30a4 | -1.213463383 | 2.116398705 |
| Q5ND52 | Mrm3 | -1.23032111 | 2.108554804 |
| Q80UW2 | Fbxo2 | -1.239461017 | 2.493164069 |
| Q9JKZ2 | Slc5a3 | -1.246091691 | 2.127577857 |
| P70698 | Ctps1 | -1.267967886 | 2.170140989 |
| P70453 | Pde7a | -1.283241251 | 2.996842248 |
| Q05915 | Gch1 | -1.290283552 | 2.427652858 |
| P23475 | Xrcc6 | -1.301236266 | 3.328376914 |
| Q8VCI5 | Pex19 | -1.315931916 | 2.904706672 |
| B2RVL6 | Zcchc24 | -1.317980116 | 2.031014058 |
| Q9WVB3 | Tle6 | -1.342304576 | 2.149070602 |
| Q8CG19 | Ltbp1 | -1.365377025 | 2.11140031 |
| Q91V09 | Wdr13 | -1.38172783 | 3.170068884 |
| Q9D8T7 | Slirp | -1.388563924 | 2.068752751 |
| Q9WTK5 | Nfkb2 | -1.462619546 | 2.65166401 |
| Q8C5R2 | Proser2 | -1.46623641 | 2.223248759 |
| Q8CDB0 | Mknk2 | -1.476748215 | 2.086066863 |
| Q80UG5 | Septin9 | -1.499604117 | 3.344960725 |
| Q61037 | Tsc2 | -1.504942322 | 3.059373679 |
| P51881 | Slc25a5 | -1.513956781 | 2.271406938 |
| Q8C0N2 | Gpat3 | -1.531120095 | 2.164178501 |
| Q91ZS8 | Adarb1 | -1.581356299 | 2.773212388 |
| Q8CFN5 | Mef2c | -1.584890685 | 2.194422606 |
| Q8R550 | Sh3kbp1 | -1.585715283 | 3.661943716 |
| Q8BXJ2 | Trerf1 | -1.589619442 | 2.137837105 |
| O88327 | Ctnnal1 | -1.594101641 | 3.371849625 |
| P20065 | Tmsb4x | -1.599790094 | 3.015735396 |
| Q3UHU5 | Mtcl1 | -1.619494078 | 2.775163213 |
| Q7TQI8 | Tspyl2 | -1.639538986 | 2.113742862 |
| P06837 | Gap43 | -1.640370396 | 2.486017612 |
| Q9D4H4 | Amotl1 | -1.640746873 | 2.764845889 |
| O09174 | Amacr | -1.646761607 | 3.512699856 |
| Q8BZB3 | Tmem266 | -1.66449054 | 2.764845889 |
| Q99J62 | Rfc4 | -1.672955099 | 2.624208669 |
| Q08874 | Mitf | -1.675282635 | 2.925243931 |
| Q3UC65 | Rsrp1 | -1.685753202 | 2.250315577 |
| E9Q555 | Rnf213 | -1.686068956 | 3.498905257 |
| Q8VBT6 | Apobr | -1.753248878 | 3.30694251 |
| P15806 | Tcf3 | -1.795084705 | 2.904706672 |
| P28359 | Hoxd10 | -1.838651708 | 2.051537194 |
| Q62393 | Tpd52 | -1.846311057 | 3.42580814 |
| Q9JLB0 | Pals2 | -1.877704529 | 2.016503794 |
| Q7M6Y3 | Picalm | -1.91423983 | 2.298512883 |
| Q9WV69 | Dmtn | -1.922375316 | 3.661943716 |
| Q7SIG6 | Asap2 | -1.973954318 | 3.498905257 |
| Q91VH1 | Adipor1 | -2.075506252 | 2.737204442 |
| Q78HU3 | Mvb12a | -2.090263664 | 3.661943716 |
| O35099 | Map3k5 | -2.163504823 | 3.128752148 |
| Q04887 | Sox9 | -2.211179403 | 3.164163452 |
| P56480 | Atp5f1b | -2.237249798 | 3.025282205 |
| Q61043 | Nin | -2.255312146 | 2.267636838 |
| Q8C8R3 | Ank2 | -2.278859188 | 3.013887794 |
| Q8R3F5 | Mcat | -2.279931754 | 2.728243041 |
| Q922T2 | Mfap3 | -2.296122303 | 2.680306258 |
| Q91YQ3 | Csdc2 | -2.331916231 | 2.361678087 |
| Q80TN7 | Nav3 | -2.355282235 | 3.738969735 |
| Q60695 | Rgl1 | -2.366252137 | 3.606422037 |
| Q7TSZ8 | Nacc1 | -2.441898448 | 2.611753457 |
| Q8VE94 | Fam110c | -2.536979311 | 3.661943716 |
| Q6PGG6 | Gnl3l | -2.540327982 | 3.183135385 |
| Q80YA9 | Cnksr2 | -2.542772332 | 2.085860358 |
| O35185 | Bhlhe40 | -2.631735391 | 2.520707546 |
| Q8BVH9 | Mettl6 | -2.712454485 | 2.698126889 |
| Q6ZWM4 | Lsm8 | -2.767412256 | 3.317008303 |
| Q9JIF7 | Copb1 | -2.774612031 | 2.281916398 |
| O70566 | Diaph2 | -2.858411656 | 3.051871581 |
| P35710 | Sox5 | -2.898865466 | 3.661943716 |
| P41164 | Etv1 | -3.056687281 | 2.817836961 |
| Q8R1B0 | Stac2 | -3.06291382 | 4.317927216 |
| Q497H0 | Zfand3 | -3.136980641 | 3.087285308 |
| P10923 | Spp1 | -3.182542372 | 4.103940296 |
| Q8BZ20 | Parp12 | -3.328133377 | 2.104526935 |
| Q61850 | Foxc2 | -3.484975304 | 4.103940296 |
| P29352 | Ptpn22 | -3.539249547 | 2.614849907 |
| Q8CC27 | Cacnb2 | -3.637558196 | 2.940729049 |
| Q3UVY1 | Gpr149 | -3.659784206 | 2.880193836 |
| P70324 | Tbx3 | -3.672719584 | 2.399028359 |
| P35546 | Ret | -3.990343851 | 3.216208563 |
| Q8R332 | Nup58 | -4.425134566 | 3.661943716 |
| Q9D030 | Twist2 | -4.923784541 | 3.078857204 |
| P70326 | Tbx5 | -5.196916167 | 2.594055016 |
| Q9D0B6 | Pbdc1 | -5.538390628 | 3.189175533 |
| Q8JZW5 | Sh2d5 | -5.69432735 | 3.661943716 |
| P97503 | Nkx3-2 | -5.873176894 | 2.397371566 |
| O54834 | Arhgap6 | -6.687829944 | 3.782573839 |
